# High-throughput complement component 4 genomic sequence analysis with C4Investigator

**DOI:** 10.1101/2023.07.18.549551

**Authors:** Wesley M. Marin, Danillo G. Augusto, Kristen J. Wade, Jill A. Hollenbach

## Abstract

The complement component 4 gene locus, composed of the *C4A* and *C4B* genes and located on chromosome 6, encodes for C4 protein, a key intermediate in the classical and lectin pathways of the complement system. The complement system is an important modulator of immune system activity and is also involved in the clearance of immune complexes and cellular debris. The *C4* gene locus exhibits copy number variation, with each composite gene varying between 0-5 copies per haplotype, *C4* genes also vary in size depending on the presence of the HERV retrovirus in intron 9, denoted by *C4(L)* for long-form and *C4(S)* for short-form, which modulates expression and is found in both *C4A* and *C4B*. Additionally, human blood group antigens Rodgers and Chido are located on the C4 protein, with the Rodger epitope generally found on C4A protein, and the Chido epitope generally found on C4B protein. *C4* copy number variation has been implicated in numerous autoimmune and pathogenic diseases. Despite the central role of C4 in immune function and regulation, high-throughput genomic sequence analysis of *C4* variants has been impeded by the high degree of sequence similarity and complex genetic variation exhibited by these genes. To investigate C4 variation using genomic sequencing data, we have developed a novel bioinformatic pipeline for comprehensive, high-throughput characterization of human *C4* sequence from short-read sequencing data, named C4Investigator. Using paired-end targeted or whole genome sequence data as input, C4Investigator determines gene copy number for overall *C4, C4A, C4B, C4(Rodger), C4(Ch), C4(L)*, and *C4(S)*, additionally, C4Ivestigator reports the full overall *C4* aligned sequence, enabling nucleotide level analysis of *C4*. To demonstrate the utility of this workflow we have analyzed *C4* variation in the 1000 Genomes Project Dataset, showing that the *C4* genes are highly poly-allelic with many variants that have the potential to impact C4 protein function.

## Introduction

The *C4* gene locus, composed of the *C4A* and *C4B* genes and located in human chromosomal region 6p21.33, encodes for complement component 4 (C4) protein, a key intermediate in the classical and lectin pathways of the complement system(1). The complement system is an important modulator of immune system activity, can activate the innate and adaptive immune response systems(2–4) and is also involved in the clearance of immune complexes and cellular debris. The *C4* gene locus exhibits copy number variation (CNV), with each composite gene varying between 0-5 copies per haplotype, and importantly, the gene copy number of *C4A* and *C4B* correlate to C4 protein levels(5). *C4* genes also vary in size depending on the presence of the HERV-K(C4) retrovirus in intron 9 (**Figure 1A**), denoted by *C4(L)* for long-form and *C4(S)* for short-form, which modulates expression and is found in both *C4A* and *C4B* resulting in four distinct genomic forms of *C4* (*C4A(L), C4B(L), C4A(S)*, and *C4B(S)*)(5).

**Figure 1.**
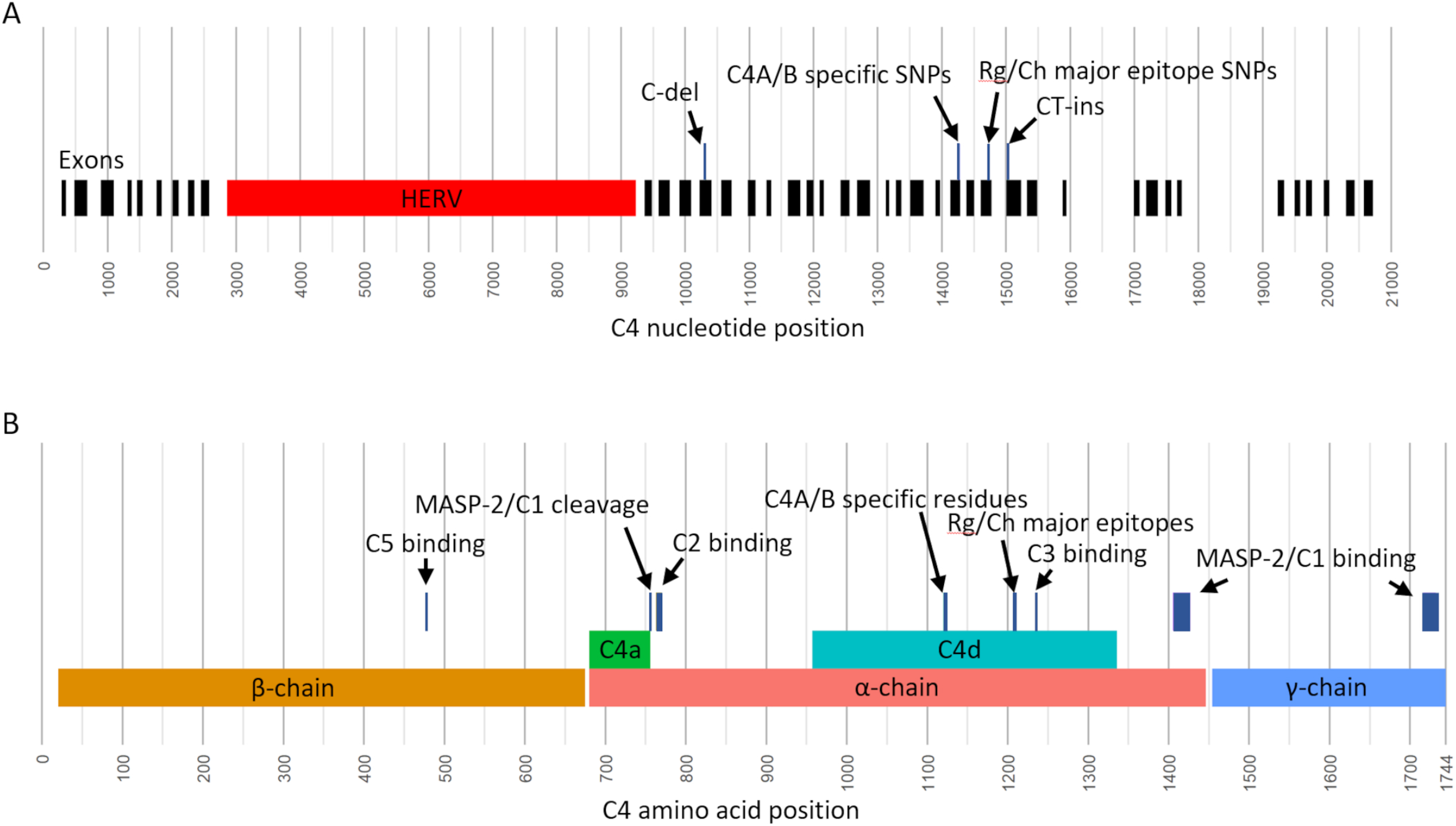
Sequence features of C4 genes and C4 proteins. **(A)** Positions of *C4A* and *C4B* genomic sequence features shown for a long-form of *C4*. Exon positions are marked in black, the HERV retroviral sequence is marked in red, and select sequence variants are shown above the exons. Positions are based on the *C4* alignment reference, which includes 5’UTR and 3’UTR sequence. The C-del variant and the CT-ins variant are frame-shift mutations that result in premature terminations. **(B)** Positions of C4A/C4B protein sequence features. The major chains, α, β, and γ, are shown in the bottom row, the cleavage products, C4a and C4d, are shown on the middle row, and important binding locations and sequence variants are shown in the top row. The amino acid positions include the leading 19 amino acid signal peptide.

C4 is mainly expressed by liver cells, white blood cells, and intestinal epithelial cells(6), but also by central nervous system cells(7). C4 is expressed as two isotypes, C4A and C4B, encoded by the *C4A* and *C4B* genes, respectively. The isotypes have nearly identical sequence but are differentiated by a short peptide sequence motif at positions 1120-1125 (**Figure 1B**), which are **PC**PV**LD** for C4A and **LS**PV**IH** for C4B. Additionally, human blood group antigens Rodgers (Rg) and Chido (Ch) are located on the C4 protein at positions 1207-1210(8–10). The Rg epitope is generally found on C4A protein, and the Ch epitope is generally found on C4B protein. The relative locations of the C4A/B specific single nucleotide polymorphisms (SNPs) and the Rg/Ch major epitope encoding SNPs are shown in **Figure 1A**.

*C4* CNV has been implicated in the neurological diseases schizophrenia(11,12) and Alzheimer’s(13), and there is a large body of evidence connecting *C4A* deficiency and the development of systemic lupus erythematosus (SLE)(14–16), an autoimmune disease. Additionally, while the role of *C4* CNV has yet to be studied in the context of COVID-19 pathology, recent studies have implicated complement hyperactivation with severe SARS-CoV-2 complications(17–19).

Currently, interrogation of *C4* CNV is accomplished through digital droplet polymerase chain reaction (ddPCR)(11,20), which is capable of quantifying gene copy number for overall *C4, C4A(L), C4A(S), C4B(L)* and *C4B(S)*. While this method produces accurate results for *C4A* and *C4B* gene copy number and phasing with long and short form, it is intractable for identifying additional sequence variation at scale, including loss of function mutations(21,22) and recombinations(23,24), and is completely blind to novel sequence variation. High-throughput genomic sequence analysis of *C4* variants has been impeded by the complex genetic variation exhibited by these genes. One recent tool for assessing *C4* sequence variation is the *C4A*/*B* analysis workflow hosted on Terra (25), which was developed using the Genome STRiP software (26) to analyze *C4* from WGS data. However, this tool is currently unpublished and is restricted to analysis of copy number variation of *C4A*/*C4B* specific SNPs and the HERV retrovirus.

Most C4 analysis workflows are targeted at characterizing the region of *C4A*/*C4B* specific SNPs, which encode for an important active site that causes C4A and C4B to have unique biochemistries. However, there are many other vital locations along C4 sequence that when mutated have drastic functional consequences (**Figure 1B**). First are amino acid positions 477 and 478; mutations at these positions can disrupt C5 convertase activity (27,28), an important step in the classical and lectin complement cascade pathways that results in the formation of the membrane attack complex (MAC). Positions 756 and 757 are the site of C1/MASP-2 cleavage(29) to produce C4a and C4b, which is the initial modification made to C4 to initiate the complement cascade. Positions 1405-1427 and 1716-1732 are binding sites for C1/MASP-2 (30,31). Positions 763-770 make up a binding site for C2a (32), an intermediary of the classical and lectin cascade pathways that binds with C4b to make a C3 convertase. Positions 1236 and 1238 are known binding positions for C3b (33), an intermediary that binds with the C4b·C2a complex to make a C5 convertase. Finally, there are known frame-shift mutations on exon 13 and 29 that both result in premature terminations (**Figure 1A**) (22).

Due to the importance of C4 in complement cascade activity, coupled with the high degree of allotypic variation (34,35), we believe that full genomic sequence characterization of *C4* is of vital importance to advancing our understanding of its in human health. To investigate *C4* variation using genomic sequencing data, we have developed a bioinformatic pipeline for comprehensive, high-throughput characterization of human *C4* copy number and sequence variation from short-read sequencing data, named C4Investigator. Using whole genome sequence data as input, C4Investigator determines gene copy number for overall *C4, C4A, C4B, C4(Rg), C4(Ch), C4(L)*, and *C4(S)*; additionally, C4Investigator reports full genomic sequence and highlights frame-shift mutations and potential recombinations.

To demonstrate the utility of C4Investigator, we have applied the workflow to the 1000 Genomes Project (1KGP) high depth 30x WGS data(36,37), a dataset consisting of 3,202 samples, characterizing *C4* copy number and sequence variation for the first time in this dataset to provide a snapshot of population-level differentiation at this important genomic region.

## Materials and Methods

### 1.1 C4Investigator overview

Due to the high degree of sequence similarity between *C4A* and *C4B*, the C4Investigator workflow combines alignments of these two genes into an overall *C4* alignment. A long-form *C4A* sequence and a short-form *C4B* sequence are used as a reference for this alignment. A custom alignment processing workflow, similar to that outlined in Marin et al.(38), was developed to integrate the *C4A* and *C4B* alignments into the overall *C4* alignment. From the overall alignment, *C4* copy number is determined by comparing the median alignment depth across *C4* to the average depth of the Tenascin XB (*TNXB)* gene, a nearby copy-stable gene. Gene copy number of *C4A, C4B, C4(Ch), C4(Rg), C4(L)* and *C4(S)* are determined by multiplying the ratios of *C4A*/*B* specific SNPs, *Rg*/*Ch* specific SNPs and the HERV insertion region, to the overall *C4* copy. *C4A-Ch* and *C4B-Rg* recombinants are identified using read-based phasing. A limitation of this approach is that because of the genomic distance between the *C4A*/*B* specific SNPs to the HERV region, this method is unable to phase *C4A*/*B* with long and short-form.

In addition to gene copy number analysis, C4Investigator outputs the full overall *C4* aligned sequence as a SNP table.

The pipeline is available at: https://github.com/hollenbach-lab/C4Investigator.

### 1.2 C4 alignment workflow

The structural variation of the *C4* gene locus and high-degree of sequence similarity between *C4A* and *C4B* necessitates a custom alignment and processing workflow. The first step of the workflow is a Bowtie2(39) alignment to a reference consisting of a short-form of *C4B*, the long-form of *C4A*, and *TNXB*, which is used as a close proximity normalizer gene. Subsequently, the reads aligned to both *C4A* and *C4B* are combined, formatted, and indexed according to the aligned read formatting procedure outlined in Marin et al. (2021) to generate an overall *C4* alignment used for downstream analysis. The output of this workflow is a *C4* depth table spanning from position -285 5’UTR to position 341 3’UTR with depths marked independently for A, T, C, G, deletions, and insertions.

### 1.3 C4 copy number determination

The median depth of the overall *C4* alignment is normalized by the median depth of *TNXB* to determine the overall *C4* gene copy number. The relative depth ratios of the *C4A* and *C4B* specific SNPs, at positions E26.129, E26.132, E26.140, E26.143, and E26.145, are multiplied by the overall *C4* gene copy number to determine the *C4A* and *C4B* gene copy number. Similarly, the *Rg* and *Ch* major epitope specific SNPs, at positions E28.111, E28.116, E28.125, and E28.126, are processed to determine the *C4(Rg)* and *C4(Ch)* gene copy number. Finally, the depth ratio of the *HERV* insertion, across positions I9.276-I9.6642, is multiplied by the overall *C4* gene copy number to determine the long-form and short-form copy number.

Exon 29 TC insertion sequence depth ratio is multiplied by the overall *C4* copy to determine the copy of loss of function alleles, this value is subtracted from *C4A* gene copy number to give the functional *C4A* copy number. While it is possible for the TC insertion to exist in a *C4B* sequence, this variant is very rare(40) and there is no solid evidence of it in the datasets we analyzed. A similar approach is utilized for the exon 13 C deletion in *C4B* to give the functional *C4B* copy number.

### 1.4 C4 sequence analysis

The overall *C4* depth table is processed to generate a SNP table for positions passing a minimum depth threshold (6 for whole genome sequence data and 20 for targeted sequence data). Heterozygous positions are identified using a depth ratio of 0.5 normalized by the determined *C4* gene copy number. The output of this step is an overall *C4* SNP table with combined sequence for *C4A* and *C4B*.

### 1.5 Targeted sequencing dataset generation

To validate the C4Investigator workflow, we applied targeted-capture next-generation sequencing (NGS) in a cohort of 38 African Americans and 37 European Americans from the United States. These healthy individuals were unrelated and part of the INDIGO (The Immunogenetics for Neurological DIseases working GrOup) cohort(41).

A total of 100 ng of high-quality DNA is fragmented using the Twist EF Kit 2.0 l (Twist Bioscience), incubating for 5 minutes at 37 °C. Subsequently, the fragmented DNA have their ends repaired, poly-A tail added, and are ligated through PCR to Illumina compatible dual index adapters uniquely barcoded. After ligation, fragments are purified with 0.8X ratio Ampure XP magnetic beads (Beckman Coulter) followed by double size selection (0.42X and 0.15X ratios) to select libraries of approximately 800 bp. Finally, libraries are amplified and purified with magnetic beads. After quantification by quantitative PCR, 60 ng of each sample are precisely pooled using ultrasonic acoustic energy, and the enrichment targeted capture is performed with hybridization kits from Twist Bioscience. Briefly, the libraries are bound to 33,620 biotinylated 120 bp probes target the entire MHC (chr6:28525013-33457522, hg38). By using streptavidin magnetic beads, the targeted fragments are captured and then amplified and purified. Enriched libraries are analyzed in BioAnalyzer (Agilent) and quantified by digital-droplet PCR. Finally, enriched libraries are sequenced using NovaSeq6000 (Illumina) with paired-end 150bp sequencing protocol.

C4Investigator was run over both targeted sequencing datasets using a minimum depth of 20 for variant calling and a ratio of 0.50, normalized by the total copy of *C4*, for heterozygous position identification. C4Investigator results were compared to ddPCR results to provide validation for C4 interpretation from targeted sequence data.

### 1.6 ddPCR genotyping

Gene copy number for *C4A, C4B, C4(L)* and *C4(S)* were determined by ddPCR as described previously(11) for 38 samples of African ancestry and 37 samples of European ancestry to provide a copy determination comparison dataset.

### 1.7 1000 Genomes Project analysis

Reads aligned to *C4* and the nearby region were extracted from GRCh38 aligned CRAM files using the coordinates outlined in **Table S1** using Samtools(42). The extracted reads were converted to paired-end FASTQ files using Bazam(43). C4Investigator was run over the paired-end fastq files using a minimum depth of 6 for variant calling and a ratio of 0.50, normalized by the total copy of *C4*, for heterozygous position identification. *C4* copy number results were stratified by superpopulation. Population totals and abbreviations are outlined in **Table 1**.

**Table 1.**
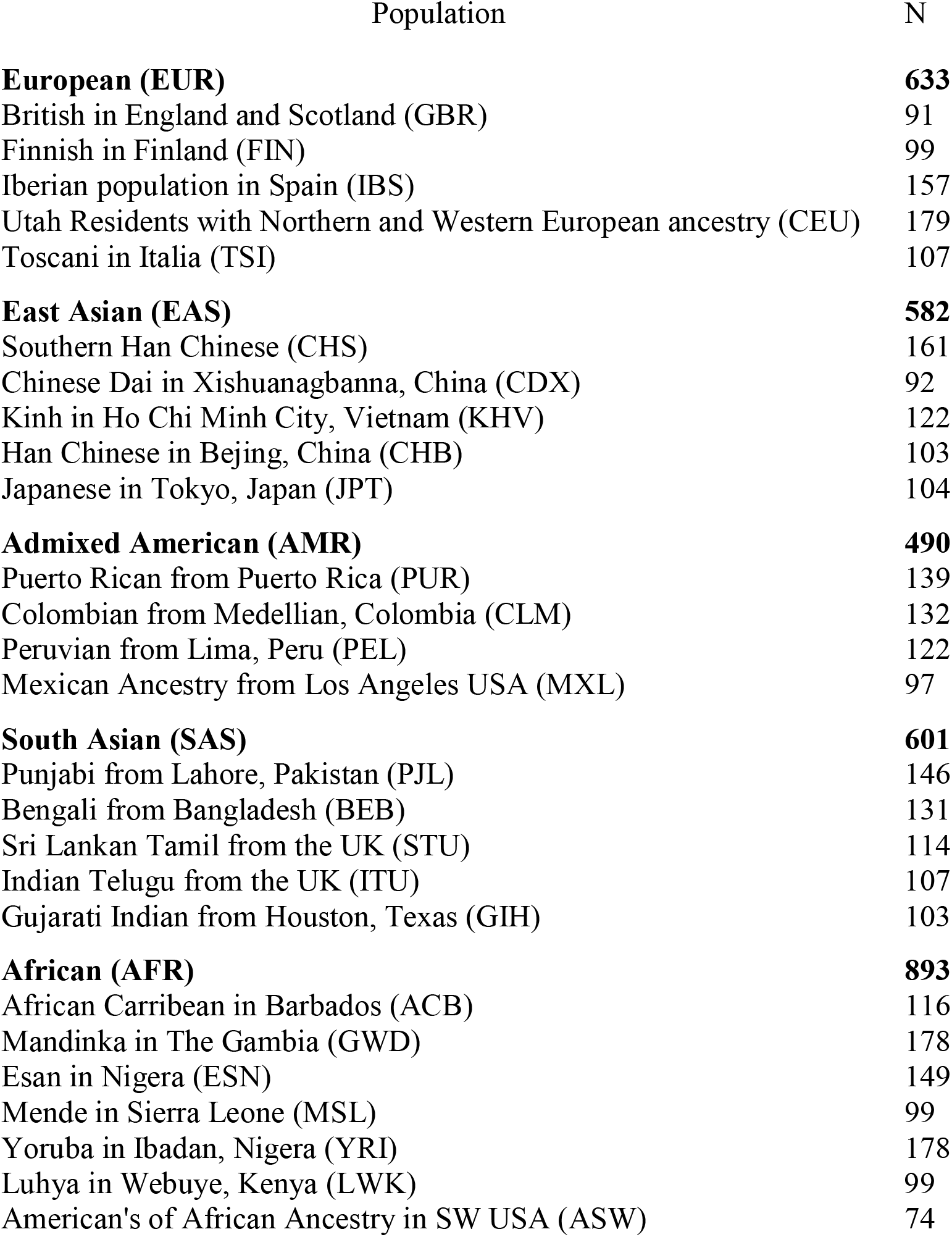
1000 Genomes Project population abbreviations and size.

### 1.8 Validation

C4Investigator performance was validated against ddPCR results for 38 samples of African ancestry and 37 samples of European ancestry. Gene copy number results determined by C4Investigator were compared to ddPCR determined results to quantify the copies of *C4A, C4B, C4(L)* and *C4(S)* that were identified by both methods.

C4Investigator copy number results for the 1KGP dataset were compared to results from the *C4A*/*B* analysis workflow utilizing Genome STRiP(36) implemented in Terra (25). Results were compared across overall *C4, C4A, C4B, C4(L)* and *C4(S)* results. For overall *C4* all results across both datasets were compared. For *C4A* and *C4B* comparison, samples marked as *C4A1, C4A2, C4B1*, or *C4R1*, which represented rare *C4* sequence variants, by the Genome STRiP Terra workflow were excluded, this excluded a total of 55 samples from comparison. For *C4(L)* and *C4(S)* all results were compared. *C4A1, C4A2, C4B1*, and *C4R1* results for C4Investigator were generated by confirming correct phase across positions E26.128 – E26.145, based on the k-mers provided for these variants by the Terra workflow, then determining the copy number of these variants based on the relative SNP depth.

## Results

### 1.9 Performance evaluation – ddPCR copy number comparison

Evaluation of C4Investigator copy number determination performance compared to ddPCR results for European and African datasets show perfect concordance between the two methods for *C4A* and *C4B* copy number determination (**Table 2**), 94% for *C4(S)* and 98% for *C4(L)* for the European dataset, and 89% for *C4(S)* and 91% for *C4(L)* for the African dataset.

**Table 2.** Evaluation of C4Investigator copy number determination performance compared to ddPCR for European and African datasets. *C4(S)* = *C4* short-form, *C4(L)* = *C4* long-form

| <u>Ancestry</u> | <u>C4A</u> | <u>C4B</u> | <u>C4(S)</u> | <u>C4(L)</u> |
| --- | --- | --- | --- | --- |
| African | 1.00 N=76 | 1.00 N=66 | 0.89 N=61 | 0.91 N=81 |
| European | 1.00 N=82 | 1.00 N=70 | 0.94 N=34 | 0.98 N=118 |

### 1.10 Performance evaluation – *C4A/B* Terra copy number comparison

To benchmark C4Investigator performance against another bioinformatic workflow, we compared results for the 1000 Genomes Project dataset (N=3199) against results from the unpublished *C4A/B* Terra workflow(25), a bioinformatic pipeline that utilizes Genome STRiP(36) to quantify *C4* copy number.

Overall *C4* copy determination performance was highly concordant with the *C4A*/*B* Terra workflow, at 99.95% (N=12977). *C4A* and *C4B* copy identification concordance was 99.12% (N=6942) for *C4A* and 98.96% (N=5976) for *C4B. C4(L)* and *C4(S)* copy identification concordance was 99.60% (N=8700). Comparing the additional *C4* variants quantified by *C4A*/*B* Terra workflow showed an overall concordance of 96.6% (N=59).

Investigation into the discordant *C4A* and *C4B* samples showed the ratios of *C4A* were near the copy thresholds for both methods (**Figure S1A**), further examination into the *C4A*/*B* Terra k-mer quality scores showed the discordant samples had a median quality of 9, while concordant samples had a median quality of 62.7 (**Figure S1B**). A similar analysis was performed for the *C4(L)* and *C4(S)* discordant samples, which showed the C4Investigator ratios were near the copy thresholds, while the *C4A*/*B* Terra workflow ratios were clustered near the center of the copy intervals (**Figure S2**).

### 1.11 1000 Genomes Project – C4 copy number analysis

Analysis of *C4* copy number variation across superpopulations showed most individuals across all superpopulations had 4 copies of *C4* overall, 2 copies of *C4A*, and 2 copies of *C4B*, and there were very few individuals with 0 copies of *C4A* or *C4B* (**Figure 2**). Outside of these similarities there were stark differences observed between the superpopulations. The African (AFR) and European (EUR) superpopulations had much higher occurrences of 3 overall copies of *C4*, almost double that observed in the other superpopulations, and much lower occurrences of 5 and 6 overall copies of *C4*. In contract, the South Asian (SAS) superpopulation had the lowest occurrence of 3 overall copies of *C4*, but the highest of 5 and 6. One of the largest differences observed was with *C4L* copy 2 for the AFR superpopulation, which was observed at over double the rate of the other superpopulations; this superpopulation also had substantially lower *C4L* copy 3 occurrence and virtually no occurrence of 4 copies. The *C4S* copy 0 occurrence for the AFR superpopulation was negligible, while other superpopulations were over 20%.

**Figure 2.**
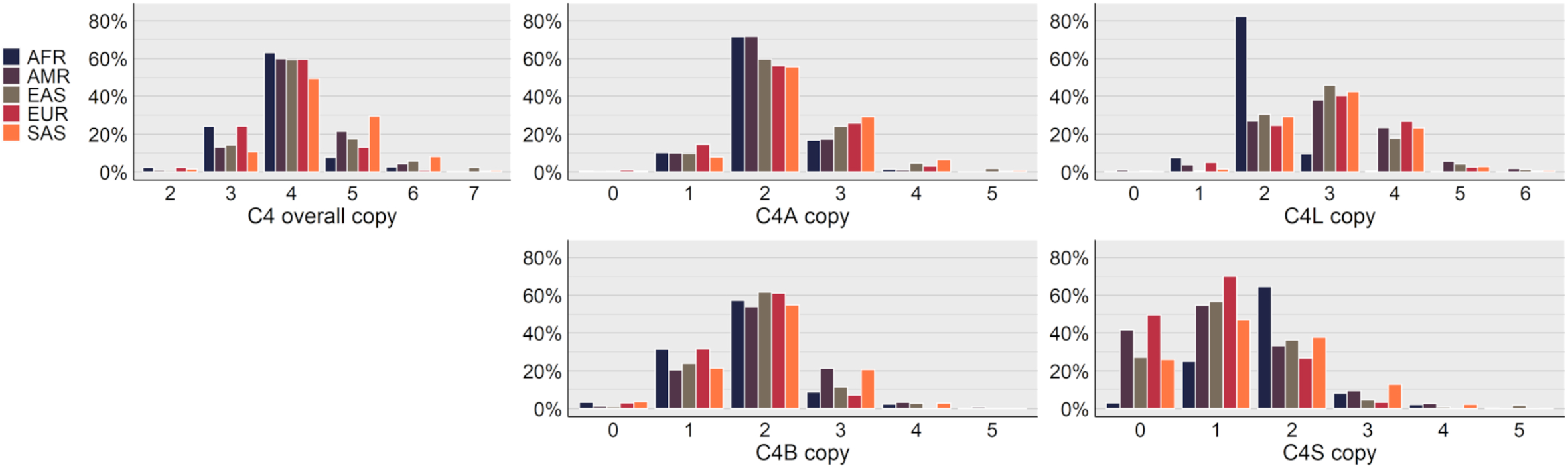
Superpopulation distributions of *C4* copy number results for the 1KGP dataset. *C4* overall copy represents the total copy number of *C4A* and *C4B, C4S* represents the total copy number for the short-forms of *C4A* and *C4B*, and *C4L* represents the total copy number for the long-forms of *C4A* and *C4B*. AFR = African, AMR = Admixed American, EAS = East Asian, EUR = European, SAS = South Asian.

### 1.12 1000 Genomes Project – SNP analysis

The SNP tables output by C4Investigator, which represent combined *C4A* and *C4B* sequence, were parsed to identify sequence variation, and any identified exonic nucleotide variants are evaluated for amino acid coding change. From these results we have summarized non-synonymous mutations in **Table 3**, and SNP variation that is not represented in the main assembly of the GRCh38 reference in **Figure 3**.

**Table 3.** Population specific minor allele frequencies for *C4A* and *C4B* unphased, non-synonymous exonic sequence variants. For this analysis we did not distinguish between *C4A* and *C4B*. This table shows amino acid frequencies, the amino acid position and nucleotide position, the nucleotide frequencies, and population allele frequencies for the minor allele. Major amino acids and nucleotides represent the most frequent global variant while minor amino acids and nucleotides represent the second most frequent variant. This data was filtered to only show variants with allele frequencies >= 2% for any population. Blank values represent absence of the variant. See Table 1 for population abbreviations.

|  | aa<br>pos | 141 | 229 | 325 | 328 | 478 | 549 | 726 | 791 | 916 | 959 | 1286 | 1413 | 1530 |
| --- | --- | --- | --- | --- | --- | --- | --- | --- | --- | --- | --- | --- | --- | --- |
|  | major<br>aa | L | T | K | M | P | H | P | R | R | E | A | A | P |
|  | minor<br>aa | V | I | M | I | L | P | L | H | Q | D | S | P | S |
|  | nuc<br>pos | E3<br>157 | E6<br>60 | E9<br>62 | E9<br>72 | E12<br>92 | E13<br>122 | E17<br>106 | E18<br>103 | E21<br>155 | E23<br>23 | E29<br>180 | E33<br>6 | E36<br>4 |
|  | major<br>nuc | C | C | A | G | C | A | C | G | G | A | G | G | C |
|  | minor<br>nuc | G | T | T | A | T | C | T | A | A | C | T | C | T |
| EUR | GBR | 6.1 |  |  |  |  | 2.6 |  |  |  |  | 7.8 |  |  |
|  | FIN | 8.4 |  |  |  |  | 5.9 |  |  |  | 2.2 | 4.5 |  |  |
|  | IBS | 3.2 |  |  |  |  | 2.1 |  |  |  |  | 4.6 |  |  |
|  | CEU | 6.1 |  |  |  |  | 5.5 |  |  |  |  | 8.2 |  |  |
|  | TSI | 4.1 |  |  |  |  | 4.1 |  |  |  |  | 3.8 |  |  |
| EAS | CHS | 35.8 |  |  |  |  | 14.3 |  |  |  | 8.6 | 3.6 |  |  |
|  | CDX | 53.3 |  |  |  |  | 18.7 |  |  |  | 17.7 |  |  |  |
|  | KHV | 33.7 | 2.1 | 4.4 | 4 |  | 10.2 |  |  |  | 13 |  |  |  |
|  | CHB | 25.8 |  | 2.9 | 2.9 |  | 12.8 |  |  |  | 4.6 | 5.6 |  |  |
|  | JPT | 13.3 |  | 4.4 | 4.2 |  | 8 |  |  |  |  | 13.1 |  |  |
| AMR | PUR | 8.7 |  |  |  |  | 3 |  |  |  |  | 3.1 |  |  |
|  | CLM | 7.7 |  |  |  |  | 3.9 |  |  |  |  | 2.4 |  |  |
|  | PEL | 16 |  |  |  |  | 6.8 |  |  |  | 2.7 |  |  |  |
|  | MXL | 15.8 |  |  |  |  | 7 |  |  |  |  | 2.2 |  |  |
| SAS | PJL | 4 |  |  |  |  | 4 |  |  |  |  | 5.7 |  |  |
|  | BEB | 16.1 |  |  |  |  | 3.8 |  |  |  | 7.4 | 4.8 |  |  |
|  | STU | 8 |  |  |  |  | 4 |  |  |  |  | 4.6 |  |  |
|  | ITU | 8.8 |  |  |  |  | 5.5 |  |  |  | 2.9 | 6.4 |  |  |
|  | GIH | 6.3 |  |  |  |  | 3.1 |  |  |  |  | 5.9 |  |  |
| AFR | ACB | 7 |  |  |  | 3.5 |  |  | 2.2 |  |  |  | 2.2 | 2.2 |
|  | GWD | 5.1 |  |  |  | 3.8 |  |  |  | 3.1 | 2.3 |  | 4.5 |  |
|  | ESN | 10.6 |  |  |  | 4.7 |  |  |  |  | 2.3 |  |  |  |
|  | MSL | 3.5 |  |  |  |  |  |  |  | 4.3 |  |  | 10.2 | 10.2 |
|  | YRI | 10.5 |  |  |  | 4.1 |  |  |  |  | 2.4 |  | 4 | 4 |
|  | LWK | 10.7 |  |  |  |  |  | 3.1 |  | 2.1 |  |  | 2.9 | 3.7 |
|  | ASW | 10.7 |  |  |  |  | 2.5 |  |  |  |  |  | 2.1 | 2.8 |

**Figure 3.**
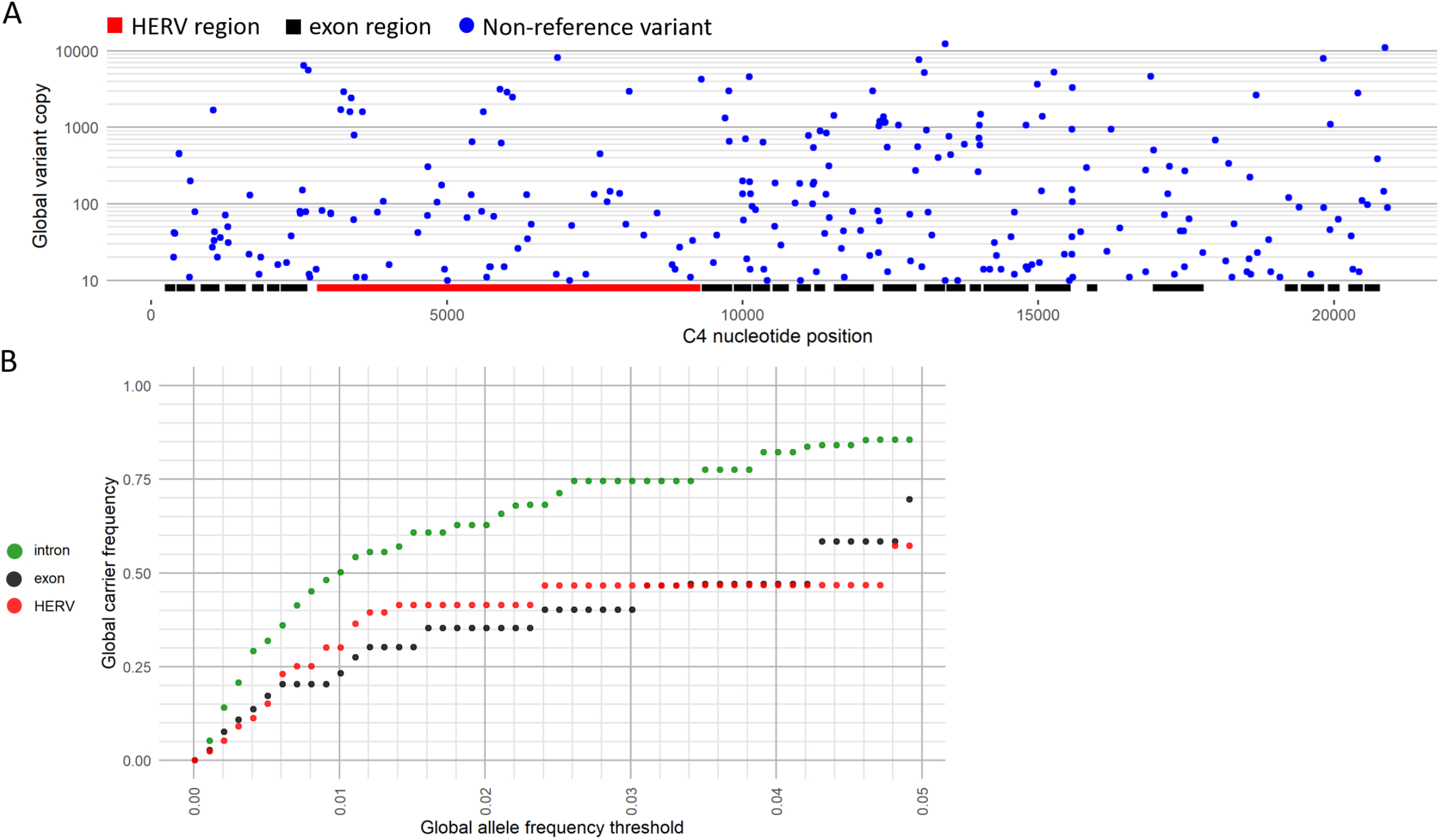
SNV variation across the 1KGP dataset. **(A)** Total copy of combined *C4A* and *C4B* non-reference variants, which are variants not represented in the main assembly of GRCh38, by *C4* position for the 1KGP dataset. The copy number of all non-reference variants for a position across the 1KGP dataset are summed to get the non-reference variant copy, which was then filtered to only show variant positions with total copy of at least 10. Positions of *C4* exon and HERV regions are marked. **(B)** Global carrier frequencies for non-reference variants in the 1KGP dataset for increasing global allele frequency thresholds from 0.00-0.05 for introns, exons, and the HERV region. The y-axis represents the total proportion of carriers that carry a non-reference allele that is at or below the global allele frequency threshold on the x-axis. For example, nearly 25% of the 1KGP dataset carried exonic variants with a global allele frequency of 1% or lower.

Analysis of allele frequencies for *C4A* and *C4B* non-synonymous exonic sequence variation showed large variations in frequencies across populations (**Table 3**). The variant p.H549P was very common in the EAS superpopulation, and was found in most populations, but very rare in the AFR superpopulation. The variant p.L141V was the major allele in the CDX population, was highly frequent across the EAS superpopulation, and was found at appreciable frequencies across all populations. The variants p.T229I, p.K325M, and p.M328I were only found in the EAS superpopulation. And the variants p.P478L, p.P726L, p.R791H, p.R916Q, p.A1413P, and p.P1530S were only found in the AFR superpopulation.

An analysis into non-reference SNVs, which are variants not represented in the main assembly of GRCh38, for the 1KGP dataset across *C4A* and *C4B* showed 251 variant positions with total non-reference variant copy of at least 10 (**Figure 3A, Table S2**). Examination of the positional distribution of these variants across *C4A* and *C4B* showed 50 exonic variant positions accounting for 0.955% of all exonic positions (N=5235), 138 intronic variant positions accounting for 1.56% of all intronic positions (N=8831, exclusive of HERV), and 59 HERV variant positions accounting for 0.927% of all HERV positions (N=6367).

An examination of the proportion of the 1KGP dataset that carry rare variants showed that almost 25% of the samples carried exonic variants with global allele frequencies at or below 1% (**Figure 3B, Table S3**), and about 50% carried intronic variants. Looking at the carrier distribution of more common variants showed that about 70% of the samples carried exonic variants with global allele frequencies below 5%, and about 85% carried intronic variants.

### 1.13 1000 Genomes Project – recombinant analysis

Analysis of carrier frequencies for *C4A*/*C4B* and Rodger/Chido recombinants, *C4A-Ch* and *C4B-Rg*, showed higher overall frequencies of the *C4A-Ch* recombinant compared to *C4B-Rg* (**Table 4**). The *C4A-Ch* recombinant was highly prominent in the AFR superpopulation, with a 37.4% carrier frequency in the MSL population, 20% in GWD and YRI, 14.1% in LWK, 13.5% in ASW, 11.2% in ACB, and 8.1% in ESN. The AMR superpopulation also showed appreciable *C4A-Ch* carrier frequencies, the highest being the PEL population at 7.4%, followed by PUR at 5.8%, MXL at 5.2% and CLM at 4.5%. While carrier frequencies of the *C4B-Rg* recombinant were generally lower overall, with many populations showing no carriers, the frequencies of this recombinant were not negligible, with 8 of the populations displaying at least 4.5% carrier frequency. The AMR and SAS superpopulations showed the highest frequencies of the *C4B-Rg* recombinant, the highest being the STU population at 7.0%, followed by CLM at 6.8%.

**Table 4.** *C4A-Ch* and *C4B-Rg* carrier frequencies by population. Carrier frequencies were calculated by the total *C4A* and *C4B* carrier count per population. *C4A-Ch* = *C4A-Chido, C4B-Rg* = *C4B-Rodger*. See Table 1 for population abbreviations.

|  |  | *C4A-Ch* | *C4B-Rg* | N |
| --- | --- | --- | --- | --- |
| EUR | GBR | 1.1 | 0 | 91 |
|  | FIN | 1.0 | 0 | 99 |
|  | IBS | 1.9 | 6.4 | 157 |
|  | CEU | 1.1 | 1.7 | 179 |
|  | TSI | 0 | 3.7 | 107 |
| EAS | CHS | 0.6 | 0 | 161 |
|  | CDX | 2.2 | 1.1 | 92 |
|  | KHV | 4.9 | 0.8 | 122 |
|  | CHB | 2.9 | 1.9 | 103 |
|  | JPT | 4.8 | 0 | 104 |
| AMR | PUR | 5.8 | 5.0 | 139 |
|  | CLM | 4.5 | 6.8 | 132 |
|  | PEL | 7.4 | 3.3 | 122 |
|  | MXL | 5.2 | 5.2 | 97 |
| SAS | PJL | 2.1 | 4.1 | 146 |
|  | BEB | 0 | 1.5 | 131 |
|  | STU | 0.9 | 7.0 | 114 |
|  | ITU | 0.9 | 4.7 | 107 |
|  | GIH | 1.0 | 4.9 | 103 |
| AFR | ACB | 11.2 | 0 | 116 |
|  | GWD | 20.2 | 4.5 | 178 |
|  | ESN | 8.1 | 0 | 149 |
|  | MSL | 37.4 | 0 | 99 |
|  | YRI | 20.2 | 0 | 178 |
|  | LWK | 14.1 | 0 | 99 |
|  | ASW | 13.5 | 2.7 | 74 |

### 1.14 Performance evaluation – C4A/C4B and Rodger/Chido phasing

Phasing completeness between the *C4A*/*C4B* specific SNP group and the *Rg*/*Ch* specific SNP group was estimated by comparing the number of samples with read-backed phasing for the non-recombinant variants, *C4A*-*Rg* and *C4B*-*Ch*, to the total number of samples carrying *C4A*-*Rg* and *C4B*-*Ch*, respectively. Phasing completeness for *C4A*-*Rg* was 97.69% (N=3167) and *C4B*-*Ch* was 96.60% (N=3113).

## Discussion

Comparison of C4Investigator *C4* copy number determination to ddPCR results showed high concordance between the two methods for *C4A* and *C4B* copy number determination across divergent populations (**Table 2**). *C4(L)* and *C4(S)* copy determination performance was acceptable for the European dataset, but poor for the African dataset.

Comparison of C4Investigator to the *C4A/B* Terra workflow, another bioinformatic pipeline, on the 1KGP WGS dataset showed high concordance between the two workflows, especially for overall *C4* copy. An investigation into discordant *C4A/B* results showed that the discordant samples had lower base quality scores on average (**Figure S1B**), with neither method showing clear copy number results for the discordant samples (**Figure S1A**). In contrast, the investigation into discordant HERV results showed a marked difference between the two methods, with the *C4A/B* Terra workflow showing clear copy numbers for these samples while C4Investigator had unclear determinations (**Figure S2**). This is likely due to the additional structural variant processing of the *C4A/B* Terra workflow, which incorporates Genome STRiP (36), a workflow specifically developed for identifying copy number variation in WGS data. The *C4A/B* Terra is strictly focused on identifying copy number variation, a task that it appears to perform very well. In contrast, C4Investigator takes a different approach, focusing on identifying nucleotide variants in a copy variable system through the utilization of custom alignment processing algorithms, which has enabled the identification and quantification of SNP variation across the *C4* genes.

An analysis into C4 copy number variation between superpopulations (**Figure 2**) demonstrated some specific patterns, such as a median overall *C4* copy number of 4, and a median copy number for *C4A* and *C4B* of 2 each, but also important distinctions between populations, such as the strikingly high number of *C4L* copy 2 genotypes in the AFR superpopulation, and the general imbalance between overall *C4* copy of 3 and 5, which was unique for each superpopulation. Differences of this nature might suggest evolutionary pressure or unique genomic makeups that are specific to the different superpopulations and modulate the fitness of different *C4* gene structures.

An essential innovation of C4Investigator is demonstrated by its capacity to reveal important differences in sequence variation between populations, with likely important functional implications. An analysis of non-synonymous exonic sequence variants demonstrated that *C4* sequence makeup can differ greatly between populations, with some variants with seemingly rare global allele frequencies showing high allele frequencies in specific populations. For example, the p.A1413P and the p.P1530S mutations were absent in most populations, but both had 10.2% allele frequency in the MSL population (**Table 3**). The fact that both mutations have the same allele frequency raises the question of if these mutations are in-phase, unfortunately, there is a 2046bp gap between these variants which was outside the scope of our phasing approach. However, an examination of the individuals that carried each mutation showed a high overlap, where 28 individuals carried both mutations compared to total 33 individuals carrying the p.A1413P mutation and 31 individuals carrying the p.P1530S mutation. A structural interrogation of C4·MASP-2 binding shows the p.A1413P mutation occurs in the middle of a MASP-2 exosite(31) (**Figure 1**), while the change from alanine to proline would not likely change the electrostatic interactions between C4 and MASP-2, it could potentially alter the structure of the binding site. Another sequence variant with potential to impact function is the p.P478L mutation, which causes severe reduction of hemolytic activity by disruption of C5 binding(28). Similar analyses in the context of disease association studies are likely to reveal important insights into immune-mediated pathogenesis.

An analysis into *C4A* and *C4B* non-reference variants demonstrated that the *C4* genes are highly poly-allelic across introns, exons and the HERV region (**Figure 3A**). Further examination into rare variant carrier frequencies demonstrated that exonic variants under 5% global allele frequency are carried by around 70% of the 1KGP samples (**Figure 3B**). This analysis demonstrates the value of nucleotide level analysis of *C4*, which reveals important features of genomic variation not otherwise evident with existing methods.

One important aspect of SNP variation identification is the ability to phase variants. However, phasing high-copy variants (gene copy number > 2) is very complex and it is difficult to be certain of phasing completeness due to the high potential for missing information. Due to the high sequence similarity between *C4A* and *C4B*, the alignments must be treated as a single gene, exacerbating the high-copy phasing problem. We have implemented read-backed phasing that enables us to determine whether two variants in proximity are in-phase, but the potential for missing information means in many cases we cannot make the determination that two variants are *not* in-phase; essentially, we can make more confident true positive phasing calls than true negative. Because of the distance between the *C4A*/*C4B* SNPs and the *Rg*/*Ch* SNPs, 440bp, we can determine presence of recombinants between the two SNP groups. An estimate of phasing completeness between *C4A-Rg* and *C4B-Ch* showed this phasing approach only missed a small percentage of samples. Utilization of this phasing approach to identify *C4A-Ch* and *C4B-Rg* recombinants showed high *C4A-Ch* carrier frequencies across the AFR superpopulation (**Table 4**), and appreciable carrier frequencies for the *C4B-Rg* recombinant and the AMR and SAS superpopulations.

In conclusion, C4Investigator fills a critical role in the investigation of *C4* variation, processing WGS data to provide *C4* copy number variation and full genomic sequence information. Here, we have demonstrated the utility of this workflow on the Thousand Genomes Project dataset, revealing that *C4* copy number varies between superpopulations, that alleles with low global allele frequencies can have high population specific frequencies, the presence and distribution of *C4* recombinant variants, and population specific carrier frequencies for rare alleles. Additionally, we have demonstrated that C4Investigator can identify *C4* variation that is known to alter C4 function. To the best of our knowledge, C4Investigator is the only bioinformatic workflow currently available for nucleotide level characterization of *C4* from WGS data, and as such, promises to contribute to our understanding of the role of this genomic region in human health and disease.

## Acknowledgments and funding

We would like to thank Michael Wilson and Mark Seielstad for constructive comments. We would like to acknowledge that this work exists as a chapter of Wesley Marin’s doctoral dissertation (44). This work was supported by the National Institutes of Health (NIH-R01AI128775). The funders had no roles in study design, data collection and analysis, decision to publish, or preparation of the manuscript.

## Author Contributions

**Conceptualization:** WMM, DGA, JAH

**Data Curation:** WMM, DGA

**Formal analysis:** WMM, KJW

**Funding acquisition:** JAH

**Investigation:** WMM

**Methodology:** WMM

**Project Administration:** JAH

**Resources:** DGA, JAH

**Software:** WMM

**Supervision:** JAH

**Validation:** WMM, DGA

**Visualization:** WMM

**Writing – Original Draft Preparation:** WMM

**Writing – Review & Editing:** WMM, DGA, JAH, KJW

## Data Availability Statement

The datasets analyzed for this study can be found in the International Genome Sample Resource data portal at https://www.internationalgenome.org/data. The C4Investigator workflow is available at https://github.com/Hollenbach-lab/C4Investigator. And the scripts used to analyze the data are available at https://github.com/wesleymarin/C4investigator_scripts.

