## Supplemental Table 1 for "High-throughput complement component 4 genomic sequence analysis with C4Investigator"

Table_S1_suppInfo

| 1000Genomes project accession resource. 1000Genomes project data is downloaded from: http://s3.amazonaws.com/1000genomes | |
|---|---|
| **Accession Number** | URL Suffix |
| **NA06985** | 1000G_2504_high_coverage/data/ERR3239276/NA06985.final.cram |
| **NA06986** | 1000G_2504_high_coverage/data/ERR3239277/NA06986.final.cram |
| **NA06994** | 1000G_2504_high_coverage/data/ERR3239278/NA06994.final.cram |
| **NA07000** | 1000G_2504_high_coverage/data/ERR3239279/NA07000.final.cram |
| **NA07037** | 1000G_2504_high_coverage/data/ERR3239280/NA07037.final.cram |
| **NA07051** | 1000G_2504_high_coverage/data/ERR3239281/NA07051.final.cram |
| **NA07347** | 1000G_2504_high_coverage/data/ERR3239282/NA07347.final.cram |
| **NA07357** | 1000G_2504_high_coverage/data/ERR3239283/NA07357.final.cram |
| **NA10847** | 1000G_2504_high_coverage/data/ERR3239284/NA10847.final.cram |
| **NA10851** | 1000G_2504_high_coverage/data/ERR3239285/NA10851.final.cram |
| **NA11829** | 1000G_2504_high_coverage/data/ERR3239286/NA11829.final.cram |
| **NA11830** | 1000G_2504_high_coverage/data/ERR3239287/NA11830.final.cram |
| **NA11831** | 1000G_2504_high_coverage/data/ERR3239288/NA11831.final.cram |
| **NA11832** | 1000G_2504_high_coverage/data/ERR3239289/NA11832.final.cram |
| **NA11840** | 1000G_2504_high_coverage/data/ERR3239290/NA11840.final.cram |
| **NA11881** | 1000G_2504_high_coverage/data/ERR3239291/NA11881.final.cram |
| **NA11894** | 1000G_2504_high_coverage/data/ERR3239292/NA11894.final.cram |
| **NA11918** | 1000G_2504_high_coverage/data/ERR3239293/NA11918.final.cram |
| **NA11919** | 1000G_2504_high_coverage/data/ERR3239294/NA11919.final.cram |
| **NA11920** | 1000G_2504_high_coverage/data/ERR3239295/NA11920.final.cram |
| **NA11931** | 1000G_2504_high_coverage/data/ERR3239296/NA11931.final.cram |
| **NA11992** | 1000G_2504_high_coverage/data/ERR3239297/NA11992.final.cram |
| **NA11994** | 1000G_2504_high_coverage/data/ERR3239298/NA11994.final.cram |
| **NA11995** | 1000G_2504_high_coverage/data/ERR3239299/NA11995.final.cram |
| **NA12003** | 1000G_2504_high_coverage/data/ERR3239300/NA12003.final.cram |
| **NA12004** | 1000G_2504_high_coverage/data/ERR3239301/NA12004.final.cram |
| **NA12005** | 1000G_2504_high_coverage/data/ERR3239302/NA12005.final.cram |
| **NA12006** | 1000G_2504_high_coverage/data/ERR3239303/NA12006.final.cram |
| **NA12043** | 1000G_2504_high_coverage/data/ERR3239304/NA12043.final.cram |
| **NA12044** | 1000G_2504_high_coverage/data/ERR3239305/NA12044.final.cram |
| **NA12045** | 1000G_2504_high_coverage/data/ERR3239306/NA12045.final.cram |
| **NA12144** | 1000G_2504_high_coverage/data/ERR3239307/NA12144.final.cram |
| **NA12154** | 1000G_2504_high_coverage/data/ERR3239308/NA12154.final.cram |
| **NA12155** | 1000G_2504_high_coverage/data/ERR3239309/NA12155.final.cram |
| **NA12156** | 1000G_2504_high_coverage/data/ERR3239310/NA12156.final.cram |
| **NA12234** | 1000G_2504_high_coverage/data/ERR3239311/NA12234.final.cram |
| **NA12249** | 1000G_2504_high_coverage/data/ERR3239312/NA12249.final.cram |
| **NA12287** | 1000G_2504_high_coverage/data/ERR3239313/NA12287.final.cram |
| **NA12414** | 1000G_2504_high_coverage/data/ERR3239314/NA12414.final.cram |
| **NA12489** | 1000G_2504_high_coverage/data/ERR3239315/NA12489.final.cram |
| **NA12716** | 1000G_2504_high_coverage/data/ERR3239316/NA12716.final.cram |
| **NA12717** | 1000G_2504_high_coverage/data/ERR3239317/NA12717.final.cram |
| **NA12749** | 1000G_2504_high_coverage/data/ERR3239318/NA12749.final.cram |
| **NA12750** | 1000G_2504_high_coverage/data/ERR3239319/NA12750.final.cram |
| **NA12751** | 1000G_2504_high_coverage/data/ERR3239320/NA12751.final.cram |
| **NA12760** | 1000G_2504_high_coverage/data/ERR3239321/NA12760.final.cram |
| **NA12761** | 1000G_2504_high_coverage/data/ERR3239322/NA12761.final.cram |
| **NA12762** | 1000G_2504_high_coverage/data/ERR3239323/NA12762.final.cram |
| **NA12763** | 1000G_2504_high_coverage/data/ERR3239324/NA12763.final.cram |
| **NA12776** | 1000G_2504_high_coverage/data/ERR3239325/NA12776.final.cram |
| **NA12812** | 1000G_2504_high_coverage/data/ERR3239326/NA12812.final.cram |
| **NA12813** | 1000G_2504_high_coverage/data/ERR3239327/NA12813.final.cram |
| **NA12814** | 1000G_2504_high_coverage/data/ERR3239328/NA12814.final.cram |
| **NA12815** | 1000G_2504_high_coverage/data/ERR3239329/NA12815.final.cram |
| **NA12828** | 1000G_2504_high_coverage/data/ERR3239330/NA12828.final.cram |
| **NA12872** | 1000G_2504_high_coverage/data/ERR3239331/NA12872.final.cram |
| **NA12873** | 1000G_2504_high_coverage/data/ERR3239332/NA12873.final.cram |
| **NA12874** | 1000G_2504_high_coverage/data/ERR3239333/NA12874.final.cram |
| **NA12878** | 1000G_2504_high_coverage/data/ERR3239334/NA12878.final.cram |
| **NA18486** | 1000G_2504_high_coverage/data/ERR3239335/NA18486.final.cram |
| **NA18489** | 1000G_2504_high_coverage/data/ERR3239336/NA18489.final.cram |
| **NA18498** | 1000G_2504_high_coverage/data/ERR3239337/NA18498.final.cram |

| | |
|---|---|
| **NA18499** | 1000G_2504_high_coverage/data/ERR3239338/NA18499.final.cram |
| **NA18501** | 1000G_2504_high_coverage/data/ERR3239339/NA18501.final.cram |
| **NA18502** | 1000G_2504_high_coverage/data/ERR3239340/NA18502.final.cram |
| **NA18504** | 1000G_2504_high_coverage/data/ERR3239341/NA18504.final.cram |
| **NA18505** | 1000G_2504_high_coverage/data/ERR3239342/NA18505.final.cram |
| **NA18507** | 1000G_2504_high_coverage/data/ERR3239343/NA18507.final.cram |
| **NA18508** | 1000G_2504_high_coverage/data/ERR3239344/NA18508.final.cram |
| **NA18510** | 1000G_2504_high_coverage/data/ERR3239345/NA18510.final.cram |
| **NA18511** | 1000G_2504_high_coverage/data/ERR3239346/NA18511.final.cram |
| **NA18516** | 1000G_2504_high_coverage/data/ERR3239347/NA18516.final.cram |
| **NA18517** | 1000G_2504_high_coverage/data/ERR3239348/NA18517.final.cram |
| **NA18519** | 1000G_2504_high_coverage/data/ERR3239349/NA18519.final.cram |
| **NA18520** | 1000G_2504_high_coverage/data/ERR3239350/NA18520.final.cram |
| **NA18522** | 1000G_2504_high_coverage/data/ERR3239351/NA18522.final.cram |
| **NA18523** | 1000G_2504_high_coverage/data/ERR3239352/NA18523.final.cram |
| **NA18526** | 1000G_2504_high_coverage/data/ERR3239353/NA18526.final.cram |
| **NA18532** | 1000G_2504_high_coverage/data/ERR3239354/NA18532.final.cram |
| **NA18537** | 1000G_2504_high_coverage/data/ERR3239355/NA18537.final.cram |
| **NA18542** | 1000G_2504_high_coverage/data/ERR3239356/NA18542.final.cram |
| **NA18545** | 1000G_2504_high_coverage/data/ERR3239357/NA18545.final.cram |
| **NA18547** | 1000G_2504_high_coverage/data/ERR3239358/NA18547.final.cram |
| **NA18550** | 1000G_2504_high_coverage/data/ERR3239359/NA18550.final.cram |
| **NA18552** | 1000G_2504_high_coverage/data/ERR3239360/NA18552.final.cram |
| **NA18555** | 1000G_2504_high_coverage/data/ERR3239361/NA18555.final.cram |
| **NA18558** | 1000G_2504_high_coverage/data/ERR3239362/NA18558.final.cram |
| **NA18561** | 1000G_2504_high_coverage/data/ERR3239363/NA18561.final.cram |
| **NA18562** | 1000G_2504_high_coverage/data/ERR3239364/NA18562.final.cram |
| **NA18563** | 1000G_2504_high_coverage/data/ERR3239365/NA18563.final.cram |
| **NA18564** | 1000G_2504_high_coverage/data/ERR3239366/NA18564.final.cram |
| **NA18566** | 1000G_2504_high_coverage/data/ERR3239367/NA18566.final.cram |
| **NA18570** | 1000G_2504_high_coverage/data/ERR3239368/NA18570.final.cram |
| **NA18571** | 1000G_2504_high_coverage/data/ERR3239369/NA18571.final.cram |
| **NA18572** | 1000G_2504_high_coverage/data/ERR3239370/NA18572.final.cram |
| **NA18573** | 1000G_2504_high_coverage/data/ERR3239371/NA18573.final.cram |
| **NA18577** | 1000G_2504_high_coverage/data/ERR3239372/NA18577.final.cram |
| **NA18579** | 1000G_2504_high_coverage/data/ERR3239373/NA18579.final.cram |
| **NA18582** | 1000G_2504_high_coverage/data/ERR3239374/NA18582.final.cram |
| **NA18592** | 1000G_2504_high_coverage/data/ERR3239375/NA18592.final.cram |
| **NA18593** | 1000G_2504_high_coverage/data/ERR3239376/NA18593.final.cram |
| **NA18603** | 1000G_2504_high_coverage/data/ERR3239377/NA18603.final.cram |
| **NA18605** | 1000G_2504_high_coverage/data/ERR3239378/NA18605.final.cram |
| **NA18608** | 1000G_2504_high_coverage/data/ERR3239379/NA18608.final.cram |
| **NA18609** | 1000G_2504_high_coverage/data/ERR3239380/NA18609.final.cram |
| **NA18638** | 1000G_2504_high_coverage/data/ERR3239381/NA18638.final.cram |
| **NA18853** | 1000G_2504_high_coverage/data/ERR3239382/NA18853.final.cram |
| **NA18856** | 1000G_2504_high_coverage/data/ERR3239383/NA18856.final.cram |
| **NA18858** | 1000G_2504_high_coverage/data/ERR3239384/NA18858.final.cram |
| **NA18861** | 1000G_2504_high_coverage/data/ERR3239385/NA18861.final.cram |
| **NA18870** | 1000G_2504_high_coverage/data/ERR3239386/NA18870.final.cram |
| **NA18871** | 1000G_2504_high_coverage/data/ERR3239387/NA18871.final.cram |
| **NA18907** | 1000G_2504_high_coverage/data/ERR3239388/NA18907.final.cram |
| **NA18909** | 1000G_2504_high_coverage/data/ERR3239389/NA18909.final.cram |
| **NA18912** | 1000G_2504_high_coverage/data/ERR3239390/NA18912.final.cram |
| **NA18916** | 1000G_2504_high_coverage/data/ERR3239391/NA18916.final.cram |
| **NA18940** | 1000G_2504_high_coverage/data/ERR3239392/NA18940.final.cram |
| **NA18942** | 1000G_2504_high_coverage/data/ERR3239393/NA18942.final.cram |
| **NA18943** | 1000G_2504_high_coverage/data/ERR3239394/NA18943.final.cram |
| **NA18944** | 1000G_2504_high_coverage/data/ERR3239395/NA18944.final.cram |
| **NA18945** | 1000G_2504_high_coverage/data/ERR3239396/NA18945.final.cram |
| **NA18947** | 1000G_2504_high_coverage/data/ERR3239397/NA18947.final.cram |
| **NA18948** | 1000G_2504_high_coverage/data/ERR3239398/NA18948.final.cram |
| **NA18949** | 1000G_2504_high_coverage/data/ERR3239399/NA18949.final.cram |
| **NA18951** | 1000G_2504_high_coverage/data/ERR3239400/NA18951.final.cram |
| **NA18952** | 1000G_2504_high_coverage/data/ERR3239401/NA18952.final.cram |
| **NA18953** | 1000G_2504_high_coverage/data/ERR3239402/NA18953.final.cram |

| | |
|---|---|
| **NA18956** | 1000G_2504_high_coverage/data/ERR3239403/NA18956.final.cram |
| **NA18959** | 1000G_2504_high_coverage/data/ERR3239404/NA18959.final.cram |
| **NA18960** | 1000G_2504_high_coverage/data/ERR3239405/NA18960.final.cram |
| **NA18961** | 1000G_2504_high_coverage/data/ERR3239406/NA18961.final.cram |
| **NA18964** | 1000G_2504_high_coverage/data/ERR3239407/NA18964.final.cram |
| **NA18965** | 1000G_2504_high_coverage/data/ERR3239408/NA18965.final.cram |
| **NA18967** | 1000G_2504_high_coverage/data/ERR3239409/NA18967.final.cram |
| **NA18968** | 1000G_2504_high_coverage/data/ERR3239410/NA18968.final.cram |
| **NA18969** | 1000G_2504_high_coverage/data/ERR3239411/NA18969.final.cram |
| **NA18970** | 1000G_2504_high_coverage/data/ERR3239412/NA18970.final.cram |
| **NA18971** | 1000G_2504_high_coverage/data/ERR3239413/NA18971.final.cram |
| **NA18972** | 1000G_2504_high_coverage/data/ERR3239414/NA18972.final.cram |
| **NA18973** | 1000G_2504_high_coverage/data/ERR3239415/NA18973.final.cram |
| **NA18974** | 1000G_2504_high_coverage/data/ERR3239416/NA18974.final.cram |
| **NA18975** | 1000G_2504_high_coverage/data/ERR3239417/NA18975.final.cram |
| **NA18976** | 1000G_2504_high_coverage/data/ERR3239418/NA18976.final.cram |
| **NA18980** | 1000G_2504_high_coverage/data/ERR3239419/NA18980.final.cram |
| **NA18981** | 1000G_2504_high_coverage/data/ERR3239420/NA18981.final.cram |
| **NA19005** | 1000G_2504_high_coverage/data/ERR3239421/NA19005.final.cram |
| **NA19093** | 1000G_2504_high_coverage/data/ERR3239422/NA19093.final.cram |
| **NA19098** | 1000G_2504_high_coverage/data/ERR3239423/NA19098.final.cram |
| **NA19099** | 1000G_2504_high_coverage/data/ERR3239424/NA19099.final.cram |
| **NA19102** | 1000G_2504_high_coverage/data/ERR3239425/NA19102.final.cram |
| **NA19108** | 1000G_2504_high_coverage/data/ERR3239426/NA19108.final.cram |
| **NA19114** | 1000G_2504_high_coverage/data/ERR3239427/NA19114.final.cram |
| **NA19116** | 1000G_2504_high_coverage/data/ERR3239428/NA19116.final.cram |
| **NA19119** | 1000G_2504_high_coverage/data/ERR3239429/NA19119.final.cram |
| **NA19129** | 1000G_2504_high_coverage/data/ERR3239430/NA19129.final.cram |
| **NA19131** | 1000G_2504_high_coverage/data/ERR3239431/NA19131.final.cram |
| **NA19137** | 1000G_2504_high_coverage/data/ERR3239432/NA19137.final.cram |
| **NA19138** | 1000G_2504_high_coverage/data/ERR3239433/NA19138.final.cram |
| **NA19141** | 1000G_2504_high_coverage/data/ERR3239434/NA19141.final.cram |
| **NA19143** | 1000G_2504_high_coverage/data/ERR3239435/NA19143.final.cram |
| **NA19144** | 1000G_2504_high_coverage/data/ERR3239436/NA19144.final.cram |
| **NA19147** | 1000G_2504_high_coverage/data/ERR3239437/NA19147.final.cram |
| **NA19152** | 1000G_2504_high_coverage/data/ERR3239438/NA19152.final.cram |
| **NA19153** | 1000G_2504_high_coverage/data/ERR3239439/NA19153.final.cram |
| **NA19159** | 1000G_2504_high_coverage/data/ERR3239440/NA19159.final.cram |
| **NA19160** | 1000G_2504_high_coverage/data/ERR3239441/NA19160.final.cram |
| **NA19171** | 1000G_2504_high_coverage/data/ERR3239442/NA19171.final.cram |
| **NA19172** | 1000G_2504_high_coverage/data/ERR3239443/NA19172.final.cram |
| **NA19190** | 1000G_2504_high_coverage/data/ERR3239444/NA19190.final.cram |
| **NA19200** | 1000G_2504_high_coverage/data/ERR3239445/NA19200.final.cram |
| **NA19201** | 1000G_2504_high_coverage/data/ERR3239446/NA19201.final.cram |
| **NA19204** | 1000G_2504_high_coverage/data/ERR3239447/NA19204.final.cram |
| **NA19206** | 1000G_2504_high_coverage/data/ERR3239448/NA19206.final.cram |
| **NA19207** | 1000G_2504_high_coverage/data/ERR3239449/NA19207.final.cram |
| **NA19209** | 1000G_2504_high_coverage/data/ERR3239450/NA19209.final.cram |
| **NA19210** | 1000G_2504_high_coverage/data/ERR3239451/NA19210.final.cram |
| **NA19225** | 1000G_2504_high_coverage/data/ERR3239452/NA19225.final.cram |
| **NA19238** | 1000G_2504_high_coverage/data/ERR3239453/NA19238.final.cram |
| **NA19239** | 1000G_2504_high_coverage/data/ERR3239454/NA19239.final.cram |
| **NA19257** | 1000G_2504_high_coverage/data/ERR3239455/NA19257.final.cram |
| **NA07056** | 1000G_2504_high_coverage/data/ERR3239456/NA07056.final.cram |
| **NA18614** | 1000G_2504_high_coverage/data/ERR3239457/NA18614.final.cram |
| **NA06984** | 1000G_2504_high_coverage/data/ERR3239458/NA06984.final.cram |
| **NA06989** | 1000G_2504_high_coverage/data/ERR3239459/NA06989.final.cram |
| **NA11843** | 1000G_2504_high_coverage/data/ERR3239460/NA11843.final.cram |
| **NA11892** | 1000G_2504_high_coverage/data/ERR3239461/NA11892.final.cram |
| **NA11893** | 1000G_2504_high_coverage/data/ERR3239462/NA11893.final.cram |
| **NA11930** | 1000G_2504_high_coverage/data/ERR3239463/NA11930.final.cram |
| **NA12058** | 1000G_2504_high_coverage/data/ERR3239464/NA12058.final.cram |
| **NA12272** | 1000G_2504_high_coverage/data/ERR3239465/NA12272.final.cram |
| **NA12273** | 1000G_2504_high_coverage/data/ERR3239466/NA12273.final.cram |
| **NA12275** | 1000G_2504_high_coverage/data/ERR3239467/NA12275.final.cram |

| | |
|---|---|
| **NA12282** | 1000G_2504_high_coverage/data/ERR3239468/NA12282.final.cram |
| **NA12283** | 1000G_2504_high_coverage/data/ERR3239469/NA12283.final.cram |
| **NA12286** | 1000G_2504_high_coverage/data/ERR3239470/NA12286.final.cram |
| **NA12340** | 1000G_2504_high_coverage/data/ERR3239471/NA12340.final.cram |
| **NA12341** | 1000G_2504_high_coverage/data/ERR3239472/NA12341.final.cram |
| **NA12342** | 1000G_2504_high_coverage/data/ERR3239473/NA12342.final.cram |
| **NA12347** | 1000G_2504_high_coverage/data/ERR3239474/NA12347.final.cram |
| **NA12348** | 1000G_2504_high_coverage/data/ERR3239475/NA12348.final.cram |
| **NA12383** | 1000G_2504_high_coverage/data/ERR3239476/NA12383.final.cram |
| **NA12399** | 1000G_2504_high_coverage/data/ERR3239477/NA12399.final.cram |
| **NA12400** | 1000G_2504_high_coverage/data/ERR3239478/NA12400.final.cram |
| **NA12413** | 1000G_2504_high_coverage/data/ERR3239479/NA12413.final.cram |
| **NA12718** | 1000G_2504_high_coverage/data/ERR3239480/NA12718.final.cram |
| **NA12748** | 1000G_2504_high_coverage/data/ERR3239481/NA12748.final.cram |
| **NA12775** | 1000G_2504_high_coverage/data/ERR3239482/NA12775.final.cram |
| **NA12777** | 1000G_2504_high_coverage/data/ERR3239483/NA12777.final.cram |
| **NA12778** | 1000G_2504_high_coverage/data/ERR3239484/NA12778.final.cram |
| **NA12827** | 1000G_2504_high_coverage/data/ERR3239485/NA12827.final.cram |
| **NA12829** | 1000G_2504_high_coverage/data/ERR3239486/NA12829.final.cram |
| **NA12842** | 1000G_2504_high_coverage/data/ERR3239487/NA12842.final.cram |
| **NA12843** | 1000G_2504_high_coverage/data/ERR3239488/NA12843.final.cram |
| **NA12889** | 1000G_2504_high_coverage/data/ERR3239489/NA12889.final.cram |
| **NA12890** | 1000G_2504_high_coverage/data/ERR3239490/NA12890.final.cram |
| **NA18488** | 1000G_2504_high_coverage/data/ERR3239491/NA18488.final.cram |
| **NA18530** | 1000G_2504_high_coverage/data/ERR3239492/NA18530.final.cram |
| **NA18534** | 1000G_2504_high_coverage/data/ERR3239493/NA18534.final.cram |
| **NA18536** | 1000G_2504_high_coverage/data/ERR3239494/NA18536.final.cram |
| **NA18543** | 1000G_2504_high_coverage/data/ERR3239495/NA18543.final.cram |
| **NA18544** | 1000G_2504_high_coverage/data/ERR3239496/NA18544.final.cram |
| **NA18546** | 1000G_2504_high_coverage/data/ERR3239497/NA18546.final.cram |
| **NA18548** | 1000G_2504_high_coverage/data/ERR3239498/NA18548.final.cram |
| **NA18549** | 1000G_2504_high_coverage/data/ERR3239499/NA18549.final.cram |
| **NA18557** | 1000G_2504_high_coverage/data/ERR3239500/NA18557.final.cram |
| **NA18559** | 1000G_2504_high_coverage/data/ERR3239501/NA18559.final.cram |
| **NA18595** | 1000G_2504_high_coverage/data/ERR3239502/NA18595.final.cram |
| **NA18596** | 1000G_2504_high_coverage/data/ERR3239503/NA18596.final.cram |
| **NA18597** | 1000G_2504_high_coverage/data/ERR3239504/NA18597.final.cram |
| **NA18599** | 1000G_2504_high_coverage/data/ERR3239505/NA18599.final.cram |
| **NA18602** | 1000G_2504_high_coverage/data/ERR3239506/NA18602.final.cram |
| **NA18606** | 1000G_2504_high_coverage/data/ERR3239507/NA18606.final.cram |
| **NA18610** | 1000G_2504_high_coverage/data/ERR3239508/NA18610.final.cram |
| **NA18611** | 1000G_2504_high_coverage/data/ERR3239509/NA18611.final.cram |
| **NA18612** | 1000G_2504_high_coverage/data/ERR3239510/NA18612.final.cram |
| **NA18613** | 1000G_2504_high_coverage/data/ERR3239511/NA18613.final.cram |
| **NA18615** | 1000G_2504_high_coverage/data/ERR3239512/NA18615.final.cram |
| **NA18616** | 1000G_2504_high_coverage/data/ERR3239513/NA18616.final.cram |
| **NA18617** | 1000G_2504_high_coverage/data/ERR3239514/NA18617.final.cram |
| **NA18618** | 1000G_2504_high_coverage/data/ERR3239515/NA18618.final.cram |
| **NA18619** | 1000G_2504_high_coverage/data/ERR3239516/NA18619.final.cram |
| **NA18620** | 1000G_2504_high_coverage/data/ERR3239517/NA18620.final.cram |
| **NA18621** | 1000G_2504_high_coverage/data/ERR3239518/NA18621.final.cram |
| **NA18622** | 1000G_2504_high_coverage/data/ERR3239519/NA18622.final.cram |
| **NA18623** | 1000G_2504_high_coverage/data/ERR3239520/NA18623.final.cram |
| **NA18624** | 1000G_2504_high_coverage/data/ERR3239521/NA18624.final.cram |
| **NA18625** | 1000G_2504_high_coverage/data/ERR3239522/NA18625.final.cram |
| **NA18626** | 1000G_2504_high_coverage/data/ERR3239523/NA18626.final.cram |
| **NA18627** | 1000G_2504_high_coverage/data/ERR3239524/NA18627.final.cram |
| **NA18628** | 1000G_2504_high_coverage/data/ERR3239525/NA18628.final.cram |
| **NA18630** | 1000G_2504_high_coverage/data/ERR3239526/NA18630.final.cram |
| **NA18631** | 1000G_2504_high_coverage/data/ERR3239527/NA18631.final.cram |
| **NA18632** | 1000G_2504_high_coverage/data/ERR3239528/NA18632.final.cram |
| **NA18633** | 1000G_2504_high_coverage/data/ERR3239529/NA18633.final.cram |
| **NA18634** | 1000G_2504_high_coverage/data/ERR3239530/NA18634.final.cram |
| **NA18635** | 1000G_2504_high_coverage/data/ERR3239531/NA18635.final.cram |
| **NA18636** | 1000G_2504_high_coverage/data/ERR3239532/NA18636.final.cram |

| | |
|---|---|
| **NA18637** | 1000G_2504_high_coverage/data/ERR3239533/NA18637.final.cram |
| **NA18639** | 1000G_2504_high_coverage/data/ERR3239534/NA18639.final.cram |
| **NA18640** | 1000G_2504_high_coverage/data/ERR3239535/NA18640.final.cram |
| **NA18641** | 1000G_2504_high_coverage/data/ERR3239536/NA18641.final.cram |
| **NA18642** | 1000G_2504_high_coverage/data/ERR3239537/NA18642.final.cram |
| **NA18643** | 1000G_2504_high_coverage/data/ERR3239538/NA18643.final.cram |
| **NA18645** | 1000G_2504_high_coverage/data/ERR3239539/NA18645.final.cram |
| **NA18647** | 1000G_2504_high_coverage/data/ERR3239540/NA18647.final.cram |
| **NA18740** | 1000G_2504_high_coverage/data/ERR3239541/NA18740.final.cram |
| **NA18745** | 1000G_2504_high_coverage/data/ERR3239542/NA18745.final.cram |
| **NA18747** | 1000G_2504_high_coverage/data/ERR3239543/NA18747.final.cram |
| **NA18748** | 1000G_2504_high_coverage/data/ERR3239544/NA18748.final.cram |
| **NA18749** | 1000G_2504_high_coverage/data/ERR3239545/NA18749.final.cram |
| **NA18757** | 1000G_2504_high_coverage/data/ERR3239546/NA18757.final.cram |
| **NA18867** | 1000G_2504_high_coverage/data/ERR3239547/NA18867.final.cram |
| **NA18868** | 1000G_2504_high_coverage/data/ERR3239548/NA18868.final.cram |
| **NA18873** | 1000G_2504_high_coverage/data/ERR3239549/NA18873.final.cram |
| **NA18908** | 1000G_2504_high_coverage/data/ERR3239550/NA18908.final.cram |
| **NA18910** | 1000G_2504_high_coverage/data/ERR3239551/NA18910.final.cram |
| **NA18917** | 1000G_2504_high_coverage/data/ERR3239552/NA18917.final.cram |
| **NA18923** | 1000G_2504_high_coverage/data/ERR3239553/NA18923.final.cram |
| **NA18924** | 1000G_2504_high_coverage/data/ERR3239554/NA18924.final.cram |
| **NA18933** | 1000G_2504_high_coverage/data/ERR3239555/NA18933.final.cram |
| **NA18934** | 1000G_2504_high_coverage/data/ERR3239556/NA18934.final.cram |
| **NA18939** | 1000G_2504_high_coverage/data/ERR3239557/NA18939.final.cram |
| **NA18941** | 1000G_2504_high_coverage/data/ERR3239558/NA18941.final.cram |
| **NA18946** | 1000G_2504_high_coverage/data/ERR3239559/NA18946.final.cram |
| **NA18954** | 1000G_2504_high_coverage/data/ERR3239560/NA18954.final.cram |
| **NA18957** | 1000G_2504_high_coverage/data/ERR3239561/NA18957.final.cram |
| **NA18962** | 1000G_2504_high_coverage/data/ERR3239562/NA18962.final.cram |
| **NA18963** | 1000G_2504_high_coverage/data/ERR3239563/NA18963.final.cram |
| **NA18966** | 1000G_2504_high_coverage/data/ERR3239564/NA18966.final.cram |
| **NA18977** | 1000G_2504_high_coverage/data/ERR3239565/NA18977.final.cram |
| **NA18978** | 1000G_2504_high_coverage/data/ERR3239566/NA18978.final.cram |
| **NA18979** | 1000G_2504_high_coverage/data/ERR3239567/NA18979.final.cram |
| **NA18987** | 1000G_2504_high_coverage/data/ERR3239568/NA18987.final.cram |
| **NA18990** | 1000G_2504_high_coverage/data/ERR3239569/NA18990.final.cram |
| **NA18991** | 1000G_2504_high_coverage/data/ERR3239570/NA18991.final.cram |
| **NA18992** | 1000G_2504_high_coverage/data/ERR3239571/NA18992.final.cram |
| **NA18993** | 1000G_2504_high_coverage/data/ERR3239572/NA18993.final.cram |
| **NA18994** | 1000G_2504_high_coverage/data/ERR3239573/NA18994.final.cram |
| **NA18995** | 1000G_2504_high_coverage/data/ERR3239574/NA18995.final.cram |
| **NA18997** | 1000G_2504_high_coverage/data/ERR3239575/NA18997.final.cram |
| **NA18998** | 1000G_2504_high_coverage/data/ERR3239576/NA18998.final.cram |
| **NA18999** | 1000G_2504_high_coverage/data/ERR3239577/NA18999.final.cram |
| **NA19000** | 1000G_2504_high_coverage/data/ERR3239578/NA19000.final.cram |
| **NA19001** | 1000G_2504_high_coverage/data/ERR3239579/NA19001.final.cram |
| **NA19002** | 1000G_2504_high_coverage/data/ERR3239580/NA19002.final.cram |
| **NA19003** | 1000G_2504_high_coverage/data/ERR3239581/NA19003.final.cram |
| **NA19007** | 1000G_2504_high_coverage/data/ERR3239582/NA19007.final.cram |
| **NA19009** | 1000G_2504_high_coverage/data/ERR3239583/NA19009.final.cram |
| **NA19010** | 1000G_2504_high_coverage/data/ERR3239584/NA19010.final.cram |
| **NA19012** | 1000G_2504_high_coverage/data/ERR3239585/NA19012.final.cram |
| **NA19054** | 1000G_2504_high_coverage/data/ERR3239586/NA19054.final.cram |
| **NA19055** | 1000G_2504_high_coverage/data/ERR3239587/NA19055.final.cram |
| **NA19056** | 1000G_2504_high_coverage/data/ERR3239588/NA19056.final.cram |
| **NA19057** | 1000G_2504_high_coverage/data/ERR3239589/NA19057.final.cram |
| **NA19058** | 1000G_2504_high_coverage/data/ERR3239590/NA19058.final.cram |
| **NA19059** | 1000G_2504_high_coverage/data/ERR3239591/NA19059.final.cram |
| **NA19060** | 1000G_2504_high_coverage/data/ERR3239592/NA19060.final.cram |
| **NA19062** | 1000G_2504_high_coverage/data/ERR3239593/NA19062.final.cram |
| **NA19063** | 1000G_2504_high_coverage/data/ERR3239594/NA19063.final.cram |
| **NA19064** | 1000G_2504_high_coverage/data/ERR3239595/NA19064.final.cram |
| **NA19065** | 1000G_2504_high_coverage/data/ERR3239596/NA19065.final.cram |
| **NA19066** | 1000G_2504_high_coverage/data/ERR3239597/NA19066.final.cram |

| | |
|---|---|
| NA19067 | 1000G_2504_high_coverage/data/ERR3239598/NA19067.final.cram |
| NA19068 | 1000G_2504_high_coverage/data/ERR3239599/NA19068.final.cram |
| NA19070 | 1000G_2504_high_coverage/data/ERR3239600/NA19070.final.cram |
| NA19072 | 1000G_2504_high_coverage/data/ERR3239601/NA19072.final.cram |
| NA19074 | 1000G_2504_high_coverage/data/ERR3239602/NA19074.final.cram |
| NA19075 | 1000G_2504_high_coverage/data/ERR3239603/NA19075.final.cram |
| NA19076 | 1000G_2504_high_coverage/data/ERR3239604/NA19076.final.cram |
| NA19077 | 1000G_2504_high_coverage/data/ERR3239605/NA19077.final.cram |
| NA19078 | 1000G_2504_high_coverage/data/ERR3239606/NA19078.final.cram |
| NA19079 | 1000G_2504_high_coverage/data/ERR3239607/NA19079.final.cram |
| NA19080 | 1000G_2504_high_coverage/data/ERR3239608/NA19080.final.cram |
| NA19081 | 1000G_2504_high_coverage/data/ERR3239609/NA19081.final.cram |
| NA19082 | 1000G_2504_high_coverage/data/ERR3239610/NA19082.final.cram |
| NA19083 | 1000G_2504_high_coverage/data/ERR3239611/NA19083.final.cram |
| NA19084 | 1000G_2504_high_coverage/data/ERR3239612/NA19084.final.cram |
| NA19085 | 1000G_2504_high_coverage/data/ERR3239613/NA19085.final.cram |
| NA19086 | 1000G_2504_high_coverage/data/ERR3239614/NA19086.final.cram |
| NA19087 | 1000G_2504_high_coverage/data/ERR3239615/NA19087.final.cram |
| NA19088 | 1000G_2504_high_coverage/data/ERR3239616/NA19088.final.cram |
| NA19092 | 1000G_2504_high_coverage/data/ERR3239617/NA19092.final.cram |
| NA19095 | 1000G_2504_high_coverage/data/ERR3239618/NA19095.final.cram |
| NA19096 | 1000G_2504_high_coverage/data/ERR3239619/NA19096.final.cram |
| NA19107 | 1000G_2504_high_coverage/data/ERR3239620/NA19107.final.cram |
| NA19113 | 1000G_2504_high_coverage/data/ERR3239621/NA19113.final.cram |
| NA19117 | 1000G_2504_high_coverage/data/ERR3239622/NA19117.final.cram |
| NA19118 | 1000G_2504_high_coverage/data/ERR3239623/NA19118.final.cram |
| NA19121 | 1000G_2504_high_coverage/data/ERR3239624/NA19121.final.cram |
| NA19130 | 1000G_2504_high_coverage/data/ERR3239625/NA19130.final.cram |
| NA19146 | 1000G_2504_high_coverage/data/ERR3239626/NA19146.final.cram |
| NA19149 | 1000G_2504_high_coverage/data/ERR3239627/NA19149.final.cram |
| NA19175 | 1000G_2504_high_coverage/data/ERR3239628/NA19175.final.cram |
| NA19184 | 1000G_2504_high_coverage/data/ERR3239629/NA19184.final.cram |
| NA19185 | 1000G_2504_high_coverage/data/ERR3239630/NA19185.final.cram |
| NA19189 | 1000G_2504_high_coverage/data/ERR3239631/NA19189.final.cram |
| NA19197 | 1000G_2504_high_coverage/data/ERR3239632/NA19197.final.cram |
| NA19198 | 1000G_2504_high_coverage/data/ERR3239633/NA19198.final.cram |
| NA19213 | 1000G_2504_high_coverage/data/ERR3239634/NA19213.final.cram |
| NA19214 | 1000G_2504_high_coverage/data/ERR3239635/NA19214.final.cram |
| NA19223 | 1000G_2504_high_coverage/data/ERR3239636/NA19223.final.cram |
| NA19235 | 1000G_2504_high_coverage/data/ERR3239637/NA19235.final.cram |
| NA19236 | 1000G_2504_high_coverage/data/ERR3239638/NA19236.final.cram |
| NA19247 | 1000G_2504_high_coverage/data/ERR3239639/NA19247.final.cram |
| NA19248 | 1000G_2504_high_coverage/data/ERR3239640/NA19248.final.cram |
| NA19256 | 1000G_2504_high_coverage/data/ERR3239641/NA19256.final.cram |
| NA07048 | 1000G_2504_high_coverage/data/ERR3239642/NA07048.final.cram |
| NA11932 | 1000G_2504_high_coverage/data/ERR3239643/NA11932.final.cram |
| NA11933 | 1000G_2504_high_coverage/data/ERR3239644/NA11933.final.cram |
| NA12046 | 1000G_2504_high_coverage/data/ERR3239645/NA12046.final.cram |
| NA18525 | 1000G_2504_high_coverage/data/ERR3239646/NA18525.final.cram |
| NA18528 | 1000G_2504_high_coverage/data/ERR3239647/NA18528.final.cram |
| NA18531 | 1000G_2504_high_coverage/data/ERR3239648/NA18531.final.cram |
| NA18533 | 1000G_2504_high_coverage/data/ERR3239649/NA18533.final.cram |
| NA18535 | 1000G_2504_high_coverage/data/ERR3239650/NA18535.final.cram |
| NA18538 | 1000G_2504_high_coverage/data/ERR3239651/NA18538.final.cram |
| NA18539 | 1000G_2504_high_coverage/data/ERR3239652/NA18539.final.cram |
| NA18541 | 1000G_2504_high_coverage/data/ERR3239653/NA18541.final.cram |
| NA18553 | 1000G_2504_high_coverage/data/ERR3239654/NA18553.final.cram |
| NA18560 | 1000G_2504_high_coverage/data/ERR3239655/NA18560.final.cram |
| NA18565 | 1000G_2504_high_coverage/data/ERR3239656/NA18565.final.cram |
| NA18567 | 1000G_2504_high_coverage/data/ERR3239657/NA18567.final.cram |
| NA18574 | 1000G_2504_high_coverage/data/ERR3239658/NA18574.final.cram |
| NA18591 | 1000G_2504_high_coverage/data/ERR3239659/NA18591.final.cram |
| NA18629 | 1000G_2504_high_coverage/data/ERR3239660/NA18629.final.cram |
| NA18644 | 1000G_2504_high_coverage/data/ERR3239661/NA18644.final.cram |
| NA18646 | 1000G_2504_high_coverage/data/ERR3239662/NA18646.final.cram |

| | |
|---|---|
| **NA18648** | 1000G_2504_high_coverage/data/ERR3239663/NA18648.final.cram |
| **NA18864** | 1000G_2504_high_coverage/data/ERR3239664/NA18864.final.cram |
| **NA18865** | 1000G_2504_high_coverage/data/ERR3239665/NA18865.final.cram |
| **NA18876** | 1000G_2504_high_coverage/data/ERR3239666/NA18876.final.cram |
| **NA18877** | 1000G_2504_high_coverage/data/ERR3239667/NA18877.final.cram |
| **NA18878** | 1000G_2504_high_coverage/data/ERR3239668/NA18878.final.cram |
| **NA18879** | 1000G_2504_high_coverage/data/ERR3239669/NA18879.final.cram |
| **NA18881** | 1000G_2504_high_coverage/data/ERR3239670/NA18881.final.cram |
| **NA18915** | 1000G_2504_high_coverage/data/ERR3239671/NA18915.final.cram |
| **NA18950** | 1000G_2504_high_coverage/data/ERR3239672/NA18950.final.cram |
| **NA18982** | 1000G_2504_high_coverage/data/ERR3239673/NA18982.final.cram |
| **NA18983** | 1000G_2504_high_coverage/data/ERR3239674/NA18983.final.cram |
| **NA18984** | 1000G_2504_high_coverage/data/ERR3239675/NA18984.final.cram |
| **NA18985** | 1000G_2504_high_coverage/data/ERR3239676/NA18985.final.cram |
| **NA18986** | 1000G_2504_high_coverage/data/ERR3239677/NA18986.final.cram |
| **NA18988** | 1000G_2504_high_coverage/data/ERR3239678/NA18988.final.cram |
| **NA18989** | 1000G_2504_high_coverage/data/ERR3239679/NA18989.final.cram |
| **NA19004** | 1000G_2504_high_coverage/data/ERR3239680/NA19004.final.cram |
| **NA19006** | 1000G_2504_high_coverage/data/ERR3239681/NA19006.final.cram |
| **NA19011** | 1000G_2504_high_coverage/data/ERR3239682/NA19011.final.cram |
| **NA19017** | 1000G_2504_high_coverage/data/ERR3239683/NA19017.final.cram |
| **NA19019** | 1000G_2504_high_coverage/data/ERR3239684/NA19019.final.cram |
| **NA19020** | 1000G_2504_high_coverage/data/ERR3239685/NA19020.final.cram |
| **NA19023** | 1000G_2504_high_coverage/data/ERR3239686/NA19023.final.cram |
| **NA19024** | 1000G_2504_high_coverage/data/ERR3239687/NA19024.final.cram |
| **NA19025** | 1000G_2504_high_coverage/data/ERR3239688/NA19025.final.cram |
| **NA19026** | 1000G_2504_high_coverage/data/ERR3239689/NA19026.final.cram |
| **NA19027** | 1000G_2504_high_coverage/data/ERR3239690/NA19027.final.cram |
| **NA19028** | 1000G_2504_high_coverage/data/ERR3239691/NA19028.final.cram |
| **NA19030** | 1000G_2504_high_coverage/data/ERR3239692/NA19030.final.cram |
| **NA19031** | 1000G_2504_high_coverage/data/ERR3239693/NA19031.final.cram |
| **NA19035** | 1000G_2504_high_coverage/data/ERR3239694/NA19035.final.cram |
| **NA19036** | 1000G_2504_high_coverage/data/ERR3239695/NA19036.final.cram |
| **NA19037** | 1000G_2504_high_coverage/data/ERR3239696/NA19037.final.cram |
| **NA19038** | 1000G_2504_high_coverage/data/ERR3239697/NA19038.final.cram |
| **NA19041** | 1000G_2504_high_coverage/data/ERR3239698/NA19041.final.cram |
| **NA19042** | 1000G_2504_high_coverage/data/ERR3239699/NA19042.final.cram |
| **NA19043** | 1000G_2504_high_coverage/data/ERR3239700/NA19043.final.cram |
| **NA19089** | 1000G_2504_high_coverage/data/ERR3239701/NA19089.final.cram |
| **NA19090** | 1000G_2504_high_coverage/data/ERR3239702/NA19090.final.cram |
| **NA19091** | 1000G_2504_high_coverage/data/ERR3239703/NA19091.final.cram |
| **NA19307** | 1000G_2504_high_coverage/data/ERR3239704/NA19307.final.cram |
| **NA19308** | 1000G_2504_high_coverage/data/ERR3239705/NA19308.final.cram |
| **NA19309** | 1000G_2504_high_coverage/data/ERR3239706/NA19309.final.cram |
| **NA19310** | 1000G_2504_high_coverage/data/ERR3239707/NA19310.final.cram |
| **NA19312** | 1000G_2504_high_coverage/data/ERR3239708/NA19312.final.cram |
| **NA19314** | 1000G_2504_high_coverage/data/ERR3239709/NA19314.final.cram |
| **NA19315** | 1000G_2504_high_coverage/data/ERR3239710/NA19315.final.cram |
| **NA19316** | 1000G_2504_high_coverage/data/ERR3239711/NA19316.final.cram |
| **NA19317** | 1000G_2504_high_coverage/data/ERR3239712/NA19317.final.cram |
| **NA19318** | 1000G_2504_high_coverage/data/ERR3239713/NA19318.final.cram |
| **NA19319** | 1000G_2504_high_coverage/data/ERR3239714/NA19319.final.cram |
| **NA19320** | 1000G_2504_high_coverage/data/ERR3239715/NA19320.final.cram |
| **NA19321** | 1000G_2504_high_coverage/data/ERR3239716/NA19321.final.cram |
| **NA19323** | 1000G_2504_high_coverage/data/ERR3239717/NA19323.final.cram |
| **NA19324** | 1000G_2504_high_coverage/data/ERR3239718/NA19324.final.cram |
| **NA19327** | 1000G_2504_high_coverage/data/ERR3239719/NA19327.final.cram |
| **NA19328** | 1000G_2504_high_coverage/data/ERR3239720/NA19328.final.cram |
| **NA19331** | 1000G_2504_high_coverage/data/ERR3239721/NA19331.final.cram |
| **NA19332** | 1000G_2504_high_coverage/data/ERR3239722/NA19332.final.cram |
| **NA19334** | 1000G_2504_high_coverage/data/ERR3239723/NA19334.final.cram |
| **NA19338** | 1000G_2504_high_coverage/data/ERR3239724/NA19338.final.cram |
| **NA19346** | 1000G_2504_high_coverage/data/ERR3239725/NA19346.final.cram |
| **NA19347** | 1000G_2504_high_coverage/data/ERR3239726/NA19347.final.cram |
| **NA19350** | 1000G_2504_high_coverage/data/ERR3239727/NA19350.final.cram |

| | |
|---|---|
| **NA19351** | 1000G_2504_high_coverage/data/ERR3239728/NA19351.final.cram |
| **NA19355** | 1000G_2504_high_coverage/data/ERR3239729/NA19355.final.cram |
| **NA19360** | 1000G_2504_high_coverage/data/ERR3239730/NA19360.final.cram |
| **NA19372** | 1000G_2504_high_coverage/data/ERR3239731/NA19372.final.cram |
| **NA19374** | 1000G_2504_high_coverage/data/ERR3239732/NA19374.final.cram |
| **NA19375** | 1000G_2504_high_coverage/data/ERR3239733/NA19375.final.cram |
| **NA19376** | 1000G_2504_high_coverage/data/ERR3239734/NA19376.final.cram |
| **NA19377** | 1000G_2504_high_coverage/data/ERR3239735/NA19377.final.cram |
| **NA19378** | 1000G_2504_high_coverage/data/ERR3239736/NA19378.final.cram |
| **NA19379** | 1000G_2504_high_coverage/data/ERR3239737/NA19379.final.cram |
| **NA19380** | 1000G_2504_high_coverage/data/ERR3239738/NA19380.final.cram |
| **NA19383** | 1000G_2504_high_coverage/data/ERR3239739/NA19383.final.cram |
| **NA19384** | 1000G_2504_high_coverage/data/ERR3239740/NA19384.final.cram |
| **NA19385** | 1000G_2504_high_coverage/data/ERR3239741/NA19385.final.cram |
| **NA19390** | 1000G_2504_high_coverage/data/ERR3239742/NA19390.final.cram |
| **NA19391** | 1000G_2504_high_coverage/data/ERR3239743/NA19391.final.cram |
| **NA19393** | 1000G_2504_high_coverage/data/ERR3239744/NA19393.final.cram |
| **NA19394** | 1000G_2504_high_coverage/data/ERR3239745/NA19394.final.cram |
| **NA19395** | 1000G_2504_high_coverage/data/ERR3239746/NA19395.final.cram |
| **NA19397** | 1000G_2504_high_coverage/data/ERR3239747/NA19397.final.cram |
| **NA19399** | 1000G_2504_high_coverage/data/ERR3239748/NA19399.final.cram |
| **NA19401** | 1000G_2504_high_coverage/data/ERR3239749/NA19401.final.cram |
| **NA19403** | 1000G_2504_high_coverage/data/ERR3239750/NA19403.final.cram |
| **NA19404** | 1000G_2504_high_coverage/data/ERR3239751/NA19404.final.cram |
| **NA19428** | 1000G_2504_high_coverage/data/ERR3239752/NA19428.final.cram |
| **NA19429** | 1000G_2504_high_coverage/data/ERR3239753/NA19429.final.cram |
| **NA19430** | 1000G_2504_high_coverage/data/ERR3239754/NA19430.final.cram |
| **NA19431** | 1000G_2504_high_coverage/data/ERR3239755/NA19431.final.cram |
| **NA19434** | 1000G_2504_high_coverage/data/ERR3239756/NA19434.final.cram |
| **NA19435** | 1000G_2504_high_coverage/data/ERR3239757/NA19435.final.cram |
| **NA19436** | 1000G_2504_high_coverage/data/ERR3239758/NA19436.final.cram |
| **NA19437** | 1000G_2504_high_coverage/data/ERR3239759/NA19437.final.cram |
| **NA19438** | 1000G_2504_high_coverage/data/ERR3239760/NA19438.final.cram |
| **NA19439** | 1000G_2504_high_coverage/data/ERR3239761/NA19439.final.cram |
| **NA19440** | 1000G_2504_high_coverage/data/ERR3239762/NA19440.final.cram |
| **NA19443** | 1000G_2504_high_coverage/data/ERR3239763/NA19443.final.cram |
| **NA19445** | 1000G_2504_high_coverage/data/ERR3239764/NA19445.final.cram |
| **NA19446** | 1000G_2504_high_coverage/data/ERR3239765/NA19446.final.cram |
| **NA19448** | 1000G_2504_high_coverage/data/ERR3239766/NA19448.final.cram |
| **NA19449** | 1000G_2504_high_coverage/data/ERR3239767/NA19449.final.cram |
| **NA19451** | 1000G_2504_high_coverage/data/ERR3239768/NA19451.final.cram |
| **NA19452** | 1000G_2504_high_coverage/data/ERR3239769/NA19452.final.cram |
| **NA19454** | 1000G_2504_high_coverage/data/ERR3239770/NA19454.final.cram |
| **NA19455** | 1000G_2504_high_coverage/data/ERR3239771/NA19455.final.cram |
| **NA19456** | 1000G_2504_high_coverage/data/ERR3239772/NA19456.final.cram |
| **NA19457** | 1000G_2504_high_coverage/data/ERR3239773/NA19457.final.cram |
| **NA19461** | 1000G_2504_high_coverage/data/ERR3239774/NA19461.final.cram |
| **NA19462** | 1000G_2504_high_coverage/data/ERR3239775/NA19462.final.cram |
| **NA19463** | 1000G_2504_high_coverage/data/ERR3239776/NA19463.final.cram |
| **NA19466** | 1000G_2504_high_coverage/data/ERR3239777/NA19466.final.cram |
| **NA19467** | 1000G_2504_high_coverage/data/ERR3239778/NA19467.final.cram |
| **NA19468** | 1000G_2504_high_coverage/data/ERR3239779/NA19468.final.cram |
| **NA19471** | 1000G_2504_high_coverage/data/ERR3239780/NA19471.final.cram |
| **NA19472** | 1000G_2504_high_coverage/data/ERR3239781/NA19472.final.cram |
| **NA19473** | 1000G_2504_high_coverage/data/ERR3239782/NA19473.final.cram |
| **NA19474** | 1000G_2504_high_coverage/data/ERR3239783/NA19474.final.cram |
| **NA19475** | 1000G_2504_high_coverage/data/ERR3239784/NA19475.final.cram |
| **NA20502** | 1000G_2504_high_coverage/data/ERR3239785/NA20502.final.cram |
| **NA20503** | 1000G_2504_high_coverage/data/ERR3239786/NA20503.final.cram |
| **NA20504** | 1000G_2504_high_coverage/data/ERR3239787/NA20504.final.cram |
| **NA20505** | 1000G_2504_high_coverage/data/ERR3239788/NA20505.final.cram |
| **NA20506** | 1000G_2504_high_coverage/data/ERR3239789/NA20506.final.cram |
| **NA20507** | 1000G_2504_high_coverage/data/ERR3239790/NA20507.final.cram |
| **NA20508** | 1000G_2504_high_coverage/data/ERR3239791/NA20508.final.cram |
| **NA20509** | 1000G_2504_high_coverage/data/ERR3239792/NA20509.final.cram |

| | |
|---|---|
| **NA20510** | 1000G_2504_high_coverage/data/ERR3239793/NA20510.final.cram |
| **NA20511** | 1000G_2504_high_coverage/data/ERR3239794/NA20511.final.cram |
| **NA20512** | 1000G_2504_high_coverage/data/ERR3239795/NA20512.final.cram |
| **NA20513** | 1000G_2504_high_coverage/data/ERR3239796/NA20513.final.cram |
| **NA20514** | 1000G_2504_high_coverage/data/ERR3239797/NA20514.final.cram |
| **NA20515** | 1000G_2504_high_coverage/data/ERR3239798/NA20515.final.cram |
| **NA20516** | 1000G_2504_high_coverage/data/ERR3239799/NA20516.final.cram |
| **NA20517** | 1000G_2504_high_coverage/data/ERR3239800/NA20517.final.cram |
| **NA20518** | 1000G_2504_high_coverage/data/ERR3239801/NA20518.final.cram |
| **NA20519** | 1000G_2504_high_coverage/data/ERR3239802/NA20519.final.cram |
| **NA20520** | 1000G_2504_high_coverage/data/ERR3239803/NA20520.final.cram |
| **NA20521** | 1000G_2504_high_coverage/data/ERR3239804/NA20521.final.cram |
| **NA20522** | 1000G_2504_high_coverage/data/ERR3239805/NA20522.final.cram |
| **NA20524** | 1000G_2504_high_coverage/data/ERR3239806/NA20524.final.cram |
| **NA20525** | 1000G_2504_high_coverage/data/ERR3239807/NA20525.final.cram |
| **NA20527** | 1000G_2504_high_coverage/data/ERR3239808/NA20527.final.cram |
| **NA20528** | 1000G_2504_high_coverage/data/ERR3239809/NA20528.final.cram |
| **NA20529** | 1000G_2504_high_coverage/data/ERR3239810/NA20529.final.cram |
| **NA20530** | 1000G_2504_high_coverage/data/ERR3239811/NA20530.final.cram |
| **NA20531** | 1000G_2504_high_coverage/data/ERR3239812/NA20531.final.cram |
| **NA20532** | 1000G_2504_high_coverage/data/ERR3239813/NA20532.final.cram |
| **NA20533** | 1000G_2504_high_coverage/data/ERR3239814/NA20533.final.cram |
| **NA20534** | 1000G_2504_high_coverage/data/ERR3239815/NA20534.final.cram |
| **NA20535** | 1000G_2504_high_coverage/data/ERR3239816/NA20535.final.cram |
| **NA20536** | 1000G_2504_high_coverage/data/ERR3239817/NA20536.final.cram |
| **NA20538** | 1000G_2504_high_coverage/data/ERR3239818/NA20538.final.cram |
| **NA20539** | 1000G_2504_high_coverage/data/ERR3239819/NA20539.final.cram |
| **NA20540** | 1000G_2504_high_coverage/data/ERR3239820/NA20540.final.cram |
| **NA20541** | 1000G_2504_high_coverage/data/ERR3239821/NA20541.final.cram |
| **NA20542** | 1000G_2504_high_coverage/data/ERR3239822/NA20542.final.cram |
| **NA20543** | 1000G_2504_high_coverage/data/ERR3239823/NA20543.final.cram |
| **NA20544** | 1000G_2504_high_coverage/data/ERR3239824/NA20544.final.cram |
| **NA20581** | 1000G_2504_high_coverage/data/ERR3239825/NA20581.final.cram |
| **NA20582** | 1000G_2504_high_coverage/data/ERR3239826/NA20582.final.cram |
| **NA20585** | 1000G_2504_high_coverage/data/ERR3239827/NA20585.final.cram |
| **NA20586** | 1000G_2504_high_coverage/data/ERR3239828/NA20586.final.cram |
| **NA20587** | 1000G_2504_high_coverage/data/ERR3239829/NA20587.final.cram |
| **NA20588** | 1000G_2504_high_coverage/data/ERR3239830/NA20588.final.cram |
| **NA20589** | 1000G_2504_high_coverage/data/ERR3239831/NA20589.final.cram |
| **NA20752** | 1000G_2504_high_coverage/data/ERR3239832/NA20752.final.cram |
| **NA20753** | 1000G_2504_high_coverage/data/ERR3239833/NA20753.final.cram |
| **NA20754** | 1000G_2504_high_coverage/data/ERR3239834/NA20754.final.cram |
| **NA20755** | 1000G_2504_high_coverage/data/ERR3239835/NA20755.final.cram |
| **NA20756** | 1000G_2504_high_coverage/data/ERR3239836/NA20756.final.cram |
| **NA20757** | 1000G_2504_high_coverage/data/ERR3239837/NA20757.final.cram |
| **NA20758** | 1000G_2504_high_coverage/data/ERR3239838/NA20758.final.cram |
| **NA20759** | 1000G_2504_high_coverage/data/ERR3239839/NA20759.final.cram |
| **NA20760** | 1000G_2504_high_coverage/data/ERR3239840/NA20760.final.cram |
| **NA20761** | 1000G_2504_high_coverage/data/ERR3239841/NA20761.final.cram |
| **NA20762** | 1000G_2504_high_coverage/data/ERR3239842/NA20762.final.cram |
| **NA20763** | 1000G_2504_high_coverage/data/ERR3239843/NA20763.final.cram |
| **NA20764** | 1000G_2504_high_coverage/data/ERR3239844/NA20764.final.cram |
| **NA20765** | 1000G_2504_high_coverage/data/ERR3239845/NA20765.final.cram |
| **NA20766** | 1000G_2504_high_coverage/data/ERR3239846/NA20766.final.cram |
| **NA20767** | 1000G_2504_high_coverage/data/ERR3239847/NA20767.final.cram |
| **NA20768** | 1000G_2504_high_coverage/data/ERR3239848/NA20768.final.cram |
| **NA20769** | 1000G_2504_high_coverage/data/ERR3239849/NA20769.final.cram |
| **NA20770** | 1000G_2504_high_coverage/data/ERR3239850/NA20770.final.cram |
| **NA20771** | 1000G_2504_high_coverage/data/ERR3239851/NA20771.final.cram |
| **NA20772** | 1000G_2504_high_coverage/data/ERR3239852/NA20772.final.cram |
| **NA20773** | 1000G_2504_high_coverage/data/ERR3239853/NA20773.final.cram |
| **NA20774** | 1000G_2504_high_coverage/data/ERR3239854/NA20774.final.cram |
| **NA20775** | 1000G_2504_high_coverage/data/ERR3239855/NA20775.final.cram |
| **NA20778** | 1000G_2504_high_coverage/data/ERR3239856/NA20778.final.cram |
| **NA20783** | 1000G_2504_high_coverage/data/ERR3239857/NA20783.final.cram |

| | |
|---|---|
| **NA20785** | 1000G_2504_high_coverage/data/ERR3239858/NA20785.final.cram |
| **NA20786** | 1000G_2504_high_coverage/data/ERR3239859/NA20786.final.cram |
| **NA20787** | 1000G_2504_high_coverage/data/ERR3239860/NA20787.final.cram |
| **NA20790** | 1000G_2504_high_coverage/data/ERR3239861/NA20790.final.cram |
| **NA20792** | 1000G_2504_high_coverage/data/ERR3239862/NA20792.final.cram |
| **NA20795** | 1000G_2504_high_coverage/data/ERR3239863/NA20795.final.cram |
| **NA20796** | 1000G_2504_high_coverage/data/ERR3239864/NA20796.final.cram |
| **NA20797** | 1000G_2504_high_coverage/data/ERR3239865/NA20797.final.cram |
| **NA20798** | 1000G_2504_high_coverage/data/ERR3239866/NA20798.final.cram |
| **NA20799** | 1000G_2504_high_coverage/data/ERR3239867/NA20799.final.cram |
| **NA20800** | 1000G_2504_high_coverage/data/ERR3239868/NA20800.final.cram |
| **NA20801** | 1000G_2504_high_coverage/data/ERR3239869/NA20801.final.cram |
| **NA20802** | 1000G_2504_high_coverage/data/ERR3239870/NA20802.final.cram |
| **NA20803** | 1000G_2504_high_coverage/data/ERR3239871/NA20803.final.cram |
| **NA20804** | 1000G_2504_high_coverage/data/ERR3239872/NA20804.final.cram |
| **NA20805** | 1000G_2504_high_coverage/data/ERR3239873/NA20805.final.cram |
| **NA20806** | 1000G_2504_high_coverage/data/ERR3239874/NA20806.final.cram |
| **NA20807** | 1000G_2504_high_coverage/data/ERR3239875/NA20807.final.cram |
| **NA20808** | 1000G_2504_high_coverage/data/ERR3239876/NA20808.final.cram |
| **NA20809** | 1000G_2504_high_coverage/data/ERR3239877/NA20809.final.cram |
| **NA20810** | 1000G_2504_high_coverage/data/ERR3239878/NA20810.final.cram |
| **NA20811** | 1000G_2504_high_coverage/data/ERR3239879/NA20811.final.cram |
| **NA20812** | 1000G_2504_high_coverage/data/ERR3239880/NA20812.final.cram |
| **NA20813** | 1000G_2504_high_coverage/data/ERR3239881/NA20813.final.cram |
| **NA20814** | 1000G_2504_high_coverage/data/ERR3239882/NA20814.final.cram |
| **NA20815** | 1000G_2504_high_coverage/data/ERR3239883/NA20815.final.cram |
| **NA20818** | 1000G_2504_high_coverage/data/ERR3239884/NA20818.final.cram |
| **NA20819** | 1000G_2504_high_coverage/data/ERR3239885/NA20819.final.cram |
| **NA20821** | 1000G_2504_high_coverage/data/ERR3239886/NA20821.final.cram |
| **NA20822** | 1000G_2504_high_coverage/data/ERR3239887/NA20822.final.cram |
| **NA20826** | 1000G_2504_high_coverage/data/ERR3239888/NA20826.final.cram |
| **NA20827** | 1000G_2504_high_coverage/data/ERR3239889/NA20827.final.cram |
| **NA20828** | 1000G_2504_high_coverage/data/ERR3239890/NA20828.final.cram |
| **NA20832** | 1000G_2504_high_coverage/data/ERR3239891/NA20832.final.cram |
| **NA19222** | 1000G_2504_high_coverage/data/ERR3239892/NA19222.final.cram |
| **NA19625** | 1000G_2504_high_coverage/data/ERR3239893/NA19625.final.cram |
| **NA19648** | 1000G_2504_high_coverage/data/ERR3239894/NA19648.final.cram |
| **NA19649** | 1000G_2504_high_coverage/data/ERR3239895/NA19649.final.cram |
| **NA19651** | 1000G_2504_high_coverage/data/ERR3239896/NA19651.final.cram |
| **NA19652** | 1000G_2504_high_coverage/data/ERR3239897/NA19652.final.cram |
| **NA19654** | 1000G_2504_high_coverage/data/ERR3239898/NA19654.final.cram |
| **NA19655** | 1000G_2504_high_coverage/data/ERR3239899/NA19655.final.cram |
| **NA19657** | 1000G_2504_high_coverage/data/ERR3239900/NA19657.final.cram |
| **NA19658** | 1000G_2504_high_coverage/data/ERR3239901/NA19658.final.cram |
| **NA19661** | 1000G_2504_high_coverage/data/ERR3239902/NA19661.final.cram |
| **NA19663** | 1000G_2504_high_coverage/data/ERR3239903/NA19663.final.cram |
| **NA19664** | 1000G_2504_high_coverage/data/ERR3239904/NA19664.final.cram |
| **NA19669** | 1000G_2504_high_coverage/data/ERR3239905/NA19669.final.cram |
| **NA19670** | 1000G_2504_high_coverage/data/ERR3239906/NA19670.final.cram |
| **NA19676** | 1000G_2504_high_coverage/data/ERR3239907/NA19676.final.cram |
| **NA19678** | 1000G_2504_high_coverage/data/ERR3239908/NA19678.final.cram |
| **NA19679** | 1000G_2504_high_coverage/data/ERR3239909/NA19679.final.cram |
| **NA19681** | 1000G_2504_high_coverage/data/ERR3239910/NA19681.final.cram |
| **NA19682** | 1000G_2504_high_coverage/data/ERR3239911/NA19682.final.cram |
| **NA19684** | 1000G_2504_high_coverage/data/ERR3239912/NA19684.final.cram |
| **NA19700** | 1000G_2504_high_coverage/data/ERR3239913/NA19700.final.cram |
| **NA19701** | 1000G_2504_high_coverage/data/ERR3239914/NA19701.final.cram |
| **NA19703** | 1000G_2504_high_coverage/data/ERR3239915/NA19703.final.cram |
| **NA19704** | 1000G_2504_high_coverage/data/ERR3239916/NA19704.final.cram |
| **NA19707** | 1000G_2504_high_coverage/data/ERR3239917/NA19707.final.cram |
| **NA19711** | 1000G_2504_high_coverage/data/ERR3239918/NA19711.final.cram |
| **NA19712** | 1000G_2504_high_coverage/data/ERR3239919/NA19712.final.cram |
| **NA19713** | 1000G_2504_high_coverage/data/ERR3239920/NA19713.final.cram |
| **NA19716** | 1000G_2504_high_coverage/data/ERR3239921/NA19716.final.cram |
| **NA19717** | 1000G_2504_high_coverage/data/ERR3239922/NA19717.final.cram |

| | |
|---|---|
| **NA19719** | 1000G_2504_high_coverage/data/ERR3239923/NA19719.final.cram |
| **NA19720** | 1000G_2504_high_coverage/data/ERR3239924/NA19720.final.cram |
| **NA19722** | 1000G_2504_high_coverage/data/ERR3239925/NA19722.final.cram |
| **NA19723** | 1000G_2504_high_coverage/data/ERR3239926/NA19723.final.cram |
| **NA19725** | 1000G_2504_high_coverage/data/ERR3239927/NA19725.final.cram |
| **NA19726** | 1000G_2504_high_coverage/data/ERR3239928/NA19726.final.cram |
| **NA19728** | 1000G_2504_high_coverage/data/ERR3239929/NA19728.final.cram |
| **NA19729** | 1000G_2504_high_coverage/data/ERR3239930/NA19729.final.cram |
| **NA19731** | 1000G_2504_high_coverage/data/ERR3239931/NA19731.final.cram |
| **NA19732** | 1000G_2504_high_coverage/data/ERR3239932/NA19732.final.cram |
| **NA19746** | 1000G_2504_high_coverage/data/ERR3239933/NA19746.final.cram |
| **NA19747** | 1000G_2504_high_coverage/data/ERR3239934/NA19747.final.cram |
| **NA19749** | 1000G_2504_high_coverage/data/ERR3239935/NA19749.final.cram |
| **NA19750** | 1000G_2504_high_coverage/data/ERR3239936/NA19750.final.cram |
| **NA19755** | 1000G_2504_high_coverage/data/ERR3239937/NA19755.final.cram |
| **NA19756** | 1000G_2504_high_coverage/data/ERR3239938/NA19756.final.cram |
| **NA19758** | 1000G_2504_high_coverage/data/ERR3239939/NA19758.final.cram |
| **NA19759** | 1000G_2504_high_coverage/data/ERR3239940/NA19759.final.cram |
| **NA19761** | 1000G_2504_high_coverage/data/ERR3239941/NA19761.final.cram |
| **NA19762** | 1000G_2504_high_coverage/data/ERR3239942/NA19762.final.cram |
| **NA19770** | 1000G_2504_high_coverage/data/ERR3239943/NA19770.final.cram |
| **NA19771** | 1000G_2504_high_coverage/data/ERR3239944/NA19771.final.cram |
| **NA19773** | 1000G_2504_high_coverage/data/ERR3239945/NA19773.final.cram |
| **NA19774** | 1000G_2504_high_coverage/data/ERR3239946/NA19774.final.cram |
| **NA19776** | 1000G_2504_high_coverage/data/ERR3239947/NA19776.final.cram |
| **NA19777** | 1000G_2504_high_coverage/data/ERR3239948/NA19777.final.cram |
| **NA19779** | 1000G_2504_high_coverage/data/ERR3239949/NA19779.final.cram |
| **NA19780** | 1000G_2504_high_coverage/data/ERR3239950/NA19780.final.cram |
| **NA19782** | 1000G_2504_high_coverage/data/ERR3239951/NA19782.final.cram |
| **NA19783** | 1000G_2504_high_coverage/data/ERR3239952/NA19783.final.cram |
| **NA19785** | 1000G_2504_high_coverage/data/ERR3239953/NA19785.final.cram |
| **NA19786** | 1000G_2504_high_coverage/data/ERR3239954/NA19786.final.cram |
| **NA19788** | 1000G_2504_high_coverage/data/ERR3239955/NA19788.final.cram |
| **NA19789** | 1000G_2504_high_coverage/data/ERR3239956/NA19789.final.cram |
| **NA19794** | 1000G_2504_high_coverage/data/ERR3239957/NA19794.final.cram |
| **NA19795** | 1000G_2504_high_coverage/data/ERR3239958/NA19795.final.cram |
| **NA19818** | 1000G_2504_high_coverage/data/ERR3239959/NA19818.final.cram |
| **NA19819** | 1000G_2504_high_coverage/data/ERR3239960/NA19819.final.cram |
| **NA19834** | 1000G_2504_high_coverage/data/ERR3239961/NA19834.final.cram |
| **NA19835** | 1000G_2504_high_coverage/data/ERR3239962/NA19835.final.cram |
| **NA19900** | 1000G_2504_high_coverage/data/ERR3239963/NA19900.final.cram |
| **NA19901** | 1000G_2504_high_coverage/data/ERR3239964/NA19901.final.cram |
| **NA19904** | 1000G_2504_high_coverage/data/ERR3239965/NA19904.final.cram |
| **NA19908** | 1000G_2504_high_coverage/data/ERR3239966/NA19908.final.cram |
| **NA19909** | 1000G_2504_high_coverage/data/ERR3239967/NA19909.final.cram |
| **NA19914** | 1000G_2504_high_coverage/data/ERR3239968/NA19914.final.cram |
| **NA19916** | 1000G_2504_high_coverage/data/ERR3239969/NA19916.final.cram |
| **NA19917** | 1000G_2504_high_coverage/data/ERR3239970/NA19917.final.cram |
| **NA19920** | 1000G_2504_high_coverage/data/ERR3239971/NA19920.final.cram |
| **NA19921** | 1000G_2504_high_coverage/data/ERR3239972/NA19921.final.cram |
| **NA19982** | 1000G_2504_high_coverage/data/ERR3239973/NA19982.final.cram |
| **NA20126** | 1000G_2504_high_coverage/data/ERR3239974/NA20126.final.cram |
| **NA20127** | 1000G_2504_high_coverage/data/ERR3239975/NA20127.final.cram |
| **NA20276** | 1000G_2504_high_coverage/data/ERR3239976/NA20276.final.cram |
| **NA20278** | 1000G_2504_high_coverage/data/ERR3239977/NA20278.final.cram |
| **NA20281** | 1000G_2504_high_coverage/data/ERR3239978/NA20281.final.cram |
| **NA20282** | 1000G_2504_high_coverage/data/ERR3239979/NA20282.final.cram |
| **NA20287** | 1000G_2504_high_coverage/data/ERR3239980/NA20287.final.cram |
| **NA20289** | 1000G_2504_high_coverage/data/ERR3239981/NA20289.final.cram |
| **NA20291** | 1000G_2504_high_coverage/data/ERR3239982/NA20291.final.cram |
| **NA20294** | 1000G_2504_high_coverage/data/ERR3239983/NA20294.final.cram |
| **NA20296** | 1000G_2504_high_coverage/data/ERR3239984/NA20296.final.cram |
| **NA20299** | 1000G_2504_high_coverage/data/ERR3239985/NA20299.final.cram |
| **NA20314** | 1000G_2504_high_coverage/data/ERR3239986/NA20314.final.cram |
| **NA20317** | 1000G_2504_high_coverage/data/ERR3239987/NA20317.final.cram |

| | |
|---|---|
| **NA20332** | 1000G_2504_high_coverage/data/ERR3239988/NA20332.final.cram |
| **NA20334** | 1000G_2504_high_coverage/data/ERR3239989/NA20334.final.cram |
| **NA20340** | 1000G_2504_high_coverage/data/ERR3239990/NA20340.final.cram |
| **NA20342** | 1000G_2504_high_coverage/data/ERR3239991/NA20342.final.cram |
| **NA20346** | 1000G_2504_high_coverage/data/ERR3239992/NA20346.final.cram |
| **NA20348** | 1000G_2504_high_coverage/data/ERR3239993/NA20348.final.cram |
| **NA20356** | 1000G_2504_high_coverage/data/ERR3239994/NA20356.final.cram |
| **NA20357** | 1000G_2504_high_coverage/data/ERR3239995/NA20357.final.cram |
| **NA20359** | 1000G_2504_high_coverage/data/ERR3239996/NA20359.final.cram |
| **NA20845** | 1000G_2504_high_coverage/data/ERR3239997/NA20845.final.cram |
| **NA20846** | 1000G_2504_high_coverage/data/ERR3239998/NA20846.final.cram |
| **NA20847** | 1000G_2504_high_coverage/data/ERR3239999/NA20847.final.cram |
| **NA20849** | 1000G_2504_high_coverage/data/ERR3240000/NA20849.final.cram |
| **NA20850** | 1000G_2504_high_coverage/data/ERR3240001/NA20850.final.cram |
| **NA20851** | 1000G_2504_high_coverage/data/ERR3240002/NA20851.final.cram |
| **NA20852** | 1000G_2504_high_coverage/data/ERR3240003/NA20852.final.cram |
| **NA20853** | 1000G_2504_high_coverage/data/ERR3240004/NA20853.final.cram |
| **NA20854** | 1000G_2504_high_coverage/data/ERR3240005/NA20854.final.cram |
| **NA20856** | 1000G_2504_high_coverage/data/ERR3240006/NA20856.final.cram |
| **NA20858** | 1000G_2504_high_coverage/data/ERR3240007/NA20858.final.cram |
| **NA20859** | 1000G_2504_high_coverage/data/ERR3240008/NA20859.final.cram |
| **NA20861** | 1000G_2504_high_coverage/data/ERR3240009/NA20861.final.cram |
| **NA20862** | 1000G_2504_high_coverage/data/ERR3240010/NA20862.final.cram |
| **NA20866** | 1000G_2504_high_coverage/data/ERR3240011/NA20866.final.cram |
| **NA20869** | 1000G_2504_high_coverage/data/ERR3240012/NA20869.final.cram |
| **NA20870** | 1000G_2504_high_coverage/data/ERR3240013/NA20870.final.cram |
| **NA20872** | 1000G_2504_high_coverage/data/ERR3240014/NA20872.final.cram |
| **NA20874** | 1000G_2504_high_coverage/data/ERR3240015/NA20874.final.cram |
| **NA20875** | 1000G_2504_high_coverage/data/ERR3240016/NA20875.final.cram |
| **NA20876** | 1000G_2504_high_coverage/data/ERR3240017/NA20876.final.cram |
| **NA20877** | 1000G_2504_high_coverage/data/ERR3240018/NA20877.final.cram |
| **NA20878** | 1000G_2504_high_coverage/data/ERR3240019/NA20878.final.cram |
| **NA20881** | 1000G_2504_high_coverage/data/ERR3240020/NA20881.final.cram |
| **NA20882** | 1000G_2504_high_coverage/data/ERR3240021/NA20882.final.cram |
| **NA20884** | 1000G_2504_high_coverage/data/ERR3240022/NA20884.final.cram |
| **NA20885** | 1000G_2504_high_coverage/data/ERR3240023/NA20885.final.cram |
| **NA20886** | 1000G_2504_high_coverage/data/ERR3240024/NA20886.final.cram |
| **NA20887** | 1000G_2504_high_coverage/data/ERR3240025/NA20887.final.cram |
| **NA20888** | 1000G_2504_high_coverage/data/ERR3240026/NA20888.final.cram |
| **NA20889** | 1000G_2504_high_coverage/data/ERR3240027/NA20889.final.cram |
| **NA20890** | 1000G_2504_high_coverage/data/ERR3240028/NA20890.final.cram |
| **NA20891** | 1000G_2504_high_coverage/data/ERR3240029/NA20891.final.cram |
| **NA20892** | 1000G_2504_high_coverage/data/ERR3240030/NA20892.final.cram |
| **NA20894** | 1000G_2504_high_coverage/data/ERR3240031/NA20894.final.cram |
| **NA20895** | 1000G_2504_high_coverage/data/ERR3240032/NA20895.final.cram |
| **NA20896** | 1000G_2504_high_coverage/data/ERR3240033/NA20896.final.cram |
| **NA20897** | 1000G_2504_high_coverage/data/ERR3240034/NA20897.final.cram |
| **NA20899** | 1000G_2504_high_coverage/data/ERR3240035/NA20899.final.cram |
| **NA20900** | 1000G_2504_high_coverage/data/ERR3240036/NA20900.final.cram |
| **NA20901** | 1000G_2504_high_coverage/data/ERR3240037/NA20901.final.cram |
| **NA20902** | 1000G_2504_high_coverage/data/ERR3240038/NA20902.final.cram |
| **NA20903** | 1000G_2504_high_coverage/data/ERR3240039/NA20903.final.cram |
| **NA20904** | 1000G_2504_high_coverage/data/ERR3240040/NA20904.final.cram |
| **NA20906** | 1000G_2504_high_coverage/data/ERR3240041/NA20906.final.cram |
| **NA20908** | 1000G_2504_high_coverage/data/ERR3240042/NA20908.final.cram |
| **NA20910** | 1000G_2504_high_coverage/data/ERR3240043/NA20910.final.cram |
| **NA20911** | 1000G_2504_high_coverage/data/ERR3240044/NA20911.final.cram |
| **NA21086** | 1000G_2504_high_coverage/data/ERR3240045/NA21086.final.cram |
| **NA21088** | 1000G_2504_high_coverage/data/ERR3240046/NA21088.final.cram |
| **NA21089** | 1000G_2504_high_coverage/data/ERR3240047/NA21089.final.cram |
| **NA21090** | 1000G_2504_high_coverage/data/ERR3240048/NA21090.final.cram |
| **NA21091** | 1000G_2504_high_coverage/data/ERR3240049/NA21091.final.cram |
| **NA21092** | 1000G_2504_high_coverage/data/ERR3240050/NA21092.final.cram |
| **NA21094** | 1000G_2504_high_coverage/data/ERR3240051/NA21094.final.cram |
| **NA21097** | 1000G_2504_high_coverage/data/ERR3240052/NA21097.final.cram |

| | |
|---|---|
| **NA21098** | 1000G_2504_high_coverage/data/ERR3240053/NA21098.final.cram |
| **NA21099** | 1000G_2504_high_coverage/data/ERR3240054/NA21099.final.cram |
| **NA21100** | 1000G_2504_high_coverage/data/ERR3240055/NA21100.final.cram |
| **NA21101** | 1000G_2504_high_coverage/data/ERR3240056/NA21101.final.cram |
| **NA21102** | 1000G_2504_high_coverage/data/ERR3240057/NA21102.final.cram |
| **NA21103** | 1000G_2504_high_coverage/data/ERR3240058/NA21103.final.cram |
| **NA21104** | 1000G_2504_high_coverage/data/ERR3240059/NA21104.final.cram |
| **NA21105** | 1000G_2504_high_coverage/data/ERR3240060/NA21105.final.cram |
| **NA21106** | 1000G_2504_high_coverage/data/ERR3240061/NA21106.final.cram |
| **NA21107** | 1000G_2504_high_coverage/data/ERR3240062/NA21107.final.cram |
| **NA21108** | 1000G_2504_high_coverage/data/ERR3240063/NA21108.final.cram |
| **NA21109** | 1000G_2504_high_coverage/data/ERR3240064/NA21109.final.cram |
| **NA21110** | 1000G_2504_high_coverage/data/ERR3240065/NA21110.final.cram |
| **NA21111** | 1000G_2504_high_coverage/data/ERR3240066/NA21111.final.cram |
| **NA21112** | 1000G_2504_high_coverage/data/ERR3240067/NA21112.final.cram |
| **NA21113** | 1000G_2504_high_coverage/data/ERR3240068/NA21113.final.cram |
| **NA21115** | 1000G_2504_high_coverage/data/ERR3240069/NA21115.final.cram |
| **NA21116** | 1000G_2504_high_coverage/data/ERR3240070/NA21116.final.cram |
| **NA21117** | 1000G_2504_high_coverage/data/ERR3240071/NA21117.final.cram |
| **NA21118** | 1000G_2504_high_coverage/data/ERR3240072/NA21118.final.cram |
| **NA21119** | 1000G_2504_high_coverage/data/ERR3240073/NA21119.final.cram |
| **NA21120** | 1000G_2504_high_coverage/data/ERR3240074/NA21120.final.cram |
| **NA21122** | 1000G_2504_high_coverage/data/ERR3240075/NA21122.final.cram |
| **NA21123** | 1000G_2504_high_coverage/data/ERR3240076/NA21123.final.cram |
| **NA21125** | 1000G_2504_high_coverage/data/ERR3240077/NA21125.final.cram |
| **NA21127** | 1000G_2504_high_coverage/data/ERR3240078/NA21127.final.cram |
| **NA21128** | 1000G_2504_high_coverage/data/ERR3240079/NA21128.final.cram |
| **NA21130** | 1000G_2504_high_coverage/data/ERR3240080/NA21130.final.cram |
| **NA21133** | 1000G_2504_high_coverage/data/ERR3240081/NA21133.final.cram |
| **NA21135** | 1000G_2504_high_coverage/data/ERR3240082/NA21135.final.cram |
| **NA21137** | 1000G_2504_high_coverage/data/ERR3240083/NA21137.final.cram |
| **NA21141** | 1000G_2504_high_coverage/data/ERR3240084/NA21141.final.cram |
| **NA21142** | 1000G_2504_high_coverage/data/ERR3240085/NA21142.final.cram |
| **NA21143** | 1000G_2504_high_coverage/data/ERR3240086/NA21143.final.cram |
| **NA21144** | 1000G_2504_high_coverage/data/ERR3240087/NA21144.final.cram |
| **NA19734** | 1000G_2504_high_coverage/data/ERR3240088/NA19734.final.cram |
| **NA19735** | 1000G_2504_high_coverage/data/ERR3240089/NA19735.final.cram |
| **NA19740** | 1000G_2504_high_coverage/data/ERR3240090/NA19740.final.cram |
| **NA19741** | 1000G_2504_high_coverage/data/ERR3240091/NA19741.final.cram |
| **NA19752** | 1000G_2504_high_coverage/data/ERR3240092/NA19752.final.cram |
| **NA19764** | 1000G_2504_high_coverage/data/ERR3240093/NA19764.final.cram |
| **NA19792** | 1000G_2504_high_coverage/data/ERR3240094/NA19792.final.cram |
| **NA19922** | 1000G_2504_high_coverage/data/ERR3240095/NA19922.final.cram |
| **NA19923** | 1000G_2504_high_coverage/data/ERR3240096/NA19923.final.cram |
| **NA19984** | 1000G_2504_high_coverage/data/ERR3240097/NA19984.final.cram |
| **NA20298** | 1000G_2504_high_coverage/data/ERR3240098/NA20298.final.cram |
| **NA20339** | 1000G_2504_high_coverage/data/ERR3240099/NA20339.final.cram |
| **NA20351** | 1000G_2504_high_coverage/data/ERR3240100/NA20351.final.cram |
| **NA20412** | 1000G_2504_high_coverage/data/ERR3240101/NA20412.final.cram |
| **NA20863** | 1000G_2504_high_coverage/data/ERR3240102/NA20863.final.cram |
| **NA20864** | 1000G_2504_high_coverage/data/ERR3240103/NA20864.final.cram |
| **NA20867** | 1000G_2504_high_coverage/data/ERR3240104/NA20867.final.cram |
| **NA20868** | 1000G_2504_high_coverage/data/ERR3240105/NA20868.final.cram |
| **NA20905** | 1000G_2504_high_coverage/data/ERR3240106/NA20905.final.cram |
| **NA21087** | 1000G_2504_high_coverage/data/ERR3240107/NA21087.final.cram |
| **NA21093** | 1000G_2504_high_coverage/data/ERR3240108/NA21093.final.cram |
| **NA21095** | 1000G_2504_high_coverage/data/ERR3240109/NA21095.final.cram |
| **NA21114** | 1000G_2504_high_coverage/data/ERR3240110/NA21114.final.cram |
| **NA21124** | 1000G_2504_high_coverage/data/ERR3240111/NA21124.final.cram |
| **NA21126** | 1000G_2504_high_coverage/data/ERR3240112/NA21126.final.cram |
| **NA21129** | 1000G_2504_high_coverage/data/ERR3240113/NA21129.final.cram |
| **HG00096** | 1000G_2504_high_coverage/data/ERR3240114/HG00096.final.cram |
| **HG00097** | 1000G_2504_high_coverage/data/ERR3240115/HG00097.final.cram |
| **HG00099** | 1000G_2504_high_coverage/data/ERR3240116/HG00099.final.cram |
| **HG00100** | 1000G_2504_high_coverage/data/ERR3240117/HG00100.final.cram |

| | |
|---|---|
| **HG00101** | 1000G_2504_high_coverage/data/ERR3240118/HG00101.final.cram |
| **HG00102** | 1000G_2504_high_coverage/data/ERR3240119/HG00102.final.cram |
| **HG00103** | 1000G_2504_high_coverage/data/ERR3240120/HG00103.final.cram |
| **HG00105** | 1000G_2504_high_coverage/data/ERR3240121/HG00105.final.cram |
| **HG00106** | 1000G_2504_high_coverage/data/ERR3240122/HG00106.final.cram |
| **HG00107** | 1000G_2504_high_coverage/data/ERR3240123/HG00107.final.cram |
| **HG00108** | 1000G_2504_high_coverage/data/ERR3240124/HG00108.final.cram |
| **HG00109** | 1000G_2504_high_coverage/data/ERR3240125/HG00109.final.cram |
| **HG00110** | 1000G_2504_high_coverage/data/ERR3240126/HG00110.final.cram |
| **HG00111** | 1000G_2504_high_coverage/data/ERR3240127/HG00111.final.cram |
| **HG00112** | 1000G_2504_high_coverage/data/ERR3240128/HG00112.final.cram |
| **HG00113** | 1000G_2504_high_coverage/data/ERR3240129/HG00113.final.cram |
| **HG00114** | 1000G_2504_high_coverage/data/ERR3240130/HG00114.final.cram |
| **HG00115** | 1000G_2504_high_coverage/data/ERR3240131/HG00115.final.cram |
| **HG00116** | 1000G_2504_high_coverage/data/ERR3240132/HG00116.final.cram |
| **HG00117** | 1000G_2504_high_coverage/data/ERR3240133/HG00117.final.cram |
| **HG00120** | 1000G_2504_high_coverage/data/ERR3240134/HG00120.final.cram |
| **HG00127** | 1000G_2504_high_coverage/data/ERR3240135/HG00127.final.cram |
| **HG00128** | 1000G_2504_high_coverage/data/ERR3240136/HG00128.final.cram |
| **HG00129** | 1000G_2504_high_coverage/data/ERR3240137/HG00129.final.cram |
| **HG00130** | 1000G_2504_high_coverage/data/ERR3240138/HG00130.final.cram |
| **HG00131** | 1000G_2504_high_coverage/data/ERR3240139/HG00131.final.cram |
| **HG00132** | 1000G_2504_high_coverage/data/ERR3240140/HG00132.final.cram |
| **HG00133** | 1000G_2504_high_coverage/data/ERR3240141/HG00133.final.cram |
| **HG00136** | 1000G_2504_high_coverage/data/ERR3240142/HG00136.final.cram |
| **HG00137** | 1000G_2504_high_coverage/data/ERR3240143/HG00137.final.cram |
| **HG00138** | 1000G_2504_high_coverage/data/ERR3240144/HG00138.final.cram |
| **HG00139** | 1000G_2504_high_coverage/data/ERR3240145/HG00139.final.cram |
| **HG00140** | 1000G_2504_high_coverage/data/ERR3240146/HG00140.final.cram |
| **HG00141** | 1000G_2504_high_coverage/data/ERR3240147/HG00141.final.cram |
| **HG00142** | 1000G_2504_high_coverage/data/ERR3240148/HG00142.final.cram |
| **HG00143** | 1000G_2504_high_coverage/data/ERR3240149/HG00143.final.cram |
| **HG00145** | 1000G_2504_high_coverage/data/ERR3240150/HG00145.final.cram |
| **HG00146** | 1000G_2504_high_coverage/data/ERR3240151/HG00146.final.cram |
| **HG00148** | 1000G_2504_high_coverage/data/ERR3240152/HG00148.final.cram |
| **HG00149** | 1000G_2504_high_coverage/data/ERR3240153/HG00149.final.cram |
| **HG00150** | 1000G_2504_high_coverage/data/ERR3240154/HG00150.final.cram |
| **HG00151** | 1000G_2504_high_coverage/data/ERR3240155/HG00151.final.cram |
| **HG00155** | 1000G_2504_high_coverage/data/ERR3240156/HG00155.final.cram |
| **HG00157** | 1000G_2504_high_coverage/data/ERR3240157/HG00157.final.cram |
| **HG00158** | 1000G_2504_high_coverage/data/ERR3240158/HG00158.final.cram |
| **HG00160** | 1000G_2504_high_coverage/data/ERR3240159/HG00160.final.cram |
| **HG00171** | 1000G_2504_high_coverage/data/ERR3240160/HG00171.final.cram |
| **HG00173** | 1000G_2504_high_coverage/data/ERR3240161/HG00173.final.cram |
| **HG00174** | 1000G_2504_high_coverage/data/ERR3240162/HG00174.final.cram |
| **HG00176** | 1000G_2504_high_coverage/data/ERR3240163/HG00176.final.cram |
| **HG00177** | 1000G_2504_high_coverage/data/ERR3240164/HG00177.final.cram |
| **HG00178** | 1000G_2504_high_coverage/data/ERR3240165/HG00178.final.cram |
| **HG00179** | 1000G_2504_high_coverage/data/ERR3240166/HG00179.final.cram |
| **HG00180** | 1000G_2504_high_coverage/data/ERR3240167/HG00180.final.cram |
| **HG00181** | 1000G_2504_high_coverage/data/ERR3240168/HG00181.final.cram |
| **HG00182** | 1000G_2504_high_coverage/data/ERR3240169/HG00182.final.cram |
| **HG00183** | 1000G_2504_high_coverage/data/ERR3240170/HG00183.final.cram |
| **HG00185** | 1000G_2504_high_coverage/data/ERR3240171/HG00185.final.cram |
| **HG00186** | 1000G_2504_high_coverage/data/ERR3240172/HG00186.final.cram |
| **HG00187** | 1000G_2504_high_coverage/data/ERR3240173/HG00187.final.cram |
| **HG00188** | 1000G_2504_high_coverage/data/ERR3240174/HG00188.final.cram |
| **HG00189** | 1000G_2504_high_coverage/data/ERR3240175/HG00189.final.cram |
| **HG00190** | 1000G_2504_high_coverage/data/ERR3240176/HG00190.final.cram |
| **HG00233** | 1000G_2504_high_coverage/data/ERR3240177/HG00233.final.cram |
| **HG00234** | 1000G_2504_high_coverage/data/ERR3240178/HG00234.final.cram |
| **HG00235** | 1000G_2504_high_coverage/data/ERR3240179/HG00235.final.cram |
| **HG00238** | 1000G_2504_high_coverage/data/ERR3240180/HG00238.final.cram |
| **HG00240** | 1000G_2504_high_coverage/data/ERR3240181/HG00240.final.cram |
| **HG00255** | 1000G_2504_high_coverage/data/ERR3240182/HG00255.final.cram |

| | |
|---|---|
| **HG00442** | 1000G_2504_high_coverage/data/ERR3240183/HG00442.final.cram |
| **HG00443** | 1000G_2504_high_coverage/data/ERR3240184/HG00443.final.cram |
| **HG00445** | 1000G_2504_high_coverage/data/ERR3240185/HG00445.final.cram |
| **HG00446** | 1000G_2504_high_coverage/data/ERR3240186/HG00446.final.cram |
| **HG00448** | 1000G_2504_high_coverage/data/ERR3240187/HG00448.final.cram |
| **HG00449** | 1000G_2504_high_coverage/data/ERR3240188/HG00449.final.cram |
| **HG00451** | 1000G_2504_high_coverage/data/ERR3240189/HG00451.final.cram |
| **HG00452** | 1000G_2504_high_coverage/data/ERR3240190/HG00452.final.cram |
| **HG00463** | 1000G_2504_high_coverage/data/ERR3240191/HG00463.final.cram |
| **HG00464** | 1000G_2504_high_coverage/data/ERR3240192/HG00464.final.cram |
| **HG00118** | 1000G_2504_high_coverage/data/ERR3240193/HG00118.final.cram |
| **HG00119** | 1000G_2504_high_coverage/data/ERR3240194/HG00119.final.cram |
| **HG00121** | 1000G_2504_high_coverage/data/ERR3240195/HG00121.final.cram |
| **HG00122** | 1000G_2504_high_coverage/data/ERR3240196/HG00122.final.cram |
| **HG00123** | 1000G_2504_high_coverage/data/ERR3240197/HG00123.final.cram |
| **HG00125** | 1000G_2504_high_coverage/data/ERR3240198/HG00125.final.cram |
| **HG00126** | 1000G_2504_high_coverage/data/ERR3240199/HG00126.final.cram |
| **HG00159** | 1000G_2504_high_coverage/data/ERR3240200/HG00159.final.cram |
| **HG00231** | 1000G_2504_high_coverage/data/ERR3240201/HG00231.final.cram |
| **HG00232** | 1000G_2504_high_coverage/data/ERR3240202/HG00232.final.cram |
| **HG00236** | 1000G_2504_high_coverage/data/ERR3240203/HG00236.final.cram |
| **HG00237** | 1000G_2504_high_coverage/data/ERR3240204/HG00237.final.cram |
| **HG00239** | 1000G_2504_high_coverage/data/ERR3240205/HG00239.final.cram |
| **HG00242** | 1000G_2504_high_coverage/data/ERR3240206/HG00242.final.cram |
| **HG00243** | 1000G_2504_high_coverage/data/ERR3240207/HG00243.final.cram |
| **HG00244** | 1000G_2504_high_coverage/data/ERR3240208/HG00244.final.cram |
| **HG00245** | 1000G_2504_high_coverage/data/ERR3240209/HG00245.final.cram |
| **HG00252** | 1000G_2504_high_coverage/data/ERR3240210/HG00252.final.cram |
| **HG00253** | 1000G_2504_high_coverage/data/ERR3240211/HG00253.final.cram |
| **HG00254** | 1000G_2504_high_coverage/data/ERR3240212/HG00254.final.cram |
| **HG00256** | 1000G_2504_high_coverage/data/ERR3240213/HG00256.final.cram |
| **HG00257** | 1000G_2504_high_coverage/data/ERR3240214/HG00257.final.cram |
| **HG00258** | 1000G_2504_high_coverage/data/ERR3240215/HG00258.final.cram |
| **HG00259** | 1000G_2504_high_coverage/data/ERR3240216/HG00259.final.cram |
| **HG00260** | 1000G_2504_high_coverage/data/ERR3240217/HG00260.final.cram |
| **HG00261** | 1000G_2504_high_coverage/data/ERR3240218/HG00261.final.cram |
| **HG00262** | 1000G_2504_high_coverage/data/ERR3240219/HG00262.final.cram |
| **HG00263** | 1000G_2504_high_coverage/data/ERR3240220/HG00263.final.cram |
| **HG00264** | 1000G_2504_high_coverage/data/ERR3240221/HG00264.final.cram |
| **HG00265** | 1000G_2504_high_coverage/data/ERR3240222/HG00265.final.cram |
| **HG00266** | 1000G_2504_high_coverage/data/ERR3240223/HG00266.final.cram |
| **HG00267** | 1000G_2504_high_coverage/data/ERR3240224/HG00267.final.cram |
| **HG00268** | 1000G_2504_high_coverage/data/ERR3240225/HG00268.final.cram |
| **HG00269** | 1000G_2504_high_coverage/data/ERR3240226/HG00269.final.cram |
| **HG00271** | 1000G_2504_high_coverage/data/ERR3240227/HG00271.final.cram |
| **HG00272** | 1000G_2504_high_coverage/data/ERR3240228/HG00272.final.cram |
| **HG00273** | 1000G_2504_high_coverage/data/ERR3240229/HG00273.final.cram |
| **HG00306** | 1000G_2504_high_coverage/data/ERR3240230/HG00306.final.cram |
| **HG00308** | 1000G_2504_high_coverage/data/ERR3240231/HG00308.final.cram |
| **HG00309** | 1000G_2504_high_coverage/data/ERR3240232/HG00309.final.cram |
| **HG00311** | 1000G_2504_high_coverage/data/ERR3240233/HG00311.final.cram |
| **HG00313** | 1000G_2504_high_coverage/data/ERR3240234/HG00313.final.cram |
| **HG00315** | 1000G_2504_high_coverage/data/ERR3240235/HG00315.final.cram |
| **HG00318** | 1000G_2504_high_coverage/data/ERR3240236/HG00318.final.cram |
| **HG00319** | 1000G_2504_high_coverage/data/ERR3240237/HG00319.final.cram |
| **HG00320** | 1000G_2504_high_coverage/data/ERR3240238/HG00320.final.cram |
| **HG00321** | 1000G_2504_high_coverage/data/ERR3240239/HG00321.final.cram |
| **HG00323** | 1000G_2504_high_coverage/data/ERR3240240/HG00323.final.cram |
| **HG00327** | 1000G_2504_high_coverage/data/ERR3240241/HG00327.final.cram |
| **HG00328** | 1000G_2504_high_coverage/data/ERR3240242/HG00328.final.cram |
| **HG00335** | 1000G_2504_high_coverage/data/ERR3240243/HG00335.final.cram |
| **HG00336** | 1000G_2504_high_coverage/data/ERR3240244/HG00336.final.cram |
| **HG00337** | 1000G_2504_high_coverage/data/ERR3240245/HG00337.final.cram |
| **HG00338** | 1000G_2504_high_coverage/data/ERR3240246/HG00338.final.cram |
| **HG00339** | 1000G_2504_high_coverage/data/ERR3240247/HG00339.final.cram |

| | |
|---|---|
| HG00342 | 1000G_2504_high_coverage/data/ERR3240248/HG00342.final.cram |
| HG00343 | 1000G_2504_high_coverage/data/ERR3240249/HG00343.final.cram |
| HG00344 | 1000G_2504_high_coverage/data/ERR3240250/HG00344.final.cram |
| HG00345 | 1000G_2504_high_coverage/data/ERR3240251/HG00345.final.cram |
| HG00346 | 1000G_2504_high_coverage/data/ERR3240252/HG00346.final.cram |
| HG00349 | 1000G_2504_high_coverage/data/ERR3240253/HG00349.final.cram |
| HG00350 | 1000G_2504_high_coverage/data/ERR3240254/HG00350.final.cram |
| HG00351 | 1000G_2504_high_coverage/data/ERR3240255/HG00351.final.cram |
| HG00353 | 1000G_2504_high_coverage/data/ERR3240256/HG00353.final.cram |
| HG00355 | 1000G_2504_high_coverage/data/ERR3240257/HG00355.final.cram |
| HG00356 | 1000G_2504_high_coverage/data/ERR3240258/HG00356.final.cram |
| HG00357 | 1000G_2504_high_coverage/data/ERR3240259/HG00357.final.cram |
| HG00358 | 1000G_2504_high_coverage/data/ERR3240260/HG00358.final.cram |
| HG00360 | 1000G_2504_high_coverage/data/ERR3240261/HG00360.final.cram |
| HG00361 | 1000G_2504_high_coverage/data/ERR3240262/HG00361.final.cram |
| HG00362 | 1000G_2504_high_coverage/data/ERR3240263/HG00362.final.cram |
| HG00364 | 1000G_2504_high_coverage/data/ERR3240264/HG00364.final.cram |
| HG00365 | 1000G_2504_high_coverage/data/ERR3240265/HG00365.final.cram |
| HG00366 | 1000G_2504_high_coverage/data/ERR3240266/HG00366.final.cram |
| HG00367 | 1000G_2504_high_coverage/data/ERR3240267/HG00367.final.cram |
| HG00368 | 1000G_2504_high_coverage/data/ERR3240268/HG00368.final.cram |
| HG00369 | 1000G_2504_high_coverage/data/ERR3240269/HG00369.final.cram |
| HG00371 | 1000G_2504_high_coverage/data/ERR3240270/HG00371.final.cram |
| HG00372 | 1000G_2504_high_coverage/data/ERR3240271/HG00372.final.cram |
| HG00373 | 1000G_2504_high_coverage/data/ERR3240272/HG00373.final.cram |
| HG00375 | 1000G_2504_high_coverage/data/ERR3240273/HG00375.final.cram |
| HG00376 | 1000G_2504_high_coverage/data/ERR3240274/HG00376.final.cram |
| HG00378 | 1000G_2504_high_coverage/data/ERR3241659/HG00378.final.cram |
| HG00379 | 1000G_2504_high_coverage/data/ERR3241660/HG00379.final.cram |
| HG00380 | 1000G_2504_high_coverage/data/ERR3241661/HG00380.final.cram |
| HG00381 | 1000G_2504_high_coverage/data/ERR3241662/HG00381.final.cram |
| HG00382 | 1000G_2504_high_coverage/data/ERR3241663/HG00382.final.cram |
| HG00384 | 1000G_2504_high_coverage/data/ERR3241664/HG00384.final.cram |
| HG00403 | 1000G_2504_high_coverage/data/ERR3241665/HG00403.final.cram |
| HG00404 | 1000G_2504_high_coverage/data/ERR3241666/HG00404.final.cram |
| HG00406 | 1000G_2504_high_coverage/data/ERR3241667/HG00406.final.cram |
| HG00407 | 1000G_2504_high_coverage/data/ERR3241668/HG00407.final.cram |
| HG00419 | 1000G_2504_high_coverage/data/ERR3241669/HG00419.final.cram |
| HG00421 | 1000G_2504_high_coverage/data/ERR3241670/HG00421.final.cram |
| HG00422 | 1000G_2504_high_coverage/data/ERR3241671/HG00422.final.cram |
| HG00428 | 1000G_2504_high_coverage/data/ERR3241672/HG00428.final.cram |
| HG00436 | 1000G_2504_high_coverage/data/ERR3241673/HG00436.final.cram |
| HG00437 | 1000G_2504_high_coverage/data/ERR3241674/HG00437.final.cram |
| HG00457 | 1000G_2504_high_coverage/data/ERR3241675/HG00457.final.cram |
| HG00458 | 1000G_2504_high_coverage/data/ERR3241676/HG00458.final.cram |
| HG00472 | 1000G_2504_high_coverage/data/ERR3241677/HG00472.final.cram |
| HG00473 | 1000G_2504_high_coverage/data/ERR3241678/HG00473.final.cram |
| HG00475 | 1000G_2504_high_coverage/data/ERR3241679/HG00475.final.cram |
| HG00476 | 1000G_2504_high_coverage/data/ERR3241680/HG00476.final.cram |
| HG00478 | 1000G_2504_high_coverage/data/ERR3241681/HG00478.final.cram |
| HG00479 | 1000G_2504_high_coverage/data/ERR3241682/HG00479.final.cram |
| HG00500 | 1000G_2504_high_coverage/data/ERR3241683/HG00500.final.cram |
| HG00513 | 1000G_2504_high_coverage/data/ERR3241684/HG00513.final.cram |
| HG00524 | 1000G_2504_high_coverage/data/ERR3241685/HG00524.final.cram |
| HG00525 | 1000G_2504_high_coverage/data/ERR3241686/HG00525.final.cram |
| HG00530 | 1000G_2504_high_coverage/data/ERR3241687/HG00530.final.cram |
| HG00531 | 1000G_2504_high_coverage/data/ERR3241688/HG00531.final.cram |
| HG00533 | 1000G_2504_high_coverage/data/ERR3241689/HG00533.final.cram |
| HG00534 | 1000G_2504_high_coverage/data/ERR3241690/HG00534.final.cram |
| HG00536 | 1000G_2504_high_coverage/data/ERR3241691/HG00536.final.cram |
| HG00537 | 1000G_2504_high_coverage/data/ERR3241692/HG00537.final.cram |
| HG00542 | 1000G_2504_high_coverage/data/ERR3241693/HG00542.final.cram |
| HG00543 | 1000G_2504_high_coverage/data/ERR3241694/HG00543.final.cram |
| HG00553 | 1000G_2504_high_coverage/data/ERR3241695/HG00553.final.cram |
| HG00554 | 1000G_2504_high_coverage/data/ERR3241696/HG00554.final.cram |

| | |
|---|---|
| **HG00556** | 1000G_2504_high_coverage/data/ERR3241697/HG00556.final.cram |
| **HG00557** | 1000G_2504_high_coverage/data/ERR3241698/HG00557.final.cram |
| **HG00559** | 1000G_2504_high_coverage/data/ERR3241699/HG00559.final.cram |
| **HG00560** | 1000G_2504_high_coverage/data/ERR3241700/HG00560.final.cram |
| **HG00565** | 1000G_2504_high_coverage/data/ERR3241701/HG00565.final.cram |
| **HG00566** | 1000G_2504_high_coverage/data/ERR3241702/HG00566.final.cram |
| **HG00580** | 1000G_2504_high_coverage/data/ERR3241703/HG00580.final.cram |
| **HG00581** | 1000G_2504_high_coverage/data/ERR3241704/HG00581.final.cram |
| **HG00583** | 1000G_2504_high_coverage/data/ERR3241705/HG00583.final.cram |
| **HG00584** | 1000G_2504_high_coverage/data/ERR3241706/HG00584.final.cram |
| **HG00589** | 1000G_2504_high_coverage/data/ERR3241707/HG00589.final.cram |
| **HG00590** | 1000G_2504_high_coverage/data/ERR3241708/HG00590.final.cram |
| **HG00592** | 1000G_2504_high_coverage/data/ERR3241709/HG00592.final.cram |
| **HG00593** | 1000G_2504_high_coverage/data/ERR3241710/HG00593.final.cram |
| **HG00595** | 1000G_2504_high_coverage/data/ERR3241711/HG00595.final.cram |
| **HG00596** | 1000G_2504_high_coverage/data/ERR3241712/HG00596.final.cram |
| **HG00607** | 1000G_2504_high_coverage/data/ERR3241713/HG00607.final.cram |
| **HG00608** | 1000G_2504_high_coverage/data/ERR3241714/HG00608.final.cram |
| **HG00610** | 1000G_2504_high_coverage/data/ERR3241715/HG00610.final.cram |
| **HG00611** | 1000G_2504_high_coverage/data/ERR3241716/HG00611.final.cram |
| **HG00613** | 1000G_2504_high_coverage/data/ERR3241717/HG00613.final.cram |
| **HG00614** | 1000G_2504_high_coverage/data/ERR3241718/HG00614.final.cram |
| **HG00619** | 1000G_2504_high_coverage/data/ERR3241719/HG00619.final.cram |
| **HG00620** | 1000G_2504_high_coverage/data/ERR3241720/HG00620.final.cram |
| **HG00625** | 1000G_2504_high_coverage/data/ERR3241721/HG00625.final.cram |
| **HG00626** | 1000G_2504_high_coverage/data/ERR3241722/HG00626.final.cram |
| **HG00628** | 1000G_2504_high_coverage/data/ERR3241723/HG00628.final.cram |
| **HG00629** | 1000G_2504_high_coverage/data/ERR3241724/HG00629.final.cram |
| **HG00634** | 1000G_2504_high_coverage/data/ERR3241725/HG00634.final.cram |
| **HG00637** | 1000G_2504_high_coverage/data/ERR3241726/HG00637.final.cram |
| **HG00638** | 1000G_2504_high_coverage/data/ERR3241727/HG00638.final.cram |
| **HG00640** | 1000G_2504_high_coverage/data/ERR3241728/HG00640.final.cram |
| **HG00641** | 1000G_2504_high_coverage/data/ERR3241729/HG00641.final.cram |
| **HG00650** | 1000G_2504_high_coverage/data/ERR3241730/HG00650.final.cram |
| **HG00651** | 1000G_2504_high_coverage/data/ERR3241731/HG00651.final.cram |
| **HG00653** | 1000G_2504_high_coverage/data/ERR3241732/HG00653.final.cram |
| **HG00654** | 1000G_2504_high_coverage/data/ERR3241733/HG00654.final.cram |
| **HG00656** | 1000G_2504_high_coverage/data/ERR3241734/HG00656.final.cram |
| **HG00657** | 1000G_2504_high_coverage/data/ERR3241735/HG00657.final.cram |
| **HG00662** | 1000G_2504_high_coverage/data/ERR3241736/HG00662.final.cram |
| **HG00663** | 1000G_2504_high_coverage/data/ERR3241737/HG00663.final.cram |
| **HG00671** | 1000G_2504_high_coverage/data/ERR3241738/HG00671.final.cram |
| **HG00672** | 1000G_2504_high_coverage/data/ERR3241739/HG00672.final.cram |
| **HG00683** | 1000G_2504_high_coverage/data/ERR3241740/HG00683.final.cram |
| **HG00684** | 1000G_2504_high_coverage/data/ERR3241741/HG00684.final.cram |
| **HG00689** | 1000G_2504_high_coverage/data/ERR3241742/HG00689.final.cram |
| **HG00690** | 1000G_2504_high_coverage/data/ERR3241743/HG00690.final.cram |
| **HG00692** | 1000G_2504_high_coverage/data/ERR3241744/HG00692.final.cram |
| **HG00693** | 1000G_2504_high_coverage/data/ERR3241745/HG00693.final.cram |
| **HG00698** | 1000G_2504_high_coverage/data/ERR3241746/HG00698.final.cram |
| **HG00699** | 1000G_2504_high_coverage/data/ERR3241747/HG00699.final.cram |
| **HG00701** | 1000G_2504_high_coverage/data/ERR3241748/HG00701.final.cram |
| **HG00704** | 1000G_2504_high_coverage/data/ERR3241749/HG00704.final.cram |
| **HG00705** | 1000G_2504_high_coverage/data/ERR3241750/HG00705.final.cram |
| **HG00707** | 1000G_2504_high_coverage/data/ERR3241751/HG00707.final.cram |
| **HG00708** | 1000G_2504_high_coverage/data/ERR3241752/HG00708.final.cram |
| **HG00717** | 1000G_2504_high_coverage/data/ERR3241753/HG00717.final.cram |
| **HG00731** | 1000G_2504_high_coverage/data/ERR3241754/HG00731.final.cram |
| **HG00732** | 1000G_2504_high_coverage/data/ERR3241755/HG00732.final.cram |
| **HG00736** | 1000G_2504_high_coverage/data/ERR3241756/HG00736.final.cram |
| **HG00737** | 1000G_2504_high_coverage/data/ERR3241757/HG00737.final.cram |
| **HG00739** | 1000G_2504_high_coverage/data/ERR3241758/HG00739.final.cram |
| **HG00740** | 1000G_2504_high_coverage/data/ERR3241759/HG00740.final.cram |
| **HG01047** | 1000G_2504_high_coverage/data/ERR3241760/HG01047.final.cram |
| **HG01048** | 1000G_2504_high_coverage/data/ERR3241761/HG01048.final.cram |

| | |
|---|---|
| **HG01049** | 1000G_2504_high_coverage/data/ERR3241762/HG01049.final.cram |
| **HG01054** | 1000G_2504_high_coverage/data/ERR3241763/HG01054.final.cram |
| **HG01055** | 1000G_2504_high_coverage/data/ERR3241764/HG01055.final.cram |
| **HG01066** | 1000G_2504_high_coverage/data/ERR3241765/HG01066.final.cram |
| **HG01067** | 1000G_2504_high_coverage/data/ERR3241766/HG01067.final.cram |
| **HG01079** | 1000G_2504_high_coverage/data/ERR3241767/HG01079.final.cram |
| **HG01080** | 1000G_2504_high_coverage/data/ERR3241768/HG01080.final.cram |
| **HG01097** | 1000G_2504_high_coverage/data/ERR3241769/HG01097.final.cram |
| **HG01098** | 1000G_2504_high_coverage/data/ERR3241770/HG01098.final.cram |
| **NA19913** | 1000G_2504_high_coverage/data/ERR3241771/NA19913.final.cram |
| **NA20274** | 1000G_2504_high_coverage/data/ERR3241772/NA20274.final.cram |
| **NA20318** | 1000G_2504_high_coverage/data/ERR3241773/NA20318.final.cram |
| **NA20320** | 1000G_2504_high_coverage/data/ERR3241774/NA20320.final.cram |
| **NA20321** | 1000G_2504_high_coverage/data/ERR3241775/NA20321.final.cram |
| **NA20355** | 1000G_2504_high_coverage/data/ERR3241776/NA20355.final.cram |
| **NA20362** | 1000G_2504_high_coverage/data/ERR3241777/NA20362.final.cram |
| **HG00734** | 1000G_2504_high_coverage/data/ERR3241778/HG00734.final.cram |
| **HG00154** | 1000G_2504_high_coverage/data/ERR3241779/HG00154.final.cram |
| **HG00246** | 1000G_2504_high_coverage/data/ERR3241780/HG00246.final.cram |
| **HG00250** | 1000G_2504_high_coverage/data/ERR3241781/HG00250.final.cram |
| **HG00251** | 1000G_2504_high_coverage/data/ERR3241782/HG00251.final.cram |
| **HG00274** | 1000G_2504_high_coverage/data/ERR3241783/HG00274.final.cram |
| **HG00275** | 1000G_2504_high_coverage/data/ERR3241784/HG00275.final.cram |
| **HG00276** | 1000G_2504_high_coverage/data/ERR3241785/HG00276.final.cram |
| **HG00277** | 1000G_2504_high_coverage/data/ERR3241786/HG00277.final.cram |
| **HG00278** | 1000G_2504_high_coverage/data/ERR3241787/HG00278.final.cram |
| **HG00280** | 1000G_2504_high_coverage/data/ERR3241788/HG00280.final.cram |
| **HG00281** | 1000G_2504_high_coverage/data/ERR3241789/HG00281.final.cram |
| **HG00282** | 1000G_2504_high_coverage/data/ERR3241790/HG00282.final.cram |
| **HG00284** | 1000G_2504_high_coverage/data/ERR3241791/HG00284.final.cram |
| **HG00285** | 1000G_2504_high_coverage/data/ERR3241792/HG00285.final.cram |
| **HG00310** | 1000G_2504_high_coverage/data/ERR3241793/HG00310.final.cram |
| **HG00324** | 1000G_2504_high_coverage/data/ERR3241794/HG00324.final.cram |
| **HG00325** | 1000G_2504_high_coverage/data/ERR3241795/HG00325.final.cram |
| **HG00326** | 1000G_2504_high_coverage/data/ERR3241796/HG00326.final.cram |
| **HG00329** | 1000G_2504_high_coverage/data/ERR3241797/HG00329.final.cram |
| **HG00330** | 1000G_2504_high_coverage/data/ERR3241798/HG00330.final.cram |
| **HG00331** | 1000G_2504_high_coverage/data/ERR3241799/HG00331.final.cram |
| **HG00332** | 1000G_2504_high_coverage/data/ERR3241800/HG00332.final.cram |
| **HG00334** | 1000G_2504_high_coverage/data/ERR3241801/HG00334.final.cram |
| **HG00341** | 1000G_2504_high_coverage/data/ERR3241802/HG00341.final.cram |
| **HG00383** | 1000G_2504_high_coverage/data/ERR3241803/HG00383.final.cram |
| **HG00551** | 1000G_2504_high_coverage/data/ERR3241804/HG00551.final.cram |
| **HG01051** | 1000G_2504_high_coverage/data/ERR3241805/HG01051.final.cram |
| **HG01052** | 1000G_2504_high_coverage/data/ERR3241806/HG01052.final.cram |
| **HG01060** | 1000G_2504_high_coverage/data/ERR3241807/HG01060.final.cram |
| **HG01061** | 1000G_2504_high_coverage/data/ERR3241808/HG01061.final.cram |
| **HG01069** | 1000G_2504_high_coverage/data/ERR3241809/HG01069.final.cram |
| **HG01070** | 1000G_2504_high_coverage/data/ERR3241810/HG01070.final.cram |
| **HG01072** | 1000G_2504_high_coverage/data/ERR3241811/HG01072.final.cram |
| **HG01073** | 1000G_2504_high_coverage/data/ERR3241812/HG01073.final.cram |
| **HG01075** | 1000G_2504_high_coverage/data/ERR3241813/HG01075.final.cram |
| **HG01082** | 1000G_2504_high_coverage/data/ERR3241814/HG01082.final.cram |
| **HG01083** | 1000G_2504_high_coverage/data/ERR3241815/HG01083.final.cram |
| **HG01085** | 1000G_2504_high_coverage/data/ERR3241816/HG01085.final.cram |
| **HG01086** | 1000G_2504_high_coverage/data/ERR3241817/HG01086.final.cram |
| **HG01094** | 1000G_2504_high_coverage/data/ERR3241818/HG01094.final.cram |
| **HG01095** | 1000G_2504_high_coverage/data/ERR3241819/HG01095.final.cram |
| **HG01101** | 1000G_2504_high_coverage/data/ERR3241820/HG01101.final.cram |
| **HG01102** | 1000G_2504_high_coverage/data/ERR3241821/HG01102.final.cram |
| **HG01104** | 1000G_2504_high_coverage/data/ERR3241822/HG01104.final.cram |
| **HG01105** | 1000G_2504_high_coverage/data/ERR3241823/HG01105.final.cram |
| **HG01107** | 1000G_2504_high_coverage/data/ERR3241824/HG01107.final.cram |
| **HG01108** | 1000G_2504_high_coverage/data/ERR3241825/HG01108.final.cram |
| **HG01110** | 1000G_2504_high_coverage/data/ERR3241826/HG01110.final.cram |

| | |
|---|---|
| **HG01111** | 1000G_2504_high_coverage/data/ERR3241827/HG01111.final.cram |
| **HG01112** | 1000G_2504_high_coverage/data/ERR3241828/HG01112.final.cram |
| **HG01113** | 1000G_2504_high_coverage/data/ERR3241829/HG01113.final.cram |
| **HG01124** | 1000G_2504_high_coverage/data/ERR3241830/HG01124.final.cram |
| **HG01125** | 1000G_2504_high_coverage/data/ERR3241831/HG01125.final.cram |
| **HG01133** | 1000G_2504_high_coverage/data/ERR3241832/HG01133.final.cram |
| **HG01134** | 1000G_2504_high_coverage/data/ERR3241833/HG01134.final.cram |
| **HG01136** | 1000G_2504_high_coverage/data/ERR3241834/HG01136.final.cram |
| **HG01137** | 1000G_2504_high_coverage/data/ERR3241835/HG01137.final.cram |
| **HG01139** | 1000G_2504_high_coverage/data/ERR3241836/HG01139.final.cram |
| **HG01140** | 1000G_2504_high_coverage/data/ERR3241837/HG01140.final.cram |
| **HG01148** | 1000G_2504_high_coverage/data/ERR3241838/HG01148.final.cram |
| **HG01149** | 1000G_2504_high_coverage/data/ERR3241839/HG01149.final.cram |
| **HG01167** | 1000G_2504_high_coverage/data/ERR3241840/HG01167.final.cram |
| **HG01168** | 1000G_2504_high_coverage/data/ERR3241841/HG01168.final.cram |
| **HG01170** | 1000G_2504_high_coverage/data/ERR3241842/HG01170.final.cram |
| **HG01171** | 1000G_2504_high_coverage/data/ERR3241843/HG01171.final.cram |
| **HG01173** | 1000G_2504_high_coverage/data/ERR3241844/HG01173.final.cram |
| **HG01174** | 1000G_2504_high_coverage/data/ERR3241845/HG01174.final.cram |
| **HG01176** | 1000G_2504_high_coverage/data/ERR3241846/HG01176.final.cram |
| **HG01177** | 1000G_2504_high_coverage/data/ERR3241847/HG01177.final.cram |
| **HG01182** | 1000G_2504_high_coverage/data/ERR3241848/HG01182.final.cram |
| **HG01183** | 1000G_2504_high_coverage/data/ERR3241849/HG01183.final.cram |
| **HG01187** | 1000G_2504_high_coverage/data/ERR3241850/HG01187.final.cram |
| **HG01188** | 1000G_2504_high_coverage/data/ERR3241851/HG01188.final.cram |
| **HG01190** | 1000G_2504_high_coverage/data/ERR3241852/HG01190.final.cram |
| **HG01191** | 1000G_2504_high_coverage/data/ERR3241853/HG01191.final.cram |
| **HG01197** | 1000G_2504_high_coverage/data/ERR3241854/HG01197.final.cram |
| **HG01198** | 1000G_2504_high_coverage/data/ERR3241855/HG01198.final.cram |
| **HG01204** | 1000G_2504_high_coverage/data/ERR3241856/HG01204.final.cram |
| **HG01205** | 1000G_2504_high_coverage/data/ERR3241857/HG01205.final.cram |
| **HG01241** | 1000G_2504_high_coverage/data/ERR3241858/HG01241.final.cram |
| **HG01242** | 1000G_2504_high_coverage/data/ERR3241859/HG01242.final.cram |
| **HG01247** | 1000G_2504_high_coverage/data/ERR3241860/HG01247.final.cram |
| **HG01248** | 1000G_2504_high_coverage/data/ERR3241861/HG01248.final.cram |
| **HG01250** | 1000G_2504_high_coverage/data/ERR3241862/HG01250.final.cram |
| **HG01251** | 1000G_2504_high_coverage/data/ERR3241863/HG01251.final.cram |
| **HG01253** | 1000G_2504_high_coverage/data/ERR3241864/HG01253.final.cram |
| **HG01254** | 1000G_2504_high_coverage/data/ERR3241865/HG01254.final.cram |
| **HG01256** | 1000G_2504_high_coverage/data/ERR3241866/HG01256.final.cram |
| **HG01257** | 1000G_2504_high_coverage/data/ERR3241867/HG01257.final.cram |
| **HG01259** | 1000G_2504_high_coverage/data/ERR3241868/HG01259.final.cram |
| **HG01260** | 1000G_2504_high_coverage/data/ERR3241869/HG01260.final.cram |
| **HG01271** | 1000G_2504_high_coverage/data/ERR3241870/HG01271.final.cram |
| **HG01272** | 1000G_2504_high_coverage/data/ERR3241871/HG01272.final.cram |
| **HG01275** | 1000G_2504_high_coverage/data/ERR3241872/HG01275.final.cram |
| **HG01277** | 1000G_2504_high_coverage/data/ERR3241873/HG01277.final.cram |
| **HG01334** | 1000G_2504_high_coverage/data/ERR3241874/HG01334.final.cram |
| **HG01344** | 1000G_2504_high_coverage/data/ERR3241875/HG01344.final.cram |
| **HG01345** | 1000G_2504_high_coverage/data/ERR3241876/HG01345.final.cram |
| **HG01348** | 1000G_2504_high_coverage/data/ERR3241877/HG01348.final.cram |
| **HG01356** | 1000G_2504_high_coverage/data/ERR3241878/HG01356.final.cram |
| **HG01357** | 1000G_2504_high_coverage/data/ERR3241879/HG01357.final.cram |
| **HG01377** | 1000G_2504_high_coverage/data/ERR3241880/HG01377.final.cram |
| **HG01378** | 1000G_2504_high_coverage/data/ERR3241881/HG01378.final.cram |
| **HG01383** | 1000G_2504_high_coverage/data/ERR3241882/HG01383.final.cram |
| **HG01384** | 1000G_2504_high_coverage/data/ERR3241883/HG01384.final.cram |
| **HG01341** | 1000G_2504_high_coverage/data/ERR3241884/HG01341.final.cram |
| **HG01342** | 1000G_2504_high_coverage/data/ERR3241885/HG01342.final.cram |
| **HG01350** | 1000G_2504_high_coverage/data/ERR3241886/HG01350.final.cram |
| **HG01351** | 1000G_2504_high_coverage/data/ERR3241887/HG01351.final.cram |
| **HG01353** | 1000G_2504_high_coverage/data/ERR3241888/HG01353.final.cram |
| **HG01354** | 1000G_2504_high_coverage/data/ERR3241889/HG01354.final.cram |
| **HG01359** | 1000G_2504_high_coverage/data/ERR3241890/HG01359.final.cram |
| **HG01360** | 1000G_2504_high_coverage/data/ERR3241891/HG01360.final.cram |

| | |
|---|---|
| **HG01365** | 1000G_2504_high_coverage/data/ERR3241892/HG01365.final.cram |
| **HG01366** | 1000G_2504_high_coverage/data/ERR3241893/HG01366.final.cram |
| **HG01374** | 1000G_2504_high_coverage/data/ERR3241894/HG01374.final.cram |
| **HG01375** | 1000G_2504_high_coverage/data/ERR3241895/HG01375.final.cram |
| **HG01389** | 1000G_2504_high_coverage/data/ERR3241896/HG01389.final.cram |
| **HG01390** | 1000G_2504_high_coverage/data/ERR3241897/HG01390.final.cram |
| **HG01437** | 1000G_2504_high_coverage/data/ERR3241898/HG01437.final.cram |
| **HG01438** | 1000G_2504_high_coverage/data/ERR3241899/HG01438.final.cram |
| **HG01440** | 1000G_2504_high_coverage/data/ERR3241900/HG01440.final.cram |
| **HG01441** | 1000G_2504_high_coverage/data/ERR3241901/HG01441.final.cram |
| **HG01455** | 1000G_2504_high_coverage/data/ERR3241902/HG01455.final.cram |
| **HG01456** | 1000G_2504_high_coverage/data/ERR3241903/HG01456.final.cram |
| **HG01461** | 1000G_2504_high_coverage/data/ERR3241904/HG01461.final.cram |
| **HG01462** | 1000G_2504_high_coverage/data/ERR3241905/HG01462.final.cram |
| **HG01464** | 1000G_2504_high_coverage/data/ERR3241906/HG01464.final.cram |
| **HG01465** | 1000G_2504_high_coverage/data/ERR3241907/HG01465.final.cram |
| **HG01488** | 1000G_2504_high_coverage/data/ERR3241908/HG01488.final.cram |
| **HG01489** | 1000G_2504_high_coverage/data/ERR3241909/HG01489.final.cram |
| **HG01491** | 1000G_2504_high_coverage/data/ERR3241910/HG01491.final.cram |
| **HG01492** | 1000G_2504_high_coverage/data/ERR3241911/HG01492.final.cram |
| **HG01494** | 1000G_2504_high_coverage/data/ERR3241912/HG01494.final.cram |
| **HG01495** | 1000G_2504_high_coverage/data/ERR3241913/HG01495.final.cram |
| **HG01497** | 1000G_2504_high_coverage/data/ERR3241914/HG01497.final.cram |
| **HG01498** | 1000G_2504_high_coverage/data/ERR3241915/HG01498.final.cram |
| **HG01500** | 1000G_2504_high_coverage/data/ERR3241916/HG01500.final.cram |
| **HG01501** | 1000G_2504_high_coverage/data/ERR3241917/HG01501.final.cram |
| **HG01503** | 1000G_2504_high_coverage/data/ERR3241918/HG01503.final.cram |
| **HG01504** | 1000G_2504_high_coverage/data/ERR3241919/HG01504.final.cram |
| **HG01506** | 1000G_2504_high_coverage/data/ERR3241920/HG01506.final.cram |
| **HG01507** | 1000G_2504_high_coverage/data/ERR3241921/HG01507.final.cram |
| **HG01509** | 1000G_2504_high_coverage/data/ERR3241922/HG01509.final.cram |
| **HG01510** | 1000G_2504_high_coverage/data/ERR3241923/HG01510.final.cram |
| **HG01512** | 1000G_2504_high_coverage/data/ERR3241924/HG01512.final.cram |
| **HG01513** | 1000G_2504_high_coverage/data/ERR3241925/HG01513.final.cram |
| **HG01515** | 1000G_2504_high_coverage/data/ERR3241926/HG01515.final.cram |
| **HG01516** | 1000G_2504_high_coverage/data/ERR3241927/HG01516.final.cram |
| **HG01518** | 1000G_2504_high_coverage/data/ERR3241928/HG01518.final.cram |
| **HG01519** | 1000G_2504_high_coverage/data/ERR3241929/HG01519.final.cram |
| **HG01521** | 1000G_2504_high_coverage/data/ERR3241930/HG01521.final.cram |
| **HG01522** | 1000G_2504_high_coverage/data/ERR3241931/HG01522.final.cram |
| **HG01524** | 1000G_2504_high_coverage/data/ERR3241932/HG01524.final.cram |
| **HG01525** | 1000G_2504_high_coverage/data/ERR3241933/HG01525.final.cram |
| **HG01527** | 1000G_2504_high_coverage/data/ERR3241934/HG01527.final.cram |
| **HG01528** | 1000G_2504_high_coverage/data/ERR3241935/HG01528.final.cram |
| **HG01530** | 1000G_2504_high_coverage/data/ERR3241936/HG01530.final.cram |
| **HG01531** | 1000G_2504_high_coverage/data/ERR3241937/HG01531.final.cram |
| **HG01536** | 1000G_2504_high_coverage/data/ERR3241938/HG01536.final.cram |
| **HG01537** | 1000G_2504_high_coverage/data/ERR3241939/HG01537.final.cram |
| **HG01550** | 1000G_2504_high_coverage/data/ERR3241940/HG01550.final.cram |
| **HG01551** | 1000G_2504_high_coverage/data/ERR3241941/HG01551.final.cram |
| **HG01602** | 1000G_2504_high_coverage/data/ERR3241942/HG01602.final.cram |
| **HG01603** | 1000G_2504_high_coverage/data/ERR3241943/HG01603.final.cram |
| **HG01605** | 1000G_2504_high_coverage/data/ERR3241944/HG01605.final.cram |
| **HG01606** | 1000G_2504_high_coverage/data/ERR3241945/HG01606.final.cram |
| **HG01607** | 1000G_2504_high_coverage/data/ERR3241946/HG01607.final.cram |
| **HG01608** | 1000G_2504_high_coverage/data/ERR3241947/HG01608.final.cram |
| **HG01610** | 1000G_2504_high_coverage/data/ERR3241948/HG01610.final.cram |
| **HG01612** | 1000G_2504_high_coverage/data/ERR3241949/HG01612.final.cram |
| **HG01613** | 1000G_2504_high_coverage/data/ERR3241950/HG01613.final.cram |
| **HG01615** | 1000G_2504_high_coverage/data/ERR3241951/HG01615.final.cram |
| **HG01617** | 1000G_2504_high_coverage/data/ERR3241952/HG01617.final.cram |
| **HG01618** | 1000G_2504_high_coverage/data/ERR3241953/HG01618.final.cram |
| **HG01619** | 1000G_2504_high_coverage/data/ERR3241954/HG01619.final.cram |
| **HG01620** | 1000G_2504_high_coverage/data/ERR3241955/HG01620.final.cram |
| **HG01623** | 1000G_2504_high_coverage/data/ERR3241956/HG01623.final.cram |

| | |
|---|---|
| **HG01624** | 1000G_2504_high_coverage/data/ERR3241957/HG01624.final.cram |
| **HG01625** | 1000G_2504_high_coverage/data/ERR3241958/HG01625.final.cram |
| **HG01626** | 1000G_2504_high_coverage/data/ERR3241959/HG01626.final.cram |
| **HG01628** | 1000G_2504_high_coverage/data/ERR3241960/HG01628.final.cram |
| **HG01630** | 1000G_2504_high_coverage/data/ERR3241961/HG01630.final.cram |
| **HG01631** | 1000G_2504_high_coverage/data/ERR3241962/HG01631.final.cram |
| **HG01632** | 1000G_2504_high_coverage/data/ERR3241963/HG01632.final.cram |
| **HG01668** | 1000G_2504_high_coverage/data/ERR3241964/HG01668.final.cram |
| **HG01669** | 1000G_2504_high_coverage/data/ERR3241965/HG01669.final.cram |
| **HG01670** | 1000G_2504_high_coverage/data/ERR3241966/HG01670.final.cram |
| **HG01672** | 1000G_2504_high_coverage/data/ERR3241967/HG01672.final.cram |
| **HG01673** | 1000G_2504_high_coverage/data/ERR3241968/HG01673.final.cram |
| **HG01675** | 1000G_2504_high_coverage/data/ERR3241969/HG01675.final.cram |
| **HG01676** | 1000G_2504_high_coverage/data/ERR3241970/HG01676.final.cram |
| **HG01678** | 1000G_2504_high_coverage/data/ERR3241971/HG01678.final.cram |
| **HG01679** | 1000G_2504_high_coverage/data/ERR3241972/HG01679.final.cram |
| **HG01680** | 1000G_2504_high_coverage/data/ERR3241973/HG01680.final.cram |
| **HG01682** | 1000G_2504_high_coverage/data/ERR3241974/HG01682.final.cram |
| **HG01684** | 1000G_2504_high_coverage/data/ERR3241975/HG01684.final.cram |
| **HG01685** | 1000G_2504_high_coverage/data/ERR3241976/HG01685.final.cram |
| **HG01686** | 1000G_2504_high_coverage/data/ERR3241977/HG01686.final.cram |
| **HG01700** | 1000G_2504_high_coverage/data/ERR3241978/HG01700.final.cram |
| **HG01702** | 1000G_2504_high_coverage/data/ERR3241979/HG01702.final.cram |
| **HG01704** | 1000G_2504_high_coverage/data/ERR3241980/HG01704.final.cram |
| **HG01705** | 1000G_2504_high_coverage/data/ERR3241981/HG01705.final.cram |
| **HG01707** | 1000G_2504_high_coverage/data/ERR3241982/HG01707.final.cram |
| **HG01708** | 1000G_2504_high_coverage/data/ERR3241983/HG01708.final.cram |
| **HG01565** | 1000G_2504_high_coverage/data/ERR3241984/HG01565.final.cram |
| **HG01566** | 1000G_2504_high_coverage/data/ERR3241985/HG01566.final.cram |
| **HG01571** | 1000G_2504_high_coverage/data/ERR3241986/HG01571.final.cram |
| **HG01572** | 1000G_2504_high_coverage/data/ERR3241987/HG01572.final.cram |
| **HG01577** | 1000G_2504_high_coverage/data/ERR3241988/HG01577.final.cram |
| **HG01578** | 1000G_2504_high_coverage/data/ERR3241989/HG01578.final.cram |
| **HG01917** | 1000G_2504_high_coverage/data/ERR3241990/HG01917.final.cram |
| **HG01918** | 1000G_2504_high_coverage/data/ERR3241991/HG01918.final.cram |
| **HG01892** | 1000G_2504_high_coverage/data/ERR3241992/HG01892.final.cram |
| **HG01893** | 1000G_2504_high_coverage/data/ERR3241993/HG01893.final.cram |
| **HG01920** | 1000G_2504_high_coverage/data/ERR3241994/HG01920.final.cram |
| **HG01921** | 1000G_2504_high_coverage/data/ERR3241995/HG01921.final.cram |
| **HG01923** | 1000G_2504_high_coverage/data/ERR3241996/HG01923.final.cram |
| **HG01924** | 1000G_2504_high_coverage/data/ERR3241997/HG01924.final.cram |
| **HG01926** | 1000G_2504_high_coverage/data/ERR3241998/HG01926.final.cram |
| **HG01927** | 1000G_2504_high_coverage/data/ERR3241999/HG01927.final.cram |
| **HG01932** | 1000G_2504_high_coverage/data/ERR3242000/HG01932.final.cram |
| **HG01933** | 1000G_2504_high_coverage/data/ERR3242001/HG01933.final.cram |
| **HG01935** | 1000G_2504_high_coverage/data/ERR3242002/HG01935.final.cram |
| **HG01936** | 1000G_2504_high_coverage/data/ERR3242003/HG01936.final.cram |
| **HG01938** | 1000G_2504_high_coverage/data/ERR3242004/HG01938.final.cram |
| **HG01939** | 1000G_2504_high_coverage/data/ERR3242005/HG01939.final.cram |
| **HG01941** | 1000G_2504_high_coverage/data/ERR3242006/HG01941.final.cram |
| **HG01942** | 1000G_2504_high_coverage/data/ERR3242007/HG01942.final.cram |
| **HG01944** | 1000G_2504_high_coverage/data/ERR3242008/HG01944.final.cram |
| **HG01945** | 1000G_2504_high_coverage/data/ERR3242009/HG01945.final.cram |
| **HG01947** | 1000G_2504_high_coverage/data/ERR3242010/HG01947.final.cram |
| **HG01948** | 1000G_2504_high_coverage/data/ERR3242011/HG01948.final.cram |
| **HG01950** | 1000G_2504_high_coverage/data/ERR3242012/HG01950.final.cram |
| **HG01951** | 1000G_2504_high_coverage/data/ERR3242013/HG01951.final.cram |
| **HG01953** | 1000G_2504_high_coverage/data/ERR3242014/HG01953.final.cram |
| **HG01954** | 1000G_2504_high_coverage/data/ERR3242015/HG01954.final.cram |
| **HG01967** | 1000G_2504_high_coverage/data/ERR3242016/HG01967.final.cram |
| **HG01968** | 1000G_2504_high_coverage/data/ERR3242017/HG01968.final.cram |
| **HG01970** | 1000G_2504_high_coverage/data/ERR3242018/HG01970.final.cram |
| **HG01971** | 1000G_2504_high_coverage/data/ERR3242019/HG01971.final.cram |
| **HG01974** | 1000G_2504_high_coverage/data/ERR3242020/HG01974.final.cram |
| **HG01973** | 1000G_2504_high_coverage/data/ERR3242021/HG01973.final.cram |

| | |
|---|---|
| **HG01977** | 1000G_2504_high_coverage/data/ERR3242022/HG01977.final.cram |
| **HG01976** | 1000G_2504_high_coverage/data/ERR3242023/HG01976.final.cram |
| **HG01979** | 1000G_2504_high_coverage/data/ERR3242024/HG01979.final.cram |
| **HG01980** | 1000G_2504_high_coverage/data/ERR3242025/HG01980.final.cram |
| **HG01991** | 1000G_2504_high_coverage/data/ERR3242026/HG01991.final.cram |
| **HG01992** | 1000G_2504_high_coverage/data/ERR3242027/HG01992.final.cram |
| **HG02008** | 1000G_2504_high_coverage/data/ERR3242028/HG02008.final.cram |
| **HG01997** | 1000G_2504_high_coverage/data/ERR3242029/HG01997.final.cram |
| **HG02002** | 1000G_2504_high_coverage/data/ERR3242030/HG02002.final.cram |
| **HG02003** | 1000G_2504_high_coverage/data/ERR3242031/HG02003.final.cram |
| **HG02090** | 1000G_2504_high_coverage/data/ERR3242032/HG02090.final.cram |
| **HG02089** | 1000G_2504_high_coverage/data/ERR3242033/HG02089.final.cram |
| **HG02104** | 1000G_2504_high_coverage/data/ERR3242034/HG02104.final.cram |
| **HG02105** | 1000G_2504_high_coverage/data/ERR3242035/HG02105.final.cram |
| **HG02017** | 1000G_2504_high_coverage/data/ERR3242036/HG02017.final.cram |
| **HG02016** | 1000G_2504_high_coverage/data/ERR3242037/HG02016.final.cram |
| **HG02020** | 1000G_2504_high_coverage/data/ERR3242038/HG02020.final.cram |
| **HG02019** | 1000G_2504_high_coverage/data/ERR3242039/HG02019.final.cram |
| **HG02023** | 1000G_2504_high_coverage/data/ERR3242040/HG02023.final.cram |
| **HG02026** | 1000G_2504_high_coverage/data/ERR3242041/HG02026.final.cram |
| **HG02025** | 1000G_2504_high_coverage/data/ERR3242042/HG02025.final.cram |
| **HG02029** | 1000G_2504_high_coverage/data/ERR3242043/HG02029.final.cram |
| **HG02028** | 1000G_2504_high_coverage/data/ERR3242044/HG02028.final.cram |
| **HG02058** | 1000G_2504_high_coverage/data/ERR3242045/HG02058.final.cram |
| **HG02057** | 1000G_2504_high_coverage/data/ERR3242046/HG02057.final.cram |
| **HG02070** | 1000G_2504_high_coverage/data/ERR3242047/HG02070.final.cram |
| **HG02069** | 1000G_2504_high_coverage/data/ERR3242048/HG02069.final.cram |
| **HG02073** | 1000G_2504_high_coverage/data/ERR3242049/HG02073.final.cram |
| **HG02072** | 1000G_2504_high_coverage/data/ERR3242050/HG02072.final.cram |
| **HG02076** | 1000G_2504_high_coverage/data/ERR3242051/HG02076.final.cram |
| **HG02075** | 1000G_2504_high_coverage/data/ERR3242052/HG02075.final.cram |
| **HG02079** | 1000G_2504_high_coverage/data/ERR3242053/HG02079.final.cram |
| **HG02078** | 1000G_2504_high_coverage/data/ERR3242054/HG02078.final.cram |
| **HG02082** | 1000G_2504_high_coverage/data/ERR3242055/HG02082.final.cram |
| **HG02081** | 1000G_2504_high_coverage/data/ERR3242056/HG02081.final.cram |
| **HG02085** | 1000G_2504_high_coverage/data/ERR3242057/HG02085.final.cram |
| **HG02084** | 1000G_2504_high_coverage/data/ERR3242058/HG02084.final.cram |
| **HG02134** | 1000G_2504_high_coverage/data/ERR3242059/HG02134.final.cram |
| **HG02133** | 1000G_2504_high_coverage/data/ERR3242060/HG02133.final.cram |
| **HG01841** | 1000G_2504_high_coverage/data/ERR3242061/HG01841.final.cram |
| **HG01595** | 1000G_2504_high_coverage/data/ERR3242062/HG01595.final.cram |
| **HG01596** | 1000G_2504_high_coverage/data/ERR3242063/HG01596.final.cram |
| **HG01597** | 1000G_2504_high_coverage/data/ERR3242064/HG01597.final.cram |
| **HG01598** | 1000G_2504_high_coverage/data/ERR3242065/HG01598.final.cram |
| **HG01599** | 1000G_2504_high_coverage/data/ERR3242066/HG01599.final.cram |
| **HG01600** | 1000G_2504_high_coverage/data/ERR3242067/HG01600.final.cram |
| **HG01840** | 1000G_2504_high_coverage/data/ERR3242068/HG01840.final.cram |
| **HG01842** | 1000G_2504_high_coverage/data/ERR3242069/HG01842.final.cram |
| **HG01843** | 1000G_2504_high_coverage/data/ERR3242070/HG01843.final.cram |
| **HG01844** | 1000G_2504_high_coverage/data/ERR3242071/HG01844.final.cram |
| **HG01845** | 1000G_2504_high_coverage/data/ERR3242072/HG01845.final.cram |
| **HG01846** | 1000G_2504_high_coverage/data/ERR3242073/HG01846.final.cram |
| **HG01847** | 1000G_2504_high_coverage/data/ERR3242074/HG01847.final.cram |
| **HG01848** | 1000G_2504_high_coverage/data/ERR3242075/HG01848.final.cram |
| **HG01849** | 1000G_2504_high_coverage/data/ERR3242076/HG01849.final.cram |
| **HG01850** | 1000G_2504_high_coverage/data/ERR3242077/HG01850.final.cram |
| **HG01851** | 1000G_2504_high_coverage/data/ERR3242078/HG01851.final.cram |
| **HG01852** | 1000G_2504_high_coverage/data/ERR3242079/HG01852.final.cram |
| **HG01853** | 1000G_2504_high_coverage/data/ERR3242080/HG01853.final.cram |
| **HG01855** | 1000G_2504_high_coverage/data/ERR3242081/HG01855.final.cram |
| **HG01857** | 1000G_2504_high_coverage/data/ERR3242082/HG01857.final.cram |
| **HG01858** | 1000G_2504_high_coverage/data/ERR3242083/HG01858.final.cram |
| **HG01859** | 1000G_2504_high_coverage/data/ERR3242084/HG01859.final.cram |
| **HG01860** | 1000G_2504_high_coverage/data/ERR3242085/HG01860.final.cram |
| **HG01861** | 1000G_2504_high_coverage/data/ERR3242086/HG01861.final.cram |

| | |
|---|---|
| **HG01862** | 1000G_2504_high_coverage/data/ERR3242087/HG01862.final.cram |
| **HG01863** | 1000G_2504_high_coverage/data/ERR3242088/HG01863.final.cram |
| **HG01864** | 1000G_2504_high_coverage/data/ERR3242089/HG01864.final.cram |
| **HG01865** | 1000G_2504_high_coverage/data/ERR3242090/HG01865.final.cram |
| **HG01866** | 1000G_2504_high_coverage/data/ERR3242091/HG01866.final.cram |
| **HG01867** | 1000G_2504_high_coverage/data/ERR3242092/HG01867.final.cram |
| **HG01868** | 1000G_2504_high_coverage/data/ERR3242093/HG01868.final.cram |
| **HG01869** | 1000G_2504_high_coverage/data/ERR3242094/HG01869.final.cram |
| **HG01870** | 1000G_2504_high_coverage/data/ERR3242095/HG01870.final.cram |
| **HG01871** | 1000G_2504_high_coverage/data/ERR3242096/HG01871.final.cram |
| **HG01872** | 1000G_2504_high_coverage/data/ERR3242097/HG01872.final.cram |
| **HG01873** | 1000G_2504_high_coverage/data/ERR3242098/HG01873.final.cram |
| **HG01874** | 1000G_2504_high_coverage/data/ERR3242099/HG01874.final.cram |
| **HG01878** | 1000G_2504_high_coverage/data/ERR3242100/HG01878.final.cram |
| **HG02048** | 1000G_2504_high_coverage/data/ERR3242101/HG02048.final.cram |
| **HG02049** | 1000G_2504_high_coverage/data/ERR3242102/HG02049.final.cram |
| **HG02086** | 1000G_2504_high_coverage/data/ERR3242103/HG02086.final.cram |
| **HG02087** | 1000G_2504_high_coverage/data/ERR3242104/HG02087.final.cram |
| **HG02088** | 1000G_2504_high_coverage/data/ERR3242105/HG02088.final.cram |
| **HG02138** | 1000G_2504_high_coverage/data/ERR3242106/HG02138.final.cram |
| **HG02139** | 1000G_2504_high_coverage/data/ERR3242107/HG02139.final.cram |
| **HG02140** | 1000G_2504_high_coverage/data/ERR3242108/HG02140.final.cram |
| **HG02142** | 1000G_2504_high_coverage/data/ERR3242109/HG02142.final.cram |
| **HG01694** | 1000G_2504_high_coverage/data/ERR3242110/HG01694.final.cram |
| **HG01695** | 1000G_2504_high_coverage/data/ERR3242111/HG01695.final.cram |
| **HG01709** | 1000G_2504_high_coverage/data/ERR3242112/HG01709.final.cram |
| **HG01710** | 1000G_2504_high_coverage/data/ERR3242113/HG01710.final.cram |
| **HG01747** | 1000G_2504_high_coverage/data/ERR3242114/HG01747.final.cram |
| **HG01746** | 1000G_2504_high_coverage/data/ERR3242115/HG01746.final.cram |
| **HG01756** | 1000G_2504_high_coverage/data/ERR3242116/HG01756.final.cram |
| **HG01757** | 1000G_2504_high_coverage/data/ERR3242117/HG01757.final.cram |
| **HG01761** | 1000G_2504_high_coverage/data/ERR3242118/HG01761.final.cram |
| **HG01762** | 1000G_2504_high_coverage/data/ERR3242119/HG01762.final.cram |
| **HG01771** | 1000G_2504_high_coverage/data/ERR3242120/HG01771.final.cram |
| **HG01770** | 1000G_2504_high_coverage/data/ERR3242121/HG01770.final.cram |
| **HG01775** | 1000G_2504_high_coverage/data/ERR3242122/HG01775.final.cram |
| **HG01773** | 1000G_2504_high_coverage/data/ERR3242123/HG01773.final.cram |
| **HG01777** | 1000G_2504_high_coverage/data/ERR3242124/HG01777.final.cram |
| **HG01776** | 1000G_2504_high_coverage/data/ERR3242125/HG01776.final.cram |
| **HG01781** | 1000G_2504_high_coverage/data/ERR3242126/HG01781.final.cram |
| **HG01779** | 1000G_2504_high_coverage/data/ERR3242127/HG01779.final.cram |
| **HG00759** | 1000G_2504_high_coverage/data/ERR3242128/HG00759.final.cram |
| **HG00766** | 1000G_2504_high_coverage/data/ERR3242129/HG00766.final.cram |
| **HG00844** | 1000G_2504_high_coverage/data/ERR3242130/HG00844.final.cram |
| **HG00851** | 1000G_2504_high_coverage/data/ERR3242131/HG00851.final.cram |
| **HG00864** | 1000G_2504_high_coverage/data/ERR3242132/HG00864.final.cram |
| **HG00867** | 1000G_2504_high_coverage/data/ERR3242133/HG00867.final.cram |
| **HG00879** | 1000G_2504_high_coverage/data/ERR3242134/HG00879.final.cram |
| **HG00881** | 1000G_2504_high_coverage/data/ERR3242135/HG00881.final.cram |
| **HG00956** | 1000G_2504_high_coverage/data/ERR3242136/HG00956.final.cram |
| **HG00978** | 1000G_2504_high_coverage/data/ERR3242137/HG00978.final.cram |
| **HG00982** | 1000G_2504_high_coverage/data/ERR3242138/HG00982.final.cram |
| **HG01028** | 1000G_2504_high_coverage/data/ERR3242139/HG01028.final.cram |
| **HG01029** | 1000G_2504_high_coverage/data/ERR3242140/HG01029.final.cram |
| **HG01031** | 1000G_2504_high_coverage/data/ERR3242141/HG01031.final.cram |
| **HG01046** | 1000G_2504_high_coverage/data/ERR3242142/HG01046.final.cram |
| **HG01794** | 1000G_2504_high_coverage/data/ERR3242143/HG01794.final.cram |
| **HG01795** | 1000G_2504_high_coverage/data/ERR3242144/HG01795.final.cram |
| **HG01796** | 1000G_2504_high_coverage/data/ERR3242145/HG01796.final.cram |
| **HG01797** | 1000G_2504_high_coverage/data/ERR3242146/HG01797.final.cram |
| **HG01798** | 1000G_2504_high_coverage/data/ERR3242147/HG01798.final.cram |
| **HG01799** | 1000G_2504_high_coverage/data/ERR3242148/HG01799.final.cram |
| **HG01800** | 1000G_2504_high_coverage/data/ERR3242149/HG01800.final.cram |
| **HG01801** | 1000G_2504_high_coverage/data/ERR3242150/HG01801.final.cram |
| **HG01802** | 1000G_2504_high_coverage/data/ERR3242151/HG01802.final.cram |

| | |
|---|---|
| **HG01804** | 1000G_2504_high_coverage/data/ERR3242152/HG01804.final.cram |
| **HG01805** | 1000G_2504_high_coverage/data/ERR3242153/HG01805.final.cram |
| **HG01806** | 1000G_2504_high_coverage/data/ERR3242154/HG01806.final.cram |
| **HG01807** | 1000G_2504_high_coverage/data/ERR3242155/HG01807.final.cram |
| **HG01808** | 1000G_2504_high_coverage/data/ERR3242156/HG01808.final.cram |
| **HG01809** | 1000G_2504_high_coverage/data/ERR3242157/HG01809.final.cram |
| **HG01810** | 1000G_2504_high_coverage/data/ERR3242158/HG01810.final.cram |
| **HG01811** | 1000G_2504_high_coverage/data/ERR3242159/HG01811.final.cram |
| **HG01812** | 1000G_2504_high_coverage/data/ERR3242160/HG01812.final.cram |
| **HG01813** | 1000G_2504_high_coverage/data/ERR3242161/HG01813.final.cram |
| **HG02151** | 1000G_2504_high_coverage/data/ERR3242162/HG02151.final.cram |
| **HG01815** | 1000G_2504_high_coverage/data/ERR3242163/HG01815.final.cram |
| **HG01816** | 1000G_2504_high_coverage/data/ERR3242164/HG01816.final.cram |
| **HG01817** | 1000G_2504_high_coverage/data/ERR3242165/HG01817.final.cram |
| **HG02152** | 1000G_2504_high_coverage/data/ERR3242166/HG02152.final.cram |
| **HG02153** | 1000G_2504_high_coverage/data/ERR3242167/HG02153.final.cram |
| **HG02154** | 1000G_2504_high_coverage/data/ERR3242168/HG02154.final.cram |
| **HG02155** | 1000G_2504_high_coverage/data/ERR3242169/HG02155.final.cram |
| **HG02156** | 1000G_2504_high_coverage/data/ERR3242170/HG02156.final.cram |
| **HG02164** | 1000G_2504_high_coverage/data/ERR3242171/HG02164.final.cram |
| **HG02165** | 1000G_2504_high_coverage/data/ERR3242172/HG02165.final.cram |
| **HG02166** | 1000G_2504_high_coverage/data/ERR3242173/HG02166.final.cram |
| **HG02178** | 1000G_2504_high_coverage/data/ERR3242174/HG02178.final.cram |
| **HG02179** | 1000G_2504_high_coverage/data/ERR3242175/HG02179.final.cram |
| **HG02180** | 1000G_2504_high_coverage/data/ERR3242176/HG02180.final.cram |
| **HG02181** | 1000G_2504_high_coverage/data/ERR3242177/HG02181.final.cram |
| **HG02182** | 1000G_2504_high_coverage/data/ERR3242178/HG02182.final.cram |
| **HG02184** | 1000G_2504_high_coverage/data/ERR3242179/HG02184.final.cram |
| **HG02185** | 1000G_2504_high_coverage/data/ERR3242180/HG02185.final.cram |
| **HG02186** | 1000G_2504_high_coverage/data/ERR3242181/HG02186.final.cram |
| **HG02187** | 1000G_2504_high_coverage/data/ERR3242182/HG02187.final.cram |
| **HG02188** | 1000G_2504_high_coverage/data/ERR3242183/HG02188.final.cram |
| **HG02190** | 1000G_2504_high_coverage/data/ERR3242184/HG02190.final.cram |
| **HG01789** | 1000G_2504_high_coverage/data/ERR3242185/HG01789.final.cram |
| **HG01790** | 1000G_2504_high_coverage/data/ERR3242186/HG01790.final.cram |
| **HG01791** | 1000G_2504_high_coverage/data/ERR3242187/HG01791.final.cram |
| **HG02215** | 1000G_2504_high_coverage/data/ERR3242188/HG02215.final.cram |
| **HG01882** | 1000G_2504_high_coverage/data/ERR3242189/HG01882.final.cram |
| **HG01883** | 1000G_2504_high_coverage/data/ERR3242190/HG01883.final.cram |
| **HG01885** | 1000G_2504_high_coverage/data/ERR3242191/HG01885.final.cram |
| **HG01956** | 1000G_2504_high_coverage/data/ERR3242192/HG01956.final.cram |
| **HG02014** | 1000G_2504_high_coverage/data/ERR3242193/HG02014.final.cram |
| **HG01886** | 1000G_2504_high_coverage/data/ERR3242194/HG01886.final.cram |
| **HG01890** | 1000G_2504_high_coverage/data/ERR3242195/HG01890.final.cram |
| **HG01889** | 1000G_2504_high_coverage/data/ERR3242196/HG01889.final.cram |
| **HG02009** | 1000G_2504_high_coverage/data/ERR3242197/HG02009.final.cram |
| **HG02010** | 1000G_2504_high_coverage/data/ERR3242198/HG02010.final.cram |
| **HG02285** | 1000G_2504_high_coverage/data/ERR3242199/HG02285.final.cram |
| **HG02286** | 1000G_2504_high_coverage/data/ERR3242200/HG02286.final.cram |
| **HG02301** | 1000G_2504_high_coverage/data/ERR3242201/HG02301.final.cram |
| **HG01783** | 1000G_2504_high_coverage/data/ERR3242202/HG01783.final.cram |
| **HG01784** | 1000G_2504_high_coverage/data/ERR3242203/HG01784.final.cram |
| **HG02221** | 1000G_2504_high_coverage/data/ERR3242204/HG02221.final.cram |
| **HG02220** | 1000G_2504_high_coverage/data/ERR3242205/HG02220.final.cram |
| **HG02236** | 1000G_2504_high_coverage/data/ERR3242206/HG02236.final.cram |
| **HG02235** | 1000G_2504_high_coverage/data/ERR3242207/HG02235.final.cram |
| **HG02113** | 1000G_2504_high_coverage/data/ERR3242208/HG02113.final.cram |
| **HG02141** | 1000G_2504_high_coverage/data/ERR3242209/HG02141.final.cram |
| **HG02122** | 1000G_2504_high_coverage/data/ERR3242210/HG02122.final.cram |
| **HG02121** | 1000G_2504_high_coverage/data/ERR3242211/HG02121.final.cram |
| **HG02128** | 1000G_2504_high_coverage/data/ERR3242212/HG02128.final.cram |
| **HG02127** | 1000G_2504_high_coverage/data/ERR3242213/HG02127.final.cram |
| **HG02373** | 1000G_2504_high_coverage/data/ERR3242214/HG02373.final.cram |
| **HG02394** | 1000G_2504_high_coverage/data/ERR3242215/HG02394.final.cram |
| **HG02396** | 1000G_2504_high_coverage/data/ERR3242216/HG02396.final.cram |

| | |
|---|---|
| **HG02397** | 1000G_2504_high_coverage/data/ERR3242217/HG02397.final.cram |
| **HG01914** | 1000G_2504_high_coverage/data/ERR3242218/HG01914.final.cram |
| **HG01915** | 1000G_2504_high_coverage/data/ERR3242219/HG01915.final.cram |
| **HG01986** | 1000G_2504_high_coverage/data/ERR3242220/HG01986.final.cram |
| **HG01958** | 1000G_2504_high_coverage/data/ERR3242221/HG01958.final.cram |
| **HG01990** | 1000G_2504_high_coverage/data/ERR3242222/HG01990.final.cram |
| **HG02012** | 1000G_2504_high_coverage/data/ERR3242223/HG02012.final.cram |
| **HG02053** | 1000G_2504_high_coverage/data/ERR3242224/HG02053.final.cram |
| **HG02054** | 1000G_2504_high_coverage/data/ERR3242225/HG02054.final.cram |
| **HG02255** | 1000G_2504_high_coverage/data/ERR3242226/HG02255.final.cram |
| **HG02256** | 1000G_2504_high_coverage/data/ERR3242227/HG02256.final.cram |
| **HG02314** | 1000G_2504_high_coverage/data/ERR3242228/HG02314.final.cram |
| **HG02315** | 1000G_2504_high_coverage/data/ERR3242229/HG02315.final.cram |
| **HG02281** | 1000G_2504_high_coverage/data/ERR3242230/HG02281.final.cram |
| **HG02282** | 1000G_2504_high_coverage/data/ERR3242231/HG02282.final.cram |
| **HG02317** | 1000G_2504_high_coverage/data/ERR3242232/HG02317.final.cram |
| **HG02318** | 1000G_2504_high_coverage/data/ERR3242233/HG02318.final.cram |
| **HG01912** | 1000G_2504_high_coverage/data/ERR3242234/HG01912.final.cram |
| **HG01894** | 1000G_2504_high_coverage/data/ERR3242235/HG01894.final.cram |
| **HG01982** | 1000G_2504_high_coverage/data/ERR3242236/HG01982.final.cram |
| **HG02146** | 1000G_2504_high_coverage/data/ERR3242237/HG02146.final.cram |
| **HG02147** | 1000G_2504_high_coverage/data/ERR3242238/HG02147.final.cram |
| **HG02259** | 1000G_2504_high_coverage/data/ERR3242239/HG02259.final.cram |
| **HG02260** | 1000G_2504_high_coverage/data/ERR3242240/HG02260.final.cram |
| **HG02271** | 1000G_2504_high_coverage/data/ERR3242241/HG02271.final.cram |
| **HG02272** | 1000G_2504_high_coverage/data/ERR3242242/HG02272.final.cram |
| **HG02277** | 1000G_2504_high_coverage/data/ERR3242243/HG02277.final.cram |
| **HG02278** | 1000G_2504_high_coverage/data/ERR3242244/HG02278.final.cram |
| **HG02291** | 1000G_2504_high_coverage/data/ERR3242245/HG02291.final.cram |
| **HG02292** | 1000G_2504_high_coverage/data/ERR3242246/HG02292.final.cram |
| **HG02299** | 1000G_2504_high_coverage/data/ERR3242247/HG02299.final.cram |
| **HG02298** | 1000G_2504_high_coverage/data/ERR3242248/HG02298.final.cram |
| **HG01699** | 1000G_2504_high_coverage/data/ERR3242249/HG01699.final.cram |
| **HG01697** | 1000G_2504_high_coverage/data/ERR3242250/HG01697.final.cram |
| **HG02224** | 1000G_2504_high_coverage/data/ERR3242251/HG02224.final.cram |
| **HG02223** | 1000G_2504_high_coverage/data/ERR3242252/HG02223.final.cram |
| **HG02231** | 1000G_2504_high_coverage/data/ERR3242253/HG02231.final.cram |
| **HG02230** | 1000G_2504_high_coverage/data/ERR3242254/HG02230.final.cram |
| **HG02238** | 1000G_2504_high_coverage/data/ERR3242255/HG02238.final.cram |
| **HG02239** | 1000G_2504_high_coverage/data/ERR3242256/HG02239.final.cram |
| **HG02032** | 1000G_2504_high_coverage/data/ERR3242257/HG02032.final.cram |
| **HG02031** | 1000G_2504_high_coverage/data/ERR3242258/HG02031.final.cram |
| **HG02050** | 1000G_2504_high_coverage/data/ERR3242259/HG02050.final.cram |
| **HG02250** | 1000G_2504_high_coverage/data/ERR3242260/HG02250.final.cram |
| **HG02351** | 1000G_2504_high_coverage/data/ERR3242261/HG02351.final.cram |
| **HG02353** | 1000G_2504_high_coverage/data/ERR3242262/HG02353.final.cram |
| **HG02355** | 1000G_2504_high_coverage/data/ERR3242263/HG02355.final.cram |
| **HG02356** | 1000G_2504_high_coverage/data/ERR3242264/HG02356.final.cram |
| **HG02360** | 1000G_2504_high_coverage/data/ERR3242265/HG02360.final.cram |
| **HG02364** | 1000G_2504_high_coverage/data/ERR3242266/HG02364.final.cram |
| **HG02367** | 1000G_2504_high_coverage/data/ERR3242267/HG02367.final.cram |
| **HG02371** | 1000G_2504_high_coverage/data/ERR3242268/HG02371.final.cram |
| **HG02374** | 1000G_2504_high_coverage/data/ERR3242269/HG02374.final.cram |
| **HG02375** | 1000G_2504_high_coverage/data/ERR3242270/HG02375.final.cram |
| **HG02379** | 1000G_2504_high_coverage/data/ERR3242271/HG02379.final.cram |
| **HG02380** | 1000G_2504_high_coverage/data/ERR3242272/HG02380.final.cram |
| **HG02382** | 1000G_2504_high_coverage/data/ERR3242273/HG02382.final.cram |
| **HG02383** | 1000G_2504_high_coverage/data/ERR3242274/HG02383.final.cram |
| **HG02384** | 1000G_2504_high_coverage/data/ERR3242275/HG02384.final.cram |
| **HG02385** | 1000G_2504_high_coverage/data/ERR3242276/HG02385.final.cram |
| **HG02386** | 1000G_2504_high_coverage/data/ERR3242277/HG02386.final.cram |
| **HG02389** | 1000G_2504_high_coverage/data/ERR3242278/HG02389.final.cram |
| **HG02390** | 1000G_2504_high_coverage/data/ERR3242279/HG02390.final.cram |
| **HG02391** | 1000G_2504_high_coverage/data/ERR3242280/HG02391.final.cram |
| **HG02392** | 1000G_2504_high_coverage/data/ERR3242281/HG02392.final.cram |

| | |
|---|---|
| **HG02395** | 1000G_2504_high_coverage/data/ERR3242282/HG02395.final.cram |
| **HG02398** | 1000G_2504_high_coverage/data/ERR3242283/HG02398.final.cram |
| **HG02399** | 1000G_2504_high_coverage/data/ERR3242284/HG02399.final.cram |
| **HG02401** | 1000G_2504_high_coverage/data/ERR3242285/HG02401.final.cram |
| **HG02402** | 1000G_2504_high_coverage/data/ERR3242286/HG02402.final.cram |
| **HG02406** | 1000G_2504_high_coverage/data/ERR3242287/HG02406.final.cram |
| **HG02407** | 1000G_2504_high_coverage/data/ERR3242288/HG02407.final.cram |
| **HG02409** | 1000G_2504_high_coverage/data/ERR3242289/HG02409.final.cram |
| **HG02233** | 1000G_2504_high_coverage/data/ERR3242290/HG02233.final.cram |
| **HG02232** | 1000G_2504_high_coverage/data/ERR3242291/HG02232.final.cram |
| **HG01879** | 1000G_2504_high_coverage/data/ERR3242292/HG01879.final.cram |
| **HG01880** | 1000G_2504_high_coverage/data/ERR3242293/HG01880.final.cram |
| **HG02013** | 1000G_2504_high_coverage/data/ERR3242294/HG02013.final.cram |
| **HG01896** | 1000G_2504_high_coverage/data/ERR3242295/HG01896.final.cram |
| **HG01985** | 1000G_2504_high_coverage/data/ERR3242296/HG01985.final.cram |
| **HG01988** | 1000G_2504_high_coverage/data/ERR3242297/HG01988.final.cram |
| **HG01989** | 1000G_2504_high_coverage/data/ERR3242298/HG01989.final.cram |
| **HG02051** | 1000G_2504_high_coverage/data/ERR3242299/HG02051.final.cram |
| **HG02052** | 1000G_2504_high_coverage/data/ERR3242300/HG02052.final.cram |
| **HG02095** | 1000G_2504_high_coverage/data/ERR3242301/HG02095.final.cram |
| **HG02332** | 1000G_2504_high_coverage/data/ERR3242302/HG02332.final.cram |
| **HG02107** | 1000G_2504_high_coverage/data/ERR3242303/HG02107.final.cram |
| **HG02108** | 1000G_2504_high_coverage/data/ERR3242304/HG02108.final.cram |
| **HG02111** | 1000G_2504_high_coverage/data/ERR3242305/HG02111.final.cram |
| **HG02143** | 1000G_2504_high_coverage/data/ERR3242306/HG02143.final.cram |
| **HG02144** | 1000G_2504_high_coverage/data/ERR3242307/HG02144.final.cram |
| **HG02307** | 1000G_2504_high_coverage/data/ERR3242308/HG02307.final.cram |
| **HG02308** | 1000G_2504_high_coverage/data/ERR3242309/HG02308.final.cram |
| **HG02283** | 1000G_2504_high_coverage/data/ERR3242310/HG02283.final.cram |
| **HG02309** | 1000G_2504_high_coverage/data/ERR3242311/HG02309.final.cram |
| **HG02284** | 1000G_2504_high_coverage/data/ERR3242312/HG02284.final.cram |
| **HG02322** | 1000G_2504_high_coverage/data/ERR3242313/HG02322.final.cram |
| **HG02323** | 1000G_2504_high_coverage/data/ERR3242314/HG02323.final.cram |
| **HG02325** | 1000G_2504_high_coverage/data/ERR3242315/HG02325.final.cram |
| **HG02337** | 1000G_2504_high_coverage/data/ERR3242316/HG02337.final.cram |
| **HG02334** | 1000G_2504_high_coverage/data/ERR3242317/HG02334.final.cram |
| **HG02330** | 1000G_2504_high_coverage/data/ERR3242318/HG02330.final.cram |
| **HG02339** | 1000G_2504_high_coverage/data/ERR3242319/HG02339.final.cram |
| **HG02343** | 1000G_2504_high_coverage/data/ERR3242320/HG02343.final.cram |
| **HG02419** | 1000G_2504_high_coverage/data/ERR3242321/HG02419.final.cram |
| **HG02420** | 1000G_2504_high_coverage/data/ERR3242322/HG02420.final.cram |
| **HG02427** | 1000G_2504_high_coverage/data/ERR3242323/HG02427.final.cram |
| **HG02429** | 1000G_2504_high_coverage/data/ERR3242324/HG02429.final.cram |
| **HG02433** | 1000G_2504_high_coverage/data/ERR3242325/HG02433.final.cram |
| **HG02442** | 1000G_2504_high_coverage/data/ERR3242326/HG02442.final.cram |
| **HG02445** | 1000G_2504_high_coverage/data/ERR3242327/HG02445.final.cram |
| **HG02489** | 1000G_2504_high_coverage/data/ERR3242328/HG02489.final.cram |
| **HG02449** | 1000G_2504_high_coverage/data/ERR3242329/HG02449.final.cram |
| **HG02450** | 1000G_2504_high_coverage/data/ERR3242330/HG02450.final.cram |
| **HG02470** | 1000G_2504_high_coverage/data/ERR3242331/HG02470.final.cram |
| **HG02471** | 1000G_2504_high_coverage/data/ERR3242332/HG02471.final.cram |
| **HG02479** | 1000G_2504_high_coverage/data/ERR3242333/HG02479.final.cram |
| **HG02484** | 1000G_2504_high_coverage/data/ERR3242334/HG02484.final.cram |
| **HG02485** | 1000G_2504_high_coverage/data/ERR3242335/HG02485.final.cram |
| **HG02511** | 1000G_2504_high_coverage/data/ERR3242336/HG02511.final.cram |
| **HG02496** | 1000G_2504_high_coverage/data/ERR3242337/HG02496.final.cram |
| **HG02497** | 1000G_2504_high_coverage/data/ERR3242338/HG02497.final.cram |
| **HG02508** | 1000G_2504_high_coverage/data/ERR3242339/HG02508.final.cram |
| **HG02537** | 1000G_2504_high_coverage/data/ERR3242340/HG02537.final.cram |
| **HG02035** | 1000G_2504_high_coverage/data/ERR3242341/HG02035.final.cram |
| **HG02040** | 1000G_2504_high_coverage/data/ERR3242342/HG02040.final.cram |
| **HG02047** | 1000G_2504_high_coverage/data/ERR3242343/HG02047.final.cram |
| **HG02061** | 1000G_2504_high_coverage/data/ERR3242344/HG02061.final.cram |
| **HG02060** | 1000G_2504_high_coverage/data/ERR3242345/HG02060.final.cram |
| **HG02064** | 1000G_2504_high_coverage/data/ERR3242346/HG02064.final.cram |

| | |
|---|---|
| **HG02067** | 1000G_2504_high_coverage/data/ERR3242347/HG02067.final.cram |
| **HG02116** | 1000G_2504_high_coverage/data/ERR3242348/HG02116.final.cram |
| **HG02131** | 1000G_2504_high_coverage/data/ERR3242349/HG02131.final.cram |
| **HG02130** | 1000G_2504_high_coverage/data/ERR3242350/HG02130.final.cram |
| **HG02137** | 1000G_2504_high_coverage/data/ERR3242351/HG02137.final.cram |
| **HG02136** | 1000G_2504_high_coverage/data/ERR3242352/HG02136.final.cram |
| **HG02512** | 1000G_2504_high_coverage/data/ERR3242353/HG02512.final.cram |
| **HG02513** | 1000G_2504_high_coverage/data/ERR3242354/HG02513.final.cram |
| **HG02521** | 1000G_2504_high_coverage/data/ERR3242355/HG02521.final.cram |
| **HG02522** | 1000G_2504_high_coverage/data/ERR3242356/HG02522.final.cram |
| **HG02461** | 1000G_2504_high_coverage/data/ERR3242357/HG02461.final.cram |
| **HG02462** | 1000G_2504_high_coverage/data/ERR3242358/HG02462.final.cram |
| **HG02464** | 1000G_2504_high_coverage/data/ERR3242359/HG02464.final.cram |
| **HG02465** | 1000G_2504_high_coverage/data/ERR3242360/HG02465.final.cram |
| **HG02561** | 1000G_2504_high_coverage/data/ERR3242361/HG02561.final.cram |
| **HG02562** | 1000G_2504_high_coverage/data/ERR3242362/HG02562.final.cram |
| **HG02570** | 1000G_2504_high_coverage/data/ERR3242363/HG02570.final.cram |
| **HG02571** | 1000G_2504_high_coverage/data/ERR3242364/HG02571.final.cram |
| **HG02573** | 1000G_2504_high_coverage/data/ERR3242365/HG02573.final.cram |
| **HG02574** | 1000G_2504_high_coverage/data/ERR3242366/HG02574.final.cram |
| **HG02582** | 1000G_2504_high_coverage/data/ERR3242367/HG02582.final.cram |
| **HG02583** | 1000G_2504_high_coverage/data/ERR3242368/HG02583.final.cram |
| **HG02585** | 1000G_2504_high_coverage/data/ERR3242369/HG02585.final.cram |
| **HG02586** | 1000G_2504_high_coverage/data/ERR3242370/HG02586.final.cram |
| **HG02588** | 1000G_2504_high_coverage/data/ERR3242371/HG02588.final.cram |
| **HG02589** | 1000G_2504_high_coverage/data/ERR3242372/HG02589.final.cram |
| **HG02594** | 1000G_2504_high_coverage/data/ERR3242373/HG02594.final.cram |
| **HG02595** | 1000G_2504_high_coverage/data/ERR3242374/HG02595.final.cram |
| **HG02613** | 1000G_2504_high_coverage/data/ERR3242375/HG02613.final.cram |
| **HG02614** | 1000G_2504_high_coverage/data/ERR3242376/HG02614.final.cram |
| **HG02620** | 1000G_2504_high_coverage/data/ERR3242377/HG02620.final.cram |
| **HG02621** | 1000G_2504_high_coverage/data/ERR3242378/HG02621.final.cram |
| **HG02623** | 1000G_2504_high_coverage/data/ERR3242379/HG02623.final.cram |
| **HG02624** | 1000G_2504_high_coverage/data/ERR3242380/HG02624.final.cram |
| **HG02628** | 1000G_2504_high_coverage/data/ERR3242381/HG02628.final.cram |
| **HG02629** | 1000G_2504_high_coverage/data/ERR3242382/HG02629.final.cram |
| **HG02642** | 1000G_2504_high_coverage/data/ERR3242383/HG02642.final.cram |
| **HG02643** | 1000G_2504_high_coverage/data/ERR3242384/HG02643.final.cram |
| **HG02645** | 1000G_2504_high_coverage/data/ERR3242385/HG02645.final.cram |
| **HG02646** | 1000G_2504_high_coverage/data/ERR3242386/HG02646.final.cram |
| **HG02666** | 1000G_2504_high_coverage/data/ERR3242387/HG02666.final.cram |
| **HG02667** | 1000G_2504_high_coverage/data/ERR3242388/HG02667.final.cram |
| **HG02675** | 1000G_2504_high_coverage/data/ERR3242389/HG02675.final.cram |
| **HG02676** | 1000G_2504_high_coverage/data/ERR3242390/HG02676.final.cram |
| **HG02678** | 1000G_2504_high_coverage/data/ERR3242391/HG02678.final.cram |
| **HG02679** | 1000G_2504_high_coverage/data/ERR3242392/HG02679.final.cram |
| **HG02702** | 1000G_2504_high_coverage/data/ERR3242393/HG02702.final.cram |
| **HG02703** | 1000G_2504_high_coverage/data/ERR3242394/HG02703.final.cram |
| **HG02721** | 1000G_2504_high_coverage/data/ERR3242395/HG02721.final.cram |
| **HG02722** | 1000G_2504_high_coverage/data/ERR3242396/HG02722.final.cram |
| **HG02490** | 1000G_2504_high_coverage/data/ERR3242397/HG02490.final.cram |
| **HG02491** | 1000G_2504_high_coverage/data/ERR3242398/HG02491.final.cram |
| **HG02600** | 1000G_2504_high_coverage/data/ERR3242399/HG02600.final.cram |
| **HG02601** | 1000G_2504_high_coverage/data/ERR3242400/HG02601.final.cram |
| **HG02603** | 1000G_2504_high_coverage/data/ERR3242401/HG02603.final.cram |
| **HG02604** | 1000G_2504_high_coverage/data/ERR3242402/HG02604.final.cram |
| **HG02654** | 1000G_2504_high_coverage/data/ERR3242403/HG02654.final.cram |
| **HG02655** | 1000G_2504_high_coverage/data/ERR3242404/HG02655.final.cram |
| **HG02657** | 1000G_2504_high_coverage/data/ERR3242405/HG02657.final.cram |
| **HG02658** | 1000G_2504_high_coverage/data/ERR3242406/HG02658.final.cram |
| **HG02660** | 1000G_2504_high_coverage/data/ERR3242407/HG02660.final.cram |
| **HG02661** | 1000G_2504_high_coverage/data/ERR3242408/HG02661.final.cram |
| **HG02684** | 1000G_2504_high_coverage/data/ERR3242409/HG02684.final.cram |
| **HG02685** | 1000G_2504_high_coverage/data/ERR3242410/HG02685.final.cram |
| **HG02687** | 1000G_2504_high_coverage/data/ERR3242411/HG02687.final.cram |

| | |
|---|---|
| **HG02688** | 1000G_2504_high_coverage/data/ERR3242412/HG02688.final.cram |
| **HG02696** | 1000G_2504_high_coverage/data/ERR3242413/HG02696.final.cram |
| **HG02697** | 1000G_2504_high_coverage/data/ERR3242414/HG02697.final.cram |
| **HG02733** | 1000G_2504_high_coverage/data/ERR3242415/HG02733.final.cram |
| **HG02734** | 1000G_2504_high_coverage/data/ERR3242416/HG02734.final.cram |
| **HG02610** | 1000G_2504_high_coverage/data/ERR3242417/HG02610.final.cram |
| **HG02611** | 1000G_2504_high_coverage/data/ERR3242418/HG02611.final.cram |
| **HG02634** | 1000G_2504_high_coverage/data/ERR3242419/HG02634.final.cram |
| **HG02635** | 1000G_2504_high_coverage/data/ERR3242420/HG02635.final.cram |
| **HG02715** | 1000G_2504_high_coverage/data/ERR3242421/HG02715.final.cram |
| **HG02716** | 1000G_2504_high_coverage/data/ERR3242422/HG02716.final.cram |
| **HG02768** | 1000G_2504_high_coverage/data/ERR3242423/HG02768.final.cram |
| **HG02769** | 1000G_2504_high_coverage/data/ERR3242424/HG02769.final.cram |
| **HG02771** | 1000G_2504_high_coverage/data/ERR3242425/HG02771.final.cram |
| **HG02772** | 1000G_2504_high_coverage/data/ERR3242426/HG02772.final.cram |
| **HG02798** | 1000G_2504_high_coverage/data/ERR3242427/HG02798.final.cram |
| **HG02799** | 1000G_2504_high_coverage/data/ERR3242428/HG02799.final.cram |
| **HG02804** | 1000G_2504_high_coverage/data/ERR3242429/HG02804.final.cram |
| **HG02805** | 1000G_2504_high_coverage/data/ERR3242430/HG02805.final.cram |
| **HG02807** | 1000G_2504_high_coverage/data/ERR3242431/HG02807.final.cram |
| **HG02808** | 1000G_2504_high_coverage/data/ERR3242432/HG02808.final.cram |
| **HG02810** | 1000G_2504_high_coverage/data/ERR3242433/HG02810.final.cram |
| **HG02811** | 1000G_2504_high_coverage/data/ERR3242434/HG02811.final.cram |
| **HG02813** | 1000G_2504_high_coverage/data/ERR3242435/HG02813.final.cram |
| **HG02814** | 1000G_2504_high_coverage/data/ERR3242436/HG02814.final.cram |
| **HG02839** | 1000G_2504_high_coverage/data/ERR3242437/HG02839.final.cram |
| **HG02840** | 1000G_2504_high_coverage/data/ERR3242438/HG02840.final.cram |
| **HG02851** | 1000G_2504_high_coverage/data/ERR3242439/HG02851.final.cram |
| **HG02852** | 1000G_2504_high_coverage/data/ERR3242440/HG02852.final.cram |
| **HG02887** | 1000G_2504_high_coverage/data/ERR3242441/HG02887.final.cram |
| **HG02888** | 1000G_2504_high_coverage/data/ERR3242442/HG02888.final.cram |
| **HG02895** | 1000G_2504_high_coverage/data/ERR3242443/HG02895.final.cram |
| **HG02896** | 1000G_2504_high_coverage/data/ERR3242444/HG02896.final.cram |
| **HG03027** | 1000G_2504_high_coverage/data/ERR3242445/HG03027.final.cram |
| **HG03028** | 1000G_2504_high_coverage/data/ERR3242446/HG03028.final.cram |
| **HG03039** | 1000G_2504_high_coverage/data/ERR3242447/HG03039.final.cram |
| **HG03040** | 1000G_2504_high_coverage/data/ERR3242448/HG03040.final.cram |
| **HG03045** | 1000G_2504_high_coverage/data/ERR3242449/HG03045.final.cram |
| **HG03046** | 1000G_2504_high_coverage/data/ERR3242450/HG03046.final.cram |
| **HG03048** | 1000G_2504_high_coverage/data/ERR3242451/HG03048.final.cram |
| **HG03049** | 1000G_2504_high_coverage/data/ERR3242452/HG03049.final.cram |
| **HG03258** | 1000G_2504_high_coverage/data/ERR3242453/HG03258.final.cram |
| **HG03259** | 1000G_2504_high_coverage/data/ERR3242454/HG03259.final.cram |
| **HG02756** | 1000G_2504_high_coverage/data/ERR3242455/HG02756.final.cram |
| **HG02757** | 1000G_2504_high_coverage/data/ERR3242456/HG02757.final.cram |
| **HG02816** | 1000G_2504_high_coverage/data/ERR3242457/HG02816.final.cram |
| **HG02817** | 1000G_2504_high_coverage/data/ERR3242458/HG02817.final.cram |
| **HG02836** | 1000G_2504_high_coverage/data/ERR3242459/HG02836.final.cram |
| **HG02837** | 1000G_2504_high_coverage/data/ERR3242460/HG02837.final.cram |
| **HG02860** | 1000G_2504_high_coverage/data/ERR3242461/HG02860.final.cram |
| **HG02861** | 1000G_2504_high_coverage/data/ERR3242462/HG02861.final.cram |
| **HG02884** | 1000G_2504_high_coverage/data/ERR3242463/HG02884.final.cram |
| **HG02885** | 1000G_2504_high_coverage/data/ERR3242464/HG02885.final.cram |
| **HG02890** | 1000G_2504_high_coverage/data/ERR3242465/HG02890.final.cram |
| **HG02891** | 1000G_2504_high_coverage/data/ERR3242466/HG02891.final.cram |
| **HG03052** | 1000G_2504_high_coverage/data/ERR3242467/HG03052.final.cram |
| **HG03057** | 1000G_2504_high_coverage/data/ERR3242468/HG03057.final.cram |
| **HG03058** | 1000G_2504_high_coverage/data/ERR3242469/HG03058.final.cram |
| **HG03078** | 1000G_2504_high_coverage/data/ERR3242470/HG03078.final.cram |
| **HG03084** | 1000G_2504_high_coverage/data/ERR3242471/HG03084.final.cram |
| **HG03085** | 1000G_2504_high_coverage/data/ERR3242472/HG03085.final.cram |
| **HG03096** | 1000G_2504_high_coverage/data/ERR3242473/HG03096.final.cram |
| **HG03097** | 1000G_2504_high_coverage/data/ERR3242474/HG03097.final.cram |
| **HG03397** | 1000G_2504_high_coverage/data/ERR3242475/HG03397.final.cram |
| **HG03451** | 1000G_2504_high_coverage/data/ERR3242476/HG03451.final.cram |

| | |
|---|---|
| **HG03452** | 1000G_2504_high_coverage/data/ERR3242477/HG03452.final.cram |
| **HG03464** | 1000G_2504_high_coverage/data/ERR3242478/HG03464.final.cram |
| **HG03469** | 1000G_2504_high_coverage/data/ERR3242479/HG03469.final.cram |
| **HG03472** | 1000G_2504_high_coverage/data/ERR3242480/HG03472.final.cram |
| **HG02923** | 1000G_2504_high_coverage/data/ERR3242481/HG02923.final.cram |
| **HG02922** | 1000G_2504_high_coverage/data/ERR3242482/HG02922.final.cram |
| **HG02944** | 1000G_2504_high_coverage/data/ERR3242483/HG02944.final.cram |
| **HG02943** | 1000G_2504_high_coverage/data/ERR3242484/HG02943.final.cram |
| **HG02974** | 1000G_2504_high_coverage/data/ERR3242485/HG02974.final.cram |
| **HG02973** | 1000G_2504_high_coverage/data/ERR3242486/HG02973.final.cram |
| **HG03100** | 1000G_2504_high_coverage/data/ERR3242487/HG03100.final.cram |
| **HG03099** | 1000G_2504_high_coverage/data/ERR3242488/HG03099.final.cram |
| **HG03109** | 1000G_2504_high_coverage/data/ERR3242489/HG03109.final.cram |
| **HG03108** | 1000G_2504_high_coverage/data/ERR3242490/HG03108.final.cram |
| **HG03115** | 1000G_2504_high_coverage/data/ERR3242491/HG03115.final.cram |
| **HG03114** | 1000G_2504_high_coverage/data/ERR3242492/HG03114.final.cram |
| **HG03121** | 1000G_2504_high_coverage/data/ERR3242493/HG03121.final.cram |
| **HG03120** | 1000G_2504_high_coverage/data/ERR3242494/HG03120.final.cram |
| **HG03124** | 1000G_2504_high_coverage/data/ERR3242495/HG03124.final.cram |
| **HG03123** | 1000G_2504_high_coverage/data/ERR3242496/HG03123.final.cram |
| **HG03133** | 1000G_2504_high_coverage/data/ERR3242497/HG03133.final.cram |
| **HG03132** | 1000G_2504_high_coverage/data/ERR3242498/HG03132.final.cram |
| **HG03136** | 1000G_2504_high_coverage/data/ERR3242499/HG03136.final.cram |
| **HG03135** | 1000G_2504_high_coverage/data/ERR3242500/HG03135.final.cram |
| **HG02724** | 1000G_2504_high_coverage/data/ERR3242501/HG02724.final.cram |
| **HG02725** | 1000G_2504_high_coverage/data/ERR3242502/HG02725.final.cram |
| **HG02727** | 1000G_2504_high_coverage/data/ERR3242503/HG02727.final.cram |
| **HG02728** | 1000G_2504_high_coverage/data/ERR3242504/HG02728.final.cram |
| **HG02783** | 1000G_2504_high_coverage/data/ERR3242505/HG02783.final.cram |
| **HG02784** | 1000G_2504_high_coverage/data/ERR3242506/HG02784.final.cram |
| **HG02786** | 1000G_2504_high_coverage/data/ERR3242507/HG02786.final.cram |
| **HG02787** | 1000G_2504_high_coverage/data/ERR3242508/HG02787.final.cram |
| **HG02789** | 1000G_2504_high_coverage/data/ERR3242509/HG02789.final.cram |
| **HG02790** | 1000G_2504_high_coverage/data/ERR3242510/HG02790.final.cram |
| **HG00409** | 1000G_2504_high_coverage/data/ERR3242511/HG00409.final.cram |
| **HG00410** | 1000G_2504_high_coverage/data/ERR3242512/HG00410.final.cram |
| **HG00598** | 1000G_2504_high_coverage/data/ERR3242513/HG00598.final.cram |
| **HG00599** | 1000G_2504_high_coverage/data/ERR3242514/HG00599.final.cram |
| **HG00728** | 1000G_2504_high_coverage/data/ERR3242515/HG00728.final.cram |
| **HG00729** | 1000G_2504_high_coverage/data/ERR3242516/HG00729.final.cram |
| **HG00622** | 1000G_2504_high_coverage/data/ERR3242517/HG00622.final.cram |
| **HG00623** | 1000G_2504_high_coverage/data/ERR3242518/HG00623.final.cram |
| **HG00631** | 1000G_2504_high_coverage/data/ERR3242519/HG00631.final.cram |
| **HG00632** | 1000G_2504_high_coverage/data/ERR3242520/HG00632.final.cram |
| **HG00674** | 1000G_2504_high_coverage/data/ERR3242521/HG00674.final.cram |
| **HG00675** | 1000G_2504_high_coverage/data/ERR3242522/HG00675.final.cram |
| **HG03024** | 1000G_2504_high_coverage/data/ERR3242523/HG03024.final.cram |
| **HG03025** | 1000G_2504_high_coverage/data/ERR3242524/HG03025.final.cram |
| **HG03240** | 1000G_2504_high_coverage/data/ERR3242525/HG03240.final.cram |
| **HG03241** | 1000G_2504_high_coverage/data/ERR3242526/HG03241.final.cram |
| **HG03246** | 1000G_2504_high_coverage/data/ERR3242527/HG03246.final.cram |
| **HG03247** | 1000G_2504_high_coverage/data/ERR3242528/HG03247.final.cram |
| **HG03538** | 1000G_2504_high_coverage/data/ERR3242529/HG03538.final.cram |
| **HG03539** | 1000G_2504_high_coverage/data/ERR3242530/HG03539.final.cram |
| **HG02819** | 1000G_2504_high_coverage/data/ERR3242531/HG02819.final.cram |
| **HG02820** | 1000G_2504_high_coverage/data/ERR3242532/HG02820.final.cram |
| **HG02854** | 1000G_2504_high_coverage/data/ERR3242533/HG02854.final.cram |
| **HG02855** | 1000G_2504_high_coverage/data/ERR3242534/HG02855.final.cram |
| **HG03470** | 1000G_2504_high_coverage/data/ERR3242535/HG03470.final.cram |
| **HG03473** | 1000G_2504_high_coverage/data/ERR3242536/HG03473.final.cram |
| **HG03081** | 1000G_2504_high_coverage/data/ERR3242537/HG03081.final.cram |
| **HG03088** | 1000G_2504_high_coverage/data/ERR3242538/HG03088.final.cram |
| **HG03209** | 1000G_2504_high_coverage/data/ERR3242539/HG03209.final.cram |
| **HG03212** | 1000G_2504_high_coverage/data/ERR3242540/HG03212.final.cram |
| **HG03055** | 1000G_2504_high_coverage/data/ERR3242541/HG03055.final.cram |

| | |
|---|---|
| **HG03376** | 1000G_2504_high_coverage/data/ERR3242542/HG03376.final.cram |
| **HG03380** | 1000G_2504_high_coverage/data/ERR3242543/HG03380.final.cram |
| **HG03385** | 1000G_2504_high_coverage/data/ERR3242544/HG03385.final.cram |
| **HG03388** | 1000G_2504_high_coverage/data/ERR3242545/HG03388.final.cram |
| **HG03401** | 1000G_2504_high_coverage/data/ERR3242546/HG03401.final.cram |
| **HG03419** | 1000G_2504_high_coverage/data/ERR3242547/HG03419.final.cram |
| **HG03061** | 1000G_2504_high_coverage/data/ERR3242548/HG03061.final.cram |
| **HG03428** | 1000G_2504_high_coverage/data/ERR3242549/HG03428.final.cram |
| **HG03433** | 1000G_2504_high_coverage/data/ERR3242550/HG03433.final.cram |
| **HG03439** | 1000G_2504_high_coverage/data/ERR3242551/HG03439.final.cram |
| **HG03442** | 1000G_2504_high_coverage/data/ERR3242552/HG03442.final.cram |
| **HG03445** | 1000G_2504_high_coverage/data/ERR3242553/HG03445.final.cram |
| **HG03446** | 1000G_2504_high_coverage/data/ERR3242554/HG03446.final.cram |
| **HG03460** | 1000G_2504_high_coverage/data/ERR3242555/HG03460.final.cram |
| **HG03461** | 1000G_2504_high_coverage/data/ERR3242556/HG03461.final.cram |
| **HG03478** | 1000G_2504_high_coverage/data/ERR3242557/HG03478.final.cram |
| **HG03479** | 1000G_2504_high_coverage/data/ERR3242558/HG03479.final.cram |
| **HG03066** | 1000G_2504_high_coverage/data/ERR3242559/HG03066.final.cram |
| **HG03069** | 1000G_2504_high_coverage/data/ERR3242560/HG03069.final.cram |
| **HG03072** | 1000G_2504_high_coverage/data/ERR3242561/HG03072.final.cram |
| **HG03410** | 1000G_2504_high_coverage/data/ERR3242562/HG03410.final.cram |
| **HG03436** | 1000G_2504_high_coverage/data/ERR3242563/HG03436.final.cram |
| **HG03437** | 1000G_2504_high_coverage/data/ERR3242564/HG03437.final.cram |
| **HG03449** | 1000G_2504_high_coverage/data/ERR3242565/HG03449.final.cram |
| **HG03458** | 1000G_2504_high_coverage/data/ERR3242566/HG03458.final.cram |
| **HG03476** | 1000G_2504_high_coverage/data/ERR3242567/HG03476.final.cram |
| **HG03484** | 1000G_2504_high_coverage/data/ERR3242568/HG03484.final.cram |
| **HG03485** | 1000G_2504_high_coverage/data/ERR3242569/HG03485.final.cram |
| **HG03547** | 1000G_2504_high_coverage/data/ERR3242570/HG03547.final.cram |
| **HG03548** | 1000G_2504_high_coverage/data/ERR3242571/HG03548.final.cram |
| **HG03556** | 1000G_2504_high_coverage/data/ERR3242572/HG03556.final.cram |
| **HG03557** | 1000G_2504_high_coverage/data/ERR3242573/HG03557.final.cram |
| **HG03391** | 1000G_2504_high_coverage/data/ERR3242574/HG03391.final.cram |
| **HG03455** | 1000G_2504_high_coverage/data/ERR3242575/HG03455.final.cram |
| **HG03559** | 1000G_2504_high_coverage/data/ERR3242576/HG03559.final.cram |
| **HG03563** | 1000G_2504_high_coverage/data/ERR3242577/HG03563.final.cram |
| **HG03565** | 1000G_2504_high_coverage/data/ERR3242578/HG03565.final.cram |
| **HG03571** | 1000G_2504_high_coverage/data/ERR3242579/HG03571.final.cram |
| **HG03572** | 1000G_2504_high_coverage/data/ERR3242580/HG03572.final.cram |
| **HG03575** | 1000G_2504_high_coverage/data/ERR3242581/HG03575.final.cram |
| **HG03577** | 1000G_2504_high_coverage/data/ERR3242582/HG03577.final.cram |
| **HG03578** | 1000G_2504_high_coverage/data/ERR3242583/HG03578.final.cram |
| **HG03583** | 1000G_2504_high_coverage/data/ERR3242584/HG03583.final.cram |
| **HG03515** | 1000G_2504_high_coverage/data/ERR3242585/HG03515.final.cram |
| **HG03514** | 1000G_2504_high_coverage/data/ERR3242586/HG03514.final.cram |
| **HG03268** | 1000G_2504_high_coverage/data/ERR3242587/HG03268.final.cram |
| **HG03267** | 1000G_2504_high_coverage/data/ERR3242588/HG03267.final.cram |
| **HG03271** | 1000G_2504_high_coverage/data/ERR3242589/HG03271.final.cram |
| **HG03270** | 1000G_2504_high_coverage/data/ERR3242590/HG03270.final.cram |
| **HG03280** | 1000G_2504_high_coverage/data/ERR3242591/HG03280.final.cram |
| **HG03279** | 1000G_2504_high_coverage/data/ERR3242592/HG03279.final.cram |
| **HG03291** | 1000G_2504_high_coverage/data/ERR3242593/HG03291.final.cram |
| **HG03295** | 1000G_2504_high_coverage/data/ERR3242594/HG03295.final.cram |
| **HG03294** | 1000G_2504_high_coverage/data/ERR3242595/HG03294.final.cram |
| **HG03301** | 1000G_2504_high_coverage/data/ERR3242596/HG03301.final.cram |
| **HG03300** | 1000G_2504_high_coverage/data/ERR3242597/HG03300.final.cram |
| **HG03367** | 1000G_2504_high_coverage/data/ERR3242598/HG03367.final.cram |
| **HG03366** | 1000G_2504_high_coverage/data/ERR3242599/HG03366.final.cram |
| **HG02947** | 1000G_2504_high_coverage/data/ERR3242600/HG02947.final.cram |
| **HG02946** | 1000G_2504_high_coverage/data/ERR3242601/HG02946.final.cram |
| **HG02971** | 1000G_2504_high_coverage/data/ERR3242602/HG02971.final.cram |
| **HG02970** | 1000G_2504_high_coverage/data/ERR3242603/HG02970.final.cram |
| **HG02953** | 1000G_2504_high_coverage/data/ERR3242604/HG02953.final.cram |
| **HG02952** | 1000G_2504_high_coverage/data/ERR3242605/HG02952.final.cram |
| **HG02977** | 1000G_2504_high_coverage/data/ERR3242606/HG02977.final.cram |

| | |
|---|---|
| **HG02976** | 1000G_2504_high_coverage/data/ERR3242607/HG02976.final.cram |
| **HG03118** | 1000G_2504_high_coverage/data/ERR3242608/HG03118.final.cram |
| **HG03117** | 1000G_2504_high_coverage/data/ERR3242609/HG03117.final.cram |
| **HG03130** | 1000G_2504_high_coverage/data/ERR3242610/HG03130.final.cram |
| **HG03129** | 1000G_2504_high_coverage/data/ERR3242611/HG03129.final.cram |
| **HG03160** | 1000G_2504_high_coverage/data/ERR3242612/HG03160.final.cram |
| **HG03159** | 1000G_2504_high_coverage/data/ERR3242613/HG03159.final.cram |
| **HG03163** | 1000G_2504_high_coverage/data/ERR3242614/HG03163.final.cram |
| **HG03162** | 1000G_2504_high_coverage/data/ERR3242615/HG03162.final.cram |
| **HG03169** | 1000G_2504_high_coverage/data/ERR3242616/HG03169.final.cram |
| **HG03168** | 1000G_2504_high_coverage/data/ERR3242617/HG03168.final.cram |
| **HG03172** | 1000G_2504_high_coverage/data/ERR3242618/HG03172.final.cram |
| **HG03196** | 1000G_2504_high_coverage/data/ERR3242619/HG03196.final.cram |
| **HG03195** | 1000G_2504_high_coverage/data/ERR3242620/HG03195.final.cram |
| **HG03199** | 1000G_2504_high_coverage/data/ERR3242621/HG03199.final.cram |
| **HG03198** | 1000G_2504_high_coverage/data/ERR3242622/HG03198.final.cram |
| **HG03343** | 1000G_2504_high_coverage/data/ERR3242623/HG03343.final.cram |
| **HG03342** | 1000G_2504_high_coverage/data/ERR3242624/HG03342.final.cram |
| **HG03370** | 1000G_2504_high_coverage/data/ERR3242625/HG03370.final.cram |
| **HG03369** | 1000G_2504_high_coverage/data/ERR3242626/HG03369.final.cram |
| **HG03234** | 1000G_2504_high_coverage/data/ERR3242627/HG03234.final.cram |
| **HG03235** | 1000G_2504_high_coverage/data/ERR3242628/HG03235.final.cram |
| **HG03488** | 1000G_2504_high_coverage/data/ERR3242629/HG03488.final.cram |
| **HG03490** | 1000G_2504_high_coverage/data/ERR3242630/HG03490.final.cram |
| **HG03491** | 1000G_2504_high_coverage/data/ERR3242631/HG03491.final.cram |
| **HG03619** | 1000G_2504_high_coverage/data/ERR3242632/HG03619.final.cram |
| **HG03634** | 1000G_2504_high_coverage/data/ERR3242633/HG03634.final.cram |
| **HG03667** | 1000G_2504_high_coverage/data/ERR3242634/HG03667.final.cram |
| **HG03668** | 1000G_2504_high_coverage/data/ERR3242635/HG03668.final.cram |
| **HG03015** | 1000G_2504_high_coverage/data/ERR3242636/HG03015.final.cram |
| **HG03016** | 1000G_2504_high_coverage/data/ERR3242637/HG03016.final.cram |
| **HG03702** | 1000G_2504_high_coverage/data/ERR3242638/HG03702.final.cram |
| **HG03703** | 1000G_2504_high_coverage/data/ERR3242639/HG03703.final.cram |
| **HG03708** | 1000G_2504_high_coverage/data/ERR3242640/HG03708.final.cram |
| **HG03709** | 1000G_2504_high_coverage/data/ERR3242641/HG03709.final.cram |
| **HG03006** | 1000G_2504_high_coverage/data/ERR3242642/HG03006.final.cram |
| **HG03007** | 1000G_2504_high_coverage/data/ERR3242643/HG03007.final.cram |
| **HG03615** | 1000G_2504_high_coverage/data/ERR3242644/HG03615.final.cram |
| **HG03616** | 1000G_2504_high_coverage/data/ERR3242645/HG03616.final.cram |
| **HG03793** | 1000G_2504_high_coverage/data/ERR3242646/HG03793.final.cram |
| **HG03796** | 1000G_2504_high_coverage/data/ERR3242647/HG03796.final.cram |
| **HG03800** | 1000G_2504_high_coverage/data/ERR3242648/HG03800.final.cram |
| **HG03803** | 1000G_2504_high_coverage/data/ERR3242649/HG03803.final.cram |
| **HG03802** | 1000G_2504_high_coverage/data/ERR3242650/HG03802.final.cram |
| **HG03815** | 1000G_2504_high_coverage/data/ERR3242651/HG03815.final.cram |
| **HG03814** | 1000G_2504_high_coverage/data/ERR3242652/HG03814.final.cram |
| **HG03824** | 1000G_2504_high_coverage/data/ERR3242653/HG03824.final.cram |
| **HG03823** | 1000G_2504_high_coverage/data/ERR3242654/HG03823.final.cram |
| **HG03826** | 1000G_2504_high_coverage/data/ERR3242655/HG03826.final.cram |
| **HG03830** | 1000G_2504_high_coverage/data/ERR3242656/HG03830.final.cram |
| **HG03829** | 1000G_2504_high_coverage/data/ERR3242657/HG03829.final.cram |
| **HG03833** | 1000G_2504_high_coverage/data/ERR3242658/HG03833.final.cram |
| **HG03832** | 1000G_2504_high_coverage/data/ERR3242659/HG03832.final.cram |
| **HG03914** | 1000G_2504_high_coverage/data/ERR3242660/HG03914.final.cram |
| **HG03913** | 1000G_2504_high_coverage/data/ERR3242661/HG03913.final.cram |
| **HG03941** | 1000G_2504_high_coverage/data/ERR3242662/HG03941.final.cram |
| **HG03940** | 1000G_2504_high_coverage/data/ERR3242663/HG03940.final.cram |
| **HG04155** | 1000G_2504_high_coverage/data/ERR3242664/HG04155.final.cram |
| **HG04156** | 1000G_2504_high_coverage/data/ERR3242665/HG04156.final.cram |
| **HG03644** | 1000G_2504_high_coverage/data/ERR3242666/HG03644.final.cram |
| **HG03643** | 1000G_2504_high_coverage/data/ERR3242667/HG03643.final.cram |
| **HG03679** | 1000G_2504_high_coverage/data/ERR3242668/HG03679.final.cram |
| **HG03642** | 1000G_2504_high_coverage/data/ERR3242669/HG03642.final.cram |
| **HG03673** | 1000G_2504_high_coverage/data/ERR3242670/HG03673.final.cram |
| **HG03684** | 1000G_2504_high_coverage/data/ERR3242671/HG03684.final.cram |

| | |
|---|---|
| HG03885 | 1000G_2504_high_coverage/data/ERR3242672/HG03885.final.cram |
| HG03886 | 1000G_2504_high_coverage/data/ERR3242673/HG03886.final.cram |
| HG03692 | 1000G_2504_high_coverage/data/ERR3242674/HG03692.final.cram |
| HG03693 | 1000G_2504_high_coverage/data/ERR3242675/HG03693.final.cram |
| HG03740 | 1000G_2504_high_coverage/data/ERR3242676/HG03740.final.cram |
| HG03741 | 1000G_2504_high_coverage/data/ERR3242677/HG03741.final.cram |
| HG03884 | 1000G_2504_high_coverage/data/ERR3242678/HG03884.final.cram |
| HG03743 | 1000G_2504_high_coverage/data/ERR3242679/HG03743.final.cram |
| HG03757 | 1000G_2504_high_coverage/data/ERR3242680/HG03757.final.cram |
| HG03746 | 1000G_2504_high_coverage/data/ERR3242681/HG03746.final.cram |
| HG03888 | 1000G_2504_high_coverage/data/ERR3242682/HG03888.final.cram |
| HG03750 | 1000G_2504_high_coverage/data/ERR3242683/HG03750.final.cram |
| HG03754 | 1000G_2504_high_coverage/data/ERR3242684/HG03754.final.cram |
| HG03756 | 1000G_2504_high_coverage/data/ERR3242685/HG03756.final.cram |
| HG03844 | 1000G_2504_high_coverage/data/ERR3242686/HG03844.final.cram |
| HG04006 | 1000G_2504_high_coverage/data/ERR3242687/HG04006.final.cram |
| HG03838 | 1000G_2504_high_coverage/data/ERR3242688/HG03838.final.cram |
| HG03849 | 1000G_2504_high_coverage/data/ERR3242689/HG03849.final.cram |
| HG03950 | 1000G_2504_high_coverage/data/ERR3242690/HG03950.final.cram |
| HG03858 | 1000G_2504_high_coverage/data/ERR3242691/HG03858.final.cram |
| HG03887 | 1000G_2504_high_coverage/data/ERR3242692/HG03887.final.cram |
| HG03986 | 1000G_2504_high_coverage/data/ERR3242693/HG03986.final.cram |
| HG03896 | 1000G_2504_high_coverage/data/ERR3242694/HG03896.final.cram |
| HG03898 | 1000G_2504_high_coverage/data/ERR3242695/HG03898.final.cram |
| HG03900 | 1000G_2504_high_coverage/data/ERR3242696/HG03900.final.cram |
| HG03943 | 1000G_2504_high_coverage/data/ERR3242697/HG03943.final.cram |
| HG03713 | 1000G_2504_high_coverage/data/ERR3242698/HG03713.final.cram |
| HG03716 | 1000G_2504_high_coverage/data/ERR3242699/HG03716.final.cram |
| HG03714 | 1000G_2504_high_coverage/data/ERR3242700/HG03714.final.cram |
| HG03717 | 1000G_2504_high_coverage/data/ERR3242701/HG03717.final.cram |
| HG03722 | 1000G_2504_high_coverage/data/ERR3242702/HG03722.final.cram |
| HG03730 | 1000G_2504_high_coverage/data/ERR3242703/HG03730.final.cram |
| HG03727 | 1000G_2504_high_coverage/data/ERR3242704/HG03727.final.cram |
| HG03729 | 1000G_2504_high_coverage/data/ERR3242705/HG03729.final.cram |
| HG03731 | 1000G_2504_high_coverage/data/ERR3242706/HG03731.final.cram |
| HG03771 | 1000G_2504_high_coverage/data/ERR3242707/HG03771.final.cram |
| HG03770 | 1000G_2504_high_coverage/data/ERR3242708/HG03770.final.cram |
| HG03882 | 1000G_2504_high_coverage/data/ERR3242709/HG03882.final.cram |
| HG03781 | 1000G_2504_high_coverage/data/ERR3242710/HG03781.final.cram |
| HG03775 | 1000G_2504_high_coverage/data/ERR3242711/HG03775.final.cram |
| HG03773 | 1000G_2504_high_coverage/data/ERR3242712/HG03773.final.cram |
| HG03772 | 1000G_2504_high_coverage/data/ERR3242713/HG03772.final.cram |
| HG03786 | 1000G_2504_high_coverage/data/ERR3242714/HG03786.final.cram |
| HG03784 | 1000G_2504_high_coverage/data/ERR3242715/HG03784.final.cram |
| HG03787 | 1000G_2504_high_coverage/data/ERR3242716/HG03787.final.cram |
| HG03788 | 1000G_2504_high_coverage/data/ERR3242717/HG03788.final.cram |
| HG03789 | 1000G_2504_high_coverage/data/ERR3242718/HG03789.final.cram |
| HG03864 | 1000G_2504_high_coverage/data/ERR3242719/HG03864.final.cram |
| HG03869 | 1000G_2504_high_coverage/data/ERR3242720/HG03869.final.cram |
| HG03862 | 1000G_2504_high_coverage/data/ERR3242721/HG03862.final.cram |
| HG03874 | 1000G_2504_high_coverage/data/ERR3242722/HG03874.final.cram |
| HG03978 | 1000G_2504_high_coverage/data/ERR3242723/HG03978.final.cram |
| HG03971 | 1000G_2504_high_coverage/data/ERR3242724/HG03971.final.cram |
| HG03974 | 1000G_2504_high_coverage/data/ERR3242725/HG03974.final.cram |
| HG03976 | 1000G_2504_high_coverage/data/ERR3242726/HG03976.final.cram |
| HG03973 | 1000G_2504_high_coverage/data/ERR3242727/HG03973.final.cram |
| HG04002 | 1000G_2504_high_coverage/data/ERR3242728/HG04002.final.cram |
| HG04017 | 1000G_2504_high_coverage/data/ERR3242729/HG04017.final.cram |
| HG04026 | 1000G_2504_high_coverage/data/ERR3242730/HG04026.final.cram |
| HG02439 | 1000G_2504_high_coverage/data/ERR3242731/HG02439.final.cram |
| HG02455 | 1000G_2504_high_coverage/data/ERR3242732/HG02455.final.cram |
| HG02476 | 1000G_2504_high_coverage/data/ERR3242733/HG02476.final.cram |
| HG02477 | 1000G_2504_high_coverage/data/ERR3242734/HG02477.final.cram |
| HG02481 | 1000G_2504_high_coverage/data/ERR3242735/HG02481.final.cram |
| HG02501 | 1000G_2504_high_coverage/data/ERR3242736/HG02501.final.cram |

| | |
|---|---|
| **HG02502** | 1000G_2504_high_coverage/data/ERR3242737/HG02502.final.cram |
| **HG02505** | 1000G_2504_high_coverage/data/ERR3242738/HG02505.final.cram |
| **HG02536** | 1000G_2504_high_coverage/data/ERR3242739/HG02536.final.cram |
| **HG02541** | 1000G_2504_high_coverage/data/ERR3242740/HG02541.final.cram |
| **HG02545** | 1000G_2504_high_coverage/data/ERR3242741/HG02545.final.cram |
| **HG02546** | 1000G_2504_high_coverage/data/ERR3242742/HG02546.final.cram |
| **HG02549** | 1000G_2504_high_coverage/data/ERR3242743/HG02549.final.cram |
| **HG02554** | 1000G_2504_high_coverage/data/ERR3242744/HG02554.final.cram |
| **HG02555** | 1000G_2504_high_coverage/data/ERR3242745/HG02555.final.cram |
| **HG02557** | 1000G_2504_high_coverage/data/ERR3242746/HG02557.final.cram |
| **HG02558** | 1000G_2504_high_coverage/data/ERR3242747/HG02558.final.cram |
| **HG02577** | 1000G_2504_high_coverage/data/ERR3242748/HG02577.final.cram |
| **HG02580** | 1000G_2504_high_coverage/data/ERR3242749/HG02580.final.cram |
| **HG01088** | 1000G_2504_high_coverage/data/ERR3242750/HG01088.final.cram |
| **HG01089** | 1000G_2504_high_coverage/data/ERR3242751/HG01089.final.cram |
| **HG01302** | 1000G_2504_high_coverage/data/ERR3242752/HG01302.final.cram |
| **HG01303** | 1000G_2504_high_coverage/data/ERR3242753/HG01303.final.cram |
| **HG01311** | 1000G_2504_high_coverage/data/ERR3242754/HG01311.final.cram |
| **HG01312** | 1000G_2504_high_coverage/data/ERR3242755/HG01312.final.cram |
| **HG01323** | 1000G_2504_high_coverage/data/ERR3242756/HG01323.final.cram |
| **HG01325** | 1000G_2504_high_coverage/data/ERR3242757/HG01325.final.cram |
| **HG01326** | 1000G_2504_high_coverage/data/ERR3242758/HG01326.final.cram |
| **HG01392** | 1000G_2504_high_coverage/data/ERR3242759/HG01392.final.cram |
| **HG01393** | 1000G_2504_high_coverage/data/ERR3242760/HG01393.final.cram |
| **HG01395** | 1000G_2504_high_coverage/data/ERR3242761/HG01395.final.cram |
| **HG01396** | 1000G_2504_high_coverage/data/ERR3242762/HG01396.final.cram |
| **HG01402** | 1000G_2504_high_coverage/data/ERR3242763/HG01402.final.cram |
| **HG01403** | 1000G_2504_high_coverage/data/ERR3242764/HG01403.final.cram |
| **HG01413** | 1000G_2504_high_coverage/data/ERR3242765/HG01413.final.cram |
| **HG01414** | 1000G_2504_high_coverage/data/ERR3242766/HG01414.final.cram |
| **HG00742** | 1000G_2504_high_coverage/data/ERR3242767/HG00742.final.cram |
| **HG00743** | 1000G_2504_high_coverage/data/ERR3242768/HG00743.final.cram |
| **HG01058** | 1000G_2504_high_coverage/data/ERR3242769/HG01058.final.cram |
| **HG01063** | 1000G_2504_high_coverage/data/ERR3242770/HG01063.final.cram |
| **HG01064** | 1000G_2504_high_coverage/data/ERR3242771/HG01064.final.cram |
| **HG01077** | 1000G_2504_high_coverage/data/ERR3242772/HG01077.final.cram |
| **HG01092** | 1000G_2504_high_coverage/data/ERR3242773/HG01092.final.cram |
| **HG01161** | 1000G_2504_high_coverage/data/ERR3242774/HG01161.final.cram |
| **HG01162** | 1000G_2504_high_coverage/data/ERR3242775/HG01162.final.cram |
| **HG01164** | 1000G_2504_high_coverage/data/ERR3242776/HG01164.final.cram |
| **HG01200** | 1000G_2504_high_coverage/data/ERR3242777/HG01200.final.cram |
| **HG01286** | 1000G_2504_high_coverage/data/ERR3242778/HG01286.final.cram |
| **HG01305** | 1000G_2504_high_coverage/data/ERR3242779/HG01305.final.cram |
| **HG01308** | 1000G_2504_high_coverage/data/ERR3242780/HG01308.final.cram |
| **HG01398** | 1000G_2504_high_coverage/data/ERR3242781/HG01398.final.cram |
| **HG01405** | 1000G_2504_high_coverage/data/ERR3242782/HG01405.final.cram |
| **HG01412** | 1000G_2504_high_coverage/data/ERR3242783/HG01412.final.cram |
| **HG01119** | 1000G_2504_high_coverage/data/ERR3242784/HG01119.final.cram |
| **HG01121** | 1000G_2504_high_coverage/data/ERR3242785/HG01121.final.cram |
| **HG01122** | 1000G_2504_high_coverage/data/ERR3242786/HG01122.final.cram |
| **HG01130** | 1000G_2504_high_coverage/data/ERR3242787/HG01130.final.cram |
| **HG01131** | 1000G_2504_high_coverage/data/ERR3242788/HG01131.final.cram |
| **HG01142** | 1000G_2504_high_coverage/data/ERR3242789/HG01142.final.cram |
| **HG01269** | 1000G_2504_high_coverage/data/ERR3242790/HG01269.final.cram |
| **HG01280** | 1000G_2504_high_coverage/data/ERR3242791/HG01280.final.cram |
| **HG01281** | 1000G_2504_high_coverage/data/ERR3242792/HG01281.final.cram |
| **HG01284** | 1000G_2504_high_coverage/data/ERR3242793/HG01284.final.cram |
| **HG01362** | 1000G_2504_high_coverage/data/ERR3242794/HG01362.final.cram |
| **HG01363** | 1000G_2504_high_coverage/data/ERR3242795/HG01363.final.cram |
| **HG01369** | 1000G_2504_high_coverage/data/ERR3242796/HG01369.final.cram |
| **HG01372** | 1000G_2504_high_coverage/data/ERR3242797/HG01372.final.cram |
| **HG01431** | 1000G_2504_high_coverage/data/ERR3242798/HG01431.final.cram |
| **HG01432** | 1000G_2504_high_coverage/data/ERR3242799/HG01432.final.cram |
| **HG01435** | 1000G_2504_high_coverage/data/ERR3242800/HG01435.final.cram |
| **HG01443** | 1000G_2504_high_coverage/data/ERR3242801/HG01443.final.cram |

| | |
|---|---|
| **HG01444** | 1000G_2504_high_coverage/data/ERR3242802/HG01444.final.cram |
| **HG01447** | 1000G_2504_high_coverage/data/ERR3242803/HG01447.final.cram |
| **HG01459** | 1000G_2504_high_coverage/data/ERR3242804/HG01459.final.cram |
| **HG01468** | 1000G_2504_high_coverage/data/ERR3242805/HG01468.final.cram |
| **HG01474** | 1000G_2504_high_coverage/data/ERR3242806/HG01474.final.cram |
| **HG01479** | 1000G_2504_high_coverage/data/ERR3242807/HG01479.final.cram |
| **HG01485** | 1000G_2504_high_coverage/data/ERR3242808/HG01485.final.cram |
| **HG01486** | 1000G_2504_high_coverage/data/ERR3242809/HG01486.final.cram |
| **HG01556** | 1000G_2504_high_coverage/data/ERR3242810/HG01556.final.cram |
| **HG01961** | 1000G_2504_high_coverage/data/ERR3242811/HG01961.final.cram |
| **HG01965** | 1000G_2504_high_coverage/data/ERR3242812/HG01965.final.cram |
| **HG02006** | 1000G_2504_high_coverage/data/ERR3242813/HG02006.final.cram |
| **HG02102** | 1000G_2504_high_coverage/data/ERR3242814/HG02102.final.cram |
| **HG02150** | 1000G_2504_high_coverage/data/ERR3242815/HG02150.final.cram |
| **HG02252** | 1000G_2504_high_coverage/data/ERR3242816/HG02252.final.cram |
| **HG02253** | 1000G_2504_high_coverage/data/ERR3242817/HG02253.final.cram |
| **HG02262** | 1000G_2504_high_coverage/data/ERR3242818/HG02262.final.cram |
| **HG02265** | 1000G_2504_high_coverage/data/ERR3242819/HG02265.final.cram |
| **HG02266** | 1000G_2504_high_coverage/data/ERR3242820/HG02266.final.cram |
| **HG02274** | 1000G_2504_high_coverage/data/ERR3242821/HG02274.final.cram |
| **HG02275** | 1000G_2504_high_coverage/data/ERR3242822/HG02275.final.cram |
| **HG02304** | 1000G_2504_high_coverage/data/ERR3242823/HG02304.final.cram |
| **HG02312** | 1000G_2504_high_coverage/data/ERR3242824/HG02312.final.cram |
| **HG02345** | 1000G_2504_high_coverage/data/ERR3242825/HG02345.final.cram |
| **HG02348** | 1000G_2504_high_coverage/data/ERR3242826/HG02348.final.cram |
| **HG02425** | 1000G_2504_high_coverage/data/ERR3242827/HG02425.final.cram |
| **HG02408** | 1000G_2504_high_coverage/data/ERR3242828/HG02408.final.cram |
| **HG02410** | 1000G_2504_high_coverage/data/ERR3242829/HG02410.final.cram |
| **HG03237** | 1000G_2504_high_coverage/data/ERR3242830/HG03237.final.cram |
| **HG03238** | 1000G_2504_high_coverage/data/ERR3242831/HG03238.final.cram |
| **HG03607** | 1000G_2504_high_coverage/data/ERR3242832/HG03607.final.cram |
| **HG03805** | 1000G_2504_high_coverage/data/ERR3242833/HG03805.final.cram |
| **HG03908** | 1000G_2504_high_coverage/data/ERR3242834/HG03908.final.cram |
| **HG03907** | 1000G_2504_high_coverage/data/ERR3242835/HG03907.final.cram |
| **HG03926** | 1000G_2504_high_coverage/data/ERR3242836/HG03926.final.cram |
| **HG03925** | 1000G_2504_high_coverage/data/ERR3242837/HG03925.final.cram |
| **HG03594** | 1000G_2504_high_coverage/data/ERR3242838/HG03594.final.cram |
| **HG03595** | 1000G_2504_high_coverage/data/ERR3242839/HG03595.final.cram |
| **HG03809** | 1000G_2504_high_coverage/data/ERR3242840/HG03809.final.cram |
| **HG03808** | 1000G_2504_high_coverage/data/ERR3242841/HG03808.final.cram |
| **HG03009** | 1000G_2504_high_coverage/data/ERR3242842/HG03009.final.cram |
| **HG03812** | 1000G_2504_high_coverage/data/ERR3242843/HG03812.final.cram |
| **HG03817** | 1000G_2504_high_coverage/data/ERR3242844/HG03817.final.cram |
| **HG03902** | 1000G_2504_high_coverage/data/ERR3242845/HG03902.final.cram |
| **HG03012** | 1000G_2504_high_coverage/data/ERR3242846/HG03012.final.cram |
| **HG03905** | 1000G_2504_high_coverage/data/ERR3242847/HG03905.final.cram |
| **HG03911** | 1000G_2504_high_coverage/data/ERR3242848/HG03911.final.cram |
| **HG03910** | 1000G_2504_high_coverage/data/ERR3242849/HG03910.final.cram |
| **HG03916** | 1000G_2504_high_coverage/data/ERR3242850/HG03916.final.cram |
| **HG03919** | 1000G_2504_high_coverage/data/ERR3242851/HG03919.final.cram |
| **HG03928** | 1000G_2504_high_coverage/data/ERR3242852/HG03928.final.cram |
| **HG03931** | 1000G_2504_high_coverage/data/ERR3242853/HG03931.final.cram |
| **HG03585** | 1000G_2504_high_coverage/data/ERR3242854/HG03585.final.cram |
| **HG03937** | 1000G_2504_high_coverage/data/ERR3242855/HG03937.final.cram |
| **HG03934** | 1000G_2504_high_coverage/data/ERR3242856/HG03934.final.cram |
| **HG04146** | 1000G_2504_high_coverage/data/ERR3242857/HG04146.final.cram |
| **HG03589** | 1000G_2504_high_coverage/data/ERR3242858/HG03589.final.cram |
| **HG04159** | 1000G_2504_high_coverage/data/ERR3242859/HG04159.final.cram |
| **HG03593** | 1000G_2504_high_coverage/data/ERR3242860/HG03593.final.cram |
| **HG04186** | 1000G_2504_high_coverage/data/ERR3242861/HG04186.final.cram |
| **HG03598** | 1000G_2504_high_coverage/data/ERR3242862/HG03598.final.cram |
| **HG03755** | 1000G_2504_high_coverage/data/ERR3242863/HG03755.final.cram |
| **HG03850** | 1000G_2504_high_coverage/data/ERR3242864/HG03850.final.cram |
| **HG03894** | 1000G_2504_high_coverage/data/ERR3242865/HG03894.final.cram |
| **HG03897** | 1000G_2504_high_coverage/data/ERR3242866/HG03897.final.cram |

| | |
|---|---|
| HG03944 | 1000G_2504_high_coverage/data/ERR3242867/HG03944.final.cram |
| HG03646 | 1000G_2504_high_coverage/data/ERR3242868/HG03646.final.cram |
| HG03645 | 1000G_2504_high_coverage/data/ERR3242869/HG03645.final.cram |
| HG03999 | 1000G_2504_high_coverage/data/ERR3242870/HG03999.final.cram |
| HG03672 | 1000G_2504_high_coverage/data/ERR3242871/HG03672.final.cram |
| HG03680 | 1000G_2504_high_coverage/data/ERR3242872/HG03680.final.cram |
| HG03681 | 1000G_2504_high_coverage/data/ERR3242873/HG03681.final.cram |
| HG03685 | 1000G_2504_high_coverage/data/ERR3242874/HG03685.final.cram |
| HG03686 | 1000G_2504_high_coverage/data/ERR3242875/HG03686.final.cram |
| HG03687 | 1000G_2504_high_coverage/data/ERR3242876/HG03687.final.cram |
| HG03689 | 1000G_2504_high_coverage/data/ERR3242877/HG03689.final.cram |
| HG03690 | 1000G_2504_high_coverage/data/ERR3242878/HG03690.final.cram |
| HG03691 | 1000G_2504_high_coverage/data/ERR3242879/HG03691.final.cram |
| HG03694 | 1000G_2504_high_coverage/data/ERR3242880/HG03694.final.cram |
| HG03695 | 1000G_2504_high_coverage/data/ERR3242881/HG03695.final.cram |
| HG03696 | 1000G_2504_high_coverage/data/ERR3242882/HG03696.final.cram |
| HG03697 | 1000G_2504_high_coverage/data/ERR3242883/HG03697.final.cram |
| HG03698 | 1000G_2504_high_coverage/data/ERR3242884/HG03698.final.cram |
| HG03711 | 1000G_2504_high_coverage/data/ERR3242885/HG03711.final.cram |
| HG03733 | 1000G_2504_high_coverage/data/ERR3242886/HG03733.final.cram |
| HG03736 | 1000G_2504_high_coverage/data/ERR3242887/HG03736.final.cram |
| HG03738 | 1000G_2504_high_coverage/data/ERR3242888/HG03738.final.cram |
| HG03744 | 1000G_2504_high_coverage/data/ERR3242889/HG03744.final.cram |
| HG03745 | 1000G_2504_high_coverage/data/ERR3242890/HG03745.final.cram |
| HG03752 | 1000G_2504_high_coverage/data/ERR3242891/HG03752.final.cram |
| HG03753 | 1000G_2504_high_coverage/data/ERR3242892/HG03753.final.cram |
| HG03760 | 1000G_2504_high_coverage/data/ERR3242893/HG03760.final.cram |
| HG03836 | 1000G_2504_high_coverage/data/ERR3242894/HG03836.final.cram |
| HG03837 | 1000G_2504_high_coverage/data/ERR3242895/HG03837.final.cram |
| HG03846 | 1000G_2504_high_coverage/data/ERR3242896/HG03846.final.cram |
| HG03848 | 1000G_2504_high_coverage/data/ERR3242897/HG03848.final.cram |
| HG03851 | 1000G_2504_high_coverage/data/ERR3242898/HG03851.final.cram |
| HG03856 | 1000G_2504_high_coverage/data/ERR3242899/HG03856.final.cram |
| HG03895 | 1000G_2504_high_coverage/data/ERR3242900/HG03895.final.cram |
| HG03890 | 1000G_2504_high_coverage/data/ERR3242901/HG03890.final.cram |
| HG04033 | 1000G_2504_high_coverage/data/ERR3242902/HG04033.final.cram |
| HG03947 | 1000G_2504_high_coverage/data/ERR3242903/HG03947.final.cram |
| HG03953 | 1000G_2504_high_coverage/data/ERR3242904/HG03953.final.cram |
| HG03949 | 1000G_2504_high_coverage/data/ERR3242905/HG03949.final.cram |
| HG04035 | 1000G_2504_high_coverage/data/ERR3242906/HG04035.final.cram |
| HG04029 | 1000G_2504_high_coverage/data/ERR3242907/HG04029.final.cram |
| HG04042 | 1000G_2504_high_coverage/data/ERR3242908/HG04042.final.cram |
| HG04099 | 1000G_2504_high_coverage/data/ERR3242909/HG04099.final.cram |
| HG03977 | 1000G_2504_high_coverage/data/ERR3242910/HG03977.final.cram |
| HG03780 | 1000G_2504_high_coverage/data/ERR3242911/HG03780.final.cram |
| HG03718 | 1000G_2504_high_coverage/data/ERR3242912/HG03718.final.cram |
| HG04018 | 1000G_2504_high_coverage/data/ERR3242913/HG04018.final.cram |
| HG03742 | 1000G_2504_high_coverage/data/ERR3242914/HG03742.final.cram |
| HG03779 | 1000G_2504_high_coverage/data/ERR3242915/HG03779.final.cram |
| HG03777 | 1000G_2504_high_coverage/data/ERR3242916/HG03777.final.cram |
| HG03785 | 1000G_2504_high_coverage/data/ERR3242917/HG03785.final.cram |
| HG03790 | 1000G_2504_high_coverage/data/ERR3242918/HG03790.final.cram |
| HG03792 | 1000G_2504_high_coverage/data/ERR3242919/HG03792.final.cram |
| HG03782 | 1000G_2504_high_coverage/data/ERR3242920/HG03782.final.cram |
| HG03873 | 1000G_2504_high_coverage/data/ERR3242921/HG03873.final.cram |
| HG03861 | 1000G_2504_high_coverage/data/ERR3242922/HG03861.final.cram |
| HG03863 | 1000G_2504_high_coverage/data/ERR3242923/HG03863.final.cram |
| HG03872 | 1000G_2504_high_coverage/data/ERR3242924/HG03872.final.cram |
| HG03870 | 1000G_2504_high_coverage/data/ERR3242925/HG03870.final.cram |
| HG03867 | 1000G_2504_high_coverage/data/ERR3242926/HG03867.final.cram |
| HG03963 | 1000G_2504_high_coverage/data/ERR3242927/HG03963.final.cram |
| HG03960 | 1000G_2504_high_coverage/data/ERR3242928/HG03960.final.cram |
| HG03968 | 1000G_2504_high_coverage/data/ERR3242929/HG03968.final.cram |
| HG03720 | 1000G_2504_high_coverage/data/ERR3242930/HG03720.final.cram |
| HG04019 | 1000G_2504_high_coverage/data/ERR3242931/HG04019.final.cram |

| | |
|---|---|
| **HG04025** | 1000G_2504_high_coverage/data/ERR3242932/HG04025.final.cram |
| **HG04020** | 1000G_2504_high_coverage/data/ERR3242933/HG04020.final.cram |
| **HG04014** | 1000G_2504_high_coverage/data/ERR3242934/HG04014.final.cram |
| **HG04015** | 1000G_2504_high_coverage/data/ERR3242935/HG04015.final.cram |
| **HG04022** | 1000G_2504_high_coverage/data/ERR3242936/HG04022.final.cram |
| **HG04063** | 1000G_2504_high_coverage/data/ERR3242937/HG04063.final.cram |
| **HG04093** | 1000G_2504_high_coverage/data/ERR3242938/HG04093.final.cram |
| **HG04118** | 1000G_2504_high_coverage/data/ERR3242939/HG04118.final.cram |
| **HG02568** | 1000G_2504_high_coverage/data/ERR3242940/HG02568.final.cram |
| **HG02759** | 1000G_2504_high_coverage/data/ERR3242941/HG02759.final.cram |
| **HG02760** | 1000G_2504_high_coverage/data/ERR3242942/HG02760.final.cram |
| **HG02763** | 1000G_2504_high_coverage/data/ERR3242943/HG02763.final.cram |
| **HG02870** | 1000G_2504_high_coverage/data/ERR3242944/HG02870.final.cram |
| **HG02878** | 1000G_2504_high_coverage/data/ERR3242945/HG02878.final.cram |
| **HG02879** | 1000G_2504_high_coverage/data/ERR3242946/HG02879.final.cram |
| **HG02881** | 1000G_2504_high_coverage/data/ERR3242947/HG02881.final.cram |
| **HG02882** | 1000G_2504_high_coverage/data/ERR3242948/HG02882.final.cram |
| **HG02982** | 1000G_2504_high_coverage/data/ERR3242949/HG02982.final.cram |
| **HG02983** | 1000G_2504_high_coverage/data/ERR3242950/HG02983.final.cram |
| **HG03054** | 1000G_2504_high_coverage/data/ERR3242951/HG03054.final.cram |
| **HG03060** | 1000G_2504_high_coverage/data/ERR3242952/HG03060.final.cram |
| **HG03063** | 1000G_2504_high_coverage/data/ERR3242953/HG03063.final.cram |
| **HG03064** | 1000G_2504_high_coverage/data/ERR3242954/HG03064.final.cram |
| **HG03073** | 1000G_2504_high_coverage/data/ERR3242955/HG03073.final.cram |
| **HG03074** | 1000G_2504_high_coverage/data/ERR3242956/HG03074.final.cram |
| **HG03077** | 1000G_2504_high_coverage/data/ERR3242957/HG03077.final.cram |
| **HG03079** | 1000G_2504_high_coverage/data/ERR3242958/HG03079.final.cram |
| **HG03082** | 1000G_2504_high_coverage/data/ERR3242959/HG03082.final.cram |
| **HG03086** | 1000G_2504_high_coverage/data/ERR3242960/HG03086.final.cram |
| **HG03091** | 1000G_2504_high_coverage/data/ERR3242961/HG03091.final.cram |
| **HG03095** | 1000G_2504_high_coverage/data/ERR3242962/HG03095.final.cram |
| **HG03224** | 1000G_2504_high_coverage/data/ERR3242963/HG03224.final.cram |
| **HG03225** | 1000G_2504_high_coverage/data/ERR3242964/HG03225.final.cram |
| **HG03378** | 1000G_2504_high_coverage/data/ERR3242965/HG03378.final.cram |
| **HG03382** | 1000G_2504_high_coverage/data/ERR3242966/HG03382.final.cram |
| **HG03394** | 1000G_2504_high_coverage/data/ERR3242967/HG03394.final.cram |
| **HG03432** | 1000G_2504_high_coverage/data/ERR3242968/HG03432.final.cram |
| **HG03457** | 1000G_2504_high_coverage/data/ERR3242969/HG03457.final.cram |
| **HG03558** | 1000G_2504_high_coverage/data/ERR3242970/HG03558.final.cram |
| **HG03567** | 1000G_2504_high_coverage/data/ERR3242971/HG03567.final.cram |
| **HG02938** | 1000G_2504_high_coverage/data/ERR3242972/HG02938.final.cram |
| **HG02941** | 1000G_2504_high_coverage/data/ERR3242973/HG02941.final.cram |
| **HG02968** | 1000G_2504_high_coverage/data/ERR3242974/HG02968.final.cram |
| **HG02979** | 1000G_2504_high_coverage/data/ERR3242975/HG02979.final.cram |
| **HG02981** | 1000G_2504_high_coverage/data/ERR3242976/HG02981.final.cram |
| **HG03103** | 1000G_2504_high_coverage/data/ERR3242977/HG03103.final.cram |
| **HG03105** | 1000G_2504_high_coverage/data/ERR3242978/HG03105.final.cram |
| **HG03111** | 1000G_2504_high_coverage/data/ERR3242979/HG03111.final.cram |
| **HG03112** | 1000G_2504_high_coverage/data/ERR3242980/HG03112.final.cram |
| **HG03126** | 1000G_2504_high_coverage/data/ERR3242981/HG03126.final.cram |
| **HG03127** | 1000G_2504_high_coverage/data/ERR3242982/HG03127.final.cram |
| **HG03139** | 1000G_2504_high_coverage/data/ERR3242983/HG03139.final.cram |
| **HG03157** | 1000G_2504_high_coverage/data/ERR3242984/HG03157.final.cram |
| **HG03166** | 1000G_2504_high_coverage/data/ERR3242985/HG03166.final.cram |
| **HG03175** | 1000G_2504_high_coverage/data/ERR3242986/HG03175.final.cram |
| **HG03189** | 1000G_2504_high_coverage/data/ERR3242987/HG03189.final.cram |
| **HG03190** | 1000G_2504_high_coverage/data/ERR3242988/HG03190.final.cram |
| **HG03193** | 1000G_2504_high_coverage/data/ERR3242989/HG03193.final.cram |
| **HG03202** | 1000G_2504_high_coverage/data/ERR3242990/HG03202.final.cram |
| **HG03265** | 1000G_2504_high_coverage/data/ERR3242991/HG03265.final.cram |
| **HG03297** | 1000G_2504_high_coverage/data/ERR3242992/HG03297.final.cram |
| **HG03298** | 1000G_2504_high_coverage/data/ERR3242993/HG03298.final.cram |
| **HG03303** | 1000G_2504_high_coverage/data/ERR3242994/HG03303.final.cram |
| **HG03304** | 1000G_2504_high_coverage/data/ERR3242995/HG03304.final.cram |
| **HG03311** | 1000G_2504_high_coverage/data/ERR3242996/HG03311.final.cram |

| | |
|---|---|
| **HG03313** | 1000G_2504_high_coverage/data/ERR3242997/HG03313.final.cram |
| **HG03351** | 1000G_2504_high_coverage/data/ERR3242998/HG03351.final.cram |
| **HG03352** | 1000G_2504_high_coverage/data/ERR3242999/HG03352.final.cram |
| **HG03354** | 1000G_2504_high_coverage/data/ERR3243000/HG03354.final.cram |
| **HG03363** | 1000G_2504_high_coverage/data/ERR3243001/HG03363.final.cram |
| **HG03372** | 1000G_2504_high_coverage/data/ERR3243002/HG03372.final.cram |
| **HG03499** | 1000G_2504_high_coverage/data/ERR3243003/HG03499.final.cram |
| **HG03511** | 1000G_2504_high_coverage/data/ERR3243004/HG03511.final.cram |
| **HG03517** | 1000G_2504_high_coverage/data/ERR3243005/HG03517.final.cram |
| **HG03518** | 1000G_2504_high_coverage/data/ERR3243006/HG03518.final.cram |
| **HG03520** | 1000G_2504_high_coverage/data/ERR3243007/HG03520.final.cram |
| **HG03521** | 1000G_2504_high_coverage/data/ERR3243008/HG03521.final.cram |
| **HG02493** | 1000G_2504_high_coverage/data/ERR3243009/HG02493.final.cram |
| **HG02494** | 1000G_2504_high_coverage/data/ERR3243010/HG02494.final.cram |
| **HG02648** | 1000G_2504_high_coverage/data/ERR3243011/HG02648.final.cram |
| **HG02649** | 1000G_2504_high_coverage/data/ERR3243012/HG02649.final.cram |
| **HG02651** | 1000G_2504_high_coverage/data/ERR3243013/HG02651.final.cram |
| **HG02652** | 1000G_2504_high_coverage/data/ERR3243014/HG02652.final.cram |
| **HG02681** | 1000G_2504_high_coverage/data/ERR3243015/HG02681.final.cram |
| **HG02682** | 1000G_2504_high_coverage/data/ERR3243016/HG02682.final.cram |
| **HG02690** | 1000G_2504_high_coverage/data/ERR3243017/HG02690.final.cram |
| **HG02691** | 1000G_2504_high_coverage/data/ERR3243018/HG02691.final.cram |
| **HG02699** | 1000G_2504_high_coverage/data/ERR3243019/HG02699.final.cram |
| **HG02700** | 1000G_2504_high_coverage/data/ERR3243020/HG02700.final.cram |
| **HG02736** | 1000G_2504_high_coverage/data/ERR3243021/HG02736.final.cram |
| **HG02737** | 1000G_2504_high_coverage/data/ERR3243022/HG02737.final.cram |
| **HG02774** | 1000G_2504_high_coverage/data/ERR3243023/HG02774.final.cram |
| **HG02775** | 1000G_2504_high_coverage/data/ERR3243024/HG02775.final.cram |
| **HG02792** | 1000G_2504_high_coverage/data/ERR3243025/HG02792.final.cram |
| **HG02793** | 1000G_2504_high_coverage/data/ERR3243026/HG02793.final.cram |
| **HG03228** | 1000G_2504_high_coverage/data/ERR3243027/HG03228.final.cram |
| **HG03229** | 1000G_2504_high_coverage/data/ERR3243028/HG03229.final.cram |
| **HG03624** | 1000G_2504_high_coverage/data/ERR3243029/HG03624.final.cram |
| **HG03625** | 1000G_2504_high_coverage/data/ERR3243030/HG03625.final.cram |
| **HG03640** | 1000G_2504_high_coverage/data/ERR3243031/HG03640.final.cram |
| **HG03649** | 1000G_2504_high_coverage/data/ERR3243032/HG03649.final.cram |
| **HG03652** | 1000G_2504_high_coverage/data/ERR3243033/HG03652.final.cram |
| **HG03653** | 1000G_2504_high_coverage/data/ERR3243034/HG03653.final.cram |
| **HG03705** | 1000G_2504_high_coverage/data/ERR3243035/HG03705.final.cram |
| **HG03706** | 1000G_2504_high_coverage/data/ERR3243036/HG03706.final.cram |
| **HG03762** | 1000G_2504_high_coverage/data/ERR3243037/HG03762.final.cram |
| **HG01593** | 1000G_2504_high_coverage/data/ERR3243038/HG01593.final.cram |
| **HG02597** | 1000G_2504_high_coverage/data/ERR3243039/HG02597.final.cram |
| **HG02778** | 1000G_2504_high_coverage/data/ERR3243040/HG02778.final.cram |
| **HG03021** | 1000G_2504_high_coverage/data/ERR3243041/HG03021.final.cram |
| **HG03636** | 1000G_2504_high_coverage/data/ERR3243042/HG03636.final.cram |
| **HG03765** | 1000G_2504_high_coverage/data/ERR3243043/HG03765.final.cram |
| **HG03767** | 1000G_2504_high_coverage/data/ERR3243044/HG03767.final.cram |
| **HG01583** | 1000G_2504_high_coverage/data/ERR3243045/HG01583.final.cram |
| **HG01586** | 1000G_2504_high_coverage/data/ERR3243046/HG01586.final.cram |
| **HG01589** | 1000G_2504_high_coverage/data/ERR3243047/HG01589.final.cram |
| **HG02694** | 1000G_2504_high_coverage/data/ERR3243048/HG02694.final.cram |
| **HG02731** | 1000G_2504_high_coverage/data/ERR3243049/HG02731.final.cram |
| **HG02780** | 1000G_2504_high_coverage/data/ERR3243050/HG02780.final.cram |
| **HG03018** | 1000G_2504_high_coverage/data/ERR3243051/HG03018.final.cram |
| **HG03019** | 1000G_2504_high_coverage/data/ERR3243052/HG03019.final.cram |
| **HG03022** | 1000G_2504_high_coverage/data/ERR3243053/HG03022.final.cram |
| **HG03629** | 1000G_2504_high_coverage/data/ERR3243054/HG03629.final.cram |
| **HG03631** | 1000G_2504_high_coverage/data/ERR3243055/HG03631.final.cram |
| **HG03660** | 1000G_2504_high_coverage/data/ERR3243056/HG03660.final.cram |
| **HG03663** | 1000G_2504_high_coverage/data/ERR3243057/HG03663.final.cram |
| **HG04158** | 1000G_2504_high_coverage/data/ERR3243058/HG04158.final.cram |
| **HG04185** | 1000G_2504_high_coverage/data/ERR3243059/HG04185.final.cram |
| **HG04131** | 1000G_2504_high_coverage/data/ERR3243060/HG04131.final.cram |
| **HG04134** | 1000G_2504_high_coverage/data/ERR3243061/HG04134.final.cram |

| | |
|---|---|
| **HG04140** | 1000G_2504_high_coverage/data/ERR3243062/HG04140.final.cram |
| **HG04141** | 1000G_2504_high_coverage/data/ERR3243063/HG04141.final.cram |
| **HG04173** | 1000G_2504_high_coverage/data/ERR3243064/HG04173.final.cram |
| **HG04182** | 1000G_2504_high_coverage/data/ERR3243065/HG04182.final.cram |
| **HG04183** | 1000G_2504_high_coverage/data/ERR3243066/HG04183.final.cram |
| **HG03821** | 1000G_2504_high_coverage/data/ERR3243067/HG03821.final.cram |
| **HG03922** | 1000G_2504_high_coverage/data/ERR3243068/HG03922.final.cram |
| **HG04180** | 1000G_2504_high_coverage/data/ERR3243069/HG04180.final.cram |
| **HG03600** | 1000G_2504_high_coverage/data/ERR3243070/HG03600.final.cram |
| **HG03603** | 1000G_2504_high_coverage/data/ERR3243071/HG03603.final.cram |
| **HG03604** | 1000G_2504_high_coverage/data/ERR3243072/HG03604.final.cram |
| **HG03611** | 1000G_2504_high_coverage/data/ERR3243073/HG03611.final.cram |
| **HG03917** | 1000G_2504_high_coverage/data/ERR3243074/HG03917.final.cram |
| **HG03920** | 1000G_2504_high_coverage/data/ERR3243075/HG03920.final.cram |
| **HG04144** | 1000G_2504_high_coverage/data/ERR3243076/HG04144.final.cram |
| **HG04152** | 1000G_2504_high_coverage/data/ERR3243077/HG04152.final.cram |
| **HG04153** | 1000G_2504_high_coverage/data/ERR3243078/HG04153.final.cram |
| **HG04161** | 1000G_2504_high_coverage/data/ERR3243079/HG04161.final.cram |
| **HG04162** | 1000G_2504_high_coverage/data/ERR3243080/HG04162.final.cram |
| **HG04164** | 1000G_2504_high_coverage/data/ERR3243081/HG04164.final.cram |
| **HG04171** | 1000G_2504_high_coverage/data/ERR3243082/HG04171.final.cram |
| **HG04176** | 1000G_2504_high_coverage/data/ERR3243083/HG04176.final.cram |
| **HG04177** | 1000G_2504_high_coverage/data/ERR3243084/HG04177.final.cram |
| **HG04188** | 1000G_2504_high_coverage/data/ERR3243085/HG04188.final.cram |
| **HG04189** | 1000G_2504_high_coverage/data/ERR3243086/HG04189.final.cram |
| **HG04194** | 1000G_2504_high_coverage/data/ERR3243087/HG04194.final.cram |
| **HG04195** | 1000G_2504_high_coverage/data/ERR3243088/HG04195.final.cram |
| **HG03995** | 1000G_2504_high_coverage/data/ERR3243089/HG03995.final.cram |
| **HG03899** | 1000G_2504_high_coverage/data/ERR3243090/HG03899.final.cram |
| **HG03951** | 1000G_2504_high_coverage/data/ERR3243091/HG03951.final.cram |
| **HG03857** | 1000G_2504_high_coverage/data/ERR3243092/HG03857.final.cram |
| **HG03854** | 1000G_2504_high_coverage/data/ERR3243093/HG03854.final.cram |
| **HG04100** | 1000G_2504_high_coverage/data/ERR3243094/HG04100.final.cram |
| **HG04210** | 1000G_2504_high_coverage/data/ERR3243095/HG04210.final.cram |
| **HG04075** | 1000G_2504_high_coverage/data/ERR3243096/HG04075.final.cram |
| **HG04229** | 1000G_2504_high_coverage/data/ERR3243097/HG04229.final.cram |
| **HG04227** | 1000G_2504_high_coverage/data/ERR3243098/HG04227.final.cram |
| **HG03945** | 1000G_2504_high_coverage/data/ERR3243099/HG03945.final.cram |
| **HG03955** | 1000G_2504_high_coverage/data/ERR3243100/HG03955.final.cram |
| **HG03990** | 1000G_2504_high_coverage/data/ERR3243101/HG03990.final.cram |
| **HG03991** | 1000G_2504_high_coverage/data/ERR3243102/HG03991.final.cram |
| **HG03985** | 1000G_2504_high_coverage/data/ERR3243103/HG03985.final.cram |
| **HG03989** | 1000G_2504_high_coverage/data/ERR3243104/HG03989.final.cram |
| **HG03998** | 1000G_2504_high_coverage/data/ERR3243105/HG03998.final.cram |
| **HG04003** | 1000G_2504_high_coverage/data/ERR3243106/HG04003.final.cram |
| **HG04038** | 1000G_2504_high_coverage/data/ERR3243107/HG04038.final.cram |
| **HG04039** | 1000G_2504_high_coverage/data/ERR3243108/HG04039.final.cram |
| **HG04047** | 1000G_2504_high_coverage/data/ERR3243109/HG04047.final.cram |
| **HG04107** | 1000G_2504_high_coverage/data/ERR3243110/HG04107.final.cram |
| **HG04106** | 1000G_2504_high_coverage/data/ERR3243111/HG04106.final.cram |
| **HG03774** | 1000G_2504_high_coverage/data/ERR3243112/HG03774.final.cram |
| **HG04001** | 1000G_2504_high_coverage/data/ERR3243113/HG04001.final.cram |
| **HG03866** | 1000G_2504_high_coverage/data/ERR3243114/HG03866.final.cram |
| **HG03969** | 1000G_2504_high_coverage/data/ERR3243115/HG03969.final.cram |
| **HG04023** | 1000G_2504_high_coverage/data/ERR3243116/HG04023.final.cram |
| **HG04060** | 1000G_2504_high_coverage/data/ERR3243117/HG04060.final.cram |
| **HG04090** | 1000G_2504_high_coverage/data/ERR3243118/HG04090.final.cram |
| **HG04056** | 1000G_2504_high_coverage/data/ERR3243119/HG04056.final.cram |
| **HG04054** | 1000G_2504_high_coverage/data/ERR3243120/HG04054.final.cram |
| **HG04061** | 1000G_2504_high_coverage/data/ERR3243121/HG04061.final.cram |
| **HG04062** | 1000G_2504_high_coverage/data/ERR3243122/HG04062.final.cram |
| **HG04059** | 1000G_2504_high_coverage/data/ERR3243123/HG04059.final.cram |
| **HG04076** | 1000G_2504_high_coverage/data/ERR3243124/HG04076.final.cram |
| **HG04080** | 1000G_2504_high_coverage/data/ERR3243125/HG04080.final.cram |
| **HG04094** | 1000G_2504_high_coverage/data/ERR3243126/HG04094.final.cram |

| | |
|---|---|
| **HG04096** | 1000G_2504_high_coverage/data/ERR3243127/HG04096.final.cram |
| **HG04098** | 1000G_2504_high_coverage/data/ERR3243128/HG04098.final.cram |
| **HG03778** | 1000G_2504_high_coverage/data/ERR3243129/HG03778.final.cram |
| **HG04209** | 1000G_2504_high_coverage/data/ERR3243130/HG04209.final.cram |
| **HG04198** | 1000G_2504_high_coverage/data/ERR3243131/HG04198.final.cram |
| **HG04200** | 1000G_2504_high_coverage/data/ERR3243132/HG04200.final.cram |
| **HG04206** | 1000G_2504_high_coverage/data/ERR3243133/HG04206.final.cram |
| **HG04202** | 1000G_2504_high_coverage/data/ERR3243134/HG04202.final.cram |
| **HG04211** | 1000G_2504_high_coverage/data/ERR3243135/HG04211.final.cram |
| **HG04212** | 1000G_2504_high_coverage/data/ERR3243136/HG04212.final.cram |
| **HG04222** | 1000G_2504_high_coverage/data/ERR3243137/HG04222.final.cram |
| **HG04214** | 1000G_2504_high_coverage/data/ERR3243138/HG04214.final.cram |
| **HG04216** | 1000G_2504_high_coverage/data/ERR3243139/HG04216.final.cram |
| **HG04239** | 1000G_2504_high_coverage/data/ERR3243140/HG04239.final.cram |
| **HG04219** | 1000G_2504_high_coverage/data/ERR3243141/HG04219.final.cram |
| **HG04225** | 1000G_2504_high_coverage/data/ERR3243142/HG04225.final.cram |
| **HG04235** | 1000G_2504_high_coverage/data/ERR3243143/HG04235.final.cram |
| **HG04238** | 1000G_2504_high_coverage/data/ERR3243144/HG04238.final.cram |
| **HG03875** | 1000G_2504_high_coverage/data/ERR3243145/HG03875.final.cram |
| **HG03965** | 1000G_2504_high_coverage/data/ERR3243146/HG03965.final.cram |
| **HG04070** | 1000G_2504_high_coverage/data/ERR3243147/HG04070.final.cram |
| **HG03868** | 1000G_2504_high_coverage/data/ERR3243148/HG03868.final.cram |
| **HG03871** | 1000G_2504_high_coverage/data/ERR3243149/HG03871.final.cram |
| **HG03967** | 1000G_2504_high_coverage/data/ERR3243150/HG03967.final.cram |
| **HG00288** | 1000G_2504_high_coverage/data/ERR3243151/HG00288.final.cram |
| **HG00290** | 1000G_2504_high_coverage/data/ERR3243152/HG00290.final.cram |
| **HG00304** | 1000G_2504_high_coverage/data/ERR3243153/HG00304.final.cram |
| **HG01765** | 1000G_2504_high_coverage/data/ERR3243154/HG01765.final.cram |
| **HG01766** | 1000G_2504_high_coverage/data/ERR3243155/HG01766.final.cram |
| **HG01767** | 1000G_2504_high_coverage/data/ERR3243156/HG01767.final.cram |
| **HG01768** | 1000G_2504_high_coverage/data/ERR3243157/HG01768.final.cram |
| **HG01785** | 1000G_2504_high_coverage/data/ERR3243158/HG01785.final.cram |
| **HG01786** | 1000G_2504_high_coverage/data/ERR3243159/HG01786.final.cram |
| **HG02219** | 1000G_2504_high_coverage/data/ERR3243160/HG02219.final.cram |
| **NA18874** | 1000G_2504_high_coverage/data/ERR3243161/NA18874.final.cram |
| **NA12830** | 1000G_2504_high_coverage/data/ERR3243162/NA12830.final.cram |
| **NA12546** | 1000G_2504_high_coverage/data/ERR3243163/NA12546.final.cram |
| **HG00405** | 1000G_2504_high_coverage/additional_698_related/data/ERR398876 |
| **HG00408** | 1000G_2504_high_coverage/additional_698_related/data/ERR398876 |
| **HG00418** | 1000G_2504_high_coverage/additional_698_related/data/ERR398876 |
| **HG00420** | 1000G_2504_high_coverage/additional_698_related/data/ERR398876 |
| **HG00423** | 1000G_2504_high_coverage/additional_698_related/data/ERR398876 |
| **HG00427** | 1000G_2504_high_coverage/additional_698_related/data/ERR398876 |
| **HG00429** | 1000G_2504_high_coverage/additional_698_related/data/ERR398876 |
| **HG00438** | 1000G_2504_high_coverage/additional_698_related/data/ERR398876 |
| **HG00444** | 1000G_2504_high_coverage/additional_698_related/data/ERR398876 |
| **HG00447** | 1000G_2504_high_coverage/additional_698_related/data/ERR398877 |
| **HG00450** | 1000G_2504_high_coverage/additional_698_related/data/ERR398877 |
| **HG00453** | 1000G_2504_high_coverage/additional_698_related/data/ERR398877 |
| **HG00459** | 1000G_2504_high_coverage/additional_698_related/data/ERR398877 |
| **HG00465** | 1000G_2504_high_coverage/additional_698_related/data/ERR398877 |
| **HG00474** | 1000G_2504_high_coverage/additional_698_related/data/ERR398877 |
| **HG00477** | 1000G_2504_high_coverage/additional_698_related/data/ERR398877 |
| **HG00480** | 1000G_2504_high_coverage/additional_698_related/data/ERR398877 |
| **HG00501** | 1000G_2504_high_coverage/additional_698_related/data/ERR398877 |
| **HG00502** | 1000G_2504_high_coverage/additional_698_related/data/ERR398877 |
| **HG00512** | 1000G_2504_high_coverage/additional_698_related/data/ERR398878 |
| **HG00514** | 1000G_2504_high_coverage/additional_698_related/data/ERR398878 |
| **HG00526** | 1000G_2504_high_coverage/additional_698_related/data/ERR398878 |
| **HG00532** | 1000G_2504_high_coverage/additional_698_related/data/ERR398878 |
| **HG00535** | 1000G_2504_high_coverage/additional_698_related/data/ERR398878 |
| **HG00538** | 1000G_2504_high_coverage/additional_698_related/data/ERR398878 |
| **HG00544** | 1000G_2504_high_coverage/additional_698_related/data/ERR398878 |
| **HG00552** | 1000G_2504_high_coverage/additional_698_related/data/ERR398878 |
| **HG00555** | 1000G_2504_high_coverage/additional_698_related/data/ERR398878 |

| | |
|---|---|
| **HG00558** | 1000G_2504_high_coverage/additional_698_related/data/ERR398878 |
| **HG00561** | 1000G_2504_high_coverage/additional_698_related/data/ERR398879 |
| **HG00567** | 1000G_2504_high_coverage/additional_698_related/data/ERR398879 |
| **HG00577** | 1000G_2504_high_coverage/additional_698_related/data/ERR398879 |
| **HG00578** | 1000G_2504_high_coverage/additional_698_related/data/ERR398879 |
| **HG00579** | 1000G_2504_high_coverage/additional_698_related/data/ERR398879 |
| **HG00582** | 1000G_2504_high_coverage/additional_698_related/data/ERR398879 |
| **HG00585** | 1000G_2504_high_coverage/additional_698_related/data/ERR398879 |
| **HG00591** | 1000G_2504_high_coverage/additional_698_related/data/ERR398879 |
| **HG00594** | 1000G_2504_high_coverage/additional_698_related/data/ERR398879 |
| **HG00597** | 1000G_2504_high_coverage/additional_698_related/data/ERR398879 |
| **HG00609** | 1000G_2504_high_coverage/additional_698_related/data/ERR398880 |
| **HG00612** | 1000G_2504_high_coverage/additional_698_related/data/ERR398880 |
| **HG00615** | 1000G_2504_high_coverage/additional_698_related/data/ERR398880 |
| **HG00621** | 1000G_2504_high_coverage/additional_698_related/data/ERR398880 |
| **HG00627** | 1000G_2504_high_coverage/additional_698_related/data/ERR398880 |
| **HG00630** | 1000G_2504_high_coverage/additional_698_related/data/ERR398880 |
| **HG00635** | 1000G_2504_high_coverage/additional_698_related/data/ERR398880 |
| **HG00636** | 1000G_2504_high_coverage/additional_698_related/data/ERR398880 |
| **HG00639** | 1000G_2504_high_coverage/additional_698_related/data/ERR398880 |
| **HG00642** | 1000G_2504_high_coverage/additional_698_related/data/ERR398880 |
| **HG00652** | 1000G_2504_high_coverage/additional_698_related/data/ERR398881 |
| **HG00655** | 1000G_2504_high_coverage/additional_698_related/data/ERR398881 |
| **HG00658** | 1000G_2504_high_coverage/additional_698_related/data/ERR398881 |
| **HG00664** | 1000G_2504_high_coverage/additional_698_related/data/ERR398881 |
| **HG00673** | 1000G_2504_high_coverage/additional_698_related/data/ERR398881 |
| **HG00685** | 1000G_2504_high_coverage/additional_698_related/data/ERR398881 |
| **HG00691** | 1000G_2504_high_coverage/additional_698_related/data/ERR398881 |
| **HG00694** | 1000G_2504_high_coverage/additional_698_related/data/ERR398881 |
| **HG00700** | 1000G_2504_high_coverage/additional_698_related/data/ERR398881 |
| **HG00702** | 1000G_2504_high_coverage/additional_698_related/data/ERR398881 |
| **HG00703** | 1000G_2504_high_coverage/additional_698_related/data/ERR398882 |
| **HG00706** | 1000G_2504_high_coverage/additional_698_related/data/ERR398882 |
| **HG00709** | 1000G_2504_high_coverage/additional_698_related/data/ERR398882 |
| **HG00733** | 1000G_2504_high_coverage/additional_698_related/data/ERR398882 |
| **HG00735** | 1000G_2504_high_coverage/additional_698_related/data/ERR398882 |
| **HG00738** | 1000G_2504_high_coverage/additional_698_related/data/ERR398882 |
| **HG00741** | 1000G_2504_high_coverage/additional_698_related/data/ERR398882 |
| **HG01050** | 1000G_2504_high_coverage/additional_698_related/data/ERR398882 |
| **HG01053** | 1000G_2504_high_coverage/additional_698_related/data/ERR398882 |
| **HG01056** | 1000G_2504_high_coverage/additional_698_related/data/ERR398882 |
| **HG01062** | 1000G_2504_high_coverage/additional_698_related/data/ERR398883 |
| **HG01068** | 1000G_2504_high_coverage/additional_698_related/data/ERR398883 |
| **HG01071** | 1000G_2504_high_coverage/additional_698_related/data/ERR398883 |
| **HG01074** | 1000G_2504_high_coverage/additional_698_related/data/ERR398883 |
| **HG01081** | 1000G_2504_high_coverage/additional_698_related/data/ERR398883 |
| **HG01084** | 1000G_2504_high_coverage/additional_698_related/data/ERR398883 |
| **HG01087** | 1000G_2504_high_coverage/additional_698_related/data/ERR398883 |
| **HG01096** | 1000G_2504_high_coverage/additional_698_related/data/ERR398883 |
| **HG01099** | 1000G_2504_high_coverage/additional_698_related/data/ERR398883 |
| **HG01100** | 1000G_2504_high_coverage/additional_698_related/data/ERR398883 |
| **HG01103** | 1000G_2504_high_coverage/additional_698_related/data/ERR398884 |
| **HG01106** | 1000G_2504_high_coverage/additional_698_related/data/ERR398884 |
| **HG01109** | 1000G_2504_high_coverage/additional_698_related/data/ERR398884 |
| **HG01114** | 1000G_2504_high_coverage/additional_698_related/data/ERR398884 |
| **HG01126** | 1000G_2504_high_coverage/additional_698_related/data/ERR398884 |
| **HG01135** | 1000G_2504_high_coverage/additional_698_related/data/ERR398884 |
| **HG01138** | 1000G_2504_high_coverage/additional_698_related/data/ERR398884 |
| **HG01141** | 1000G_2504_high_coverage/additional_698_related/data/ERR398884 |
| **HG01150** | 1000G_2504_high_coverage/additional_698_related/data/ERR398884 |
| **HG01169** | 1000G_2504_high_coverage/additional_698_related/data/ERR398884 |
| **HG01172** | 1000G_2504_high_coverage/additional_698_related/data/ERR398885 |
| **HG01175** | 1000G_2504_high_coverage/additional_698_related/data/ERR398885 |
| **HG01178** | 1000G_2504_high_coverage/additional_698_related/data/ERR398885 |
| **HG01184** | 1000G_2504_high_coverage/additional_698_related/data/ERR398885 |

| | |
|---|---|
| **HG01189** | 1000G_2504_high_coverage/additional_698_related/data/ERR398885 |
| **HG01192** | 1000G_2504_high_coverage/additional_698_related/data/ERR398885 |
| **HG01199** | 1000G_2504_high_coverage/additional_698_related/data/ERR398885 |
| **HG01206** | 1000G_2504_high_coverage/additional_698_related/data/ERR398885 |
| **HG01243** | 1000G_2504_high_coverage/additional_698_related/data/ERR398885 |
| **HG01249** | 1000G_2504_high_coverage/additional_698_related/data/ERR398885 |
| **HG01252** | 1000G_2504_high_coverage/additional_698_related/data/ERR398886 |
| **HG01255** | 1000G_2504_high_coverage/additional_698_related/data/ERR398886 |
| **HG01258** | 1000G_2504_high_coverage/additional_698_related/data/ERR398886 |
| **HG01261** | 1000G_2504_high_coverage/additional_698_related/data/ERR398886 |
| **HG01273** | 1000G_2504_high_coverage/additional_698_related/data/ERR398886 |
| **HG01274** | 1000G_2504_high_coverage/additional_698_related/data/ERR398886 |
| **HG01276** | 1000G_2504_high_coverage/additional_698_related/data/ERR398886 |
| **HG01278** | 1000G_2504_high_coverage/additional_698_related/data/ERR398886 |
| **HG01279** | 1000G_2504_high_coverage/additional_698_related/data/ERR398886 |
| **HG01343** | 1000G_2504_high_coverage/additional_698_related/data/ERR398886 |
| **HG01346** | 1000G_2504_high_coverage/additional_698_related/data/ERR398887 |
| **HG01347** | 1000G_2504_high_coverage/additional_698_related/data/ERR398887 |
| **HG01349** | 1000G_2504_high_coverage/additional_698_related/data/ERR398887 |
| **HG01352** | 1000G_2504_high_coverage/additional_698_related/data/ERR398887 |
| **HG01355** | 1000G_2504_high_coverage/additional_698_related/data/ERR398887 |
| **HG01358** | 1000G_2504_high_coverage/additional_698_related/data/ERR398887 |
| **HG01361** | 1000G_2504_high_coverage/additional_698_related/data/ERR398887 |
| **HG01367** | 1000G_2504_high_coverage/additional_698_related/data/ERR398887 |
| **HG01376** | 1000G_2504_high_coverage/additional_698_related/data/ERR398887 |
| **HG01379** | 1000G_2504_high_coverage/additional_698_related/data/ERR398887 |
| **HG01385** | 1000G_2504_high_coverage/additional_698_related/data/ERR398888 |
| **HG01391** | 1000G_2504_high_coverage/additional_698_related/data/ERR398888 |
| **HG01433** | 1000G_2504_high_coverage/additional_698_related/data/ERR398888 |
| **HG01439** | 1000G_2504_high_coverage/additional_698_related/data/ERR398888 |
| **HG01457** | 1000G_2504_high_coverage/additional_698_related/data/ERR398888 |
| **HG01463** | 1000G_2504_high_coverage/additional_698_related/data/ERR398888 |
| **HG01466** | 1000G_2504_high_coverage/additional_698_related/data/ERR398888 |
| **HG01490** | 1000G_2504_high_coverage/additional_698_related/data/ERR398888 |
| **HG01493** | 1000G_2504_high_coverage/additional_698_related/data/ERR398888 |
| **HG01496** | 1000G_2504_high_coverage/additional_698_related/data/ERR398888 |
| **HG01499** | 1000G_2504_high_coverage/additional_698_related/data/ERR398889 |
| **HG01502** | 1000G_2504_high_coverage/additional_698_related/data/ERR398889 |
| **HG01505** | 1000G_2504_high_coverage/additional_698_related/data/ERR398889 |
| **HG01508** | 1000G_2504_high_coverage/additional_698_related/data/ERR398889 |
| **HG01511** | 1000G_2504_high_coverage/additional_698_related/data/ERR398889 |
| **HG01514** | 1000G_2504_high_coverage/additional_698_related/data/ERR398889 |
| **HG01517** | 1000G_2504_high_coverage/additional_698_related/data/ERR398889 |
| **HG01520** | 1000G_2504_high_coverage/additional_698_related/data/ERR398889 |
| **HG01523** | 1000G_2504_high_coverage/additional_698_related/data/ERR398889 |
| **HG01526** | 1000G_2504_high_coverage/additional_698_related/data/ERR398889 |
| **HG01529** | 1000G_2504_high_coverage/additional_698_related/data/ERR398890 |
| **HG01532** | 1000G_2504_high_coverage/additional_698_related/data/ERR398890 |
| **HG01538** | 1000G_2504_high_coverage/additional_698_related/data/ERR398890 |
| **HG01552** | 1000G_2504_high_coverage/additional_698_related/data/ERR398890 |
| **HG01567** | 1000G_2504_high_coverage/additional_698_related/data/ERR398890 |
| **HG01573** | 1000G_2504_high_coverage/additional_698_related/data/ERR398890 |
| **HG01579** | 1000G_2504_high_coverage/additional_698_related/data/ERR398890 |
| **HG01601** | 1000G_2504_high_coverage/additional_698_related/data/ERR398890 |
| **HG01604** | 1000G_2504_high_coverage/additional_698_related/data/ERR398890 |
| **HG01609** | 1000G_2504_high_coverage/additional_698_related/data/ERR398890 |
| **HG01611** | 1000G_2504_high_coverage/additional_698_related/data/ERR398891 |
| **HG01614** | 1000G_2504_high_coverage/additional_698_related/data/ERR398891 |
| **HG01616** | 1000G_2504_high_coverage/additional_698_related/data/ERR398891 |
| **HG01621** | 1000G_2504_high_coverage/additional_698_related/data/ERR398891 |
| **HG01622** | 1000G_2504_high_coverage/additional_698_related/data/ERR398891 |
| **HG01627** | 1000G_2504_high_coverage/additional_698_related/data/ERR398891 |
| **HG01629** | 1000G_2504_high_coverage/additional_698_related/data/ERR398891 |
| **HG01633** | 1000G_2504_high_coverage/additional_698_related/data/ERR398891 |
| **HG01667** | 1000G_2504_high_coverage/additional_698_related/data/ERR398891 |

| | |
|---|---|
| **HG01671** | 1000G_2504_high_coverage/additional_698_related/data/ERR398891 |
| **HG01674** | 1000G_2504_high_coverage/additional_698_related/data/ERR398892 |
| **HG01677** | 1000G_2504_high_coverage/additional_698_related/data/ERR398892 |
| **HG01681** | 1000G_2504_high_coverage/additional_698_related/data/ERR398892 |
| **HG01683** | 1000G_2504_high_coverage/additional_698_related/data/ERR398892 |
| **HG01687** | 1000G_2504_high_coverage/additional_698_related/data/ERR398892 |
| **HG01696** | 1000G_2504_high_coverage/additional_698_related/data/ERR398892 |
| **HG01698** | 1000G_2504_high_coverage/additional_698_related/data/ERR398892 |
| **HG01701** | 1000G_2504_high_coverage/additional_698_related/data/ERR398892 |
| **HG01703** | 1000G_2504_high_coverage/additional_698_related/data/ERR398892 |
| **HG01706** | 1000G_2504_high_coverage/additional_698_related/data/ERR398892 |
| **HG01711** | 1000G_2504_high_coverage/additional_698_related/data/ERR398893 |
| **HG01748** | 1000G_2504_high_coverage/additional_698_related/data/ERR398893 |
| **HG01755** | 1000G_2504_high_coverage/additional_698_related/data/ERR398893 |
| **HG01763** | 1000G_2504_high_coverage/additional_698_related/data/ERR398893 |
| **HG01772** | 1000G_2504_high_coverage/additional_698_related/data/ERR398893 |
| **HG01774** | 1000G_2504_high_coverage/additional_698_related/data/ERR398893 |
| **HG01778** | 1000G_2504_high_coverage/additional_698_related/data/ERR398893 |
| **HG01780** | 1000G_2504_high_coverage/additional_698_related/data/ERR398893 |
| **HG01782** | 1000G_2504_high_coverage/additional_698_related/data/ERR398893 |
| **HG01881** | 1000G_2504_high_coverage/additional_698_related/data/ERR398893 |
| **HG01884** | 1000G_2504_high_coverage/additional_698_related/data/ERR398894 |
| **HG01887** | 1000G_2504_high_coverage/additional_698_related/data/ERR398894 |
| **HG01888** | 1000G_2504_high_coverage/additional_698_related/data/ERR398894 |
| **HG01891** | 1000G_2504_high_coverage/additional_698_related/data/ERR398894 |
| **HG01895** | 1000G_2504_high_coverage/additional_698_related/data/ERR398894 |
| **HG01897** | 1000G_2504_high_coverage/additional_698_related/data/ERR398894 |
| **HG01898** | 1000G_2504_high_coverage/additional_698_related/data/ERR398894 |
| **HG01916** | 1000G_2504_high_coverage/additional_698_related/data/ERR398894 |
| **HG01919** | 1000G_2504_high_coverage/additional_698_related/data/ERR398894 |
| **HG01922** | 1000G_2504_high_coverage/additional_698_related/data/ERR398894 |
| **HG01925** | 1000G_2504_high_coverage/additional_698_related/data/ERR398895 |
| **HG01928** | 1000G_2504_high_coverage/additional_698_related/data/ERR398895 |
| **HG01934** | 1000G_2504_high_coverage/additional_698_related/data/ERR398895 |
| **HG01937** | 1000G_2504_high_coverage/additional_698_related/data/ERR398895 |
| **HG01940** | 1000G_2504_high_coverage/additional_698_related/data/ERR398895 |
| **HG01943** | 1000G_2504_high_coverage/additional_698_related/data/ERR398895 |
| **HG01946** | 1000G_2504_high_coverage/additional_698_related/data/ERR398895 |
| **HG01949** | 1000G_2504_high_coverage/additional_698_related/data/ERR398895 |
| **HG01952** | 1000G_2504_high_coverage/additional_698_related/data/ERR398895 |
| **HG01955** | 1000G_2504_high_coverage/additional_698_related/data/ERR398895 |
| **HG01959** | 1000G_2504_high_coverage/additional_698_related/data/ERR398896 |
| **HG01960** | 1000G_2504_high_coverage/additional_698_related/data/ERR398896 |
| **HG01969** | 1000G_2504_high_coverage/additional_698_related/data/ERR398896 |
| **HG01972** | 1000G_2504_high_coverage/additional_698_related/data/ERR398896 |
| **HG01975** | 1000G_2504_high_coverage/additional_698_related/data/ERR398896 |
| **HG01978** | 1000G_2504_high_coverage/additional_698_related/data/ERR398896 |
| **HG01981** | 1000G_2504_high_coverage/additional_698_related/data/ERR398896 |
| **HG01983** | 1000G_2504_high_coverage/additional_698_related/data/ERR398896 |
| **HG01984** | 1000G_2504_high_coverage/additional_698_related/data/ERR398896 |
| **HG01987** | 1000G_2504_high_coverage/additional_698_related/data/ERR398896 |
| **HG01993** | 1000G_2504_high_coverage/additional_698_related/data/ERR398897 |
| **HG01998** | 1000G_2504_high_coverage/additional_698_related/data/ERR398897 |
| **HG02004** | 1000G_2504_high_coverage/additional_698_related/data/ERR398897 |
| **HG02011** | 1000G_2504_high_coverage/additional_698_related/data/ERR398897 |
| **HG02015** | 1000G_2504_high_coverage/additional_698_related/data/ERR398897 |
| **HG02018** | 1000G_2504_high_coverage/additional_698_related/data/ERR398897 |
| **HG02024** | 1000G_2504_high_coverage/additional_698_related/data/ERR398897 |
| **HG02027** | 1000G_2504_high_coverage/additional_698_related/data/ERR398897 |
| **HG02030** | 1000G_2504_high_coverage/additional_698_related/data/ERR398897 |
| **HG02055** | 1000G_2504_high_coverage/additional_698_related/data/ERR398897 |
| **HG02056** | 1000G_2504_high_coverage/additional_698_related/data/ERR398898 |
| **HG02059** | 1000G_2504_high_coverage/additional_698_related/data/ERR398898 |
| **HG02068** | 1000G_2504_high_coverage/additional_698_related/data/ERR398898 |
| **HG02071** | 1000G_2504_high_coverage/additional_698_related/data/ERR398898 |

| | |
|---|---|
| **HG02074** | 1000G_2504_high_coverage/additional_698_related/data/ERR398898 |
| **HG02077** | 1000G_2504_high_coverage/additional_698_related/data/ERR398898 |
| **HG02080** | 1000G_2504_high_coverage/additional_698_related/data/ERR398898 |
| **HG02083** | 1000G_2504_high_coverage/additional_698_related/data/ERR398898 |
| **HG02091** | 1000G_2504_high_coverage/additional_698_related/data/ERR398898 |
| **HG02106** | 1000G_2504_high_coverage/additional_698_related/data/ERR398898 |
| **HG02120** | 1000G_2504_high_coverage/additional_698_related/data/ERR398899 |
| **HG02126** | 1000G_2504_high_coverage/additional_698_related/data/ERR398899 |
| **HG02129** | 1000G_2504_high_coverage/additional_698_related/data/ERR398899 |
| **HG02132** | 1000G_2504_high_coverage/additional_698_related/data/ERR398899 |
| **HG02135** | 1000G_2504_high_coverage/additional_698_related/data/ERR398899 |
| **HG02145** | 1000G_2504_high_coverage/additional_698_related/data/ERR398899 |
| **HG02148** | 1000G_2504_high_coverage/additional_698_related/data/ERR398899 |
| **HG02222** | 1000G_2504_high_coverage/additional_698_related/data/ERR398899 |
| **HG02225** | 1000G_2504_high_coverage/additional_698_related/data/ERR398899 |
| **HG02229** | 1000G_2504_high_coverage/additional_698_related/data/ERR398899 |
| **HG02234** | 1000G_2504_high_coverage/additional_698_related/data/ERR398900 |
| **HG02237** | 1000G_2504_high_coverage/additional_698_related/data/ERR398900 |
| **HG02240** | 1000G_2504_high_coverage/additional_698_related/data/ERR398900 |
| **HG02257** | 1000G_2504_high_coverage/additional_698_related/data/ERR398900 |
| **HG02258** | 1000G_2504_high_coverage/additional_698_related/data/ERR398900 |
| **HG02261** | 1000G_2504_high_coverage/additional_698_related/data/ERR398900 |
| **HG02273** | 1000G_2504_high_coverage/additional_698_related/data/ERR398900 |
| **HG02279** | 1000G_2504_high_coverage/additional_698_related/data/ERR398900 |
| **HG02280** | 1000G_2504_high_coverage/additional_698_related/data/ERR398900 |
| **HG02287** | 1000G_2504_high_coverage/additional_698_related/data/ERR398900 |
| **HG02293** | 1000G_2504_high_coverage/additional_698_related/data/ERR398901 |
| **HG02300** | 1000G_2504_high_coverage/additional_698_related/data/ERR398901 |
| **HG02302** | 1000G_2504_high_coverage/additional_698_related/data/ERR398901 |
| **HG02303** | 1000G_2504_high_coverage/additional_698_related/data/ERR398901 |
| **HG02316** | 1000G_2504_high_coverage/additional_698_related/data/ERR398901 |
| **HG02321** | 1000G_2504_high_coverage/additional_698_related/data/ERR398901 |
| **HG02451** | 1000G_2504_high_coverage/additional_698_related/data/ERR398901 |
| **HG02463** | 1000G_2504_high_coverage/additional_698_related/data/ERR398901 |
| **HG02466** | 1000G_2504_high_coverage/additional_698_related/data/ERR398901 |
| **HG02492** | 1000G_2504_high_coverage/additional_698_related/data/ERR398901 |
| **HG02495** | 1000G_2504_high_coverage/additional_698_related/data/ERR398902 |
| **HG02514** | 1000G_2504_high_coverage/additional_698_related/data/ERR398902 |
| **HG02523** | 1000G_2504_high_coverage/additional_698_related/data/ERR398902 |
| **HG02524** | 1000G_2504_high_coverage/additional_698_related/data/ERR398902 |
| **HG02525** | 1000G_2504_high_coverage/additional_698_related/data/ERR398902 |
| **HG02526** | 1000G_2504_high_coverage/additional_698_related/data/ERR398902 |
| **HG02563** | 1000G_2504_high_coverage/additional_698_related/data/ERR398902 |
| **HG02569** | 1000G_2504_high_coverage/additional_698_related/data/ERR398902 |
| **HG02572** | 1000G_2504_high_coverage/additional_698_related/data/ERR398902 |
| **HG02575** | 1000G_2504_high_coverage/additional_698_related/data/ERR398902 |
| **HG02584** | 1000G_2504_high_coverage/additional_698_related/data/ERR398903 |
| **HG02587** | 1000G_2504_high_coverage/additional_698_related/data/ERR398903 |
| **HG02590** | 1000G_2504_high_coverage/additional_698_related/data/ERR398903 |
| **HG02596** | 1000G_2504_high_coverage/additional_698_related/data/ERR398903 |
| **HG02602** | 1000G_2504_high_coverage/additional_698_related/data/ERR398903 |
| **HG02605** | 1000G_2504_high_coverage/additional_698_related/data/ERR398903 |
| **HG02612** | 1000G_2504_high_coverage/additional_698_related/data/ERR398903 |
| **HG02615** | 1000G_2504_high_coverage/additional_698_related/data/ERR398903 |
| **HG02622** | 1000G_2504_high_coverage/additional_698_related/data/ERR398903 |
| **HG02625** | 1000G_2504_high_coverage/additional_698_related/data/ERR398903 |
| **HG02630** | 1000G_2504_high_coverage/additional_698_related/data/ERR398904 |
| **HG02636** | 1000G_2504_high_coverage/additional_698_related/data/ERR398904 |
| **HG02644** | 1000G_2504_high_coverage/additional_698_related/data/ERR398904 |
| **HG02647** | 1000G_2504_high_coverage/additional_698_related/data/ERR398904 |
| **HG02650** | 1000G_2504_high_coverage/additional_698_related/data/ERR398904 |
| **HG02653** | 1000G_2504_high_coverage/additional_698_related/data/ERR398904 |
| **HG02656** | 1000G_2504_high_coverage/additional_698_related/data/ERR398904 |
| **HG02659** | 1000G_2504_high_coverage/additional_698_related/data/ERR398904 |
| **HG02662** | 1000G_2504_high_coverage/additional_698_related/data/ERR398904 |

| | |
|---|---|
| **HG02668** | 1000G_2504_high_coverage/additional_698_related/data/ERR398904 |
| **HG02677** | 1000G_2504_high_coverage/additional_698_related/data/ERR398905 |
| **HG02680** | 1000G_2504_high_coverage/additional_698_related/data/ERR398905 |
| **HG02683** | 1000G_2504_high_coverage/additional_698_related/data/ERR398905 |
| **HG02686** | 1000G_2504_high_coverage/additional_698_related/data/ERR398905 |
| **HG02689** | 1000G_2504_high_coverage/additional_698_related/data/ERR398905 |
| **HG02692** | 1000G_2504_high_coverage/additional_698_related/data/ERR398905 |
| **HG02698** | 1000G_2504_high_coverage/additional_698_related/data/ERR398905 |
| **HG02701** | 1000G_2504_high_coverage/additional_698_related/data/ERR398905 |
| **HG02704** | 1000G_2504_high_coverage/additional_698_related/data/ERR398905 |
| **HG02717** | 1000G_2504_high_coverage/additional_698_related/data/ERR398905 |
| **HG02723** | 1000G_2504_high_coverage/additional_698_related/data/ERR398906 |
| **HG02726** | 1000G_2504_high_coverage/additional_698_related/data/ERR398906 |
| **HG02729** | 1000G_2504_high_coverage/additional_698_related/data/ERR398906 |
| **HG02735** | 1000G_2504_high_coverage/additional_698_related/data/ERR398906 |
| **HG02738** | 1000G_2504_high_coverage/additional_698_related/data/ERR398906 |
| **HG02761** | 1000G_2504_high_coverage/additional_698_related/data/ERR398906 |
| **HG02762** | 1000G_2504_high_coverage/additional_698_related/data/ERR398906 |
| **HG02764** | 1000G_2504_high_coverage/additional_698_related/data/ERR398906 |
| **HG02770** | 1000G_2504_high_coverage/additional_698_related/data/ERR398906 |
| **HG02773** | 1000G_2504_high_coverage/additional_698_related/data/ERR398906 |
| **HG02776** | 1000G_2504_high_coverage/additional_698_related/data/ERR398907 |
| **HG02785** | 1000G_2504_high_coverage/additional_698_related/data/ERR398907 |
| **HG02788** | 1000G_2504_high_coverage/additional_698_related/data/ERR398907 |
| **HG02791** | 1000G_2504_high_coverage/additional_698_related/data/ERR398907 |
| **HG02794** | 1000G_2504_high_coverage/additional_698_related/data/ERR398907 |
| **HG02800** | 1000G_2504_high_coverage/additional_698_related/data/ERR398907 |
| **HG02806** | 1000G_2504_high_coverage/additional_698_related/data/ERR398907 |
| **HG02809** | 1000G_2504_high_coverage/additional_698_related/data/ERR398907 |
| **HG02812** | 1000G_2504_high_coverage/additional_698_related/data/ERR398907 |
| **HG02815** | 1000G_2504_high_coverage/additional_698_related/data/ERR398907 |
| **HG02818** | 1000G_2504_high_coverage/additional_698_related/data/ERR398908 |
| **HG02821** | 1000G_2504_high_coverage/additional_698_related/data/ERR398908 |
| **HG02838** | 1000G_2504_high_coverage/additional_698_related/data/ERR398908 |
| **HG02841** | 1000G_2504_high_coverage/additional_698_related/data/ERR398908 |
| **HG02853** | 1000G_2504_high_coverage/additional_698_related/data/ERR398908 |
| **HG02856** | 1000G_2504_high_coverage/additional_698_related/data/ERR398908 |
| **HG02862** | 1000G_2504_high_coverage/additional_698_related/data/ERR398908 |
| **HG02869** | 1000G_2504_high_coverage/additional_698_related/data/ERR398908 |
| **HG02871** | 1000G_2504_high_coverage/additional_698_related/data/ERR398908 |
| **HG02880** | 1000G_2504_high_coverage/additional_698_related/data/ERR398908 |
| **HG02883** | 1000G_2504_high_coverage/additional_698_related/data/ERR398909 |
| **HG02886** | 1000G_2504_high_coverage/additional_698_related/data/ERR398909 |
| **HG02889** | 1000G_2504_high_coverage/additional_698_related/data/ERR398909 |
| **HG02892** | 1000G_2504_high_coverage/additional_698_related/data/ERR398909 |
| **HG02897** | 1000G_2504_high_coverage/additional_698_related/data/ERR398909 |
| **HG02924** | 1000G_2504_high_coverage/additional_698_related/data/ERR398909 |
| **HG02945** | 1000G_2504_high_coverage/additional_698_related/data/ERR398909 |
| **HG02948** | 1000G_2504_high_coverage/additional_698_related/data/ERR398909 |
| **HG02954** | 1000G_2504_high_coverage/additional_698_related/data/ERR398909 |
| **HG02964** | 1000G_2504_high_coverage/additional_698_related/data/ERR398909 |
| **HG02965** | 1000G_2504_high_coverage/additional_698_related/data/ERR398910 |
| **HG02966** | 1000G_2504_high_coverage/additional_698_related/data/ERR398910 |
| **HG02972** | 1000G_2504_high_coverage/additional_698_related/data/ERR398910 |
| **HG02975** | 1000G_2504_high_coverage/additional_698_related/data/ERR398910 |
| **HG02978** | 1000G_2504_high_coverage/additional_698_related/data/ERR398910 |
| **HG02980** | 1000G_2504_high_coverage/additional_698_related/data/ERR398910 |
| **HG02984** | 1000G_2504_high_coverage/additional_698_related/data/ERR398910 |
| **HG03008** | 1000G_2504_high_coverage/additional_698_related/data/ERR398910 |
| **HG03017** | 1000G_2504_high_coverage/additional_698_related/data/ERR398910 |
| **HG03026** | 1000G_2504_high_coverage/additional_698_related/data/ERR398910 |
| **HG03029** | 1000G_2504_high_coverage/additional_698_related/data/ERR398911 |
| **HG03033** | 1000G_2504_high_coverage/additional_698_related/data/ERR398911 |
| **HG03034** | 1000G_2504_high_coverage/additional_698_related/data/ERR398911 |
| **HG03035** | 1000G_2504_high_coverage/additional_698_related/data/ERR398911 |

| | |
|---|---|
| **HG03041** | 1000G_2504_high_coverage/additional_698_related/data/ERR398911 |
| **HG03047** | 1000G_2504_high_coverage/additional_698_related/data/ERR398911 |
| **HG03050** | 1000G_2504_high_coverage/additional_698_related/data/ERR398911 |
| **HG03056** | 1000G_2504_high_coverage/additional_698_related/data/ERR398911 |
| **HG03065** | 1000G_2504_high_coverage/additional_698_related/data/ERR398911 |
| **HG03098** | 1000G_2504_high_coverage/additional_698_related/data/ERR398911 |
| **HG03101** | 1000G_2504_high_coverage/additional_698_related/data/ERR398912 |
| **HG03110** | 1000G_2504_high_coverage/additional_698_related/data/ERR398912 |
| **HG03113** | 1000G_2504_high_coverage/additional_698_related/data/ERR398912 |
| **HG03116** | 1000G_2504_high_coverage/additional_698_related/data/ERR398912 |
| **HG03119** | 1000G_2504_high_coverage/additional_698_related/data/ERR398912 |
| **HG03122** | 1000G_2504_high_coverage/additional_698_related/data/ERR398912 |
| **HG03125** | 1000G_2504_high_coverage/additional_698_related/data/ERR398912 |
| **HG03128** | 1000G_2504_high_coverage/additional_698_related/data/ERR398912 |
| **HG03131** | 1000G_2504_high_coverage/additional_698_related/data/ERR398912 |
| **HG03134** | 1000G_2504_high_coverage/additional_698_related/data/ERR398912 |
| **HG03137** | 1000G_2504_high_coverage/additional_698_related/data/ERR398913 |
| **HG03161** | 1000G_2504_high_coverage/additional_698_related/data/ERR398913 |
| **HG03164** | 1000G_2504_high_coverage/additional_698_related/data/ERR398913 |
| **HG03170** | 1000G_2504_high_coverage/additional_698_related/data/ERR398913 |
| **HG03191** | 1000G_2504_high_coverage/additional_698_related/data/ERR398913 |
| **HG03197** | 1000G_2504_high_coverage/additional_698_related/data/ERR398913 |
| **HG03200** | 1000G_2504_high_coverage/additional_698_related/data/ERR398913 |
| **HG03230** | 1000G_2504_high_coverage/additional_698_related/data/ERR398913 |
| **HG03236** | 1000G_2504_high_coverage/additional_698_related/data/ERR398913 |
| **HG03239** | 1000G_2504_high_coverage/additional_698_related/data/ERR398913 |
| **HG03242** | 1000G_2504_high_coverage/additional_698_related/data/ERR398914 |
| **HG03248** | 1000G_2504_high_coverage/additional_698_related/data/ERR398914 |
| **HG03249** | 1000G_2504_high_coverage/additional_698_related/data/ERR398914 |
| **HG03250** | 1000G_2504_high_coverage/additional_698_related/data/ERR398914 |
| **HG03251** | 1000G_2504_high_coverage/additional_698_related/data/ERR398914 |
| **HG03260** | 1000G_2504_high_coverage/additional_698_related/data/ERR398914 |
| **HG03269** | 1000G_2504_high_coverage/additional_698_related/data/ERR398914 |
| **HG03272** | 1000G_2504_high_coverage/additional_698_related/data/ERR398914 |
| **HG03296** | 1000G_2504_high_coverage/additional_698_related/data/ERR398914 |
| **HG03299** | 1000G_2504_high_coverage/additional_698_related/data/ERR398914 |
| **HG03302** | 1000G_2504_high_coverage/additional_698_related/data/ERR398915 |
| **HG03305** | 1000G_2504_high_coverage/additional_698_related/data/ERR398915 |
| **HG03306** | 1000G_2504_high_coverage/additional_698_related/data/ERR398915 |
| **HG03307** | 1000G_2504_high_coverage/additional_698_related/data/ERR398915 |
| **HG03308** | 1000G_2504_high_coverage/additional_698_related/data/ERR398915 |
| **HG03309** | 1000G_2504_high_coverage/additional_698_related/data/ERR398915 |
| **HG03310** | 1000G_2504_high_coverage/additional_698_related/data/ERR398915 |
| **HG03312** | 1000G_2504_high_coverage/additional_698_related/data/ERR398915 |
| **HG03314** | 1000G_2504_high_coverage/additional_698_related/data/ERR398915 |
| **HG03344** | 1000G_2504_high_coverage/additional_698_related/data/ERR398915 |
| **HG03350** | 1000G_2504_high_coverage/additional_698_related/data/ERR398916 |
| **HG03368** | 1000G_2504_high_coverage/additional_698_related/data/ERR398916 |
| **HG03371** | 1000G_2504_high_coverage/additional_698_related/data/ERR398916 |
| **HG03373** | 1000G_2504_high_coverage/additional_698_related/data/ERR398916 |
| **HG03374** | 1000G_2504_high_coverage/additional_698_related/data/ERR398916 |
| **HG03438** | 1000G_2504_high_coverage/additional_698_related/data/ERR398916 |
| **HG03453** | 1000G_2504_high_coverage/additional_698_related/data/ERR398916 |
| **HG03454** | 1000G_2504_high_coverage/additional_698_related/data/ERR398916 |
| **HG03456** | 1000G_2504_high_coverage/additional_698_related/data/ERR398916 |
| **HG03480** | 1000G_2504_high_coverage/additional_698_related/data/ERR398916 |
| **HG03486** | 1000G_2504_high_coverage/additional_698_related/data/ERR398917 |
| **HG03487** | 1000G_2504_high_coverage/additional_698_related/data/ERR398917 |
| **HG03489** | 1000G_2504_high_coverage/additional_698_related/data/ERR398917 |
| **HG03492** | 1000G_2504_high_coverage/additional_698_related/data/ERR398917 |
| **HG03516** | 1000G_2504_high_coverage/additional_698_related/data/ERR398917 |
| **HG03519** | 1000G_2504_high_coverage/additional_698_related/data/ERR398917 |
| **HG03522** | 1000G_2504_high_coverage/additional_698_related/data/ERR398917 |
| **HG03540** | 1000G_2504_high_coverage/additional_698_related/data/ERR398917 |
| **HG03574** | 1000G_2504_high_coverage/additional_698_related/data/ERR398917 |

| | |
|---|---|
| **HG03576** | 1000G_2504_high_coverage/additional_698_related/data/ERR398917 |
| **HG03579** | 1000G_2504_high_coverage/additional_698_related/data/ERR398918 |
| **HG03582** | 1000G_2504_high_coverage/additional_698_related/data/ERR398918 |
| **HG03584** | 1000G_2504_high_coverage/additional_698_related/data/ERR398918 |
| **HG03596** | 1000G_2504_high_coverage/additional_698_related/data/ERR398918 |
| **HG03605** | 1000G_2504_high_coverage/additional_698_related/data/ERR398918 |
| **HG03606** | 1000G_2504_high_coverage/additional_698_related/data/ERR398918 |
| **HG03617** | 1000G_2504_high_coverage/additional_698_related/data/ERR398918 |
| **HG03618** | 1000G_2504_high_coverage/additional_698_related/data/ERR398918 |
| **HG03620** | 1000G_2504_high_coverage/additional_698_related/data/ERR398918 |
| **HG03626** | 1000G_2504_high_coverage/additional_698_related/data/ERR398918 |
| **HG03633** | 1000G_2504_high_coverage/additional_698_related/data/ERR398919 |
| **HG03635** | 1000G_2504_high_coverage/additional_698_related/data/ERR398919 |
| **HG03639** | 1000G_2504_high_coverage/additional_698_related/data/ERR398919 |
| **HG03641** | 1000G_2504_high_coverage/additional_698_related/data/ERR398919 |
| **HG03650** | 1000G_2504_high_coverage/additional_698_related/data/ERR398919 |
| **HG03651** | 1000G_2504_high_coverage/additional_698_related/data/ERR398919 |
| **HG03654** | 1000G_2504_high_coverage/additional_698_related/data/ERR398919 |
| **HG03669** | 1000G_2504_high_coverage/additional_698_related/data/ERR398919 |
| **HG03682** | 1000G_2504_high_coverage/additional_698_related/data/ERR398919 |
| **HG03683** | 1000G_2504_high_coverage/additional_698_related/data/ERR398919 |
| **HG03688** | 1000G_2504_high_coverage/additional_698_related/data/ERR398920 |
| **HG03699** | 1000G_2504_high_coverage/additional_698_related/data/ERR398920 |
| **HG03700** | 1000G_2504_high_coverage/additional_698_related/data/ERR398920 |
| **HG03701** | 1000G_2504_high_coverage/additional_698_related/data/ERR398920 |
| **HG03704** | 1000G_2504_high_coverage/additional_698_related/data/ERR398920 |
| **HG03707** | 1000G_2504_high_coverage/additional_698_related/data/ERR398920 |
| **HG03710** | 1000G_2504_high_coverage/additional_698_related/data/ERR398920 |
| **HG03719** | 1000G_2504_high_coverage/additional_698_related/data/ERR398920 |
| **HG03721** | 1000G_2504_high_coverage/additional_698_related/data/ERR398920 |
| **HG03725** | 1000G_2504_high_coverage/additional_698_related/data/ERR398920 |
| **HG03732** | 1000G_2504_high_coverage/additional_698_related/data/ERR398921 |
| **HG03761** | 1000G_2504_high_coverage/additional_698_related/data/ERR398921 |
| **HG03763** | 1000G_2504_high_coverage/additional_698_related/data/ERR398921 |
| **HG03794** | 1000G_2504_high_coverage/additional_698_related/data/ERR398921 |
| **HG03795** | 1000G_2504_high_coverage/additional_698_related/data/ERR398921 |
| **HG03797** | 1000G_2504_high_coverage/additional_698_related/data/ERR398921 |
| **HG03798** | 1000G_2504_high_coverage/additional_698_related/data/ERR398921 |
| **HG03799** | 1000G_2504_high_coverage/additional_698_related/data/ERR398921 |
| **HG03801** | 1000G_2504_high_coverage/additional_698_related/data/ERR398921 |
| **HG03804** | 1000G_2504_high_coverage/additional_698_related/data/ERR398921 |
| **HG03806** | 1000G_2504_high_coverage/additional_698_related/data/ERR398922 |
| **HG03807** | 1000G_2504_high_coverage/additional_698_related/data/ERR398922 |
| **HG03816** | 1000G_2504_high_coverage/additional_698_related/data/ERR398922 |
| **HG03825** | 1000G_2504_high_coverage/additional_698_related/data/ERR398922 |
| **HG03831** | 1000G_2504_high_coverage/additional_698_related/data/ERR398922 |
| **HG03834** | 1000G_2504_high_coverage/additional_698_related/data/ERR398922 |
| **HG03904** | 1000G_2504_high_coverage/additional_698_related/data/ERR398922 |
| **HG03906** | 1000G_2504_high_coverage/additional_698_related/data/ERR398922 |
| **HG03909** | 1000G_2504_high_coverage/additional_698_related/data/ERR398922 |
| **HG03915** | 1000G_2504_high_coverage/additional_698_related/data/ERR398922 |
| **HG03927** | 1000G_2504_high_coverage/additional_698_related/data/ERR398923 |
| **HG03929** | 1000G_2504_high_coverage/additional_698_related/data/ERR398923 |
| **HG03930** | 1000G_2504_high_coverage/additional_698_related/data/ERR398923 |
| **HG03942** | 1000G_2504_high_coverage/additional_698_related/data/ERR398923 |
| **HG03992** | 1000G_2504_high_coverage/additional_698_related/data/ERR398923 |
| **HG04036** | 1000G_2504_high_coverage/additional_698_related/data/ERR398923 |
| **HG04115** | 1000G_2504_high_coverage/additional_698_related/data/ERR398923 |
| **HG04122** | 1000G_2504_high_coverage/additional_698_related/data/ERR398923 |
| **HG04127** | 1000G_2504_high_coverage/additional_698_related/data/ERR398923 |
| **HG04132** | 1000G_2504_high_coverage/additional_698_related/data/ERR398923 |
| **HG04133** | 1000G_2504_high_coverage/additional_698_related/data/ERR398924 |
| **HG04135** | 1000G_2504_high_coverage/additional_698_related/data/ERR398924 |
| **HG04136** | 1000G_2504_high_coverage/additional_698_related/data/ERR398924 |
| **HG04142** | 1000G_2504_high_coverage/additional_698_related/data/ERR398924 |

| | |
|---|---|
| **HG04147** | 1000G_2504_high_coverage/additional_698_related/data/ERR398924 |
| **HG04148** | 1000G_2504_high_coverage/additional_698_related/data/ERR398924 |
| **HG04149** | 1000G_2504_high_coverage/additional_698_related/data/ERR398924 |
| **HG04150** | 1000G_2504_high_coverage/additional_698_related/data/ERR398924 |
| **HG04151** | 1000G_2504_high_coverage/additional_698_related/data/ERR398924 |
| **HG04157** | 1000G_2504_high_coverage/additional_698_related/data/ERR398924 |
| **HG04160** | 1000G_2504_high_coverage/additional_698_related/data/ERR398925 |
| **HG04174** | 1000G_2504_high_coverage/additional_698_related/data/ERR398925 |
| **HG04175** | 1000G_2504_high_coverage/additional_698_related/data/ERR398925 |
| **HG04184** | 1000G_2504_high_coverage/additional_698_related/data/ERR398925 |
| **HG04187** | 1000G_2504_high_coverage/additional_698_related/data/ERR398925 |
| **HG04191** | 1000G_2504_high_coverage/additional_698_related/data/ERR398925 |
| **HG04192** | 1000G_2504_high_coverage/additional_698_related/data/ERR398925 |
| **HG04193** | 1000G_2504_high_coverage/additional_698_related/data/ERR398925 |
| **HG04199** | 1000G_2504_high_coverage/additional_698_related/data/ERR398925 |
| **HG04204** | 1000G_2504_high_coverage/additional_698_related/data/ERR398925 |
| **HG04215** | 1000G_2504_high_coverage/additional_698_related/data/ERR398926 |
| **HG04217** | 1000G_2504_high_coverage/additional_698_related/data/ERR398926 |
| **HG04228** | 1000G_2504_high_coverage/additional_698_related/data/ERR398926 |
| **NA06991** | 1000G_2504_high_coverage/additional_698_related/data/ERR398926 |
| **NA06993** | 1000G_2504_high_coverage/additional_698_related/data/ERR398926 |
| **NA06995** | 1000G_2504_high_coverage/additional_698_related/data/ERR398926 |
| **NA06997** | 1000G_2504_high_coverage/additional_698_related/data/ERR398926 |
| **NA07014** | 1000G_2504_high_coverage/additional_698_related/data/ERR398926 |
| **NA07019** | 1000G_2504_high_coverage/additional_698_related/data/ERR398926 |
| **NA07022** | 1000G_2504_high_coverage/additional_698_related/data/ERR398926 |
| **NA07029** | 1000G_2504_high_coverage/additional_698_related/data/ERR398927 |
| **NA07031** | 1000G_2504_high_coverage/additional_698_related/data/ERR398927 |
| **NA07034** | 1000G_2504_high_coverage/additional_698_related/data/ERR398927 |
| **NA07045** | 1000G_2504_high_coverage/additional_698_related/data/ERR398927 |
| **NA07055** | 1000G_2504_high_coverage/additional_698_related/data/ERR398927 |
| **NA07345** | 1000G_2504_high_coverage/additional_698_related/data/ERR398927 |
| **NA07346** | 1000G_2504_high_coverage/additional_698_related/data/ERR398927 |
| **NA07348** | 1000G_2504_high_coverage/additional_698_related/data/ERR398927 |
| **NA07349** | 1000G_2504_high_coverage/additional_698_related/data/ERR398927 |
| **NA07435** | 1000G_2504_high_coverage/additional_698_related/data/ERR398927 |
| **NA10830** | 1000G_2504_high_coverage/additional_698_related/data/ERR398928 |
| **NA10831** | 1000G_2504_high_coverage/additional_698_related/data/ERR398928 |
| **NA10835** | 1000G_2504_high_coverage/additional_698_related/data/ERR398928 |
| **NA10836** | 1000G_2504_high_coverage/additional_698_related/data/ERR398928 |
| **NA10837** | 1000G_2504_high_coverage/additional_698_related/data/ERR398928 |
| **NA10838** | 1000G_2504_high_coverage/additional_698_related/data/ERR398928 |
| **NA10839** | 1000G_2504_high_coverage/additional_698_related/data/ERR398928 |
| **NA10840** | 1000G_2504_high_coverage/additional_698_related/data/ERR398928 |
| **NA10842** | 1000G_2504_high_coverage/additional_698_related/data/ERR398928 |
| **NA10843** | 1000G_2504_high_coverage/additional_698_related/data/ERR398928 |
| **NA10845** | 1000G_2504_high_coverage/additional_698_related/data/ERR398929 |
| **NA10846** | 1000G_2504_high_coverage/additional_698_related/data/ERR398929 |
| **NA10852** | 1000G_2504_high_coverage/additional_698_related/data/ERR398929 |
| **NA10854** | 1000G_2504_high_coverage/additional_698_related/data/ERR398929 |
| **NA10855** | 1000G_2504_high_coverage/additional_698_related/data/ERR398929 |
| **NA10856** | 1000G_2504_high_coverage/additional_698_related/data/ERR398929 |
| **NA10857** | 1000G_2504_high_coverage/additional_698_related/data/ERR398929 |
| **NA10859** | 1000G_2504_high_coverage/additional_698_related/data/ERR398929 |
| **NA10860** | 1000G_2504_high_coverage/additional_698_related/data/ERR398929 |
| **NA10861** | 1000G_2504_high_coverage/additional_698_related/data/ERR398929 |
| **NA10863** | 1000G_2504_high_coverage/additional_698_related/data/ERR398930 |
| **NA10864** | 1000G_2504_high_coverage/additional_698_related/data/ERR398930 |
| **NA10865** | 1000G_2504_high_coverage/additional_698_related/data/ERR398930 |
| **NA11839** | 1000G_2504_high_coverage/additional_698_related/data/ERR398930 |
| **NA11882** | 1000G_2504_high_coverage/additional_698_related/data/ERR398930 |
| **NA11891** | 1000G_2504_high_coverage/additional_698_related/data/ERR398930 |
| **NA11917** | 1000G_2504_high_coverage/additional_698_related/data/ERR398930 |
| **NA11993** | 1000G_2504_high_coverage/additional_698_related/data/ERR398930 |
| **NA12056** | 1000G_2504_high_coverage/additional_698_related/data/ERR398930 |

| | |
|---|---|
| **NA12057** | 1000G_2504_high_coverage/additional_698_related/data/ERR398930 |
| **NA12145** | 1000G_2504_high_coverage/additional_698_related/data/ERR398931 |
| **NA12146** | 1000G_2504_high_coverage/additional_698_related/data/ERR398931 |
| **NA12236** | 1000G_2504_high_coverage/additional_698_related/data/ERR398931 |
| **NA12239** | 1000G_2504_high_coverage/additional_698_related/data/ERR398931 |
| **NA12248** | 1000G_2504_high_coverage/additional_698_related/data/ERR398931 |
| **NA12264** | 1000G_2504_high_coverage/additional_698_related/data/ERR398931 |
| **NA12274** | 1000G_2504_high_coverage/additional_698_related/data/ERR398931 |
| **NA12329** | 1000G_2504_high_coverage/additional_698_related/data/ERR398931 |
| **NA12335** | 1000G_2504_high_coverage/additional_698_related/data/ERR398931 |
| **NA12336** | 1000G_2504_high_coverage/additional_698_related/data/ERR398931 |
| **NA12343** | 1000G_2504_high_coverage/additional_698_related/data/ERR398932 |
| **NA12344** | 1000G_2504_high_coverage/additional_698_related/data/ERR398932 |
| **NA12376** | 1000G_2504_high_coverage/additional_698_related/data/ERR398932 |
| **NA12386** | 1000G_2504_high_coverage/additional_698_related/data/ERR398932 |
| **NA12485** | 1000G_2504_high_coverage/additional_698_related/data/ERR398932 |
| **NA12707** | 1000G_2504_high_coverage/additional_698_related/data/ERR398932 |
| **NA12739** | 1000G_2504_high_coverage/additional_698_related/data/ERR398932 |
| **NA12740** | 1000G_2504_high_coverage/additional_698_related/data/ERR398932 |
| **NA12752** | 1000G_2504_high_coverage/additional_698_related/data/ERR398932 |
| **NA12753** | 1000G_2504_high_coverage/additional_698_related/data/ERR398932 |
| **NA12766** | 1000G_2504_high_coverage/additional_698_related/data/ERR398933 |
| **NA12767** | 1000G_2504_high_coverage/additional_698_related/data/ERR398933 |
| **NA12801** | 1000G_2504_high_coverage/additional_698_related/data/ERR398933 |
| **NA12802** | 1000G_2504_high_coverage/additional_698_related/data/ERR398933 |
| **NA12817** | 1000G_2504_high_coverage/additional_698_related/data/ERR398933 |
| **NA12818** | 1000G_2504_high_coverage/additional_698_related/data/ERR398933 |
| **NA12832** | 1000G_2504_high_coverage/additional_698_related/data/ERR398933 |
| **NA12864** | 1000G_2504_high_coverage/additional_698_related/data/ERR398933 |
| **NA12865** | 1000G_2504_high_coverage/additional_698_related/data/ERR398933 |
| **NA12875** | 1000G_2504_high_coverage/additional_698_related/data/ERR398933 |
| **NA12877** | 1000G_2504_high_coverage/additional_698_related/data/ERR398934 |
| **NA12891** | 1000G_2504_high_coverage/additional_698_related/data/ERR398934 |
| **NA12892** | 1000G_2504_high_coverage/additional_698_related/data/ERR398934 |
| **NA18484** | 1000G_2504_high_coverage/additional_698_related/data/ERR398934 |
| **NA18485** | 1000G_2504_high_coverage/additional_698_related/data/ERR398934 |
| **NA18487** | 1000G_2504_high_coverage/additional_698_related/data/ERR398934 |
| **NA18497** | 1000G_2504_high_coverage/additional_698_related/data/ERR398934 |
| **NA18500** | 1000G_2504_high_coverage/additional_698_related/data/ERR398934 |
| **NA18503** | 1000G_2504_high_coverage/additional_698_related/data/ERR398934 |
| **NA18506** | 1000G_2504_high_coverage/additional_698_related/data/ERR398934 |
| **NA18515** | 1000G_2504_high_coverage/additional_698_related/data/ERR398935 |
| **NA18518** | 1000G_2504_high_coverage/additional_698_related/data/ERR398935 |
| **NA18521** | 1000G_2504_high_coverage/additional_698_related/data/ERR398935 |
| **NA18852** | 1000G_2504_high_coverage/additional_698_related/data/ERR398935 |
| **NA18854** | 1000G_2504_high_coverage/additional_698_related/data/ERR398935 |
| **NA18855** | 1000G_2504_high_coverage/additional_698_related/data/ERR398935 |
| **NA18857** | 1000G_2504_high_coverage/additional_698_related/data/ERR398935 |
| **NA18859** | 1000G_2504_high_coverage/additional_698_related/data/ERR398935 |
| **NA18860** | 1000G_2504_high_coverage/additional_698_related/data/ERR398935 |
| **NA18862** | 1000G_2504_high_coverage/additional_698_related/data/ERR398935 |
| **NA18863** | 1000G_2504_high_coverage/additional_698_related/data/ERR398936 |
| **NA18869** | 1000G_2504_high_coverage/additional_698_related/data/ERR398936 |
| **NA18872** | 1000G_2504_high_coverage/additional_698_related/data/ERR398936 |
| **NA18875** | 1000G_2504_high_coverage/additional_698_related/data/ERR398936 |
| **NA18906** | 1000G_2504_high_coverage/additional_698_related/data/ERR398936 |
| **NA18911** | 1000G_2504_high_coverage/additional_698_related/data/ERR398936 |
| **NA18913** | 1000G_2504_high_coverage/additional_698_related/data/ERR398936 |
| **NA18914** | 1000G_2504_high_coverage/additional_698_related/data/ERR398936 |
| **NA18925** | 1000G_2504_high_coverage/additional_698_related/data/ERR398936 |
| **NA18930** | 1000G_2504_high_coverage/additional_698_related/data/ERR398936 |
| **NA18935** | 1000G_2504_high_coverage/additional_698_related/data/ERR398937 |
| **NA19094** | 1000G_2504_high_coverage/additional_698_related/data/ERR398937 |
| **NA19097** | 1000G_2504_high_coverage/additional_698_related/data/ERR398937 |
| **NA19100** | 1000G_2504_high_coverage/additional_698_related/data/ERR398937 |

| | |
|---|---|
| **NA19101** | 1000G_2504_high_coverage/additional_698_related/data/ERR398937 |
| **NA19103** | 1000G_2504_high_coverage/additional_698_related/data/ERR398937 |
| **NA19109** | 1000G_2504_high_coverage/additional_698_related/data/ERR398937 |
| **NA19115** | 1000G_2504_high_coverage/additional_698_related/data/ERR398937 |
| **NA19120** | 1000G_2504_high_coverage/additional_698_related/data/ERR398937 |
| **NA19122** | 1000G_2504_high_coverage/additional_698_related/data/ERR398937 |
| **NA19123** | 1000G_2504_high_coverage/additional_698_related/data/ERR398938 |
| **NA19127** | 1000G_2504_high_coverage/additional_698_related/data/ERR398938 |
| **NA19128** | 1000G_2504_high_coverage/additional_698_related/data/ERR398938 |
| **NA19132** | 1000G_2504_high_coverage/additional_698_related/data/ERR398938 |
| **NA19139** | 1000G_2504_high_coverage/additional_698_related/data/ERR398938 |
| **NA19140** | 1000G_2504_high_coverage/additional_698_related/data/ERR398938 |
| **NA19142** | 1000G_2504_high_coverage/additional_698_related/data/ERR398938 |
| **NA19145** | 1000G_2504_high_coverage/additional_698_related/data/ERR398938 |
| **NA19148** | 1000G_2504_high_coverage/additional_698_related/data/ERR398938 |
| **NA19150** | 1000G_2504_high_coverage/additional_698_related/data/ERR398938 |
| **NA19151** | 1000G_2504_high_coverage/additional_698_related/data/ERR398939 |
| **NA19154** | 1000G_2504_high_coverage/additional_698_related/data/ERR398939 |
| **NA19161** | 1000G_2504_high_coverage/additional_698_related/data/ERR398939 |
| **NA19173** | 1000G_2504_high_coverage/additional_698_related/data/ERR398939 |
| **NA19174** | 1000G_2504_high_coverage/additional_698_related/data/ERR398939 |
| **NA19176** | 1000G_2504_high_coverage/additional_698_related/data/ERR398939 |
| **NA19177** | 1000G_2504_high_coverage/additional_698_related/data/ERR398939 |
| **NA19186** | 1000G_2504_high_coverage/additional_698_related/data/ERR398939 |
| **NA19191** | 1000G_2504_high_coverage/additional_698_related/data/ERR398939 |
| **NA19199** | 1000G_2504_high_coverage/additional_698_related/data/ERR398939 |
| **NA19202** | 1000G_2504_high_coverage/additional_698_related/data/ERR398940 |
| **NA19203** | 1000G_2504_high_coverage/additional_698_related/data/ERR398940 |
| **NA19205** | 1000G_2504_high_coverage/additional_698_related/data/ERR398940 |
| **NA19208** | 1000G_2504_high_coverage/additional_698_related/data/ERR398940 |
| **NA19211** | 1000G_2504_high_coverage/additional_698_related/data/ERR398940 |
| **NA19215** | 1000G_2504_high_coverage/additional_698_related/data/ERR398940 |
| **NA19221** | 1000G_2504_high_coverage/additional_698_related/data/ERR398940 |
| **NA19224** | 1000G_2504_high_coverage/additional_698_related/data/ERR398940 |
| **NA19226** | 1000G_2504_high_coverage/additional_698_related/data/ERR398940 |
| **NA19237** | 1000G_2504_high_coverage/additional_698_related/data/ERR398940 |
| **NA19240** | 1000G_2504_high_coverage/additional_698_related/data/ERR398941 |
| **NA19249** | 1000G_2504_high_coverage/additional_698_related/data/ERR398941 |
| **NA19258** | 1000G_2504_high_coverage/additional_698_related/data/ERR398941 |
| **NA19650** | 1000G_2504_high_coverage/additional_698_related/data/ERR398941 |
| **NA19653** | 1000G_2504_high_coverage/additional_698_related/data/ERR398941 |
| **NA19656** | 1000G_2504_high_coverage/additional_698_related/data/ERR398941 |
| **NA19659** | 1000G_2504_high_coverage/additional_698_related/data/ERR398941 |
| **NA19660** | 1000G_2504_high_coverage/additional_698_related/data/ERR398941 |
| **NA19662** | 1000G_2504_high_coverage/additional_698_related/data/ERR398941 |
| **NA19665** | 1000G_2504_high_coverage/additional_698_related/data/ERR398941 |
| **NA19671** | 1000G_2504_high_coverage/additional_698_related/data/ERR398942 |
| **NA19675** | 1000G_2504_high_coverage/additional_698_related/data/ERR398942 |
| **NA19677** | 1000G_2504_high_coverage/additional_698_related/data/ERR398942 |
| **NA19680** | 1000G_2504_high_coverage/additional_698_related/data/ERR398942 |
| **NA19683** | 1000G_2504_high_coverage/additional_698_related/data/ERR398942 |
| **NA19685** | 1000G_2504_high_coverage/additional_698_related/data/ERR398942 |
| **NA19686** | 1000G_2504_high_coverage/additional_698_related/data/ERR398942 |
| **NA19702** | 1000G_2504_high_coverage/additional_698_related/data/ERR398942 |
| **NA19705** | 1000G_2504_high_coverage/additional_698_related/data/ERR398942 |
| **NA19718** | 1000G_2504_high_coverage/additional_698_related/data/ERR398942 |
| **NA19721** | 1000G_2504_high_coverage/additional_698_related/data/ERR398943 |
| **NA19724** | 1000G_2504_high_coverage/additional_698_related/data/ERR398943 |
| **NA19727** | 1000G_2504_high_coverage/additional_698_related/data/ERR398943 |
| **NA19730** | 1000G_2504_high_coverage/additional_698_related/data/ERR398943 |
| **NA19733** | 1000G_2504_high_coverage/additional_698_related/data/ERR398943 |
| **NA19748** | 1000G_2504_high_coverage/additional_698_related/data/ERR398943 |
| **NA19751** | 1000G_2504_high_coverage/additional_698_related/data/ERR398943 |
| **NA19757** | 1000G_2504_high_coverage/additional_698_related/data/ERR398943 |
| **NA19760** | 1000G_2504_high_coverage/additional_698_related/data/ERR398943 |

| | |
|---|---|
| **NA19763** | 1000G_2504_high_coverage/additional_698_related/data/ERR398943 |
| **NA19772** | 1000G_2504_high_coverage/additional_698_related/data/ERR398944 |
| **NA19775** | 1000G_2504_high_coverage/additional_698_related/data/ERR398944 |
| **NA19778** | 1000G_2504_high_coverage/additional_698_related/data/ERR398944 |
| **NA19781** | 1000G_2504_high_coverage/additional_698_related/data/ERR398944 |
| **NA19784** | 1000G_2504_high_coverage/additional_698_related/data/ERR398944 |
| **NA19787** | 1000G_2504_high_coverage/additional_698_related/data/ERR398944 |
| **NA19790** | 1000G_2504_high_coverage/additional_698_related/data/ERR398944 |
| **NA19796** | 1000G_2504_high_coverage/additional_698_related/data/ERR398944 |
| **NA19828** | 1000G_2504_high_coverage/additional_698_related/data/ERR398944 |
| **NA19836** | 1000G_2504_high_coverage/additional_698_related/data/ERR398944 |
| **NA19902** | 1000G_2504_high_coverage/additional_698_related/data/ERR398945 |
| **NA19918** | 1000G_2504_high_coverage/additional_698_related/data/ERR398945 |
| **NA19919** | 1000G_2504_high_coverage/additional_698_related/data/ERR398945 |
| **NA19924** | 1000G_2504_high_coverage/additional_698_related/data/ERR398945 |
| **NA19983** | 1000G_2504_high_coverage/additional_698_related/data/ERR398945 |
| **NA20128** | 1000G_2504_high_coverage/additional_698_related/data/ERR398945 |
| **NA20129** | 1000G_2504_high_coverage/additional_698_related/data/ERR398945 |
| **NA20279** | 1000G_2504_high_coverage/additional_698_related/data/ERR398945 |
| **NA20358** | 1000G_2504_high_coverage/additional_698_related/data/ERR398945 |
