## Supplemental Table 2 for "High-throughput complement component 4 genomic sequence analysis with C4Investigator"

Table_S2_suppInfo

| Sample name | Sex | Biosample ID | Population code | Population name | Superpopulation code | Superpopulation name | Population elastic ID | Data collections |
|---|---|---|---|---|---|---|---|---|
| **HG00315** | female | SAME124335 | FIN | Finnish | EUR | European Ancestry | FIN | 1000 Genomes on GRCh38,1000 Genomes 30x on GRCh38,1000 Genomes phase 3 release,1000 Genomes phase 1 release,Geuvadis |
| **HG00327** | female | SAME123791 | FIN | Finnish | EUR | European Ancestry | FIN | 1000 Genomes on GRCh38,1000 Genomes 30x on GRCh38,1000 Genomes phase 3 release,1000 Genomes phase 1 release,Geuvadis |
| **HG00334** | female | SAME123981 | FIN | Finnish | EUR | European Ancestry | FIN | 1000 Genomes on GRCh38,1000 Genomes 30x on GRCh38,1000 Genomes phase 3 release,1000 Genomes phase 1 release,Geuvadis |
| **HG00339** | female | SAME123991 | FIN | Finnish | EUR | European Ancestry | FIN | 1000 Genomes on GRCh38,1000 Genomes 30x on GRCh38,1000 Genomes phase 3 release,1000 Genomes phase 1 release,Geuvadis |
| **HG00341** | male | SAME124922 | FIN | Finnish | EUR | European Ancestry | FIN | 1000 Genomes on GRCh38,1000 Genomes 30x on GRCh38,1000 Genomes phase 3 release,1000 Genomes phase 1 release,Geuvadis |
| **HG00346** | female | SAME125264 | FIN | Finnish | EUR | European Ancestry | FIN | 1000 Genomes on GRCh38,1000 Genomes 30x on GRCh38,1000 Genomes phase 3 release,1000 Genomes phase 1 release,Geuvadis |
| **HG00353** | female | SAME125120 | FIN | Finnish | EUR | European Ancestry | FIN | 1000 Genomes on GRCh38,1000 Genomes 30x on GRCh38,1000 Genomes phase 3 release,1000 Genomes phase 1 release,Geuvadis |
| **HG00358** | male | SAME125126 | FIN | Finnish | EUR | European Ancestry | FIN | 1000 Genomes on GRCh38,1000 Genomes 30x on GRCh38,1000 Genomes phase 3 release,1000 Genomes phase 1 release,Geuvadis |
| **HG00360** | male | SAME124539 | FIN | Finnish,Finnish | EUR | European Ancestry,West Eurasia (SGDP) | FIN,FinnishSGDP | 1000 Genomes on GRCh38,Simons Genome Diversity Project,1000 Genomes 30x on GRCh38,1000 Genomes phase 3 release,1000 Genomes phase 1 release,Geuvadis |
| **HG00365** | female | SAME124542 | FIN | Finnish | EUR | European Ancestry | FIN | 1000 Genomes on GRCh38,1000 Genomes 30x on GRCh38,1000 Genomes phase 3 release,1000 Genomes phase 1 release,Geuvadis |
| **HG00372** | male | SAME124748 | FIN | Finnish | EUR | European Ancestry | FIN | 1000 Genomes on GRCh38,1000 Genomes 30x on GRCh38,1000 Genomes phase 3 release,1000 Genomes phase 1 release,Geuvadis |
| **HG00384** | female | SAME123034 | FIN | Finnish | EUR | European Ancestry | FIN | 1000 Genomes on GRCh38,1000 Genomes 30x on GRCh38,1000 Genomes phase 3 release,1000 Genomes phase 1 release,Geuvadis |
| **HG00404** | female | SAME123158 | CHS | Southern Han Chinese | EAS | East Asian Ancestry | CHS | 1000 Genomes on GRCh38,1000 Genomes 30x on GRCh38,1000 Genomes phase 3 release,1000 Genomes phase 1 release |
| **HG00409** | male | SAME1840026 | CHS | Southern Han Chinese | EAS | East Asian Ancestry | CHS | 1000 Genomes on GRCh38,1000 Genomes 30x on GRCh38,1000 Genomes phase 3 release |
| **HG00106** | female | SAME123948 | GBR | British | EUR | European Ancestry | GBR | 1000 Genomes on GRCh38,1000 Genomes 30x on GRCh38,1000 Genomes phase 3 release,1000 Genomes phase 1 release,Geuvadis |
| **HG00113** | male | SAME125342 | GBR | British | EUR | European Ancestry | GBR | 1000 Genomes on GRCh38,1000 Genomes 30x on GRCh38,1000 Genomes phase 3 release,1000 Genomes phase 1 release |
| **HG00118** | female | SAME125347 | GBR | British | EUR | European Ancestry | GBR | 1000 Genomes on GRCh38,1000 Genomes 30x on GRCh38,1000 Genomes phase 3 release,1000 Genomes phase 1 release,Geuvadis |
| **HG00120** | female | SAME122874 | GBR | British | EUR | European Ancestry | GBR | 1000 Genomes on GRCh38,1000 Genomes 30x on GRCh38,1000 Genomes phase 3 release,1000 Genomes phase 1 release,Geuvadis |
| **HG00125** | female | SAME122869 | GBR | British | EUR | European Ancestry | GBR | 1000 Genomes on GRCh38,1000 Genomes 30x on GRCh38,1000 Genomes phase 3 release,1000 Genomes phase 1 release,Geuvadis |
| **HG00132** | female | SAME123061 | GBR | British | EUR | European Ancestry | GBR | 1000 Genomes on GRCh38,1000 Genomes 30x on GRCh38,1000 Genomes phase 3 release,1000 Genomes phase 1 release,Geuvadis |
| **HG00137** | female | SAME123066 | GBR | British | EUR | European Ancestry | GBR | 1000 Genomes on GRCh38,1000 Genomes 30x on GRCh38,1000 Genomes phase 3 release,1000 Genomes phase 1 release,Geuvadis |
| **HG00149** | male | SAME124387 | GBR | British | EUR | European Ancestry | GBR | 1000 Genomes on GRCh38,1000 Genomes 30x on GRCh38,1000 Genomes phase 3 release,1000 Genomes phase 1 release,Geuvadis |
| **HG00151** | male | SAME124592 | GBR | British | EUR | European Ancestry | GBR | 1000 Genomes on GRCh38,1000 Genomes 30x on GRCh38,1000 Genomes phase 3 release,1000 Genomes phase 1 release,Geuvadis |
| **HG00182** | male | SAME123643 | FIN | Finnish | EUR | European Ancestry | FIN | 1000 Genomes on GRCh38,1000 Genomes 30x on GRCh38,1000 Genomes phase 3 release,1000 Genomes phase 1 release,Geuvadis |
| **HG00187** | male | SAME123646 | FIN | Finnish | EUR | European Ancestry | FIN | 1000 Genomes on GRCh38,1000 Genomes 30x on GRCh38,1000 Genomes phase 3 release,1000 Genomes phase 1 release,Geuvadis |
| **HG00233** | female | SAME124127 | GBR | British | EUR | European Ancestry | GBR | 1000 Genomes on GRCh38,1000 Genomes 30x on GRCh38,1000 Genomes phase 3 release,1000 Genomes phase 1 release,Geuvadis |
| **HG00238** | female | SAME124131 | GBR | British | EUR | European Ancestry | GBR | 1000 Genomes on GRCh38,1000 Genomes 30x on GRCh38,1000 Genomes phase 3 release,1000 Genomes phase 1 release,Geuvadis |
| **HG00240** | female | SAME123221 | GBR | British | EUR | European Ancestry | GBR | 1000 Genomes on GRCh38,1000 Genomes 30x on GRCh38,1000 Genomes phase 3 release,1000 Genomes phase 1 release,Geuvadis |
| **HG00245** | female | SAME123218 | GBR | British | EUR | European Ancestry | GBR | 1000 Genomes on GRCh38,1000 Genomes 30x on GRCh38,1000 Genomes phase 3 release,1000 Genomes phase 1 release,Geuvadis |
| **HG00252** | male | SAME123052 | GBR | British | EUR | European Ancestry | GBR | 1000 Genomes on GRCh38,1000 Genomes 30x on GRCh38,1000 Genomes phase 3 release,1000 Genomes phase 1 release,Geuvadis |
| **HG00257** | female | SAME123047 | GBR | British | EUR | European Ancestry | GBR | 1000 Genomes on GRCh38,1000 Genomes 30x on GRCh38,1000 Genomes phase 3 release,1000 Genomes phase 1 release,Geuvadis |
| **HG00097** | female | SAME123267 | GBR | British | EUR | European Ancestry | GBR | 1000 Genomes on GRCh38,1000 Genomes 30x on GRCh38,1000 Genomes phase 3 release,1000 Genomes phase 1 release,Geuvadis |
| **HG00264** | male | SAME123578 | GBR | British | EUR | European Ancestry | GBR | 1000 Genomes on GRCh38,1000 Genomes 30x on GRCh38,1000 Genomes phase 3 release,1000 Genomes phase 1 release,Geuvadis |
| **HG00100** | female | SAME125154 | GBR | British | EUR | European Ancestry | GBR | 1000 Genomes on GRCh38,1000 Genomes 30x on GRCh38,1000 Genomes phase 3 release,1000 Genomes phase 1 release,Geuvadis |
| **HG00269** | female | SAME123575 | FIN | Finnish | EUR | European Ancestry | FIN | 1000 Genomes on GRCh38,1000 Genomes 30x on GRCh38,1000 Genomes phase 3 release,1000 Genomes phase 1 release,Geuvadis |
| **HG00105** | male | SAME123949 | GBR | British | EUR | European Ancestry | GBR | 1000 Genomes on GRCh38,1000 Genomes 30x on GRCh38,1000 Genomes phase 3 release,1000 Genomes phase 1 release,Geuvadis |
| **HG00271** | male | SAME123417 | FIN | Finnish | EUR | European Ancestry | FIN | 1000 Genomes on GRCh38,1000 Genomes 30x on GRCh38,1000 Genomes phase 3 release,1000 Genomes phase 1 release,Geuvadis |
| **HG00112** | male | SAME125341 | GBR | British | EUR | European Ancestry | GBR | 1000 Genomes on GRCh38,1000 Genomes 30x on GRCh38,1000 Genomes phase 3 release,1000 Genomes phase 1 release,Geuvadis |
| **HG00276** | female | SAME123424 | FIN | Finnish | EUR | European Ancestry | FIN | 1000 Genomes on GRCh38,1000 Genomes 30x on GRCh38,1000 Genomes phase 3 release,1000 Genomes phase 1 release,Geuvadis |
| **HG00117** | male | SAME125346 | GBR | British | EUR | European Ancestry | GBR | 1000 Genomes on GRCh38,1000 Genomes 30x on GRCh38,1000 Genomes phase 3 release,1000 Genomes phase 1 release,Geuvadis |
| **HG00288** | female | SAME1839246 | FIN | Finnish | EUR | European Ancestry | FIN | 1000 Genomes on GRCh38,1000 Genomes 30x on GRCh38,1000 Genomes phase 3 release |
| **HG00290** | male | SAME1839057 | FIN | Finnish | EUR | European Ancestry | FIN | 1000 Genomes on GRCh38,1000 Genomes 30x on GRCh38,1000 Genomes phase 3 release |
| **HG00129** | male | SAME122867 | GBR | British | EUR | European Ancestry | GBR | 1000 Genomes on GRCh38,1000 Genomes 30x on GRCh38,1000 Genomes phase 3 release,1000 Genomes phase 1 release,Geuvadis |
| **HG00131** | male | SAME123064 | GBR | British | EUR | European Ancestry | GBR | 1000 Genomes on GRCh38,1000 Genomes 30x on GRCh38,1000 Genomes phase 3 release,1000 Genomes phase 1 release,Geuvadis |
| **HG00308** | male | SAME124161 | FIN | Finnish | EUR | European Ancestry | FIN | 1000 Genomes on GRCh38,1000 Genomes 30x on GRCh38,1000 Genomes phase 3 release,Geuvadis |
| **HG00136** | male | SAME123065 | GBR | British | EUR | European Ancestry | GBR | 1000 Genomes on GRCh38,1000 Genomes 30x on GRCh38,1000 Genomes phase 3 release,1000 Genomes phase 1 release,Geuvadis |
| **HG00310** | male | SAME124338 | FIN | Finnish | EUR | European Ancestry | FIN | 1000 Genomes on GRCh38,1000 Genomes 30x on GRCh38,1000 Genomes phase 3 release,1000 Genomes phase 1 release,Geuvadis |
| **HG00143** | male | SAME124393 | GBR | British | EUR | European Ancestry | GBR | 1000 Genomes on GRCh38,1000 Genomes 30x on GRCh38,1000 Genomes phase 3 release,1000 Genomes phase 1 release,Geuvadis |
| **HG00148** | male | SAME124388 | GBR | British | EUR | European Ancestry | GBR | 1000 Genomes on GRCh38,1000 Genomes 30x on GRCh38,1000 Genomes phase 3 release,1000 Genomes phase 1 release,Geuvadis |
| **HG00101** | male | SAME125153 | GBR | British | EUR | European Ancestry | GBR | 1000 Genomes on GRCh38,1000 Genomes 30x on GRCh38,1000 Genomes phase 3 release,1000 Genomes phase 1 release,Geuvadis |
| **HG00150** | female | SAME124591 | GBR | British | EUR | European Ancestry | GBR | 1000 Genomes on GRCh38,1000 Genomes 30x on GRCh38,1000 Genomes phase 3 release,1000 Genomes phase 1 release,Geuvadis |
| **HG00423** | female | SAME122810 | CHS | Southern Han Chinese | EAS | East Asian Ancestry | CHS | 1000 Genomes 30x on GRCh38,1000 Genomes phase 3 release |
| **HG00155** | male | SAME124588 | GBR | British | EUR | European Ancestry | GBR | 1000 Genomes on GRCh38,1000 Genomes 30x on GRCh38,1000 Genomes phase 3 release,1000 Genomes phase 1 release,Geuvadis |
| **HG00428** | female | SAME122811 | CHS | Southern Han Chinese | EAS | East Asian Ancestry | CHS | 1000 Genomes on GRCh38,1000 Genomes 30x on GRCh38,1000 Genomes phase 3 release,1000 Genomes phase 1 release,90 Han Chinese high coverage genomes |
| **HG00174** | female | SAME124958 | FIN | Finnish,Finnish | EUR | European Ancestry,West Eurasia (SGDP) | FIN,FinnishSGDP | 1000 Genomes on GRCh38,Simons Genome Diversity Project,1000 Genomes 30x on GRCh38,1000 Genomes phase 3 release,1000 Genomes phase 1 release,Geuvadis |
| **HG00442** | male | SAME124912 | CHS | Southern Han Chinese | EAS | East Asian Ancestry | CHS | 1000 Genomes on GRCh38,1000 Genomes 30x on GRCh38,1000 Genomes phase 3 release,1000 Genomes phase 1 release,90 Han Chinese high coverage genomes |
| **HG00179** | female | SAME124965 | FIN | Finnish | EUR | European Ancestry | FIN | 1000 Genomes on GRCh38,1000 Genomes 30x on GRCh38,1000 Genomes phase 3 release,1000 Genomes phase 1 release,Geuvadis |
| **HG00447** | female | SAME124953 | CHS | Southern Han Chinese | EAS | East Asian Ancestry | CHS | 1000 Genomes 30x on GRCh38,1000 Genomes phase 3 release |
| **HG00181** | male | SAME123644 | FIN | Finnish | EUR | European Ancestry | FIN | 1000 Genomes on GRCh38,1000 Genomes 30x on GRCh38,1000 Genomes phase 3 release,1000 Genomes phase 1 release,Geuvadis |
| **HG00459** | male | SAME125269 | CHS | Southern Han Chinese | EAS | East Asian Ancestry | CHS | 1000 Genomes 30x on GRCh38,1000 Genomes phase 3 release |
| **HG00186** | male | SAME123647 | FIN | Finnish | EUR | European Ancestry | FIN | 1000 Genomes on GRCh38,1000 Genomes 30x on GRCh38,1000 Genomes phase 3 release,1000 Genomes phase 1 release,Geuvadis |
| **HG00473** | female | SAME123338 | CHS | Southern Han Chinese | EAS | East Asian Ancestry | CHS | 1000 Genomes on GRCh38,1000 Genomes 30x on GRCh38,1000 Genomes phase 3 release,1000 Genomes phase 1 release,90 Han Chinese high coverage genomes |
| **HG00232** | female | SAME124128 | GBR | British | EUR | European Ancestry | GBR | 1000 Genomes on GRCh38,1000 Genomes 30x on GRCh38,1000 Genomes phase 3 release,1000 Genomes phase 1 release,Geuvadis |
| **HG00478** | male | SAME123335 | CHS | Southern Han Chinese | EAS | East Asian Ancestry | CHS | 1000 Genomes on GRCh38,1000 Genomes 30x on GRCh38,1000 Genomes phase 3 release,1000 Genomes phase 1 release,90 Han Chinese high coverage genomes |
| **HG00237** | female | SAME124124 | GBR | British | EUR | European Ancestry | GBR | 1000 Genomes on GRCh38,1000 Genomes 30x on GRCh38,1000 Genomes phase 3 release,1000 Genomes phase 1 release,Geuvadis |
| **HG00480** | male | SAME124617 | CHS | Southern Han Chinese | EAS | East Asian Ancestry | CHS | 1000 Genomes 30x on GRCh38,1000 Genomes phase 3 release |
| **HG00244** | male | SAME123217 | GBR | British | EUR | European Ancestry | GBR | 1000 Genomes on GRCh38,1000 Genomes 30x on GRCh38,1000 Genomes phase 3 release,1000 Genomes phase 1 release,Geuvadis |
| **HG00500** | male | SAME123363 | CHS | Southern Han Chinese | EAS | East Asian Ancestry | CHS | 1000 Genomes on GRCh38,1000 Genomes 30x on GRCh38,1000 Genomes phase 3 release,1000 Genomes phase 1 release,90 Han Chinese high coverage genomes |
| **HG00319** | female | SAME124334 | FIN | Finnish | EUR | European Ancestry | FIN | 1000 Genomes on GRCh38,1000 Genomes 30x on GRCh38,1000 Genomes phase 3 release,1000 Genomes phase 1 release,Geuvadis |
| **HG00512** | male | SAME123525 | CHS | Southern Han Chinese | EAS | East Asian Ancestry | CHS | 1000 Genomes on GRCh38,1000 Genomes 30x on GRCh38,Human Genome Structural Variation Consortium, Phase 2,The Human Genome Structural Variation Consortiu |
| **HG00321** | male | SAME123796 | FIN | Finnish | EUR | European Ancestry | FIN | 1000 Genomes on GRCh38,1000 Genomes 30x on GRCh38,1000 Genomes phase 3 release,1000 Genomes phase 1 release,Geuvadis |
| **HG00524** | male | SAME123724 | CHS | Southern Han Chinese | EAS | East Asian Ancestry | CHS | 1000 Genomes on GRCh38,1000 Genomes 30x on GRCh38,1000 Genomes phase 3 release,1000 Genomes phase 1 release,90 Han Chinese high coverage genomes |
| **HG00326** | female | SAME123792 | FIN | Finnish | EUR | European Ancestry | FIN | 1000 Genomes on GRCh38,1000 Genomes 30x on GRCh38,1000 Genomes phase 3 release,1000 Genomes phase 1 release,Geuvadis |
| **HG00531** | female | SAME125381 | CHS | Southern Han Chinese | EAS | East Asian Ancestry | CHS | 1000 Genomes on GRCh38,1000 Genomes 30x on GRCh38,1000 Genomes phase 3 release,1000 Genomes phase 1 release,90 Han Chinese high coverage genomes |
| **HG00338** | male | SAME123990 | FIN | Finnish | EUR | European Ancestry | FIN | 1000 Genomes on GRCh38,1000 Genomes 30x on GRCh38,1000 Genomes phase 3 release,1000 Genomes phase 1 release,Geuvadis |
| **HG00536** | male | SAME124344 | CHS | Southern Han Chinese | EAS | East Asian Ancestry | CHS | 1000 Genomes on GRCh38,1000 Genomes 30x on GRCh38,1000 Genomes phase 3 release,1000 Genomes phase 1 release,90 Han Chinese high coverage genomes |
| **HG00345** | male | SAME124919 | FIN | Finnish | EUR | European Ancestry | FIN | 1000 Genomes on GRCh38,1000 Genomes 30x on GRCh38,1000 Genomes phase 3 release,1000 Genomes phase 1 release,Geuvadis |
| **HG00733** | female | SAME125399 | PUR | Puerto Rican | AMR | American Ancestry | PUR | 1000 Genomes on GRCh38,1000 Genomes 30x on GRCh38,Human Genome Structural Variation Consortium, Phase 2,The Human Genome Structural Variation Consortiu |
| **HG00357** | female | SAME125124 | FIN | Finnish | EUR | European Ancestry | FIN | 1000 Genomes on GRCh38,1000 Genomes 30x on GRCh38,1000 Genomes phase 3 release,1000 Genomes phase 1 release |
| **HG00738** | male | SAME125368 | PUR | Puerto Rican | AMR | American Ancestry | PUR | 1000 Genomes 30x on GRCh38,1000 Genomes phase 3 release |
| **HG00364** | female | SAME124543 | FIN | Finnish | EUR | European Ancestry | FIN | 1000 Genomes on GRCh38,1000 Genomes 30x on GRCh38,1000 Genomes phase 3 release,1000 Genomes phase 1 release,Geuvadis |
| **HG00740** | female | SAME124834 | PUR | Puerto Rican | AMR | American Ancestry | PUR | 1000 Genomes on GRCh38,1000 Genomes 30x on GRCh38,1000 Genomes phase 3 release,1000 Genomes phase 1 release |
| **HG00369** | male | SAME124534 | FIN | Finnish | EUR | European Ancestry | FIN | 1000 Genomes on GRCh38,1000 Genomes 30x on GRCh38,1000 Genomes phase 3 release,1000 Genomes phase 1 release,Geuvadis |
| **HG00978** | female | SAME124727 | CDX | Dai Chinese | EAS | East Asian Ancestry | CDX | 1000 Genomes on GRCh38,1000 Genomes 30x on GRCh38,1000 Genomes phase 3 release |
| **HG00371** | male | SAME124747 | FIN | Finnish | EUR | European Ancestry | FIN | 1000 Genomes on GRCh38,1000 Genomes 30x on GRCh38,1000 Genomes phase 3 release,1000 Genomes phase 1 release,Geuvadis |
| **HG01192** | male | SAME124298 | PUR | Puerto Rican | AMR | American Ancestry | PUR | 1000 Genomes 30x on GRCh38,1000 Genomes phase 3 release |
| **HG00376** | female | SAME124751 | FIN | Finnish | EUR | European Ancestry | FIN | 1000 Genomes on GRCh38,1000 Genomes 30x on GRCh38,1000 Genomes phase 3 release,1000 Genomes phase 1 release,Geuvadis |
| **HG01197** | male | SAME124237 | PUR | Puerto Rican | AMR | American Ancestry | PUR | 1000 Genomes on GRCh38,1000 Genomes 30x on GRCh38,1000 Genomes phase 3 release,1000 Genomes phase 1 release |
| **HG00383** | female | SAME123032 | FIN | Finnish | EUR | European Ancestry | FIN | 1000 Genomes on GRCh38,1000 Genomes 30x on GRCh38,1000 Genomes phase 3 release,1000 Genomes phase 1 release,Geuvadis |
| **HG01200** | male | SAME1840109 | PUR | Puerto Rican | AMR | American Ancestry | PUR | 1000 Genomes on GRCh38,1000 Genomes 30x on GRCh38,1000 Genomes phase 3 release |
| **HG00403** | male | SAME123160 | CHS | Southern Han Chinese | EAS | East Asian Ancestry | CHS | 1000 Genomes on GRCh38,1000 Genomes 30x on GRCh38,1000 Genomes phase 3 release,1000 Genomes phase 1 release |
| **HG01205** | female | SAME124296 | PUR | Puerto Rican | AMR | American Ancestry | PUR | 1000 Genomes on GRCh38,1000 Genomes 30x on GRCh38,1000 Genomes phase 3 release,1000 Genomes phase 1 release |
| **HG00408** | female | SAME123155 | CHS | Southern Han Chinese | EAS | East Asian Ancestry | CHS | 1000 Genomes 30x on GRCh38,1000 Genomes phase 3 release |
| **HG01243** | male | SAME125063 | PUR | Puerto Rican | AMR | American Ancestry | PUR | 1000 Genomes 30x on GRCh38,1000 Genomes phase 3 release |
| **HG00410** | female | SAME1839734 | CHS | Southern Han Chinese | EAS | East Asian Ancestry | CHS | 1000 Genomes on GRCh38,1000 Genomes 30x on GRCh38,1000 Genomes phase 3 release |
| **HG01248** | female | SAME125058 | PUR | Puerto Rican | AMR | American Ancestry | PUR | 1000 Genomes on GRCh38,1000 Genomes 30x on GRCh38,1000 Genomes phase 3 release,1000 Genomes phase 1 release |
| **HG00422** | female | SAME122809 | CHS | Southern Han Chinese | EAS | East Asian Ancestry | CHS | 1000 Genomes on GRCh38,1000 Genomes 30x on GRCh38,1000 Genomes phase 3 release,1000 Genomes phase 1 release,90 Han Chinese high coverage genomes |
| **HG01250** | male | SAME125281 | CLM | Colombian | AMR | American Ancestry | CLM | 1000 Genomes on GRCh38,1000 Genomes 30x on GRCh38,1000 Genomes phase 3 release,1000 Genomes phase 1 release |
| **HG00542** | male | SAME124352 | CHS | Southern Han Chinese | EAS | East Asian Ancestry | CHS | 1000 Genomes on GRCh38,1000 Genomes 30x on GRCh38,1000 Genomes phase 3 release,1000 Genomes phase 1 release,90 Han Chinese high coverage genomes |
| **HG01255** | male | SAME125286 | CLM | Colombian | AMR | American Ancestry | CLM | 1000 Genomes 30x on GRCh38,1000 Genomes phase 3 release |
| **HG00554** | female | SAME123969 | PUR | Puerto Rican | AMR | American Ancestry | PUR | 1000 Genomes on GRCh38,1000 Genomes 30x on GRCh38,1000 Genomes phase 3 release,1000 Genomes phase 1 release |
| **HG01274** | male | SAME123001 | CLM | Colombian | AMR | American Ancestry | CLM | 1000 Genomes on GRCh38,1000 Genomes 30x on GRCh38,1000 Genomes phase 3 release,1000 Genomes phase 1 release |
| **HG00559** | male | SAME124248 | CHS | Southern Han Chinese | EAS | East Asian Ancestry | CHS | 1000 Genomes on GRCh38,1000 Genomes 30x on GRCh38,1000 Genomes phase 3 release,1000 Genomes phase 1 release |
| **HG01279** | female | SAME123004 | CLM | Colombian | AMR | American Ancestry | CLM | 1000 Genomes 30x on GRCh38,1000 Genomes phase 3 release |
| **HG00561** | female | SAME124455 | CHS | Southern Han Chinese | EAS | East Asian Ancestry | CHS | 1000 Genomes 30x on GRCh38,1000 Genomes phase 3 release |
| **HG00620** | female | SAME123081 | CHS | Southern Han Chinese | EAS | East Asian Ancestry | CHS | 1000 Genomes on GRCh38,1000 Genomes 30x on GRCh38,1000 Genomes phase 3 release,1000 Genomes phase 1 release,90 Han Chinese high coverage genomes |
| **HG00566** | female | SAME125048 | CHS | Southern Han Chinese | EAS | East Asian Ancestry | CHS | 1000 Genomes on GRCh38,1000 Genomes 30x on GRCh38,1000 Genomes phase 3 release,1000 Genomes phase 1 release |
| **HG00625** | male | SAME123080 | CHS | Southern Han Chinese | EAS | East Asian Ancestry | CHS | 1000 Genomes on GRCh38,1000 Genomes 30x on GRCh38,1000 Genomes phase 3 release,1000 Genomes phase 1 release |
| **HG00578** | female | SAME123528 | CHS | Southern Han Chinese | EAS | East Asian Ancestry | CHS | 1000 Genomes on GRCh38,1000 Genomes 30x on GRCh38,1000 Genomes phase 3 release,1000 Genomes phase 1 release |

| | | | | | | | | |
|---|---|---|---|---|---|---|---|---|
| **HG00632** | female | SAME1839151 | CHS | Southern Han Chinese | EAS | East Asian Ancestry | CHS | 1000 Genomes on GRCh38,1000 Genomes 30x on GRCh38,1000 Genomes phase 3 release |
| **HG00580** | male | SAME124832 | CHS | Southern Han Chinese | EAS | East Asian Ancestry | CHS | 1000 Genomes on GRCh38,1000 Genomes 30x on GRCh38,1000 Genomes phase 3 release,1000 Genomes phase 1 release,90 Han Chinese high coverage genomes |
| **HG00637** | male | SAME123710 | PUR | Puerto Rican | AMR | American Ancestry | PUR | 1000 Genomes on GRCh38,1000 Genomes 30x on GRCh38,1000 Genomes phase 3 release,1000 Genomes phase 1 release |
| **HG00585** | female | SAME124722 | CHS | Southern Han Chinese | EAS | East Asian Ancestry | CHS | 1000 Genomes 30x on GRCh38,1000 Genomes phase 3 release |
| **HG00651** | female | SAME123966 | CHS | Southern Han Chinese | EAS | East Asian Ancestry | CHS | 1000 Genomes on GRCh38,1000 Genomes 30x on GRCh38,1000 Genomes phase 3 release,1000 Genomes phase 1 release |
| **HG00592** | male | SAME125011 | CHS | Southern Han Chinese | EAS | East Asian Ancestry | CHS | 1000 Genomes on GRCh38,1000 Genomes 30x on GRCh38,1000 Genomes phase 3 release,1000 Genomes phase 1 release |
| **HG00656** | male | SAME123233 | CHS | Southern Han Chinese | EAS | East Asian Ancestry | CHS | 1000 Genomes on GRCh38,1000 Genomes 30x on GRCh38,1000 Genomes phase 3 release,1000 Genomes phase 1 release,90 Han Chinese high coverage genomes |
| **HG00597** | female | SAME125008 | CHS | Southern Han Chinese | EAS | East Asian Ancestry | CHS | 1000 Genomes 30x on GRCh38,1000 Genomes phase 3 release |
| **HG00663** | female | SAME123813 | CHS | Southern Han Chinese | EAS | East Asian Ancestry | CHS | 1000 Genomes on GRCh38,1000 Genomes 30x on GRCh38,1000 Genomes phase 3 release,1000 Genomes phase 1 release,90 Han Chinese high coverage genomes |
| **HG00543** | female | SAME122857 | CHS | Southern Han Chinese | EAS | East Asian Ancestry | CHS | 1000 Genomes on GRCh38,1000 Genomes 30x on GRCh38,1000 Genomes phase 3 release,1000 Genomes phase 1 release |
| **HG00427** | male | SAME122806 | CHS | Southern Han Chinese | EAS | East Asian Ancestry | CHS | 1000 Genomes on GRCh38,1000 Genomes 30x on GRCh38,1000 Genomes phase 3 release,1000 Genomes phase 1 release,90 Han Chinese high coverage genomes |
| **HG00555** | female | SAME122856 | PUR | Puerto Rican | AMR | American Ancestry | PUR | 1000 Genomes 30x on GRCh38,1000 Genomes phase 3 release |
| **HG00446** | female | SAME123880 | CHS | Southern Han Chinese | EAS | East Asian Ancestry | CHS | 1000 Genomes on GRCh38,1000 Genomes 30x on GRCh38,1000 Genomes phase 3 release,1000 Genomes phase 1 release |
| **HG00567** | male | SAME124453 | CHS | Southern Han Chinese | EAS | East Asian Ancestry | CHS | 1000 Genomes 30x on GRCh38,1000 Genomes phase 3 release |
| **HG00453** | female | SAME123113 | CHS | Southern Han Chinese | EAS | East Asian Ancestry | CHS | 1000 Genomes 30x on GRCh38,1000 Genomes phase 3 release |
| **HG00579** | female | SAME123530 | CHS | Southern Han Chinese | EAS | East Asian Ancestry | CHS | 1000 Genomes 30x on GRCh38,1000 Genomes phase 3 release |
| **HG00458** | female | SAME125278 | CHS | Southern Han Chinese | EAS | East Asian Ancestry | CHS | 1000 Genomes on GRCh38,1000 Genomes 30x on GRCh38,1000 Genomes phase 3 release,1000 Genomes phase 1 release |
| **HG00581** | female | SAME124831 | CHS | Southern Han Chinese | EAS | East Asian Ancestry | CHS | 1000 Genomes on GRCh38,1000 Genomes 30x on GRCh38,1000 Genomes phase 3 release,1000 Genomes phase 1 release |
| **HG00465** | male | SAME123498 | CHS | Southern Han Chinese | EAS | East Asian Ancestry | CHS | 1000 Genomes 30x on GRCh38,1000 Genomes phase 3 release |
| **HG00593** | female | SAME125012 | CHS | Southern Han Chinese | EAS | East Asian Ancestry | CHS | 1000 Genomes on GRCh38,1000 Genomes 30x on GRCh38,1000 Genomes phase 3 release,1000 Genomes phase 1 release |
| **HG00472** | male | SAME123339 | CHS | Southern Han Chinese | EAS | East Asian Ancestry | CHS | 1000 Genomes on GRCh38,1000 Genomes 30x on GRCh38,1000 Genomes phase 3 release,1000 Genomes phase 1 release,90 Han Chinese high coverage genomes |
| **HG00598** | male | SAME1839713 | CHS | Southern Han Chinese | EAS | East Asian Ancestry | CHS | 1000 Genomes on GRCh38,1000 Genomes 30x on GRCh38,1000 Genomes phase 3 release |
| **HG00477** | male | SAME123340 | CHS | Southern Han Chinese | EAS | East Asian Ancestry | CHS | 1000 Genomes 30x on GRCh38,1000 Genomes phase 3 release |
| **HG00613** | male | SAME125196 | CHS | Southern Han Chinese | EAS | East Asian Ancestry | CHS | 1000 Genomes on GRCh38,1000 Genomes 30x on GRCh38,1000 Genomes phase 3 release,1000 Genomes phase 1 release |
| **HG00530** | male | SAME124356 | CHS | Southern Han Chinese | EAS | East Asian Ancestry | CHS | 1000 Genomes on GRCh38,1000 Genomes 30x on GRCh38,1000 Genomes phase 3 release,1000 Genomes phase 1 release,90 Han Chinese high coverage genomes |
| **HG01077** | female | SAME1839581 | PUR | Puerto Rican | AMR | American Ancestry | PUR | 1000 Genomes on GRCh38,1000 Genomes 30x on GRCh38,1000 Genomes phase 3 release |
| **HG00535** | male | SAME124348 | CHS | Southern Han Chinese | EAS | East Asian Ancestry | CHS | 1000 Genomes 30x on GRCh38,1000 Genomes phase 3 release |
| **HG01084** | female | SAME125392 | PUR | Puerto Rican | AMR | American Ancestry | PUR | 1000 Genomes 30x on GRCh38,1000 Genomes phase 3 release |
| **HG00732** | female | SAME125396 | PUR | Puerto Rican | AMR | American Ancestry | PUR | 1000 Genomes on GRCh38,1000 Genomes 30x on GRCh38,Human Genome Structural Variation Consortium, Phase 2,The Human Genome Structural Variation Consortiu |
| **HG01089** | female | SAME1839432 | PUR | Puerto Rican | AMR | American Ancestry | PUR | 1000 Genomes on GRCh38,1000 Genomes 30x on GRCh38,1000 Genomes phase 3 release |
| **HG00737** | female | SAME125416 | PUR | Puerto Rican | AMR | American Ancestry | PUR | 1000 Genomes on GRCh38,1000 Genomes 30x on GRCh38,1000 Genomes phase 3 release,1000 Genomes phase 1 release |
| **HG01096** | male | SAME123776 | PUR | Puerto Rican | AMR | American Ancestry | PUR | 1000 Genomes 30x on GRCh38,1000 Genomes phase 3 release |
| **HG00864** | female | SAME123547 | CDX | Dai Chinese | EAS | East Asian Ancestry | CDX | 1000 Genomes on GRCh38,1000 Genomes 30x on GRCh38,Human Genome Structural Variation Consortium, Phase 2,1000 Genomes phase 3 release |
| **HG01104** | male | SAME123672 | PUR | Puerto Rican | AMR | American Ancestry | PUR | 1000 Genomes on GRCh38,1000 Genomes 30x on GRCh38,1000 Genomes phase 3 release,1000 Genomes phase 1 release |
| **HG01109** | male | SAME123669 | PUR | Puerto Rican | AMR | American Ancestry | PUR | 1000 Genomes 30x on GRCh38,1000 Genomes phase 3 release |
| **HG00251** | male | SAME123053 | GBR | British | EUR | European Ancestry | GBR | 1000 Genomes on GRCh38,1000 Genomes 30x on GRCh38,1000 Genomes phase 3 release,1000 Genomes phase 1 release,Geuvadis |
| **HG01111** | female | SAME123486 | PUR | Puerto Rican | AMR | American Ancestry | PUR | 1000 Genomes on GRCh38,1000 Genomes 30x on GRCh38,1000 Genomes phase 3 release,1000 Genomes phase 1 release |
| **HG00256** | male | SAME123048 | GBR | British | EUR | European Ancestry | GBR | 1000 Genomes on GRCh38,1000 Genomes 30x on GRCh38,1000 Genomes phase 3 release,1000 Genomes phase 1 release,Geuvadis |
| **HG00263** | female | SAME123581 | GBR | British | EUR | European Ancestry | GBR | 1000 Genomes on GRCh38,1000 Genomes 30x on GRCh38,1000 Genomes phase 3 release,1000 Genomes phase 1 release,Geuvadis |
| **HG00675** | female | SAME1840076 | CHS | Southern Han Chinese | EAS | East Asian Ancestry | CHS | 1000 Genomes on GRCh38,1000 Genomes 30x on GRCh38,1000 Genomes phase 3 release |
| **HG00268** | female | SAME123574 | FIN | Finnish | EUR | European Ancestry | FIN | 1000 Genomes on GRCh38,1000 Genomes 30x on GRCh38,Human Genome Structural Variation Consortium, Phase 2,1000 Genomes phase 3 release,1000 Genomes ph |
| **HG00694** | male | SAME123154 | CHS | Southern Han Chinese | EAS | East Asian Ancestry | CHS | 1000 Genomes 30x on GRCh38,1000 Genomes phase 3 release |
| **HG00699** | female | SAME123165 | CHS | Southern Han Chinese | EAS | East Asian Ancestry | CHS | 1000 Genomes on GRCh38,1000 Genomes 30x on GRCh38,1000 Genomes phase 3 release,1000 Genomes phase 1 release,90 Han Chinese high coverage genomes |
| **HG00275** | female | SAME123421 | FIN | Finnish | EUR | European Ancestry | FIN | 1000 Genomes on GRCh38,1000 Genomes 30x on GRCh38,1000 Genomes phase 3 release,1000 Genomes phase 1 release,Geuvadis |
| **HG00702** | female | SAME122934 | CHS | Southern Han Chinese | EAS | East Asian Ancestry | CHS | 1000 Genomes on GRCh38,1000 Genomes 30x on GRCh38,1000 Genomes phase 3 release,1000 Genomes phase 1 release |
| **HG00282** | female | SAME122833 | FIN | Finnish | EUR | European Ancestry | FIN | 1000 Genomes on GRCh38,1000 Genomes 30x on GRCh38,1000 Genomes phase 3 release,1000 Genomes phase 1 release,Geuvadis |
| **HG00707** | male | SAME122929 | CHS | Southern Han Chinese | EAS | East Asian Ancestry | CHS | 1000 Genomes on GRCh38,1000 Genomes 30x on GRCh38,1000 Genomes phase 3 release,1000 Genomes phase 1 release |
| **HG01130** | male | SAME1839995 | CLM | Colombian | AMR | American Ancestry | CLM | 1000 Genomes on GRCh38,1000 Genomes 30x on GRCh38,1000 Genomes phase 3 release |
| **HG00612** | female | SAME125197 | CHS | Southern Han Chinese | EAS | East Asian Ancestry | CHS | 1000 Genomes 30x on GRCh38,1000 Genomes phase 3 release |
| **HG01135** | male | SAME123886 | CLM | Colombian | AMR | American Ancestry | CLM | 1000 Genomes 30x on GRCh38,1000 Genomes phase 3 release |
| **HG01142** | male | SAME1839731 | CLM | Colombian | AMR | American Ancestry | CLM | 1000 Genomes on GRCh38,1000 Genomes 30x on GRCh38,1000 Genomes phase 3 release |
| **HG00629** | female | SAME123086 | CHS | Southern Han Chinese | EAS | East Asian Ancestry | CHS | 1000 Genomes on GRCh38,1000 Genomes 30x on GRCh38,1000 Genomes phase 3 release,1000 Genomes phase 1 release |
| **HG01161** | male | SAME1839615 | PUR | Puerto Rican | AMR | American Ancestry | PUR | 1000 Genomes on GRCh38,1000 Genomes 30x on GRCh38,1000 Genomes phase 3 release |
| **HG00631** | male | SAME1839060 | CHS | Southern Han Chinese | EAS | East Asian Ancestry | CHS | 1000 Genomes on GRCh38,1000 Genomes 30x on GRCh38,1000 Genomes phase 3 release |
| **HG01173** | male | SAME124624 | PUR | Puerto Rican | AMR | American Ancestry | PUR | 1000 Genomes on GRCh38,1000 Genomes 30x on GRCh38,1000 Genomes phase 3 release,1000 Genomes phase 1 release |
| **HG00636** | male | SAME123711 | CHS | Southern Han Chinese | EAS | East Asian Ancestry | CHS | 1000 Genomes 30x on GRCh38,1000 Genomes phase 3 release |
| **HG01178** | male | SAME124616 | PUR | Puerto Rican | AMR | American Ancestry | PUR | 1000 Genomes 30x on GRCh38,1000 Genomes phase 3 release |
| **HG00650** | male | SAME123254 | CHS | Southern Han Chinese | EAS | East Asian Ancestry | CHS | 1000 Genomes on GRCh38,1000 Genomes 30x on GRCh38,1000 Genomes phase 3 release,1000 Genomes phase 1 release |
| **HG01046** | female | SAME123167 | CDX | Dai Chinese | EAS | East Asian Ancestry | CDX | 1000 Genomes on GRCh38,1000 Genomes 30x on GRCh38,1000 Genomes phase 3 release |
| **HG00655** | female | SAME123257 | CHS | Southern Han Chinese | EAS | East Asian Ancestry | CHS | 1000 Genomes 30x on GRCh38,1000 Genomes phase 3 release |
| **HG01053** | female | SAME124866 | PUR | Puerto Rican | AMR | American Ancestry | PUR | 1000 Genomes 30x on GRCh38,1000 Genomes phase 3 release |
| **HG00662** | male | SAME123812 | CHS | Southern Han Chinese | EAS | East Asian Ancestry | CHS | 1000 Genomes on GRCh38,1000 Genomes 30x on GRCh38,1000 Genomes phase 3 release,1000 Genomes phase 1 release |
| **HG01058** | female | SAME1840086 | PUR | Puerto Rican | AMR | American Ancestry | PUR | 1000 Genomes on GRCh38,1000 Genomes 30x on GRCh38,1000 Genomes phase 3 release |
| **HG00674** | male | SAME1840080 | CHS | Southern Han Chinese | EAS | East Asian Ancestry | CHS | 1000 Genomes on GRCh38,1000 Genomes 30x on GRCh38,1000 Genomes phase 3 release |
| **HG01060** | male | SAME123015 | PUR | Puerto Rican | AMR | American Ancestry | PUR | 1000 Genomes on GRCh38,1000 Genomes 30x on GRCh38,1000 Genomes phase 3 release,1000 Genomes phase 1 release |
| **HG00693** | female | SAME123683 | CHS | Southern Han Chinese | EAS | East Asian Ancestry | CHS | 1000 Genomes on GRCh38,1000 Genomes 30x on GRCh38,1000 Genomes phase 3 release,1000 Genomes phase 1 release |
| **HG01072** | male | SAME124091 | PUR | Puerto Rican | AMR | American Ancestry | PUR | 1000 Genomes on GRCh38,1000 Genomes 30x on GRCh38,1000 Genomes phase 3 release,1000 Genomes phase 1 release |
| **HG00698** | male | SAME123693 | CHS | Southern Han Chinese | EAS | East Asian Ancestry | CHS | 1000 Genomes on GRCh38,1000 Genomes 30x on GRCh38,1000 Genomes phase 3 release,1000 Genomes phase 1 release,90 Han Chinese high coverage genomes |
| **HG00701** | male | SAME122931 | CHS | Southern Han Chinese | EAS | East Asian Ancestry | CHS | 1000 Genomes on GRCh38,1000 Genomes 30x on GRCh38,1000 Genomes phase 3 release,1000 Genomes phase 1 release |
| **HG01488** | male | SAME123973 | CLM | Colombian | AMR | American Ancestry | CLM | 1000 Genomes on GRCh38,1000 Genomes 30x on GRCh38,1000 Genomes phase 3 release,1000 Genomes phase 1 release |
| **HG00706** | male | SAME122930 | CHS | Southern Han Chinese | EAS | East Asian Ancestry | CHS | 1000 Genomes 30x on GRCh38,1000 Genomes phase 3 release |
| **HG01490** | female | SAME124139 | CLM | Colombian | AMR | American Ancestry | CLM | 1000 Genomes 30x on GRCh38,1000 Genomes phase 3 release |
| **HG01495** | female | SAME124136 | CLM | Colombian | AMR | American Ancestry | CLM | 1000 Genomes on GRCh38,1000 Genomes 30x on GRCh38,1000 Genomes phase 3 release,1000 Genomes phase 1 release |
| **HG01052** | female | SAME123368 | PUR | Puerto Rican | AMR | American Ancestry | PUR | 1000 Genomes on GRCh38,1000 Genomes 30x on GRCh38,1000 Genomes phase 3 release,1000 Genomes phase 1 release |
| **HG01503** | male | SAME124944 | IBS | Iberian,Spanish | EUR | European Ancestry,West Eurasia (SGDP) | IBS,SpanishSGDP | 1000 Genomes on GRCh38,Simons Genome Diversity Project,1000 Genomes 30x on GRCh38,1000 Genomes phase 3 release |
| **HG01064** | female | SAME1839759 | PUR | Puerto Rican | AMR | American Ancestry | PUR | 1000 Genomes on GRCh38,1000 Genomes 30x on GRCh38,1000 Genomes phase 3 release |
| **HG01508** | male | SAME124939 | IBS | Iberian | EUR | European Ancestry | IBS | 1000 Genomes 30x on GRCh38,1000 Genomes phase 3 release |
| **HG01069** | male | SAME122825 | PUR | Puerto Rican | AMR | American Ancestry | PUR | 1000 Genomes on GRCh38,1000 Genomes 30x on GRCh38,1000 Genomes phase 3 release,1000 Genomes phase 1 release |
| **HG01565** | male | SAME124566 | PEL | Peruvian | AMR | American Ancestry | PEL | 1000 Genomes on GRCh38,1000 Genomes 30x on GRCh38,1000 Genomes phase 3 release |
| **HG01071** | female | SAME123009 | PUR | Puerto Rican | AMR | American Ancestry | PUR | 1000 Genomes 30x on GRCh38,1000 Genomes phase 3 release |
| **HG01572** | female | SAME124705 | PEL | Peruvian | AMR | American Ancestry | PEL | 1000 Genomes on GRCh38,1000 Genomes 30x on GRCh38,1000 Genomes phase 3 release |
| **HG01191** | female | SAME123651 | PUR | Puerto Rican | AMR | American Ancestry | PUR | 1000 Genomes on GRCh38,1000 Genomes 30x on GRCh38,1000 Genomes phase 3 release,1000 Genomes phase 1 release |
| **HG01577** | male | SAME124349 | PEL | Peruvian | AMR | American Ancestry | PEL | 1000 Genomes on GRCh38,1000 Genomes 30x on GRCh38,1000 Genomes phase 3 release |
| **HG01204** | male | SAME124297 | PUR | Puerto Rican | AMR | American Ancestry | PUR | 1000 Genomes on GRCh38,1000 Genomes 30x on GRCh38,1000 Genomes phase 3 release,1000 Genomes phase 1 release |
| **HG01589** | male | SAME1839859 | PJL | Punjabi | SAS | South Asian Ancestry | PJL | 1000 Genomes on GRCh38,1000 Genomes 30x on GRCh38,1000 Genomes phase 3 release |
| **HG01242** | female | SAME125064 | PUR | Puerto Rican | AMR | American Ancestry | PUR | 1000 Genomes on GRCh38,1000 Genomes 30x on GRCh38,1000 Genomes phase 3 release,1000 Genomes phase 1 release |
| **HG01596** | male | SAME125227 | KHV | Kinh Vietnamese | EAS | East Asian Ancestry | KHV | 1000 Genomes on GRCh38,1000 Genomes 30x on GRCh38,Human Genome Structural Variation Consortium, Phase 2,1000 Genomes phase 3 release |
| **HG01247** | male | SAME125067 | PUR | Puerto Rican | AMR | American Ancestry | PUR | 1000 Genomes on GRCh38,1000 Genomes 30x on GRCh38,1000 Genomes phase 3 release,1000 Genomes phase 1 release |
| **HG01349** | female | SAME124865 | CLM | Colombian | AMR | American Ancestry | CLM | 1000 Genomes 30x on GRCh38,1000 Genomes phase 3 release |
| **HG01254** | female | SAME125285 | CLM | Colombian | AMR | American Ancestry | CLM | 1000 Genomes on GRCh38,1000 Genomes 30x on GRCh38,1000 Genomes phase 3 release,1000 Genomes phase 1 release |
| **HG01351** | female | SAME124683 | CLM | Colombian | AMR | American Ancestry | CLM | 1000 Genomes on GRCh38,1000 Genomes 30x on GRCh38,1000 Genomes phase 3 release,1000 Genomes phase 1 release |
| **HG01259** | male | SAME125280 | CLM | Colombian | AMR | American Ancestry | CLM | 1000 Genomes on GRCh38,1000 Genomes 30x on GRCh38,1000 Genomes phase 3 release,1000 Genomes phase 1 release |
| **HG01356** | male | SAME124682 | CLM | Colombian | AMR | American Ancestry | CLM | 1000 Genomes on GRCh38,1000 Genomes 30x on GRCh38,1000 Genomes phase 3 release,1000 Genomes phase 1 release |
| **HG01261** | male | SAME122814 | CLM | Colombian | AMR | American Ancestry | CLM | 1000 Genomes 30x on GRCh38,1000 Genomes phase 3 release |
| **HG01363** | female | SAME1839059 | CLM | Colombian | AMR | American Ancestry | CLM | 1000 Genomes on GRCh38,1000 Genomes 30x on GRCh38,1000 Genomes phase 3 release |
| **HG01273** | male | SAME124046 | CLM | Colombian | AMR | American Ancestry | CLM | 1000 Genomes 30x on GRCh38,1000 Genomes phase 3 release |
| **HG01375** | female | SAME124290 | CLM | Colombian | AMR | American Ancestry | CLM | 1000 Genomes on GRCh38,1000 Genomes 30x on GRCh38,1000 Genomes phase 3 release,1000 Genomes phase 1 release |
| **HG01278** | female | SAME123003 | CLM | Colombian | AMR | American Ancestry | CLM | 1000 Genomes on GRCh38,1000 Genomes 30x on GRCh38,1000 Genomes phase 3 release,1000 Genomes phase 1 release |
| **HG01280** | male | SAME1839113 | CLM | Colombian | AMR | American Ancestry | CLM | 1000 Genomes on GRCh38,1000 Genomes 30x on GRCh38,1000 Genomes phase 3 release |
| **HG01402** | male | SAME1839873 | PUR | Puerto Rican | AMR | American Ancestry | PUR | 1000 Genomes on GRCh38,1000 Genomes 30x on GRCh38,1000 Genomes phase 3 release |
| **HG01083** | female | SAME125397 | PUR | Puerto Rican | AMR | American Ancestry | PUR | 1000 Genomes on GRCh38,1000 Genomes 30x on GRCh38,1000 Genomes phase 3 release,1000 Genomes phase 1 release |
| **HG01414** | female | SAME1839703 | PUR | Puerto Rican | AMR | American Ancestry | PUR | 1000 Genomes on GRCh38,1000 Genomes 30x on GRCh38,1000 Genomes phase 3 release |
| **HG01088** | male | SAME1839361 | PUR | Puerto Rican | AMR | American Ancestry | PUR | 1000 Genomes on GRCh38,1000 Genomes 30x on GRCh38,1000 Genomes phase 3 release |
| **HG01433** | male | SAME1839252 | CLM | Colombian | AMR | American Ancestry | CLM | 1000 Genomes 30x on GRCh38 |
| **HG01438** | female | SAME124971 | CLM | Colombian | AMR | American Ancestry | CLM | 1000 Genomes on GRCh38,1000 Genomes 30x on GRCh38,1000 Genomes phase 3 release,1000 Genomes phase 1 release |
| **HG01095** | female | SAME122926 | PUR | Puerto Rican | AMR | American Ancestry | PUR | 1000 Genomes on GRCh38,1000 Genomes 30x on GRCh38,1000 Genomes phase 3 release,1000 Genomes phase 1 release |
| **HG01440** | male | SAME123237 | CLM | Colombian | AMR | American Ancestry | CLM | 1000 Genomes on GRCh38,1000 Genomes 30x on GRCh38,1000 Genomes phase 3 release,1000 Genomes phase 1 release |
| **HG01103** | female | SAME123679 | PUR | Puerto Rican | AMR | American Ancestry | PUR | 1000 Genomes 30x on GRCh38,1000 Genomes phase 3 release |
| **HG01281** | female | SAME1839107 | CLM | Colombian | AMR | American Ancestry | CLM | 1000 Genomes on GRCh38,1000 Genomes 30x on GRCh38,1000 Genomes phase 3 release |
| **HG01108** | female | SAME123103 | PUR | Puerto Rican | AMR | American Ancestry | PUR | 1000 Genomes on GRCh38,1000 Genomes 30x on GRCh38,1000 Genomes phase 3 release,1000 Genomes phase 1 release |
| **HG01286** | male | SAME1839176 | PUR | Puerto Rican | AMR | American Ancestry | PUR | 1000 Genomes on GRCh38,1000 Genomes 30x on GRCh38,1000 Genomes phase 3 release |

| | | | | | | | | |
|---|---|---|---|---|---|---|---|---|
| **HG01110** | male | SAME123487 | PUR | Puerto Rican | AMR | American Ancestry | PUR | 1000 Genomes on GRCh38,1000 Genomes 30x on GRCh38,1000 Genomes phase 3 release,1000 Genomes phase 1 release |
| **HG01122** | female | SAME1840203 | CLM | Colombian | AMR | American Ancestry | CLM | 1000 Genomes on GRCh38,1000 Genomes 30x on GRCh38,1000 Genomes phase 3 release |
| **HG01325** | male | SAME1839604 | PUR | Puerto Rican | AMR | American Ancestry | PUR | 1000 Genomes on GRCh38,1000 Genomes 30x on GRCh38,1000 Genomes phase 3 release |
| **HG01350** | male | SAME124684 | CLM | Colombian | AMR | American Ancestry | CLM | 1000 Genomes on GRCh38,1000 Genomes 30x on GRCh38,1000 Genomes phase 3 release,1000 Genomes phase 1 release |
| **HG01344** | male | SAME124862 | CLM | Colombian | AMR | American Ancestry | CLM | 1000 Genomes on GRCh38,1000 Genomes 30x on GRCh38,1000 Genomes phase 3 release,1000 Genomes phase 1 release |
| **HG01355** | female | SAME124679 | CLM | Colombian | AMR | American Ancestry | CLM | 1000 Genomes 30x on GRCh38,1000 Genomes phase 3 release |
| **HG01604** | female | SAME123460 | IBS | Iberian | EUR | European Ancestry | IBS | 1000 Genomes 30x on GRCh38,1000 Genomes phase 3 release |
| **HG01362** | male | SAME1839066 | CLM | Colombian | AMR | American Ancestry | CLM | 1000 Genomes on GRCh38,1000 Genomes 30x on GRCh38,1000 Genomes phase 3 release |
| **HG01609** | male | SAME123461 | IBS | Iberian | EUR | European Ancestry | IBS | 1000 Genomes 30x on GRCh38,1000 Genomes phase 3 release |
| **HG01367** | female | SAME125266 | CLM | Colombian | AMR | American Ancestry | CLM | 1000 Genomes 30x on GRCh38,1000 Genomes phase 3 release |
| **HG01611** | male | SAME123624 | IBS | Iberian | EUR | European Ancestry | IBS | 1000 Genomes 30x on GRCh38,1000 Genomes phase 3 release |
| **HG01374** | male | SAME124291 | CLM | Colombian | AMR | American Ancestry | CLM | 1000 Genomes on GRCh38,1000 Genomes 30x on GRCh38,1000 Genomes phase 3 release,1000 Genomes phase 1 release |
| **HG01616** | female | SAME123629 | IBS | Iberian | EUR | European Ancestry | IBS | 1000 Genomes 30x on GRCh38,1000 Genomes phase 3 release |
| **HG01379** | female | SAME124283 | CLM | Colombian | AMR | American Ancestry | CLM | 1000 Genomes 30x on GRCh38,1000 Genomes phase 3 release |
| **HG01623** | female | SAME123845 | IBS | Iberian | EUR | European Ancestry | IBS | 1000 Genomes on GRCh38,1000 Genomes 30x on GRCh38,1000 Genomes phase 3 release,1000 Genomes phase 1 release |
| **HG01393** | female | SAME1840222 | PUR | Puerto Rican | AMR | American Ancestry | PUR | 1000 Genomes on GRCh38,1000 Genomes 30x on GRCh38,1000 Genomes phase 3 release |
| **HG01134** | female | SAME123887 | CLM | Colombian | AMR | American Ancestry | CLM | 1000 Genomes on GRCh38,1000 Genomes 30x on GRCh38,1000 Genomes phase 3 release,1000 Genomes phase 1 release |
| **HG01139** | male | SAME123876 | CLM | Colombian | AMR | American Ancestry | CLM | 1000 Genomes on GRCh38,1000 Genomes 30x on GRCh38,1000 Genomes phase 3 release,1000 Genomes phase 1 release |
| **HG01457** | male | SAME123428 | CLM | Colombian | AMR | American Ancestry | CLM | 1000 Genomes 30x on GRCh38,1000 Genomes phase 3 release |
| **HG01141** | female | SAME124423 | CLM | Colombian | AMR | American Ancestry | CLM | 1000 Genomes 30x on GRCh38,1000 Genomes phase 3 release |
| **HG01464** | male | SAME122883 | CLM | Colombian | AMR | American Ancestry | CLM | 1000 Genomes on GRCh38,1000 Genomes 30x on GRCh38,1000 Genomes phase 3 release,1000 Genomes phase 1 release |
| **HG01172** | male | SAME124625 | PUR | Puerto Rican | AMR | American Ancestry | PUR | 1000 Genomes 30x on GRCh38,1000 Genomes phase 3 release |
| **HG01177** | female | SAME124620 | PUR | Puerto Rican | AMR | American Ancestry | PUR | 1000 Genomes on GRCh38,1000 Genomes 30x on GRCh38,1000 Genomes phase 3 release,1000 Genomes phase 1 release |
| **HG01628** | female | SAME123837 | IBS | Iberian | EUR | European Ancestry | IBS | 1000 Genomes on GRCh38,1000 Genomes 30x on GRCh38,1000 Genomes phase 3 release |
| **HG01184** | female | SAME124321 | PUR | Puerto Rican | AMR | American Ancestry | PUR | 1000 Genomes 30x on GRCh38,1000 Genomes phase 3 release |
| **HG01630** | male | SAME124034 | IBS | Iberian | EUR | European Ancestry | IBS | 1000 Genomes on GRCh38,1000 Genomes 30x on GRCh38,1000 Genomes phase 3 release |
| **HG01189** | male | SAME124878 | PUR | Puerto Rican | AMR | American Ancestry | PUR | 1000 Genomes 30x on GRCh38,1000 Genomes phase 3 release |
| **HG01673** | female | SAME123289 | IBS | Iberian | EUR | European Ancestry | IBS | 1000 Genomes on GRCh38,1000 Genomes 30x on GRCh38,1000 Genomes phase 3 release |
| **HG01678** | male | SAME123292 | IBS | Iberian | EUR | European Ancestry | IBS | 1000 Genomes on GRCh38,1000 Genomes 30x on GRCh38,1000 Genomes phase 3 release |
| **HG01680** | male | SAME124656 | IBS | Iberian | EUR | European Ancestry | IBS | 1000 Genomes on GRCh38,1000 Genomes 30x on GRCh38,1000 Genomes phase 3 release |
| **HG01494** | male | SAME124135 | CLM | Colombian | AMR | American Ancestry | CLM | 1000 Genomes on GRCh38,1000 Genomes 30x on GRCh38,1000 Genomes phase 3 release,1000 Genomes phase 1 release |
| **HG01510** | female | SAME124771 | IBS | Iberian | EUR | European Ancestry | IBS | 1000 Genomes on GRCh38,1000 Genomes 30x on GRCh38,1000 Genomes phase 3 release |
| **HG01499** | male | SAME124143 | CLM | Colombian | AMR | American Ancestry | CLM | 1000 Genomes 30x on GRCh38,1000 Genomes phase 3 release |
| **HG01515** | male | SAME124774 | IBS | Iberian | EUR | European Ancestry | IBS | 1000 Genomes on GRCh38,1000 Genomes 30x on GRCh38,1000 Genomes phase 3 release,1000 Genomes phase 1 release |
| **HG01502** | male | SAME124943 | IBS | Iberian | EUR | European Ancestry | IBS | 1000 Genomes 30x on GRCh38,1000 Genomes phase 3 release |
| **HG01522** | female | SAME125321 | IBS | Iberian | EUR | European Ancestry | IBS | 1000 Genomes on GRCh38,1000 Genomes 30x on GRCh38,1000 Genomes phase 3 release,1000 Genomes phase 1 release |
| **HG01507** | female | SAME124948 | IBS | Iberian | EUR | European Ancestry | IBS | 1000 Genomes on GRCh38,1000 Genomes 30x on GRCh38,1000 Genomes phase 3 release |
| **HG01527** | male | SAME125326 | IBS | Iberian | EUR | European Ancestry | IBS | 1000 Genomes on GRCh38,1000 Genomes 30x on GRCh38,1000 Genomes phase 3 release |
| **HG01305** | male | SAME1840020 | PUR | Puerto Rican | AMR | American Ancestry | PUR | 1000 Genomes on GRCh38,1000 Genomes 30x on GRCh38,1000 Genomes phase 3 release |
| **HG01894** | female | SAME1840357 | ACB | African Caribbean | AFR | African Ancestry | ACB | 1000 Genomes on GRCh38,1000 Genomes 30x on GRCh38,1000 Genomes phase 3 release |
| **HG01312** | female | SAME1839791 | PUR | Puerto Rican | AMR | American Ancestry | PUR | 1000 Genomes on GRCh38,1000 Genomes 30x on GRCh38,1000 Genomes phase 3 release |
| **HG01914** | male | SAME1839768 | ACB | African Caribbean | AFR | African Ancestry | ACB | 1000 Genomes on GRCh38,1000 Genomes 30x on GRCh38,1000 Genomes phase 3 release |
| **HG01919** | female | SAME123239 | PEL | Peruvian | AMR | American Ancestry | PEL | 1000 Genomes 30x on GRCh38,1000 Genomes phase 3 release |
| **HG01343** | female | SAME124857 | CLM | Colombian | AMR | American Ancestry | CLM | 1000 Genomes 30x on GRCh38,1000 Genomes phase 3 release |
| **HG01921** | female | SAME123790 | PEL | Peruvian | AMR | American Ancestry | PEL | 1000 Genomes on GRCh38,1000 Genomes 30x on GRCh38,1000 Genomes phase 3 release |
| **HG01348** | female | SAME124864 | CLM | Colombian | AMR | American Ancestry | CLM | 1000 Genomes on GRCh38,1000 Genomes 30x on GRCh38,1000 Genomes phase 3 release |
| **HG01868** | female | SAME123193 | KHV | Kinh Vietnamese | EAS | East Asian Ancestry | KHV | 1000 Genomes on GRCh38,1000 Genomes 30x on GRCh38,1000 Genomes phase 3 release |
| **HG01398** | male | SAME1839950 | PUR | Puerto Rican | AMR | American Ancestry | PUR | 1000 Genomes on GRCh38,1000 Genomes 30x on GRCh38,1000 Genomes phase 3 release |
| **HG01870** | female | SAME123397 | KHV | Kinh Vietnamese | EAS | East Asian Ancestry | KHV | 1000 Genomes on GRCh38,1000 Genomes 30x on GRCh38,1000 Genomes phase 3 release |
| **HG01413** | male | SAME1839612 | PUR | Puerto Rican | AMR | American Ancestry | PUR | 1000 Genomes on GRCh38,1000 Genomes 30x on GRCh38,1000 Genomes phase 3 release |
| **HG01882** | male | SAME122849 | ACB | African Caribbean | AFR | African Ancestry | ACB | 1000 Genomes on GRCh38,1000 Genomes 30x on GRCh38,1000 Genomes phase 3 release |
| **HG01432** | female | SAME1839256 | CLM | Colombian | AMR | American Ancestry | CLM | 1000 Genomes on GRCh38,1000 Genomes 30x on GRCh38,1000 Genomes phase 3 release |
| **HG01887** | male | SAME122846 | ACB | African Caribbean | AFR | African Ancestry | ACB | 1000 Genomes 30x on GRCh38,1000 Genomes phase 3 release |
| **HG01437** | male | SAME124966 | CLM | Colombian | AMR | American Ancestry | CLM | 1000 Genomes on GRCh38,1000 Genomes 30x on GRCh38,1000 Genomes phase 3 release,1000 Genomes phase 1 release |
| **HG01926** | male | SAME123783 | PEL | Peruvian | AMR | American Ancestry | PEL | 1000 Genomes on GRCh38,1000 Genomes 30x on GRCh38,1000 Genomes phase 3 release |
| **HG01444** | female | SAME1839026 | CLM | Colombian | AMR | American Ancestry | CLM | 1000 Genomes on GRCh38,1000 Genomes 30x on GRCh38,1000 Genomes phase 3 release |
| **HG01933** | female | SAME123586 | PEL | Peruvian | AMR | American Ancestry | PEL | 1000 Genomes on GRCh38,1000 Genomes 30x on GRCh38,1000 Genomes phase 3 release |
| **HG01456** | female | SAME123427 | CLM | Colombian | AMR | American Ancestry | CLM | 1000 Genomes on GRCh38,1000 Genomes 30x on GRCh38,1000 Genomes phase 3 release,1000 Genomes phase 1 release |
| **HG01938** | male | SAME123593 | PEL | Peruvian | AMR | American Ancestry | PEL | 1000 Genomes on GRCh38,1000 Genomes 30x on GRCh38,1000 Genomes phase 3 release |
| **HG01463** | female | SAME123537 | CLM | Colombian | AMR | American Ancestry | CLM | 1000 Genomes 30x on GRCh38,1000 Genomes phase 3 release |
| **HG01940** | female | SAME124148 | PEL | Peruvian | AMR | American Ancestry | PEL | 1000 Genomes 30x on GRCh38,1000 Genomes phase 3 release |
| **HG01468** | female | SAME1840382 | CLM | Colombian | AMR | American Ancestry | CLM | 1000 Genomes on GRCh38,1000 Genomes 30x on GRCh38,1000 Genomes phase 3 release |
| **HG01945** | female | SAME124153 | PEL | Peruvian | AMR | American Ancestry | PEL | 1000 Genomes on GRCh38,1000 Genomes 30x on GRCh38,1000 Genomes phase 3 release |
| **HG01514** | female | SAME124775 | IBS | Iberian | EUR | European Ancestry | IBS | 1000 Genomes 30x on GRCh38,1000 Genomes phase 3 release |
| **HG01748** | male | SAME125191 | IBS | Iberian | EUR | European Ancestry | IBS | 1000 Genomes 30x on GRCh38,1000 Genomes phase 3 release |
| **HG01519** | female | SAME124765 | IBS | Iberian | EUR | European Ancestry | IBS | 1000 Genomes on GRCh38,1000 Genomes 30x on GRCh38,1000 Genomes phase 3 release,1000 Genomes phase 1 release |
| **HG01755** | female | SAME124986 | IBS | Iberian | EUR | European Ancestry | IBS | 1000 Genomes 30x on GRCh38,1000 Genomes phase 3 release |
| **HG01521** | male | SAME125324 | IBS | Iberian | EUR | European Ancestry | IBS | 1000 Genomes on GRCh38,1000 Genomes 30x on GRCh38,1000 Genomes phase 3 release,1000 Genomes phase 1 release |
| **HG01762** | female | SAME124811 | IBS | Iberian | EUR | European Ancestry | IBS | 1000 Genomes on GRCh38,1000 Genomes 30x on GRCh38,1000 Genomes phase 3 release |
| **HG01526** | male | SAME125325 | IBS | Iberian | EUR | European Ancestry | IBS | 1000 Genomes 30x on GRCh38,1000 Genomes phase 3 release |
| **HG01767** | male | SAME1839671 | IBS | Iberian | EUR | European Ancestry | IBS | 1000 Genomes on GRCh38,1000 Genomes 30x on GRCh38,1000 Genomes phase 3 release |
| **HG01538** | female | SAME125141 | IBS | Iberian | EUR | European Ancestry | IBS | 1000 Genomes 30x on GRCh38,1000 Genomes phase 3 release |
| **HG01774** | male | SAME124634 | IBS | Iberian | EUR | European Ancestry | IBS | 1000 Genomes 30x on GRCh38,1000 Genomes phase 3 release |
| **HG01552** | male | SAME124007 | CLM | Colombian | AMR | American Ancestry | CLM | 1000 Genomes 30x on GRCh38,1000 Genomes phase 3 release |
| **HG01779** | female | SAME124629 | IBS | Iberian | EUR | European Ancestry | IBS | 1000 Genomes on GRCh38,1000 Genomes 30x on GRCh38,1000 Genomes phase 3 release |
| **HG01571** | male | SAME124346 | PEL | Peruvian | AMR | American Ancestry | PEL | 1000 Genomes on GRCh38,1000 Genomes 30x on GRCh38,1000 Genomes phase 3 release |
| **HG01781** | male | SAME124921 | IBS | Iberian | EUR | European Ancestry | IBS | 1000 Genomes on GRCh38,1000 Genomes 30x on GRCh38,1000 Genomes phase 3 release |
| **HG01583** | male | SAME1839895 | PJL | Punjabi | SAS | South Asian Ancestry | PJL | 1000 Genomes on GRCh38,1000 Genomes 30x on GRCh38,1000 Genomes phase 3 release |
| **HG01786** | female | SAME1839312 | IBS | Iberian | EUR | European Ancestry | IBS | 1000 Genomes on GRCh38,1000 Genomes 30x on GRCh38,1000 Genomes phase 3 release |
| **HG01952** | male | SAME123975 | PEL | Peruvian | AMR | American Ancestry | PEL | 1000 Genomes 30x on GRCh38,1000 Genomes phase 3 release |
| **HG01595** | female | SAME123569 | KHV | Kinh Vietnamese | EAS | East Asian Ancestry | KHV | 1000 Genomes on GRCh38,1000 Genomes 30x on GRCh38,1000 Genomes phase 3 release |
| **HG01969** | female | SAME124533 | PEL | Peruvian | AMR | American Ancestry | PEL | 1000 Genomes 30x on GRCh38,1000 Genomes phase 3 release |
| **HG01867** | male | SAME123194 | KHV | Kinh Vietnamese | EAS | East Asian Ancestry | KHV | 1000 Genomes on GRCh38,1000 Genomes 30x on GRCh38,1000 Genomes phase 3 release |
| **HG01971** | female | SAME124326 | PEL | Peruvian | AMR | American Ancestry | PEL | 1000 Genomes on GRCh38,1000 Genomes 30x on GRCh38,1000 Genomes phase 3 release |
| **HG01874** | female | SAME123396 | KHV | Kinh Vietnamese | EAS | East Asian Ancestry | KHV | 1000 Genomes on GRCh38,1000 Genomes 30x on GRCh38,1000 Genomes phase 3 release |
| **HG01685** | female | SAME124659 | IBS | Iberian | EUR | European Ancestry | IBS | 1000 Genomes on GRCh38,1000 Genomes 30x on GRCh38,1000 Genomes phase 3 release |
| **HG01879** | male | SAME1839037 | ACB | African Caribbean | AFR | African Ancestry | ACB | 1000 Genomes on GRCh38,1000 Genomes 30x on GRCh38,1000 Genomes phase 3 release |
| **HG01697** | female | SAME1839223 | IBS | Iberian | EUR | European Ancestry | IBS | 1000 Genomes on GRCh38,1000 Genomes 30x on GRCh38,1000 Genomes phase 3 release |
| **HG01881** | female | SAME1839193 | ACB | African Caribbean | AFR | African Ancestry | ACB | 1000 Genomes 30x on GRCh38,1000 Genomes phase 3 release |
| **HG01700** | male | SAME123261 | IBS | Iberian | EUR | European Ancestry | IBS | 1000 Genomes on GRCh38,1000 Genomes 30x on GRCh38,1000 Genomes phase 3 release |
| **HG01886** | female | SAME122847 | ACB | African Caribbean | AFR | African Ancestry | ACB | 1000 Genomes on GRCh38,1000 Genomes 30x on GRCh38,1000 Genomes phase 3 release |
| **HG01705** | male | SAME123266 | IBS | Iberian | EUR | European Ancestry | IBS | 1000 Genomes on GRCh38,1000 Genomes 30x on GRCh38,1000 Genomes phase 3 release |
| **HG01709** | male | SAME123229 | IBS | Iberian | EUR | European Ancestry | IBS | 1000 Genomes on GRCh38,1000 Genomes 30x on GRCh38,1000 Genomes phase 3 release |
| **HG01844** | male | SAME123556 | KHV | Kinh Vietnamese | EAS | East Asian Ancestry | KHV | 1000 Genomes on GRCh38,1000 Genomes 30x on GRCh38,1000 Genomes phase 3 release |
| **HG01711** | female | SAME123083 | IBS | Iberian | EUR | European Ancestry | IBS | 1000 Genomes 30x on GRCh38,1000 Genomes phase 3 release |
| **HG01849** | male | SAME125110 | KHV | Kinh Vietnamese | EAS | East Asian Ancestry | KHV | 1000 Genomes on GRCh38,1000 Genomes 30x on GRCh38,1000 Genomes phase 3 release |
| **HG01747** | male | SAME125181 | IBS | Iberian | EUR | European Ancestry | IBS | 1000 Genomes on GRCh38,1000 Genomes 30x on GRCh38,1000 Genomes phase 3 release |
| **HG01851** | female | SAME123753 | KHV | Kinh Vietnamese | EAS | East Asian Ancestry | KHV | 1000 Genomes on GRCh38,1000 Genomes 30x on GRCh38,1000 Genomes phase 3 release |
| **HG01761** | male | SAME124810 | IBS | Iberian | EUR | European Ancestry | IBS | 1000 Genomes on GRCh38,1000 Genomes 30x on GRCh38,1000 Genomes phase 3 release |
| **HG01863** | female | SAME123198 | KHV | Kinh Vietnamese | EAS | East Asian Ancestry | KHV | 1000 Genomes on GRCh38,1000 Genomes 30x on GRCh38,1000 Genomes phase 3 release |
| **HG01766** | female | SAME1839673 | IBS | Iberian | EUR | European Ancestry | IBS | 1000 Genomes on GRCh38,1000 Genomes 30x on GRCh38,1000 Genomes phase 3 release |
| **HG01798** | female | SAME124228 | CDX | Dai Chinese | EAS | East Asian Ancestry | CDX | 1000 Genomes on GRCh38,1000 Genomes 30x on GRCh38,1000 Genomes phase 3 release |
| **HG01773** | female | SAME124635 | IBS | Iberian | EUR | European Ancestry | IBS | 1000 Genomes on GRCh38,1000 Genomes 30x on GRCh38,1000 Genomes phase 3 release |
| **HG01801** | female | SAME124305 | CDX | Dai Chinese | EAS | East Asian Ancestry | CDX | 1000 Genomes on GRCh38,1000 Genomes 30x on GRCh38,1000 Genomes phase 3 release |
| **HG01603** | male | SAME123453 | IBS | Iberian | EUR | European Ancestry | IBS | 1000 Genomes on GRCh38,1000 Genomes 30x on GRCh38,1000 Genomes phase 3 release |
| **HG01806** | female | SAME124304 | CDX | Dai Chinese | EAS | East Asian Ancestry | CDX | 1000 Genomes on GRCh38,1000 Genomes 30x on GRCh38,1000 Genomes phase 3 release |
| **HG01608** | male | SAME123462 | IBS | Iberian | EUR | European Ancestry | IBS | 1000 Genomes on GRCh38,1000 Genomes 30x on GRCh38,1000 Genomes phase 3 release |
| **HG01813** | female | SAME124518 | CDX | Dai Chinese | EAS | East Asian Ancestry | CDX | 1000 Genomes on GRCh38,1000 Genomes 30x on GRCh38,1000 Genomes phase 3 release |
| **HG01610** | male | SAME123623 | IBS | Iberian | EUR | European Ancestry | IBS | 1000 Genomes on GRCh38,1000 Genomes 30x on GRCh38,1000 Genomes phase 3 release |
| **HG02006** | female | SAME1840279 | PEL | Peruvian | AMR | American Ancestry | PEL | 1000 Genomes on GRCh38,1000 Genomes 30x on GRCh38,1000 Genomes phase 3 release |
| **HG01615** | male | SAME123628 | IBS | Iberian | EUR | European Ancestry | IBS | 1000 Genomes on GRCh38,1000 Genomes 30x on GRCh38,1000 Genomes phase 3 release |

| | | | | | | | | |
|---|---|---|---|---|---|---|---|---|
| **HG02013** | male | SAME1840396 | ACB | African Caribbean | AFR | African Ancestry | ACB | 1000 Genomes on GRCh38,1000 Genomes 30x on GRCh38,1000 Genomes phase 3 release |
| **HG01622** | female | SAME123846 | IBS | Iberian | EUR | European Ancestry | IBS | 1000 Genomes 30x on GRCh38,1000 Genomes phase 3 release |
| **HG02018** | female | SAME123451 | KHV | Kinh Vietnamese | EAS | East Asian Ancestry | KHV | 1000 Genomes 30x on GRCh38,Human Genome Structural Variation Consortium, Phase 2,1000 Genomes phase 3 release |
| **HG01627** | male | SAME123841 | IBS | Iberian | EUR | European Ancestry | IBS | 1000 Genomes 30x on GRCh38,1000 Genomes phase 3 release |
| **HG02020** | male | SAME125168 | KHV | Kinh Vietnamese | EAS | East Asian Ancestry | KHV | 1000 Genomes on GRCh38,1000 Genomes 30x on GRCh38,1000 Genomes phase 3 release |
| **HG01778** | male | SAME124630 | IBS | Iberian | EUR | European Ancestry | IBS | 1000 Genomes 30x on GRCh38,1000 Genomes phase 3 release |
| **HG01976** | female | SAME124325 | PEL | Peruvian | AMR | American Ancestry | PEL | 1000 Genomes on GRCh38,1000 Genomes 30x on GRCh38,1000 Genomes phase 3 release |
| **HG01780** | male | SAME124428 | IBS | Iberian | EUR | European Ancestry | IBS | 1000 Genomes 30x on GRCh38,1000 Genomes phase 3 release |
| **HG01983** | female | SAME1840106 | PEL | Peruvian | AMR | American Ancestry | PEL | 1000 Genomes on GRCh38,1000 Genomes 30x on GRCh38,1000 Genomes phase 3 release |
| **HG01785** | male | SAME1839313 | IBS | Iberian | EUR | European Ancestry | IBS | 1000 Genomes on GRCh38,1000 Genomes 30x on GRCh38,1000 Genomes phase 3 release |
| **HG01988** | male | SAME1840175 | ACB | African Caribbean | AFR | African Ancestry | ACB | 1000 Genomes on GRCh38,1000 Genomes 30x on GRCh38,1000 Genomes phase 3 release |
| **HG01797** | female | SAME124219 | CDX | Dai Chinese | EAS | East Asian Ancestry | CDX | 1000 Genomes on GRCh38,1000 Genomes 30x on GRCh38,1000 Genomes phase 3 release |
| **HG01990** | male | SAME1840235 | ACB | African Caribbean | AFR | African Ancestry | ACB | 1000 Genomes on GRCh38,1000 Genomes 30x on GRCh38,1000 Genomes phase 3 release |
| **HG01684** | female | SAME124660 | IBS | Iberian | EUR | European Ancestry | IBS | 1000 Genomes on GRCh38,1000 Genomes 30x on GRCh38,1000 Genomes phase 3 release |
| **HG01696** | male | SAME124715 | IBS | Iberian | EUR | European Ancestry | IBS | 1000 Genomes 30x on GRCh38,1000 Genomes phase 3 release |
| **HG02025** | female | SAME125335 | KHV | Kinh Vietnamese | EAS | East Asian Ancestry | KHV | 1000 Genomes on GRCh38,1000 Genomes 30x on GRCh38,1000 Genomes phase 3 release |
| **HG01704** | female | SAME123265 | IBS | Iberian | EUR | European Ancestry | IBS | 1000 Genomes on GRCh38,1000 Genomes 30x on GRCh38,1000 Genomes phase 3 release |
| **HG02032** | male | SAME1839959 | KHV | Kinh Vietnamese | EAS | East Asian Ancestry | KHV | 1000 Genomes on GRCh38,1000 Genomes 30x on GRCh38,1000 Genomes phase 3 release |
| **HG01843** | female | SAME123553 | KHV | Kinh Vietnamese | EAS | East Asian Ancestry | KHV | 1000 Genomes on GRCh38,1000 Genomes 30x on GRCh38,1000 Genomes phase 3 release |
| **HG02049** | female | SAME123773 | KHV | Kinh Vietnamese | EAS | East Asian Ancestry | KHV | 1000 Genomes on GRCh38,1000 Genomes 30x on GRCh38,1000 Genomes phase 3 release |
| **HG01848** | female | SAME123548 | KHV | Kinh Vietnamese | EAS | East Asian Ancestry | KHV | 1000 Genomes on GRCh38,1000 Genomes 30x on GRCh38,1000 Genomes phase 3 release |
| **HG02051** | male | SAME1839549 | ACB | African Caribbean | AFR | African Ancestry | ACB | 1000 Genomes on GRCh38,1000 Genomes 30x on GRCh38,1000 Genomes phase 3 release |
| **HG01850** | female | SAME123752 | KHV | Kinh Vietnamese | EAS | East Asian Ancestry | KHV | 1000 Genomes on GRCh38,1000 Genomes 30x on GRCh38,1000 Genomes phase 3 release |
| **HG02082** | male | SAME124950 | KHV | Kinh Vietnamese | EAS | East Asian Ancestry | KHV | 1000 Genomes on GRCh38,1000 Genomes 30x on GRCh38,1000 Genomes phase 3 release |
| **HG01855** | female | SAME123749 | KHV | Kinh Vietnamese | EAS | East Asian Ancestry | KHV | 1000 Genomes on GRCh38,1000 Genomes 30x on GRCh38,1000 Genomes phase 3 release |
| **HG02087** | female | SAME124678 | KHV | Kinh Vietnamese | EAS | East Asian Ancestry | KHV | 1000 Genomes on GRCh38,1000 Genomes 30x on GRCh38,1000 Genomes phase 3 release |
| **HG01862** | female | SAME123199 | KHV | Kinh Vietnamese | EAS | East Asian Ancestry | KHV | 1000 Genomes on GRCh38,1000 Genomes 30x on GRCh38,1000 Genomes phase 3 release |
| **HG02102** | female | SAME1840155 | PEL | Peruvian | AMR | American Ancestry | PEL | 1000 Genomes on GRCh38,1000 Genomes 30x on GRCh38,1000 Genomes phase 3 release |
| **HG01672** | male | SAME123288 | IBS | Iberian | EUR | European Ancestry | IBS | 1000 Genomes on GRCh38,1000 Genomes 30x on GRCh38,1000 Genomes phase 3 release |
| **HG02107** | male | SAME1840206 | ACB | African Caribbean | AFR | African Ancestry | ACB | 1000 Genomes on GRCh38,1000 Genomes 30x on GRCh38,1000 Genomes phase 3 release |
| **HG01677** | female | SAME123285 | IBS | Iberian | EUR | European Ancestry | IBS | 1000 Genomes 30x on GRCh38,1000 Genomes phase 3 release |
| **HG02145** | male | SAME1839616 | ACB | African Caribbean | AFR | African Ancestry | ACB | 1000 Genomes 30x on GRCh38,1000 Genomes phase 3 release |
| **HG01800** | female | SAME124306 | CDX | Dai Chinese | EAS | East Asian Ancestry | CDX | 1000 Genomes on GRCh38,1000 Genomes 30x on GRCh38,1000 Genomes phase 3 release |
| **HG02152** | female | SAME124790 | CDX | Dai Chinese | EAS | East Asian Ancestry | CDX | 1000 Genomes on GRCh38,1000 Genomes 30x on GRCh38,1000 Genomes phase 3 release |
| **HG01805** | female | SAME124301 | CDX | Dai Chinese | EAS | East Asian Ancestry | CDX | 1000 Genomes on GRCh38,1000 Genomes 30x on GRCh38,1000 Genomes phase 3 release |
| **HG02164** | female | SAME124209 | CDX | Dai Chinese | EAS | East Asian Ancestry | CDX | 1000 Genomes on GRCh38,1000 Genomes 30x on GRCh38,1000 Genomes phase 3 release |
| **HG01812** | female | SAME124517 | CDX | Dai Chinese | EAS | East Asian Ancestry | CDX | 1000 Genomes on GRCh38,1000 Genomes 30x on GRCh38,1000 Genomes phase 3 release |
| **HG01817** | female | SAME124522 | CDX | Dai Chinese | EAS | East Asian Ancestry | CDX | 1000 Genomes on GRCh38,1000 Genomes 30x on GRCh38,1000 Genomes phase 3 release |
| **HG02121** | female | SAME123479 | KHV | Kinh Vietnamese | EAS | East Asian Ancestry | KHV | 1000 Genomes on GRCh38,1000 Genomes 30x on GRCh38,1000 Genomes phase 3 release |
| **HG02079** | male | SAME123098 | KHV | Kinh Vietnamese | EAS | East Asian Ancestry | KHV | 1000 Genomes on GRCh38,1000 Genomes 30x on GRCh38,1000 Genomes phase 3 release |
| **HG02126** | female | SAME123477 | KHV | Kinh Vietnamese | EAS | East Asian Ancestry | KHV | 1000 Genomes 30x on GRCh38,1000 Genomes phase 3 release |
| **HG02081** | female | SAME124826 | KHV | Kinh Vietnamese | EAS | East Asian Ancestry | KHV | 1000 Genomes on GRCh38,1000 Genomes 30x on GRCh38,1000 Genomes phase 3 release |
| **HG02133** | female | SAME1839882 | KHV | Kinh Vietnamese | EAS | East Asian Ancestry | KHV | 1000 Genomes on GRCh38,1000 Genomes 30x on GRCh38,1000 Genomes phase 3 release |
| **HG02086** | female | SAME124675 | KHV | Kinh Vietnamese | EAS | East Asian Ancestry | KHV | 1000 Genomes on GRCh38,1000 Genomes 30x on GRCh38,1000 Genomes phase 3 release |
| **HG02138** | male | SAME123659 | KHV | Kinh Vietnamese | EAS | East Asian Ancestry | KHV | 1000 Genomes on GRCh38,1000 Genomes 30x on GRCh38,1000 Genomes phase 3 release |
| **HG02106** | female | SAME123875 | PEL | Peruvian | AMR | American Ancestry | PEL | 1000 Genomes 30x on GRCh38,Human Genome Structural Variation Consortium, Phase 2,1000 Genomes phase 3 release |
| **HG02140** | female | SAME124612 | KHV | Kinh Vietnamese | EAS | East Asian Ancestry | KHV | 1000 Genomes on GRCh38,1000 Genomes 30x on GRCh38,1000 Genomes phase 3 release |
| **HG01949** | female | SAME124145 | PEL | Peruvian | AMR | American Ancestry | PEL | 1000 Genomes 30x on GRCh38,1000 Genomes phase 3 release |
| **HG02222** | female | SAME123717 | IBS | Iberian | EUR | European Ancestry | IBS | 1000 Genomes 30x on GRCh38,1000 Genomes phase 3 release |
| **HG01951** | female | SAME123976 | PEL | Peruvian | AMR | American Ancestry | PEL | 1000 Genomes on GRCh38,1000 Genomes 30x on GRCh38,1000 Genomes phase 3 release |
| **HG02234** | male | SAME1839367 | IBS | Iberian | EUR | European Ancestry | IBS | 1000 Genomes 30x on GRCh38,1000 Genomes phase 3 release |
| **HG01956** | female | SAME123978 | ACB | African Caribbean | AFR | African Ancestry | ACB | 1000 Genomes on GRCh38,1000 Genomes 30x on GRCh38,1000 Genomes phase 3 release |
| **HG02239** | female | SAME1839420 | IBS | Iberian | EUR | European Ancestry | IBS | 1000 Genomes on GRCh38,1000 Genomes 30x on GRCh38,1000 Genomes phase 3 release |
| **HG01968** | female | SAME124532 | PEL | Peruvian | AMR | American Ancestry | PEL | 1000 Genomes on GRCh38,1000 Genomes 30x on GRCh38,1000 Genomes phase 3 release |
| **HG02253** | male | SAME1839347 | PEL | Peruvian | AMR | American Ancestry | PEL | 1000 Genomes on GRCh38,1000 Genomes 30x on GRCh38,1000 Genomes phase 3 release |
| **HG01970** | male | SAME124327 | PEL | Peruvian | AMR | American Ancestry | PEL | 1000 Genomes on GRCh38,1000 Genomes 30x on GRCh38,1000 Genomes phase 3 release |
| **HG02056** | male | SAME124162 | KHV | Kinh Vietnamese | EAS | East Asian Ancestry | KHV | 1000 Genomes 30x on GRCh38,1000 Genomes phase 3 release |
| **HG02024** | female | SAME125182 | KHV | Kinh Vietnamese | EAS | East Asian Ancestry | KHV | 1000 Genomes on GRCh38,1000 Genomes 30x on GRCh38,1000 Genomes phase 3 release |
| **HG02068** | female | SAME123374 | KHV | Kinh Vietnamese | EAS | East Asian Ancestry | KHV | 1000 Genomes 30x on GRCh38,1000 Genomes phase 3 release |
| **HG02029** | male | SAME124021 | KHV | Kinh Vietnamese | EAS | East Asian Ancestry | KHV | 1000 Genomes on GRCh38,1000 Genomes 30x on GRCh38,1000 Genomes phase 3 release |
| **HG02070** | male | SAME124908 | KHV | Kinh Vietnamese | EAS | East Asian Ancestry | KHV | 1000 Genomes on GRCh38,1000 Genomes 30x on GRCh38,1000 Genomes phase 3 release |
| **HG02031** | female | SAME1839964 | KHV | Kinh Vietnamese | EAS | East Asian Ancestry | KHV | 1000 Genomes on GRCh38,1000 Genomes 30x on GRCh38,1000 Genomes phase 3 release |
| **HG02075** | female | SAME123106 | KHV | Kinh Vietnamese | EAS | East Asian Ancestry | KHV | 1000 Genomes on GRCh38,1000 Genomes 30x on GRCh38,1000 Genomes phase 3 release |
| **HG02048** | female | SAME123772 | KHV | Kinh Vietnamese | EAS | East Asian Ancestry | KHV | 1000 Genomes on GRCh38,1000 Genomes 30x on GRCh38,1000 Genomes phase 3 release |
| **HG02050** | male | SAME1839555 | KHV | Kinh Vietnamese | EAS | East Asian Ancestry | KHV | 1000 Genomes on GRCh38,1000 Genomes 30x on GRCh38,1000 Genomes phase 3 release |
| **HG02188** | female | SAME122829 | CDX | Dai Chinese | EAS | East Asian Ancestry | CDX | 1000 Genomes on GRCh38,1000 Genomes 30x on GRCh38,1000 Genomes phase 3 release |
| **HG02055** | male | SAME1839531 | ACB | African Caribbean | AFR | African Ancestry | ACB | 1000 Genomes 30x on GRCh38,1000 Genomes phase 3 release |
| **HG02190** | female | SAME125259 | CDX | Dai Chinese | EAS | East Asian Ancestry | CDX | 1000 Genomes on GRCh38,1000 Genomes 30x on GRCh38,1000 Genomes phase 3 release |
| **HG02067** | male | SAME1839388 | KHV | Kinh Vietnamese | EAS | East Asian Ancestry | KHV | 1000 Genomes on GRCh38,1000 Genomes 30x on GRCh38,1000 Genomes phase 3 release |
| **HG02215** | female | SAME123904 | GBR | British | EUR | European Ancestry | GBR | 1000 Genomes on GRCh38,1000 Genomes 30x on GRCh38,1000 Genomes phase 3 release,Geuvadis |
| **HG02074** | male | SAME123107 | KHV | Kinh Vietnamese | EAS | East Asian Ancestry | KHV | 1000 Genomes 30x on GRCh38,1000 Genomes phase 3 release |
| **HG02563** | male | SAME1839824 | GWD | Gambian Mandinka | AFR | African Ancestry | GWD | 1000 Genomes 30x on GRCh38 |
| **HG01925** | female | SAME123786 | PEL | Peruvian | AMR | American Ancestry | PEL | 1000 Genomes 30x on GRCh38,1000 Genomes phase 3 release |
| **HG02568** | female | SAME1839899 | GWD | Gambian Mandinka | AFR | African Ancestry | GWD | 1000 Genomes on GRCh38,1000 Genomes 30x on GRCh38,1000 Genomes phase 3 release,Gambian Genome Variation Project (GRCh38) |
| **HG01932** | male | SAME123587 | PEL | Peruvian | AMR | American Ancestry | PEL | 1000 Genomes on GRCh38,1000 Genomes 30x on GRCh38,1000 Genomes phase 3 release |
| **HG02570** | male | SAME1839582 | GWD | Gambian Mandinka | AFR | African Ancestry | GWD | 1000 Genomes on GRCh38,1000 Genomes 30x on GRCh38,1000 Genomes phase 3 release,Gambian Genome Variation Project (GRCh38) |
| **HG01937** | male | SAME123590 | PEL | Peruvian | AMR | American Ancestry | PEL | 1000 Genomes 30x on GRCh38,1000 Genomes phase 3 release |
| **HG02575** | male | SAME1839650 | GWD | Gambian Mandinka | AFR | African Ancestry | GWD | 1000 Genomes 30x on GRCh38 |
| **HG01944** | male | SAME124152 | PEL | Peruvian | AMR | American Ancestry | PEL | 1000 Genomes on GRCh38,1000 Genomes 30x on GRCh38,1000 Genomes phase 3 release |
| **HG02582** | male | SAME1839431 | GWD | Gambian Mandinka | AFR | African Ancestry | GWD | 1000 Genomes on GRCh38,1000 Genomes 30x on GRCh38,1000 Genomes phase 3 release,Gambian Genome Variation Project (GRCh38) |
| **HG01975** | female | SAME124322 | PEL | Peruvian | AMR | American Ancestry | PEL | 1000 Genomes 30x on GRCh38,1000 Genomes phase 3 release |
| **HG02587** | female | SAME1839409 | GWD | Gambian Mandinka | AFR | African Ancestry | GWD | 1000 Genomes 30x on GRCh38,Human Genome Structural Variation Consortium, Phase 2 |
| **HG01982** | male | SAME1840111 | PEL | Peruvian | AMR | American Ancestry | PEL | 1000 Genomes on GRCh38,1000 Genomes 30x on GRCh38,1000 Genomes phase 3 release |
| **HG02594** | male | SAME1839208 | GWD | Gambian Mandinka | AFR | African Ancestry | GWD | 1000 Genomes on GRCh38,1000 Genomes 30x on GRCh38,1000 Genomes phase 3 release,Gambian Genome Variation Project (GRCh38) |
| **HG01987** | male | SAME1840089 | ACB | African Caribbean | AFR | African Ancestry | ACB | 1000 Genomes 30x on GRCh38,1000 Genomes phase 3 release |
| **HG02113** | female | SAME124044 | KHV | Kinh Vietnamese | EAS | East Asian Ancestry | KHV | 1000 Genomes on GRCh38,1000 Genomes 30x on GRCh38,1000 Genomes phase 3 release |
| **HG02602** | male | SAME1840290 | PJL | Punjabi | SAS | South Asian Ancestry | PJL | 1000 Genomes 30x on GRCh38,1000 Genomes phase 3 release |
| **HG02120** | male | SAME123480 | KHV | Kinh Vietnamese | EAS | East Asian Ancestry | KHV | 1000 Genomes 30x on GRCh38,1000 Genomes phase 3 release |
| **HG02429** | male | SAME1839021 | ACB | African Caribbean | AFR | African Ancestry | ACB | 1000 Genomes on GRCh38,1000 Genomes 30x on GRCh38,1000 Genomes phase 3 release |
| **HG02132** | male | SAME1839889 | KHV | Kinh Vietnamese | EAS | East Asian Ancestry | KHV | 1000 Genomes 30x on GRCh38,1000 Genomes phase 3 release |
| **HG02137** | male | SAME1839862 | KHV | Kinh Vietnamese | EAS | East Asian Ancestry | KHV | 1000 Genomes on GRCh38,1000 Genomes 30x on GRCh38,1000 Genomes phase 3 release |
| **HG02450** | female | SAME1840144 | ACB | African Caribbean | AFR | African Ancestry | ACB | 1000 Genomes on GRCh38,1000 Genomes 30x on GRCh38,1000 Genomes phase 3 release |
| **HG01893** | female | SAME123027 | PEL | Peruvian | AMR | American Ancestry | PEL | 1000 Genomes on GRCh38,1000 Genomes 30x on GRCh38,1000 Genomes phase 3 release |
| **HG02455** | male | SAME1840198 | ACB | African Caribbean | AFR | African Ancestry | ACB | 1000 Genomes on GRCh38,1000 Genomes 30x on GRCh38,1000 Genomes phase 3 release |
| **HG01898** | female | SAME124383 | PEL | Peruvian | AMR | American Ancestry | PEL | 1000 Genomes 30x on GRCh38,1000 Genomes phase 3 release |
| **HG02462** | female | SAME1839991 | GWD | Gambian Mandinka | AFR | African Ancestry | GWD | 1000 Genomes on GRCh38,1000 Genomes 30x on GRCh38,1000 Genomes phase 3 release,Gambian Genome Variation Project (GRCh38) |
| **HG01918** | female | SAME123240 | PEL | Peruvian | AMR | American Ancestry | PEL | 1000 Genomes on GRCh38,1000 Genomes 30x on GRCh38,1000 Genomes phase 3 release |
| **HG02258** | male | SAME1839322 | ACB | African Caribbean | AFR | African Ancestry | ACB | 1000 Genomes 30x on GRCh38,1000 Genomes phase 3 release |
| **HG01920** | male | SAME123789 | PEL | Peruvian | AMR | American Ancestry | PEL | 1000 Genomes on GRCh38,1000 Genomes 30x on GRCh38,1000 Genomes phase 3 release |
| **HG02260** | female | SAME1839139 | PEL | Peruvian | AMR | American Ancestry | PEL | 1000 Genomes on GRCh38,1000 Genomes 30x on GRCh38,1000 Genomes phase 3 release |
| **HG02012** | female | SAME1840403 | ACB | African Caribbean | AFR | African Ancestry | ACB | 1000 Genomes on GRCh38,1000 Genomes 30x on GRCh38,1000 Genomes phase 3 release |
| **HG02265** | male | SAME1839117 | PEL | Peruvian | AMR | American Ancestry | PEL | 1000 Genomes on GRCh38,1000 Genomes 30x on GRCh38,1000 Genomes phase 3 release |
| **HG02017** | male | SAME123445 | KHV | Kinh Vietnamese | EAS | East Asian Ancestry | KHV | 1000 Genomes on GRCh38,1000 Genomes 30x on GRCh38,1000 Genomes phase 3 release |
| **HG02272** | female | SAME1840314 | PEL | Peruvian | AMR | American Ancestry | PEL | 1000 Genomes on GRCh38,1000 Genomes 30x on GRCh38,1000 Genomes phase 3 release |
| **HG02277** | male | SAME1840370 | PEL | Peruvian | AMR | American Ancestry | PEL | 1000 Genomes on GRCh38,1000 Genomes 30x on GRCh38,1000 Genomes phase 3 release |
| **HG02442** | male | SAME1840352 | ACB | African Caribbean | AFR | African Ancestry | ACB | 1000 Genomes on GRCh38,1000 Genomes 30x on GRCh38,1000 Genomes phase 3 release |
| **HG02479** | female | SAME1839798 | ACB | African Caribbean | AFR | African Ancestry | ACB | 1000 Genomes on GRCh38,1000 Genomes 30x on GRCh38,1000 Genomes phase 3 release |
| **HG02461** | male | SAME1839993 | GWD | Gambian Mandinka | AFR | African Ancestry | GWD | 1000 Genomes on GRCh38,1000 Genomes 30x on GRCh38,1000 Genomes phase 3 release,Gambian Genome Variation Project (GRCh38) |
| **HG02481** | male | SAME1839870 | ACB | African Caribbean | AFR | African Ancestry | ACB | 1000 Genomes on GRCh38,1000 Genomes 30x on GRCh38,1000 Genomes phase 3 release |
| **HG02466** | male | SAME1839970 | GWD | Gambian Mandinka | AFR | African Ancestry | GWD | 1000 Genomes 30x on GRCh38 |
| **HG02493** | male | SAME1839704 | PJL | Punjabi | SAS | South Asian Ancestry | PJL | 1000 Genomes on GRCh38,1000 Genomes 30x on GRCh38,1000 Genomes phase 3 release |

| | | | | | | | | |
|---|---|---|---|---|---|---|---|---|
| **HG02182** | female | SAME125035 | CDX | Dai Chinese | EAS | East Asian Ancestry | CDX | 1000 Genomes on GRCh38,1000 Genomes 30x on GRCh38,1000 Genomes phase 3 release |
| **HG02501** | male | SAME1839281 | ACB | African Caribbean | AFR | African Ancestry | ACB | 1000 Genomes on GRCh38,1000 Genomes 30x on GRCh38,1000 Genomes phase 3 release |
| **HG02187** | female | SAME125038 | CDX | Dai Chinese | EAS | East Asian Ancestry | CDX | 1000 Genomes on GRCh38,1000 Genomes 30x on GRCh38,1000 Genomes phase 3 release |
| **HG02513** | female | SAME1839138 | KHV | Kinh Vietnamese | EAS | East Asian Ancestry | KHV | 1000 Genomes on GRCh38,1000 Genomes 30x on GRCh38,1000 Genomes phase 3 release |
| **HG02219** | male | SAME1839813 | IBS | Iberian | EUR | European Ancestry | IBS | 1000 Genomes on GRCh38,1000 Genomes 30x on GRCh38,1000 Genomes phase 3 release |
| **HG02525** | female | SAME1840313 | KHV | Kinh Vietnamese | EAS | East Asian Ancestry | KHV | 1000 Genomes on GRCh38,1000 Genomes 30x on GRCh38 |
| **HG02221** | male | SAME123716 | IBS | Iberian | EUR | European Ancestry | IBS | 1000 Genomes on GRCh38,1000 Genomes 30x on GRCh38,1000 Genomes phase 3 release |
| **HG02537** | female | SAME1840165 | ACB | African Caribbean | AFR | African Ancestry | ACB | 1000 Genomes on GRCh38,1000 Genomes 30x on GRCh38,1000 Genomes phase 3 release |
| **HG02283** | male | SAME1840173 | ACB | African Caribbean | AFR | African Ancestry | ACB | 1000 Genomes on GRCh38,1000 Genomes 30x on GRCh38,1000 Genomes phase 3 release |
| **HG02549** | female | SAME1839917 | ACB | African Caribbean | AFR | African Ancestry | ACB | 1000 Genomes on GRCh38,1000 Genomes 30x on GRCh38,1000 Genomes phase 3 release |
| **HG02284** | male | SAME1840168 | ACB | African Caribbean | AFR | African Ancestry | ACB | 1000 Genomes on GRCh38,1000 Genomes 30x on GRCh38,1000 Genomes phase 3 release |
| **HG02303** | female | SAME124278 | PEL | Peruvian | AMR | American Ancestry | PEL | 1000 Genomes 30x on GRCh38,1000 Genomes phase 3 release |
| **HG02291** | male | SAME1839948 | PEL | Peruvian | AMR | American Ancestry | PEL | 1000 Genomes on GRCh38,1000 Genomes 30x on GRCh38,1000 Genomes phase 3 release |
| **HG02308** | female | SAME1839529 | ACB | African Caribbean | AFR | African Ancestry | ACB | 1000 Genomes on GRCh38,1000 Genomes 30x on GRCh38,1000 Genomes phase 3 release |
| **HG02304** | male | SAME1839548 | PEL | Peruvian | AMR | American Ancestry | PEL | 1000 Genomes on GRCh38,1000 Genomes 30x on GRCh38,1000 Genomes phase 3 release |
| **HG02353** | male | SAME1840276 | CDX | Dai Chinese | EAS | East Asian Ancestry | CDX | 1000 Genomes on GRCh38,1000 Genomes 30x on GRCh38,1000 Genomes phase 3 release |
| **HG02309** | female | SAME1839525 | ACB | African Caribbean | AFR | African Ancestry | ACB | 1000 Genomes on GRCh38,1000 Genomes 30x on GRCh38,1000 Genomes phase 3 release |
| **HG02316** | male | SAME1839661 | ACB | African Caribbean | AFR | African Ancestry | ACB | 1000 Genomes 30x on GRCh38,1000 Genomes phase 3 release |
| **HG02360** | male | SAME1840391 | CDX | Dai Chinese | EAS | East Asian Ancestry | CDX | 1000 Genomes on GRCh38,1000 Genomes 30x on GRCh38,1000 Genomes phase 3 release |
| **HG02323** | male | SAME1839440 | ACB | African Caribbean | AFR | African Ancestry | ACB | 1000 Genomes on GRCh38,1000 Genomes 30x on GRCh38,1000 Genomes phase 3 release |
| **HG02330** | male | SAME1839237 | ACB | African Caribbean | AFR | African Ancestry | ACB | 1000 Genomes on GRCh38,1000 Genomes 30x on GRCh38,1000 Genomes phase 3 release |
| **HG02384** | male | SAME1840011 | CDX | Dai Chinese | EAS | East Asian Ancestry | CDX | 1000 Genomes on GRCh38,1000 Genomes 30x on GRCh38,1000 Genomes phase 3 release |
| **HG02373** | male | SAME124104 | CDX | Dai Chinese | EAS | East Asian Ancestry | CDX | 1000 Genomes on GRCh38,1000 Genomes 30x on GRCh38,1000 Genomes phase 3 release |
| **HG02144** | female | SAME1839624 | ACB | African Caribbean | AFR | African Ancestry | ACB | 1000 Genomes on GRCh38,1000 Genomes 30x on GRCh38,1000 Genomes phase 3 release |
| **HG02380** | male | SAME1840045 | CDX | Dai Chinese | EAS | East Asian Ancestry | CDX | 1000 Genomes on GRCh38,1000 Genomes 30x on GRCh38,1000 Genomes phase 3 release |
| **HG02151** | female | SAME1839403 | CDX | Dai Chinese | EAS | East Asian Ancestry | CDX | 1000 Genomes on GRCh38,1000 Genomes 30x on GRCh38,1000 Genomes phase 3 release |
| **HG02385** | male | SAME1840006 | CDX | Dai Chinese | EAS | East Asian Ancestry | CDX | 1000 Genomes on GRCh38,1000 Genomes 30x on GRCh38,1000 Genomes phase 3 release |
| **HG02156** | female | SAME124793 | CDX | Dai Chinese | EAS | East Asian Ancestry | CDX | 1000 Genomes on GRCh38,1000 Genomes 30x on GRCh38,1000 Genomes phase 3 release |
| **HG02392** | male | SAME1839776 | CDX | Dai Chinese | EAS | East Asian Ancestry | CDX | 1000 Genomes on GRCh38,1000 Genomes 30x on GRCh38,1000 Genomes phase 3 release |
| **HG02397** | male | SAME123308 | CDX | Dai Chinese | EAS | East Asian Ancestry | CDX | 1000 Genomes on GRCh38,1000 Genomes 30x on GRCh38,1000 Genomes phase 3 release |
| **HG02233** | male | SAME1839370 | IBS | Iberian | EUR | European Ancestry | IBS | 1000 Genomes on GRCh38,1000 Genomes 30x on GRCh38,1000 Genomes phase 3 release |
| **HG02238** | male | SAME1839423 | IBS | Iberian | EUR | European Ancestry | IBS | 1000 Genomes on GRCh38,1000 Genomes 30x on GRCh38,1000 Genomes phase 3 release |
| **HG02789** | male | SAME1840280 | PJL | Punjabi | SAS | South Asian Ancestry | PJL | 1000 Genomes on GRCh38,1000 Genomes 30x on GRCh38,1000 Genomes phase 3 release |
| **HG02240** | female | SAME1839481 | IBS | Iberian | EUR | European Ancestry | IBS | 1000 Genomes 30x on GRCh38,1000 Genomes phase 3 release |
| **HG02791** | male | SAME1840342 | PJL | Punjabi | SAS | South Asian Ancestry | PJL | 1000 Genomes 30x on GRCh38,1000 Genomes phase 3 release |
| **HG02252** | female | SAME1839354 | PEL | Peruvian | AMR | American Ancestry | PEL | 1000 Genomes on GRCh38,1000 Genomes 30x on GRCh38,1000 Genomes phase 3 release |
| **HG02804** | male | SAME1840084 | GWD | Gambian Mandinka | AFR | African Ancestry | GWD | 1000 Genomes on GRCh38,1000 Genomes 30x on GRCh38,1000 Genomes phase 3 release,Gambian Genome Variation Project (GRCh38) |
| **HG02257** | female | SAME1839326 | ACB | African Caribbean | AFR | African Ancestry | ACB | 1000 Genomes 30x on GRCh38,1000 Genomes phase 3 release |
| **HG02809** | female | SAME1840067 | GWD | Gambian Mandinka | AFR | African Ancestry | GWD | 1000 Genomes 30x on GRCh38 |
| **HG02271** | male | SAME1840319 | PEL | Peruvian | AMR | American Ancestry | PEL | 1000 Genomes on GRCh38,1000 Genomes 30x on GRCh38,1000 Genomes phase 3 release |
| **HG02811** | female | SAME1839849 | GWD | Gambian Mandinka | AFR | African Ancestry | GWD | 1000 Genomes on GRCh38,1000 Genomes 30x on GRCh38,1000 Genomes phase 3 release,Gambian Genome Variation Project (GRCh38) |
| **HG02816** | male | SAME1839822 | GWD | Gambian Mandinka | AFR | African Ancestry | GWD | 1000 Genomes on GRCh38,1000 Genomes 30x on GRCh38,1000 Genomes phase 3 release,Gambian Genome Variation Project (GRCh38) |
| **HG02315** | female | SAME1839666 | ACB | African Caribbean | AFR | African Ancestry | ACB | 1000 Genomes on GRCh38,1000 Genomes 30x on GRCh38,1000 Genomes phase 3 release |
| **HG02854** | male | SAME1839336 | GWD | Gambian Mandinka | AFR | African Ancestry | GWD | 1000 Genomes on GRCh38,1000 Genomes 30x on GRCh38,1000 Genomes phase 3 release,Gambian Genome Variation Project (GRCh38) |
| **HG02322** | female | SAME1839444 | ACB | African Caribbean | AFR | African Ancestry | ACB | 1000 Genomes on GRCh38,1000 Genomes 30x on GRCh38,1000 Genomes phase 3 release |
| **HG02861** | female | SAME1839125 | GWD | Gambian Mandinka | AFR | African Ancestry | GWD | 1000 Genomes on GRCh38,1000 Genomes 30x on GRCh38,1000 Genomes phase 3 release,Gambian Genome Variation Project (GRCh38) |
| **HG02334** | male | SAME1839307 | ACB | African Caribbean | AFR | African Ancestry | ACB | 1000 Genomes on GRCh38,1000 Genomes 30x on GRCh38,1000 Genomes phase 3 release |
| **HG02878** | male | SAME1840363 | GWD | Gambian Mandinka | AFR | African Ancestry | GWD | 1000 Genomes on GRCh38,1000 Genomes 30x on GRCh38,1000 Genomes phase 3 release,Gambian Genome Variation Project (GRCh38) |
| **HG02339** | female | SAME1839288 | ACB | African Caribbean | AFR | African Ancestry | ACB | 1000 Genomes on GRCh38,1000 Genomes 30x on GRCh38,1000 Genomes phase 3 release |
| **HG02880** | female | SAME1840090 | GWD | Gambian Mandinka | AFR | African Ancestry | GWD | 1000 Genomes 30x on GRCh38 |
| **HG02389** | male | SAME1840085 | CDX | Dai Chinese | EAS | East Asian Ancestry | CDX | 1000 Genomes on GRCh38,1000 Genomes 30x on GRCh38,1000 Genomes phase 3 release |
| **HG02885** | female | SAME1840157 | GWD | Gambian Mandinka | AFR | African Ancestry | GWD | 1000 Genomes on GRCh38,1000 Genomes 30x on GRCh38,1000 Genomes phase 3 release,Gambian Genome Variation Project (GRCh38) |
| **HG02391** | male | SAME1839784 | CDX | Dai Chinese | EAS | East Asian Ancestry | CDX | 1000 Genomes on GRCh38,1000 Genomes 30x on GRCh38,1000 Genomes phase 3 release |
| **HG02892** | male | SAME1839935 | GWD | Gambian Mandinka | AFR | African Ancestry | GWD | 1000 Genomes 30x on GRCh38 |
| **HG02396** | male | SAME125357 | CDX | Dai Chinese | EAS | East Asian Ancestry | CDX | 1000 Genomes on GRCh38,1000 Genomes 30x on GRCh38,1000 Genomes phase 3 release |
| **HG02897** | female | SAME1839910 | GWD | Gambian Mandinka | AFR | African Ancestry | GWD | 1000 Genomes 30x on GRCh38 |
| **HG02409** | male | SAME1839457 | CDX | Dai Chinese | EAS | East Asian Ancestry | CDX | 1000 Genomes on GRCh38,1000 Genomes 30x on GRCh38,1000 Genomes phase 3 release |
| **HG02924** | male | SAME1839516 | ESN | Esan | AFR | African Ancestry | ESN | 1000 Genomes 30x on GRCh38 |
| **HG02733** | male | SAME1839874 | PJL | Punjabi | SAS | South Asian Ancestry | PJL | 1000 Genomes on GRCh38,1000 Genomes 30x on GRCh38,1000 Genomes phase 3 release |
| **HG02974** | female | SAME1840241 | ESN | Esan | AFR | African Ancestry | ESN | 1000 Genomes on GRCh38,1000 Genomes 30x on GRCh38,1000 Genomes phase 3 release |
| **HG02738** | male | SAME1839560 | PJL | Punjabi | SAS | South Asian Ancestry | PJL | 1000 Genomes 30x on GRCh38 |
| **HG02979** | female | SAME1840209 | ESN | Esan | AFR | African Ancestry | ESN | 1000 Genomes on GRCh38,1000 Genomes 30x on GRCh38,1000 Genomes phase 3 release |
| **HG02757** | female | SAME1839451 | GWD | Gambian Mandinka | AFR | African Ancestry | GWD | 1000 Genomes on GRCh38,1000 Genomes 30x on GRCh38,1000 Genomes phase 3 release,Gambian Genome Variation Project (GRCh38) |
| **HG02981** | male | SAME1840024 | ESN | Esan | AFR | African Ancestry | ESN | 1000 Genomes on GRCh38,1000 Genomes 30x on GRCh38,1000 Genomes phase 3 release |
| **HG02764** | male | SAME1839250 | GWD | Gambian Mandinka | AFR | African Ancestry | GWD | 1000 Genomes 30x on GRCh38 |
| **HG03009** | male | SAME1840256 | BEB | Bengali | SAS | South Asian Ancestry | BEB | 1000 Genomes on GRCh38,1000 Genomes 30x on GRCh38,Human Genome Structural Variation Consortium, Phase 2,1000 Genomes phase 3 release |
| **HG02769** | female | SAME1839231 | GWD | Gambian Mandinka | AFR | African Ancestry | GWD | 1000 Genomes on GRCh38,1000 Genomes 30x on GRCh38,1000 Genomes phase 3 release,Gambian Genome Variation Project (GRCh38) |
| **HG03016** | female | SAME1839952 | PJL | Punjabi | SAS | South Asian Ancestry | PJL | 1000 Genomes on GRCh38,1000 Genomes 30x on GRCh38,1000 Genomes phase 3 release |
| **HG02771** | male | SAME1839045 | GWD | Gambian Mandinka | AFR | African Ancestry | GWD | 1000 Genomes on GRCh38,1000 Genomes 30x on GRCh38,1000 Genomes phase 3 release,Gambian Genome Variation Project (GRCh38) |
| **HG02776** | female | SAME1839017 | PJL | Punjabi | SAS | South Asian Ancestry | PJL | 1000 Genomes 30x on GRCh38 |
| **HG02734** | female | SAME1839869 | PJL | Punjabi | SAS | South Asian Ancestry | PJL | 1000 Genomes on GRCh38,1000 Genomes 30x on GRCh38,1000 Genomes phase 3 release |
| **HG02783** | male | SAME1839162 | PJL | Punjabi,Punjabi | SAS | South Asia (SGDP),South Asian Ancestry | PunjabiSGDP,PJL | 1000 Genomes on GRCh38,Simons Genome Diversity Project,1000 Genomes 30x on GRCh38,1000 Genomes phase 3 release |
| **HG02562** | female | SAME1839832 | GWD | Gambian Mandinka | AFR | African Ancestry | GWD | 1000 Genomes on GRCh38,1000 Genomes 30x on GRCh38,1000 Genomes phase 3 release,Gambian Genome Variation Project (GRCh38) |
| **HG02760** | female | SAME1839267 | GWD | Gambian Mandinka | AFR | African Ancestry | GWD | 1000 Genomes on GRCh38,1000 Genomes 30x on GRCh38,1000 Genomes phase 3 release,Gambian Genome Variation Project (GRCh38) |
| **HG02772** | female | SAME1839039 | GWD | Gambian Mandinka | AFR | African Ancestry | GWD | 1000 Genomes on GRCh38,1000 Genomes 30x on GRCh38,1000 Genomes phase 3 release,Gambian Genome Variation Project (GRCh38) |
| **HG02574** | female | SAME1839652 | GWD | Gambian Mandinka,Gar | AFR | African Ancestry,Africa (SGDP) | GWD,GambianSGDP | 1000 Genomes on GRCh38,Simons Genome Diversity Project,1000 Genomes 30x on GRCh38,1000 Genomes phase 3 release,Gambian Genome Variation Project (GRCh |
| **HG02784** | female | SAME1839157 | PJL | Punjabi | SAS | South Asian Ancestry | PJL | 1000 Genomes on GRCh38,1000 Genomes 30x on GRCh38,1000 Genomes phase 3 release |
| **HG02586** | female | SAME1839414 | GWD | Gambian Mandinka | AFR | African Ancestry | GWD | 1000 Genomes on GRCh38,1000 Genomes 30x on GRCh38,1000 Genomes phase 3 release,Gambian Genome Variation Project (GRCh38) |
| **HG02690** | male | SAME1839088 | PJL | Punjabi | SAS | South Asian Ancestry | PJL | 1000 Genomes on GRCh38,1000 Genomes 30x on GRCh38,1000 Genomes phase 3 release |
| **HG02601** | female | SAME1840293 | PJL | Punjabi | SAS | South Asian Ancestry | PJL | 1000 Genomes on GRCh38,1000 Genomes 30x on GRCh38,1000 Genomes phase 3 release |
| **HG02703** | female | SAME1840142 | GWD | Gambian Mandinka | AFR | African Ancestry | GWD | 1000 Genomes on GRCh38,1000 Genomes 30x on GRCh38,1000 Genomes phase 3 release,Gambian Genome Variation Project (GRCh38) |
| **HG02788** | female | SAME1840285 | PJL | Punjabi | SAS | South Asian Ancestry | PJL | 1000 Genomes 30x on GRCh38 |
| **HG02715** | male | SAME1839987 | GWD | Gambian Mandinka | AFR | African Ancestry | GWD | 1000 Genomes on GRCh38,1000 Genomes 30x on GRCh38,1000 Genomes phase 3 release,Gambian Genome Variation Project (GRCh38) |
| **HG02790** | female | SAME1840345 | PJL | Punjabi,Punjabi | SAS | South Asia (SGDP),South Asian Ancestry | PunjabiSGDP,PJL | 1000 Genomes on GRCh38,Simons Genome Diversity Project,1000 Genomes 30x on GRCh38,1000 Genomes phase 3 release |
| **HG02722** | female | SAME1839747 | GWD | Gambian Mandinka | AFR | African Ancestry | GWD | 1000 Genomes on GRCh38,1000 Genomes 30x on GRCh38,1000 Genomes phase 3 release,Gambian Genome Variation Project (GRCh38) |
| **HG02808** | female | SAME1840069 | GWD | Gambian Mandinka | AFR | African Ancestry | GWD | 1000 Genomes on GRCh38,1000 Genomes 30x on GRCh38,1000 Genomes phase 3 release,Gambian Genome Variation Project (GRCh38) |
| **HG02727** | male | SAME1839727 | PJL | Punjabi | SAS | South Asian Ancestry | PJL | 1000 Genomes on GRCh38,1000 Genomes 30x on GRCh38,1000 Genomes phase 3 release |
| **HG02810** | male | SAME1839854 | GWD | Gambian Mandinka | AFR | African Ancestry | GWD | 1000 Genomes on GRCh38,1000 Genomes 30x on GRCh38,1000 Genomes phase 3 release,Gambian Genome Variation Project (GRCh38) |
| **HG03028** | female | SAME1839729 | GWD | Gambian Mandinka | AFR | African Ancestry | GWD | 1000 Genomes on GRCh38,1000 Genomes 30x on GRCh38,1000 Genomes phase 3 release,Gambian Genome Variation Project (GRCh38) |
| **HG02815** | male | SAME1839829 | GWD | Gambian Mandinka | AFR | African Ancestry | GWD | 1000 Genomes 30x on GRCh38 |
| **HG03035** | male | SAME1839573 | GWD | Gambian Mandinka | AFR | African Ancestry | GWD | 1000 Genomes 30x on GRCh38 |
| **HG02839** | male | SAME1839413 | GWD | Gambian Mandinka | AFR | African Ancestry | GWD | 1000 Genomes on GRCh38,1000 Genomes 30x on GRCh38,1000 Genomes phase 3 release,Gambian Genome Variation Project (GRCh38) |
| **HG03047** | female | SAME1839644 | GWD | Gambian Mandinka | AFR | African Ancestry | GWD | 1000 Genomes 30x on GRCh38 |
| **HG02841** | female | SAME1839470 | GWD | Gambian Mandinka | AFR | African Ancestry | GWD | 1000 Genomes 30x on GRCh38 |
| **HG03054** | male | SAME1839507 | MSL | Mende | AFR | African Ancestry | MSL | 1000 Genomes on GRCh38,1000 Genomes 30x on GRCh38,1000 Genomes phase 3 release |
| **HG02853** | male | SAME1839342 | GWD | Gambian Mandinka | AFR | African Ancestry | GWD | 1000 Genomes 30x on GRCh38 |
| **HG02860** | male | SAME1839132 | GWD | Gambian Mandinka | AFR | African Ancestry | GWD | 1000 Genomes on GRCh38,1000 Genomes 30x on GRCh38,1000 Genomes phase 3 release,Gambian Genome Variation Project (GRCh38) |
| **HG03061** | female | SAME1839215 | MSL | Mende | AFR | African Ancestry | MSL | 1000 Genomes on GRCh38,1000 Genomes 30x on GRCh38,1000 Genomes phase 3 release |
| **HG02884** | male | SAME1840161 | GWD | Gambian Mandinka | AFR | African Ancestry | GWD | 1000 Genomes on GRCh38,1000 Genomes 30x on GRCh38,1000 Genomes phase 3 release,Gambian Genome Variation Project (GRCh38) |
| **HG02657** | male | SAME1839600 | PJL | Punjabi | SAS | South Asian Ancestry | PJL | 1000 Genomes on GRCh38,1000 Genomes 30x on GRCh38,1000 Genomes phase 3 release |
| **HG02889** | female | SAME1840133 | GWD | Gambian Mandinka | AFR | African Ancestry | GWD | 1000 Genomes 30x on GRCh38 |
| **HG02676** | female | SAME1839491 | GWD | Gambian Mandinka | AFR | African Ancestry | GWD | 1000 Genomes on GRCh38,1000 Genomes 30x on GRCh38,1000 Genomes phase 3 release,Gambian Genome Variation Project (GRCh38) |
| **HG02891** | female | SAME1839938 | GWD | Gambian Mandinka | AFR | African Ancestry | GWD | 1000 Genomes on GRCh38,1000 Genomes 30x on GRCh38,1000 Genomes phase 3 release,Gambian Genome Variation Project (GRCh38) |
| **HG02683** | female | SAME1839298 | PJL | Punjabi | SAS | South Asian Ancestry | PJL | 1000 Genomes 30x on GRCh38 |
| **HG02896** | female | SAME1839916 | GWD | Gambian Mandinka | AFR | African Ancestry | GWD | 1000 Genomes on GRCh38,1000 Genomes 30x on GRCh38,1000 Genomes phase 3 release,Gambian Genome Variation Project (GRCh38) |
| **HG02688** | female | SAME1839358 | PJL | Punjabi | SAS | South Asian Ancestry | PJL | 1000 Genomes on GRCh38,1000 Genomes 30x on GRCh38,1000 Genomes phase 3 release |
| **HG02923** | male | SAME1839436 | ESN | Esan | AFR | African Ancestry | ESN | 1000 Genomes on GRCh38,1000 Genomes 30x on GRCh38,1000 Genomes phase 3 release |
| **HG02614** | female | SAME1840385 | GWD | Gambian Mandinka | AFR | African Ancestry | GWD | 1000 Genomes on GRCh38,1000 Genomes 30x on GRCh38,1000 Genomes phase 3 release,Gambian Genome Variation Project (GRCh38) |
| **HG02621** | female | SAME1840187 | GWD | Gambian Mandinka | AFR | African Ancestry | GWD | 1000 Genomes on GRCh38,1000 Genomes 30x on GRCh38,1000 Genomes phase 3 release,Gambian Genome Variation Project (GRCh38) |
| **HG02645** | male | SAME1839774 | GWD | Gambian Mandinka | AFR | African Ancestry | GWD | 1000 Genomes on GRCh38,1000 Genomes 30x on GRCh38,1000 Genomes phase 3 release,Gambian Genome Variation Project (GRCh38) |
| **HG02485** | female | SAME1839564 | ACB | African Caribbean | AFR | African Ancestry | ACB | 1000 Genomes on GRCh38,1000 Genomes 30x on GRCh38,1000 Genomes phase 3 release |

| | | | | | | | | |
|---|---|---|---|---|---|---|---|---|
| **HG02652** | female | SAME1839541 | PJL | Punjabi | SAS | South Asian Ancestry | PJL | 1000 Genomes on GRCh38,1000 Genomes 30x on GRCh38,1000 Genomes phase 3 release |
| **HG02492** | male | SAME1839610 | PJL | Punjabi | SAS | South Asian Ancestry | PJL | 1000 Genomes 30x on GRCh38,Human Genome Structural Variation Consortium, Phase 2,1000 Genomes phase 3 release |
| **HG02943** | female | SAME1839085 | ESN | Esan,Esan | AFR | African Ancestry,Africa (SGDP) | ESN,EsanSGDP | 1000 Genomes on GRCh38,Simons Genome Diversity Project,1000 Genomes 30x on GRCh38,1000 Genomes phase 3 release |
| **HG02497** | female | SAME1839682 | ACB | African Caribbean | AFR | African Ancestry | ACB | 1000 Genomes on GRCh38,1000 Genomes 30x on GRCh38,1000 Genomes phase 3 release |
| **HG02948** | male | SAME1839153 | ESN | Esan | AFR | African Ancestry | ESN | 1000 Genomes 30x on GRCh38 |
| **HG02505** | female | SAME1839351 | ACB | African Caribbean | AFR | African Ancestry | ACB | 1000 Genomes on GRCh38,1000 Genomes 30x on GRCh38,1000 Genomes phase 3 release |
| **HG03066** | male | SAME1839324 | MSL | Mende | AFR | African Ancestry | MSL | 1000 Genomes on GRCh38,1000 Genomes 30x on GRCh38,1000 Genomes phase 3 release |
| **HG02656** | male | SAME1839607 | PJL | Punjabi | SAS | South Asian Ancestry | PJL | 1000 Genomes 30x on GRCh38,1000 Genomes phase 3 release |
| **HG03073** | female | SAME1839019 | MSL | Mende | AFR | African Ancestry | MSL | 1000 Genomes on GRCh38,1000 Genomes 30x on GRCh38,1000 Genomes phase 3 release |
| **HG02668** | male | SAME1839377 | GWD | Gambian Mandinka | AFR | African Ancestry | GWD | 1000 Genomes 30x on GRCh38 |
| **HG03078** | male | SAME1839047 | MSL | Mende,Mende | AFR | African Ancestry,Africa (SGDP) | MSL,MendeSGDP | 1000 Genomes on GRCh38,Simons Genome Diversity Project,1000 Genomes 30x on GRCh38,1000 Genomes phase 3 release |
| **HG02675** | male | SAME1839495 | GWD | Gambian Mandinka | AFR | African Ancestry | GWD | 1000 Genomes on GRCh38,1000 Genomes 30x on GRCh38,1000 Genomes phase 3 release,Gambian Genome Variation Project (GRCh38) |
| **HG02682** | female | SAME1839300 | PJL | Punjabi | SAS | South Asian Ancestry | PJL | 1000 Genomes on GRCh38,1000 Genomes 30x on GRCh38,1000 Genomes phase 3 release |
| **HG03085** | female | SAME1840328 | MSL | Mende,Mende | AFR | African Ancestry,Africa (SGDP) | MSL,MendeSGDP | 1000 Genomes on GRCh38,Simons Genome Diversity Project,1000 Genomes 30x on GRCh38,1000 Genomes phase 3 release |
| **HG02687** | male | SAME1839270 | PJL | Punjabi | SAS | South Asian Ancestry | PJL | 1000 Genomes on GRCh38,1000 Genomes 30x on GRCh38,1000 Genomes phase 3 release |
| **HG03097** | female | SAME1840130 | MSL | Mende | AFR | African Ancestry | MSL | 1000 Genomes on GRCh38,1000 Genomes 30x on GRCh38,1000 Genomes phase 3 release |
| **HG02613** | male | SAME1840390 | GWD | Gambian Mandinka | AFR | African Ancestry | GWD | 1000 Genomes on GRCh38,1000 Genomes 30x on GRCh38,1000 Genomes phase 3 release,Gambian Genome Variation Project (GRCh38) |
| **HG03249** | male | SAME1839183 | GWD | Gambian Mandinka | AFR | African Ancestry | GWD | 1000 Genomes on GRCh38,1000 Genomes 30x on GRCh38,Gambian Genome Variation Project (GRCh38) |
| **HG02620** | male | SAME1840191 | GWD | Gambian Mandinka | AFR | African Ancestry | GWD | 1000 Genomes on GRCh38,1000 Genomes 30x on GRCh38,1000 Genomes phase 3 release,Gambian Genome Variation Project (GRCh38) |
| **HG03251** | female | SAME1840266 | GWD | Gambian Mandinka | AFR | African Ancestry | GWD | 1000 Genomes 30x on GRCh38 |
| **HG02625** | male | SAME1840257 | GWD | Gambian Mandinka | AFR | African Ancestry | GWD | 1000 Genomes 30x on GRCh38 |
| **HG03268** | male | SAME1840093 | ESN | Esan | AFR | African Ancestry | ESN | 1000 Genomes on GRCh38,1000 Genomes 30x on GRCh38,1000 Genomes phase 3 release |
| **HG02644** | male | SAME1839782 | GWD | Gambian Mandinka | AFR | African Ancestry | GWD | 1000 Genomes 30x on GRCh38 |
| **HG03270** | female | SAME1840194 | ESN | Esan | AFR | African Ancestry | ESN | 1000 Genomes on GRCh38,1000 Genomes 30x on GRCh38,1000 Genomes phase 3 release |
| **HG02649** | female | SAME1839852 | PJL | Punjabi | SAS | South Asian Ancestry | PJL | 1000 Genomes on GRCh38,1000 Genomes 30x on GRCh38,1000 Genomes phase 3 release |
| **HG03294** | female | SAME1839877 | ESN | Esan | AFR | African Ancestry | ESN | 1000 Genomes on GRCh38,1000 Genomes 30x on GRCh38,1000 Genomes phase 3 release |
| **HG02651** | male | SAME1839544 | PJL | Punjabi | SAS | South Asian Ancestry | PJL | 1000 Genomes on GRCh38,1000 Genomes 30x on GRCh38,1000 Genomes phase 3 release |
| **HG03100** | male | SAME1840003 | ESN | Esan,Esan | AFR | African Ancestry,Africa (SGDP) | ESN,EsanSGDP | 1000 Genomes on GRCh38,Simons Genome Diversity Project,1000 Genomes 30x on GRCh38,1000 Genomes phase 3 release |
| **HG02694** | female | SAME1839065 | PJL | Punjabi | SAS | South Asian Ancestry | PJL | 1000 Genomes on GRCh38,1000 Genomes 30x on GRCh38,1000 Genomes phase 3 release |
| **HG03105** | female | SAME1840041 | ESN | Esan | AFR | African Ancestry | ESN | 1000 Genomes on GRCh38,1000 Genomes 30x on GRCh38,1000 Genomes phase 3 release |
| **HG02699** | male | SAME1839129 | PJL | Punjabi | SAS | South Asian Ancestry | PJL | 1000 Genomes on GRCh38,1000 Genomes 30x on GRCh38,1000 Genomes phase 3 release |
| **HG03112** | male | SAME1839893 | ESN | Esan | AFR | African Ancestry | ESN | 1000 Genomes on GRCh38,1000 Genomes 30x on GRCh38,1000 Genomes phase 3 release |
| **HG02702** | male | SAME1840149 | GWD | Gambian Mandinka | AFR | African Ancestry | GWD | 1000 Genomes on GRCh38,1000 Genomes 30x on GRCh38,1000 Genomes phase 3 release,Gambian Genome Variation Project (GRCh38) |
| **HG03117** | female | SAME1839821 | ESN | Esan | AFR | African Ancestry | ESN | 1000 Genomes on GRCh38,1000 Genomes 30x on GRCh38,1000 Genomes phase 3 release |
| **HG02721** | male | SAME1839749 | GWD | Gambian Mandinka | AFR | African Ancestry | GWD | 1000 Genomes on GRCh38,1000 Genomes 30x on GRCh38,1000 Genomes phase 3 release,Gambian Genome Variation Project (GRCh38) |
| **HG03352** | male | SAME1840243 | ESN | Esan | AFR | African Ancestry | ESN | 1000 Genomes on GRCh38,1000 Genomes 30x on GRCh38,1000 Genomes phase 3 release |
| **HG02726** | female | SAME1839732 | PJL | Punjabi | SAS | South Asian Ancestry | PJL | 1000 Genomes 30x on GRCh38,1000 Genomes phase 3 release |
| **HG03369** | female | SAME1840078 | ESN | Esan | AFR | African Ancestry | ESN | 1000 Genomes on GRCh38,1000 Genomes 30x on GRCh38,1000 Genomes phase 3 release |
| **HG02512** | male | SAME1839146 | KHV | Kinh Vietnamese | EAS | East Asian Ancestry | KHV | 1000 Genomes on GRCh38,1000 Genomes 30x on GRCh38,1000 Genomes phase 3 release |
| **HG03371** | male | SAME1839712 | ESN | Esan | AFR | African Ancestry | ESN | 1000 Genomes 30x on GRCh38,Human Genome Structural Variation Consortium, Phase 2 |
| **HG02524** | male | SAME1840317 | KHV | Kinh Vietnamese | EAS | East Asian Ancestry | KHV | 1000 Genomes on GRCh38,1000 Genomes 30x on GRCh38 |
| **HG03376** | male | SAME1839736 | MSL | Mende | AFR | African Ancestry | MSL | 1000 Genomes on GRCh38,1000 Genomes 30x on GRCh38,1000 Genomes phase 3 release |
| **HG02536** | male | SAME1840172 | ACB | African Caribbean | AFR | African Ancestry | ACB | 1000 Genomes on GRCh38,1000 Genomes 30x on GRCh38,1000 Genomes phase 3 release |
| **HG02555** | female | SAME1840070 | ACB | African Caribbean | AFR | African Ancestry | ACB | 1000 Genomes on GRCh38,1000 Genomes 30x on GRCh38,1000 Genomes phase 3 release |
| **HG03388** | male | SAME1839524 | MSL | Mende | AFR | African Ancestry | MSL | 1000 Genomes on GRCh38,1000 Genomes 30x on GRCh38,1000 Genomes phase 3 release |
| **HG03027** | male | SAME1839725 | GWD | Gambian Mandinka | AFR | African Ancestry | GWD | 1000 Genomes on GRCh38,1000 Genomes 30x on GRCh38,1000 Genomes phase 3 release |
| **HG03034** | female | SAME1839568 | GWD | Gambian Mandinka | AFR | African Ancestry | GWD | 1000 Genomes on GRCh38,1000 Genomes 30x on GRCh38,Gambian Genome Variation Project (GRCh38) |
| **HG03039** | male | SAME1839593 | GWD | Gambian Mandinka | AFR | African Ancestry | GWD | 1000 Genomes on GRCh38,1000 Genomes 30x on GRCh38,1000 Genomes phase 3 release,Gambian Genome Variation Project (GRCh38) |
| **HG03410** | female | SAME1839182 | MSL | Mende | AFR | African Ancestry | MSL | 1000 Genomes on GRCh38,1000 Genomes 30x on GRCh38,1000 Genomes phase 3 release |
| **HG03041** | female | SAME1839620 | GWD | Gambian Mandinka | AFR | African Ancestry | GWD | 1000 Genomes 30x on GRCh38 |
| **HG03198** | female | SAME1839933 | ESN | Esan | AFR | African Ancestry | ESN | 1000 Genomes on GRCh38,1000 Genomes 30x on GRCh38,1000 Genomes phase 3 release |
| **HG03046** | female | SAME1839639 | GWD | Gambian Mandinka | AFR | African Ancestry | GWD | 1000 Genomes on GRCh38,1000 Genomes 30x on GRCh38,1000 Genomes phase 3 release,Gambian Genome Variation Project (GRCh38) |
| **HG03225** | male | SAME1839460 | MSL | Mende | AFR | African Ancestry | MSL | 1000 Genomes on GRCh38,1000 Genomes 30x on GRCh38,1000 Genomes phase 3 release |
| **HG03237** | male | SAME1839282 | PJL | Punjabi | SAS | South Asian Ancestry | PJL | 1000 Genomes on GRCh38,1000 Genomes 30x on GRCh38,1000 Genomes phase 3 release |
| **HG03058** | female | SAME1839442 | MSL | Mende | AFR | African Ancestry | MSL | 1000 Genomes on GRCh38,1000 Genomes 30x on GRCh38,1000 Genomes phase 3 release |
| **HG03124** | male | SAME1839669 | ESN | Esan | AFR | African Ancestry | ESN | 1000 Genomes on GRCh38,1000 Genomes 30x on GRCh38,1000 Genomes phase 3 release |
| **HG03060** | male | SAME1839211 | MSL | Mende | AFR | African Ancestry | MSL | 1000 Genomes on GRCh38,1000 Genomes 30x on GRCh38,1000 Genomes phase 3 release |
| **HG03129** | female | SAME1839699 | ESN | Esan | AFR | African Ancestry | ESN | 1000 Genomes on GRCh38,1000 Genomes 30x on GRCh38,1000 Genomes phase 3 release |
| **HG03065** | male | SAME1839318 | MSL | Mende | AFR | African Ancestry | MSL | 1000 Genomes 30x on GRCh38,Human Genome Structural Variation Consortium, Phase 2 |
| **HG03131** | male | SAME1839368 | ESN | Esan | AFR | African Ancestry | ESN | 1000 Genomes 30x on GRCh38 |
| **HG03072** | male | SAME1839016 | MSL | Mende | AFR | African Ancestry | MSL | 1000 Genomes on GRCh38,1000 Genomes 30x on GRCh38,1000 Genomes phase 3 release |
| **HG03136** | male | SAME1839387 | ESN | Esan | AFR | African Ancestry | ESN | 1000 Genomes on GRCh38,1000 Genomes 30x on GRCh38,1000 Genomes phase 3 release |
| **HG03077** | male | SAME1839044 | MSL | Mende | AFR | African Ancestry | MSL | 1000 Genomes on GRCh38,1000 Genomes 30x on GRCh38,1000 Genomes phase 3 release |
| **HG03299** | male | SAME1839805 | ESN | Esan | AFR | African Ancestry | ESN | 1000 Genomes 30x on GRCh38 |
| **HG03084** | male | SAME1840324 | MSL | Mende | AFR | African Ancestry | MSL | 1000 Genomes on GRCh38,1000 Genomes 30x on GRCh38,1000 Genomes phase 3 release |
| **HG03302** | male | SAME1839479 | ESN | Esan | AFR | African Ancestry | ESN | 1000 Genomes 30x on GRCh38 |
| **HG03091** | female | SAME1840360 | MSL | Mende | AFR | African Ancestry | MSL | 1000 Genomes on GRCh38,1000 Genomes 30x on GRCh38,1000 Genomes phase 3 release |
| **HG03307** | male | SAME1839509 | ESN | Esan | AFR | African Ancestry | ESN | 1000 Genomes on GRCh38,1000 Genomes 30x on GRCh38 |
| **HG03096** | male | SAME1840126 | MSL | Mende | AFR | African Ancestry | MSL | 1000 Genomes on GRCh38,1000 Genomes 30x on GRCh38,1000 Genomes phase 3 release |
| **HG03314** | male | SAME1839217 | ESN | Esan | AFR | African Ancestry | ESN | 1000 Genomes 30x on GRCh38 |
| **HG03248** | male | SAME1839180 | GWD | Gambian Mandinka | AFR | African Ancestry | GWD | 1000 Genomes 30x on GRCh38 |
| **HG03162** | female | SAME1839112 | ESN | Esan | AFR | African Ancestry | ESN | 1000 Genomes on GRCh38,1000 Genomes 30x on GRCh38,1000 Genomes phase 3 release |
| **HG03250** | female | SAME1840262 | GWD | Gambian Mandinka | AFR | African Ancestry | GWD | 1000 Genomes on GRCh38,1000 Genomes 30x on GRCh38,Gambian Genome Variation Project (GRCh38) |
| **HG03267** | female | SAME1840088 | ESN | Esan | AFR | African Ancestry | ESN | 1000 Genomes on GRCh38,1000 Genomes 30x on GRCh38,1000 Genomes phase 3 release |
| **HG03193** | male | SAME1839909 | ESN | Esan | AFR | African Ancestry | ESN | 1000 Genomes on GRCh38,1000 Genomes 30x on GRCh38,1000 Genomes phase 3 release |
| **HG03279** | female | SAME1839972 | ESN | Esan | AFR | African Ancestry | ESN | 1000 Genomes on GRCh38,1000 Genomes 30x on GRCh38,1000 Genomes phase 3 release |
| **HG03605** | female | SAME1840240 | BEB | Bengali | SAS | South Asian Ancestry | BEB | 1000 Genomes 30x on GRCh38 |
| **HG03617** | female | SAME1840056 | BEB | Bengali | SAS | South Asian Ancestry | BEB | 1000 Genomes 30x on GRCh38 |
| **HG03197** | male | SAME1839926 | ESN | Esan | AFR | African Ancestry | ESN | 1000 Genomes 30x on GRCh38 |
| **HG03624** | male | SAME1839709 | PJL | Punjabi | SAS | South Asian Ancestry | PJL | 1000 Genomes on GRCh38,1000 Genomes 30x on GRCh38,1000 Genomes phase 3 release |
| **HG03200** | female | SAME1839589 | ESN | Esan | AFR | African Ancestry | ESN | 1000 Genomes 30x on GRCh38 |
| **HG03629** | male | SAME1839738 | PJL | Punjabi | SAS | South Asian Ancestry | PJL | 1000 Genomes on GRCh38,1000 Genomes 30x on GRCh38,1000 Genomes phase 3 release |
| **HG03212** | female | SAME1839399 | MSL | Mende | AFR | African Ancestry | MSL | 1000 Genomes on GRCh38,1000 Genomes 30x on GRCh38,1000 Genomes phase 3 release |
| **HG03631** | female | SAME1839855 | PJL | Punjabi | SAS | South Asian Ancestry | PJL | 1000 Genomes on GRCh38,1000 Genomes 30x on GRCh38,1000 Genomes phase 3 release |
| **HG03224** | male | SAME1839455 | MSL | Mende | AFR | African Ancestry | MSL | 1000 Genomes on GRCh38,1000 Genomes 30x on GRCh38,1000 Genomes phase 3 release |
| **HG03636** | male | SAME1839586 | PJL | Punjabi | SAS | South Asian Ancestry | PJL | 1000 Genomes on GRCh38,1000 Genomes 30x on GRCh38,1000 Genomes phase 3 release |
| **HG03229** | female | SAME1839234 | PJL | Punjabi | SAS | South Asian Ancestry | PJL | 1000 Genomes on GRCh38,1000 Genomes 30x on GRCh38,1000 Genomes phase 3 release |
| **HG03698** | female | SAME1840145 | STU | Tamil | SAS | South Asian Ancestry | STU | 1000 Genomes on GRCh38,1000 Genomes 30x on GRCh38,1000 Genomes phase 3 release |
| **HG03236** | female | SAME1839276 | PJL | Punjabi | SAS | South Asian Ancestry | PJL | 1000 Genomes 30x on GRCh38 |
| **HG03701** | male | SAME1840016 | PJL | Punjabi | SAS | South Asian Ancestry | PJL | 1000 Genomes 30x on GRCh38 |
| **HG02973** | male | SAME1840246 | ESN | Esan | AFR | African Ancestry | ESN | 1000 Genomes on GRCh38,1000 Genomes 30x on GRCh38,1000 Genomes phase 3 release |
| **HG03706** | female | SAME1839758 | PJL | Punjabi | SAS | South Asian Ancestry | PJL | 1000 Genomes on GRCh38,1000 Genomes 30x on GRCh38,1000 Genomes phase 3 release |
| **HG02978** | female | SAME1840216 | ESN | Esan | AFR | African Ancestry | ESN | 1000 Genomes 30x on GRCh38 |
| **HG03713** | male | SAME1839807 | ITU | Telugu | SAS | South Asian Ancestry | ITU | 1000 Genomes on GRCh38,1000 Genomes 30x on GRCh38,1000 Genomes phase 3 release |
| **HG02980** | male | SAME1840034 | ESN | Esan | AFR | African Ancestry | ESN | 1000 Genomes 30x on GRCh38 |
| **HG03489** | male | SAME1839494 | PJL | Punjabi | SAS | South Asian Ancestry | PJL | 1000 Genomes 30x on GRCh38 |
| **HG03008** | male | SAME1840253 | BEB | Bengali | SAS | South Asian Ancestry | BEB | 1000 Genomes 30x on GRCh38 |
| **HG03491** | female | SAME1839264 | PJL | Punjabi | SAS | South Asian Ancestry | PJL | 1000 Genomes on GRCh38,1000 Genomes 30x on GRCh38,1000 Genomes phase 3 release |
| **HG03015** | male | SAME1839940 | PJL | Punjabi | SAS | South Asian Ancestry | PJL | 1000 Genomes on GRCh38,1000 Genomes 30x on GRCh38,1000 Genomes phase 3 release |
| **HG03511** | female | SAME1840131 | ESN | Esan | AFR | African Ancestry | ESN | 1000 Genomes on GRCh38,1000 Genomes 30x on GRCh38,1000 Genomes phase 3 release |
| **HG03022** | female | SAME1839795 | PJL | Punjabi | SAS | South Asian Ancestry | PJL | 1000 Genomes on GRCh38,1000 Genomes 30x on GRCh38,1000 Genomes phase 3 release |
| **HG03516** | female | SAME1840160 | ESN | Esan | AFR | African Ancestry | ESN | 1000 Genomes 30x on GRCh38 |
| **HG03123** | female | SAME1839663 | ESN | Esan | AFR | African Ancestry | ESN | 1000 Genomes on GRCh38,1000 Genomes 30x on GRCh38,1000 Genomes phase 3 release |
| **HG03460** | male | SAME1839908 | MSL | Mende | AFR | African Ancestry | MSL | 1000 Genomes on GRCh38,1000 Genomes 30x on GRCh38,1000 Genomes phase 3 release |
| **HG03128** | male | SAME1839692 | ESN | Esan | AFR | African Ancestry | ESN | 1000 Genomes 30x on GRCh38 |
| **HG03130** | male | SAME1839365 | ESN | Esan | AFR | African Ancestry | ESN | 1000 Genomes on GRCh38,1000 Genomes 30x on GRCh38,1000 Genomes phase 3 release |
| **HG03472** | male | SAME1839678 | MSL | Mende | AFR | African Ancestry | MSL | 1000 Genomes on GRCh38,1000 Genomes 30x on GRCh38,1000 Genomes phase 3 release |
| **HG03135** | female | SAME1839385 | ESN | Esan | AFR | African Ancestry | ESN | 1000 Genomes on GRCh38,1000 Genomes 30x on GRCh38,1000 Genomes phase 3 release |
| **HG03159** | female | SAME1839092 | ESN | Esan | AFR | African Ancestry | ESN | 1000 Genomes on GRCh38,1000 Genomes 30x on GRCh38,1000 Genomes phase 3 release |
| **HG03484** | male | SAME1839393 | MSL | Mende | AFR | African Ancestry | MSL | 1000 Genomes on GRCh38,1000 Genomes 30x on GRCh38,1000 Genomes phase 3 release |
| **HG03439** | male | SAME1840213 | MSL | Mende | AFR | African Ancestry | MSL | 1000 Genomes on GRCh38,1000 Genomes 30x on GRCh38,1000 Genomes phase 3 release |

| | | | | | | | | |
|---|---|---|---|---|---|---|---|---|
| **HG03109** | male | SAME1839765 | ESN | Esan | AFR | African Ancestry | ESN | 1000 Genomes on GRCh38,1000 Genomes 30x on GRCh38,1000 Genomes phase 3 release |
| **HG03446** | female | SAME1839911 | MSL | Mende | AFR | African Ancestry | MSL | 1000 Genomes on GRCh38,1000 Genomes 30x on GRCh38,1000 Genomes phase 3 release |
| **HG03111** | female | SAME1839884 | ESN | Esan | AFR | African Ancestry | ESN | 1000 Genomes on GRCh38,1000 Genomes 30x on GRCh38,1000 Genomes phase 3 release |
| **HG03453** | female | SAME1839750 | MSL | Mende | AFR | African Ancestry | MSL | 1000 Genomes 30x on GRCh38 |
| **HG03116** | female | SAME1839816 | ESN | Esan | AFR | African Ancestry | ESN | 1000 Genomes 30x on GRCh38 |
| **HG03458** | female | SAME1839786 | MSL | Mende | AFR | African Ancestry | MSL | 1000 Genomes on GRCh38,1000 Genomes 30x on GRCh38,1000 Genomes phase 3 release |
| **HG03643** | female | SAME1839633 | STU | Tamil | SAS | South Asian Ancestry | STU | 1000 Genomes on GRCh38,1000 Genomes 30x on GRCh38,1000 Genomes phase 3 release |
| **HG02947** | male | SAME1839061 | ESN | Esan | AFR | African Ancestry | ESN | 1000 Genomes on GRCh38,1000 Genomes 30x on GRCh38,1000 Genomes phase 3 release |
| **HG03650** | female | SAME1839488 | PJL | Punjabi | SAS | South Asian Ancestry | PJL | 1000 Genomes on GRCh38,1000 Genomes 30x on GRCh38 |
| **HG02954** | male | SAME1840264 | ESN | Esan | AFR | African Ancestry | ESN | 1000 Genomes 30x on GRCh38 |
| **HG03667** | male | SAME1839331 | PJL | Punjabi | SAS | South Asian Ancestry | PJL | 1000 Genomes on GRCh38,1000 Genomes 30x on GRCh38,1000 Genomes phase 3 release |
| **HG02966** | male | SAME1840114 | ESN | Esan | AFR | African Ancestry | ESN | 1000 Genomes 30x on GRCh38 |
| **HG03578** | female | SAME1839245 | MSL | Mende | AFR | African Ancestry | MSL | 1000 Genomes on GRCh38,1000 Genomes 30x on GRCh38,1000 Genomes phase 3 release |
| **HG03161** | male | SAME1839104 | ESN | Esan | AFR | African Ancestry | ESN | 1000 Genomes 30x on GRCh38 |
| **HG03585** | male | SAME1839293 | BEB | Bengali | SAS | South Asian Ancestry | BEB | 1000 Genomes on GRCh38,1000 Genomes 30x on GRCh38,1000 Genomes phase 3 release |
| **HG03166** | male | SAME1839127 | ESN | Esan | AFR | African Ancestry | ESN | 1000 Genomes on GRCh38,1000 Genomes 30x on GRCh38,1000 Genomes phase 3 release |
| **HG03600** | male | SAME1840215 | BEB | Bengali | SAS | South Asian Ancestry | BEB | 1000 Genomes on GRCh38,1000 Genomes 30x on GRCh38,1000 Genomes phase 3 release |
| **HG03679** | male | SAME1839055 | STU | Tamil | SAS | South Asian Ancestry | STU | 1000 Genomes on GRCh38,1000 Genomes 30x on GRCh38,1000 Genomes phase 3 release |
| **HG03522** | female | SAME1840193 | ESN | Esan | AFR | African Ancestry | ESN | 1000 Genomes 30x on GRCh38 |
| **HG03681** | male | SAME1840321 | STU | Tamil | SAS | South Asian Ancestry | STU | 1000 Genomes on GRCh38,1000 Genomes 30x on GRCh38,1000 Genomes phase 3 release |
| **HG03539** | female | SAME1840014 | GWD | Gambian Mandinka | AFR | African Ancestry | GWD | 1000 Genomes on GRCh38,1000 Genomes 30x on GRCh38,1000 Genomes phase 3 release,Gambian Genome Variation Project (GRCh38) |
| **HG03686** | male | SAME1840335 | STU | Tamil | SAS | South Asian Ancestry | STU | 1000 Genomes on GRCh38,1000 Genomes 30x on GRCh38,1000 Genomes phase 3 release |
| **HG03558** | female | SAME1839561 | MSL | Mende | AFR | African Ancestry | MSL | 1000 Genomes on GRCh38,1000 Genomes 30x on GRCh38,1000 Genomes phase 3 release |
| **HG03693** | male | SAME1840372 | STU | Tamil | SAS | South Asian Ancestry | STU | 1000 Genomes on GRCh38,1000 Genomes 30x on GRCh38,1000 Genomes phase 3 release |
| **HG03565** | male | SAME1839430 | MSL | Mende | AFR | African Ancestry | MSL | 1000 Genomes on GRCh38,1000 Genomes 30x on GRCh38,1000 Genomes phase 3 release |
| **HG03547** | male | SAME1839878 | MSL | Mende | AFR | African Ancestry | MSL | 1000 Genomes on GRCh38,1000 Genomes 30x on GRCh38,1000 Genomes phase 3 release |
| **HG03572** | female | SAME1839463 | MSL | Mende | AFR | African Ancestry | MSL | 1000 Genomes on GRCh38,1000 Genomes 30x on GRCh38,1000 Genomes phase 3 release |
| **HG03559** | male | SAME1839654 | MSL | Mende | AFR | African Ancestry | MSL | 1000 Genomes on GRCh38,1000 Genomes 30x on GRCh38,1000 Genomes phase 3 release |
| **HG03577** | male | SAME1839242 | MSL | Mende | AFR | African Ancestry | MSL | 1000 Genomes on GRCh38,1000 Genomes 30x on GRCh38,1000 Genomes phase 3 release |
| **HG03584** | female | SAME1839290 | MSL | Mende | AFR | African Ancestry | MSL | 1000 Genomes 30x on GRCh38,1000 Genomes phase 3 release |
| **HG03718** | male | SAME1839834 | ITU | Telugu | SAS | South Asian Ancestry | ITU | 1000 Genomes on GRCh38,1000 Genomes 30x on GRCh38,1000 Genomes phase 3 release |
| **HG03589** | female | SAME1839311 | BEB | Bengali | SAS | South Asian Ancestry | BEB | 1000 Genomes on GRCh38,1000 Genomes 30x on GRCh38,1000 Genomes phase 3 release |
| **HG03720** | male | SAME1839562 | ITU | Telugu | SAS | South Asian Ancestry | ITU | 1000 Genomes on GRCh38,1000 Genomes 30x on GRCh38,1000 Genomes phase 3 release |
| **HG03596** | male | SAME1839188 | BEB | Bengali | SAS | South Asian Ancestry | BEB | 1000 Genomes 30x on GRCh38 |
| **HG03725** | male | SAME1839680 | ITU | Telugu | SAS | South Asian Ancestry | ITU | 1000 Genomes on GRCh38,1000 Genomes 30x on GRCh38 |
| **HG03604** | female | SAME1840236 | BEB | Bengali | SAS | South Asian Ancestry | BEB | 1000 Genomes on GRCh38,1000 Genomes 30x on GRCh38,1000 Genomes phase 3 release |
| **HG03732** | male | SAME1839378 | ITU | Telugu | SAS | South Asian Ancestry | ITU | 1000 Genomes 30x on GRCh38,Human Genome Structural Variation Consortium, Phase 2 |
| **HG03488** | female | SAME1839486 | PJL | Punjabi | SAS | South Asian Ancestry | PJL | 1000 Genomes on GRCh38,1000 Genomes 30x on GRCh38,1000 Genomes phase 3 release |
| **HG03744** | male | SAME1839265 | STU | Tamil | SAS | South Asian Ancestry | STU | 1000 Genomes on GRCh38,1000 Genomes 30x on GRCh38,1000 Genomes phase 3 release |
| **HG03490** | male | SAME1839259 | PJL | Punjabi | SAS | South Asian Ancestry | PJL | 1000 Genomes on GRCh38,1000 Genomes 30x on GRCh38,1000 Genomes phase 3 release |
| **HG03821** | male | SAME1839449 | BEB | Bengali | SAS | South Asian Ancestry | BEB | 1000 Genomes on GRCh38,1000 Genomes 30x on GRCh38,1000 Genomes phase 3 release |
| **HG03826** | female | SAME1839465 | BEB | Bengali | SAS | South Asian Ancestry | BEB | 1000 Genomes on GRCh38,1000 Genomes 30x on GRCh38,1000 Genomes phase 3 release |
| **HG03833** | male | SAME1839356 | BEB | Bengali | SAS | South Asian Ancestry | BEB | 1000 Genomes on GRCh38,1000 Genomes 30x on GRCh38,1000 Genomes phase 3 release |
| **HG03515** | male | SAME1840154 | ESN | Esan | AFR | African Ancestry | ESN | 1000 Genomes on GRCh38,1000 Genomes 30x on GRCh38,1000 Genomes phase 3 release |
| **HG03838** | female | SAME1839297 | STU | Tamil | SAS | South Asian Ancestry | STU | 1000 Genomes on GRCh38,1000 Genomes 30x on GRCh38,1000 Genomes phase 3 release |
| **HG03433** | male | SAME1840110 | MSL | Mende | AFR | African Ancestry | MSL | 1000 Genomes on GRCh38,1000 Genomes 30x on GRCh38,1000 Genomes phase 3 release |
| **HG03438** | female | SAME1840208 | MSL | Mende | AFR | African Ancestry | MSL | 1000 Genomes 30x on GRCh38 |
| **HG03445** | male | SAME1839996 | MSL | Mende | AFR | African Ancestry | MSL | 1000 Genomes on GRCh38,1000 Genomes 30x on GRCh38,1000 Genomes phase 3 release |
| **HG03934** | female | SAME1840318 | BEB | Bengali | SAS | South Asian Ancestry | BEB | 1000 Genomes on GRCh38,1000 Genomes 30x on GRCh38,1000 Genomes phase 3 release |
| **HG03452** | female | SAME1840046 | MSL | Mende | AFR | African Ancestry | MSL | 1000 Genomes on GRCh38,1000 Genomes 30x on GRCh38,1000 Genomes phase 3 release |
| **HG03457** | male | SAME1839781 | MSL | Mende | AFR | African Ancestry | MSL | 1000 Genomes on GRCh38,1000 Genomes 30x on GRCh38,1000 Genomes phase 3 release |
| **HG03941** | male | SAME1840353 | BEB | Bengali | SAS | South Asian Ancestry | BEB | 1000 Genomes on GRCh38,1000 Genomes 30x on GRCh38,1000 Genomes phase 3 release |
| **HG03344** | male | SAME1840364 | ESN | Esan | AFR | African Ancestry | ESN | 1000 Genomes 30x on GRCh38 |
| **HG03953** | male | SAME1840252 | STU | Tamil | SAS | South Asian Ancestry | STU | 1000 Genomes on GRCh38,1000 Genomes 30x on GRCh38,1000 Genomes phase 3 release |
| **HG03351** | female | SAME1840231 | ESN | Esan | AFR | African Ancestry | ESN | 1000 Genomes on GRCh38,1000 Genomes 30x on GRCh38,1000 Genomes phase 3 release |
| **HG03960** | male | SAME1839945 | ITU | Telugu | SAS | South Asian Ancestry | ITU | 1000 Genomes on GRCh38,1000 Genomes 30x on GRCh38,1000 Genomes phase 3 release |
| **HG03363** | female | SAME1839957 | ESN | Esan | AFR | African Ancestry | ESN | 1000 Genomes on GRCh38,1000 Genomes 30x on GRCh38,1000 Genomes phase 3 release |
| **HG03368** | male | SAME1840074 | ESN | Esan | AFR | African Ancestry | ESN | 1000 Genomes 30x on GRCh38 |
| **HG03908** | male | SAME1839522 | BEB | Bengali | SAS | South Asian Ancestry | BEB | 1000 Genomes on GRCh38,1000 Genomes 30x on GRCh38,1000 Genomes phase 3 release |
| **HG03370** | male | SAME1839800 | ESN | Esan | AFR | African Ancestry | ESN | 1000 Genomes on GRCh38,1000 Genomes 30x on GRCh38,1000 Genomes phase 3 release |
| **HG03910** | female | SAME1839205 | BEB | Bengali | SAS | South Asian Ancestry | BEB | 1000 Genomes on GRCh38,1000 Genomes 30x on GRCh38,1000 Genomes phase 3 release |
| **HG03382** | male | SAME1839580 | MSL | Mende | AFR | African Ancestry | MSL | 1000 Genomes on GRCh38,1000 Genomes 30x on GRCh38,1000 Genomes phase 3 release |
| **HG03915** | female | SAME1839227 | BEB | Bengali | SAS | South Asian Ancestry | BEB | 1000 Genomes 30x on GRCh38 |
| **HG03394** | male | SAME1839648 | MSL | Mende | AFR | African Ancestry | MSL | 1000 Genomes on GRCh38,1000 Genomes 30x on GRCh38,1000 Genomes phase 3 release |
| **HG03922** | female | SAME1839103 | BEB | Bengali | SAS | South Asian Ancestry | BEB | 1000 Genomes on GRCh38,1000 Genomes 30x on GRCh38,1000 Genomes phase 3 release |
| **HG03927** | female | SAME1839035 | BEB | Bengali | SAS | South Asian Ancestry | BEB | 1000 Genomes 30x on GRCh38 |
| **HG03799** | female | SAME1839944 | BEB | Bengali | SAS | South Asian Ancestry | BEB | 1000 Genomes on GRCh38,1000 Genomes 30x on GRCh38 |
| **HG03419** | female | SAME1839130 | MSL | Mende | AFR | African Ancestry | MSL | 1000 Genomes on GRCh38,1000 Genomes 30x on GRCh38,1000 Genomes phase 3 release |
| **HG03802** | female | SAME1839608 | BEB | Bengali | SAS | South Asian Ancestry | BEB | 1000 Genomes on GRCh38,1000 Genomes 30x on GRCh38,1000 Genomes phase 3 release |
| **HG03464** | female | SAME1839827 | MSL | Mende | AFR | African Ancestry | MSL | 1000 Genomes on GRCh38,1000 Genomes 30x on GRCh38,1000 Genomes phase 3 release |
| **HG03807** | female | SAME1839547 | BEB | Bengali | SAS | South Asian Ancestry | BEB | 1000 Genomes 30x on GRCh38,Human Genome Structural Variation Consortium, Phase 2 |
| **HG03469** | male | SAME1839850 | MSL | Mende | AFR | African Ancestry | MSL | 1000 Genomes on GRCh38,1000 Genomes 30x on GRCh38,1000 Genomes phase 3 release |
| **HG03814** | female | SAME1839411 | BEB | Bengali | SAS | South Asian Ancestry | BEB | 1000 Genomes on GRCh38,1000 Genomes 30x on GRCh38,1000 Genomes phase 3 release |
| **HG03775** | male | SAME1840413 | ITU | Telugu | SAS | South Asian Ancestry | ITU | 1000 Genomes on GRCh38,1000 Genomes 30x on GRCh38,1000 Genomes phase 3 release |
| **HG03476** | female | SAME1839702 | MSL | Mende | AFR | African Ancestry | MSL | 1000 Genomes on GRCh38,1000 Genomes 30x on GRCh38,1000 Genomes phase 3 release |
| **HG03782** | female | SAME1840119 | ITU | Telugu | SAS | South Asian Ancestry | ITU | 1000 Genomes on GRCh38,1000 Genomes 30x on GRCh38,1000 Genomes phase 3 release |
| **HG03298** | male | SAME1839903 | ESN | Esan | AFR | African Ancestry | ESN | 1000 Genomes on GRCh38,1000 Genomes 30x on GRCh38,1000 Genomes phase 3 release |
| **HG03787** | female | SAME1840224 | ITU | Telugu | SAS | South Asian Ancestry | ITU | 1000 Genomes on GRCh38,1000 Genomes 30x on GRCh38,1000 Genomes phase 3 release |
| **HG03301** | male | SAME1839472 | ESN | Esan | AFR | African Ancestry | ESN | 1000 Genomes on GRCh38,1000 Genomes 30x on GRCh38,1000 Genomes phase 3 release |
| **HG03794** | male | SAME1839920 | BEB | Bengali | SAS | South Asian Ancestry | BEB | 1000 Genomes on GRCh38,1000 Genomes 30x on GRCh38 |
| **HG03306** | female | SAME1839504 | ESN | Esan | AFR | African Ancestry | ESN | 1000 Genomes on GRCh38,1000 Genomes 30x on GRCh38 |
| **HG03756** | female | SAME1839080 | STU | Tamil | SAS | South Asian Ancestry | STU | 1000 Genomes on GRCh38,1000 Genomes 30x on GRCh38,1000 Genomes phase 3 release |
| **HG03313** | male | SAME1839213 | ESN | Esan | AFR | African Ancestry | ESN | 1000 Genomes on GRCh38,1000 Genomes 30x on GRCh38,1000 Genomes phase 3 release |
| **HG03763** | male | SAME1839122 | PJL | Punjabi | SAS | South Asian Ancestry | PJL | 1000 Genomes 30x on GRCh38 |
| **HG03611** | female | SAME1839929 | BEB | Bengali | SAS | South Asian Ancestry | BEB | 1000 Genomes on GRCh38,1000 Genomes 30x on GRCh38,1000 Genomes phase 3 release |
| **HG03770** | female | SAME1840388 | ITU | Telugu | SAS | South Asian Ancestry | ITU | 1000 Genomes on GRCh38,1000 Genomes 30x on GRCh38,1000 Genomes phase 3 release |
| **HG03616** | female | SAME1839955 | BEB | Bengali | SAS | South Asian Ancestry | BEB | 1000 Genomes on GRCh38,1000 Genomes 30x on GRCh38,1000 Genomes phase 3 release |
| **HG03635** | female | SAME1839578 | PJL | Punjabi | SAS | South Asian Ancestry | PJL | 1000 Genomes 30x on GRCh38 |
| **HG03888** | female | SAME1840039 | STU | Tamil | SAS | South Asian Ancestry | STU | 1000 Genomes on GRCh38,1000 Genomes 30x on GRCh38,1000 Genomes phase 3 release |
| **HG03697** | male | SAME1840137 | STU | Tamil | SAS | South Asian Ancestry | STU | 1000 Genomes on GRCh38,1000 Genomes 30x on GRCh38,1000 Genomes phase 3 release |
| **HG03890** | male | SAME1839864 | STU | Tamil | SAS | South Asian Ancestry | STU | 1000 Genomes on GRCh38,1000 Genomes 30x on GRCh38,1000 Genomes phase 3 release |
| **HG03700** | female | SAME1840010 | PJL | Punjabi | SAS | South Asian Ancestry | PJL | 1000 Genomes on GRCh38,1000 Genomes 30x on GRCh38 |
| **HG03895** | female | SAME1839888 | STU | Tamil | SAS | South Asian Ancestry | STU | 1000 Genomes on GRCh38,1000 Genomes 30x on GRCh38,1000 Genomes phase 3 release |
| **HG03705** | male | SAME1840051 | PJL | Punjabi | SAS | South Asian Ancestry | PJL | 1000 Genomes on GRCh38,1000 Genomes 30x on GRCh38,1000 Genomes phase 3 release |
| **HG03857** | female | SAME1840294 | STU | Tamil | SAS | South Asian Ancestry | STU | 1000 Genomes on GRCh38,1000 Genomes 30x on GRCh38,1000 Genomes phase 3 release |
| **HG03717** | female | SAME1839831 | ITU | Telugu | SAS | South Asian Ancestry | ITU | 1000 Genomes on GRCh38,1000 Genomes 30x on GRCh38,1000 Genomes phase 3 release |
| **HG03864** | male | SAME1840170 | ITU | Telugu | SAS | South Asian Ancestry | ITU | 1000 Genomes on GRCh38,1000 Genomes 30x on GRCh38,1000 Genomes phase 3 release |
| **HG03774** | female | SAME1840408 | ITU | Telugu | SAS | South Asian Ancestry | ITU | 1000 Genomes on GRCh38,1000 Genomes 30x on GRCh38,1000 Genomes phase 3 release |
| **HG03869** | male | SAME1840104 | ITU | Telugu | SAS | South Asian Ancestry | ITU | 1000 Genomes on GRCh38,1000 Genomes 30x on GRCh38,1000 Genomes phase 3 release |
| **HG03779** | male | SAME1840351 | ITU | Telugu | SAS | South Asian Ancestry | ITU | 1000 Genomes on GRCh38,1000 Genomes 30x on GRCh38,1000 Genomes phase 3 release |
| **HG03871** | male | SAME1840200 | ITU | Telugu | SAS | South Asian Ancestry | ITU | 1000 Genomes on GRCh38,1000 Genomes 30x on GRCh38,1000 Genomes phase 3 release |
| **HG03781** | female | SAME1840117 | ITU | Telugu | SAS | South Asian Ancestry | ITU | 1000 Genomes on GRCh38,1000 Genomes 30x on GRCh38,1000 Genomes phase 3 release |
| **HG04228** | male | SAME1839161 | STU | Tamil | SAS | South Asian Ancestry | STU | 1000 Genomes 30x on GRCh38 |
| **HG03786** | male | SAME1840218 | ITU | Telugu | SAS | South Asian Ancestry | ITU | 1000 Genomes on GRCh38,1000 Genomes 30x on GRCh38,1000 Genomes phase 3 release |
| **HG04235** | male | SAME1840263 | ITU | Telugu | SAS | South Asian Ancestry | ITU | 1000 Genomes on GRCh38,1000 Genomes 30x on GRCh38,1000 Genomes phase 3 release |
| **HG03793** | female | SAME1839915 | BEB | Bengali | SAS | South Asian Ancestry | BEB | 1000 Genomes on GRCh38,1000 Genomes 30x on GRCh38,1000 Genomes phase 3 release |
| **NA06984** | male | SAME124763 | CEU | CEPH | EUR | European Ancestry | CEU | 1000 Genomes on GRCh38,1000 Genomes 30x on GRCh38,1000 Genomes phase 3 release,1000 Genomes phase 1 release,Geuvadis |
| **HG03825** | female | SAME1839462 | BEB | Bengali | SAS | South Asian Ancestry | BEB | 1000 Genomes 30x on GRCh38 |
| **NA06989** | female | SAME124760 | CEU | CEPH | EUR | European Ancestry | CEU | 1000 Genomes on GRCh38,1000 Genomes 30x on GRCh38,1000 Genomes phase 3 release,1000 Genomes phase 1 release,Geuvadis |
| **HG03832** | female | SAME1839353 | BEB | Bengali | SAS | South Asian Ancestry | BEB | 1000 Genomes on GRCh38,1000 Genomes 30x on GRCh38,1000 Genomes phase 3 release |
| **NA06991** | female | SAME124936 | CEU | CEPH | EUR | European Ancestry | CEU | 1000 Genomes 30x on GRCh38,1000 Genomes phase 3 release |

| | | | | | | | | |
|---|---|---|---|---|---|---|---|---|
| **HG03837** | male | SAME1839291 | STU | Tamil | SAS | South Asian Ancestry | STU | 1000 Genomes on GRCh38,1000 Genomes 30x on GRCh38,1000 Genomes phase 3 release |
| **NA07014** | female | SAME125162 | CEU | CEPH | EUR | European Ancestry | CEU | 1000 Genomes 30x on GRCh38,1000 Genomes phase 3 release |
| **HG03844** | male | SAME1839166 | STU | Tamil | SAS | South Asian Ancestry | STU | 1000 Genomes on GRCh38,1000 Genomes 30x on GRCh38,1000 Genomes phase 3 release |
| **NA07019** | female | SAME125152 | CEU | CEPH | EUR | European Ancestry | CEU | 1000 Genomes 30x on GRCh38,1000 Genomes phase 3 release |
| **HG03750** | male | SAME1839309 | STU | Tamil | SAS | South Asian Ancestry | STU | 1000 Genomes on GRCh38,1000 Genomes 30x on GRCh38,1000 Genomes phase 3 release |
| **HG04090** | female | SAME1839927 | ITU | Telugu | SAS | South Asian Ancestry | ITU | 1000 Genomes on GRCh38,1000 Genomes 30x on GRCh38,1000 Genomes phase 3 release |
| **HG03755** | male | SAME1839074 | STU | Tamil | SAS | South Asian Ancestry | STU | 1000 Genomes on GRCh38,1000 Genomes 30x on GRCh38,1000 Genomes phase 3 release |
| **HG04115** | male | SAME1839362 | STU | Tamil | SAS | South Asian Ancestry | STU | 1000 Genomes 30x on GRCh38 |
| **HG03762** | female | SAME1839119 | PJL | Punjabi | SAS | South Asian Ancestry | PJL | 1000 Genomes on GRCh38,1000 Genomes 30x on GRCh38,1000 Genomes phase 3 release |
| **HG04122** | male | SAME1839230 | STU | Tamil | SAS | South Asian Ancestry | STU | 1000 Genomes 30x on GRCh38 |
| **HG03767** | male | SAME1839141 | PJL | Punjabi | SAS | South Asian Ancestry | PJL | 1000 Genomes on GRCh38,1000 Genomes 30x on GRCh38,1000 Genomes phase 3 release |
| **HG04127** | male | SAME1839249 | STU | Tamil | SAS | South Asian Ancestry | STU | 1000 Genomes on GRCh38,1000 Genomes 30x on GRCh38 |
| **HG03849** | female | SAME1839190 | STU | Tamil | SAS | South Asian Ancestry | STU | 1000 Genomes on GRCh38,1000 Genomes 30x on GRCh38,1000 Genomes phase 3 release |
| **HG04134** | male | SAME1839303 | BEB | Bengali | SAS | South Asian Ancestry | BEB | 1000 Genomes on GRCh38,1000 Genomes 30x on GRCh38,1000 Genomes phase 3 release |
| **HG03851** | male | SAME1840272 | STU | Tamil | SAS | South Asian Ancestry | STU | 1000 Genomes on GRCh38,1000 Genomes 30x on GRCh38,1000 Genomes phase 3 release |
| **NA07045** | female | SAME124187 | CEU | CEPH | EUR | European Ancestry | CEU | 1000 Genomes 30x on GRCh38,1000 Genomes phase 3 release |
| **HG03856** | male | SAME1840291 | STU | Tamil | SAS | South Asian Ancestry | STU | 1000 Genomes on GRCh38,1000 Genomes 30x on GRCh38,1000 Genomes phase 3 release |
| **NA07348** | female | SAME123567 | CEU | CEPH | EUR | European Ancestry | CEU | 1000 Genomes 30x on GRCh38,1000 Genomes phase 3 release |
| **HG03863** | female | SAME1840162 | ITU | Telugu | SAS | South Asian Ancestry | ITU | 1000 Genomes on GRCh38,1000 Genomes 30x on GRCh38,1000 Genomes phase 3 release |
| **NA10839** | female | SAME124551 | CEU | CEPH | EUR | European Ancestry | CEU | 1000 Genomes 30x on GRCh38,1000 Genomes phase 3 release |
| **HG03868** | female | SAME1840097 | ITU | Telugu | SAS | South Asian Ancestry | ITU | 1000 Genomes on GRCh38,1000 Genomes 30x on GRCh38,1000 Genomes phase 3 release |
| **NA10846** | male | SAME124758 | CEU | CEPH | EUR | European Ancestry | CEU | 1000 Genomes 30x on GRCh38,1000 Genomes phase 3 release |
| **HG03870** | male | SAME1840196 | ITU | Telugu | SAS | South Asian Ancestry | ITU | 1000 Genomes on GRCh38,1000 Genomes 30x on GRCh38,1000 Genomes phase 3 release |
| **HG04141** | female | SAME1839178 | BEB | Bengali | SAS | South Asian Ancestry | BEB | 1000 Genomes on GRCh38,1000 Genomes 30x on GRCh38,1000 Genomes phase 3 release |
| **HG03673** | female | SAME1839030 | STU | Tamil | SAS | South Asian Ancestry | STU | 1000 Genomes on GRCh38,1000 Genomes 30x on GRCh38,1000 Genomes phase 3 release |
| **HG04146** | male | SAME1839108 | BEB | Bengali | SAS | South Asian Ancestry | BEB | 1000 Genomes on GRCh38,1000 Genomes 30x on GRCh38,1000 Genomes phase 3 release |
| **HG03680** | male | SAME1840311 | STU | Tamil | SAS | South Asian Ancestry | STU | 1000 Genomes on GRCh38,1000 Genomes 30x on GRCh38,1000 Genomes phase 3 release |
| **HG04153** | female | SAME1840299 | BEB | Bengali | SAS | South Asian Ancestry | BEB | 1000 Genomes on GRCh38,1000 Genomes 30x on GRCh38,1000 Genomes phase 3 release |
| **HG03685** | male | SAME1840333 | STU | Tamil | SAS | South Asian Ancestry | STU | 1000 Genomes on GRCh38,1000 Genomes 30x on GRCh38,1000 Genomes phase 3 release |
| **HG04158** | male | SAME1840400 | BEB | Bengali | SAS | South Asian Ancestry | BEB | 1000 Genomes on GRCh38,1000 Genomes 30x on GRCh38,1000 Genomes phase 3 release |
| **HG03692** | female | SAME1840371 | STU | Tamil | SAS | South Asian Ancestry | STU | 1000 Genomes on GRCh38,1000 Genomes 30x on GRCh38,1000 Genomes phase 3 release |
| **HG04160** | male | SAME1840179 | BEB | Bengali | SAS | South Asian Ancestry | BEB | 1000 Genomes 30x on GRCh38 |
| **HG03642** | female | SAME1839627 | STU | Tamil | SAS | South Asian Ancestry | STU | 1000 Genomes on GRCh38,1000 Genomes 30x on GRCh38,1000 Genomes phase 3 release |
| **HG04204** | male | SAME1839437 | STU | Tamil | SAS | South Asian Ancestry | STU | 1000 Genomes 30x on GRCh38 |
| **HG03654** | male | SAME1839513 | PJL | Punjabi | SAS | South Asian Ancestry | PJL | 1000 Genomes 30x on GRCh38 |
| **HG04209** | female | SAME1839452 | ITU | Telugu | SAS | South Asian Ancestry | ITU | 1000 Genomes on GRCh38,1000 Genomes 30x on GRCh38,1000 Genomes phase 3 release |
| **HG03798** | male | SAME1839941 | BEB | Bengali | SAS | South Asian Ancestry | BEB | 1000 Genomes 30x on GRCh38 |
| **HG04211** | male | SAME1839321 | ITU | Telugu | SAS | South Asian Ancestry | ITU | 1000 Genomes on GRCh38,1000 Genomes 30x on GRCh38,1000 Genomes phase 3 release |
| **HG03801** | female | SAME1839606 | BEB | Bengali | SAS | South Asian Ancestry | BEB | 1000 Genomes 30x on GRCh38 |
| **HG04216** | female | SAME1839339 | ITU | Telugu | SAS | South Asian Ancestry | ITU | 1000 Genomes on GRCh38,1000 Genomes 30x on GRCh38,1000 Genomes phase 3 release |
| **HG03806** | male | SAME1839542 | BEB | Bengali | SAS | South Asian Ancestry | BEB | 1000 Genomes on GRCh38,1000 Genomes 30x on GRCh38 |
| **HG04177** | female | SAME1839992 | BEB | Bengali | SAS | South Asian Ancestry | BEB | 1000 Genomes on GRCh38,1000 Genomes 30x on GRCh38,1000 Genomes phase 3 release |
| **HG04184** | female | SAME1840049 | BEB | Bengali | SAS | South Asian Ancestry | BEB | 1000 Genomes 30x on GRCh38 |
| **HG03729** | male | SAME1839701 | ITU | Telugu | SAS | South Asian Ancestry | ITU | 1000 Genomes on GRCh38,1000 Genomes 30x on GRCh38,1000 Genomes phase 3 release |
| **HG04189** | female | SAME1839783 | BEB | Bengali | SAS | South Asian Ancestry | BEB | 1000 Genomes on GRCh38,1000 Genomes 30x on GRCh38,1000 Genomes phase 3 release |
| **HG03731** | female | SAME1839372 | ITU | Telugu | SAS | South Asian Ancestry | ITU | 1000 Genomes on GRCh38,1000 Genomes 30x on GRCh38,1000 Genomes phase 3 release |
| **HG04191** | male | SAME1839897 | BEB | Bengali | SAS | South Asian Ancestry | BEB | 1000 Genomes on GRCh38,1000 Genomes 30x on GRCh38 |
| **HG03736** | female | SAME1839389 | STU | Tamil | SAS | South Asian Ancestry | STU | 1000 Genomes on GRCh38,1000 Genomes 30x on GRCh38,1000 Genomes phase 3 release |
| **HG03965** | male | SAME1840063 | ITU | Telugu | SAS | South Asian Ancestry | ITU | 1000 Genomes on GRCh38,1000 Genomes 30x on GRCh38,1000 Genomes phase 3 release |
| **HG03743** | male | SAME1839261 | STU | Tamil | SAS | South Asian Ancestry | STU | 1000 Genomes on GRCh38,1000 Genomes 30x on GRCh38,1000 Genomes phase 3 release |
| **HG04094** | male | SAME1839953 | ITU | Telugu | SAS | South Asian Ancestry | ITU | 1000 Genomes on GRCh38,1000 Genomes 30x on GRCh38,1000 Genomes phase 3 release |
| **HG03977** | female | SAME1839746 | ITU | Telugu | SAS | South Asian Ancestry | ITU | 1000 Genomes on GRCh38,1000 Genomes 30x on GRCh38,1000 Genomes phase 3 release |
| **HG04099** | female | SAME1840065 | STU | Tamil | SAS | South Asian Ancestry | STU | 1000 Genomes on GRCh38,1000 Genomes 30x on GRCh38,1000 Genomes phase 3 release |
| **HG03989** | female | SAME1839532 | STU | Tamil | SAS | South Asian Ancestry | STU | 1000 Genomes on GRCh38,1000 Genomes 30x on GRCh38,1000 Genomes phase 3 release |
| **HG04107** | male | SAME1839660 | STU | Tamil | SAS | South Asian Ancestry | STU | 1000 Genomes on GRCh38,1000 Genomes 30x on GRCh38,1000 Genomes phase 3 release |
| **HG03991** | male | SAME1839640 | STU | Tamil | SAS | South Asian Ancestry | STU | 1000 Genomes on GRCh38,1000 Genomes 30x on GRCh38,1000 Genomes phase 3 release |
| **HG04038** | female | SAME1839499 | STU | Tamil | SAS | South Asian Ancestry | STU | 1000 Genomes on GRCh38,1000 Genomes 30x on GRCh38,1000 Genomes phase 3 release |
| **HG04133** | female | SAME1839294 | BEB | Bengali | SAS | South Asian Ancestry | BEB | 1000 Genomes 30x on GRCh38 |
| **HG04076** | female | SAME1840358 | ITU | Telugu | SAS | South Asian Ancestry | ITU | 1000 Genomes on GRCh38,1000 Genomes 30x on GRCh38,1000 Genomes phase 3 release |
| **HG04056** | male | SAME1839102 | ITU | Telugu | SAS | South Asian Ancestry | ITU | 1000 Genomes on GRCh38,1000 Genomes 30x on GRCh38,1000 Genomes phase 3 release |
| **HG04002** | male | SAME1839769 | ITU | Telugu | SAS | South Asian Ancestry | ITU | 1000 Genomes on GRCh38,1000 Genomes 30x on GRCh38,1000 Genomes phase 3 release |
| **HG04063** | female | SAME1839155 | ITU | Telugu | SAS | South Asian Ancestry | ITU | 1000 Genomes on GRCh38,1000 Genomes 30x on GRCh38,1000 Genomes phase 3 release |
| **HG04014** | female | SAME1839844 | ITU | Telugu | SAS | South Asian Ancestry | ITU | 1000 Genomes on GRCh38,1000 Genomes 30x on GRCh38,1000 Genomes phase 3 release |
| **HG04070** | female | SAME1840411 | ITU | Telugu | SAS | South Asian Ancestry | ITU | 1000 Genomes on GRCh38,1000 Genomes 30x on GRCh38,1000 Genomes phase 3 release |
| **HG04019** | male | SAME1839572 | ITU | Telugu | SAS | South Asian Ancestry | ITU | 1000 Genomes on GRCh38,1000 Genomes 30x on GRCh38,1000 Genomes phase 3 release |
| **HG04075** | female | SAME1840359 | STU | Tamil | SAS | South Asian Ancestry | STU | 1000 Genomes on GRCh38,1000 Genomes 30x on GRCh38,1000 Genomes phase 3 release |
| **HG04026** | female | SAME1839622 | ITU | Telugu | SAS | South Asian Ancestry | ITU | 1000 Genomes on GRCh38,1000 Genomes 30x on GRCh38,1000 Genomes phase 3 release |
| **HG03875** | male | SAME1839965 | ITU | Telugu | SAS | South Asian Ancestry | ITU | 1000 Genomes on GRCh38,1000 Genomes 30x on GRCh38,1000 Genomes phase 3 release |
| **HG04033** | male | SAME1839466 | STU | Tamil | SAS | South Asian Ancestry | STU | 1000 Genomes on GRCh38,1000 Genomes 30x on GRCh38,1000 Genomes phase 3 release |
| **HG03882** | female | SAME1839999 | ITU | Telugu | SAS | South Asian Ancestry | ITU | 1000 Genomes on GRCh38,1000 Genomes 30x on GRCh38,1000 Genomes phase 3 release |
| **NA12878** | female | SAME123392 | CEU | CEPH | EUR | European Ancestry | CEU | 1000 Genomes on GRCh38,1000 Genomes 30x on GRCh38,Human Genome Structural Variation Consortium, Phase 2,Illumina Platinum pedigree,1000 Genomes phase 3 |
| **HG03887** | male | SAME1840028 | STU | Tamil | SAS | South Asian Ancestry | STU | 1000 Genomes on GRCh38,1000 Genomes 30x on GRCh38,1000 Genomes phase 3 release |
| **NA12892** | female | SAME123181 | CEU | CEPH | EUR | European Ancestry | CEU | 1000 Genomes on GRCh38,1000 Genomes 30x on GRCh38,Illumina Platinum pedigree,1000 Genomes phase 3 release,1000 Genomes phase 1 release |
| **HG03894** | female | SAME1839885 | STU | Tamil | SAS | South Asian Ancestry | STU | 1000 Genomes on GRCh38,1000 Genomes 30x on GRCh38,1000 Genomes phase 3 release |
| **NA18484** | female | SAME124101 | YRI | Yoruba | AFR | African Ancestry | YRI | 1000 Genomes 30x on GRCh38,1000 Genomes phase 3 release |
| **HG03899** | male | SAME1839814 | STU | Tamil | SAS | South Asian Ancestry | STU | 1000 Genomes on GRCh38,1000 Genomes 30x on GRCh38,1000 Genomes phase 3 release |
| **NA18489** | female | SAME124102 | YRI | Yoruba | AFR | African Ancestry | YRI | 1000 Genomes on GRCh38,1000 Genomes 30x on GRCh38,1000 Genomes phase 3 release,1000 Genomes phase 1 release,Geuvadis |
| **HG03969** | male | SAME1840082 | ITU | Telugu | SAS | South Asian Ancestry | ITU | 1000 Genomes on GRCh38,1000 Genomes 30x on GRCh38,1000 Genomes phase 3 release |
| **NA18504** | male | SAME125061 | YRI | Yoruba | AFR | African Ancestry | YRI | 1000 Genomes on GRCh38,1000 Genomes 30x on GRCh38,1000 Genomes phase 3 release,1000 Genomes phase 1 release |
| **HG03971** | male | SAME1839718 | ITU | Telugu | SAS | South Asian Ancestry | ITU | 1000 Genomes on GRCh38,1000 Genomes 30x on GRCh38,1000 Genomes phase 3 release |
| **HG03976** | male | SAME1839739 | ITU | Telugu | SAS | South Asian Ancestry | ITU | 1000 Genomes on GRCh38,1000 Genomes 30x on GRCh38,1000 Genomes phase 3 release |
| **NA18511** | female | SAME124885 | YRI | Yoruba | AFR | African Ancestry | YRI | 1000 Genomes on GRCh38,1000 Genomes 30x on GRCh38,1000 Genomes phase 3 release,1000 Genomes phase 1 release,Geuvadis |
| **NA18516** | male | SAME124884 | YRI | Yoruba | AFR | African Ancestry | YRI | 1000 Genomes on GRCh38,1000 Genomes 30x on GRCh38,1000 Genomes phase 3 release,1000 Genomes phase 1 release |
| **HG03990** | male | SAME1839637 | STU | Tamil | SAS | South Asian Ancestry | STU | 1000 Genomes on GRCh38,1000 Genomes 30x on GRCh38,1000 Genomes phase 3 release |
| **NA18523** | female | SAME124670 | YRI | Yoruba | AFR | African Ancestry | YRI | 1000 Genomes on GRCh38,1000 Genomes 30x on GRCh38,1000 Genomes phase 3 release,1000 Genomes phase 1 release |
| **HG03995** | female | SAME1839395 | STU | Tamil | SAS | South Asian Ancestry | STU | 1000 Genomes on GRCh38,1000 Genomes 30x on GRCh38,1000 Genomes phase 3 release |
| **NA18528** | female | SAME124661 | CHB | Han Chinese | EAS | East Asian Ancestry | CHB | 1000 Genomes on GRCh38,1000 Genomes 30x on GRCh38,1000 Genomes phase 3 release,1000 Genomes phase 1 release |
| **HG03902** | male | SAME1839484 | BEB | Bengali | SAS | South Asian Ancestry | BEB | 1000 Genomes on GRCh38,1000 Genomes 30x on GRCh38,1000 Genomes phase 3 release |
| **NA18530** | male | SAME124484 | CHB | Han Chinese | EAS | East Asian Ancestry | CHB | 1000 Genomes on GRCh38,1000 Genomes 30x on GRCh38,1000 Genomes phase 3 release,1000 Genomes phase 1 release |
| **HG03907** | female | SAME1839517 | BEB | Bengali | SAS | South Asian Ancestry | BEB | 1000 Genomes on GRCh38,1000 Genomes 30x on GRCh38,1000 Genomes phase 3 release |
| **NA18535** | female | SAME124487 | CHB | Han Chinese | EAS | East Asian Ancestry | CHB | 1000 Genomes on GRCh38,1000 Genomes 30x on GRCh38,1000 Genomes phase 3 release,1000 Genomes phase 1 release |
| **HG03914** | male | SAME1839224 | BEB | Bengali | SAS | South Asian Ancestry | BEB | 1000 Genomes on GRCh38,1000 Genomes 30x on GRCh38,1000 Genomes phase 3 release |
| **NA18542** | female | SAME124267 | CHB | Han Chinese | EAS | East Asian Ancestry | CHB | 1000 Genomes on GRCh38,1000 Genomes 30x on GRCh38,1000 Genomes phase 3 release,1000 Genomes phase 1 release,90 Han Chinese high coverage genomes |
| **HG03919** | female | SAME1839329 | BEB | Bengali | SAS | South Asian Ancestry | BEB | 1000 Genomes on GRCh38,1000 Genomes 30x on GRCh38,1000 Genomes phase 3 release |
| **NA18547** | female | SAME124264 | CHB | Han Chinese | EAS | East Asian Ancestry | CHB | 1000 Genomes on GRCh38,1000 Genomes 30x on GRCh38,1000 Genomes phase 3 release,1000 Genomes phase 1 release,90 Han Chinese high coverage genomes |
| **HG03926** | male | SAME1839033 | BEB | Bengali | SAS | South Asian Ancestry | BEB | 1000 Genomes on GRCh38,1000 Genomes 30x on GRCh38,1000 Genomes phase 3 release |
| **NA18559** | male | SAME124093 | CHB | Han Chinese | EAS | East Asian Ancestry | CHB | 1000 Genomes on GRCh38,1000 Genomes 30x on GRCh38,1000 Genomes phase 3 release,1000 Genomes phase 1 release |
| **HG04164** | male | SAME1840102 | BEB | Bengali | SAS | South Asian Ancestry | BEB | 1000 Genomes on GRCh38,1000 Genomes 30x on GRCh38,1000 Genomes phase 3 release |
| **NA18561** | male | SAME123926 | CHB | Han Chinese | EAS | East Asian Ancestry | CHB | 1000 Genomes on GRCh38,1000 Genomes 30x on GRCh38,1000 Genomes phase 3 release,1000 Genomes phase 1 release,90 Han Chinese high coverage genomes |
| **HG04171** | female | SAME1839968 | BEB | Bengali | SAS | South Asian Ancestry | BEB | 1000 Genomes on GRCh38,1000 Genomes 30x on GRCh38,1000 Genomes phase 3 release |
| **NA18566** | female | SAME123923 | CHB | Han Chinese | EAS | East Asian Ancestry | CHB | 1000 Genomes on GRCh38,1000 Genomes 30x on GRCh38,1000 Genomes phase 3 release,1000 Genomes phase 1 release,90 Han Chinese high coverage genomes |
| **HG04176** | male | SAME1839986 | BEB | Bengali | SAS | South Asian Ancestry | BEB | 1000 Genomes on GRCh38,1000 Genomes 30x on GRCh38,1000 Genomes phase 3 release |
| **NA18573** | female | SAME123738 | CHB | Han Chinese | EAS | East Asian Ancestry | CHB | 1000 Genomes on GRCh38,1000 Genomes 30x on GRCh38,1000 Genomes phase 3 release,1000 Genomes phase 1 release,90 Han Chinese high coverage genomes |
| **HG04183** | female | SAME1840044 | BEB | Bengali | SAS | South Asian Ancestry | BEB | 1000 Genomes on GRCh38,1000 Genomes 30x on GRCh38,1000 Genomes phase 3 release |
| **NA18592** | female | SAME123721 | CHB | Han Chinese | EAS | East Asian Ancestry | CHB | 1000 Genomes on GRCh38,1000 Genomes 30x on GRCh38,1000 Genomes phase 3 release,1000 Genomes phase 1 release,90 Han Chinese high coverage genomes |
| **HG04188** | male | SAME1839771 | BEB | Bengali | SAS | South Asian Ancestry | BEB | 1000 Genomes on GRCh38,1000 Genomes 30x on GRCh38,1000 Genomes phase 3 release |
| **NA18597** | female | SAME123232 | CHB | Han Chinese | EAS | East Asian Ancestry | CHB | 1000 Genomes on GRCh38,1000 Genomes 30x on GRCh38,1000 Genomes phase 3 release,1000 Genomes phase 1 release |
| **HG04001** | female | SAME1839752 | ITU | Telugu | SAS | South Asian Ancestry | ITU | 1000 Genomes on GRCh38,1000 Genomes 30x on GRCh38,1000 Genomes phase 3 release |
| **NA18605** | male | SAME124877 | CHB | Han Chinese | EAS | East Asian Ancestry | CHB | 1000 Genomes on GRCh38,1000 Genomes 30x on GRCh38,1000 Genomes phase 3 release,1000 Genomes phase 1 release,90 Han Chinese high coverage genomes |
| **HG04006** | male | SAME1839787 | STU | Tamil | SAS | South Asian Ancestry | STU | 1000 Genomes on GRCh38,1000 Genomes 30x on GRCh38,1000 Genomes phase 3 release |
| **NA18612** | male | SAME125036 | CHB | Han Chinese | EAS | East Asian Ancestry | CHB | 1000 Genomes on GRCh38,1000 Genomes 30x on GRCh38,1000 Genomes phase 3 release,1000 Genomes phase 1 release,90 Han Chinese high coverage genomes |

| | | | | | | | | |
|---|---|---|---|---|---|---|---|---|
| **HG04018** | female | SAME1839565 | ITU | Telugu | SAS | South Asian Ancestry | ITU | 1000 Genomes on GRCh38,1000 Genomes 30x on GRCh38,1000 Genomes phase 3 release |
| **NA18617** | female | SAME125033 | CHB | Han Chinese | EAS | East Asian Ancestry | CHB | 1000 Genomes on GRCh38,1000 Genomes 30x on GRCh38,1000 Genomes phase 3 release,1000 Genomes phase 1 release |
| **HG04020** | male | SAME1839685 | ITU | Telugu | SAS | South Asian Ancestry | ITU | 1000 Genomes on GRCh38,1000 Genomes 30x on GRCh38,1000 Genomes phase 3 release |
| **NA18624** | male | SAME124506 | CHB | Han Chinese | EAS | East Asian Ancestry | CHB | 1000 Genomes on GRCh38,1000 Genomes 30x on GRCh38,1000 Genomes phase 3 release,1000 Genomes phase 1 release,90 Han Chinese high coverage genomes |
| **HG04025** | female | SAME1839614 | ITU | Telugu | SAS | South Asian Ancestry | ITU | 1000 Genomes on GRCh38,1000 Genomes 30x on GRCh38,1000 Genomes phase 3 release |
| **NA18629** | male | SAME124501 | CHB | Han Chinese | EAS | East Asian Ancestry | CHB | 1000 Genomes on GRCh38,1000 Genomes 30x on GRCh38,1000 Genomes phase 3 release |
| **HG04140** | male | SAME1839173 | BEB | Bengali | SAS | South Asian Ancestry | BEB | 1000 Genomes on GRCh38,1000 Genomes 30x on GRCh38,1000 Genomes phase 3 release |
| **NA18631** | female | SAME124056 | CHB | Han Chinese | EAS | East Asian Ancestry | CHB | 1000 Genomes on GRCh38,1000 Genomes 30x on GRCh38,1000 Genomes phase 3 release,1000 Genomes phase 1 release |
| **HG04152** | male | SAME1840300 | BEB | Bengali | SAS | South Asian Ancestry | BEB | 1000 Genomes on GRCh38,1000 Genomes 30x on GRCh38,1000 Genomes phase 3 release |
| **NA18636** | male | SAME124320 | CHB | Han Chinese | EAS | East Asian Ancestry | CHB | 1000 Genomes on GRCh38,1000 Genomes 30x on GRCh38,1000 Genomes phase 3 release,1000 Genomes phase 1 release,90 Han Chinese high coverage genomes |
| **HG04157** | male | SAME1840395 | BEB | Bengali | SAS | South Asian Ancestry | BEB | 1000 Genomes 30x on GRCh38 |
| **NA18643** | male | SAME122981 | CHB | Han Chinese | EAS | East Asian Ancestry | CHB | 1000 Genomes on GRCh38,1000 Genomes 30x on GRCh38,1000 Genomes phase 3 release,1000 Genomes phase 1 release |
| **HG03940** | female | SAME1840346 | BEB | Bengali | SAS | South Asian Ancestry | BEB | 1000 Genomes on GRCh38,1000 Genomes 30x on GRCh38,1000 Genomes phase 3 release |
| **NA18648** | male | SAME122974 | CHB | Han Chinese | EAS | East Asian Ancestry | CHB | 1000 Genomes on GRCh38,1000 Genomes 30x on GRCh38,1000 Genomes phase 3 release |
| **HG03945** | female | SAME1840369 | STU | Tamil | SAS | South Asian Ancestry | STU | 1000 Genomes on GRCh38,1000 Genomes 30x on GRCh38,1000 Genomes phase 3 release |
| **NA18908** | male | SAME123560 | YRI | Yoruba | AFR | African Ancestry | YRI | 1000 Genomes on GRCh38,1000 Genomes 30x on GRCh38,1000 Genomes phase 3 release,1000 Genomes phase 1 release,Geuvadis |
| **NA18534** | male | SAME124488 | CHB | Han Chinese | EAS | East Asian Ancestry | CHB | 1000 Genomes on GRCh38,1000 Genomes 30x on GRCh38,Human Genome Structural Variation Consortium, Phase 2,1000 Genomes phase 3 release,1000 Genomes ph |
| **NA18910** | male | SAME123402 | YRI | Yoruba | AFR | African Ancestry | YRI | 1000 Genomes on GRCh38,1000 Genomes 30x on GRCh38,1000 Genomes phase 3 release,1000 Genomes phase 1 release,Geuvadis |
| **NA18539** | female | SAME124475 | CHB | Han Chinese | EAS | East Asian Ancestry | CHB | 1000 Genomes on GRCh38,1000 Genomes 30x on GRCh38,1000 Genomes phase 3 release,1000 Genomes phase 1 release |
| **NA18915** | male | SAME123405 | YRI | Yoruba | AFR | African Ancestry | YRI | 1000 Genomes on GRCh38,1000 Genomes 30x on GRCh38,1000 Genomes phase 3 release |
| **NA18541** | female | SAME124270 | CHB | Han Chinese | EAS | East Asian Ancestry | CHB | 1000 Genomes on GRCh38,1000 Genomes 30x on GRCh38,1000 Genomes phase 3 release,1000 Genomes phase 1 release |
| **NA18934** | male | SAME123036 | YRI | Yoruba | AFR | African Ancestry | YRI | 1000 Genomes on GRCh38,1000 Genomes 30x on GRCh38,1000 Genomes phase 3 release,1000 Genomes phase 1 release,Geuvadis |
| **NA18546** | male | SAME124263 | CHB | Han Chinese | EAS | East Asian Ancestry | CHB | 1000 Genomes on GRCh38,1000 Genomes 30x on GRCh38,1000 Genomes phase 3 release,1000 Genomes phase 1 release |
| **NA18939** | female | SAME123041 | JPT | Japanese | EAS | East Asian Ancestry | JPT | 1000 Genomes on GRCh38,1000 Genomes 30x on GRCh38,Human Genome Structural Variation Consortium, Phase 2,1000 Genomes phase 3 release,1000 Genomes ph |
| **NA18553** | female | SAME124088 | CHB | Han Chinese | EAS | East Asian Ancestry | CHB | 1000 Genomes on GRCh38,1000 Genomes 30x on GRCh38,1000 Genomes phase 3 release,1000 Genomes phase 1 release |
| **NA18941** | female | SAME124311 | JPT | Japanese | EAS | East Asian Ancestry | JPT | 1000 Genomes on GRCh38,1000 Genomes 30x on GRCh38,1000 Genomes phase 3 release,1000 Genomes phase 1 release |
| **NA18558** | male | SAME124094 | CHB | Han Chinese | EAS | East Asian Ancestry | CHB | 1000 Genomes on GRCh38,1000 Genomes 30x on GRCh38,1000 Genomes phase 3 release,1000 Genomes phase 1 release,90 Han Chinese high coverage genomes |
| **NA18946** | female | SAME124312 | JPT | Japanese | EAS | East Asian Ancestry | JPT | 1000 Genomes on GRCh38,1000 Genomes 30x on GRCh38,1000 Genomes phase 3 release,1000 Genomes phase 1 release |
| **NA18560** | female | SAME123925 | CHB | Han Chinese | EAS | East Asian Ancestry | CHB | 1000 Genomes on GRCh38,1000 Genomes 30x on GRCh38,1000 Genomes phase 3 release,1000 Genomes phase 1 release |
| **NA18953** | male | SAME124113 | JPT | Japanese | EAS | East Asian Ancestry | JPT | 1000 Genomes on GRCh38,1000 Genomes 30x on GRCh38,1000 Genomes phase 3 release,1000 Genomes phase 1 release |
| **NA18565** | female | SAME123922 | CHB | Han Chinese | EAS | East Asian Ancestry | CHB | 1000 Genomes on GRCh38,1000 Genomes 30x on GRCh38,1000 Genomes phase 3 release,1000 Genomes phase 1 release |
| **NA18960** | male | SAME123956 | JPT | Japanese | EAS | East Asian Ancestry | JPT | 1000 Genomes on GRCh38,1000 Genomes 30x on GRCh38,1000 Genomes phase 3 release,1000 Genomes phase 1 release |
| **NA18572** | male | SAME123739 | CHB | Han Chinese | EAS | East Asian Ancestry | CHB | 1000 Genomes on GRCh38,1000 Genomes 30x on GRCh38,1000 Genomes phase 3 release,1000 Genomes phase 1 release,90 Han Chinese high coverage genomes |
| **NA18965** | male | SAME123961 | JPT | Japanese | EAS | East Asian Ancestry | JPT | 1000 Genomes on GRCh38,1000 Genomes 30x on GRCh38,1000 Genomes phase 3 release,1000 Genomes phase 1 release |
| **NA18577** | female | SAME123734 | CHB | Han Chinese | EAS | East Asian Ancestry | CHB | 1000 Genomes on GRCh38,1000 Genomes 30x on GRCh38,1000 Genomes phase 3 release,1000 Genomes phase 1 release,90 Han Chinese high coverage genomes |
| **NA18591** | female | SAME123719 | CHB | Han Chinese | EAS | East Asian Ancestry | CHB | 1000 Genomes on GRCh38,1000 Genomes 30x on GRCh38,1000 Genomes phase 3 release |
| **NA10860** | male | SAME125131 | CEU | CEPH | EUR | European Ancestry | CEU | 1000 Genomes 30x on GRCh38,1000 Genomes phase 3 release |
| **NA18596** | female | SAME123718 | CHB | Han Chinese | EAS | East Asian Ancestry | CHB | 1000 Genomes on GRCh38,1000 Genomes 30x on GRCh38,1000 Genomes phase 3 release,1000 Genomes phase 1 release |
| **NA10865** | male | SAME125128 | CEU | CEPH | EUR | European Ancestry | CEU | 1000 Genomes 30x on GRCh38,1000 Genomes phase 3 release |
| **NA18609** | male | SAME124867 | CHB | Han Chinese | EAS | East Asian Ancestry | CHB | 1000 Genomes on GRCh38,1000 Genomes 30x on GRCh38,1000 Genomes phase 3 release,1000 Genomes phase 1 release,90 Han Chinese high coverage genomes |
| **NA11832** | female | SAME124554 | CEU | CEPH | EUR | European Ancestry | CEU | 1000 Genomes on GRCh38,1000 Genomes 30x on GRCh38,1000 Genomes phase 3 release,1000 Genomes phase 1 release,Geuvadis |
| **NA18611** | male | SAME123225 | CHB | Han Chinese | EAS | East Asian Ancestry | CHB | 1000 Genomes on GRCh38,1000 Genomes 30x on GRCh38,1000 Genomes phase 3 release,1000 Genomes phase 1 release,90 Han Chinese high coverage genomes |
| **NA11882** | female | SAME123210 | CEU | CEPH | EUR | European Ancestry | CEU | 1000 Genomes 30x on GRCh38,1000 Genomes phase 3 release |
| **NA18616** | female | SAME125032 | CHB | Han Chinese | EAS | East Asian Ancestry | CHB | 1000 Genomes on GRCh38,1000 Genomes 30x on GRCh38,1000 Genomes phase 3 release,1000 Genomes phase 1 release |
| **NA11894** | female | SAME123043 | CEU | CEPH | EUR | European Ancestry | CEU | 1000 Genomes on GRCh38,1000 Genomes 30x on GRCh38,1000 Genomes phase 3 release,1000 Genomes phase 1 release,Geuvadis |
| **NA18623** | male | SAME124507 | CHB | Han Chinese | EAS | East Asian Ancestry | CHB | 1000 Genomes on GRCh38,1000 Genomes 30x on GRCh38,1000 Genomes phase 3 release,1000 Genomes phase 1 release,90 Han Chinese high coverage genomes |
| **NA11919** | male | SAME122834 | CEU | CEPH | EUR | European Ancestry | CEU | 1000 Genomes on GRCh38,1000 Genomes 30x on GRCh38,1000 Genomes phase 3 release,1000 Genomes phase 1 release |
| **NA18628** | female | SAME124502 | CHB | Han Chinese | EAS | East Asian Ancestry | CHB | 1000 Genomes on GRCh38,1000 Genomes 30x on GRCh38,1000 Genomes phase 3 release,1000 Genomes phase 1 release |
| **NA11933** | female | SAME125098 | CEU | CEPH | EUR | European Ancestry | CEU | 1000 Genomes on GRCh38,1000 Genomes 30x on GRCh38,1000 Genomes phase 3 release,1000 Genomes phase 1 release |
| **NA18630** | female | SAME124895 | CHB | Han Chinese | EAS | East Asian Ancestry | CHB | 1000 Genomes on GRCh38,1000 Genomes 30x on GRCh38,1000 Genomes phase 3 release,1000 Genomes phase 1 release |
| **NA11995** | female | SAME123968 | CEU | CEPH | EUR | European Ancestry | CEU | 1000 Genomes on GRCh38,1000 Genomes 30x on GRCh38,1000 Genomes phase 3 release,1000 Genomes phase 1 release,Geuvadis |
| **NA18635** | male | SAME123661 | CHB | Han Chinese | EAS | East Asian Ancestry | CHB | 1000 Genomes on GRCh38,1000 Genomes 30x on GRCh38,1000 Genomes phase 3 release,1000 Genomes phase 1 release,90 Han Chinese high coverage genomes |
| **NA12006** | female | SAME125145 | CEU | CEPH | EUR | European Ancestry | CEU | 1000 Genomes on GRCh38,1000 Genomes 30x on GRCh38,1000 Genomes phase 3 release,1000 Genomes phase 1 release,Geuvadis |
| **NA18642** | female | SAME122982 | CHB | Han Chinese | EAS | East Asian Ancestry | CHB | 1000 Genomes on GRCh38,1000 Genomes 30x on GRCh38,1000 Genomes phase 3 release,1000 Genomes phase 1 release |
| **NA12044** | female | SAME123212 | CEU | CEPH | EUR | European Ancestry | CEU | 1000 Genomes on GRCh38,1000 Genomes 30x on GRCh38,1000 Genomes phase 3 release,1000 Genomes phase 1 release,Geuvadis |
| **NA18647** | male | SAME122977 | CHB | Han Chinese | EAS | East Asian Ancestry | CHB | 1000 Genomes on GRCh38,1000 Genomes 30x on GRCh38,1000 Genomes phase 3 release,1000 Genomes phase 1 release |
| **NA12056** | male | SAME123414 | CEU | CEPH | EUR | European Ancestry | CEU | 1000 Genomes 30x on GRCh38,1000 Genomes phase 3 release |
| **NA10852** | female | SAME125320 | CEU | CEPH | EUR | European Ancestry | CEU | 1000 Genomes 30x on GRCh38,1000 Genomes phase 3 release |
| **NA12145** | female | SAME124157 | CEU | CEPH | EUR | European Ancestry | CEU | 1000 Genomes 30x on GRCh38,1000 Genomes phase 3 release |
| **NA10857** | male | SAME125332 | CEU | CEPH | EUR | European Ancestry | CEU | 1000 Genomes 30x on GRCh38,1000 Genomes phase 3 release |
| **NA12234** | female | SAME124003 | CEU | CEPH | EUR | European Ancestry | CEU | 1000 Genomes on GRCh38,1000 Genomes 30x on GRCh38,1000 Genomes phase 3 release,1000 Genomes phase 1 release,Geuvadis |
| **NA10864** | female | SAME125129 | CEU | CEPH | EUR | European Ancestry | CEU | 1000 Genomes 30x on GRCh38,1000 Genomes phase 3 release |
| **NA12239** | female | SAME123999 | CEU | CEPH | EUR | European Ancestry | CEU | 1000 Genomes 30x on GRCh38,1000 Genomes phase 3 release |
| **NA11829** | male | SAME124761 | CEU | CEPH | EUR | European Ancestry | CEU | 1000 Genomes on GRCh38,1000 Genomes 30x on GRCh38,1000 Genomes phase 3 release,1000 Genomes phase 1 release,Geuvadis |
| **NA12272** | male | SAME123231 | CEU | CEPH | EUR | European Ancestry | CEU | 1000 Genomes on GRCh38,1000 Genomes 30x on GRCh38,1000 Genomes phase 3 release,1000 Genomes phase 1 release,Geuvadis |
| **NA11831** | male | SAME124552 | CEU | CEPH | EUR | European Ancestry | CEU | 1000 Genomes on GRCh38,1000 Genomes 30x on GRCh38,1000 Genomes phase 3 release,1000 Genomes phase 1 release,Geuvadis |
| **NA12335** | male | SAME124097 | CEU | CEPH | EUR | European Ancestry | CEU | 1000 Genomes 30x on GRCh38,1000 Genomes phase 3 release |
| **NA11843** | male | SAME125140 | CEU | CEPH | EUR | European Ancestry | CEU | 1000 Genomes on GRCh38,1000 Genomes 30x on GRCh38,1000 Genomes phase 3 release,1000 Genomes phase 1 release,Geuvadis |
| **NA12342** | male | SAME125155 | CEU | CEPH | EUR | European Ancestry | CEU | 1000 Genomes on GRCh38,1000 Genomes 30x on GRCh38,1000 Genomes phase 3 release,1000 Genomes phase 1 release,Geuvadis |
| **NA11881** | male | SAME123209 | CEU | CEPH | EUR | European Ancestry | CEU | 1000 Genomes on GRCh38,1000 Genomes 30x on GRCh38,1000 Genomes phase 3 release,1000 Genomes phase 1 release,Geuvadis |
| **NA12347** | male | SAME125316 | CEU | CEPH | EUR | European Ancestry | CEU | 1000 Genomes on GRCh38,1000 Genomes 30x on GRCh38,1000 Genomes phase 3 release,1000 Genomes phase 1 release,Geuvadis |
| **NA11893** | male | SAME123044 | CEU | CEPH | EUR | European Ancestry | CEU | 1000 Genomes on GRCh38,1000 Genomes 30x on GRCh38,1000 Genomes phase 3 release,1000 Genomes phase 1 release,Geuvadis |
| **NA12400** | female | SAME123881 | CEU | CEPH | EUR | European Ancestry | CEU | 1000 Genomes on GRCh38,1000 Genomes 30x on GRCh38,1000 Genomes phase 3 release,1000 Genomes phase 1 release,Geuvadis |
| **NA11918** | female | SAME123135 | CEU | CEPH | EUR | European Ancestry | CEU | 1000 Genomes on GRCh38,1000 Genomes 30x on GRCh38,1000 Genomes phase 3 release,1000 Genomes phase 1 release,Geuvadis |
| **NA11920** | female | SAME124901 | CEU | CEPH | EUR | European Ancestry | CEU | 1000 Genomes on GRCh38,1000 Genomes 30x on GRCh38,1000 Genomes phase 3 release,1000 Genomes phase 1 release,Geuvadis |
| **NA12739** | male | SAME123529 | CEU | CEPH | EUR | European Ancestry | CEU | 1000 Genomes 30x on GRCh38,1000 Genomes phase 3 release |
| **NA11932** | male | SAME125097 | CEU | CEPH | EUR | European Ancestry | CEU | 1000 Genomes on GRCh38,1000 Genomes 30x on GRCh38,1000 Genomes phase 3 release,1000 Genomes phase 1 release |
| **NA12753** | female | SAME124647 | CEU | CEPH | EUR | European Ancestry | CEU | 1000 Genomes 30x on GRCh38,1000 Genomes phase 3 release |
| **NA11994** | male | SAME123967 | CEU | CEPH | EUR | European Ancestry | CEU | 1000 Genomes on GRCh38,1000 Genomes 30x on GRCh38,1000 Genomes phase 3 release,1000 Genomes phase 1 release,Geuvadis |
| **NA12760** | male | SAME124465 | CEU | CEPH | EUR | European Ancestry | CEU | 1000 Genomes on GRCh38,1000 Genomes 30x on GRCh38,1000 Genomes phase 3 release,1000 Genomes phase 1 release,Geuvadis |
| **NA12005** | male | SAME125143 | CEU | CEPH | EUR | European Ancestry | CEU | 1000 Genomes on GRCh38,1000 Genomes 30x on GRCh38,1000 Genomes phase 3 release,1000 Genomes phase 1 release,Geuvadis |
| **NA12777** | male | SAME125252 | CEU | CEPH | EUR | European Ancestry | CEU | 1000 Genomes on GRCh38,1000 Genomes 30x on GRCh38,1000 Genomes phase 3 release,1000 Genomes phase 1 release,Geuvadis |
| **NA12877** | male | SAME123391 | CEU | CEPH | EUR | European Ancestry | CEU | 1000 Genomes 30x on GRCh38,Illumina Platinum pedigree,1000 Genomes phase 3 release |
| **NA12828** | female | SAME125076 | CEU | CEPH | EUR | European Ancestry | CEU | 1000 Genomes on GRCh38,1000 Genomes 30x on GRCh38,1000 Genomes phase 3 release,1000 Genomes phase 1 release |
| **NA12889** | male | SAME123951 | CEU | CEPH | EUR | European Ancestry | CEU | 1000 Genomes on GRCh38,1000 Genomes 30x on GRCh38,Illumina Platinum pedigree,1000 Genomes phase 3 release,1000 Genomes phase 1 release,Geuvadis |
| **NA12830** | female | SAME125289 | CEU | CEPH | EUR | European Ancestry | CEU | 1000 Genomes on GRCh38,1000 Genomes 30x on GRCh38,1000 Genomes phase 3 release,1000 Genomes phase 1 release,Geuvadis |
| **NA12891** | male | SAME124117 | CEU | CEPH | EUR | European Ancestry | CEU | 1000 Genomes on GRCh38,1000 Genomes 30x on GRCh38,Illumina Platinum pedigree,1000 Genomes phase 3 release,1000 Genomes phase 1 release |
| **NA12842** | male | SAME123542 | CEU | CEPH | EUR | European Ancestry | CEU | 1000 Genomes on GRCh38,1000 Genomes 30x on GRCh38,1000 Genomes phase 3 release,1000 Genomes phase 1 release,Geuvadis |
| **NA18488** | female | SAME124103 | YRI | Yoruba | AFR | African Ancestry | YRI | 1000 Genomes on GRCh38,1000 Genomes 30x on GRCh38,1000 Genomes phase 3 release,1000 Genomes phase 1 release,Geuvadis |
| **NA12873** | female | SAME123388 | CEU | CEPH | EUR | European Ancestry | CEU | 1000 Genomes on GRCh38,1000 Genomes 30x on GRCh38,1000 Genomes phase 3 release,1000 Genomes phase 1 release,Geuvadis |
| **NA18503** | male | SAME125056 | YRI | Yoruba | AFR | African Ancestry | YRI | 1000 Genomes 30x on GRCh38,1000 Genomes phase 3 release |
| **NA19026** | male | SAME124849 | LWK | Luhya | AFR | African Ancestry | LWK | 1000 Genomes on GRCh38,1000 Genomes 30x on GRCh38,1000 Genomes phase 3 release |
| **NA18508** | female | SAME124318 | YRI | Yoruba | AFR | African Ancestry | YRI | 1000 Genomes on GRCh38,1000 Genomes 30x on GRCh38,1000 Genomes phase 3 release,1000 Genomes phase 1 release,Geuvadis |
| **NA19038** | female | SAME124668 | LWK | Luhya | AFR | African Ancestry | LWK | 1000 Genomes on GRCh38,1000 Genomes 30x on GRCh38,1000 Genomes phase 3 release,1000 Genomes phase 1 release |
| **NA18510** | male | SAME124886 | YRI | Yoruba | AFR | African Ancestry | YRI | 1000 Genomes on GRCh38,1000 Genomes 30x on GRCh38,1000 Genomes phase 3 release,1000 Genomes phase 1 release,Geuvadis |
| **NA19057** | female | SAME125025 | JPT | Japanese | EAS | East Asian Ancestry | JPT | 1000 Genomes on GRCh38,1000 Genomes 30x on GRCh38,1000 Genomes phase 3 release,1000 Genomes phase 1 release |
| **NA18515** | male | SAME124881 | YRI | Yoruba | AFR | African Ancestry | YRI | 1000 Genomes 30x on GRCh38,1000 Genomes phase 3 release |
| **NA19064** | female | SAME123010 | JPT | Japanese | EAS | East Asian Ancestry | JPT | 1000 Genomes on GRCh38,1000 Genomes 30x on GRCh38,1000 Genomes phase 3 release,1000 Genomes phase 1 release |
| **NA18522** | male | SAME124669 | YRI | Yoruba | AFR | African Ancestry | YRI | 1000 Genomes on GRCh38,1000 Genomes 30x on GRCh38,1000 Genomes phase 3 release,1000 Genomes phase 1 release |
| **NA19076** | male | SAME122819 | JPT | Japanese | EAS | East Asian Ancestry | JPT | 1000 Genomes on GRCh38,1000 Genomes 30x on GRCh38,1000 Genomes phase 3 release,1000 Genomes phase 1 release |
| **NA19083** | male | SAME123371 | JPT | Japanese | EAS | East Asian Ancestry | JPT | 1000 Genomes on GRCh38,1000 Genomes 30x on GRCh38,1000 Genomes phase 3 release,1000 Genomes phase 1 release |
| **NA12707** | male | SAME124080 | CEU | CEPH | EUR | European Ancestry | CEU | 1000 Genomes 30x on GRCh38,1000 Genomes phase 3 release |
| **NA19088** | male | SAME123360 | JPT | Japanese | EAS | East Asian Ancestry | JPT | 1000 Genomes on GRCh38,1000 Genomes 30x on GRCh38,1000 Genomes phase 3 release,1000 Genomes phase 1 release |
| **NA12740** | female | SAME124822 | CEU | CEPH | EUR | European Ancestry | CEU | 1000 Genomes 30x on GRCh38,1000 Genomes phase 3 release |
| **NA19090** | female | SAME123175 | JPT | Japanese | EAS | East Asian Ancestry | JPT | 1000 Genomes on GRCh38,1000 Genomes 30x on GRCh38,1000 Genomes phase 3 release |
| **NA12752** | male | SAME123518 | CEU | CEPH | EUR | European Ancestry | CEU | 1000 Genomes 30x on GRCh38,1000 Genomes phase 3 release |
| **NA19095** | female | SAME123170 | YRI | Yoruba | AFR | African Ancestry | YRI | 1000 Genomes on GRCh38,1000 Genomes 30x on GRCh38,1000 Genomes phase 3 release,1000 Genomes phase 1 release,Geuvadis |
| **NA12776** | female | SAME124256 | CEU | CEPH | EUR | European Ancestry | CEU | 1000 Genomes on GRCh38,1000 Genomes 30x on GRCh38,1000 Genomes phase 3 release,1000 Genomes phase 1 release,Geuvadis |
| **NA18749** | male | SAME123910 | CHB | Han Chinese | EAS | East Asian Ancestry | CHB | 1000 Genomes on GRCh38,1000 Genomes 30x on GRCh38,1000 Genomes phase 3 release,1000 Genomes phase 1 release |
| **NA12815** | female | SAME123022 | CEU | CEPH | EUR | European Ancestry | CEU | 1000 Genomes on GRCh38,1000 Genomes 30x on GRCh38,1000 Genomes phase 3 release,1000 Genomes phase 1 release,Geuvadis |
| **NA12827** | male | SAME125077 | CEU | CEPH | EUR | European Ancestry | CEU | 1000 Genomes on GRCh38,1000 Genomes 30x on GRCh38,1000 Genomes phase 3 release,1000 Genomes phase 1 release,Geuvadis |

| | | | | | | | | |
|---|---|---|---|---|---|---|---|---|
| **NA18852** | female | SAME124732 | YRI | Yoruba | AFR | African Ancestry | YRI | 1000 Genomes 30x on GRCh38,1000 Genomes phase 3 release |
| **NA12865** | female | SAME123190 | CEU | CEPH | EUR | European Ancestry | CEU | 1000 Genomes 30x on GRCh38,1000 Genomes phase 3 release |
| **NA18857** | male | SAME124737 | YRI | Yoruba | AFR | African Ancestry | YRI | 1000 Genomes 30x on GRCh38,1000 Genomes phase 3 release |
| **NA12872** | male | SAME123387 | CEU | CEPH | EUR | European Ancestry | CEU | 1000 Genomes on GRCh38,1000 Genomes 30x on GRCh38,1000 Genomes phase 3 release,1000 Genomes phase 1 release,Geuvadis |
| **NA18864** | female | SAME124903 | YRI | Yoruba | AFR | African Ancestry | YRI | 1000 Genomes on GRCh38,1000 Genomes 30x on GRCh38,1000 Genomes phase 3 release |
| **NA12043** | male | SAME123214 | CEU | CEPH | EUR | European Ancestry | CEU | 1000 Genomes on GRCh38,1000 Genomes 30x on GRCh38,1000 Genomes phase 3 release,1000 Genomes phase 1 release,Geuvadis |
| **NA18869** | male | SAME124909 | YRI | Yoruba | AFR | African Ancestry | YRI | 1000 Genomes 30x on GRCh38,1000 Genomes phase 3 release |
| **NA12144** | male | SAME124156 | CEU | CEPH | EUR | European Ancestry | CEU | 1000 Genomes on GRCh38,1000 Genomes 30x on GRCh38,1000 Genomes phase 3 release,1000 Genomes phase 1 release,Geuvadis |
| **NA18871** | male | SAME125102 | YRI | Yoruba | AFR | African Ancestry | YRI | 1000 Genomes on GRCh38,1000 Genomes 30x on GRCh38,1000 Genomes phase 3 release,1000 Genomes phase 1 release |
| **NA12156** | female | SAME124952 | CEU | CEPH | EUR | European Ancestry | CEU | 1000 Genomes on GRCh38,1000 Genomes 30x on GRCh38,1000 Genomes phase 3 release,1000 Genomes phase 1 release,Geuvadis |
| **NA18876** | female | SAME125103 | YRI | Yoruba | AFR | African Ancestry | YRI | 1000 Genomes on GRCh38,1000 Genomes 30x on GRCh38,1000 Genomes phase 3 release |
| **NA12264** | male | SAME123087 | CEU | CEPH | EUR | European Ancestry | CEU | 1000 Genomes 30x on GRCh38,1000 Genomes phase 3 release |
| **NA18972** | female | SAME123762 | JPT | Japanese | EAS | East Asian Ancestry | JPT | 1000 Genomes on GRCh38,1000 Genomes 30x on GRCh38,1000 Genomes phase 3 release,1000 Genomes phase 1 release |
| **NA12283** | female | SAME124639 | CEU | CEPH | EUR | European Ancestry | CEU | 1000 Genomes on GRCh38,1000 Genomes 30x on GRCh38,1000 Genomes phase 3 release,1000 Genomes phase 1 release,Geuvadis |
| **NA18977** | male | SAME123765 | JPT | Japanese | EAS | East Asian Ancestry | JPT | 1000 Genomes on GRCh38,1000 Genomes 30x on GRCh38,1000 Genomes phase 3 release,1000 Genomes phase 1 release |
| **NA12341** | female | SAME125158 | CEU | CEPH | EUR | European Ancestry | CEU | 1000 Genomes on GRCh38,1000 Genomes 30x on GRCh38,1000 Genomes phase 3 release,1000 Genomes phase 1 release,Geuvadis |
| **NA18984** | male | SAME125071 | JPT | Japanese | EAS | East Asian Ancestry | JPT | 1000 Genomes on GRCh38,1000 Genomes 30x on GRCh38,1000 Genomes phase 3 release,1000 Genomes phase 1 release |
| **NA12485** | male | SAME125360 | CEU | CEPH | EUR | European Ancestry | CEU | 1000 Genomes 30x on GRCh38,1000 Genomes phase 3 release |
| **NA18989** | male | SAME125079 | JPT | Japanese | EAS | East Asian Ancestry | JPT | 1000 Genomes on GRCh38,1000 Genomes 30x on GRCh38,1000 Genomes phase 3 release,1000 Genomes phase 1 release |
| **NA07037** | female | SAME122862 | CEU | CEPH | EUR | European Ancestry | CEU | 1000 Genomes on GRCh38,1000 Genomes 30x on GRCh38,1000 Genomes phase 3 release,1000 Genomes phase 1 release,Geuvadis |
| **NA18991** | female | SAME124892 | JPT | Japanese | EAS | East Asian Ancestry | JPT | 1000 Genomes on GRCh38,1000 Genomes 30x on GRCh38,1000 Genomes phase 3 release,1000 Genomes phase 1 release |
| **NA07051** | male | SAME124378 | CEU | CEPH | EUR | European Ancestry | CEU | 1000 Genomes on GRCh38,1000 Genomes 30x on GRCh38,1000 Genomes phase 3 release,1000 Genomes phase 1 release,Geuvadis |
| **NA19002** | female | SAME124479 | JPT | Japanese | EAS | East Asian Ancestry | JPT | 1000 Genomes on GRCh38,1000 Genomes 30x on GRCh38,1000 Genomes phase 3 release,1000 Genomes phase 1 release |
| **NA07056** | female | SAME124375 | CEU | CEPH | EUR | European Ancestry | CEU | 1000 Genomes on GRCh38,1000 Genomes 30x on GRCh38,1000 Genomes phase 3 release,1000 Genomes phase 1 release,Geuvadis |
| **NA19007** | male | SAME124476 | JPT | Japanese | EAS | East Asian Ancestry | JPT | 1000 Genomes on GRCh38,1000 Genomes 30x on GRCh38,1000 Genomes phase 3 release,1000 Genomes phase 1 release |
| **NA07347** | male | SAME123572 | CEU | CEPH | EUR | European Ancestry | CEU | 1000 Genomes on GRCh38,1000 Genomes 30x on GRCh38,1000 Genomes phase 3 release,1000 Genomes phase 1 release,Geuvadis |
| **NA19019** | female | SAME124000 | LWK | Luhya | AFR | African Ancestry | LWK | 1000 Genomes on GRCh38,1000 Genomes 30x on GRCh38,1000 Genomes phase 3 release |
| **NA10838** | male | SAME124550 | CEU | CEPH | EUR | European Ancestry | CEU | 1000 Genomes 30x on GRCh38,1000 Genomes phase 3 release |
| **NA19153** | male | SAME123019 | YRI | Yoruba | AFR | African Ancestry | YRI | 1000 Genomes on GRCh38,1000 Genomes 30x on GRCh38,1000 Genomes phase 3 release,1000 Genomes phase 1 release,Geuvadis |
| **NA10840** | female | SAME124756 | CEU | CEPH | EUR | European Ancestry | CEU | 1000 Genomes 30x on GRCh38,1000 Genomes phase 3 release |
| **NA19160** | male | SAME123758 | YRI | Yoruba | AFR | African Ancestry | YRI | 1000 Genomes on GRCh38,1000 Genomes 30x on GRCh38,1000 Genomes phase 3 release,1000 Genomes phase 1 release,Geuvadis |
| **NA10845** | male | SAME124759 | CEU | CEPH | EUR | European Ancestry | CEU | 1000 Genomes 30x on GRCh38,1000 Genomes phase 3 release |
| **NA19172** | female | SAME124420 | YRI | Yoruba | AFR | African Ancestry | YRI | 1000 Genomes on GRCh38,1000 Genomes 30x on GRCh38,1000 Genomes phase 3 release,1000 Genomes phase 1 release,Geuvadis |
| **NA18748** | male | SAME123909 | CHB | Han Chinese | EAS | East Asian Ancestry | CHB | 1000 Genomes on GRCh38,1000 Genomes 30x on GRCh38,1000 Genomes phase 3 release,1000 Genomes phase 1 release |
| **NA19177** | male | SAME124417 | YRI | Yoruba | AFR | African Ancestry | YRI | 1000 Genomes 30x on GRCh38,1000 Genomes phase 3 release |
| **NA18856** | male | SAME124736 | YRI | Yoruba | AFR | African Ancestry | YRI | 1000 Genomes on GRCh38,1000 Genomes 30x on GRCh38,1000 Genomes phase 3 release,1000 Genomes phase 1 release |
| **NA19184** | male | SAME123859 | YRI | Yoruba | AFR | African Ancestry | YRI | 1000 Genomes on GRCh38,1000 Genomes 30x on GRCh38,1000 Genomes phase 3 release,1000 Genomes phase 1 release,Geuvadis |
| **NA18863** | male | SAME124907 | YRI | Yoruba | AFR | African Ancestry | YRI | 1000 Genomes 30x on GRCh38,1000 Genomes phase 3 release |
| **NA19189** | male | SAME123869 | YRI | Yoruba | AFR | African Ancestry | YRI | 1000 Genomes on GRCh38,1000 Genomes 30x on GRCh38,1000 Genomes phase 3 release,1000 Genomes phase 1 release,Geuvadis |
| **NA18868** | male | SAME124910 | YRI | Yoruba | AFR | African Ancestry | YRI | 1000 Genomes on GRCh38,1000 Genomes 30x on GRCh38,1000 Genomes phase 3 release,1000 Genomes phase 1 release,Geuvadis |
| **NA19191** | male | SAME124042 | YRI | Yoruba | AFR | African Ancestry | YRI | 1000 Genomes 30x on GRCh38,1000 Genomes phase 3 release |
| **NA18870** | female | SAME125101 | YRI | Yoruba | AFR | African Ancestry | YRI | 1000 Genomes on GRCh38,1000 Genomes 30x on GRCh38,1000 Genomes phase 3 release,1000 Genomes phase 1 release,Geuvadis |
| **NA18875** | female | SAME123545 | YRI | Yoruba | AFR | African Ancestry | YRI | 1000 Genomes 30x on GRCh38,1000 Genomes phase 3 release |
| **NA19204** | female | SAME122985 | YRI | Yoruba | AFR | African Ancestry | YRI | 1000 Genomes on GRCh38,1000 Genomes 30x on GRCh38,1000 Genomes phase 3 release,1000 Genomes phase 1 release,Geuvadis |
| **HG04222** | male | SAME1839040 | ITU | Telugu | SAS | South Asian Ancestry | ITU | 1000 Genomes on GRCh38,1000 Genomes 30x on GRCh38,1000 Genomes phase 3 release |
| **NA19209** | female | SAME122994 | YRI | Yoruba | AFR | African Ancestry | YRI | 1000 Genomes on GRCh38,1000 Genomes 30x on GRCh38,1000 Genomes phase 3 release,1000 Genomes phase 1 release,Geuvadis |
| **HG04227** | female | SAME1839056 | STU | Tamil | SAS | South Asian Ancestry | STU | 1000 Genomes on GRCh38,1000 Genomes 30x on GRCh38,1000 Genomes phase 3 release |
| **NA19211** | male | SAME125382 | YRI | Yoruba | AFR | African Ancestry | YRI | 1000 Genomes 30x on GRCh38,1000 Genomes phase 3 release |
| **HG04239** | male | SAME1840281 | ITU | Telugu | SAS | South Asian Ancestry | ITU | 1000 Genomes on GRCh38,1000 Genomes 30x on GRCh38,1000 Genomes phase 3 release |
| **NA19350** | male | SAME122853 | LWK | Luhya | AFR | African Ancestry | LWK | 1000 Genomes on GRCh38,1000 Genomes 30x on GRCh38,1000 Genomes phase 3 release,1000 Genomes phase 1 release |
| **NA06995** | male | SAME123994 | CEU | CEPH | EUR | European Ancestry | CEU | 1000 Genomes 30x on GRCh38,1000 Genomes phase 3 release |
| **NA19355** | female | SAME124077 | LWK | Luhya | AFR | African Ancestry | LWK | 1000 Genomes on GRCh38,1000 Genomes 30x on GRCh38,1000 Genomes phase 3 release,1000 Genomes phase 1 release |
| **HG04195** | female | SAME1839823 | BEB | Bengali | SAS | South Asian Ancestry | BEB | 1000 Genomes on GRCh38,1000 Genomes 30x on GRCh38,1000 Genomes phase 3 release |
| **NA19374** | male | SAME125115 | LWK | Luhya | AFR | African Ancestry | LWK | 1000 Genomes on GRCh38,1000 Genomes 30x on GRCh38,1000 Genomes phase 3 release,1000 Genomes phase 1 release |
| **NA19379** | female | SAME125092 | LWK | Luhya | AFR | African Ancestry | LWK | 1000 Genomes on GRCh38,1000 Genomes 30x on GRCh38,1000 Genomes phase 3 release,1000 Genomes phase 1 release |
| **HG04210** | male | SAME1839225 | STU | Tamil | SAS | South Asian Ancestry | STU | 1000 Genomes on GRCh38,1000 Genomes 30x on GRCh38,1000 Genomes phase 3 release |
| **HG04215** | male | SAME1839333 | STU | Tamil | SAS | South Asian Ancestry | STU | 1000 Genomes 30x on GRCh38 |
| **NA19393** | male | SAME124496 | LWK | Luhya | AFR | African Ancestry | LWK | 1000 Genomes on GRCh38,1000 Genomes 30x on GRCh38,1000 Genomes phase 3 release,1000 Genomes phase 1 release |
| **NA19037** | female | SAME124665 | LWK | Luhya | AFR | African Ancestry | LWK | 1000 Genomes on GRCh38,1000 Genomes 30x on GRCh38,1000 Genomes phase 3 release |
| **NA19401** | female | SAME123634 | LWK | Luhya | AFR | African Ancestry | LWK | 1000 Genomes on GRCh38,1000 Genomes 30x on GRCh38,1000 Genomes phase 3 release,1000 Genomes phase 1 release |
| **NA19056** | male | SAME125026 | JPT | Japanese | EAS | East Asian Ancestry | JPT | 1000 Genomes on GRCh38,1000 Genomes 30x on GRCh38,1000 Genomes phase 3 release,1000 Genomes phase 1 release |
| **NA19063** | male | SAME123006 | JPT | Japanese | EAS | East Asian Ancestry | JPT | 1000 Genomes on GRCh38,1000 Genomes 30x on GRCh38,1000 Genomes phase 3 release,1000 Genomes phase 1 release |
| **NA19437** | female | SAME123851 | LWK | Luhya | AFR | African Ancestry | LWK | 1000 Genomes on GRCh38,1000 Genomes 30x on GRCh38,1000 Genomes phase 3 release,1000 Genomes phase 1 release |
| **NA19068** | male | SAME123012 | JPT | Japanese | EAS | East Asian Ancestry | JPT | 1000 Genomes on GRCh38,1000 Genomes 30x on GRCh38,1000 Genomes phase 3 release,1000 Genomes phase 1 release |
| **NA19070** | male | SAME122820 | JPT | Japanese | EAS | East Asian Ancestry | JPT | 1000 Genomes on GRCh38,1000 Genomes 30x on GRCh38,1000 Genomes phase 3 release,1000 Genomes phase 1 release |
| **NA19449** | female | SAME122946 | LWK | Luhya | AFR | African Ancestry | LWK | 1000 Genomes on GRCh38,1000 Genomes 30x on GRCh38,1000 Genomes phase 3 release,1000 Genomes phase 1 release |
| **NA19075** | male | SAME122816 | JPT | Japanese | EAS | East Asian Ancestry | JPT | 1000 Genomes on GRCh38,1000 Genomes 30x on GRCh38,1000 Genomes phase 3 release,1000 Genomes phase 1 release |
| **NA19451** | male | SAME125415 | LWK | Luhya | AFR | African Ancestry | LWK | 1000 Genomes on GRCh38,1000 Genomes 30x on GRCh38,1000 Genomes phase 3 release,1000 Genomes phase 1 release |
| **NA19082** | male | SAME123370 | JPT | Japanese | EAS | East Asian Ancestry | JPT | 1000 Genomes on GRCh38,1000 Genomes 30x on GRCh38,1000 Genomes phase 3 release,1000 Genomes phase 1 release |
| **NA19456** | female | SAME125410 | LWK | Luhya | AFR | African Ancestry | LWK | 1000 Genomes on GRCh38,1000 Genomes 30x on GRCh38,1000 Genomes phase 3 release,1000 Genomes phase 1 release |
| **NA19087** | female | SAME123367 | JPT | Japanese | EAS | East Asian Ancestry | JPT | 1000 Genomes on GRCh38,1000 Genomes 30x on GRCh38,1000 Genomes phase 3 release,1000 Genomes phase 1 release |
| **NA19463** | female | SAME123297 | LWK | Luhya | AFR | African Ancestry | LWK | 1000 Genomes on GRCh38,1000 Genomes 30x on GRCh38,1000 Genomes phase 3 release,1000 Genomes phase 1 release |
| **NA19094** | female | SAME123171 | YRI | Yoruba | AFR | African Ancestry | YRI | 1000 Genomes 30x on GRCh38,1000 Genomes phase 3 release |
| **NA19468** | female | SAME123299 | LWK | Luhya | AFR | African Ancestry | LWK | 1000 Genomes on GRCh38,1000 Genomes 30x on GRCh38,1000 Genomes phase 3 release,1000 Genomes phase 1 release |
| **NA19099** | female | SAME125240 | YRI | Yoruba | AFR | African Ancestry | YRI | 1000 Genomes on GRCh38,1000 Genomes 30x on GRCh38,1000 Genomes phase 3 release,1000 Genomes phase 1 release,Geuvadis |
| **NA19222** | female | SAME124147 | YRI | Yoruba | AFR | African Ancestry | YRI | 1000 Genomes on GRCh38,1000 Genomes 30x on GRCh38,1000 Genomes phase 3 release,1000 Genomes phase 1 release,Geuvadis |
| **NA19475** | female | SAME123121 | LWK | Luhya | AFR | African Ancestry | LWK | 1000 Genomes on GRCh38,1000 Genomes 30x on GRCh38,1000 Genomes phase 3 release |
| **NA19239** | male | SAME125081 | YRI | Yoruba | AFR | African Ancestry | YRI | 1000 Genomes on GRCh38,1000 Genomes 30x on GRCh38,Human Genome Structural Variation Consortium, Phase 2,The Human Genome Structural Variation Consortiu |
| **NA19653** | female | SAME123125 | MXL | Mexican Ancestry | AMR | American Ancestry | MXL | 1000 Genomes 30x on GRCh38,1000 Genomes phase 3 release |
| **NA19258** | male | SAME125045 | YRI | Yoruba | AFR | African Ancestry | YRI | 1000 Genomes 30x on GRCh38,1000 Genomes phase 3 release |
| **NA19658** | male | SAME123132 | MXL | Mexican Ancestry | AMR | American Ancestry | MXL | 1000 Genomes on GRCh38,1000 Genomes 30x on GRCh38,1000 Genomes phase 3 release,1000 Genomes phase 1 release |
| **NA19309** | male | SAME124984 | LWK | Luhya | AFR | African Ancestry | LWK | 1000 Genomes on GRCh38,1000 Genomes 30x on GRCh38,1000 Genomes phase 3 release,1000 Genomes phase 1 release |
| **NA19660** | female | SAME122958 | MXL | Mexican Ancestry | AMR | American Ancestry | MXL | 1000 Genomes on GRCh38,1000 Genomes 30x on GRCh38,1000 Genomes phase 3 release,1000 Genomes phase 1 release |
| **NA19665** | female | SAME122956 | MXL | Mexican Ancestry | AMR | American Ancestry | MXL | 1000 Genomes 30x on GRCh38,1000 Genomes phase 3 release |
| **NA19316** | female | SAME123311 | LWK | Luhya | AFR | African Ancestry | LWK | 1000 Genomes on GRCh38,1000 Genomes 30x on GRCh38,1000 Genomes phase 3 release,1000 Genomes phase 1 release |
| **NA19323** | female | SAME123471 | LWK | Luhya | AFR | African Ancestry | LWK | 1000 Genomes on GRCh38,1000 Genomes 30x on GRCh38,1000 Genomes phase 3 release |
| **NA19677** | female | SAME122789 | MXL | Mexican Ancestry | AMR | American Ancestry | MXL | 1000 Genomes 30x on GRCh38,1000 Genomes phase 3 release |
| **NA19328** | female | SAME123463 | LWK | Luhya | AFR | African Ancestry | LWK | 1000 Genomes on GRCh38,1000 Genomes 30x on GRCh38,1000 Genomes phase 3 release,1000 Genomes phase 1 release |
| **NA19684** | female | SAME122941 | MXL | Mexican Ancestry | AMR | American Ancestry | MXL | 1000 Genomes on GRCh38,1000 Genomes 30x on GRCh38,1000 Genomes phase 3 release,1000 Genomes phase 1 release |
| **NA19347** | male | SAME124340 | LWK | Luhya | AFR | African Ancestry | LWK | 1000 Genomes on GRCh38,1000 Genomes 30x on GRCh38,1000 Genomes phase 3 release,1000 Genomes phase 1 release |
| **NA19704** | female | SAME124236 | ASW | African Ancestry SW | AFR | African Ancestry | ASW | 1000 Genomes on GRCh38,1000 Genomes 30x on GRCh38,1000 Genomes phase 3 release,1000 Genomes phase 1 release |
| **NA19711** | male | SAME124446 | ASW | African Ancestry SW | AFR | African Ancestry | ASW | 1000 Genomes on GRCh38,1000 Genomes 30x on GRCh38,1000 Genomes phase 3 release,1000 Genomes phase 1 release |
| **NA19716** | female | SAME124450 | MXL | Mexican Ancestry | AMR | American Ancestry | MXL | 1000 Genomes on GRCh38,1000 Genomes 30x on GRCh38,1000 Genomes phase 3 release,1000 Genomes phase 1 release |
| **NA19378** | female | SAME125091 | LWK | Luhya | AFR | African Ancestry | LWK | 1000 Genomes on GRCh38,1000 Genomes 30x on GRCh38,1000 Genomes phase 3 release |
| **NA19723** | male | SAME124695 | MXL | Mexican Ancestry | AMR | American Ancestry | MXL | 1000 Genomes on GRCh38,1000 Genomes 30x on GRCh38,1000 Genomes phase 3 release,1000 Genomes phase 1 release |
| **NA19380** | male | SAME124276 | LWK | Luhya | AFR | African Ancestry | LWK | 1000 Genomes on GRCh38,1000 Genomes 30x on GRCh38,1000 Genomes phase 3 release,1000 Genomes phase 1 release |
| **NA19728** | female | SAME124702 | MXL | Mexican Ancestry | AMR | American Ancestry | MXL | 1000 Genomes on GRCh38,1000 Genomes 30x on GRCh38,1000 Genomes phase 3 release,1000 Genomes phase 1 release |
| **NA19385** | male | SAME124271 | LWK | Luhya | AFR | African Ancestry | LWK | 1000 Genomes on GRCh38,1000 Genomes 30x on GRCh38,1000 Genomes phase 3 release,1000 Genomes phase 1 release |
| **NA19730** | female | SAME124869 | MXL | Mexican Ancestry | AMR | American Ancestry | MXL | 1000 Genomes 30x on GRCh38,1000 Genomes phase 3 release |
| **NA19397** | male | SAME124492 | LWK | Luhya | AFR | African Ancestry | LWK | 1000 Genomes on GRCh38,1000 Genomes 30x on GRCh38,1000 Genomes phase 3 release,1000 Genomes phase 1 release |
| **NA19735** | male | SAME124874 | MXL | Mexican Ancestry | AMR | American Ancestry | MXL | 1000 Genomes on GRCh38,1000 Genomes 30x on GRCh38,1000 Genomes phase 3 release,1000 Genomes phase 1 release |
| **NA19429** | male | SAME124037 | LWK | Luhya | AFR | African Ancestry | LWK | 1000 Genomes on GRCh38,1000 Genomes 30x on GRCh38,1000 Genomes phase 3 release,1000 Genomes phase 1 release |
| **NA19785** | female | SAME123150 | MXL | Mexican Ancestry | AMR | American Ancestry | MXL | 1000 Genomes on GRCh38,1000 Genomes 30x on GRCh38,1000 Genomes phase 3 release,1000 Genomes phase 1 release |
| **NA19431** | female | SAME123853 | LWK | Luhya | AFR | African Ancestry | LWK | 1000 Genomes on GRCh38,1000 Genomes 30x on GRCh38,1000 Genomes phase 3 release,1000 Genomes phase 1 release |
| **NA19792** | male | SAME123327 | MXL | Mexican Ancestry | AMR | American Ancestry | MXL | 1000 Genomes on GRCh38,1000 Genomes 30x on GRCh38,1000 Genomes phase 3 release |
| **NA19436** | female | SAME123852 | LWK | Luhya | AFR | African Ancestry | LWK | 1000 Genomes on GRCh38,1000 Genomes 30x on GRCh38,1000 Genomes phase 3 release,1000 Genomes phase 1 release |
| **NA19443** | male | SAME122954 | LWK | Luhya | AFR | African Ancestry | LWK | 1000 Genomes on GRCh38,1000 Genomes 30x on GRCh38,1000 Genomes phase 3 release,1000 Genomes phase 1 release |
| **NA19836** | female | SAME125031 | ASW | African Ancestry SW | AFR | African Ancestry | ASW | 1000 Genomes 30x on GRCh38,1000 Genomes phase 3 release |
| **NA19448** | male | SAME122945 | LWK | Luhya | AFR | African Ancestry | LWK | 1000 Genomes on GRCh38,1000 Genomes 30x on GRCh38,1000 Genomes phase 3 release,1000 Genomes phase 1 release |
| **NA19901** | female | SAME125049 | ASW | African Ancestry SW | AFR | African Ancestry | ASW | 1000 Genomes on GRCh38,1000 Genomes 30x on GRCh38,1000 Genomes phase 3 release,1000 Genomes phase 1 release |
| **NA18907** | female | SAME123563 | YRI | Yoruba | AFR | African Ancestry | YRI | 1000 Genomes on GRCh38,1000 Genomes 30x on GRCh38,1000 Genomes phase 3 release,1000 Genomes phase 1 release,Geuvadis |

| | | | | | | | | |
|---|---|---|---|---|---|---|---|---|
| NA19913 | female | SAME124432 | ASW | African Ancestry SW | AFR | African Ancestry | ASW | 1000 Genomes on GRCh38,1000 Genomes 30x on GRCh38,1000 Genomes phase 3 release |
| NA18914 | male | SAME123406 | YRI | Yoruba | AFR | African Ancestry | YRI | 1000 Genomes 30x on GRCh38,1000 Genomes phase 3 release |
| NA19918 | male | SAME125275 | ASW | African Ancestry SW | AFR | African Ancestry | ASW | 1000 Genomes 30x on GRCh38,1000 Genomes phase 3 release |
| NA18933 | female | SAME123037 | YRI | Yoruba | AFR | African Ancestry | YRI | 1000 Genomes on GRCh38,1000 Genomes 30x on GRCh38,1000 Genomes phase 3 release,1000 Genomes phase 1 release,Geuvadis |
| NA19920 | male | SAME124713 | ASW | African Ancestry SW | AFR | African Ancestry | ASW | 1000 Genomes on GRCh38,1000 Genomes 30x on GRCh38,1000 Genomes phase 3 release,1000 Genomes phase 1 release |
| NA18940 | male | SAME124310 | JPT | Japanese,Japanese | EAS | East Asia (SGDP),East Asian Ancestry | JapaneseSGDP,JPT | 1000 Genomes on GRCh38,Simons Genome Diversity Project,1000 Genomes 30x on GRCh38,1000 Genomes phase 3 release,1000 Genomes phase 1 release |
| NA19982 | male | SAME124293 | ASW | African Ancestry SW | AFR | African Ancestry | ASW | 1000 Genomes on GRCh38,1000 Genomes 30x on GRCh38,1000 Genomes phase 3 release,1000 Genomes phase 1 release |
| NA18945 | male | SAME124315 | JPT | Japanese | EAS | East Asian Ancestry | JPT | 1000 Genomes on GRCh38,1000 Genomes 30x on GRCh38,1000 Genomes phase 3 release,1000 Genomes phase 1 release |
| NA19223 | male | SAME125223 | YRI | Yoruba | AFR | African Ancestry | YRI | 1000 Genomes on GRCh38,1000 Genomes 30x on GRCh38,1000 Genomes phase 3 release,1000 Genomes phase 1 release,Geuvadis |
| NA18952 | male | SAME124114 | JPT | Japanese | EAS | East Asian Ancestry | JPT | 1000 Genomes on GRCh38,1000 Genomes 30x on GRCh38,1000 Genomes phase 3 release,1000 Genomes phase 1 release |
| NA19235 | female | SAME125002 | YRI | Yoruba | AFR | African Ancestry | YRI | 1000 Genomes on GRCh38,1000 Genomes 30x on GRCh38,1000 Genomes phase 3 release,1000 Genomes phase 1 release,Geuvadis |
| NA18957 | female | SAME124109 | JPT | Japanese | EAS | East Asian Ancestry | JPT | 1000 Genomes on GRCh38,1000 Genomes 30x on GRCh38,1000 Genomes phase 3 release,1000 Genomes phase 1 release |
| NA19247 | female | SAME123691 | YRI | Yoruba | AFR | African Ancestry | YRI | 1000 Genomes on GRCh38,1000 Genomes 30x on GRCh38,1000 Genomes phase 3 release,1000 Genomes phase 1 release,Geuvadis |
| NA18964 | female | SAME123960 | JPT | Japanese | EAS | East Asian Ancestry | JPT | 1000 Genomes on GRCh38,1000 Genomes 30x on GRCh38,1000 Genomes phase 3 release,1000 Genomes phase 1 release |
| NA18969 | female | SAME123954 | JPT | Japanese | EAS | East Asian Ancestry | JPT | 1000 Genomes on GRCh38,1000 Genomes 30x on GRCh38,1000 Genomes phase 3 release,1000 Genomes phase 1 release |
| NA19312 | male | SAME123315 | LWK | Luhya | AFR | African Ancestry | LWK | 1000 Genomes on GRCh38,1000 Genomes 30x on GRCh38,1000 Genomes phase 3 release,1000 Genomes phase 1 release |
| NA18971 | male | SAME123763 | JPT | Japanese | EAS | East Asian Ancestry | JPT | 1000 Genomes on GRCh38,1000 Genomes 30x on GRCh38,1000 Genomes phase 3 release,1000 Genomes phase 1 release |
| NA19317 | male | SAME123312 | LWK | Luhya | AFR | African Ancestry | LWK | 1000 Genomes on GRCh38,1000 Genomes 30x on GRCh38,1000 Genomes phase 3 release,1000 Genomes phase 1 release |
| NA18976 | female | SAME123766 | JPT | Japanese | EAS | East Asian Ancestry | JPT | 1000 Genomes on GRCh38,1000 Genomes 30x on GRCh38,1000 Genomes phase 3 release,1000 Genomes phase 1 release |
| NA19324 | female | SAME123472 | LWK | Luhya | AFR | African Ancestry | LWK | 1000 Genomes on GRCh38,1000 Genomes 30x on GRCh38,1000 Genomes phase 3 release,1000 Genomes phase 1 release |
| NA18983 | male | SAME125074 | JPT | Japanese | EAS | East Asian Ancestry | JPT | 1000 Genomes on GRCh38,1000 Genomes 30x on GRCh38,1000 Genomes phase 3 release,1000 Genomes phase 1 release |
| NA19331 | male | SAME123250 | LWK | Luhya | AFR | African Ancestry | LWK | 1000 Genomes on GRCh38,1000 Genomes 30x on GRCh38,1000 Genomes phase 3 release,1000 Genomes phase 1 release |
| NA18988 | male | SAME125078 | JPT | Japanese | EAS | East Asian Ancestry | JPT | 1000 Genomes on GRCh38,1000 Genomes 30x on GRCh38,1000 Genomes phase 3 release,1000 Genomes phase 1 release |
| NA19103 | male | SAME124330 | YRI | Yoruba | AFR | African Ancestry | YRI | 1000 Genomes 30x on GRCh38,1000 Genomes phase 3 release |
| NA18990 | male | SAME124893 | JPT | Japanese | EAS | East Asian Ancestry | JPT | 1000 Genomes on GRCh38,1000 Genomes 30x on GRCh38,1000 Genomes phase 3 release,1000 Genomes phase 1 release |
| NA19108 | female | SAME125378 | YRI | Yoruba | AFR | African Ancestry | YRI | 1000 Genomes on GRCh38,1000 Genomes 30x on GRCh38,1000 Genomes phase 3 release,1000 Genomes phase 1 release,Geuvadis |
| NA18995 | male | SAME124896 | JPT | Japanese | EAS | East Asian Ancestry | JPT | 1000 Genomes on GRCh38,1000 Genomes 30x on GRCh38,1000 Genomes phase 3 release,1000 Genomes phase 1 release |
| NA19115 | female | SAME123857 | YRI | Yoruba | AFR | African Ancestry | YRI | 1000 Genomes 30x on GRCh38,1000 Genomes phase 3 release |
| NA19001 | female | SAME124478 | JPT | Japanese | EAS | East Asian Ancestry | JPT | 1000 Genomes on GRCh38,1000 Genomes 30x on GRCh38,1000 Genomes phase 3 release,1000 Genomes phase 1 release |
| NA19122 | female | SAME125014 | YRI | Yoruba | AFR | African Ancestry | YRI | 1000 Genomes 30x on GRCh38,1000 Genomes phase 3 release |
| NA19006 | male | SAME125119 | JPT | Japanese | EAS | East Asian Ancestry | JPT | 1000 Genomes on GRCh38,1000 Genomes 30x on GRCh38,1000 Genomes phase 3 release |
| NA19127 | female | SAME125017 | YRI | Yoruba | AFR | African Ancestry | YRI | 1000 Genomes 30x on GRCh38,1000 Genomes phase 3 release |
| NA19020 | male | SAME124855 | LWK | Luhya | AFR | African Ancestry | LWK | 1000 Genomes on GRCh38,1000 Genomes 30x on GRCh38,1000 Genomes phase 3 release,1000 Genomes phase 1 release |
| NA19139 | male | SAME125237 | YRI | Yoruba | AFR | African Ancestry | YRI | 1000 Genomes 30x on GRCh38,1000 Genomes phase 3 release |
| NA19025 | male | SAME124852 | LWK | Luhya | AFR | African Ancestry | LWK | 1000 Genomes on GRCh38,1000 Genomes 30x on GRCh38,1000 Genomes phase 3 release |
| NA19141 | male | SAME124607 | YRI | Yoruba | AFR | African Ancestry | YRI | 1000 Genomes on GRCh38,1000 Genomes 30x on GRCh38,1000 Genomes phase 3 release,1000 Genomes phase 1 release,Geuvadis |
| NA19152 | female | SAME123017 | YRI | Yoruba | AFR | African Ancestry | YRI | 1000 Genomes on GRCh38,1000 Genomes 30x on GRCh38,1000 Genomes phase 3 release,1000 Genomes phase 1 release,Geuvadis |
| NA19146 | male | SAME125080 | YRI | Yoruba | AFR | African Ancestry | YRI | 1000 Genomes on GRCh38,1000 Genomes 30x on GRCh38,1000 Genomes phase 3 release,1000 Genomes phase 1 release,Geuvadis |
| NA19171 | male | SAME124419 | YRI | Yoruba | AFR | African Ancestry | YRI | 1000 Genomes on GRCh38,1000 Genomes 30x on GRCh38,1000 Genomes phase 3 release,1000 Genomes phase 1 release,Geuvadis |
| NA19176 | female | SAME124416 | YRI | Yoruba | AFR | African Ancestry | YRI | 1000 Genomes 30x on GRCh38,1000 Genomes phase 3 release |
| NA19747 | male | SAME123501 | MXL | Mexican Ancestry | AMR | American Ancestry | MXL | 1000 Genomes on GRCh38,1000 Genomes 30x on GRCh38,1000 Genomes phase 3 release,1000 Genomes phase 1 release |
| NA19190 | female | SAME124041 | YRI | Yoruba | AFR | African Ancestry | YRI | 1000 Genomes on GRCh38,1000 Genomes 30x on GRCh38,1000 Genomes phase 3 release,1000 Genomes phase 1 release,Geuvadis |
| NA19759 | male | SAME123707 | MXL | Mexican Ancestry | AMR | American Ancestry | MXL | 1000 Genomes on GRCh38,1000 Genomes 30x on GRCh38,1000 Genomes phase 3 release,1000 Genomes phase 1 release |
| NA19203 | male | SAME122988 | YRI | Yoruba | AFR | African Ancestry | YRI | 1000 Genomes 30x on GRCh38,1000 Genomes phase 3 release |
| NA19761 | female | SAME123894 | MXL | Mexican Ancestry | AMR | American Ancestry | MXL | 1000 Genomes on GRCh38,1000 Genomes 30x on GRCh38,1000 Genomes phase 3 release,1000 Genomes phase 1 release |
| NA19208 | male | SAME122993 | YRI | Yoruba | AFR | African Ancestry | YRI | 1000 Genomes 30x on GRCh38,1000 Genomes phase 3 release |
| NA19210 | male | SAME125383 | YRI | Yoruba | AFR | African Ancestry | YRI | 1000 Genomes on GRCh38,1000 Genomes 30x on GRCh38,1000 Genomes phase 3 release,1000 Genomes phase 1 release,Geuvadis |
| NA19773 | female | SAME124067 | MXL | Mexican Ancestry | AMR | American Ancestry | MXL | 1000 Genomes on GRCh38,1000 Genomes 30x on GRCh38,1000 Genomes phase 3 release,1000 Genomes phase 1 release |
| NA19215 | female | SAME125385 | YRI | Yoruba | AFR | African Ancestry | YRI | 1000 Genomes 30x on GRCh38,1000 Genomes phase 3 release |
| NA19778 | male | SAME124060 | MXL | Mexican Ancestry | AMR | American Ancestry | MXL | 1000 Genomes 30x on GRCh38,1000 Genomes phase 3 release |
| NA19455 | male | SAME125411 | LWK | Luhya | AFR | African Ancestry | LWK | 1000 Genomes on GRCh38,1000 Genomes 30x on GRCh38,1000 Genomes phase 3 release,1000 Genomes phase 1 release |
| NA19780 | male | SAME123147 | MXL | Mexican Ancestry | AMR | American Ancestry | MXL | 1000 Genomes on GRCh38,1000 Genomes 30x on GRCh38,1000 Genomes phase 3 release,1000 Genomes phase 1 release |
| NA19462 | female | SAME123296 | LWK | Luhya | AFR | African Ancestry | LWK | 1000 Genomes on GRCh38,1000 Genomes 30x on GRCh38,1000 Genomes phase 3 release,1000 Genomes phase 1 release |
| NA20129 | female | SAME122886 | ASW | African Ancestry SW | AFR | African Ancestry | ASW | 1000 Genomes 30x on GRCh38,1000 Genomes phase 3 release |
| NA19467 | female | SAME123295 | LWK | Luhya | AFR | African Ancestry | LWK | 1000 Genomes on GRCh38,1000 Genomes 30x on GRCh38,1000 Genomes phase 3 release,1000 Genomes phase 1 release |
| NA20282 | female | SAME124583 | ASW | African Ancestry SW | AFR | African Ancestry | ASW | 1000 Genomes on GRCh38,1000 Genomes 30x on GRCh38,1000 Genomes phase 3 release,1000 Genomes phase 1 release |
| NA19474 | female | SAME123122 | LWK | Luhya | AFR | African Ancestry | LWK | 1000 Genomes on GRCh38,1000 Genomes 30x on GRCh38,1000 Genomes phase 3 release,1000 Genomes phase 1 release |
| NA20287 | female | SAME124580 | ASW | African Ancestry SW | AFR | African Ancestry | ASW | 1000 Genomes on GRCh38,1000 Genomes 30x on GRCh38,1000 Genomes phase 3 release,1000 Genomes phase 1 release |
| NA19652 | male | SAME124978 | MXL | Mexican Ancestry | AMR | American Ancestry | MXL | 1000 Genomes on GRCh38,1000 Genomes 30x on GRCh38,1000 Genomes phase 3 release,1000 Genomes phase 1 release |
| NA20294 | female | SAME124776 | ASW | African Ancestry SW | AFR | African Ancestry | ASW | 1000 Genomes on GRCh38,1000 Genomes 30x on GRCh38,1000 Genomes phase 3 release,1000 Genomes phase 1 release |
| NA19657 | female | SAME123129 | MXL | Mexican Ancestry | AMR | American Ancestry | MXL | 1000 Genomes on GRCh38,1000 Genomes 30x on GRCh38,1000 Genomes phase 3 release,1000 Genomes phase 1 release |
| NA20299 | female | SAME124381 | ASW | African Ancestry SW | AFR | African Ancestry | ASW | 1000 Genomes on GRCh38,1000 Genomes 30x on GRCh38,1000 Genomes phase 3 release,1000 Genomes phase 1 release |
| NA19664 | male | SAME122955 | MXL | Mexican Ancestry | AMR | American Ancestry | MXL | 1000 Genomes on GRCh38,1000 Genomes 30x on GRCh38,1000 Genomes phase 3 release,1000 Genomes phase 1 release |
| NA19669 | female | SAME123903 | MXL | Mexican Ancestry | AMR | American Ancestry | MXL | 1000 Genomes on GRCh38,1000 Genomes 30x on GRCh38,1000 Genomes phase 3 release,1000 Genomes phase 1 release |
| NA20314 | female | SAME125304 | ASW | African Ancestry SW | AFR | African Ancestry | ASW | 1000 Genomes on GRCh38,1000 Genomes 30x on GRCh38,1000 Genomes phase 3 release,1000 Genomes phase 1 release |
| NA19671 | female | SAME122794 | MXL | Mexican Ancestry | AMR | American Ancestry | MXL | 1000 Genomes 30x on GRCh38,1000 Genomes phase 3 release |
| NA19102 | female | SAME125112 | YRI | Yoruba | AFR | African Ancestry | YRI | 1000 Genomes on GRCh38,1000 Genomes 30x on GRCh38,1000 Genomes phase 3 release,1000 Genomes phase 1 release,Geuvadis |
| NA19107 | male | SAME123393 | YRI | Yoruba | AFR | African Ancestry | YRI | 1000 Genomes on GRCh38,1000 Genomes 30x on GRCh38,1000 Genomes phase 3 release,1000 Genomes phase 1 release,Geuvadis |
| NA20504 | female | SAME124360 | TSI | Toscani | EUR | European Ancestry | TSI | 1000 Genomes on GRCh38,1000 Genomes 30x on GRCh38,1000 Genomes phase 3 release,1000 Genomes phase 1 release,Geuvadis |
| NA19114 | female | SAME122953 | YRI | Yoruba | AFR | African Ancestry | YRI | 1000 Genomes on GRCh38,1000 Genomes 30x on GRCh38,1000 Genomes phase 3 release,1000 Genomes phase 1 release,Geuvadis |
| NA20509 | male | SAME124354 | TSI | Toscani | EUR | European Ancestry | TSI | 1000 Genomes on GRCh38,1000 Genomes 30x on GRCh38,Human Genome Structural Variation Consortium, Phase 2,1000 Genomes phase 3 release,1000 Genomes ph |
| NA19119 | male | SAME122944 | YRI | Yoruba | AFR | African Ancestry | YRI | 1000 Genomes on GRCh38,1000 Genomes 30x on GRCh38,1000 Genomes phase 3 release,1000 Genomes phase 1 release,Geuvadis |
| NA20511 | male | SAME124563 | TSI | Toscani | EUR | European Ancestry | TSI | 1000 Genomes on GRCh38,1000 Genomes 30x on GRCh38,1000 Genomes phase 3 release |
| NA19121 | male | SAME125015 | YRI | Yoruba | AFR | African Ancestry | YRI | 1000 Genomes on GRCh38,1000 Genomes 30x on GRCh38,1000 Genomes phase 3 release,1000 Genomes phase 1 release,Geuvadis |
| NA20516 | male | SAME124556 | TSI | Toscani | EUR | European Ancestry | TSI | 1000 Genomes on GRCh38,1000 Genomes 30x on GRCh38,1000 Genomes phase 3 release,1000 Genomes phase 1 release,Geuvadis |
| NA19138 | male | SAME125236 | YRI | Yoruba | AFR | African Ancestry | YRI | 1000 Genomes on GRCh38,1000 Genomes 30x on GRCh38,1000 Genomes phase 3 release,1000 Genomes phase 1 release,Geuvadis |
| NA20528 | male | SAME124012 | TSI | Toscani | EUR | European Ancestry | TSI | 1000 Genomes on GRCh38,1000 Genomes 30x on GRCh38,1000 Genomes phase 3 release,1000 Genomes phase 1 release,Geuvadis |
| NA19140 | female | SAME124608 | YRI | Yoruba | AFR | African Ancestry | YRI | 1000 Genomes 30x on GRCh38,1000 Genomes phase 3 release |
| NA20530 | female | SAME124169 | TSI | Toscani | EUR | European Ancestry | TSI | 1000 Genomes on GRCh38,1000 Genomes 30x on GRCh38,1000 Genomes phase 3 release,1000 Genomes phase 1 release,Geuvadis |
| NA19145 | male | SAME123372 | YRI | Yoruba | AFR | African Ancestry | YRI | 1000 Genomes 30x on GRCh38,1000 Genomes phase 3 release |
| NA20535 | female | SAME124166 | TSI | Toscani | EUR | European Ancestry | TSI | 1000 Genomes on GRCh38,1000 Genomes 30x on GRCh38,1000 Genomes phase 3 release,1000 Genomes phase 1 release,Geuvadis |
| NA19676 | male | SAME122790 | MXL | Mexican Ancestry | AMR | American Ancestry | MXL | 1000 Genomes on GRCh38,1000 Genomes 30x on GRCh38,1000 Genomes phase 3 release,1000 Genomes phase 1 release |
| NA20542 | female | SAME125150 | TSI | Toscani | EUR | European Ancestry | TSI | 1000 Genomes on GRCh38,1000 Genomes 30x on GRCh38,1000 Genomes phase 3 release,1000 Genomes phase 1 release,Geuvadis |
| NA19683 | female | SAME122936 | MXL | Mexican Ancestry | AMR | American Ancestry | MXL | 1000 Genomes 30x on GRCh38,1000 Genomes phase 3 release |
| NA20585 | female | SAME122861 | TSI | Toscani | EUR | European Ancestry | TSI | 1000 Genomes on GRCh38,1000 Genomes 30x on GRCh38,1000 Genomes phase 3 release,1000 Genomes phase 1 release,Geuvadis |
| NA19703 | male | SAME124230 | ASW | African Ancestry SW | AFR | African Ancestry | ASW | 1000 Genomes on GRCh38,1000 Genomes 30x on GRCh38,1000 Genomes phase 3 release,1000 Genomes phase 1 release |
| NA20756 | female | SAME124175 | TSI | Toscani | EUR | European Ancestry | TSI | 1000 Genomes on GRCh38,1000 Genomes 30x on GRCh38,1000 Genomes phase 3 release,1000 Genomes phase 1 release,Geuvadis |
| NA20763 | male | SAME124365 | TSI | Toscani | EUR | European Ancestry | TSI | 1000 Genomes on GRCh38,1000 Genomes 30x on GRCh38,1000 Genomes phase 3 release |
| NA19722 | female | SAME124696 | MXL | Mexican Ancestry | AMR | American Ancestry | MXL | 1000 Genomes on GRCh38,1000 Genomes 30x on GRCh38,1000 Genomes phase 3 release,1000 Genomes phase 1 release |
| NA20768 | female | SAME124372 | TSI | Toscani | EUR | European Ancestry | TSI | 1000 Genomes on GRCh38,1000 Genomes 30x on GRCh38,1000 Genomes phase 3 release,1000 Genomes phase 1 release,Geuvadis |
| NA19727 | male | SAME123639 | MXL | Mexican Ancestry | AMR | American Ancestry | MXL | 1000 Genomes 30x on GRCh38,1000 Genomes phase 3 release |
| NA20770 | male | SAME124576 | TSI | Toscani | EUR | European Ancestry | TSI | 1000 Genomes on GRCh38,1000 Genomes 30x on GRCh38,1000 Genomes phase 3 release,1000 Genomes phase 1 release,Geuvadis |
| NA19734 | female | SAME124873 | MXL | Mexican Ancestry | AMR | American Ancestry | MXL | 1000 Genomes on GRCh38,1000 Genomes 30x on GRCh38,1000 Genomes phase 3 release,1000 Genomes phase 1 release |
| NA20775 | female | SAME124573 | TSI | Toscani | EUR | European Ancestry | TSI | 1000 Genomes on GRCh38,1000 Genomes 30x on GRCh38,1000 Genomes phase 3 release,1000 Genomes phase 1 release |
| NA20787 | male | SAME123609 | TSI | Toscani | EUR | European Ancestry | TSI | 1000 Genomes on GRCh38,1000 Genomes 30x on GRCh38,1000 Genomes phase 3 release,1000 Genomes phase 1 release,Geuvadis |
| NA20503 | female | SAME124357 | TSI | Toscani | EUR | European Ancestry | TSI | 1000 Genomes on GRCh38,1000 Genomes 30x on GRCh38,1000 Genomes phase 3 release,1000 Genomes phase 1 release,Geuvadis |
| NA20799 | female | SAME123822 | TSI | Toscani | EUR | European Ancestry | TSI | 1000 Genomes on GRCh38,1000 Genomes 30x on GRCh38,1000 Genomes phase 3 release,1000 Genomes phase 1 release,Geuvadis |
| NA20508 | female | SAME124355 | TSI | Toscani | EUR | European Ancestry | TSI | 1000 Genomes on GRCh38,1000 Genomes 30x on GRCh38,1000 Genomes phase 3 release,1000 Genomes phase 1 release,Geuvadis |
| NA20802 | female | SAME125366 | TSI | Toscani | EUR | European Ancestry | TSI | 1000 Genomes on GRCh38,1000 Genomes 30x on GRCh38,1000 Genomes phase 3 release,1000 Genomes phase 1 release,Geuvadis |
| NA20510 | male | SAME124562 | TSI | Toscani | EUR | European Ancestry | TSI | 1000 Genomes on GRCh38,1000 Genomes 30x on GRCh38,1000 Genomes phase 3 release,1000 Genomes phase 1 release,Geuvadis |
| NA20807 | female | SAME125363 | TSI | Toscani | EUR | European Ancestry | TSI | 1000 Genomes on GRCh38,1000 Genomes 30x on GRCh38,1000 Genomes phase 3 release,1000 Genomes phase 1 release,Geuvadis |
| NA20515 | male | SAME124559 | TSI | Toscani | EUR | European Ancestry | TSI | 1000 Genomes on GRCh38,1000 Genomes 30x on GRCh38,1000 Genomes phase 3 release,1000 Genomes phase 1 release,Geuvadis |
| NA20814 | male | SAME125188 | TSI | Toscani | EUR | European Ancestry | TSI | 1000 Genomes on GRCh38,1000 Genomes 30x on GRCh38,1000 Genomes phase 3 release,1000 Genomes phase 1 release,Geuvadis |
| NA20522 | female | SAME124017 | TSI | Toscani | EUR | European Ancestry | TSI | 1000 Genomes on GRCh38,1000 Genomes 30x on GRCh38,1000 Genomes phase 3 release,1000 Genomes phase 1 release |
| NA20819 | female | SAME125183 | TSI | Toscani | EUR | European Ancestry | TSI | 1000 Genomes on GRCh38,1000 Genomes 30x on GRCh38,1000 Genomes phase 3 release,1000 Genomes phase 1 release,Geuvadis |
| NA20527 | male | SAME124013 | TSI | Toscani | EUR | European Ancestry | TSI | 1000 Genomes on GRCh38,1000 Genomes 30x on GRCh38,1000 Genomes phase 3 release,1000 Genomes phase 1 release,Geuvadis |
| NA20821 | female | SAME123075 | TSI | Toscani | EUR | European Ancestry | TSI | 1000 Genomes on GRCh38,1000 Genomes 30x on GRCh38,1000 Genomes phase 3 release |
| NA20534 | male | SAME124165 | TSI | Toscani | EUR | European Ancestry | TSI | 1000 Genomes on GRCh38,1000 Genomes 30x on GRCh38,1000 Genomes phase 3 release,1000 Genomes phase 1 release,Geuvadis |
| NA20826 | female | SAME123070 | TSI | Toscani | EUR | European Ancestry | TSI | 1000 Genomes on GRCh38,1000 Genomes 30x on GRCh38,1000 Genomes phase 3 release,1000 Genomes phase 1 release,Geuvadis |
| NA20539 | male | SAME124164 | TSI | Toscani | EUR | European Ancestry | TSI | 1000 Genomes on GRCh38,1000 Genomes 30x on GRCh38,1000 Genomes phase 3 release,1000 Genomes phase 1 release,Geuvadis |
| NA20845 | male | SAME123430 | GIH | Gujarati | SAS | South Asian Ancestry | GIH | 1000 Genomes on GRCh38,1000 Genomes 30x on GRCh38,1000 Genomes phase 3 release |

| | | | | | | | | |
|---|---|---|---|---|---|---|---|---|
| **NA20541** | female | SAME125147 | TSI | Toscani | EUR | European Ancestry | TSI | 1000 Genomes on GRCh38,1000 Genomes 30x on GRCh38,1000 Genomes phase 3 release,1000 Genomes phase 1 release,Geuvadis |
| **NA20589** | female | SAME122865 | TSI | Toscani | EUR | European Ancestry | TSI | 1000 Genomes on GRCh38,1000 Genomes 30x on GRCh38,1000 Genomes phase 3 release,1000 Genomes phase 1 release,Geuvadis |
| **NA21141** | female | SAME125205 | GIH | Gujarati | SAS | South Asian Ancestry | GIH | 1000 Genomes on GRCh38,1000 Genomes 30x on GRCh38,1000 Genomes phase 3 release |
| **NA20755** | male | SAME124174 | TSI | Toscani | EUR | European Ancestry | TSI | 1000 Genomes on GRCh38,1000 Genomes 30x on GRCh38,1000 Genomes phase 3 release,1000 Genomes phase 1 release |
| **NA19784** | male | SAME123151 | MXL | Mexican Ancestry | AMR | American Ancestry | MXL | 1000 Genomes 30x on GRCh38,1000 Genomes phase 3 release |
| **NA19789** | male | SAME123152 | MXL | Mexican Ancestry | AMR | American Ancestry | MXL | 1000 Genomes on GRCh38,1000 Genomes 30x on GRCh38,1000 Genomes phase 3 release,1000 Genomes phase 1 release |
| **NA20321** | female | SAME123996 | ASW | African Ancestry SW | AFR | African Ancestry | ASW | 1000 Genomes on GRCh38,1000 Genomes 30x on GRCh38,1000 Genomes phase 3 release |
| **NA19796** | male | SAME123330 | MXL | Mexican Ancestry | AMR | American Ancestry | MXL | 1000 Genomes 30x on GRCh38,1000 Genomes phase 3 release |
| **NA19828** | male | SAME125246 | ASW | African Ancestry SW | AFR | African Ancestry | ASW | 1000 Genomes 30x on GRCh38,1000 Genomes phase 3 release |
| **NA20340** | male | SAME124400 | ASW | African Ancestry SW | AFR | African Ancestry | ASW | 1000 Genomes on GRCh38,1000 Genomes 30x on GRCh38,1000 Genomes phase 3 release,1000 Genomes phase 1 release |
| **NA19835** | female | SAME125029 | ASW | African Ancestry SW | AFR | African Ancestry | ASW | 1000 Genomes on GRCh38,1000 Genomes 30x on GRCh38,1000 Genomes phase 3 release,1000 Genomes phase 1 release |
| **NA19900** | male | SAME125050 | ASW | African Ancestry SW | AFR | African Ancestry | ASW | 1000 Genomes on GRCh38,1000 Genomes 30x on GRCh38,1000 Genomes phase 3 release,1000 Genomes phase 1 release |
| **NA20357** | female | SAME124599 | ASW | African Ancestry SW | AFR | African Ancestry | ASW | 1000 Genomes on GRCh38,1000 Genomes 30x on GRCh38,1000 Genomes phase 3 release,1000 Genomes phase 1 release |
| **NA20895** | male | SAME123666 | GIH | Gujarati | SAS | South Asian Ancestry | GIH | 1000 Genomes on GRCh38,1000 Genomes 30x on GRCh38,1000 Genomes phase 3 release |
| **NA19917** | female | SAME125272 | ASW | African Ancestry SW | AFR | African Ancestry | ASW | 1000 Genomes on GRCh38,1000 Genomes 30x on GRCh38,1000 Genomes phase 3 release,1000 Genomes phase 1 release |
| **NA20903** | male | SAME122921 | GIH | Gujarati | SAS | South Asian Ancestry | GIH | 1000 Genomes on GRCh38,1000 Genomes 30x on GRCh38,1000 Genomes phase 3 release |
| **NA19924** | female | SAME124714 | ASW | African Ancestry SW | AFR | African Ancestry | ASW | 1000 Genomes 30x on GRCh38,1000 Genomes phase 3 release |
| **NA20908** | female | SAME123777 | GIH | Gujarati | SAS | South Asian Ancestry | GIH | 1000 Genomes on GRCh38,1000 Genomes 30x on GRCh38,1000 Genomes phase 3 release |
| **NA20762** | male | SAME124744 | TSI | Toscani | EUR | European Ancestry | TSI | 1000 Genomes on GRCh38,1000 Genomes 30x on GRCh38,1000 Genomes phase 3 release |
| **NA20910** | female | SAME123100 | GIH | Gujarati | SAS | South Asian Ancestry | GIH | 1000 Genomes on GRCh38,1000 Genomes 30x on GRCh38,1000 Genomes phase 3 release |
| **NA20767** | male | SAME124369 | TSI | Toscani | EUR | European Ancestry | TSI | 1000 Genomes on GRCh38,1000 Genomes 30x on GRCh38,1000 Genomes phase 3 release |
| **NA21088** | female | SAME123143 | GIH | Gujarati | SAS | South Asian Ancestry | GIH | 1000 Genomes on GRCh38,1000 Genomes 30x on GRCh38,1000 Genomes phase 3 release |
| **NA20774** | female | SAME124572 | TSI | Toscani | EUR | European Ancestry | TSI | 1000 Genomes on GRCh38,1000 Genomes 30x on GRCh38,1000 Genomes phase 3 release,1000 Genomes phase 1 release,Geuvadis |
| **NA20852** | male | SAME123244 | GIH | Gujarati | SAS | South Asian Ancestry | GIH | 1000 Genomes on GRCh38,1000 Genomes 30x on GRCh38,1000 Genomes phase 3 release |
| **NA20786** | female | SAME123610 | TSI | Toscani | EUR | European Ancestry | TSI | 1000 Genomes on GRCh38,1000 Genomes 30x on GRCh38,1000 Genomes phase 3 release,1000 Genomes phase 1 release,Geuvadis |
| **NA20864** | male | SAME123944 | GIH | Gujarati | SAS | South Asian Ancestry | GIH | 1000 Genomes on GRCh38,1000 Genomes 30x on GRCh38,1000 Genomes phase 3 release |
| **NA20798** | male | SAME123821 | TSI | Toscani | EUR | European Ancestry | TSI | 1000 Genomes on GRCh38,1000 Genomes 30x on GRCh38,1000 Genomes phase 3 release,1000 Genomes phase 1 release,Geuvadis |
| **NA20869** | female | SAME123801 | GIH | Gujarati | SAS | South Asian Ancestry | GIH | 1000 Genomes on GRCh38,1000 Genomes 30x on GRCh38,1000 Genomes phase 3 release |
| **NA20801** | male | SAME125365 | TSI | Toscani | EUR | European Ancestry | TSI | 1000 Genomes on GRCh38,1000 Genomes 30x on GRCh38,1000 Genomes phase 3 release,1000 Genomes phase 1 release,Geuvadis |
| **NA20806** | male | SAME125362 | TSI | Toscani | EUR | European Ancestry | TSI | 1000 Genomes on GRCh38,1000 Genomes 30x on GRCh38,1000 Genomes phase 3 release,1000 Genomes phase 1 release,Geuvadis |
| **NA21108** | female | SAME124442 | GIH | Gujarati | SAS | South Asian Ancestry | GIH | 1000 Genomes on GRCh38,1000 Genomes 30x on GRCh38,1000 Genomes phase 3 release |
| **NA20813** | female | SAME125189 | TSI | Toscani | EUR | European Ancestry | TSI | 1000 Genomes on GRCh38,1000 Genomes 30x on GRCh38,1000 Genomes phase 3 release,1000 Genomes phase 1 release,Geuvadis |
| **NA21110** | female | SAME124180 | GIH | Gujarati | SAS | South Asian Ancestry | GIH | 1000 Genomes on GRCh38,1000 Genomes 30x on GRCh38,1000 Genomes phase 3 release |
| **NA20818** | female | SAME125184 | TSI | Toscani | EUR | European Ancestry | TSI | 1000 Genomes on GRCh38,1000 Genomes 30x on GRCh38,1000 Genomes phase 3 release,1000 Genomes phase 1 release |
| **NA21115** | male | SAME124183 | GIH | Gujarati | SAS | South Asian Ancestry | GIH | 1000 Genomes on GRCh38,1000 Genomes 30x on GRCh38,1000 Genomes phase 3 release |
| **NA20832** | female | SAME122894 | TSI | Toscani | EUR | European Ancestry | TSI | 1000 Genomes on GRCh38,1000 Genomes 30x on GRCh38,1000 Genomes phase 3 release |
| **NA21122** | female | SAME124030 | GIH | Gujarati | SAS | South Asian Ancestry | GIH | 1000 Genomes on GRCh38,1000 Genomes 30x on GRCh38,1000 Genomes phase 3 release |
| **NA20128** | female | SAME122887 | ASW | African Ancestry SW | AFR | African Ancestry | ASW | 1000 Genomes 30x on GRCh38,1000 Genomes phase 3 release |
| **NA21127** | male | SAME124027 | GIH | Gujarati | SAS | South Asian Ancestry | GIH | 1000 Genomes on GRCh38,1000 Genomes 30x on GRCh38,1000 Genomes phase 3 release |
| **NA20274** | female | SAME123615 | ASW | African Ancestry SW | AFR | African Ancestry | ASW | 1000 Genomes on GRCh38,1000 Genomes 30x on GRCh38,1000 Genomes phase 3 release |
| **NA20876** | female | SAME123599 | GIH | Gujarati | SAS | South Asian Ancestry | GIH | 1000 Genomes on GRCh38,1000 Genomes 30x on GRCh38,1000 Genomes phase 3 release |
| **NA20279** | male | SAME123616 | ASW | African Ancestry SW | AFR | African Ancestry | ASW | 1000 Genomes 30x on GRCh38,1000 Genomes phase 3 release |
| **NA20281** | male | SAME124584 | ASW | African Ancestry SW | AFR | African Ancestry | ASW | 1000 Genomes on GRCh38,1000 Genomes 30x on GRCh38,1000 Genomes phase 3 release,1000 Genomes phase 1 release |
| **NA20888** | female | SAME125258 | GIH | Gujarati | SAS | South Asian Ancestry | GIH | 1000 Genomes on GRCh38,1000 Genomes 30x on GRCh38,1000 Genomes phase 3 release |
| **NA20298** | male | SAME124382 | ASW | African Ancestry SW | AFR | African Ancestry | ASW | 1000 Genomes on GRCh38,1000 Genomes 30x on GRCh38,1000 Genomes phase 3 release,1000 Genomes phase 1 release |
| **NA20890** | male | SAME123671 | GIH | Gujarati | SAS | South Asian Ancestry | GIH | 1000 Genomes on GRCh38,1000 Genomes 30x on GRCh38,1000 Genomes phase 3 release |
| **NA21090** | male | SAME123320 | GIH | Gujarati | SAS | South Asian Ancestry | GIH | 1000 Genomes on GRCh38,1000 Genomes 30x on GRCh38,1000 Genomes phase 3 release |
| **NA21095** | male | SAME123318 | GIH | Gujarati | SAS | South Asian Ancestry | GIH | 1000 Genomes on GRCh38,1000 Genomes 30x on GRCh38,1000 Genomes phase 3 release |
| **NA20318** | male | SAME123802 | ASW | African Ancestry SW | AFR | African Ancestry | ASW | 1000 Genomes on GRCh38,1000 Genomes 30x on GRCh38,1000 Genomes phase 3 release |
| **NA21103** | female | SAME124439 | GIH | Gujarati | SAS | South Asian Ancestry | GIH | 1000 Genomes on GRCh38,1000 Genomes 30x on GRCh38,1000 Genomes phase 3 release |
| **NA20320** | female | SAME123997 | ASW | African Ancestry SW | AFR | African Ancestry | ASW | 1000 Genomes on GRCh38,1000 Genomes 30x on GRCh38,1000 Genomes phase 3 release |
| **NA20332** | female | SAME124198 | ASW | African Ancestry SW | AFR | African Ancestry | ASW | 1000 Genomes on GRCh38,1000 Genomes 30x on GRCh38,1000 Genomes phase 3 release,1000 Genomes phase 1 release |
| **NA20351** | male | SAME124596 | ASW | African Ancestry SW | AFR | African Ancestry | ASW | 1000 Genomes on GRCh38,1000 Genomes 30x on GRCh38,1000 Genomes phase 3 release,1000 Genomes phase 1 release |
| **NA20356** | male | SAME124598 | ASW | African Ancestry SW | AFR | African Ancestry | ASW | 1000 Genomes on GRCh38,1000 Genomes 30x on GRCh38,1000 Genomes phase 3 release,1000 Genomes phase 1 release |
| **NA19741** | male | SAME124718 | MXL | Mexican Ancestry | AMR | American Ancestry | MXL | 1000 Genomes on GRCh38,1000 Genomes 30x on GRCh38,1000 Genomes phase 3 release,1000 Genomes phase 1 release |
| **NA19746** | female | SAME123502 | MXL | Mexican Ancestry | AMR | American Ancestry | MXL | 1000 Genomes on GRCh38,1000 Genomes 30x on GRCh38,1000 Genomes phase 3 release,1000 Genomes phase 1 release |
| **NA19758** | female | SAME123706 | MXL | Mexican Ancestry | AMR | American Ancestry | MXL | 1000 Genomes on GRCh38,1000 Genomes 30x on GRCh38,1000 Genomes phase 3 release,1000 Genomes phase 1 release |
| **NA19760** | female | SAME123895 | MXL | Mexican Ancestry | AMR | American Ancestry | MXL | 1000 Genomes 30x on GRCh38,1000 Genomes phase 3 release |
| **NA19772** | male | SAME124066 | MXL | Mexican Ancestry | AMR | American Ancestry | MXL | 1000 Genomes 30x on GRCh38,1000 Genomes phase 3 release |
| **NA19777** | male | SAME124063 | MXL | Mexican Ancestry | AMR | American Ancestry | MXL | 1000 Genomes on GRCh38,1000 Genomes 30x on GRCh38,1000 Genomes phase 3 release,1000 Genomes phase 1 release |
| **NA20849** | female | SAME123433 | GIH | Gujarati | SAS | South Asian Ancestry | GIH | 1000 Genomes on GRCh38,1000 Genomes 30x on GRCh38,1000 Genomes phase 3 release |
| **NA20851** | female | SAME123245 | GIH | Gujarati | SAS | South Asian Ancestry | GIH | 1000 Genomes on GRCh38,1000 Genomes 30x on GRCh38,1000 Genomes phase 3 release |
| **NA20856** | female | SAME123247 | GIH | Gujarati | SAS | South Asian Ancestry | GIH | 1000 Genomes on GRCh38,1000 Genomes 30x on GRCh38,1000 Genomes phase 3 release |
| **NA20863** | male | SAME123805 | GIH | Gujarati | SAS | South Asian Ancestry | GIH | 1000 Genomes on GRCh38,1000 Genomes 30x on GRCh38,1000 Genomes phase 3 release |
| **NA20868** | female | SAME123800 | GIH | Gujarati | SAS | South Asian Ancestry | GIH | 1000 Genomes on GRCh38,1000 Genomes 30x on GRCh38,1000 Genomes phase 3 release |
| **NA20870** | male | SAME122839 | GIH | Gujarati | SAS | South Asian Ancestry | GIH | 1000 Genomes on GRCh38,1000 Genomes 30x on GRCh38,1000 Genomes phase 3 release |
| **NA20894** | female | SAME123667 | GIH | Gujarati | SAS | South Asian Ancestry | GIH | 1000 Genomes on GRCh38,1000 Genomes 30x on GRCh38,1000 Genomes phase 3 release |
| **NA20899** | female | SAME123140 | GIH | Gujarati | SAS | South Asian Ancestry | GIH | 1000 Genomes on GRCh38,1000 Genomes 30x on GRCh38,1000 Genomes phase 3 release |
| **NA20902** | female | SAME122922 | GIH | Gujarati | SAS | South Asian Ancestry | GIH | 1000 Genomes on GRCh38,1000 Genomes 30x on GRCh38,1000 Genomes phase 3 release |
| **NA21087** | male | SAME123138 | GIH | Gujarati | SAS | South Asian Ancestry | GIH | 1000 Genomes on GRCh38,1000 Genomes 30x on GRCh38,1000 Genomes phase 3 release |
| **NA20875** | female | SAME123596 | GIH | Gujarati | SAS | South Asian Ancestry | GIH | 1000 Genomes on GRCh38,1000 Genomes 30x on GRCh38,1000 Genomes phase 3 release |
| **NA20882** | female | SAME123882 | GIH | Gujarati | SAS | South Asian Ancestry | GIH | 1000 Genomes on GRCh38,1000 Genomes 30x on GRCh38,1000 Genomes phase 3 release |
| **NA20887** | male | SAME123879 | GIH | Gujarati | SAS | South Asian Ancestry | GIH | 1000 Genomes on GRCh38,1000 Genomes 30x on GRCh38,1000 Genomes phase 3 release |
| **NA21133** | male | SAME123831 | GIH | Gujarati | SAS | South Asian Ancestry | GIH | 1000 Genomes on GRCh38,1000 Genomes 30x on GRCh38,1000 Genomes phase 3 release |
| **NA21114** | male | SAME124184 | GIH | Gujarati | SAS | South Asian Ancestry | GIH | 1000 Genomes on GRCh38,1000 Genomes 30x on GRCh38,1000 Genomes phase 3 release |
| **NA21119** | male | SAME123584 | GIH | Gujarati | SAS | South Asian Ancestry | GIH | 1000 Genomes on GRCh38,1000 Genomes 30x on GRCh38,1000 Genomes phase 3 release |
| **NA21126** | male | SAME124026 | GIH | Gujarati | SAS | South Asian Ancestry | GIH | 1000 Genomes on GRCh38,1000 Genomes 30x on GRCh38,1000 Genomes phase 3 release |
| **NA21094** | male | SAME123317 | GIH | Gujarati | SAS | South Asian Ancestry | GIH | 1000 Genomes on GRCh38,1000 Genomes 30x on GRCh38,1000 Genomes phase 3 release |
| **NA21099** | male | SAME123325 | GIH | Gujarati | SAS | South Asian Ancestry | GIH | 1000 Genomes on GRCh38,1000 Genomes 30x on GRCh38,1000 Genomes phase 3 release |
| **NA21102** | female | SAME124438 | GIH | Gujarati | SAS | South Asian Ancestry | GIH | 1000 Genomes on GRCh38,1000 Genomes 30x on GRCh38,1000 Genomes phase 3 release |
| **NA21107** | male | SAME124435 | GIH | Gujarati | SAS | South Asian Ancestry | GIH | 1000 Genomes on GRCh38,1000 Genomes 30x on GRCh38,1000 Genomes phase 3 release |
| **HG00157** | male | SAME124590 | GBR | British | EUR | European Ancestry | GBR | 1000 Genomes on GRCh38,1000 Genomes 30x on GRCh38,1000 Genomes phase 3 release,1000 Genomes phase 1 release,Geuvadis |
| **HG00171** | female | SAME124961 | FIN | Finnish | EUR | European Ancestry | FIN | 1000 Genomes on GRCh38,1000 Genomes 30x on GRCh38,Human Genome Structural Variation Consortium, Phase 2,1000 Genomes phase 3 release,1000 Genomes ph |
| **HG00176** | female | SAME124956 | FIN | Finnish | EUR | European Ancestry | FIN | 1000 Genomes on GRCh38,1000 Genomes 30x on GRCh38,1000 Genomes phase 3 release,1000 Genomes phase 1 release,Geuvadis |
| **HG00183** | male | SAME123642 | FIN | Finnish | EUR | European Ancestry | FIN | 1000 Genomes on GRCh38,1000 Genomes 30x on GRCh38,1000 Genomes phase 3 release,1000 Genomes phase 1 release,Geuvadis |
| **HG00188** | male | SAME123641 | FIN | Finnish | EUR | European Ancestry | FIN | 1000 Genomes on GRCh38,1000 Genomes 30x on GRCh38,1000 Genomes phase 3 release,1000 Genomes phase 1 release,Geuvadis |
| **HG00190** | male | SAME123856 | FIN | Finnish,Finnish | EUR | European Ancestry,West Eurasia (SGDP) | FIN,FinnishSGDP | 1000 Genomes on GRCh38,Simons Genome Diversity Project,1000 Genomes 30x on GRCh38,1000 Genomes phase 3 release,1000 Genomes phase 1 release |
| **HG00234** | male | SAME124123 | GBR | British | EUR | European Ancestry | GBR | 1000 Genomes on GRCh38,1000 Genomes 30x on GRCh38,1000 Genomes phase 3 release,1000 Genomes phase 1 release,Geuvadis |
| **HG00239** | female | SAME124130 | GBR | British | EUR | European Ancestry | GBR | 1000 Genomes on GRCh38,1000 Genomes 30x on GRCh38,1000 Genomes phase 3 release,1000 Genomes phase 1 release,Geuvadis |
| **HG00246** | male | SAME123219 | GBR | British | EUR | European Ancestry | GBR | 1000 Genomes on GRCh38,1000 Genomes 30x on GRCh38,1000 Genomes phase 3 release,1000 Genomes phase 1 release,Geuvadis |
| **HG00253** | female | SAME123051 | GBR | British | EUR | European Ancestry | GBR | 1000 Genomes on GRCh38,1000 Genomes 30x on GRCh38,1000 Genomes phase 3 release,1000 Genomes phase 1 release,Geuvadis |
| **HG00258** | female | SAME123057 | GBR | British | EUR | European Ancestry | GBR | 1000 Genomes on GRCh38,1000 Genomes 30x on GRCh38,1000 Genomes phase 3 release,1000 Genomes phase 1 release,Geuvadis |
| **HG00260** | male | SAME123582 | GBR | British | EUR | European Ancestry | GBR | 1000 Genomes on GRCh38,1000 Genomes 30x on GRCh38,1000 Genomes phase 3 release,1000 Genomes phase 1 release,Geuvadis |
| **HG00265** | male | SAME123579 | GBR | British | EUR | European Ancestry | GBR | 1000 Genomes on GRCh38,1000 Genomes 30x on GRCh38,1000 Genomes phase 3 release,1000 Genomes phase 1 release,Geuvadis |
| **HG00272** | female | SAME123420 | FIN | Finnish | EUR | European Ancestry | FIN | 1000 Genomes on GRCh38,1000 Genomes 30x on GRCh38,1000 Genomes phase 3 release,1000 Genomes phase 1 release,Geuvadis |
| **HG00277** | male | SAME123423 | FIN | Finnish | EUR | European Ancestry | FIN | 1000 Genomes on GRCh38,1000 Genomes 30x on GRCh38,1000 Genomes phase 3 release,1000 Genomes phase 1 release,Geuvadis |
| **HG00284** | male | SAME122835 | FIN | Finnish | EUR | European Ancestry | FIN | 1000 Genomes on GRCh38,1000 Genomes 30x on GRCh38,1000 Genomes phase 3 release,1000 Genomes phase 1 release,Geuvadis |
| **HG00479** | female | SAME123334 | CHS | Southern Han Chinese | EAS | East Asian Ancestry | CHS | 1000 Genomes on GRCh38,1000 Genomes 30x on GRCh38,1000 Genomes phase 3 release,1000 Genomes phase 1 release,90 Han Chinese high coverage genomes |
| **HG00501** | female | SAME123362 | CHS | Southern Han Chinese | EAS | East Asian Ancestry | CHS | 1000 Genomes on GRCh38,1000 Genomes 30x on GRCh38,1000 Genomes phase 3 release,1000 Genomes phase 1 release |
| **HG00513** | female | SAME123526 | CHS | Southern Han Chinese | EAS | East Asian Ancestry | CHS | 1000 Genomes on GRCh38,1000 Genomes 30x on GRCh38,Human Genome Structural Variation Consortium, Phase 2,The Human Genome Structural Variation Consortiu |
| **HG00525** | female | SAME123242 | CHS | Southern Han Chinese | EAS | East Asian Ancestry | CHS | 1000 Genomes on GRCh38,1000 Genomes 30x on GRCh38,1000 Genomes phase 3 release,1000 Genomes phase 1 release,90 Han Chinese high coverage genomes |
| **HG00532** | male | SAME125268 | CHS | Southern Han Chinese | EAS | East Asian Ancestry | CHS | 1000 Genomes 30x on GRCh38,1000 Genomes phase 3 release |
| **HG00537** | female | SAME124345 | CHS | Southern Han Chinese | EAS | East Asian Ancestry | CHS | 1000 Genomes on GRCh38,1000 Genomes 30x on GRCh38,1000 Genomes phase 3 release,1000 Genomes phase 1 release,90 Han Chinese high coverage genomes |
| **HG00109** | male | SAME125160 | GBR | British | EUR | European Ancestry | GBR | 1000 Genomes on GRCh38,1000 Genomes 30x on GRCh38,1000 Genomes phase 3 release,1000 Genomes phase 1 release,Geuvadis |
| **HG00544** | female | SAME124086 | CHS | Southern Han Chinese | EAS | East Asian Ancestry | CHS | 1000 Genomes 30x on GRCh38,1000 Genomes phase 3 release |
| **HG00111** | female | SAME125340 | GBR | British | EUR | European Ancestry | GBR | 1000 Genomes on GRCh38,1000 Genomes 30x on GRCh38,1000 Genomes phase 3 release,1000 Genomes phase 1 release,Geuvadis |
| **HG00551** | female | SAME124252 | PUR | Puerto Rican | AMR | American Ancestry | PUR | 1000 Genomes on GRCh38,1000 Genomes 30x on GRCh38,1000 Genomes phase 3 release,1000 Genomes phase 1 release |
| **HG00116** | male | SAME125345 | GBR | British | EUR | European Ancestry | GBR | 1000 Genomes on GRCh38,1000 Genomes 30x on GRCh38,1000 Genomes phase 3 release,1000 Genomes phase 1 release,Geuvadis |
| **HG00556** | male | SAME123971 | CHS | Southern Han Chinese | EAS | East Asian Ancestry | CHS | 1000 Genomes on GRCh38,1000 Genomes 30x on GRCh38,1000 Genomes phase 3 release,1000 Genomes phase 1 release,90 Han Chinese high coverage genomes |
| **HG00123** | female | SAME122875 | GBR | British | EUR | European Ancestry | GBR | 1000 Genomes on GRCh38,1000 Genomes 30x on GRCh38,1000 Genomes phase 3 release,1000 Genomes phase 1 release,Geuvadis |
| **HG00582** | female | SAME124830 | CHS | Southern Han Chinese | EAS | East Asian Ancestry | CHS | 1000 Genomes 30x on GRCh38,1000 Genomes phase 3 release |

| | | | | | | | | |
|---|---|---|---|---|---|---|---|---|
| **HG00128** | female | SAME122868 | GBR | British,English | EUR | European Ancestry,West Eurasia (SGDP) | GBR,EnglishSGDP | 1000 Genomes on GRCh38,Simons Genome Diversity Project,1000 Genomes 30x on GRCh38,1000 Genomes phase 3 release,1000 Genomes phase 1 release,Geuvadis |
| **HG00099** | female | SAME123271 | GBR | British | EUR | European Ancestry | GBR | 1000 Genomes on GRCh38,1000 Genomes 30x on GRCh38,1000 Genomes phase 3 release,1000 Genomes phase 1 release,Geuvadis |
| **HG00130** | female | SAME123063 | GBR | British | EUR | European Ancestry | GBR | 1000 Genomes on GRCh38,1000 Genomes 30x on GRCh38,1000 Genomes phase 3 release,1000 Genomes phase 1 release,Geuvadis |
| **HG00102** | female | SAME123945 | GBR | British | EUR | European Ancestry | GBR | 1000 Genomes on GRCh38,1000 Genomes 30x on GRCh38,1000 Genomes phase 3 release,1000 Genomes phase 1 release,Geuvadis |
| **HG00107** | male | SAME123947 | GBR | British | EUR | European Ancestry | GBR | 1000 Genomes on GRCh38,1000 Genomes 30x on GRCh38,1000 Genomes phase 3 release,1000 Genomes phase 1 release,Geuvadis |
| **HG00142** | male | SAME124818 | GBR | British | EUR | European Ancestry | GBR | 1000 Genomes on GRCh38,1000 Genomes 30x on GRCh38,1000 Genomes phase 3 release,1000 Genomes phase 1 release,Geuvadis |
| **HG00114** | male | SAME125343 | GBR | British | EUR | European Ancestry | GBR | 1000 Genomes on GRCh38,1000 Genomes 30x on GRCh38,1000 Genomes phase 3 release,1000 Genomes phase 1 release,Geuvadis |
| **HG00119** | male | SAME125348 | GBR | British | EUR | European Ancestry | GBR | 1000 Genomes on GRCh38,1000 Genomes 30x on GRCh38,1000 Genomes phase 3 release,1000 Genomes phase 1 release,Geuvadis |
| **HG00154** | female | SAME124587 | GBR | British | EUR | European Ancestry | GBR | 1000 Genomes on GRCh38,1000 Genomes 30x on GRCh38,1000 Genomes phase 3 release,1000 Genomes phase 1 release,Geuvadis |
| **HG00121** | female | SAME122873 | GBR | British | EUR | European Ancestry | GBR | 1000 Genomes on GRCh38,1000 Genomes 30x on GRCh38,1000 Genomes phase 3 release,1000 Genomes phase 1 release,Geuvadis |
| **HG00159** | male | SAME124586 | GBR | British | EUR | European Ancestry | GBR | 1000 Genomes on GRCh38,1000 Genomes 30x on GRCh38,1000 Genomes phase 3 release,1000 Genomes phase 1 release,Geuvadis |
| **HG00126** | male | SAME122872 | GBR | British,English | EUR | European Ancestry,West Eurasia (SGDP) | GBR,EnglishSGDP | 1000 Genomes on GRCh38,Simons Genome Diversity Project,1000 Genomes 30x on GRCh38,1000 Genomes phase 3 release,1000 Genomes phase 1 release,Geuvadis |
| **HG00173** | female | SAME124960 | FIN | Finnish | EUR | European Ancestry | FIN | 1000 Genomes on GRCh38,1000 Genomes 30x on GRCh38,1000 Genomes phase 3 release,1000 Genomes phase 1 release,Geuvadis |
| **HG00133** | female | SAME123062 | GBR | British | EUR | European Ancestry | GBR | 1000 Genomes on GRCh38,1000 Genomes 30x on GRCh38,1000 Genomes phase 3 release,1000 Genomes phase 1 release,Geuvadis |
| **HG00178** | female | SAME124964 | FIN | Finnish | EUR | European Ancestry | FIN | 1000 Genomes on GRCh38,1000 Genomes 30x on GRCh38,1000 Genomes phase 3 release,1000 Genomes phase 1 release,Geuvadis |
| **HG00138** | male | SAME123059 | GBR | British | EUR | European Ancestry | GBR | 1000 Genomes on GRCh38,1000 Genomes 30x on GRCh38,1000 Genomes phase 3 release,1000 Genomes phase 1 release,Geuvadis |
| **HG00180** | female | SAME123645 | FIN | Finnish | EUR | European Ancestry | FIN | 1000 Genomes on GRCh38,1000 Genomes 30x on GRCh38,1000 Genomes phase 3 release,1000 Genomes phase 1 release,Geuvadis |
| **HG00140** | male | SAME124396 | GBR | British | EUR | European Ancestry | GBR | 1000 Genomes on GRCh38,1000 Genomes 30x on GRCh38,1000 Genomes phase 3 release,1000 Genomes phase 1 release |
| **HG00185** | male | SAME123648 | FIN | Finnish | EUR | European Ancestry | FIN | 1000 Genomes on GRCh38,1000 Genomes 30x on GRCh38,1000 Genomes phase 3 release,1000 Genomes phase 1 release,Geuvadis |
| **HG00145** | male | SAME124391 | GBR | British | EUR | European Ancestry | GBR | 1000 Genomes on GRCh38,1000 Genomes 30x on GRCh38,1000 Genomes phase 3 release,1000 Genomes phase 1 release,Geuvadis |
| **HG00231** | female | SAME124126 | GBR | British | EUR | European Ancestry | GBR | 1000 Genomes on GRCh38,1000 Genomes 30x on GRCh38,1000 Genomes phase 3 release,1000 Genomes phase 1 release,Geuvadis |
| **HG00304** | female | SAME1840118 | FIN | Finnish | EUR | European Ancestry | FIN | 1000 Genomes on GRCh38,1000 Genomes 30x on GRCh38,1000 Genomes phase 3 release |
| **HG00236** | female | SAME123273 | GBR | British | EUR | European Ancestry | GBR | 1000 Genomes on GRCh38,1000 Genomes 30x on GRCh38,1000 Genomes phase 3 release,1000 Genomes phase 1 release,Geuvadis |
| **HG00309** | female | SAME124160 | FIN | Finnish | EUR | European Ancestry | FIN | 1000 Genomes on GRCh38,1000 Genomes 30x on GRCh38,1000 Genomes phase 3 release,1000 Genomes phase 1 release,Geuvadis |
| **HG00243** | male | SAME123224 | GBR | British | EUR | European Ancestry | GBR | 1000 Genomes on GRCh38,1000 Genomes 30x on GRCh38,1000 Genomes phase 3 release,1000 Genomes phase 1 release,Geuvadis |
| **HG00311** | male | SAME124339 | FIN | Finnish | EUR | European Ancestry | FIN | 1000 Genomes on GRCh38,1000 Genomes 30x on GRCh38,1000 Genomes phase 3 release,1000 Genomes phase 1 release,Geuvadis |
| **HG00318** | female | SAME124333 | FIN | Finnish | EUR | European Ancestry | FIN | 1000 Genomes on GRCh38,1000 Genomes 30x on GRCh38,1000 Genomes phase 3 release,1000 Genomes phase 1 release |
| **HG00323** | female | SAME123795 | FIN | Finnish | EUR | European Ancestry | FIN | 1000 Genomes on GRCh38,1000 Genomes 30x on GRCh38,1000 Genomes phase 3 release,1000 Genomes phase 1 release,Geuvadis |
| **HG00320** | female | SAME123797 | FIN | Finnish | EUR | European Ancestry | FIN | 1000 Genomes on GRCh38,1000 Genomes 30x on GRCh38,1000 Genomes phase 3 release,1000 Genomes phase 1 release,Geuvadis |
| **HG00328** | female | SAME123799 | FIN | Finnish | EUR | European Ancestry | FIN | 1000 Genomes on GRCh38,1000 Genomes 30x on GRCh38,1000 Genomes phase 3 release,1000 Genomes phase 1 release,Geuvadis |
| **HG00325** | male | SAME123793 | FIN | Finnish | EUR | European Ancestry | FIN | 1000 Genomes on GRCh38,1000 Genomes 30x on GRCh38,1000 Genomes phase 3 release,1000 Genomes phase 1 release,Geuvadis |
| **HG00330** | female | SAME123985 | FIN | Finnish | EUR | European Ancestry | FIN | 1000 Genomes on GRCh38,1000 Genomes 30x on GRCh38,1000 Genomes phase 3 release,1000 Genomes phase 1 release,Geuvadis |
| **HG00332** | female | SAME123987 | FIN | Finnish | EUR | European Ancestry | FIN | 1000 Genomes on GRCh38,1000 Genomes 30x on GRCh38,1000 Genomes phase 3 release,1000 Genomes phase 1 release,Geuvadis |
| **HG00335** | male | SAME123982 | FIN | Finnish | EUR | European Ancestry | FIN | 1000 Genomes on GRCh38,1000 Genomes 30x on GRCh38,1000 Genomes phase 3 release,1000 Genomes phase 1 release,Geuvadis |
| **HG00337** | female | SAME123984 | FIN | Finnish | EUR | European Ancestry | FIN | 1000 Genomes on GRCh38,1000 Genomes 30x on GRCh38,1000 Genomes phase 3 release,1000 Genomes phase 1 release,Geuvadis |
| **HG00342** | male | SAME124924 | FIN | Finnish | EUR | European Ancestry | FIN | 1000 Genomes on GRCh38,1000 Genomes 30x on GRCh38,1000 Genomes phase 3 release,1000 Genomes phase 1 release,Geuvadis |
| **HG00344** | female | SAME125255 | FIN | Finnish | EUR | European Ancestry | FIN | 1000 Genomes on GRCh38,1000 Genomes 30x on GRCh38,1000 Genomes phase 3 release,1000 Genomes phase 1 release,Geuvadis |
| **HG00349** | female | SAME124925 | FIN | Finnish | EUR | European Ancestry | FIN | 1000 Genomes on GRCh38,1000 Genomes 30x on GRCh38,1000 Genomes phase 3 release,1000 Genomes phase 1 release,Geuvadis |
| **HG00361** | female | SAME124538 | FIN | Finnish | EUR | European Ancestry | FIN | 1000 Genomes on GRCh38,1000 Genomes 30x on GRCh38,1000 Genomes phase 3 release,1000 Genomes phase 1 release,Geuvadis |
| **HG00351** | male | SAME125122 | FIN | Finnish | EUR | European Ancestry | FIN | 1000 Genomes on GRCh38,1000 Genomes 30x on GRCh38,1000 Genomes phase 3 release,1000 Genomes phase 1 release,Geuvadis |
| **HG00366** | male | SAME124541 | FIN | Finnish | EUR | European Ancestry | FIN | 1000 Genomes on GRCh38,1000 Genomes 30x on GRCh38,1000 Genomes phase 3 release,1000 Genomes phase 1 release,Geuvadis |
| **HG00356** | female | SAME125123 | FIN | Finnish | EUR | European Ancestry | FIN | 1000 Genomes on GRCh38,1000 Genomes 30x on GRCh38,1000 Genomes phase 3 release,1000 Genomes phase 1 release,Geuvadis |
| **HG00373** | female | SAME124749 | FIN | Finnish | EUR | European Ancestry | FIN | 1000 Genomes on GRCh38,1000 Genomes 30x on GRCh38,1000 Genomes phase 3 release,1000 Genomes phase 1 release,Geuvadis |
| **HG00368** | female | SAME124535 | FIN | Finnish | EUR | European Ancestry | FIN | 1000 Genomes on GRCh38,1000 Genomes 30x on GRCh38,1000 Genomes phase 3 release,1000 Genomes phase 1 release |
| **HG00378** | female | SAME124745 | FIN | Finnish | EUR | European Ancestry | FIN | 1000 Genomes on GRCh38,1000 Genomes 30x on GRCh38,1000 Genomes phase 3 release,1000 Genomes phase 1 release,Geuvadis |
| **HG00375** | male | SAME124750 | FIN | Finnish | EUR | European Ancestry | FIN | 1000 Genomes on GRCh38,1000 Genomes 30x on GRCh38,1000 Genomes phase 3 release,1000 Genomes phase 1 release,Geuvadis |
| **HG00380** | female | SAME123031 | FIN | Finnish | EUR | European Ancestry | FIN | 1000 Genomes on GRCh38,1000 Genomes 30x on GRCh38,1000 Genomes phase 3 release,1000 Genomes phase 1 release,Geuvadis |
| **HG00382** | male | SAME123033 | FIN | Finnish | EUR | European Ancestry | FIN | 1000 Genomes on GRCh38,1000 Genomes 30x on GRCh38,1000 Genomes phase 3 release,1000 Genomes phase 1 release,Geuvadis |
| **HG00594** | male | SAME125005 | CHS | Southern Han Chinese | EAS | East Asian Ancestry | CHS | 1000 Genomes 30x on GRCh38,1000 Genomes phase 3 release |
| **HG00407** | female | SAME125106 | CHS | Southern Han Chinese | EAS | East Asian Ancestry | CHS | 1000 Genomes on GRCh38,1000 Genomes 30x on GRCh38,1000 Genomes phase 3 release,1000 Genomes phase 1 release |
| **HG00599** | female | SAME1839802 | CHS | Southern Han Chinese | EAS | East Asian Ancestry | CHS | 1000 Genomes on GRCh38,1000 Genomes 30x on GRCh38,1000 Genomes phase 3 release |
| **HG00419** | female | SAME122991 | CHS | Southern Han Chinese | EAS | East Asian Ancestry | CHS | 1000 Genomes on GRCh38,1000 Genomes 30x on GRCh38,1000 Genomes phase 3 release,1000 Genomes phase 1 release |
| **HG00607** | male | SAME125373 | CHS | Southern Han Chinese | EAS | East Asian Ancestry | CHS | 1000 Genomes on GRCh38,1000 Genomes 30x on GRCh38,1000 Genomes phase 3 release,1000 Genomes phase 1 release |
| **HG00421** | male | SAME122808 | CHS | Southern Han Chinese | EAS | East Asian Ancestry | CHS | 1000 Genomes on GRCh38,1000 Genomes 30x on GRCh38,1000 Genomes phase 3 release,1000 Genomes phase 1 release,90 Han Chinese high coverage genomes |
| **HG00614** | female | SAME125195 | CHS | Southern Han Chinese | EAS | East Asian Ancestry | CHS | 1000 Genomes on GRCh38,1000 Genomes 30x on GRCh38,1000 Genomes phase 3 release,1000 Genomes phase 1 release |
| **HG00553** | male | SAME124254 | PUR | Puerto Rican | AMR | American Ancestry | PUR | 1000 Genomes on GRCh38,1000 Genomes 30x on GRCh38,1000 Genomes phase 3 release,1000 Genomes phase 1 release |
| **HG00619** | male | SAME125193 | CHS | Southern Han Chinese | EAS | East Asian Ancestry | CHS | 1000 Genomes on GRCh38,1000 Genomes 30x on GRCh38,1000 Genomes phase 3 release,1000 Genomes phase 1 release,90 Han Chinese high coverage genomes |
| **HG00558** | female | SAME124247 | CHS | Southern Han Chinese | EAS | East Asian Ancestry | CHS | 1000 Genomes 30x on GRCh38,1000 Genomes phase 3 release |
| **HG00621** | male | SAME123082 | CHS | Southern Han Chinese | EAS | East Asian Ancestry | CHS | 1000 Genomes 30x on GRCh38,1000 Genomes phase 3 release |
| **HG00560** | female | SAME124456 | CHS | Southern Han Chinese | EAS | East Asian Ancestry | CHS | 1000 Genomes on GRCh38,1000 Genomes 30x on GRCh38,1000 Genomes phase 3 release,1000 Genomes phase 1 release |
| **HG00626** | female | SAME123078 | CHS | Southern Han Chinese | EAS | East Asian Ancestry | CHS | 1000 Genomes on GRCh38,1000 Genomes 30x on GRCh38,1000 Genomes phase 3 release,1000 Genomes phase 1 release |
| **HG00565** | male | SAME124452 | CHS | Southern Han Chinese | EAS | East Asian Ancestry | CHS | 1000 Genomes on GRCh38,1000 Genomes 30x on GRCh38,1000 Genomes phase 3 release,1000 Genomes phase 1 release |
| **HG00577** | male | SAME123516 | CHS | Southern Han Chinese | EAS | East Asian Ancestry | CHS | 1000 Genomes on GRCh38,1000 Genomes 30x on GRCh38,1000 Genomes phase 3 release,1000 Genomes phase 1 release |
| **HG00405** | female | SAME123159 | CHS | Southern Han Chinese | EAS | East Asian Ancestry | CHS | 1000 Genomes 30x on GRCh38,1000 Genomes phase 3 release |
| **HG00584** | female | SAME124663 | CHS | Southern Han Chinese | EAS | East Asian Ancestry | CHS | 1000 Genomes on GRCh38,1000 Genomes 30x on GRCh38,1000 Genomes phase 3 release,1000 Genomes phase 1 release,90 Han Chinese high coverage genomes |
| **HG00429** | male | SAME122812 | CHS | Southern Han Chinese | EAS | East Asian Ancestry | CHS | 1000 Genomes 30x on GRCh38,1000 Genomes phase 3 release |
| **HG00589** | male | SAME124824 | CHS | Southern Han Chinese | EAS | East Asian Ancestry | CHS | 1000 Genomes on GRCh38,1000 Genomes 30x on GRCh38,1000 Genomes phase 3 release,1000 Genomes phase 1 release |
| **HG00436** | male | SAME125262 | CHS | Southern Han Chinese | EAS | East Asian Ancestry | CHS | 1000 Genomes on GRCh38,1000 Genomes 30x on GRCh38,1000 Genomes phase 3 release,1000 Genomes phase 1 release,90 Han Chinese high coverage genomes |
| **HG00591** | male | SAME125010 | CHS | Southern Han Chinese | EAS | East Asian Ancestry | CHS | 1000 Genomes 30x on GRCh38,1000 Genomes phase 3 release |
| **HG00443** | female | SAME124920 | CHS | Southern Han Chinese | EAS | East Asian Ancestry | CHS | 1000 Genomes on GRCh38,1000 Genomes 30x on GRCh38,1000 Genomes phase 3 release,1000 Genomes phase 1 release,90 Han Chinese high coverage genomes |
| **HG00596** | female | SAME125007 | CHS | Southern Han Chinese | EAS | East Asian Ancestry | CHS | 1000 Genomes on GRCh38,1000 Genomes 30x on GRCh38,1000 Genomes phase 3 release,1000 Genomes phase 1 release |
| **HG00448** | male | SAME124200 | CHS | Southern Han Chinese | EAS | East Asian Ancestry | CHS | 1000 Genomes on GRCh38,1000 Genomes 30x on GRCh38,1000 Genomes phase 3 release,1000 Genomes phase 1 release,90 Han Chinese high coverage genomes |
| **HG00609** | male | SAME125370 | CHS | Southern Han Chinese | EAS | East Asian Ancestry | CHS | 1000 Genomes 30x on GRCh38,1000 Genomes phase 3 release |
| **HG00450** | female | SAME125167 | CHS | Southern Han Chinese | EAS | East Asian Ancestry | CHS | 1000 Genomes 30x on GRCh38,1000 Genomes phase 3 release |
| **HG00611** | female | SAME125198 | CHS | Southern Han Chinese | EAS | East Asian Ancestry | CHS | 1000 Genomes on GRCh38,1000 Genomes 30x on GRCh38,1000 Genomes phase 3 release,1000 Genomes phase 1 release |
| **HG00474** | male | SAME123343 | CHS | Southern Han Chinese | EAS | East Asian Ancestry | CHS | 1000 Genomes 30x on GRCh38,1000 Genomes phase 3 release |
| **HG00638** | female | SAME123652 | PUR | Puerto Rican | AMR | American Ancestry | PUR | 1000 Genomes on GRCh38,1000 Genomes 30x on GRCh38,1000 Genomes phase 3 release,1000 Genomes phase 1 release |
| **HG00250** | female | SAME123054 | GBR | British | EUR | European Ancestry | GBR | 1000 Genomes on GRCh38,1000 Genomes 30x on GRCh38,1000 Genomes phase 3 release,1000 Genomes phase 1 release,Geuvadis |
| **HG00640** | male | SAME123434 | PUR | Puerto Rican | AMR | American Ancestry | PUR | 1000 Genomes on GRCh38,1000 Genomes 30x on GRCh38,1000 Genomes phase 3 release,1000 Genomes phase 1 release |
| **HG00255** | female | SAME123049 | GBR | British | EUR | European Ancestry | GBR | 1000 Genomes on GRCh38,1000 Genomes 30x on GRCh38,1000 Genomes phase 3 release,1000 Genomes phase 1 release,Geuvadis |
| **HG00652** | female | SAME123252 | CHS | Southern Han Chinese | EAS | East Asian Ancestry | CHS | 1000 Genomes 30x on GRCh38,1000 Genomes phase 3 release |
| **HG00262** | female | SAME123580 | GBR | British | EUR | European Ancestry | GBR | 1000 Genomes on GRCh38,1000 Genomes 30x on GRCh38,1000 Genomes phase 3 release,1000 Genomes phase 1 release,Geuvadis |
| **HG00657** | female | SAME123255 | CHS | Southern Han Chinese | EAS | East Asian Ancestry | CHS | 1000 Genomes on GRCh38,1000 Genomes 30x on GRCh38,1000 Genomes phase 3 release,1000 Genomes phase 1 release,90 Han Chinese high coverage genomes |
| **HG00267** | male | SAME123577 | FIN | Finnish | EUR | European Ancestry | FIN | 1000 Genomes on GRCh38,1000 Genomes 30x on GRCh38,1000 Genomes phase 3 release,1000 Genomes phase 1 release,Geuvadis |
| **HG00664** | male | SAME123814 | CHS | Southern Han Chinese | EAS | East Asian Ancestry | CHS | 1000 Genomes 30x on GRCh38,1000 Genomes phase 3 release |
| **HG00274** | female | SAME123422 | FIN | Finnish | EUR | European Ancestry | FIN | 1000 Genomes on GRCh38,1000 Genomes 30x on GRCh38,1000 Genomes phase 3 release,1000 Genomes phase 1 release,Geuvadis |
| **HG00671** | male | SAME123604 | CHS | Southern Han Chinese | EAS | East Asian Ancestry | CHS | 1000 Genomes on GRCh38,1000 Genomes 30x on GRCh38,1000 Genomes phase 3 release,1000 Genomes phase 1 release,90 Han Chinese high coverage genomes |
| **HG00281** | female | SAME122832 | FIN | Finnish | EUR | European Ancestry | FIN | 1000 Genomes on GRCh38,1000 Genomes 30x on GRCh38,1000 Genomes phase 3 release,1000 Genomes phase 1 release,Geuvadis |
| **HG00306** | female | SAME124159 | FIN | Finnish | EUR | European Ancestry | FIN | 1000 Genomes on GRCh38,1000 Genomes 30x on GRCh38,1000 Genomes phase 3 release,1000 Genomes phase 1 release,Geuvadis |
| **HG00683** | male | SAME123892 | CHS | Southern Han Chinese | EAS | East Asian Ancestry | CHS | 1000 Genomes on GRCh38,1000 Genomes 30x on GRCh38,1000 Genomes phase 3 release,1000 Genomes phase 1 release,90 Han Chinese high coverage genomes |
| **HG00313** | female | SAME124337 | FIN | Finnish | EUR | European Ancestry | FIN | 1000 Genomes on GRCh38,1000 Genomes 30x on GRCh38,1000 Genomes phase 3 release,1000 Genomes phase 1 release,Geuvadis |
| **HG00690** | female | SAME123686 | CHS | Southern Han Chinese | EAS | East Asian Ancestry | CHS | 1000 Genomes on GRCh38,1000 Genomes 30x on GRCh38,1000 Genomes phase 3 release,1000 Genomes phase 1 release,90 Han Chinese high coverage genomes |
| **HG00438** | female | SAME125267 | CHS | Southern Han Chinese | EAS | East Asian Ancestry | CHS | 1000 Genomes 30x on GRCh38,1000 Genomes phase 3 release |
| **HG01326** | female | SAME1839598 | PUR | Puerto Rican | AMR | American Ancestry | PUR | 1000 Genomes on GRCh38,1000 Genomes 30x on GRCh38,1000 Genomes phase 3 release |
| **HG00445** | male | SAME124975 | CHS | Southern Han Chinese | EAS | East Asian Ancestry | CHS | 1000 Genomes on GRCh38,1000 Genomes 30x on GRCh38,1000 Genomes phase 3 release,1000 Genomes phase 1 release |
| **HG01345** | female | SAME124863 | CLM | Colombian | AMR | American Ancestry | CLM | 1000 Genomes on GRCh38,1000 Genomes 30x on GRCh38,1000 Genomes phase 3 release,1000 Genomes phase 1 release |
| **HG00452** | female | SAME123115 | CHS | Southern Han Chinese | EAS | East Asian Ancestry | CHS | 1000 Genomes on GRCh38,1000 Genomes 30x on GRCh38,1000 Genomes phase 3 release,1000 Genomes phase 1 release |
| **HG01352** | female | SAME124686 | CLM | Colombian | AMR | American Ancestry | CLM | 1000 Genomes 30x on GRCh38,Human Genome Structural Variation Consortium, Phase 2,1000 Genomes phase 3 release |
| **HG00457** | male | SAME123095 | CHS | Southern Han Chinese | EAS | East Asian Ancestry | CHS | 1000 Genomes on GRCh38,1000 Genomes 30x on GRCh38,1000 Genomes phase 3 release,1000 Genomes phase 1 release |
| **HG01357** | female | SAME124681 | CLM | Colombian | AMR | American Ancestry | CLM | 1000 Genomes on GRCh38,1000 Genomes 30x on GRCh38,1000 Genomes phase 3 release,1000 Genomes phase 1 release |
| **HG00464** | female | SAME123497 | CHS | Southern Han Chinese | EAS | East Asian Ancestry | CHS | 1000 Genomes on GRCh38,1000 Genomes 30x on GRCh38,1000 Genomes phase 3 release,1000 Genomes phase 1 release |
| **HG00476** | female | SAME123341 | CHS | Southern Han Chinese | EAS | East Asian Ancestry | CHS | 1000 Genomes on GRCh38,1000 Genomes 30x on GRCh38,1000 Genomes phase 3 release,1000 Genomes phase 1 release,90 Han Chinese high coverage genomes |
| **HG01369** | female | SAME1839124 | CLM | Colombian | AMR | American Ancestry | CLM | 1000 Genomes on GRCh38,1000 Genomes 30x on GRCh38,1000 Genomes phase 3 release |
| **HG00534** | female | SAME123915 | CHS | Southern Han Chinese | EAS | East Asian Ancestry | CHS | 1000 Genomes on GRCh38,1000 Genomes 30x on GRCh38,1000 Genomes phase 3 release,1000 Genomes phase 1 release,90 Han Chinese high coverage genomes |
| **HG01376** | female | SAME124289 | CLM | Colombian | AMR | American Ancestry | CLM | 1000 Genomes 30x on GRCh38,1000 Genomes phase 3 release |
| **HG00096** | male | SAME123268 | GBR | British | EUR | European Ancestry | GBR | 1000 Genomes on GRCh38,1000 Genomes 30x on GRCh38,Human Genome Structural Variation Consortium, Phase 2,1000 Genomes phase 3 release,1000 Genomes ph |
| **HG01112** | male | SAME123485 | CLM | Colombian | AMR | American Ancestry | CLM | 1000 Genomes on GRCh38,1000 Genomes 30x on GRCh38,1000 Genomes phase 3 release,1000 Genomes phase 1 release |
| **HG00729** | female | SAME1840273 | CHS | Southern Han Chinese | EAS | East Asian Ancestry | CHS | 1000 Genomes on GRCh38,1000 Genomes 30x on GRCh38,1000 Genomes phase 3 release |
| **HG01124** | male | SAME124053 | CLM | Colombian | AMR | American Ancestry | CLM | 1000 Genomes on GRCh38,1000 Genomes 30x on GRCh38,1000 Genomes phase 3 release,1000 Genomes phase 1 release |
| **HG00731** | male | SAME125390 | PUR | Puerto Rican | AMR | American Ancestry | PUR | 1000 Genomes on GRCh38,1000 Genomes 30x on GRCh38,Human Genome Structural Variation Consortium, Phase 2,The Human Genome Structural Variation Consortiu |
| **HG01131** | female | SAME1839985 | CLM | Colombian | AMR | American Ancestry | CLM | 1000 Genomes on GRCh38,1000 Genomes 30x on GRCh38,1000 Genomes phase 3 release |

| | | | | | | | | |
|---|---|---|---|---|---|---|---|---|
| **HG00736** | male | SAME123058 | PUR | Puerto Rican | AMR | American Ancestry | PUR | 1000 Genomes on GRCh38,1000 Genomes 30x on GRCh38,1000 Genomes phase 3 release,1000 Genomes phase 1 release |
| **HG01136** | male | SAME123889 | CLM | Colombian | AMR | American Ancestry | CLM | 1000 Genomes on GRCh38,1000 Genomes 30x on GRCh38,1000 Genomes phase 3 release,1000 Genomes phase 1 release |
| **HG00743** | female | SAME1840184 | PUR | Puerto Rican | AMR | American Ancestry | PUR | 1000 Genomes on GRCh38,1000 Genomes 30x on GRCh38,1000 Genomes phase 3 release |
| **HG01148** | male | SAME124424 | CLM | Colombian | AMR | American Ancestry | CLM | 1000 Genomes on GRCh38,1000 Genomes 30x on GRCh38,1000 Genomes phase 3 release,1000 Genomes phase 1 release |
| **HG00844** | male | SAME123952 | CDX | Dai Chinese | EAS | East Asian Ancestry | CDX | 1000 Genomes on GRCh38,1000 Genomes 30x on GRCh38,1000 Genomes phase 3 release |
| **HG01150** | female | SAME124216 | CLM | Colombian | AMR | American Ancestry | CLM | 1000 Genomes 30x on GRCh38,1000 Genomes phase 3 release |
| **HG00851** | female | SAME123756 | CDX | Dai Chinese | EAS | East Asian Ancestry | CDX | 1000 Genomes on GRCh38,1000 Genomes 30x on GRCh38,1000 Genomes phase 3 release |
| **HG01162** | female | SAME1839707 | PUR | Puerto Rican | AMR | American Ancestry | PUR | 1000 Genomes on GRCh38,1000 Genomes 30x on GRCh38,1000 Genomes phase 3 release |
| **HG01167** | male | SAME124798 | PUR | Puerto Rican | AMR | American Ancestry | PUR | 1000 Genomes on GRCh38,1000 Genomes 30x on GRCh38,1000 Genomes phase 3 release,1000 Genomes phase 1 release |
| **HG01082** | male | SAME125398 | PUR | Puerto Rican | AMR | American Ancestry | PUR | 1000 Genomes on GRCh38,1000 Genomes 30x on GRCh38,1000 Genomes phase 3 release,1000 Genomes phase 1 release |
| **HG01066** | male | SAME124294 | PUR | Puerto Rican | AMR | American Ancestry | PUR | 1000 Genomes on GRCh38,1000 Genomes 30x on GRCh38,1000 Genomes phase 3 release,1000 Genomes phase 1 release |
| **HG01087** | female | SAME125393 | PUR | Puerto Rican | AMR | American Ancestry | PUR | 1000 Genomes 30x on GRCh38,1000 Genomes phase 3 release |
| **HG01073** | female | SAME124095 | PUR | Puerto Rican | AMR | American Ancestry | PUR | 1000 Genomes on GRCh38,1000 Genomes 30x on GRCh38,1000 Genomes phase 3 release,1000 Genomes phase 1 release |
| **HG01094** | male | SAME122925 | PUR | Puerto Rican | AMR | American Ancestry | PUR | 1000 Genomes on GRCh38,1000 Genomes 30x on GRCh38,1000 Genomes phase 3 release,1000 Genomes phase 1 release |
| **HG01080** | female | SAME125333 | PUR | Puerto Rican | AMR | American Ancestry | PUR | 1000 Genomes on GRCh38,1000 Genomes 30x on GRCh38,1000 Genomes phase 3 release,1000 Genomes phase 1 release |
| **HG01099** | male | SAME123771 | PUR | Puerto Rican | AMR | American Ancestry | PUR | 1000 Genomes 30x on GRCh38,1000 Genomes phase 3 release |
| **HG01085** | male | SAME125391 | PUR | Puerto Rican | AMR | American Ancestry | PUR | 1000 Genomes on GRCh38,1000 Genomes 30x on GRCh38,1000 Genomes phase 3 release,1000 Genomes phase 1 release |
| **HG01102** | female | SAME123678 | PUR | Puerto Rican | AMR | American Ancestry | PUR | 1000 Genomes on GRCh38,1000 Genomes 30x on GRCh38,1000 Genomes phase 3 release,1000 Genomes phase 1 release |
| **HG01092** | female | SAME1839489 | PUR | Puerto Rican | AMR | American Ancestry | PUR | 1000 Genomes on GRCh38,1000 Genomes 30x on GRCh38,1000 Genomes phase 3 release |
| **HG01107** | male | SAME123675 | PUR | Puerto Rican | AMR | American Ancestry | PUR | 1000 Genomes on GRCh38,1000 Genomes 30x on GRCh38,1000 Genomes phase 3 release,1000 Genomes phase 1 release |
| **HG01097** | male | SAME122924 | PUR | Puerto Rican | AMR | American Ancestry | PUR | 1000 Genomes on GRCh38,1000 Genomes 30x on GRCh38,1000 Genomes phase 3 release,1000 Genomes phase 1 release |
| **HG01114** | female | SAME123483 | CLM | Colombian | AMR | American Ancestry | CLM | 1000 Genomes 30x on GRCh38,Human Genome Structural Variation Consortium, Phase 2,1000 Genomes phase 3 release |
| **HG01100** | female | SAME123676 | PUR | Puerto Rican | AMR | American Ancestry | PUR | 1000 Genomes 30x on GRCh38,1000 Genomes phase 3 release |
| **HG01119** | female | SAME1840402 | CLM | Colombian | AMR | American Ancestry | CLM | 1000 Genomes on GRCh38,1000 Genomes 30x on GRCh38,1000 Genomes phase 3 release |
| **HG01105** | female | SAME123673 | PUR | Puerto Rican | AMR | American Ancestry | PUR | 1000 Genomes on GRCh38,1000 Genomes 30x on GRCh38,1000 Genomes phase 3 release,1000 Genomes phase 1 release |
| **HG01121** | male | SAME1840127 | CLM | Colombian | AMR | American Ancestry | CLM | 1000 Genomes on GRCh38,1000 Genomes 30x on GRCh38,1000 Genomes phase 3 release |
| **HG01174** | female | SAME124619 | PUR | Puerto Rican | AMR | American Ancestry | PUR | 1000 Genomes on GRCh38,1000 Genomes 30x on GRCh38,1000 Genomes phase 3 release,1000 Genomes phase 1 release |
| **HG01126** | male | SAME124052 | CLM | Colombian | AMR | American Ancestry | CLM | 1000 Genomes 30x on GRCh38,1000 Genomes phase 3 release |
| **HG01198** | female | SAME125228 | PUR | Puerto Rican | AMR | American Ancestry | PUR | 1000 Genomes on GRCh38,1000 Genomes 30x on GRCh38,1000 Genomes phase 3 release,1000 Genomes phase 1 release |
| **HG01133** | male | SAME123884 | CLM | Colombian | AMR | American Ancestry | CLM | 1000 Genomes on GRCh38,1000 Genomes 30x on GRCh38,1000 Genomes phase 3 release,1000 Genomes phase 1 release |
| **HG01206** | male | SAME124295 | PUR | Puerto Rican | AMR | American Ancestry | PUR | 1000 Genomes 30x on GRCh38,1000 Genomes phase 3 release |
| **HG01138** | male | SAME123877 | CLM | Colombian | AMR | American Ancestry | CLM | 1000 Genomes 30x on GRCh38,1000 Genomes phase 3 release |
| **HG01249** | female | SAME125057 | PUR | Puerto Rican | AMR | American Ancestry | PUR | 1000 Genomes 30x on GRCh38,1000 Genomes phase 3 release |
| **HG01140** | female | SAME124422 | CLM | Colombian | AMR | American Ancestry | CLM | 1000 Genomes on GRCh38,1000 Genomes 30x on GRCh38,1000 Genomes phase 3 release,1000 Genomes phase 1 release |
| **HG01251** | female | SAME125282 | CLM | Colombian | AMR | American Ancestry | CLM | 1000 Genomes on GRCh38,1000 Genomes 30x on GRCh38,1000 Genomes phase 3 release,1000 Genomes phase 1 release |
| **HG01164** | male | SAME1839690 | PUR | Puerto Rican | AMR | American Ancestry | PUR | 1000 Genomes on GRCh38,1000 Genomes 30x on GRCh38,1000 Genomes phase 3 release |
| **HG01256** | male | SAME125287 | CLM | Colombian | AMR | American Ancestry | CLM | 1000 Genomes on GRCh38,1000 Genomes 30x on GRCh38,1000 Genomes phase 3 release,1000 Genomes phase 1 release |
| **HG01169** | female | SAME124797 | PUR | Puerto Rican | AMR | American Ancestry | PUR | 1000 Genomes 30x on GRCh38,1000 Genomes phase 3 release |
| **HG00703** | male | SAME122933 | CHS | Southern Han Chinese | EAS | East Asian Ancestry | CHS | 1000 Genomes 30x on GRCh38,1000 Genomes phase 3 release |
| **HG01171** | female | SAME124622 | PUR | Puerto Rican | AMR | American Ancestry | PUR | 1000 Genomes on GRCh38,1000 Genomes 30x on GRCh38,1000 Genomes phase 3 release,1000 Genomes phase 1 release |
| **HG00708** | female | SAME122940 | CHS | Southern Han Chinese | EAS | East Asian Ancestry | CHS | 1000 Genomes on GRCh38,1000 Genomes 30x on GRCh38,1000 Genomes phase 3 release,1000 Genomes phase 1 release |
| **HG01176** | male | SAME124621 | PUR | Puerto Rican | AMR | American Ancestry | PUR | 1000 Genomes on GRCh38,1000 Genomes 30x on GRCh38,1000 Genomes phase 3 release,1000 Genomes phase 1 release |
| **HG00734** | female | SAME123016 | PUR | Puerto Rican | AMR | American Ancestry | PUR | 1000 Genomes on GRCh38,1000 Genomes 30x on GRCh38,1000 Genomes phase 3 release,1000 Genomes phase 1 release |
| **HG01183** | female | SAME124792 | PUR | Puerto Rican | AMR | American Ancestry | PUR | 1000 Genomes on GRCh38,1000 Genomes 30x on GRCh38,1000 Genomes phase 3 release,1000 Genomes phase 1 release |
| **HG00739** | male | SAME125113 | PUR | Puerto Rican | AMR | American Ancestry | PUR | 1000 Genomes on GRCh38,1000 Genomes 30x on GRCh38,1000 Genomes phase 3 release,1000 Genomes phase 1 release |
| **HG01188** | female | SAME124927 | PUR | Puerto Rican | AMR | American Ancestry | PUR | 1000 Genomes on GRCh38,1000 Genomes 30x on GRCh38,1000 Genomes phase 3 release,1000 Genomes phase 1 release |
| **HG00741** | female | SAME124833 | PUR | Puerto Rican | AMR | American Ancestry | PUR | 1000 Genomes 30x on GRCh38,1000 Genomes phase 3 release |
| **HG01190** | male | SAME123654 | PUR | Puerto Rican | AMR | American Ancestry | PUR | 1000 Genomes on GRCh38,1000 Genomes 30x on GRCh38,1000 Genomes phase 3 release,1000 Genomes phase 1 release |
| **HG00623** | female | SAME1839285 | CHS | Southern Han Chinese | EAS | East Asian Ancestry | CHS | 1000 Genomes on GRCh38,1000 Genomes 30x on GRCh38,1000 Genomes phase 3 release |
| **HG01028** | male | SAME123524 | CDX | Dai Chinese | EAS | East Asian Ancestry | CDX | 1000 Genomes on GRCh38,1000 Genomes 30x on GRCh38,1000 Genomes phase 3 release |
| **HG00628** | male | SAME123085 | CHS | Southern Han Chinese | EAS | East Asian Ancestry | CHS | 1000 Genomes on GRCh38,1000 Genomes 30x on GRCh38,1000 Genomes phase 3 release,1000 Genomes phase 1 release |
| **HG01047** | male | SAME123166 | PUR | Puerto Rican | AMR | American Ancestry | PUR | 1000 Genomes on GRCh38,1000 Genomes 30x on GRCh38,1000 Genomes phase 3 release,1000 Genomes phase 1 release |
| **HG00630** | male | SAME122901 | CHS | Southern Han Chinese | EAS | East Asian Ancestry | CHS | 1000 Genomes 30x on GRCh38,1000 Genomes phase 3 release |
| **HG01054** | male | SAME123663 | PUR | Puerto Rican | AMR | American Ancestry | PUR | 1000 Genomes on GRCh38,1000 Genomes 30x on GRCh38,1000 Genomes phase 3 release,1000 Genomes phase 1 release |
| **HG00635** | female | SAME122902 | CHS | Southern Han Chinese | EAS | East Asian Ancestry | CHS | 1000 Genomes on GRCh38,1000 Genomes 30x on GRCh38,1000 Genomes phase 3 release,1000 Genomes phase 1 release |
| **HG01061** | female | SAME122821 | PUR | Puerto Rican | AMR | American Ancestry | PUR | 1000 Genomes on GRCh38,1000 Genomes 30x on GRCh38,1000 Genomes phase 3 release,1000 Genomes phase 1 release |
| **HG00642** | male | SAME123436 | PUR | Puerto Rican | AMR | American Ancestry | PUR | 1000 Genomes 30x on GRCh38,1000 Genomes phase 3 release |
| **HG01383** | male | SAME122971 | CLM | Colombian | AMR | American Ancestry | CLM | 1000 Genomes on GRCh38,1000 Genomes 30x on GRCh38,1000 Genomes phase 3 release,1000 Genomes phase 1 release |
| **HG00654** | female | SAME123234 | CHS | Southern Han Chinese | EAS | East Asian Ancestry | CHS | 1000 Genomes on GRCh38,1000 Genomes 30x on GRCh38,1000 Genomes phase 3 release,1000 Genomes phase 1 release |
| **HG01390** | female | SAME122805 | CLM | Colombian | AMR | American Ancestry | CLM | 1000 Genomes on GRCh38,1000 Genomes 30x on GRCh38,1000 Genomes phase 3 release,1000 Genomes phase 1 release |
| **HG01395** | male | SAME1840207 | PUR | Puerto Rican | AMR | American Ancestry | PUR | 1000 Genomes on GRCh38,1000 Genomes 30x on GRCh38,1000 Genomes phase 3 release |
| **HG01241** | male | SAME125065 | PUR | Puerto Rican | AMR | American Ancestry | PUR | 1000 Genomes on GRCh38,1000 Genomes 30x on GRCh38,1000 Genomes phase 3 release,1000 Genomes phase 1 release |
| **HG01403** | female | SAME1839867 | PUR | Puerto Rican | AMR | American Ancestry | PUR | 1000 Genomes on GRCh38,1000 Genomes 30x on GRCh38,1000 Genomes phase 3 release |
| **HG01253** | male | SAME125284 | CLM | Colombian | AMR | American Ancestry | CLM | 1000 Genomes on GRCh38,1000 Genomes 30x on GRCh38,1000 Genomes phase 3 release,1000 Genomes phase 1 release |
| **HG01258** | male | SAME125279 | CLM | Colombian | AMR | American Ancestry | CLM | 1000 Genomes 30x on GRCh38,1000 Genomes phase 3 release |
| **HG01275** | female | SAME123002 | CLM | Colombian | AMR | American Ancestry | CLM | 1000 Genomes on GRCh38,1000 Genomes 30x on GRCh38,1000 Genomes phase 3 release,1000 Genomes phase 1 release |
| **HG01260** | female | SAME122815 | CLM | Colombian | AMR | American Ancestry | CLM | 1000 Genomes on GRCh38,1000 Genomes 30x on GRCh38,1000 Genomes phase 3 release,1000 Genomes phase 1 release |
| **HG01302** | male | SAME1840038 | PUR | Puerto Rican | AMR | American Ancestry | PUR | 1000 Genomes on GRCh38,1000 Genomes 30x on GRCh38,1000 Genomes phase 3 release |
| **HG01272** | female | SAME122996 | CLM | Colombian | AMR | American Ancestry | CLM | 1000 Genomes on GRCh38,1000 Genomes 30x on GRCh38,1000 Genomes phase 3 release,1000 Genomes phase 1 release |
| **HG01504** | female | SAME124945 | IBS | Iberian,Spanish | EUR | European Ancestry,West Eurasia (SGDP) | IBS,SpanishSGDP | 1000 Genomes on GRCh38,Simons Genome Diversity Project,1000 Genomes 30x on GRCh38,1000 Genomes phase 3 release |
| **HG01277** | male | SAME123000 | CLM | Colombian | AMR | American Ancestry | CLM | 1000 Genomes on GRCh38,1000 Genomes 30x on GRCh38,1000 Genomes phase 3 release,1000 Genomes phase 1 release |
| **HG01509** | male | SAME124940 | IBS | Iberian | EUR | European Ancestry | IBS | 1000 Genomes on GRCh38,1000 Genomes 30x on GRCh38,1000 Genomes phase 3 release |
| **HG01354** | female | SAME124680 | CLM | Colombian | AMR | American Ancestry | CLM | 1000 Genomes on GRCh38,1000 Genomes 30x on GRCh38,1000 Genomes phase 3 release,1000 Genomes phase 1 release |
| **HG01511** | female | SAME124770 | IBS | Iberian | EUR | European Ancestry | IBS | 1000 Genomes 30x on GRCh38,1000 Genomes phase 3 release |
| **HG01359** | male | SAME124687 | CLM | Colombian | AMR | American Ancestry | CLM | 1000 Genomes on GRCh38,1000 Genomes 30x on GRCh38,1000 Genomes phase 3 release,1000 Genomes phase 1 release |
| **HG01516** | female | SAME124773 | IBS | Iberian | EUR | European Ancestry | IBS | 1000 Genomes on GRCh38,1000 Genomes 30x on GRCh38,1000 Genomes phase 3 release,1000 Genomes phase 1 release |
| **HG01361** | female | SAME124504 | CLM | Colombian | AMR | American Ancestry | CLM | 1000 Genomes 30x on GRCh38,1000 Genomes phase 3 release |
| **HG01523** | male | SAME125322 | IBS | Iberian | EUR | European Ancestry | IBS | 1000 Genomes 30x on GRCh38,1000 Genomes phase 3 release |
| **HG01366** | female | SAME125265 | CLM | Colombian | AMR | American Ancestry | CLM | 1000 Genomes on GRCh38,1000 Genomes 30x on GRCh38,1000 Genomes phase 3 release,1000 Genomes phase 1 release |
| **HG01528** | female | SAME125329 | IBS | Iberian | EUR | European Ancestry | IBS | 1000 Genomes on GRCh38,1000 Genomes 30x on GRCh38,1000 Genomes phase 3 release |
| **HG01378** | female | SAME124076 | CLM | Colombian | AMR | American Ancestry | CLM | 1000 Genomes on GRCh38,1000 Genomes 30x on GRCh38,1000 Genomes phase 3 release,1000 Genomes phase 1 release |
| **HG01597** | female | SAME125226 | KHV | Kinh Vietnamese | EAS | East Asian Ancestry | KHV | 1000 Genomes on GRCh38,1000 Genomes 30x on GRCh38,1000 Genomes phase 3 release |
| **HG01385** | female | SAME122970 | CLM | Colombian | AMR | American Ancestry | CLM | 1000 Genomes 30x on GRCh38,1000 Genomes phase 3 release |
| **HG01600** | female | SAME123456 | KHV | Kinh,Kinh Vietnamese | EAS | East Asia (SGDP),East Asian Ancestry | KinhSGDP,KHV | 1000 Genomes on GRCh38,Simons Genome Diversity Project,1000 Genomes 30x on GRCh38,1000 Genomes phase 3 release |
| **HG01392** | male | SAME1840234 | PUR | Puerto Rican | AMR | American Ancestry | PUR | 1000 Genomes on GRCh38,1000 Genomes 30x on GRCh38,1000 Genomes phase 3 release |
| **HG01605** | female | SAME123459 | IBS | Iberian | EUR | European Ancestry | IBS | 1000 Genomes on GRCh38,1000 Genomes 30x on GRCh38,1000 Genomes phase 3 release |
| **HG01049** | female | SAME125248 | PUR | Puerto Rican | AMR | American Ancestry | PUR | 1000 Genomes on GRCh38,1000 Genomes 30x on GRCh38,1000 Genomes phase 3 release,1000 Genomes phase 1 release |
| **HG01612** | female | SAME123625 | IBS | Iberian | EUR | European Ancestry | IBS | 1000 Genomes on GRCh38,1000 Genomes 30x on GRCh38,1000 Genomes phase 3 release |
| **HG01051** | male | SAME123712 | PUR | Puerto Rican | AMR | American Ancestry | PUR | 1000 Genomes on GRCh38,1000 Genomes 30x on GRCh38,1000 Genomes phase 3 release,1000 Genomes phase 1 release |
| **HG01530** | male | SAME125137 | IBS | Iberian | EUR | European Ancestry | IBS | 1000 Genomes on GRCh38,1000 Genomes 30x on GRCh38,1000 Genomes phase 3 release |
| **HG01056** | male | SAME123658 | PUR | Puerto Rican | AMR | American Ancestry | PUR | 1000 Genomes 30x on GRCh38,1000 Genomes phase 3 release |
| **HG01566** | female | SAME124564 | PEL | Peruvian | AMR | American Ancestry | PEL | 1000 Genomes on GRCh38,1000 Genomes 30x on GRCh38,1000 Genomes phase 3 release |
| **HG01063** | male | SAME1839767 | PUR | Puerto Rican | AMR | American Ancestry | PUR | 1000 Genomes on GRCh38,1000 Genomes 30x on GRCh38,1000 Genomes phase 3 release |
| **HG01573** | female | SAME124347 | PEL | Peruvian | AMR | American Ancestry | PEL | 1000 Genomes 30x on GRCh38,Human Genome Structural Variation Consortium, Phase 2,1000 Genomes phase 3 release |
| **HG01068** | female | SAME122826 | PUR | Puerto Rican | AMR | American Ancestry | PUR | 1000 Genomes 30x on GRCh38,1000 Genomes phase 3 release |
| **HG01578** | female | SAME124351 | PEL | Peruvian | AMR | American Ancestry | PEL | 1000 Genomes on GRCh38,1000 Genomes 30x on GRCh38,1000 Genomes phase 3 release |
| **HG01070** | female | SAME123993 | PUR | Puerto Rican | AMR | American Ancestry | PUR | 1000 Genomes on GRCh38,1000 Genomes 30x on GRCh38,1000 Genomes phase 3 release,1000 Genomes phase 1 release |
| **HG01617** | male | SAME123630 | IBS | Iberian | EUR | European Ancestry | IBS | 1000 Genomes on GRCh38,1000 Genomes 30x on GRCh38,1000 Genomes phase 3 release,1000 Genomes phase 1 release |
| **HG01075** | male | SAME124071 | PUR | Puerto Rican | AMR | American Ancestry | PUR | 1000 Genomes on GRCh38,1000 Genomes 30x on GRCh38,1000 Genomes phase 3 release,1000 Genomes phase 1 release |
| **HG01624** | male | SAME123840 | IBS | Iberian | EUR | European Ancestry | IBS | 1000 Genomes on GRCh38,1000 Genomes 30x on GRCh38,1000 Genomes phase 3 release,1000 Genomes phase 1 release |
| **HG00673** | male | SAME123605 | CHS | Southern Han Chinese | EAS | East Asian Ancestry | CHS | 1000 Genomes 30x on GRCh38,1000 Genomes phase 3 release |
| **HG01629** | male | SAME123836 | IBS | Iberian | EUR | European Ancestry | IBS | 1000 Genomes 30x on GRCh38,1000 Genomes phase 3 release |
| **HG00685** | male | SAME123891 | CHS | Southern Han Chinese | EAS | East Asian Ancestry | CHS | 1000 Genomes 30x on GRCh38,1000 Genomes phase 3 release |
| **HG01631** | male | SAME124035 | IBS | Iberian | EUR | European Ancestry | IBS | 1000 Genomes on GRCh38,1000 Genomes 30x on GRCh38,1000 Genomes phase 3 release |
| **HG00692** | male | SAME123684 | CHS | Southern Han Chinese | EAS | East Asian Ancestry | CHS | 1000 Genomes on GRCh38,1000 Genomes 30x on GRCh38,1000 Genomes phase 3 release,1000 Genomes phase 1 release |
| **HG01667** | male | SAME123116 | IBS | Iberian | EUR | European Ancestry | IBS | 1000 Genomes 30x on GRCh38,1000 Genomes phase 3 release |
| **HG00700** | female | SAME122932 | CHS | Southern Han Chinese | EAS | East Asian Ancestry | CHS | 1000 Genomes 30x on GRCh38,1000 Genomes phase 3 release |
| **HG01674** | male | SAME123286 | IBS | Iberian | EUR | European Ancestry | IBS | 1000 Genomes 30x on GRCh38,1000 Genomes phase 3 release |
| **HG00705** | female | SAME122927 | CHS | Southern Han Chinese | EAS | East Asian Ancestry | CHS | 1000 Genomes on GRCh38,1000 Genomes 30x on GRCh38,1000 Genomes phase 3 release,1000 Genomes phase 1 release |
| **HG00717** | female | SAME124926 | CHS | Southern Han Chinese | EAS | East Asian Ancestry | CHS | 1000 Genomes on GRCh38,1000 Genomes 30x on GRCh38,1000 Genomes phase 3 release |
| **HG01284** | female | SAME1839191 | CLM | Colombian | AMR | American Ancestry | CLM | 1000 Genomes on GRCh38,1000 Genomes 30x on GRCh38,1000 Genomes phase 3 release |

| | | | | | | | | |
|---|---|---|---|---|---|---|---|---|
| **HG01489** | female | SAME123972 | CLM | Colombian | AMR | American Ancestry | CLM | 1000 Genomes on GRCh38,1000 Genomes 30x on GRCh38,1000 Genomes phase 3 release,1000 Genomes phase 1 release |
| **HG01311** | male | SAME1839796 | PUR | Puerto Rican | AMR | American Ancestry | PUR | 1000 Genomes on GRCh38,1000 Genomes 30x on GRCh38,1000 Genomes phase 3 release |
| **HG01491** | male | SAME124140 | CLM | Colombian | AMR | American Ancestry | CLM | 1000 Genomes on GRCh38,1000 Genomes 30x on GRCh38,1000 Genomes phase 3 release,1000 Genomes phase 1 release |
| **HG01323** | female | SAME1839534 | PUR | Puerto Rican | AMR | American Ancestry | PUR | 1000 Genomes on GRCh38,1000 Genomes 30x on GRCh38,1000 Genomes phase 3 release |
| **HG01496** | female | SAME124133 | CLM | Colombian | AMR | American Ancestry | CLM | 1000 Genomes 30x on GRCh38,1000 Genomes phase 3 release |
| **HG01342** | female | SAME124856 | CLM | Colombian | AMR | American Ancestry | CLM | 1000 Genomes on GRCh38,1000 Genomes 30x on GRCh38,1000 Genomes phase 3 release,1000 Genomes phase 1 release |
| **HG01679** | female | SAME123293 | IBS | Iberian | EUR | European Ancestry | IBS | 1000 Genomes on GRCh38,1000 Genomes 30x on GRCh38,1000 Genomes phase 3 release |
| **HG01347** | male | SAME124861 | CLM | Colombian | AMR | American Ancestry | CLM | 1000 Genomes on GRCh38,1000 Genomes 30x on GRCh38,1000 Genomes phase 3 release,1000 Genomes phase 1 release |
| **HG01681** | female | SAME124655 | IBS | Iberian | EUR | European Ancestry | IBS | 1000 Genomes 30x on GRCh38,1000 Genomes phase 3 release |
| **HG01599** | female | SAME125238 | KHV | Kinh Vietnamese | EAS | East Asian Ancestry | KHV | 1000 Genomes on GRCh38,1000 Genomes 30x on GRCh38,1000 Genomes phase 3 release |
| **HG01686** | male | SAME124658 | IBS | Iberian | EUR | European Ancestry | IBS | 1000 Genomes on GRCh38,1000 Genomes 30x on GRCh38,1000 Genomes phase 3 release |
| **HG01602** | female | SAME123454 | IBS | Iberian | EUR | European Ancestry | IBS | 1000 Genomes on GRCh38,1000 Genomes 30x on GRCh38,1000 Genomes phase 3 release |
| **HG01698** | male | SAME1839214 | IBS | Iberian | EUR | European Ancestry | IBS | 1000 Genomes 30x on GRCh38,1000 Genomes phase 3 release |
| **HG01607** | female | SAME123457 | IBS | Iberian | EUR | European Ancestry | IBS | 1000 Genomes on GRCh38,1000 Genomes 30x on GRCh38,1000 Genomes phase 3 release |
| **HG01701** | male | SAME123262 | IBS | Iberian | EUR | European Ancestry | IBS | 1000 Genomes 30x on GRCh38,1000 Genomes phase 3 release |
| **HG01614** | female | SAME123627 | IBS | Iberian | EUR | European Ancestry | IBS | 1000 Genomes 30x on GRCh38,1000 Genomes phase 3 release |
| **HG01768** | female | SAME1839665 | IBS | Iberian | EUR | European Ancestry | IBS | 1000 Genomes on GRCh38,1000 Genomes 30x on GRCh38,1000 Genomes phase 3 release |
| **HG01619** | male | SAME123632 | IBS | Iberian | EUR | European Ancestry | IBS | 1000 Genomes on GRCh38,1000 Genomes 30x on GRCh38,1000 Genomes phase 3 release,1000 Genomes phase 1 release |
| **HG01770** | female | SAME124638 | IBS | Iberian | EUR | European Ancestry | IBS | 1000 Genomes on GRCh38,1000 Genomes 30x on GRCh38,1000 Genomes phase 3 release |
| **HG01621** | male | SAME123843 | IBS | Iberian | EUR | European Ancestry | IBS | 1000 Genomes 30x on GRCh38,1000 Genomes phase 3 release |
| **HG01775** | male | SAME124633 | IBS | Iberian | EUR | European Ancestry | IBS | 1000 Genomes on GRCh38,1000 Genomes 30x on GRCh38,1000 Genomes phase 3 release |
| **HG01626** | female | SAME123842 | IBS | Iberian | EUR | European Ancestry | IBS | 1000 Genomes on GRCh38,1000 Genomes 30x on GRCh38,1000 Genomes phase 3 release,1000 Genomes phase 1 release |
| **HG01782** | female | SAME124426 | IBS | Iberian | EUR | European Ancestry | IBS | 1000 Genomes 30x on GRCh38,1000 Genomes phase 3 release |
| **HG01486** | female | SAME1840048 | CLM | Colombian | AMR | American Ancestry | CLM | 1000 Genomes on GRCh38,1000 Genomes 30x on GRCh38,1000 Genomes phase 3 release |
| **HG01493** | female | SAME124138 | CLM | Colombian | AMR | American Ancestry | CLM | 1000 Genomes 30x on GRCh38,1000 Genomes phase 3 release |
| **HG01439** | female | SAME124972 | CLM | Colombian | AMR | American Ancestry | CLM | 1000 Genomes 30x on GRCh38,1000 Genomes phase 3 release |
| **HG01498** | female | SAME124142 | CLM | Colombian | AMR | American Ancestry | CLM | 1000 Genomes on GRCh38,1000 Genomes 30x on GRCh38,1000 Genomes phase 3 release,1000 Genomes phase 1 release |
| **HG01441** | female | SAME123236 | CLM | Colombian | AMR | American Ancestry | CLM | 1000 Genomes on GRCh38,1000 Genomes 30x on GRCh38,1000 Genomes phase 3 release,1000 Genomes phase 1 release |
| **HG01501** | female | SAME124942 | IBS | Iberian | EUR | European Ancestry | IBS | 1000 Genomes on GRCh38,1000 Genomes 30x on GRCh38,1000 Genomes phase 3 release |
| **HG01506** | male | SAME124947 | IBS | Iberian | EUR | European Ancestry | IBS | 1000 Genomes on GRCh38,1000 Genomes 30x on GRCh38,1000 Genomes phase 3 release |
| **HG01465** | female | SAME122882 | CLM | Colombian | AMR | American Ancestry | CLM | 1000 Genomes on GRCh38,1000 Genomes 30x on GRCh38,1000 Genomes phase 3 release,1000 Genomes phase 1 release |
| **HG01513** | female | SAME124768 | IBS | Iberian | EUR | European Ancestry | IBS | 1000 Genomes on GRCh38,1000 Genomes 30x on GRCh38,1000 Genomes phase 3 release |
| **HG01706** | female | SAME123263 | IBS | Iberian | EUR | European Ancestry | IBS | 1000 Genomes 30x on GRCh38,1000 Genomes phase 3 release |
| **HG01518** | male | SAME124766 | IBS | Iberian | EUR | European Ancestry | IBS | 1000 Genomes on GRCh38,1000 Genomes 30x on GRCh38,1000 Genomes phase 3 release,1000 Genomes phase 1 release |
| **HG01756** | male | SAME124988 | IBS | Iberian | EUR | European Ancestry | IBS | 1000 Genomes on GRCh38,1000 Genomes 30x on GRCh38,1000 Genomes phase 3 release |
| **HG01520** | female | SAME125323 | IBS | Iberian | EUR | European Ancestry | IBS | 1000 Genomes 30x on GRCh38,1000 Genomes phase 3 release |
| **HG01763** | female | SAME124812 | IBS | Iberian | EUR | European Ancestry | IBS | 1000 Genomes 30x on GRCh38,1000 Genomes phase 3 release |
| **HG01525** | female | SAME125328 | IBS | Iberian | EUR | European Ancestry | IBS | 1000 Genomes on GRCh38,1000 Genomes 30x on GRCh38,1000 Genomes phase 3 release |
| **HG01946** | male | SAME124154 | PEL | Peruvian | AMR | American Ancestry | PEL | 1000 Genomes 30x on GRCh38,1000 Genomes phase 3 release |
| **HG01532** | male | SAME125138 | IBS | Iberian | EUR | European Ancestry | IBS | 1000 Genomes 30x on GRCh38,1000 Genomes phase 3 release |
| **HG01953** | male | SAME123974 | PEL | Peruvian | AMR | American Ancestry | PEL | 1000 Genomes on GRCh38,1000 Genomes 30x on GRCh38,1000 Genomes phase 3 release |
| **HG01537** | female | SAME125134 | IBS | Iberian | EUR | European Ancestry | IBS | 1000 Genomes on GRCh38,1000 Genomes 30x on GRCh38,1000 Genomes phase 3 release |
| **HG01958** | female | SAME1839349 | ACB | African Caribbean | AFR | African Ancestry | ACB | 1000 Genomes on GRCh38,1000 Genomes 30x on GRCh38,1000 Genomes phase 3 release |
| **HG01960** | female | SAME1839076 | ACB | African Caribbean | AFR | African Ancestry | ACB | 1000 Genomes 30x on GRCh38,1000 Genomes phase 3 release |
| **HG01405** | male | SAME1839863 | PUR | Puerto Rican | AMR | American Ancestry | PUR | 1000 Genomes on GRCh38,1000 Genomes 30x on GRCh38,1000 Genomes phase 3 release |
| **HG01965** | female | SAME1839145 | PEL | Peruvian | AMR | American Ancestry | PEL | 1000 Genomes on GRCh38,1000 Genomes 30x on GRCh38,1000 Genomes phase 3 release |
| **HG01412** | male | SAME1839619 | PUR | Puerto Rican | AMR | American Ancestry | PUR | 1000 Genomes on GRCh38,1000 Genomes 30x on GRCh38,1000 Genomes phase 3 release |
| **HG01972** | female | SAME124329 | PEL | Peruvian | AMR | American Ancestry | PEL | 1000 Genomes 30x on GRCh38,1000 Genomes phase 3 release |
| **HG01431** | male | SAME1839260 | CLM | Colombian | AMR | American Ancestry | CLM | 1000 Genomes on GRCh38,1000 Genomes 30x on GRCh38,1000 Genomes phase 3 release |
| **HG01977** | male | SAME124324 | PEL | Peruvian | AMR | American Ancestry | PEL | 1000 Genomes on GRCh38,1000 Genomes 30x on GRCh38,1000 Genomes phase 3 release |
| **HG01443** | male | SAME1839032 | CLM | Colombian | AMR | American Ancestry | CLM | 1000 Genomes on GRCh38,1000 Genomes 30x on GRCh38,1000 Genomes phase 3 release |
| **HG01984** | male | SAME1840105 | PEL | Peruvian | AMR | American Ancestry | PEL | 1000 Genomes 30x on GRCh38,1000 Genomes phase 3 release |
| **HG01455** | male | SAME123429 | CLM | Colombian | AMR | American Ancestry | CLM | 1000 Genomes on GRCh38,1000 Genomes 30x on GRCh38,1000 Genomes phase 3 release,1000 Genomes phase 1 release |
| **HG01989** | female | SAME1840166 | ACB | African Caribbean | AFR | African Ancestry | ACB | 1000 Genomes on GRCh38,1000 Genomes 30x on GRCh38,1000 Genomes phase 3 release |
| **HG01462** | female | SAME123539 | CLM | Colombian | AMR | American Ancestry | CLM | 1000 Genomes on GRCh38,1000 Genomes 30x on GRCh38,1000 Genomes phase 3 release,1000 Genomes phase 1 release |
| **HG01922** | female | SAME123787 | PEL | Peruvian | AMR | American Ancestry | PEL | 1000 Genomes 30x on GRCh38,1000 Genomes phase 3 release |
| **HG01474** | female | SAME1840189 | CLM | Colombian | AMR | American Ancestry | CLM | 1000 Genomes on GRCh38,1000 Genomes 30x on GRCh38,1000 Genomes phase 3 release |
| **HG01927** | female | SAME123784 | PEL | Peruvian | AMR | American Ancestry | PEL | 1000 Genomes on GRCh38,1000 Genomes 30x on GRCh38,1000 Genomes phase 3 release |
| **HG01479** | male | SAME1840251 | CLM | Colombian | AMR | American Ancestry | CLM | 1000 Genomes on GRCh38,1000 Genomes 30x on GRCh38,1000 Genomes phase 3 release |
| **HG01934** | male | SAME123589 | PEL | Peruvian | AMR | American Ancestry | PEL | 1000 Genomes 30x on GRCh38,1000 Genomes phase 3 release |
| **HG01939** | female | SAME123592 | PEL | Peruvian | AMR | American Ancestry | PEL | 1000 Genomes on GRCh38,1000 Genomes 30x on GRCh38,1000 Genomes phase 3 release |
| **HG01551** | female | SAME124008 | CLM | Colombian | AMR | American Ancestry | CLM | 1000 Genomes on GRCh38,1000 Genomes 30x on GRCh38,1000 Genomes phase 3 release,1000 Genomes phase 1 release |
| **HG01941** | male | SAME124149 | PEL | Peruvian | AMR | American Ancestry | PEL | 1000 Genomes on GRCh38,1000 Genomes 30x on GRCh38,1000 Genomes phase 3 release |
| **HG01556** | male | SAME1840141 | CLM | Colombian | AMR | American Ancestry | CLM | 1000 Genomes on GRCh38,1000 Genomes 30x on GRCh38,1000 Genomes phase 3 release |
| **HG01864** | male | SAME123197 | KHV | Kinh Vietnamese | EAS | East Asian Ancestry | KHV | 1000 Genomes on GRCh38,1000 Genomes 30x on GRCh38,1000 Genomes phase 3 release |
| **HG01869** | female | SAME123192 | KHV | Kinh Vietnamese | EAS | East Asian Ancestry | KHV | 1000 Genomes on GRCh38,1000 Genomes 30x on GRCh38,1000 Genomes phase 3 release |
| **HG01633** | male | SAME124033 | IBS | Iberian | EUR | European Ancestry | IBS | 1000 Genomes 30x on GRCh38,1000 Genomes phase 3 release |
| **HG01871** | female | SAME123398 | KHV | Kinh Vietnamese | EAS | East Asian Ancestry | KHV | 1000 Genomes on GRCh38,1000 Genomes 30x on GRCh38,1000 Genomes phase 3 release |
| **HG01669** | male | SAME123110 | IBS | Iberian | EUR | European Ancestry | IBS | 1000 Genomes on GRCh38,1000 Genomes 30x on GRCh38,1000 Genomes phase 3 release |
| **HG01883** | female | SAME122848 | ACB | African Caribbean | AFR | African Ancestry | ACB | 1000 Genomes on GRCh38,1000 Genomes 30x on GRCh38,1000 Genomes phase 3 release |
| **HG01671** | male | SAME123291 | IBS | Iberian | EUR | European Ancestry | IBS | 1000 Genomes 30x on GRCh38,1000 Genomes phase 3 release |
| **HG02019** | female | SAME123450 | KHV | Kinh Vietnamese | EAS | East Asian Ancestry | KHV | 1000 Genomes on GRCh38,1000 Genomes 30x on GRCh38,1000 Genomes phase 3 release |
| **HG01676** | female | SAME123284 | IBS | Iberian | EUR | European Ancestry | IBS | 1000 Genomes on GRCh38,1000 Genomes 30x on GRCh38,1000 Genomes phase 3 release |
| **HG01683** | male | SAME124653 | IBS | Iberian | EUR | European Ancestry | IBS | 1000 Genomes 30x on GRCh38,1000 Genomes phase 3 release |
| **HG02026** | male | SAME125315 | KHV | Kinh Vietnamese | EAS | East Asian Ancestry | KHV | 1000 Genomes on GRCh38,1000 Genomes 30x on GRCh38,1000 Genomes phase 3 release |
| **HG01695** | female | SAME124845 | IBS | Iberian | EUR | European Ancestry | IBS | 1000 Genomes on GRCh38,1000 Genomes 30x on GRCh38,1000 Genomes phase 3 release |
| **HG02040** | male | SAME1839803 | KHV | Kinh Vietnamese | EAS | East Asian Ancestry | KHV | 1000 Genomes on GRCh38,1000 Genomes 30x on GRCh38,1000 Genomes phase 3 release |
| **HG01703** | male | SAME123260 | IBS | Iberian | EUR | European Ancestry | IBS | 1000 Genomes 30x on GRCh38,1000 Genomes phase 3 release |
| **HG01888** | male | SAME122852 | ACB | African Caribbean | AFR | African Ancestry | ACB | 1000 Genomes 30x on GRCh38,1000 Genomes phase 3 release |
| **HG01708** | male | SAME123253 | IBS | Iberian | EUR | European Ancestry | IBS | 1000 Genomes on GRCh38,1000 Genomes 30x on GRCh38,1000 Genomes phase 3 release |
| **HG01890** | male | SAME123028 | ACB | African Caribbean | AFR | African Ancestry | ACB | 1000 Genomes on GRCh38,1000 Genomes 30x on GRCh38,1000 Genomes phase 3 release |
| **HG01710** | female | SAME123084 | IBS | Iberian | EUR | European Ancestry | IBS | 1000 Genomes on GRCh38,1000 Genomes 30x on GRCh38,1000 Genomes phase 3 release |
| **HG01895** | male | SAME1840349 | ACB | African Caribbean | AFR | African Ancestry | ACB | 1000 Genomes 30x on GRCh38,1000 Genomes phase 3 release |
| **HG01746** | female | SAME125180 | IBS | Iberian | EUR | European Ancestry | IBS | 1000 Genomes on GRCh38,1000 Genomes 30x on GRCh38,1000 Genomes phase 3 release |
| **HG01915** | female | SAME1839757 | ACB | African Caribbean | AFR | African Ancestry | ACB | 1000 Genomes on GRCh38,1000 Genomes 30x on GRCh38,1000 Genomes phase 3 release |
| **HG01765** | male | SAME1839681 | IBS | Iberian | EUR | European Ancestry | IBS | 1000 Genomes on GRCh38,1000 Genomes 30x on GRCh38,1000 Genomes phase 3 release |
| **HG01991** | male | SAME124743 | PEL | Peruvian | AMR | American Ancestry | PEL | 1000 Genomes on GRCh38,1000 Genomes 30x on GRCh38,1000 Genomes phase 3 release |
| **HG01772** | female | SAME124636 | IBS | Iberian | EUR | European Ancestry | IBS | 1000 Genomes 30x on GRCh38,1000 Genomes phase 3 release |
| **HG02002** | male | SAME123618 | PEL | Peruvian | AMR | American Ancestry | PEL | 1000 Genomes on GRCh38,1000 Genomes 30x on GRCh38,1000 Genomes phase 3 release |
| **HG01998** | male | SAME124739 | PEL | Peruvian | AMR | American Ancestry | PEL | 1000 Genomes 30x on GRCh38,1000 Genomes phase 3 release |
| **HG02014** | male | SAME123448 | ACB | African Caribbean | AFR | African Ancestry | ACB | 1000 Genomes on GRCh38,1000 Genomes 30x on GRCh38,1000 Genomes phase 3 release |
| **HG02004** | female | SAME123620 | PEL | Peruvian | AMR | American Ancestry | PEL | 1000 Genomes 30x on GRCh38,1000 Genomes phase 3 release |
| **HG01840** | male | SAME123552 | KHV | Kinh Vietnamese | EAS | East Asian Ancestry | KHV | 1000 Genomes on GRCh38,1000 Genomes 30x on GRCh38,1000 Genomes phase 3 release |
| **HG02009** | male | SAME123622 | ACB | African Caribbean | AFR | African Ancestry | ACB | 1000 Genomes on GRCh38,1000 Genomes 30x on GRCh38,1000 Genomes phase 3 release |
| **HG01845** | female | SAME123555 | KHV | Kinh Vietnamese | EAS | East Asian Ancestry | KHV | 1000 Genomes on GRCh38,1000 Genomes 30x on GRCh38,1000 Genomes phase 3 release |
| **HG02011** | male | SAME123443 | ACB | African Caribbean | AFR | African Ancestry | ACB | 1000 Genomes 30x on GRCh38,Human Genome Structural Variation Consortium, Phase 2,1000 Genomes phase 3 release |
| **HG01852** | male | SAME123750 | KHV | Kinh Vietnamese | EAS | East Asian Ancestry | KHV | 1000 Genomes on GRCh38,1000 Genomes 30x on GRCh38,1000 Genomes phase 3 release |
| **HG02016** | female | SAME123446 | KHV | Kinh Vietnamese | EAS | East Asian Ancestry | KHV | 1000 Genomes on GRCh38,1000 Genomes 30x on GRCh38,1000 Genomes phase 3 release |
| **HG01857** | female | SAME123748 | KHV | Kinh Vietnamese | EAS | East Asian Ancestry | KHV | 1000 Genomes on GRCh38,1000 Genomes 30x on GRCh38,1000 Genomes phase 3 release |
| **HG02023** | male | SAME125307 | KHV | Kinh Vietnamese | EAS | East Asian Ancestry | KHV | 1000 Genomes on GRCh38,1000 Genomes 30x on GRCh38,1000 Genomes phase 3 release |
| **HG01794** | female | SAME124218 | CDX | Dai Chinese | EAS | East Asian Ancestry | CDX | 1000 Genomes on GRCh38,1000 Genomes 30x on GRCh38,1000 Genomes phase 3 release |
| **HG01777** | male | SAME124631 | IBS | Iberian | EUR | European Ancestry | IBS | 1000 Genomes on GRCh38,1000 Genomes 30x on GRCh38,1000 Genomes phase 3 release |
| **HG01799** | female | SAME124227 | CDX | Dai Chinese | EAS | East Asian Ancestry | CDX | 1000 Genomes on GRCh38,1000 Genomes 30x on GRCh38,1000 Genomes phase 3 release |
| **HG01784** | female | SAME124431 | IBS | Iberian | EUR | European Ancestry | IBS | 1000 Genomes on GRCh38,1000 Genomes 30x on GRCh38,1000 Genomes phase 3 release |
| **HG01802** | female | SAME124307 | CDX | Dai Chinese | EAS | East Asian Ancestry | CDX | 1000 Genomes on GRCh38,1000 Genomes 30x on GRCh38,1000 Genomes phase 3 release |
| **HG01789** | male | SAME124433 | GBR | British | EUR | European Ancestry | GBR | 1000 Genomes on GRCh38,1000 Genomes 30x on GRCh38,1000 Genomes phase 3 release,Geuvadis |
| **HG01807** | female | SAME124303 | CDX | Dai Chinese | EAS | East Asian Ancestry | CDX | 1000 Genomes on GRCh38,1000 Genomes 30x on GRCh38,1000 Genomes phase 3 release |
| **HG01791** | male | SAME124221 | GBR | British | EUR | European Ancestry | GBR | 1000 Genomes on GRCh38,1000 Genomes 30x on GRCh38,1000 Genomes phase 3 release,Geuvadis |
| **HG02076** | male | SAME124913 | KHV | Kinh Vietnamese | EAS | East Asian Ancestry | KHV | 1000 Genomes on GRCh38,1000 Genomes 30x on GRCh38,1000 Genomes phase 3 release |
| **HG01796** | female | SAME124220 | CDX | Dai Chinese | EAS | East Asian Ancestry | CDX | 1000 Genomes on GRCh38,1000 Genomes 30x on GRCh38,1000 Genomes phase 3 release |
| **HG02083** | male | SAME124962 | KHV | Kinh Vietnamese | EAS | East Asian Ancestry | KHV | 1000 Genomes 30x on GRCh38,1000 Genomes phase 3 release |

| | | | | | | | | |
|---|---|---|---|---|---|---|---|---|
| **HG01974** | male | SAME124323 | PEL | Peruvian | AMR | American Ancestry | PEL | 1000 Genomes on GRCh38,1000 Genomes 30x on GRCh38,1000 Genomes phase 3 release |
| **HG02088** | male | SAME124820 | KHV | Kinh Vietnamese | EAS | East Asian Ancestry | KHV | 1000 Genomes on GRCh38,1000 Genomes 30x on GRCh38,1000 Genomes phase 3 release |
| **HG01979** | male | SAME124331 | PEL | Peruvian | AMR | American Ancestry | PEL | 1000 Genomes on GRCh38,1000 Genomes 30x on GRCh38,1000 Genomes phase 3 release |
| **HG02090** | male | SAME124651 | PEL | Peruvian | AMR | American Ancestry | PEL | 1000 Genomes on GRCh38,1000 Genomes 30x on GRCh38,1000 Genomes phase 3 release |
| **HG01981** | female | SAME125247 | PEL | Peruvian | AMR | American Ancestry | PEL | 1000 Genomes 30x on GRCh38,1000 Genomes phase 3 release |
| **HG02095** | female | SAME1839064 | ACB | African Caribbean | AFR | African Ancestry | ACB | 1000 Genomes on GRCh38,1000 Genomes 30x on GRCh38,1000 Genomes phase 3 release |
| **HG01986** | male | SAME1840091 | ACB | African Caribbean | AFR | African Ancestry | ACB | 1000 Genomes on GRCh38,1000 Genomes 30x on GRCh38,1000 Genomes phase 3 release |
| **HG02278** | female | SAME1840367 | PEL | Peruvian | AMR | American Ancestry | PEL | 1000 Genomes on GRCh38,1000 Genomes 30x on GRCh38,1000 Genomes phase 3 release |
| **HG01993** | female | SAME124741 | PEL | Peruvian | AMR | American Ancestry | PEL | 1000 Genomes 30x on GRCh38,1000 Genomes phase 3 release |
| **HG02280** | female | SAME1840096 | ACB | African Caribbean | AFR | African Ancestry | ACB | 1000 Genomes 30x on GRCh38,1000 Genomes phase 3 release |
| **HG01892** | male | SAME123026 | PEL | Peruvian | AMR | American Ancestry | PEL | 1000 Genomes on GRCh38,1000 Genomes 30x on GRCh38,1000 Genomes phase 3 release |
| **HG02285** | male | SAME122914 | PEL | Peruvian | AMR | American Ancestry | PEL | 1000 Genomes on GRCh38,1000 Genomes 30x on GRCh38,1000 Genomes phase 3 release |
| **HG01897** | male | SAME1840343 | ACB | African Caribbean | AFR | African Ancestry | ACB | 1000 Genomes 30x on GRCh38,1000 Genomes phase 3 release |
| **HG02292** | female | SAME1839943 | PEL | Peruvian | AMR | American Ancestry | PEL | 1000 Genomes on GRCh38,1000 Genomes 30x on GRCh38,1000 Genomes phase 3 release |
| **HG01912** | male | SAME1839779 | ACB | African Caribbean | AFR | African Ancestry | ACB | 1000 Genomes on GRCh38,1000 Genomes 30x on GRCh38,1000 Genomes phase 3 release |
| **HG02300** | male | SAME1839856 | PEL | Peruvian | AMR | American Ancestry | PEL | 1000 Genomes 30x on GRCh38,1000 Genomes phase 3 release |
| **HG01917** | male | SAME123235 | PEL | Peruvian | AMR | American Ancestry | PEL | 1000 Genomes on GRCh38,1000 Genomes 30x on GRCh38,1000 Genomes phase 3 release |
| **HG02139** | female | SAME123660 | KHV | Kinh Vietnamese | EAS | East Asian Ancestry | KHV | 1000 Genomes on GRCh38,1000 Genomes 30x on GRCh38,1000 Genomes phase 3 release |
| **HG01924** | female | SAME123785 | PEL | Peruvian | AMR | American Ancestry | PEL | 1000 Genomes on GRCh38,1000 Genomes 30x on GRCh38,1000 Genomes phase 3 release |
| **HG02141** | male | SAME124611 | KHV | Kinh Vietnamese | EAS | East Asian Ancestry | KHV | 1000 Genomes on GRCh38,1000 Genomes 30x on GRCh38,1000 Genomes phase 3 release |
| **HG01866** | male | SAME123195 | KHV | Kinh Vietnamese | EAS | East Asian Ancestry | KHV | 1000 Genomes on GRCh38,1000 Genomes 30x on GRCh38,1000 Genomes phase 3 release |
| **HG02146** | male | SAME1839613 | PEL | Peruvian | AMR | American Ancestry | PEL | 1000 Genomes on GRCh38,1000 Genomes 30x on GRCh38,1000 Genomes phase 3 release |
| **HG01873** | male | SAME123400 | KHV | Kinh Vietnamese | EAS | East Asian Ancestry | KHV | 1000 Genomes on GRCh38,1000 Genomes 30x on GRCh38,1000 Genomes phase 3 release |
| **HG02153** | female | SAME124791 | CDX | Dai Chinese | EAS | East Asian Ancestry | CDX | 1000 Genomes on GRCh38,1000 Genomes 30x on GRCh38,1000 Genomes phase 3 release |
| **HG01878** | female | SAME123394 | KHV | Kinh Vietnamese | EAS | East Asian Ancestry | KHV | 1000 Genomes on GRCh38,1000 Genomes 30x on GRCh38,1000 Genomes phase 3 release |
| **HG02052** | female | SAME1839545 | ACB | African Caribbean | AFR | African Ancestry | ACB | 1000 Genomes on GRCh38,1000 Genomes 30x on GRCh38,1000 Genomes phase 3 release |
| **HG01880** | female | SAME1839109 | ACB | African Caribbean | AFR | African Ancestry | ACB | 1000 Genomes on GRCh38,1000 Genomes 30x on GRCh38,1000 Genomes phase 3 release |
| **HG02057** | female | SAME124146 | KHV | Kinh Vietnamese | EAS | East Asian Ancestry | KHV | 1000 Genomes on GRCh38,1000 Genomes 30x on GRCh38,1000 Genomes phase 3 release |
| **HG01885** | male | SAME122844 | ACB | African Caribbean | AFR | African Ancestry | ACB | 1000 Genomes on GRCh38,1000 Genomes 30x on GRCh38,1000 Genomes phase 3 release |
| **HG02064** | male | SAME1839658 | KHV | Kinh Vietnamese | EAS | East Asian Ancestry | KHV | 1000 Genomes on GRCh38,1000 Genomes 30x on GRCh38,1000 Genomes phase 3 release |
| **HG01948** | female | SAME124144 | PEL | Peruvian | AMR | American Ancestry | PEL | 1000 Genomes on GRCh38,1000 Genomes 30x on GRCh38,1000 Genomes phase 3 release |
| **HG02069** | female | SAME123375 | KHV | Kinh Vietnamese | EAS | East Asian Ancestry | KHV | 1000 Genomes on GRCh38,1000 Genomes 30x on GRCh38,1000 Genomes phase 3 release |
| **HG01950** | male | SAME123977 | PEL | Peruvian | AMR | American Ancestry | PEL | 1000 Genomes on GRCh38,1000 Genomes 30x on GRCh38,1000 Genomes phase 3 release |
| **HG02071** | male | SAME123102 | KHV | Kinh Vietnamese | EAS | East Asian Ancestry | KHV | 1000 Genomes 30x on GRCh38,1000 Genomes phase 3 release |
| **HG01955** | female | SAME123979 | PEL | Peruvian | AMR | American Ancestry | PEL | 1000 Genomes 30x on GRCh38,1000 Genomes phase 3 release |
| **HG01967** | male | SAME124530 | PEL | Peruvian | AMR | American Ancestry | PEL | 1000 Genomes on GRCh38,1000 Genomes 30x on GRCh38,1000 Genomes phase 3 release |
| **HG02259** | male | SAME1839319 | PEL | Peruvian | AMR | American Ancestry | PEL | 1000 Genomes on GRCh38,1000 Genomes 30x on GRCh38,1000 Genomes phase 3 release |
| **HG01936** | female | SAME123591 | PEL | Peruvian | AMR | American Ancestry | PEL | 1000 Genomes on GRCh38,1000 Genomes 30x on GRCh38,1000 Genomes phase 3 release |
| **HG02261** | male | SAME1839134 | PEL | Peruvian | AMR | American Ancestry | PEL | 1000 Genomes 30x on GRCh38,1000 Genomes phase 3 release |
| **HG00158** | female | SAME124585 | GBR | British | EUR | European Ancestry | GBR | 1000 Genomes on GRCh38,1000 Genomes 30x on GRCh38,1000 Genomes phase 3 release,1000 Genomes phase 1 release,Geuvadis |
| **HG01943** | male | SAME124151 | PEL | Peruvian | AMR | American Ancestry | PEL | 1000 Genomes 30x on GRCh38,1000 Genomes phase 3 release |
| **HG02266** | female | SAME1839110 | PEL | Peruvian | AMR | American Ancestry | PEL | 1000 Genomes on GRCh38,1000 Genomes 30x on GRCh38,1000 Genomes phase 3 release |
| **HG00160** | male | SAME124780 | GBR | British | EUR | European Ancestry | GBR | 1000 Genomes on GRCh38,1000 Genomes 30x on GRCh38,1000 Genomes phase 3 release,1000 Genomes phase 1 release,Geuvadis |
| **HG01842** | male | SAME123554 | KHV | Kinh Vietnamese | EAS | East Asian Ancestry | KHV | 1000 Genomes on GRCh38,1000 Genomes 30x on GRCh38,1000 Genomes phase 3 release |
| **HG02273** | female | SAME1840309 | PEL | Peruvian | AMR | American Ancestry | PEL | 1000 Genomes 30x on GRCh38,1000 Genomes phase 3 release |
| **HG00177** | female | SAME124957 | FIN | Finnish | EUR | European Ancestry | FIN | 1000 Genomes on GRCh38,1000 Genomes 30x on GRCh38,1000 Genomes phase 3 release,1000 Genomes phase 1 release,Geuvadis |
| **HG01847** | female | SAME123557 | KHV | Kinh Vietnamese | EAS | East Asian Ancestry | KHV | 1000 Genomes on GRCh38,1000 Genomes 30x on GRCh38,1000 Genomes phase 3 release |
| **HG02108** | female | SAME1840201 | ACB | African Caribbean | AFR | African Ancestry | ACB | 1000 Genomes on GRCh38,1000 Genomes 30x on GRCh38,1000 Genomes phase 3 release |
| **HG00189** | male | SAME123640 | FIN | Finnish | EUR | European Ancestry | FIN | 1000 Genomes on GRCh38,1000 Genomes 30x on GRCh38,1000 Genomes phase 3 release,1000 Genomes phase 1 release,Geuvadis |
| **HG01859** | female | SAME123535 | KHV | Kinh Vietnamese | EAS | East Asian Ancestry | KHV | 1000 Genomes on GRCh38,1000 Genomes 30x on GRCh38,1000 Genomes phase 3 release |
| **HG02122** | male | SAME123478 | KHV | Kinh Vietnamese | EAS | East Asian Ancestry | KHV | 1000 Genomes on GRCh38,1000 Genomes 30x on GRCh38,1000 Genomes phase 3 release |
| **HG00235** | female | SAME124122 | GBR | British | EUR | European Ancestry | GBR | 1000 Genomes on GRCh38,1000 Genomes 30x on GRCh38,1000 Genomes phase 3 release,1000 Genomes phase 1 release,Geuvadis |
| **HG01861** | male | SAME123200 | KHV | Kinh Vietnamese | EAS | East Asian Ancestry | KHV | 1000 Genomes on GRCh38,1000 Genomes 30x on GRCh38,1000 Genomes phase 3 release |
| **HG02127** | female | SAME123476 | KHV | Kinh Vietnamese | EAS | East Asian Ancestry | KHV | 1000 Genomes on GRCh38,1000 Genomes 30x on GRCh38,1000 Genomes phase 3 release |
| **HG00242** | male | SAME123223 | GBR | British | EUR | European Ancestry | GBR | 1000 Genomes on GRCh38,1000 Genomes 30x on GRCh38,1000 Genomes phase 3 release,1000 Genomes phase 1 release,Geuvadis |
| **HG01804** | female | SAME124302 | CDX | Dai Chinese | EAS | East Asian Ancestry | CDX | 1000 Genomes on GRCh38,1000 Genomes 30x on GRCh38,1000 Genomes phase 3 release |
| **HG02134** | male | SAME1839875 | KHV | Kinh Vietnamese | EAS | East Asian Ancestry | KHV | 1000 Genomes on GRCh38,1000 Genomes 30x on GRCh38,1000 Genomes phase 3 release |
| **HG01809** | female | SAME124299 | CDX | Dai Chinese | EAS | East Asian Ancestry | CDX | 1000 Genomes on GRCh38,1000 Genomes 30x on GRCh38,1000 Genomes phase 3 release |
| **HG02312** | female | SAME1839684 | PEL | Peruvian | AMR | American Ancestry | PEL | 1000 Genomes on GRCh38,1000 Genomes 30x on GRCh38,1000 Genomes phase 3 release |
| **HG00254** | female | SAME123050 | GBR | British | EUR | European Ancestry | GBR | 1000 Genomes on GRCh38,1000 Genomes 30x on GRCh38,1000 Genomes phase 3 release,1000 Genomes phase 1 release |
| **HG01811** | male | SAME125380 | CDX | Dai Chinese | EAS | East Asian Ancestry | CDX | 1000 Genomes on GRCh38,1000 Genomes 30x on GRCh38,1000 Genomes phase 3 release |
| **HG02317** | male | SAME1839656 | ACB | African Caribbean | AFR | African Ancestry | ACB | 1000 Genomes on GRCh38,1000 Genomes 30x on GRCh38,1000 Genomes phase 3 release |
| **HG00259** | female | SAME123056 | GBR | British | EUR | European Ancestry | GBR | 1000 Genomes on GRCh38,1000 Genomes 30x on GRCh38,1000 Genomes phase 3 release,1000 Genomes phase 1 release,Geuvadis |
| **HG01816** | male | SAME124521 | CDX | Dai Chinese | EAS | East Asian Ancestry | CDX | 1000 Genomes on GRCh38,1000 Genomes 30x on GRCh38,1000 Genomes phase 3 release |
| **HG02343** | male | SAME1839091 | ACB | African Caribbean | AFR | African Ancestry | ACB | 1000 Genomes on GRCh38,1000 Genomes 30x on GRCh38,1000 Genomes phase 3 release |
| **HG00261** | female | SAME123583 | GBR | British | EUR | European Ancestry | GBR | 1000 Genomes on GRCh38,1000 Genomes 30x on GRCh38,1000 Genomes phase 3 release,1000 Genomes phase 1 release,Geuvadis |
| **HG02282** | female | SAME1840177 | ACB | African Caribbean | AFR | African Ancestry | ACB | 1000 Genomes on GRCh38,1000 Genomes 30x on GRCh38,1000 Genomes phase 3 release |
| **HG02223** | female | SAME1839584 | IBS | Iberian | EUR | European Ancestry | IBS | 1000 Genomes on GRCh38,1000 Genomes 30x on GRCh38,1000 Genomes phase 3 release |
| **HG00266** | female | SAME123576 | FIN | Finnish | EUR | European Ancestry | FIN | 1000 Genomes on GRCh38,1000 Genomes 30x on GRCh38,1000 Genomes phase 3 release,1000 Genomes phase 1 release,Geuvadis |
| **HG02287** | female | SAME123755 | PEL | Peruvian | AMR | American Ancestry | PEL | 1000 Genomes 30x on GRCh38,1000 Genomes phase 3 release |
| **HG02230** | female | SAME1839383 | IBS | Iberian | EUR | European Ancestry | IBS | 1000 Genomes on GRCh38,1000 Genomes 30x on GRCh38,1000 Genomes phase 3 release |
| **HG00273** | male | SAME123419 | FIN | Finnish | EUR | European Ancestry | FIN | 1000 Genomes on GRCh38,1000 Genomes 30x on GRCh38,1000 Genomes phase 3 release,1000 Genomes phase 1 release,Geuvadis |
| **HG02299** | male | SAME1839994 | PEL | Peruvian | AMR | American Ancestry | PEL | 1000 Genomes on GRCh38,1000 Genomes 30x on GRCh38,1000 Genomes phase 3 release |
| **HG02235** | female | SAME123521 | IBS | Iberian | EUR | European Ancestry | IBS | 1000 Genomes on GRCh38,1000 Genomes 30x on GRCh38,1000 Genomes phase 3 release |
| **HG00278** | male | SAME123425 | FIN | Finnish | EUR | European Ancestry | FIN | 1000 Genomes on GRCh38,1000 Genomes 30x on GRCh38,1000 Genomes phase 3 release,1000 Genomes phase 1 release,Geuvadis |
| **HG02302** | male | SAME124279 | PEL | Peruvian | AMR | American Ancestry | PEL | 1000 Genomes 30x on GRCh38,1000 Genomes phase 3 release |
| **HG02165** | female | SAME124208 | CDX | Dai Chinese | EAS | East Asian Ancestry | CDX | 1000 Genomes on GRCh38,1000 Genomes 30x on GRCh38,1000 Genomes phase 3 release |
| **HG00280** | male | SAME122831 | FIN | Finnish | EUR | European Ancestry | FIN | 1000 Genomes on GRCh38,1000 Genomes 30x on GRCh38,1000 Genomes phase 3 release,1000 Genomes phase 1 release,Geuvadis |
| **HG02307** | male | SAME1839535 | ACB | African Caribbean | AFR | African Ancestry | ACB | 1000 Genomes on GRCh38,1000 Genomes 30x on GRCh38,1000 Genomes phase 3 release |
| **HG02184** | female | SAME123216 | CDX | Dai Chinese | EAS | East Asian Ancestry | CDX | 1000 Genomes on GRCh38,1000 Genomes 30x on GRCh38,1000 Genomes phase 3 release |
| **HG00285** | female | SAME122836 | FIN | Finnish | EUR | European Ancestry | FIN | 1000 Genomes on GRCh38,1000 Genomes 30x on GRCh38,1000 Genomes phase 3 release,1000 Genomes phase 1 release,Geuvadis |
| **HG02143** | male | SAME1839628 | ACB | African Caribbean | AFR | African Ancestry | ACB | 1000 Genomes on GRCh38,1000 Genomes 30x on GRCh38,1000 Genomes phase 3 release |
| **HG00103** | male | SAME125151 | GBR | British | EUR | European Ancestry | GBR | 1000 Genomes on GRCh38,1000 Genomes 30x on GRCh38,1000 Genomes phase 3 release,1000 Genomes phase 1 release,Geuvadis |
| **HG02148** | female | SAME1839697 | PEL | Peruvian | AMR | American Ancestry | PEL | 1000 Genomes 30x on GRCh38,1000 Genomes phase 3 release |
| **HG02689** | male | SAME1839355 | PJL | Punjabi | SAS | South Asian Ancestry | PJL | 1000 Genomes 30x on GRCh38,1000 Genomes phase 3 release |
| **HG00108** | male | SAME125161 | GBR | British | EUR | European Ancestry | GBR | 1000 Genomes on GRCh38,1000 Genomes 30x on GRCh38,1000 Genomes phase 3 release,1000 Genomes phase 1 release,Geuvadis |
| **HG02150** | male | SAME1839407 | PEL | Peruvian | AMR | American Ancestry | PEL | 1000 Genomes on GRCh38,1000 Genomes 30x on GRCh38,1000 Genomes phase 3 release |
| **HG02691** | female | SAME1839084 | PJL | Punjabi | SAS | South Asian Ancestry | PJL | 1000 Genomes on GRCh38,1000 Genomes 30x on GRCh38,1000 Genomes phase 3 release |
| **HG00110** | female | SAME125339 | GBR | British | EUR | European Ancestry | GBR | 1000 Genomes on GRCh38,1000 Genomes 30x on GRCh38,1000 Genomes phase 3 release,1000 Genomes phase 1 release,Geuvadis |
| **HG02155** | female | SAME124795 | CDX | Dai Chinese | EAS | East Asian Ancestry | CDX | 1000 Genomes on GRCh38,1000 Genomes 30x on GRCh38,1000 Genomes phase 3 release |
| **HG02696** | male | SAME1839149 | PJL | Punjabi | SAS | South Asian Ancestry | PJL | 1000 Genomes on GRCh38,1000 Genomes 30x on GRCh38,1000 Genomes phase 3 release |
| **HG00115** | male | SAME125344 | GBR | British | EUR | European Ancestry | GBR | 1000 Genomes on GRCh38,1000 Genomes 30x on GRCh38,1000 Genomes phase 3 release,1000 Genomes phase 1 release,Geuvadis |
| **HG02028** | female | SAME124020 | KHV | Kinh Vietnamese | EAS | East Asian Ancestry | KHV | 1000 Genomes on GRCh38,1000 Genomes 30x on GRCh38,1000 Genomes phase 3 release |
| **HG02704** | female | SAME1840134 | GWD | Gambian Mandinka | AFR | African Ancestry | GWD | 1000 Genomes 30x on GRCh38 |
| **HG00122** | female | SAME122876 | GBR | British | EUR | European Ancestry | GBR | 1000 Genomes on GRCh38,1000 Genomes 30x on GRCh38,1000 Genomes phase 3 release,1000 Genomes phase 1 release,Geuvadis |
| **HG02030** | male | SAME1839966 | KHV | Kinh Vietnamese | EAS | East Asian Ancestry | KHV | 1000 Genomes 30x on GRCh38,1000 Genomes phase 3 release |
| **HG02716** | female | SAME1839981 | GWD | Gambian Mandinka | AFR | African Ancestry | GWD | 1000 Genomes on GRCh38,1000 Genomes 30x on GRCh38,1000 Genomes phase 3 release,Gambian Genome Variation Project (GRCh38) |
| **HG00127** | female | SAME122871 | GBR | British | EUR | European Ancestry | GBR | 1000 Genomes on GRCh38,1000 Genomes 30x on GRCh38,1000 Genomes phase 3 release,1000 Genomes phase 1 release,Geuvadis |
| **HG02035** | male | SAME1840035 | KHV | Kinh Vietnamese | EAS | East Asian Ancestry | KHV | 1000 Genomes on GRCh38,1000 Genomes 30x on GRCh38,1000 Genomes phase 3 release |
| **HG02723** | female | SAME1839742 | GWD | Gambian Mandinka | AFR | African Ancestry | GWD | 1000 Genomes 30x on GRCh38 |
| **HG02047** | male | SAME1839772 | KHV | Kinh Vietnamese | EAS | East Asian Ancestry | KHV | 1000 Genomes on GRCh38,1000 Genomes 30x on GRCh38,1000 Genomes phase 3 release |
| **HG02728** | female | SAME1839722 | PJL | Punjabi | SAS | South Asian Ancestry | PJL | 1000 Genomes on GRCh38,1000 Genomes 30x on GRCh38,1000 Genomes phase 3 release |
| **HG00139** | male | SAME123060 | GBR | British | EUR | European Ancestry | GBR | 1000 Genomes on GRCh38,1000 Genomes 30x on GRCh38,1000 Genomes phase 3 release,1000 Genomes phase 1 release,Geuvadis |
| **HG02054** | female | SAME1839538 | ACB | African Caribbean | AFR | African Ancestry | ACB | 1000 Genomes on GRCh38,1000 Genomes 30x on GRCh38,1000 Genomes phase 3 release |
| **HG02603** | male | SAME1840287 | PJL | Punjabi | SAS | South Asian Ancestry | PJL | 1000 Genomes on GRCh38,1000 Genomes 30x on GRCh38,1000 Genomes phase 3 release |
| **HG00141** | male | SAME124395 | GBR | British | EUR | European Ancestry | GBR | 1000 Genomes on GRCh38,1000 Genomes 30x on GRCh38,1000 Genomes phase 3 release,1000 Genomes phase 1 release,Geuvadis |
| **HG02059** | female | SAME1839596 | KHV | Kinh Vietnamese | EAS | East Asian Ancestry | KHV | 1000 Genomes 30x on GRCh38,Human Genome Structural Variation Consortium, Phase 2,1000 Genomes phase 3 release |
| **HG02610** | male | SAME1840405 | GWD | Gambian Mandinka | AFR | African Ancestry | GWD | 1000 Genomes on GRCh38,1000 Genomes 30x on GRCh38,1000 Genomes phase 3 release,Gambian Genome Variation Project (GRCh38) |
| **HG00146** | female | SAME124390 | GBR | British | EUR | European Ancestry | GBR | 1000 Genomes on GRCh38,1000 Genomes 30x on GRCh38,1000 Genomes phase 3 release,1000 Genomes phase 1 release,Geuvadis |
| **HG02061** | male | SAME1839676 | KHV | Kinh Vietnamese | EAS | East Asian Ancestry | KHV | 1000 Genomes on GRCh38,1000 Genomes 30x on GRCh38,1000 Genomes phase 3 release |
| **HG02615** | female | SAME1840380 | GWD | Gambian Mandinka | AFR | African Ancestry | GWD | 1000 Genomes 30x on GRCh38 |

| | | | | | | | | |
|---|---|---|---|---|---|---|---|---|
| **HG02073** | male | SAME123104 | KHV | Kinh Vietnamese | EAS | East Asian Ancestry | KHV | 1000 Genomes on GRCh38,1000 Genomes 30x on GRCh38,1000 Genomes phase 3 release |
| **HG02622** | female | SAME1840185 | GWD | Gambian Mandinka | AFR | African Ancestry | GWD | 1000 Genomes 30x on GRCh38 |
| **HG00324** | female | SAME123794 | FIN | Finnish | EUR | European Ancestry | FIN | 1000 Genomes on GRCh38,1000 Genomes 30x on GRCh38,1000 Genomes phase 3 release,1000 Genomes phase 1 release,Geuvadis |
| **HG02078** | female | SAME123099 | KHV | Kinh Vietnamese | EAS | East Asian Ancestry | KHV | 1000 Genomes on GRCh38,1000 Genomes 30x on GRCh38,1000 Genomes phase 3 release |
| **HG02634** | male | SAME1840031 | GWD | Gambian Mandinka | AFR | African Ancestry | GWD | 1000 Genomes on GRCh38,1000 Genomes 30x on GRCh38,1000 Genomes phase 3 release,Gambian Genome Variation Project (GRCh38) |
| **HG00329** | male | SAME123798 | FIN | Finnish | EUR | European Ancestry | FIN | 1000 Genomes on GRCh38,1000 Genomes 30x on GRCh38,1000 Genomes phase 3 release,1000 Genomes phase 1 release,Geuvadis |
| **HG02179** | female | SAME124415 | CDX | Dai Chinese | EAS | East Asian Ancestry | CDX | 1000 Genomes on GRCh38,1000 Genomes 30x on GRCh38,1000 Genomes phase 3 release |
| **HG02646** | female | SAME1839770 | GWD | Gambian Mandinka | AFR | African Ancestry | GWD | 1000 Genomes on GRCh38,1000 Genomes 30x on GRCh38,1000 Genomes phase 3 release,Gambian Genome Variation Project (GRCh38) |
| **HG00331** | female | SAME123986 | FIN | Finnish | EUR | European Ancestry | FIN | 1000 Genomes on GRCh38,1000 Genomes 30x on GRCh38,1000 Genomes phase 3 release,1000 Genomes phase 1 release,Geuvadis |
| **HG02181** | female | SAME123206 | CDX | Dai Chinese | EAS | East Asian Ancestry | CDX | 1000 Genomes on GRCh38,1000 Genomes 30x on GRCh38,1000 Genomes phase 3 release |
| **HG02557** | male | SAME1840061 | ACB | African Caribbean | AFR | African Ancestry | ACB | 1000 Genomes on GRCh38,1000 Genomes 30x on GRCh38,1000 Genomes phase 3 release |
| **HG00336** | male | SAME123983 | FIN | Finnish | EUR | European Ancestry | FIN | 1000 Genomes on GRCh38,1000 Genomes 30x on GRCh38,1000 Genomes phase 3 release,1000 Genomes phase 1 release,Geuvadis |
| **HG02186** | female | SAME125039 | CDX | Dai Chinese | EAS | East Asian Ancestry | CDX | 1000 Genomes on GRCh38,1000 Genomes 30x on GRCh38,1000 Genomes phase 3 release |
| **HG02569** | male | SAME1839892 | GWD | Gambian Mandinka | AFR | African Ancestry | GWD | 1000 Genomes 30x on GRCh38 |
| **HG00343** | female | SAME124923 | FIN | Finnish | EUR | European Ancestry | FIN | 1000 Genomes on GRCh38,1000 Genomes 30x on GRCh38,1000 Genomes phase 3 release,1000 Genomes phase 1 release,Geuvadis |
| **HG02571** | female | SAME1839577 | GWD | Gambian Mandinka | AFR | African Ancestry | GWD | 1000 Genomes on GRCh38,1000 Genomes 30x on GRCh38,1000 Genomes phase 3 release,Gambian Genome Variation Project (GRCh38) |
| **HG00350** | female | SAME125121 | FIN | Finnish | EUR | European Ancestry | FIN | 1000 Genomes on GRCh38,1000 Genomes 30x on GRCh38,1000 Genomes phase 3 release,1000 Genomes phase 1 release,Geuvadis |
| **HG02220** | female | SAME123715 | IBS | Iberian | EUR | European Ancestry | IBS | 1000 Genomes on GRCh38,1000 Genomes 30x on GRCh38,1000 Genomes phase 3 release |
| **HG02583** | female | SAME1839429 | GWD | Gambian Mandinka | AFR | African Ancestry | GWD | 1000 Genomes on GRCh38,1000 Genomes 30x on GRCh38,1000 Genomes phase 3 release,Gambian Genome Variation Project (GRCh38) |
| **HG00355** | female | SAME125125 | FIN | Finnish | EUR | European Ancestry | FIN | 1000 Genomes on GRCh38,1000 Genomes 30x on GRCh38,1000 Genomes phase 3 release,1000 Genomes phase 1 release,Geuvadis |
| **HG02080** | female | SAME124825 | KHV | Kinh Vietnamese | EAS | East Asian Ancestry | KHV | 1000 Genomes 30x on GRCh38,1000 Genomes phase 3 release |
| **HG02588** | male | SAME1839404 | GWD | Gambian Mandinka | AFR | African Ancestry | GWD | 1000 Genomes on GRCh38,1000 Genomes 30x on GRCh38,1000 Genomes phase 3 release,Gambian Genome Variation Project (GRCh38) |
| **HG00362** | female | SAME124537 | FIN | Finnish | EUR | European Ancestry | FIN | 1000 Genomes on GRCh38,1000 Genomes 30x on GRCh38,1000 Genomes phase 3 release,1000 Genomes phase 1 release,Geuvadis |
| **HG02085** | male | SAME124979 | KHV | Kinh Vietnamese | EAS | East Asian Ancestry | KHV | 1000 Genomes on GRCh38,1000 Genomes 30x on GRCh38,1000 Genomes phase 3 release |
| **HG02590** | male | SAME1839222 | GWD | Gambian Mandinka | AFR | African Ancestry | GWD | 1000 Genomes 30x on GRCh38 |
| **HG00367** | female | SAME124540 | FIN | Finnish | EUR | European Ancestry | FIN | 1000 Genomes on GRCh38,1000 Genomes 30x on GRCh38,1000 Genomes phase 3 release,1000 Genomes phase 1 release,Geuvadis |
| **HG02105** | female | SAME123873 | PEL | Peruvian | AMR | American Ancestry | PEL | 1000 Genomes on GRCh38,1000 Genomes 30x on GRCh38,1000 Genomes phase 3 release |
| **HG02595** | female | SAME1839204 | GWD | Gambian Mandinka | AFR | African Ancestry | GWD | 1000 Genomes on GRCh38,1000 Genomes 30x on GRCh38,1000 Genomes phase 3 release,Gambian Genome Variation Project (GRCh38) |
| **HG00475** | male | SAME123342 | CHS | Southern Han Chinese | EAS | East Asian Ancestry | CHS | 1000 Genomes on GRCh38,1000 Genomes 30x on GRCh38,1000 Genomes phase 3 release,1000 Genomes phase 1 release,90 Han Chinese high coverage genomes |
| **HG02129** | male | SAME1839724 | KHV | Kinh Vietnamese | EAS | East Asian Ancestry | KHV | 1000 Genomes 30x on GRCh38,1000 Genomes phase 3 release |
| **HG02420** | male | SAME1839320 | ACB | African Caribbean | AFR | African Ancestry | ACB | 1000 Genomes on GRCh38,1000 Genomes 30x on GRCh38,1000 Genomes phase 3 release |
| **HG00502** | male | SAME123361 | CHS | Southern Han Chinese | EAS | East Asian Ancestry | CHS | 1000 Genomes 30x on GRCh38,1000 Genomes phase 3 release |
| **HG02131** | male | SAME1839896 | KHV | Kinh Vietnamese | EAS | East Asian Ancestry | KHV | 1000 Genomes on GRCh38,1000 Genomes 30x on GRCh38,1000 Genomes phase 3 release |
| **HG02425** | female | SAME1839043 | PEL | Peruvian | AMR | American Ancestry | PEL | 1000 Genomes on GRCh38,1000 Genomes 30x on GRCh38,1000 Genomes phase 3 release |
| **HG00514** | female | SAME123522 | CHS | Southern Han Chinese | EAS | East Asian Ancestry | CHS | 1000 Genomes 30x on GRCh38,Human Genome Structural Variation Consortium, Phase 2,The Human Genome Structural Variation Consortium,1000 Genomes phase 3 re |
| **HG02136** | female | SAME1839865 | KHV | Kinh Vietnamese | EAS | East Asian Ancestry | KHV | 1000 Genomes on GRCh38,1000 Genomes 30x on GRCh38,1000 Genomes phase 3 release |
| **HG02449** | male | SAME1840404 | ACB | African Caribbean | AFR | African Ancestry | ACB | 1000 Genomes on GRCh38,1000 Genomes 30x on GRCh38,1000 Genomes phase 3 release |
| **HG00526** | male | SAME123395 | CHS | Southern Han Chinese | EAS | East Asian Ancestry | CHS | 1000 Genomes 30x on GRCh38,1000 Genomes phase 3 release |
| **HG02225** | female | SAME1839576 | IBS | Iberian | EUR | European Ancestry | IBS | 1000 Genomes 30x on GRCh38,1000 Genomes phase 3 release |
| **HG02451** | female | SAME1840138 | ACB | African Caribbean | AFR | African Ancestry | ACB | 1000 Genomes 30x on GRCh38,1000 Genomes phase 3 release |
| **HG00533** | male | SAME124353 | CHS | Southern Han Chinese | EAS | East Asian Ancestry | CHS | 1000 Genomes on GRCh38,1000 Genomes 30x on GRCh38,1000 Genomes phase 3 release,1000 Genomes phase 1 release,90 Han Chinese high coverage genomes |
| **HG02232** | female | SAME1839374 | IBS | Iberian | EUR | European Ancestry | IBS | 1000 Genomes on GRCh38,1000 Genomes 30x on GRCh38,1000 Genomes phase 3 release |
| **HG02463** | male | SAME1839983 | GWD | Gambian Mandinka | AFR | African Ancestry | GWD | 1000 Genomes 30x on GRCh38 |
| **HG00538** | male | SAME124343 | CHS | Southern Han Chinese | EAS | East Asian Ancestry | CHS | 1000 Genomes 30x on GRCh38,1000 Genomes phase 3 release |
| **HG02237** | female | SAME123519 | IBS | Iberian | EUR | European Ancestry | IBS | 1000 Genomes 30x on GRCh38,1000 Genomes phase 3 release |
| **HG02348** | female | SAME1839063 | PEL | Peruvian | AMR | American Ancestry | PEL | 1000 Genomes on GRCh38,1000 Genomes 30x on GRCh38,1000 Genomes phase 3 release |
| **HG00552** | male | SAME124253 | PUR | Puerto Rican | AMR | American Ancestry | PUR | 1000 Genomes 30x on GRCh38,1000 Genomes phase 3 release |
| **HG02256** | female | SAME1839332 | ACB | African Caribbean | AFR | African Ancestry | ACB | 1000 Genomes on GRCh38,1000 Genomes 30x on GRCh38,1000 Genomes phase 3 release |
| **HG02355** | male | SAME1840269 | CDX | Dai Chinese | EAS | East Asian Ancestry | CDX | 1000 Genomes on GRCh38,1000 Genomes 30x on GRCh38,1000 Genomes phase 3 release |
| **HG00557** | female | SAME124258 | CHS | Southern Han Chinese | EAS | East Asian Ancestry | CHS | 1000 Genomes on GRCh38,1000 Genomes 30x on GRCh38,1000 Genomes phase 3 release,1000 Genomes phase 1 release,90 Han Chinese high coverage genomes |
| **HG02275** | female | SAME1840376 | PEL | Peruvian | AMR | American Ancestry | PEL | 1000 Genomes on GRCh38,1000 Genomes 30x on GRCh38,1000 Genomes phase 3 release |
| **HG02367** | male | SAME1840113 | CDX | Dai Chinese | EAS | East Asian Ancestry | CDX | 1000 Genomes on GRCh38,1000 Genomes 30x on GRCh38,1000 Genomes phase 3 release |
| **HG00583** | male | SAME124829 | CHS | Southern Han Chinese | EAS | East Asian Ancestry | CHS | 1000 Genomes on GRCh38,1000 Genomes 30x on GRCh38,1000 Genomes phase 3 release,1000 Genomes phase 1 release,90 Han Chinese high coverage genomes |
| **HG02605** | female | SAME1840277 | PJL | Punjabi | SAS | South Asian Ancestry | PJL | 1000 Genomes 30x on GRCh38,1000 Genomes phase 3 release |
| **HG02374** | male | SAME1840247 | CDX | Dai Chinese | EAS | East Asian Ancestry | CDX | 1000 Genomes on GRCh38,1000 Genomes 30x on GRCh38,1000 Genomes phase 3 release |
| **HG00379** | female | SAME124746 | FIN | Finnish | EUR | European Ancestry | FIN | 1000 Genomes on GRCh38,1000 Genomes 30x on GRCh38,1000 Genomes phase 3 release,1000 Genomes phase 1 release,Geuvadis |
| **HG02612** | male | SAME1840394 | GWD | Gambian Mandinka | AFR | African Ancestry | GWD | 1000 Genomes 30x on GRCh38 |
| **HG02379** | male | SAME1840217 | CDX | Dai Chinese | EAS | East Asian Ancestry | CDX | 1000 Genomes on GRCh38,1000 Genomes 30x on GRCh38,1000 Genomes phase 3 release |
| **HG00381** | female | SAME123030 | FIN | Finnish | EUR | European Ancestry | FIN | 1000 Genomes on GRCh38,1000 Genomes 30x on GRCh38,1000 Genomes phase 3 release,1000 Genomes phase 1 release,Geuvadis |
| **HG02624** | female | SAME1840259 | GWD | Gambian Mandinka | AFR | African Ancestry | GWD | 1000 Genomes on GRCh38,1000 Genomes 30x on GRCh38,1000 Genomes phase 3 release,Gambian Genome Variation Project (GRCh38) |
| **HG00406** | male | SAME123156 | CHS | Southern Han Chinese | EAS | East Asian Ancestry | CHS | 1000 Genomes on GRCh38,1000 Genomes 30x on GRCh38,1000 Genomes phase 3 release,1000 Genomes phase 1 release |
| **HG02629** | female | SAME1840230 | GWD | Gambian Mandinka | AFR | African Ancestry | GWD | 1000 Genomes on GRCh38,1000 Genomes 30x on GRCh38,1000 Genomes phase 3 release,Gambian Genome Variation Project (GRCh38) |
| **HG02653** | female | SAME1839537 | PJL | Punjabi | SAS | South Asian Ancestry | PJL | 1000 Genomes 30x on GRCh38 |
| **HG00418** | male | SAME122992 | CHS | Southern Han Chinese | EAS | East Asian Ancestry | CHS | 1000 Genomes on GRCh38,1000 Genomes 30x on GRCh38,1000 Genomes phase 3 release,1000 Genomes phase 1 release,90 Han Chinese high coverage genomes |
| **HG02636** | male | SAME1840017 | GWD | Gambian Mandinka | AFR | African Ancestry | GWD | 1000 Genomes 30x on GRCh38 |
| **HG02658** | female | SAME1839597 | PJL | Punjabi | SAS | South Asian Ancestry | PJL | 1000 Genomes on GRCh38,1000 Genomes 30x on GRCh38,1000 Genomes phase 3 release |
| **HG00420** | male | SAME122807 | CHS | Southern Han Chinese | EAS | East Asian Ancestry | CHS | 1000 Genomes 30x on GRCh38,1000 Genomes phase 3 release |
| **HG02643** | female | SAME1839785 | GWD | Gambian Mandinka | AFR | African Ancestry | GWD | 1000 Genomes on GRCh38,1000 Genomes 30x on GRCh38,1000 Genomes phase 3 release,Gambian Genome Variation Project (GRCh38) |
| **HG02660** | male | SAME1839672 | PJL | Punjabi | SAS | South Asian Ancestry | PJL | 1000 Genomes on GRCh38,1000 Genomes 30x on GRCh38,1000 Genomes phase 3 release |
| **HG00437** | female | SAME125261 | CHS | Southern Han Chinese | EAS | East Asian Ancestry | CHS | 1000 Genomes on GRCh38,1000 Genomes 30x on GRCh38,1000 Genomes phase 3 release,1000 Genomes phase 1 release,90 Han Chinese high coverage genomes |
| **HG02648** | male | SAME1839751 | PJL | Punjabi | SAS | South Asian Ancestry | PJL | 1000 Genomes on GRCh38,1000 Genomes 30x on GRCh38,1000 Genomes phase 3 release |
| **HG02677** | female | SAME1839483 | GWD | Gambian Mandinka | AFR | African Ancestry | GWD | 1000 Genomes 30x on GRCh38 |
| **HG00444** | male | SAME124190 | CHS | Southern Han Chinese | EAS | East Asian Ancestry | CHS | 1000 Genomes 30x on GRCh38,1000 Genomes phase 3 release |
| **HG02650** | male | SAME1839550 | PJL | Punjabi | SAS | South Asian Ancestry | PJL | 1000 Genomes 30x on GRCh38 |
| **HG02684** | male | SAME1839292 | PJL | Punjabi | SAS | South Asian Ancestry | PJL | 1000 Genomes on GRCh38,1000 Genomes 30x on GRCh38,1000 Genomes phase 3 release |
| **HG00449** | female | SAME123885 | CHS | Southern Han Chinese | EAS | East Asian Ancestry | CHS | 1000 Genomes on GRCh38,1000 Genomes 30x on GRCh38,1000 Genomes phase 3 release,1000 Genomes phase 1 release |
| **HG02655** | female | SAME1839526 | PJL | Punjabi | SAS | South Asian Ancestry | PJL | 1000 Genomes on GRCh38,1000 Genomes 30x on GRCh38,1000 Genomes phase 3 release |
| **HG02502** | female | SAME1839272 | ACB | African Caribbean | AFR | African Ancestry | ACB | 1000 Genomes on GRCh38,1000 Genomes 30x on GRCh38,1000 Genomes phase 3 release |
| **HG00451** | male | SAME125159 | CHS | Southern Han Chinese | EAS | East Asian Ancestry | CHS | 1000 Genomes on GRCh38,1000 Genomes 30x on GRCh38,1000 Genomes phase 3 release,1000 Genomes phase 1 release |
| **HG02514** | male | SAME1839133 | KHV | Kinh Vietnamese | EAS | East Asian Ancestry | KHV | 1000 Genomes 30x on GRCh38,1000 Genomes phase 3 release |
| **HG00463** | male | SAME123496 | CHS | Southern Han Chinese | EAS | East Asian Ancestry | CHS | 1000 Genomes on GRCh38,1000 Genomes 30x on GRCh38,1000 Genomes phase 3 release,1000 Genomes phase 1 release |
| **HG02561** | male | SAME1839838 | GWD | Gambian Mandinka | AFR | African Ancestry | GWD | 1000 Genomes on GRCh38,1000 Genomes 30x on GRCh38,1000 Genomes phase 3 release,Gambian Genome Variation Project (GRCh38) |
| **HG02521** | male | SAME1840327 | KHV | Kinh Vietnamese | EAS | East Asian Ancestry | KHV | 1000 Genomes on GRCh38,1000 Genomes 30x on GRCh38,1000 Genomes phase 3 release |
| **HG00639** | female | SAME123650 | PUR | Puerto Rican | AMR | American Ancestry | PUR | 1000 Genomes 30x on GRCh38,1000 Genomes phase 3 release |
| **HG02573** | male | SAME1839567 | GWD | Gambian Mandinka | AFR | African Ancestry | GWD | 1000 Genomes on GRCh38,1000 Genomes 30x on GRCh38,1000 Genomes phase 3 release,Gambian Genome Variation Project (GRCh38) |
| **HG02526** | female | SAME1840308 | KHV | Kinh Vietnamese | EAS | East Asian Ancestry | KHV | 1000 Genomes 30x on GRCh38 |
| **HG00641** | female | SAME123435 | PUR | Puerto Rican | AMR | American Ancestry | PUR | 1000 Genomes on GRCh38,1000 Genomes 30x on GRCh38,1000 Genomes phase 3 release,1000 Genomes phase 1 release |
| **HG02580** | female | SAME1839366 | ACB | African Caribbean | AFR | African Ancestry | ACB | 1000 Genomes on GRCh38,1000 Genomes 30x on GRCh38,1000 Genomes phase 3 release |
| **HG02545** | male | SAME1839939 | ACB | African Caribbean | AFR | African Ancestry | ACB | 1000 Genomes on GRCh38,1000 Genomes 30x on GRCh38,1000 Genomes phase 3 release |
| **HG00653** | male | SAME123251 | CHS | Southern Han Chinese | EAS | East Asian Ancestry | CHS | 1000 Genomes on GRCh38,1000 Genomes 30x on GRCh38,1000 Genomes phase 3 release,1000 Genomes phase 1 release,90 Han Chinese high coverage genomes |
| **HG02585** | male | SAME1839418 | GWD | Gambian Mandinka | AFR | African Ancestry | GWD | 1000 Genomes on GRCh38,1000 Genomes 30x on GRCh38,1000 Genomes phase 3 release,Gambian Genome Variation Project (GRCh38) |
| **HG02470** | male | SAME1839744 | ACB | African Caribbean | AFR | African Ancestry | ACB | 1000 Genomes on GRCh38,1000 Genomes 30x on GRCh38,1000 Genomes phase 3 release |
| **HG00658** | male | SAME123241 | CHS | Southern Han Chinese | EAS | East Asian Ancestry | CHS | 1000 Genomes 30x on GRCh38,1000 Genomes phase 3 release |
| **HG02597** | male | SAME1839199 | PJL | Punjabi | SAS | South Asian Ancestry | PJL | 1000 Genomes on GRCh38,1000 Genomes 30x on GRCh38,1000 Genomes phase 3 release |
| **HG02494** | female | SAME1839698 | PJL | Punjabi,Punjabi | SAS | South Asia (SGDP),South Asian Ancestry | PunjabiSGDP,PJL | 1000 Genomes on GRCh38,Simons Genome Diversity Project,1000 Genomes 30x on GRCh38,1000 Genomes phase 3 release |
| **HG00672** | female | SAME123606 | CHS | Southern Han Chinese | EAS | East Asian Ancestry | CHS | 1000 Genomes on GRCh38,1000 Genomes 30x on GRCh38,1000 Genomes phase 3 release,1000 Genomes phase 1 release,90 Han Chinese high coverage genomes |
| **HG02600** | male | SAME1840302 | PJL | Punjabi | SAS | South Asian Ancestry | PJL | 1000 Genomes on GRCh38,1000 Genomes 30x on GRCh38,1000 Genomes phase 3 release |
| **HG02386** | male | SAME1840001 | CDX | Dai Chinese | EAS | East Asian Ancestry | CDX | 1000 Genomes on GRCh38,1000 Genomes 30x on GRCh38,1000 Genomes phase 3 release |
| **HG00684** | female | SAME123890 | CHS | Southern Han Chinese | EAS | East Asian Ancestry | CHS | 1000 Genomes on GRCh38,1000 Genomes 30x on GRCh38,1000 Genomes phase 3 release,1000 Genomes phase 1 release,90 Han Chinese high coverage genomes |
| **HG02662** | female | SAME1839662 | PJL | Punjabi | SAS | South Asian Ancestry | PJL | 1000 Genomes 30x on GRCh38,1000 Genomes phase 3 release |
| **HG02398** | male | SAME1839845 | CDX | Dai Chinese | EAS | East Asian Ancestry | CDX | 1000 Genomes on GRCh38,1000 Genomes 30x on GRCh38,1000 Genomes phase 3 release |
| **HG00689** | male | SAME123893 | CHS | Southern Han Chinese | EAS | East Asian Ancestry | CHS | 1000 Genomes on GRCh38,1000 Genomes 30x on GRCh38,1000 Genomes phase 3 release,1000 Genomes phase 1 release |
| **HG02667** | female | SAME1839384 | GWD | Gambian Mandinka | AFR | African Ancestry | GWD | 1000 Genomes on GRCh38,1000 Genomes 30x on GRCh38,1000 Genomes phase 3 release,Gambian Genome Variation Project (GRCh38) |
| **HG02401** | male | SAME1839415 | CDX | Dai Chinese | EAS | East Asian Ancestry | CDX | 1000 Genomes on GRCh38,1000 Genomes 30x on GRCh38,1000 Genomes phase 3 release |
| **HG00691** | male | SAME123161 | CHS | Southern Han Chinese | EAS | East Asian Ancestry | CHS | 1000 Genomes 30x on GRCh38,1000 Genomes phase 3 release |
| **HG02679** | female | SAME1839471 | GWD | Gambian Mandinka | AFR | African Ancestry | GWD | 1000 Genomes on GRCh38,1000 Genomes 30x on GRCh38,1000 Genomes phase 3 release,Gambian Genome Variation Project (GRCh38) |
| **HG02406** | male | SAME1839394 | CDX | Dai Chinese | EAS | East Asian Ancestry | CDX | 1000 Genomes on GRCh38,1000 Genomes 30x on GRCh38,1000 Genomes phase 3 release |
| **HG00867** | female | SAME123550 | CDX | Dai Chinese | EAS | East Asian Ancestry | CDX | 1000 Genomes on GRCh38,1000 Genomes 30x on GRCh38,1000 Genomes phase 3 release |
| **HG02681** | male | SAME1839304 | PJL | Punjabi | SAS | South Asian Ancestry | PJL | 1000 Genomes on GRCh38,1000 Genomes 30x on GRCh38,1000 Genomes phase 3 release |
| **HG02785** | female | SAME1840298 | PJL | Punjabi | SAS | South Asian Ancestry | PJL | 1000 Genomes 30x on GRCh38,1000 Genomes phase 3 release |
| **HG00879** | female | SAME123399 | CDX | Dai Chinese | EAS | East Asian Ancestry | CDX | 1000 Genomes on GRCh38,1000 Genomes 30x on GRCh38,1000 Genomes phase 3 release |

| | | | | | | | | |
|---|---|---|---|---|---|---|---|---|
| **HG02686** | male | SAME1839277 | PJL | Punjabi | SAS | South Asian Ancestry | PJL | 1000 Genomes 30x on GRCh38,1000 Genomes phase 3 release |
| **HG02792** | male | SAME1840416 | PJL | Punjabi | SAS | South Asian Ancestry | PJL | 1000 Genomes on GRCh38,1000 Genomes 30x on GRCh38,1000 Genomes phase 3 release |
| **HG00881** | male | SAME123540 | CDX | Dai Chinese | EAS | East Asian Ancestry | CDX | 1000 Genomes on GRCh38,1000 Genomes 30x on GRCh38,1000 Genomes phase 3 release |
| **HG02800** | male | SAME1840000 | GWD | Gambian Mandinka | AFR | African Ancestry | GWD | 1000 Genomes 30x on GRCh38 |
| **HG00956** | female | SAME124319 | CDX | Dai Chinese | EAS | East Asian Ancestry | CDX | 1000 Genomes on GRCh38,1000 Genomes 30x on GRCh38,1000 Genomes phase 3 release |
| **HG02427** | female | SAME1839036 | ACB | African Caribbean | AFR | African Ancestry | ACB | 1000 Genomes on GRCh38,1000 Genomes 30x on GRCh38,1000 Genomes phase 3 release |
| **HG02805** | female | SAME1840081 | GWD | Gambian Mandinka | AFR | African Ancestry | GWD | 1000 Genomes on GRCh38,1000 Genomes 30x on GRCh38,1000 Genomes phase 3 release,Gambian Genome Variation Project (GRCh38) |
| **HG00982** | male | SAME123770 | CDX | Dai Chinese | EAS | East Asian Ancestry | CDX | 1000 Genomes on GRCh38,1000 Genomes 30x on GRCh38,1000 Genomes phase 3 release |
| **HG02439** | male | SAME1840297 | ACB | African Caribbean | AFR | African Ancestry | ACB | 1000 Genomes on GRCh38,1000 Genomes 30x on GRCh38,1000 Genomes phase 3 release |
| **HG02812** | male | SAME1839846 | GWD | Gambian Mandinka | AFR | African Ancestry | GWD | 1000 Genomes 30x on GRCh38 |
| **HG01029** | female | SAME123523 | CDX | Dai Chinese | EAS | East Asian Ancestry | CDX | 1000 Genomes on GRCh38,1000 Genomes 30x on GRCh38,1000 Genomes phase 3 release |
| **HG02465** | female | SAME1839973 | GWD | Gambian Mandinka | AFR | African Ancestry | GWD | 1000 Genomes on GRCh38,1000 Genomes 30x on GRCh38,1000 Genomes phase 3 release,Gambian Genome Variation Project (GRCh38) |
| **HG02817** | female | SAME1839819 | GWD | Gambian Mandinka | AFR | African Ancestry | GWD | 1000 Genomes on GRCh38,1000 Genomes 30x on GRCh38,1000 Genomes phase 3 release,Gambian Genome Variation Project (GRCh38) |
| **HG01031** | male | SAME123728 | CDX | Dai Chinese | EAS | East Asian Ancestry | CDX | 1000 Genomes on GRCh38,1000 Genomes 30x on GRCh38,1000 Genomes phase 3 release |
| **HG02477** | female | SAME1839710 | ACB | African Caribbean | AFR | African Ancestry | ACB | 1000 Genomes on GRCh38,1000 Genomes 30x on GRCh38,1000 Genomes phase 3 release |
| **HG02735** | male | SAME1839866 | PJL | Punjabi | SAS | South Asian Ancestry | PJL | 1000 Genomes 30x on GRCh38,1000 Genomes phase 3 release |
| **HG01048** | male | SAME123169 | PUR | Puerto Rican | AMR | American Ancestry | PUR | 1000 Genomes on GRCh38,1000 Genomes 30x on GRCh38,1000 Genomes phase 3 release,1000 Genomes phase 1 release |
| **HG02484** | male | SAME1839858 | ACB | African Caribbean | AFR | African Ancestry | ACB | 1000 Genomes on GRCh38,1000 Genomes 30x on GRCh38,1000 Genomes phase 3 release |
| **HG02759** | male | SAME1839445 | GWD | Gambian Mandinka | AFR | African Ancestry | GWD | 1000 Genomes on GRCh38,1000 Genomes 30x on GRCh38,1000 Genomes phase 3 release,Gambian Genome Variation Project (GRCh38) |
| **HG01050** | female | SAME123180 | PUR | Puerto Rican | AMR | American Ancestry | PUR | 1000 Genomes 30x on GRCh38,1000 Genomes phase 3 release |
| **HG02489** | male | SAME1839546 | ACB | African Caribbean | AFR | African Ancestry | ACB | 1000 Genomes on GRCh38,1000 Genomes 30x on GRCh38,1000 Genomes phase 3 release |
| **HG02761** | female | SAME1839262 | GWD | Gambian Mandinka | AFR | African Ancestry | GWD | 1000 Genomes 30x on GRCh38 |
| **HG01055** | female | SAME123638 | PUR | Puerto Rican | AMR | American Ancestry | PUR | 1000 Genomes on GRCh38,1000 Genomes 30x on GRCh38,1000 Genomes phase 3 release,1000 Genomes phase 1 release |
| **HG02491** | female | SAME1839617 | PJL | Punjabi | SAS | South Asian Ancestry | PJL | 1000 Genomes on GRCh38,1000 Genomes 30x on GRCh38,1000 Genomes phase 3 release |
| **HG02773** | male | SAME1839034 | GWD | Gambian Mandinka | AFR | African Ancestry | GWD | 1000 Genomes 30x on GRCh38 |
| **HG01062** | female | SAME123014 | PUR | Puerto Rican | AMR | American Ancestry | PUR | 1000 Genomes 30x on GRCh38,1000 Genomes phase 3 release |
| **HG02496** | male | SAME1839688 | ACB | African Caribbean | AFR | African Ancestry | ACB | 1000 Genomes on GRCh38,1000 Genomes 30x on GRCh38,1000 Genomes phase 3 release |
| **HG02778** | female | SAME1839100 | PJL | Punjabi | SAS | South Asian Ancestry | PJL | 1000 Genomes on GRCh38,1000 Genomes 30x on GRCh38,1000 Genomes phase 3 release |
| **HG00704** | male | SAME122928 | CHS | Southern Han Chinese | EAS | East Asian Ancestry | CHS | 1000 Genomes on GRCh38,1000 Genomes 30x on GRCh38,1000 Genomes phase 3 release,1000 Genomes phase 1 release |
| **HG02511** | female | SAME1839150 | ACB | African Caribbean | AFR | African Ancestry | ACB | 1000 Genomes on GRCh38,1000 Genomes 30x on GRCh38,1000 Genomes phase 3 release |
| **HG02780** | male | SAME1839169 | PJL | Punjabi | SAS | South Asian Ancestry | PJL | 1000 Genomes on GRCh38,1000 Genomes 30x on GRCh38,1000 Genomes phase 3 release |
| **HG00709** | male | SAME125334 | CHS | Southern Han Chinese | EAS | East Asian Ancestry | CHS | 1000 Genomes 30x on GRCh38,1000 Genomes phase 3 release |
| **HG02523** | female | SAME1840322 | KHV | Kinh Vietnamese | EAS | East Asian Ancestry | KHV | 1000 Genomes 30x on GRCh38,1000 Genomes phase 3 release |
| **HG02975** | male | SAME1840232 | ESN | Esan | AFR | African Ancestry | ESN | 1000 Genomes 30x on GRCh38 |
| **HG02982** | male | SAME1840015 | GWD | Gambian Mandinka | AFR | African Ancestry | GWD | 1000 Genomes on GRCh38,1000 Genomes 30x on GRCh38,1000 Genomes phase 3 release,Gambian Genome Variation Project (GRCh38) |
| **HG00728** | male | SAME1840267 | CHS | Southern Han Chinese | EAS | East Asian Ancestry | CHS | 1000 Genomes on GRCh38,1000 Genomes 30x on GRCh38,1000 Genomes phase 3 release |
| **HG02554** | male | SAME1840075 | ACB | African Caribbean | AFR | African Ancestry | ACB | 1000 Genomes on GRCh38,1000 Genomes 30x on GRCh38,1000 Genomes phase 3 release |
| **HG03012** | male | SAME1839924 | BEB | Bengali | SAS | South Asian Ancestry | BEB | 1000 Genomes on GRCh38,1000 Genomes 30x on GRCh38,1000 Genomes phase 3 release |
| **HG03017** | male | SAME1839956 | PJL | Punjabi | SAS | South Asian Ancestry | PJL | 1000 Genomes 30x on GRCh38 |
| **HG00735** | female | SAME123042 | PUR | Puerto Rican | AMR | American Ancestry | PUR | 1000 Genomes 30x on GRCh38,1000 Genomes phase 3 release |
| **HG02390** | male | SAME1839788 | CDX | Dai Chinese | EAS | East Asian Ancestry | CDX | 1000 Genomes on GRCh38,1000 Genomes 30x on GRCh38,1000 Genomes phase 3 release |
| **HG02836** | male | SAME1839428 | GWD | Gambian Mandinka | AFR | African Ancestry | GWD | 1000 Genomes on GRCh38,1000 Genomes 30x on GRCh38,1000 Genomes phase 3 release,Gambian Genome Variation Project (GRCh38) |
| **HG00742** | male | SAME1840188 | PUR | Puerto Rican | AMR | American Ancestry | PUR | 1000 Genomes on GRCh38,1000 Genomes 30x on GRCh38,1000 Genomes phase 3 release |
| **HG02395** | male | SAME1839754 | CDX | Dai Chinese | EAS | East Asian Ancestry | CDX | 1000 Genomes on GRCh38,1000 Genomes 30x on GRCh38,1000 Genomes phase 3 release |
| **HG02855** | female | SAME1839330 | GWD | Gambian Mandinka | AFR | African Ancestry | GWD | 1000 Genomes on GRCh38,1000 Genomes 30x on GRCh38,1000 Genomes phase 3 release,Gambian Genome Variation Project (GRCh38) |
| **HG00759** | female | SAME125020 | CDX | Dai Chinese | EAS | East Asian Ancestry | CDX | 1000 Genomes on GRCh38,1000 Genomes 30x on GRCh38,1000 Genomes phase 3 release |
| **HG02408** | male | SAME1839461 | CDX | Dai Chinese | EAS | East Asian Ancestry | CDX | 1000 Genomes on GRCh38,1000 Genomes 30x on GRCh38,1000 Genomes phase 3 release |
| **HG02862** | female | SAME1839123 | GWD | Gambian Mandinka | AFR | African Ancestry | GWD | 1000 Genomes 30x on GRCh38 |
| **HG00766** | female | SAME124458 | CDX | Dai Chinese | EAS | East Asian Ancestry | CDX | 1000 Genomes on GRCh38,1000 Genomes 30x on GRCh38,1000 Genomes phase 3 release |
| **HG02410** | male | SAME1839207 | CDX | Dai Chinese | EAS | East Asian Ancestry | CDX | 1000 Genomes on GRCh38,1000 Genomes 30x on GRCh38,1000 Genomes phase 3 release |
| **HG02879** | female | SAME1840355 | GWD | Gambian Mandinka | AFR | African Ancestry | GWD | 1000 Genomes on GRCh38,1000 Genomes 30x on GRCh38,1000 Genomes phase 3 release,Gambian Genome Variation Project (GRCh38) |
| **HG00590** | female | SAME125009 | CHS | Southern Han Chinese | EAS | East Asian Ancestry | CHS | 1000 Genomes on GRCh38,1000 Genomes 30x on GRCh38,1000 Genomes phase 3 release,1000 Genomes phase 1 release |
| **HG02314** | male | SAME1839674 | ACB | African Caribbean | AFR | African Ancestry | ACB | 1000 Genomes on GRCh38,1000 Genomes 30x on GRCh38,1000 Genomes phase 3 release |
| **HG02881** | male | SAME1840178 | GWD | Gambian Mandinka | AFR | African Ancestry | GWD | 1000 Genomes on GRCh38,1000 Genomes 30x on GRCh38,1000 Genomes phase 3 release,Gambian Genome Variation Project (GRCh38) |
| **HG00595** | male | SAME125006 | CHS | Southern Han Chinese | EAS | East Asian Ancestry | CHS | 1000 Genomes on GRCh38,1000 Genomes 30x on GRCh38,1000 Genomes phase 3 release,1000 Genomes phase 1 release |
| **HG02321** | female | SAME1839446 | ACB | African Caribbean | AFR | African Ancestry | ACB | 1000 Genomes 30x on GRCh38,1000 Genomes phase 3 release |
| **HG02886** | female | SAME1840151 | GWD | Gambian Mandinka | AFR | African Ancestry | GWD | 1000 Genomes 30x on GRCh38 |
| **HG00608** | female | SAME125369 | CHS | Southern Han Chinese | EAS | East Asian Ancestry | CHS | 1000 Genomes on GRCh38,1000 Genomes 30x on GRCh38,1000 Genomes phase 3 release,1000 Genomes phase 1 release |
| **HG02345** | female | SAME1839081 | PEL | Peruvian | AMR | American Ancestry | PEL | 1000 Genomes on GRCh38,1000 Genomes 30x on GRCh38,1000 Genomes phase 3 release |
| **HG03024** | male | SAME1839799 | GWD | Gambian Mandinka | AFR | African Ancestry | GWD | 1000 Genomes on GRCh38,1000 Genomes 30x on GRCh38,1000 Genomes phase 3 release,Gambian Genome Variation Project (GRCh38) |
| **HG00610** | male | SAME125199 | CHS | Southern Han Chinese | EAS | East Asian Ancestry | CHS | 1000 Genomes on GRCh38,1000 Genomes 30x on GRCh38,1000 Genomes phase 3 release,1000 Genomes phase 1 release |
| **HG02364** | male | SAME1840125 | CDX | Dai Chinese | EAS | East Asian Ancestry | CDX | 1000 Genomes on GRCh38,1000 Genomes 30x on GRCh38,1000 Genomes phase 3 release |
| **HG03029** | male | SAME1839733 | GWD | Gambian Mandinka | AFR | African Ancestry | GWD | 1000 Genomes 30x on GRCh38 |
| **HG00615** | female | SAME125194 | CHS | Southern Han Chinese | EAS | East Asian Ancestry | CHS | 1000 Genomes 30x on GRCh38,1000 Genomes phase 3 release |
| **HG02371** | male | SAME1840180 | CDX | Dai Chinese | EAS | East Asian Ancestry | CDX | 1000 Genomes on GRCh38,1000 Genomes 30x on GRCh38,1000 Genomes phase 3 release |
| **HG03048** | male | SAME1839647 | GWD | Gambian Mandinka | AFR | African Ancestry | GWD | 1000 Genomes on GRCh38,1000 Genomes 30x on GRCh38,1000 Genomes phase 3 release,Gambian Genome Variation Project (GRCh38) |
| **HG00622** | male | SAME1839278 | CHS | Southern Han Chinese | EAS | East Asian Ancestry | CHS | 1000 Genomes on GRCh38,1000 Genomes 30x on GRCh38,1000 Genomes phase 3 release |
| **HG02383** | male | SAME1840019 | CDX | Dai Chinese | EAS | East Asian Ancestry | CDX | 1000 Genomes on GRCh38,1000 Genomes 30x on GRCh38,1000 Genomes phase 3 release |
| **HG03050** | male | SAME1839482 | GWD | Gambian Mandinka | AFR | African Ancestry | GWD | 1000 Genomes 30x on GRCh38 |
| **HG00627** | male | SAME123079 | CHS | Southern Han Chinese | EAS | East Asian Ancestry | CHS | 1000 Genomes 30x on GRCh38,1000 Genomes phase 3 release |
| **HG02819** | male | SAME1839804 | GWD | Gambian Mandinka | AFR | African Ancestry | GWD | 1000 Genomes on GRCh38,1000 Genomes 30x on GRCh38,1000 Genomes phase 3 release,Gambian Genome Variation Project (GRCh38) |
| **HG03055** | female | SAME1839514 | MSL | Mende | AFR | African Ancestry | MSL | 1000 Genomes on GRCh38,1000 Genomes 30x on GRCh38,1000 Genomes phase 3 release |
| **HG00634** | male | SAME122903 | CHS | Southern Han Chinese | EAS | East Asian Ancestry | CHS | 1000 Genomes on GRCh38,1000 Genomes 30x on GRCh38,1000 Genomes phase 3 release,1000 Genomes phase 1 release |
| **HG02821** | male | SAME1839588 | GWD | Gambian Mandinka | AFR | African Ancestry | GWD | 1000 Genomes 30x on GRCh38 |
| **HG03098** | male | SAME1840136 | MSL | Mende | AFR | African Ancestry | MSL | 1000 Genomes 30x on GRCh38 |
| **HG01170** | male | SAME124623 | PUR | Puerto Rican | AMR | American Ancestry | PUR | 1000 Genomes on GRCh38,1000 Genomes 30x on GRCh38,1000 Genomes phase 3 release,1000 Genomes phase 1 release |
| **HG02838** | male | SAME1839417 | GWD | Gambian Mandinka | AFR | African Ancestry | GWD | 1000 Genomes 30x on GRCh38 |
| **HG03101** | male | SAME1840008 | ESN | Esan | AFR | African Ancestry | ESN | 1000 Genomes 30x on GRCh38 |
| **HG01175** | female | SAME124618 | PUR | Puerto Rican | AMR | American Ancestry | PUR | 1000 Genomes 30x on GRCh38,1000 Genomes phase 3 release |
| **HG02840** | female | SAME1839475 | GWD | Gambian Mandinka | AFR | African Ancestry | GWD | 1000 Genomes on GRCh38,1000 Genomes 30x on GRCh38,1000 Genomes phase 3 release,Gambian Genome Variation Project (GRCh38) |
| **HG03113** | female | SAME1839901 | ESN | Esan | AFR | African Ancestry | ESN | 1000 Genomes 30x on GRCh38 |
| **HG01182** | male | SAME124789 | PUR | Puerto Rican | AMR | American Ancestry | PUR | 1000 Genomes on GRCh38,1000 Genomes 30x on GRCh38,1000 Genomes phase 3 release |
| **HG02852** | female | SAME1839346 | GWD | Gambian Mandinka | AFR | African Ancestry | GWD | 1000 Genomes on GRCh38,1000 Genomes 30x on GRCh38,1000 Genomes phase 3 release,Gambian Genome Variation Project (GRCh38) |
| **HG03118** | male | SAME1839830 | ESN | Esan | AFR | African Ancestry | ESN | 1000 Genomes on GRCh38,1000 Genomes 30x on GRCh38,1000 Genomes phase 3 release |
| **HG01187** | male | SAME124783 | PUR | Puerto Rican | AMR | American Ancestry | PUR | 1000 Genomes on GRCh38,1000 Genomes 30x on GRCh38,1000 Genomes phase 3 release,1000 Genomes phase 1 release |
| **HG02869** | male | SAME1839184 | GWD | Gambian Mandinka | AFR | African Ancestry | GWD | 1000 Genomes on GRCh38,1000 Genomes 30x on GRCh38,Gambian Genome Variation Project (GRCh38) |
| **HG03120** | male | SAME1839557 | ESN | Esan | AFR | African Ancestry | ESN | 1000 Genomes on GRCh38,1000 Genomes 30x on GRCh38,1000 Genomes phase 3 release |
| **HG01199** | male | SAME124366 | PUR | Puerto Rican | AMR | American Ancestry | PUR | 1000 Genomes 30x on GRCh38,1000 Genomes phase 3 release |
| **HG02871** | female | SAME1840312 | GWD | Gambian Mandinka | AFR | African Ancestry | GWD | 1000 Genomes 30x on GRCh38 |
| **HG03125** | female | SAME1839675 | ESN | Esan | AFR | African Ancestry | ESN | 1000 Genomes 30x on GRCh38,Human Genome Structural Variation Consortium, Phase 2 |
| **HG01252** | male | SAME125283 | CLM | Colombian | AMR | American Ancestry | CLM | 1000 Genomes 30x on GRCh38,1000 Genomes phase 3 release |
| **HG02883** | male | SAME1840164 | GWD | Gambian Mandinka | AFR | African Ancestry | GWD | 1000 Genomes 30x on GRCh38 |
| **HG03132** | female | SAME1839373 | ESN | Esan | AFR | African Ancestry | ESN | 1000 Genomes on GRCh38,1000 Genomes 30x on GRCh38,1000 Genomes phase 3 release |
| **HG01257** | female | SAME125288 | CLM | Colombian | AMR | American Ancestry | CLM | 1000 Genomes on GRCh38,1000 Genomes 30x on GRCh38,1000 Genomes phase 3 release,1000 Genomes phase 1 release |
| **HG02888** | female | SAME1840139 | GWD | Gambian Mandinka | AFR | African Ancestry | GWD | 1000 Genomes on GRCh38,1000 Genomes 30x on GRCh38,1000 Genomes phase 3 release,Gambian Genome Variation Project (GRCh38) |
| **HG03137** | male | SAME1839390 | ESN | Esan | AFR | African Ancestry | ESN | 1000 Genomes 30x on GRCh38 |
| **HG01067** | female | SAME122997 | PUR | Puerto Rican | AMR | American Ancestry | PUR | 1000 Genomes on GRCh38,1000 Genomes 30x on GRCh38,1000 Genomes phase 3 release,1000 Genomes phase 1 release |
| **HG02890** | male | SAME1839947 | GWD | Gambian Mandinka | AFR | African Ancestry | GWD | 1000 Genomes on GRCh38,1000 Genomes 30x on GRCh38,1000 Genomes phase 3 release,Gambian Genome Variation Project (GRCh38) |
| **HG01074** | male | SAME124070 | PUR | Puerto Rican | AMR | American Ancestry | PUR | 1000 Genomes 30x on GRCh38,1000 Genomes phase 3 release |
| **HG02895** | male | SAME1839918 | GWD | Gambian Mandinka | AFR | African Ancestry | GWD | 1000 Genomes on GRCh38,1000 Genomes 30x on GRCh38,1000 Genomes phase 3 release,Gambian Genome Variation Project (GRCh38) |
| **HG03074** | male | SAME1839029 | MSL | Mende | AFR | African Ancestry | MSL | 1000 Genomes on GRCh38,1000 Genomes 30x on GRCh38,1000 Genomes phase 3 release |
| **HG01079** | male | SAME123013 | PUR | Puerto Rican | AMR | American Ancestry | PUR | 1000 Genomes on GRCh38,1000 Genomes 30x on GRCh38,1000 Genomes phase 3 release,1000 Genomes phase 1 release |
| **HG02922** | female | SAME1839441 | ESN | Esan | AFR | African Ancestry | ESN | 1000 Genomes on GRCh38,1000 Genomes 30x on GRCh38,1000 Genomes phase 3 release |
| **HG03079** | female | SAME1839051 | MSL | Mende | AFR | African Ancestry | MSL | 1000 Genomes on GRCh38,1000 Genomes 30x on GRCh38,1000 Genomes phase 3 release |
| **HG01081** | female | SAME125395 | PUR | Puerto Rican | AMR | American Ancestry | PUR | 1000 Genomes 30x on GRCh38,1000 Genomes phase 3 release |
| **HG02972** | male | SAME1840250 | ESN | Esan | AFR | African Ancestry | ESN | 1000 Genomes 30x on GRCh38 |
| **HG03081** | male | SAME1840310 | MSL | Mende | AFR | African Ancestry | MSL | 1000 Genomes on GRCh38,1000 Genomes 30x on GRCh38,1000 Genomes phase 3 release |
| **HG01086** | female | SAME125394 | PUR | Puerto Rican | AMR | American Ancestry | PUR | 1000 Genomes on GRCh38,1000 Genomes 30x on GRCh38,1000 Genomes phase 3 release,1000 Genomes phase 1 release |
| **HG02977** | male | SAME1840221 | ESN | Esan | AFR | African Ancestry | ESN | 1000 Genomes on GRCh38,1000 Genomes 30x on GRCh38,1000 Genomes phase 3 release |
| **HG03086** | female | SAME1840331 | MSL | Mende | AFR | African Ancestry | MSL | 1000 Genomes on GRCh38,1000 Genomes 30x on GRCh38,1000 Genomes phase 3 release |
| **HG01098** | female | SAME123490 | PUR | Puerto Rican | AMR | American Ancestry | PUR | 1000 Genomes on GRCh38,1000 Genomes 30x on GRCh38,1000 Genomes phase 3 release,1000 Genomes phase 1 release |

| | | | | | | | | |
|---|---|---|---|---|---|---|---|---|
| **HG02984** | male | SAME1840004 | GWD | Gambian Mandinka | AFR | African Ancestry | GWD | 1000 Genomes 30x on GRCh38 |
| **HG02944** | male | SAME1839082 | ESN | Esan | AFR | African Ancestry | ESN | 1000 Genomes on GRCh38,1000 Genomes 30x on GRCh38,1000 Genomes phase 3 release |
| **HG01101** | male | SAME123677 | PUR | Puerto Rican | AMR | American Ancestry | PUR | 1000 Genomes on GRCh38,1000 Genomes 30x on GRCh38,1000 Genomes phase 3 release,1000 Genomes phase 1 release |
| **HG03007** | female | SAME1840245 | BEB | Bengali,Bengali | SAS | South Asia (SGDP),South Asian Ancestry | BengaliSGDP,BEB | 1000 Genomes on GRCh38,Simons Genome Diversity Project,1000 Genomes 30x on GRCh38,1000 Genomes phase 3 release |
| **HG02968** | male | SAME1840108 | ESN | Esan | AFR | African Ancestry | ESN | 1000 Genomes on GRCh38,1000 Genomes 30x on GRCh38,1000 Genomes phase 3 release |
| **HG01106** | male | SAME123674 | PUR | Puerto Rican | AMR | American Ancestry | PUR | 1000 Genomes 30x on GRCh38,1000 Genomes phase 3 release |
| **HG03019** | female | SAME1840060 | PJL | Punjabi | SAS | South Asian Ancestry | PJL | 1000 Genomes on GRCh38,1000 Genomes 30x on GRCh38,1000 Genomes phase 3 release |
| **HG02970** | female | SAME1840260 | ESN | Esan | AFR | African Ancestry | ESN | 1000 Genomes on GRCh38,1000 Genomes 30x on GRCh38,1000 Genomes phase 3 release |
| **HG01113** | female | SAME123484 | CLM | Colombian | AMR | American Ancestry | CLM | 1000 Genomes on GRCh38,1000 Genomes 30x on GRCh38,1000 Genomes phase 3 release,1000 Genomes phase 1 release |
| **HG03021** | male | SAME1839789 | PJL | Punjabi | SAS | South Asian Ancestry | PJL | 1000 Genomes on GRCh38,1000 Genomes 30x on GRCh38,1000 Genomes phase 3 release |
| **HG01125** | female | SAME124054 | CLM | Colombian | AMR | American Ancestry | CLM | 1000 Genomes on GRCh38,1000 Genomes 30x on GRCh38,1000 Genomes phase 3 release,1000 Genomes phase 1 release |
| **HG03026** | female | SAME1839714 | GWD | Gambian Mandinka | AFR | African Ancestry | GWD | 1000 Genomes 30x on GRCh38 |
| **HG03461** | female | SAME1839808 | MSL | Mende | AFR | African Ancestry | MSL | 1000 Genomes on GRCh38,1000 Genomes 30x on GRCh38,1000 Genomes phase 3 release |
| **HG01137** | female | SAME123888 | CLM | Colombian | AMR | American Ancestry | CLM | 1000 Genomes on GRCh38,1000 Genomes 30x on GRCh38,1000 Genomes phase 3 release,1000 Genomes phase 1 release |
| **HG03033** | male | SAME1839566 | GWD | Gambian Mandinka | AFR | African Ancestry | GWD | 1000 Genomes on GRCh38,1000 Genomes 30x on GRCh38,Gambian Genome Variation Project (GRCh38) |
| **HG01149** | female | SAME124425 | CLM | Colombian | AMR | American Ancestry | CLM | 1000 Genomes on GRCh38,1000 Genomes 30x on GRCh38,1000 Genomes phase 3 release,1000 Genomes phase 1 release |
| **HG03040** | female | SAME1839611 | GWD | Gambian Mandinka | AFR | African Ancestry | GWD | 1000 Genomes on GRCh38,1000 Genomes 30x on GRCh38,1000 Genomes phase 3 release,Gambian Genome Variation Project (GRCh38) |
| **HG03473** | female | SAME1839687 | MSL | Mende | AFR | African Ancestry | MSL | 1000 Genomes on GRCh38,1000 Genomes 30x on GRCh38,1000 Genomes phase 3 release |
| **HG01168** | female | SAME124796 | PUR | Puerto Rican | AMR | American Ancestry | PUR | 1000 Genomes on GRCh38,1000 Genomes 30x on GRCh38,1000 Genomes phase 3 release,1000 Genomes phase 1 release |
| **HG03045** | male | SAME1839635 | GWD | Gambian Mandinka | AFR | African Ancestry | GWD | 1000 Genomes on GRCh38,1000 Genomes 30x on GRCh38,1000 Genomes phase 3 release,Gambian Genome Variation Project (GRCh38) |
| **HG03478** | male | SAME1839618 | MSL | Mende | AFR | African Ancestry | MSL | 1000 Genomes on GRCh38,1000 Genomes 30x on GRCh38,1000 Genomes phase 3 release |
| **HG03052** | female | SAME1839496 | MSL | Mende | AFR | African Ancestry | MSL | 1000 Genomes on GRCh38,1000 Genomes 30x on GRCh38,1000 Genomes phase 3 release |
| **HG03480** | female | SAME1839381 | MSL | Mende | AFR | African Ancestry | MSL | 1000 Genomes 30x on GRCh38 |
| **HG01334** | male | SAME123533 | GBR | British | EUR | European Ancestry | GBR | 1000 Genomes on GRCh38,1000 Genomes 30x on GRCh38,1000 Genomes phase 3 release,1000 Genomes phase 1 release,Geuvadis |
| **HG03057** | male | SAME1839434 | MSL | Mende | AFR | African Ancestry | MSL | 1000 Genomes on GRCh38,1000 Genomes 30x on GRCh38,1000 Genomes phase 3 release |
| **HG03485** | female | SAME1839469 | MSL | Mende | AFR | African Ancestry | MSL | 1000 Genomes on GRCh38,1000 Genomes 30x on GRCh38,1000 Genomes phase 3 release |
| **HG01341** | male | SAME124859 | CLM | Colombian | AMR | American Ancestry | CLM | 1000 Genomes on GRCh38,1000 Genomes 30x on GRCh38,1000 Genomes phase 3 release,1000 Genomes phase 1 release |
| **HG02737** | female | SAME1839857 | PJL | Punjabi | SAS | South Asian Ancestry | PJL | 1000 Genomes on GRCh38,1000 Genomes 30x on GRCh38,1000 Genomes phase 3 release |
| **HG03240** | male | SAME1839050 | GWD | Gambian Mandinka | AFR | African Ancestry | GWD | 1000 Genomes on GRCh38,1000 Genomes 30x on GRCh38,1000 Genomes phase 3 release,Gambian Genome Variation Project (GRCh38) |
| **HG01346** | female | SAME124860 | CLM | Colombian | AMR | American Ancestry | CLM | 1000 Genomes 30x on GRCh38,1000 Genomes phase 3 release |
| **HG02756** | male | SAME1839454 | GWD | Gambian Mandinka | AFR | African Ancestry | GWD | 1000 Genomes on GRCh38,1000 Genomes 30x on GRCh38,1000 Genomes phase 3 release,Gambian Genome Variation Project (GRCh38) |
| **HG03269** | male | SAME1840100 | ESN | Esan | AFR | African Ancestry | ESN | 1000 Genomes 30x on GRCh38 |
| **HG01353** | male | SAME124685 | CLM | Colombian | AMR | American Ancestry | CLM | 1000 Genomes on GRCh38,1000 Genomes 30x on GRCh38,1000 Genomes phase 3 release,1000 Genomes phase 1 release |
| **HG02763** | female | SAME1839254 | GWD | Gambian Mandinka | AFR | African Ancestry | GWD | 1000 Genomes on GRCh38,1000 Genomes 30x on GRCh38,1000 Genomes phase 3 release,Gambian Genome Variation Project (GRCh38) |
| **HG03271** | male | SAME1840195 | ESN | Esan | AFR | African Ancestry | ESN | 1000 Genomes on GRCh38,1000 Genomes 30x on GRCh38,1000 Genomes phase 3 release |
| **HG01358** | male | SAME124688 | CLM | Colombian | AMR | American Ancestry | CLM | 1000 Genomes 30x on GRCh38,1000 Genomes phase 3 release |
| **HG02768** | male | SAME1839235 | GWD | Gambian Mandinka | AFR | African Ancestry | GWD | 1000 Genomes on GRCh38,1000 Genomes 30x on GRCh38,1000 Genomes phase 3 release,Gambian Genome Variation Project (GRCh38) |
| **HG03295** | male | SAME1839881 | ESN | Esan | AFR | African Ancestry | ESN | 1000 Genomes on GRCh38,1000 Genomes 30x on GRCh38,1000 Genomes phase 3 release |
| **HG01360** | female | SAME122855 | CLM | Colombian | AMR | American Ancestry | CLM | 1000 Genomes on GRCh38,1000 Genomes 30x on GRCh38,1000 Genomes phase 3 release,1000 Genomes phase 1 release |
| **HG02770** | female | SAME1839049 | GWD | Gambian Mandinka | AFR | African Ancestry | GWD | 1000 Genomes 30x on GRCh38 |
| **HG03303** | female | SAME1839485 | ESN | Esan | AFR | African Ancestry | ESN | 1000 Genomes on GRCh38,1000 Genomes 30x on GRCh38,1000 Genomes phase 3 release |
| **HG01365** | male | SAME124508 | CLM | Colombian | AMR | American Ancestry | CLM | 1000 Genomes on GRCh38,1000 Genomes 30x on GRCh38,1000 Genomes phase 3 release,1000 Genomes phase 1 release |
| **HG02775** | female | SAME1839022 | PJL | Punjabi | SAS | South Asian Ancestry | PJL | 1000 Genomes on GRCh38,1000 Genomes 30x on GRCh38,1000 Genomes phase 3 release |
| **HG03308** | male | SAME1839515 | ESN | Esan | AFR | African Ancestry | ESN | 1000 Genomes 30x on GRCh38 |
| **HG01372** | female | SAME1840339 | CLM | Colombian | AMR | American Ancestry | CLM | 1000 Genomes on GRCh38,1000 Genomes 30x on GRCh38,1000 Genomes phase 3 release |
| **HG02787** | female | SAME1840288 | PJL | Punjabi | SAS | South Asian Ancestry | PJL | 1000 Genomes on GRCh38,1000 Genomes 30x on GRCh38,1000 Genomes phase 3 release |
| **HG03310** | female | SAME1839202 | ESN | Esan | AFR | African Ancestry | ESN | 1000 Genomes 30x on GRCh38 |
| **HG01377** | male | SAME124288 | CLM | Colombian | AMR | American Ancestry | CLM | 1000 Genomes on GRCh38,1000 Genomes 30x on GRCh38,1000 Genomes phase 3 release,1000 Genomes phase 1 release |
| **HG02794** | male | SAME1840410 | PJL | Punjabi | SAS | South Asian Ancestry | PJL | 1000 Genomes 30x on GRCh38 |
| **HG01529** | male | SAME125330 | IBS | Iberian | EUR | European Ancestry | IBS | 1000 Genomes 30x on GRCh38,1000 Genomes phase 3 release |
| **HG02799** | female | SAME1840384 | GWD | Gambian Mandinka | AFR | African Ancestry | GWD | 1000 Genomes on GRCh38,1000 Genomes 30x on GRCh38,1000 Genomes phase 3 release,Gambian Genome Variation Project (GRCh38) |
| **HG03163** | male | SAME1839118 | ESN | Esan | AFR | African Ancestry | ESN | 1000 Genomes on GRCh38,1000 Genomes 30x on GRCh38,1000 Genomes phase 3 release |
| **HG01531** | female | SAME125136 | IBS | Iberian | EUR | European Ancestry | IBS | 1000 Genomes on GRCh38,1000 Genomes 30x on GRCh38,1000 Genomes phase 3 release |
| **HG02807** | male | SAME1840073 | GWD | Gambian Mandinka | AFR | African Ancestry | GWD | 1000 Genomes on GRCh38,1000 Genomes 30x on GRCh38,1000 Genomes phase 3 release,Gambian Genome Variation Project (GRCh38) |
| **HG03168** | female | SAME1839143 | ESN | Esan | AFR | African Ancestry | ESN | 1000 Genomes on GRCh38,1000 Genomes 30x on GRCh38,1000 Genomes phase 3 release |
| **HG01536** | male | SAME125135 | IBS | Iberian | EUR | European Ancestry | IBS | 1000 Genomes on GRCh38,1000 Genomes 30x on GRCh38,1000 Genomes phase 3 release |
| **HG02814** | female | SAME1839837 | GWD | Gambian Mandinka | AFR | African Ancestry | GWD | 1000 Genomes on GRCh38,1000 Genomes 30x on GRCh38,1000 Genomes phase 3 release,Gambian Genome Variation Project (GRCh38) |
| **HG03170** | male | SAME1840383 | ESN | Esan | AFR | African Ancestry | ESN | 1000 Genomes 30x on GRCh38 |
| **HG01550** | male | SAME124009 | CLM | Colombian | AMR | American Ancestry | CLM | 1000 Genomes on GRCh38,1000 Genomes 30x on GRCh38,1000 Genomes phase 3 release,1000 Genomes phase 1 release |
| **HG03064** | female | SAME1839228 | MSL | Mende | AFR | African Ancestry | MSL | 1000 Genomes on GRCh38,1000 Genomes 30x on GRCh38,1000 Genomes phase 3 release |
| **HG03175** | male | SAME1840407 | ESN | Esan | AFR | African Ancestry | ESN | 1000 Genomes on GRCh38,1000 Genomes 30x on GRCh38,1000 Genomes phase 3 release |
| **HG01567** | female | SAME124565 | PEL | Peruvian | AMR | American Ancestry | PEL | 1000 Genomes 30x on GRCh38,1000 Genomes phase 3 release |
| **HG03069** | male | SAME1839340 | MSL | Mende | AFR | African Ancestry | MSL | 1000 Genomes on GRCh38,1000 Genomes 30x on GRCh38,1000 Genomes phase 3 release |
| **HG01579** | male | SAME124350 | PEL | Peruvian | AMR | American Ancestry | PEL | 1000 Genomes 30x on GRCh38,1000 Genomes phase 3 release |
| **HG03199** | male | SAME1839937 | ESN | Esan | AFR | African Ancestry | ESN | 1000 Genomes on GRCh38,1000 Genomes 30x on GRCh38,1000 Genomes phase 3 release |
| **HG01586** | male | SAME1839879 | PJL | Punjabi | SAS | South Asian Ancestry | PJL | 1000 Genomes on GRCh38,1000 Genomes 30x on GRCh38,1000 Genomes phase 3 release |
| **HG03088** | female | SAME1840338 | MSL | Mende | AFR | African Ancestry | MSL | 1000 Genomes on GRCh38,1000 Genomes 30x on GRCh38,1000 Genomes phase 3 release |
| **HG03202** | male | SAME1839602 | ESN | Esan | AFR | African Ancestry | ESN | 1000 Genomes on GRCh38,1000 Genomes 30x on GRCh38,1000 Genomes phase 3 release |
| **HG01435** | female | SAME1839239 | CLM | Colombian | AMR | American Ancestry | CLM | 1000 Genomes on GRCh38,1000 Genomes 30x on GRCh38,1000 Genomes phase 3 release |
| **HG03095** | female | SAME1840374 | MSL | Mende | AFR | African Ancestry | MSL | 1000 Genomes on GRCh38,1000 Genomes 30x on GRCh38,1000 Genomes phase 3 release |
| **HG03238** | female | SAME1839289 | PJL | Punjabi | SAS | South Asian Ancestry | PJL | 1000 Genomes on GRCh38,1000 Genomes 30x on GRCh38,1000 Genomes phase 3 release |
| **HG03492** | male | SAME1839196 | PJL | Punjabi | SAS | South Asian Ancestry | PJL | 1000 Genomes 30x on GRCh38,1000 Genomes phase 3 release |
| **HG01447** | female | SAME1839099 | CLM | Colombian | AMR | American Ancestry | CLM | 1000 Genomes on GRCh38,1000 Genomes 30x on GRCh38,1000 Genomes phase 3 release |
| **HG02941** | male | SAME1839096 | ESN | Esan | AFR | African Ancestry | ESN | 1000 Genomes on GRCh38,1000 Genomes 30x on GRCh38,1000 Genomes phase 3 release |
| **HG03517** | female | SAME1840163 | ESN | Esan | AFR | African Ancestry | ESN | 1000 Genomes on GRCh38,1000 Genomes 30x on GRCh38,1000 Genomes phase 3 release |
| **HG02946** | female | SAME1839068 | ESN | Esan | AFR | African Ancestry | ESN | 1000 Genomes on GRCh38,1000 Genomes 30x on GRCh38,1000 Genomes phase 3 release |
| **HG03428** | female | SAME1840409 | MSL | Mende | AFR | African Ancestry | MSL | 1000 Genomes on GRCh38,1000 Genomes 30x on GRCh38,1000 Genomes phase 3 release |
| **HG01459** | female | SAME1840278 | CLM | Colombian | AMR | American Ancestry | CLM | 1000 Genomes on GRCh38,1000 Genomes 30x on GRCh38,1000 Genomes phase 3 release |
| **HG02953** | male | SAME1840271 | ESN | Esan | AFR | African Ancestry | ESN | 1000 Genomes on GRCh38,1000 Genomes 30x on GRCh38,1000 Genomes phase 3 release |
| **HG03442** | male | SAME1839979 | MSL | Mende | AFR | African Ancestry | MSL | 1000 Genomes on GRCh38,1000 Genomes 30x on GRCh38,1000 Genomes phase 3 release |
| **HG01461** | male | SAME122879 | CLM | Colombian | AMR | American Ancestry | CLM | 1000 Genomes on GRCh38,1000 Genomes 30x on GRCh38,1000 Genomes phase 3 release,1000 Genomes phase 1 release |
| **HG02965** | male | SAME1840116 | ESN | Esan | AFR | African Ancestry | ESN | 1000 Genomes on GRCh38,1000 Genomes 30x on GRCh38 |
| **HG03454** | male | SAME1839764 | MSL | Mende | AFR | African Ancestry | MSL | 1000 Genomes on GRCh38,1000 Genomes 30x on GRCh38 |
| **HG01466** | female | SAME122881 | CLM | Colombian | AMR | American Ancestry | CLM | 1000 Genomes 30x on GRCh38,1000 Genomes phase 3 release |
| **HG02698** | male | SAME1839137 | PJL | Punjabi | SAS | South Asian Ancestry | PJL | 1000 Genomes 30x on GRCh38,1000 Genomes phase 3 release |
| **HG01593** | female | SAME1839634 | PJL | Punjabi | SAS | South Asian Ancestry | PJL | 1000 Genomes on GRCh38,1000 Genomes 30x on GRCh38,1000 Genomes phase 3 release |
| **HG02701** | male | SAME1840152 | PJL | Punjabi | SAS | South Asian Ancestry | PJL | 1000 Genomes 30x on GRCh38 |
| **HG01598** | female | SAME125239 | KHV | Kinh Vietnamese | EAS | East Asian Ancestry | KHV | 1000 Genomes on GRCh38,1000 Genomes 30x on GRCh38,1000 Genomes phase 3 release |
| **HG02725** | female | SAME1839737 | PJL | Punjabi | SAS | South Asian Ancestry | PJL | 1000 Genomes on GRCh38,1000 Genomes 30x on GRCh38,1000 Genomes phase 3 release |
| **HG03372** | female | SAME1839716 | ESN | Esan | AFR | African Ancestry | ESN | 1000 Genomes on GRCh38,1000 Genomes 30x on GRCh38,1000 Genomes phase 3 release |
| **HG01601** | male | SAME123455 | IBS | Iberian | EUR | European Ancestry | IBS | 1000 Genomes 30x on GRCh38,1000 Genomes phase 3 release |
| **HG03103** | male | SAME1840022 | ESN | Esan | AFR | African Ancestry | ESN | 1000 Genomes on GRCh38,1000 Genomes 30x on GRCh38,1000 Genomes phase 3 release |
| **HG01606** | male | SAME123458 | IBS | Iberian | EUR | European Ancestry | IBS | 1000 Genomes on GRCh38,1000 Genomes 30x on GRCh38,1000 Genomes phase 3 release |
| **HG03108** | female | SAME1839760 | ESN | Esan | AFR | African Ancestry | ESN | 1000 Genomes on GRCh38,1000 Genomes 30x on GRCh38,1000 Genomes phase 3 release |
| **HG03391** | male | SAME1839636 | MSL | Mende | AFR | African Ancestry | MSL | 1000 Genomes on GRCh38,1000 Genomes 30x on GRCh38,1000 Genomes phase 3 release |
| **HG01613** | female | SAME123626 | IBS | Iberian | EUR | European Ancestry | IBS | 1000 Genomes on GRCh38,1000 Genomes 30x on GRCh38,1000 Genomes phase 3 release |
| **HG03110** | female | SAME1839880 | ESN | Esan | AFR | African Ancestry | ESN | 1000 Genomes 30x on GRCh38 |
| **HG01384** | female | SAME122969 | CLM | Colombian | AMR | American Ancestry | CLM | 1000 Genomes on GRCh38,1000 Genomes 30x on GRCh38,1000 Genomes phase 3 release,1000 Genomes phase 1 release |
| **HG03115** | male | SAME1839812 | ESN | Esan | AFR | African Ancestry | ESN | 1000 Genomes on GRCh38,1000 Genomes 30x on GRCh38,1000 Genomes phase 3 release |
| **HG03644** | male | SAME1839638 | STU | Tamil | SAS | South Asian Ancestry | STU | 1000 Genomes on GRCh38,1000 Genomes 30x on GRCh38,1000 Genomes phase 3 release |
| **HG01389** | male | SAME122972 | CLM | Colombian | AMR | American Ancestry | CLM | 1000 Genomes on GRCh38,1000 Genomes 30x on GRCh38,1000 Genomes phase 3 release,1000 Genomes phase 1 release |
| **HG03242** | male | SAME1839058 | GWD | Gambian Mandinka | AFR | African Ancestry | GWD | 1000 Genomes 30x on GRCh38 |
| **HG03649** | male | SAME1839396 | PJL | Punjabi | SAS | South Asian Ancestry | PJL | 1000 Genomes on GRCh38,1000 Genomes 30x on GRCh38,1000 Genomes phase 3 release |
| **HG01391** | female | SAME122804 | CLM | Colombian | AMR | American Ancestry | CLM | 1000 Genomes 30x on GRCh38,1000 Genomes phase 3 release |
| **HG03247** | female | SAME1839174 | GWD | Gambian Mandinka | AFR | African Ancestry | GWD | 1000 Genomes on GRCh38,1000 Genomes 30x on GRCh38,1000 Genomes phase 3 release,Gambian Genome Variation Project (GRCh38) |
| **HG03651** | male | SAME1839498 | PJL | Punjabi | SAS | South Asian Ancestry | PJL | 1000 Genomes 30x on GRCh38 |
| **HG01396** | female | SAME1839954 | PUR | Puerto Rican | AMR | American Ancestry | PUR | 1000 Genomes on GRCh38,1000 Genomes 30x on GRCh38,1000 Genomes phase 3 release |
| **HG03259** | female | SAME1840303 | GWD | Gambian Mandinka | AFR | African Ancestry | GWD | 1000 Genomes on GRCh38,1000 Genomes 30x on GRCh38,1000 Genomes phase 3 release,Gambian Genome Variation Project (GRCh38) |
| **HG03663** | male | SAME1839317 | PJL | Punjabi | SAS | South Asian Ancestry | PJL | 1000 Genomes on GRCh38,1000 Genomes 30x on GRCh38,1000 Genomes phase 3 release |
| **HG03280** | male | SAME1840083 | ESN | Esan | AFR | African Ancestry | ESN | 1000 Genomes on GRCh38,1000 Genomes 30x on GRCh38,1000 Genomes phase 3 release |
| **HG03668** | female | SAME1839338 | PJL | Punjabi | SAS | South Asian Ancestry | PJL | 1000 Genomes on GRCh38,1000 Genomes 30x on GRCh38,1000 Genomes phase 3 release |

| | | | | | | | | |
|---|---|---|---|---|---|---|---|---|
| **HG01269** | female | SAME1839257 | CLM | Colombian | AMR | American Ancestry | CLM | 1000 Genomes on GRCh38,1000 Genomes 30x on GRCh38,1000 Genomes phase 3 release |
| **HG03196** | male | SAME1839921 | ESN | Esan | AFR | African Ancestry | ESN | 1000 Genomes on GRCh38,1000 Genomes 30x on GRCh38,1000 Genomes phase 3 release |
| **HG03719** | female | SAME1839843 | ITU | Telugu | SAS | South Asian Ancestry | ITU | 1000 Genomes 30x on GRCh38 |
| **HG01271** | male | SAME122998 | CLM | Colombian | AMR | American Ancestry | CLM | 1000 Genomes on GRCh38,1000 Genomes 30x on GRCh38,1000 Genomes phase 3 release,1000 Genomes phase 1 release |
| **HG03209** | male | SAME1839551 | MSL | Mende | AFR | African Ancestry | MSL | 1000 Genomes on GRCh38,1000 Genomes 30x on GRCh38,1000 Genomes phase 3 release |
| **HG03721** | female | SAME1839659 | ITU | Telugu | SAS | South Asian Ancestry | ITU | 1000 Genomes on GRCh38,1000 Genomes 30x on GRCh38 |
| **HG01276** | female | SAME122999 | CLM | Colombian | AMR | American Ancestry | CLM | 1000 Genomes 30x on GRCh38,1000 Genomes phase 3 release |
| **HG03228** | male | SAME1839232 | PJL | Punjabi | SAS | South Asian Ancestry | PJL | 1000 Genomes on GRCh38,1000 Genomes 30x on GRCh38,1000 Genomes phase 3 release |
| **HG03733** | female | SAME1839382 | STU | Tamil | SAS | South Asian Ancestry | STU | 1000 Genomes on GRCh38,1000 Genomes 30x on GRCh38,1000 Genomes phase 3 release |
| **HG01303** | female | SAME1840027 | PUR | Puerto Rican | AMR | American Ancestry | PUR | 1000 Genomes on GRCh38,1000 Genomes 30x on GRCh38,1000 Genomes phase 3 release |
| **HG03230** | female | SAME1839335 | PJL | Punjabi | SAS | South Asian Ancestry | PJL | 1000 Genomes 30x on GRCh38 |
| **HG03738** | male | SAME1839473 | STU | Tamil | SAS | South Asian Ancestry | STU | 1000 Genomes on GRCh38,1000 Genomes 30x on GRCh38,1000 Genomes phase 3 release |
| **HG01308** | male | SAME1840002 | PUR | Puerto Rican | AMR | American Ancestry | PUR | 1000 Genomes on GRCh38,1000 Genomes 30x on GRCh38,1000 Genomes phase 3 release |
| **HG03235** | female | SAME1839360 | PJL | Punjabi | SAS | South Asian Ancestry | PJL | 1000 Genomes on GRCh38,1000 Genomes 30x on GRCh38,1000 Genomes phase 3 release |
| **HG03740** | male | SAME1839248 | STU | Tamil | SAS | South Asian Ancestry | STU | 1000 Genomes on GRCh38,1000 Genomes 30x on GRCh38,1000 Genomes phase 3 release |
| **HG03432** | male | SAME1840107 | MSL | Mende | AFR | African Ancestry | MSL | 1000 Genomes on GRCh38,1000 Genomes 30x on GRCh38,1000 Genomes phase 3 release |
| **HG03745** | male | SAME1839195 | STU | Tamil | SAS | South Asian Ancestry | STU | 1000 Genomes on GRCh38,1000 Genomes 30x on GRCh38,1000 Genomes phase 3 release |
| **HG03437** | female | SAME1840123 | MSL | Mende | AFR | African Ancestry | MSL | 1000 Genomes on GRCh38,1000 Genomes 30x on GRCh38,1000 Genomes phase 3 release |
| **HG03606** | male | SAME1840248 | BEB | Bengali | SAS | South Asian Ancestry | BEB | 1000 Genomes on GRCh38,1000 Genomes 30x on GRCh38,1000 Genomes phase 3 release |
| **HG03449** | female | SAME1839923 | MSL | Mende | AFR | African Ancestry | MSL | 1000 Genomes on GRCh38,1000 Genomes 30x on GRCh38,1000 Genomes phase 3 release |
| **HG03618** | male | SAME1840062 | PJL | Punjabi | SAS | South Asian Ancestry | PJL | 1000 Genomes on GRCh38,1000 Genomes 30x on GRCh38 |
| **HG01485** | male | SAME1840047 | CLM | Colombian | AMR | American Ancestry | CLM | 1000 Genomes on GRCh38,1000 Genomes 30x on GRCh38,1000 Genomes phase 3 release |
| **HG03451** | male | SAME1840043 | MSL | Mende | AFR | African Ancestry | MSL | 1000 Genomes on GRCh38,1000 Genomes 30x on GRCh38,1000 Genomes phase 3 release |
| **HG03620** | female | SAME1839790 | PJL | Punjabi | SAS | South Asian Ancestry | PJL | 1000 Genomes 30x on GRCh38 |
| **HG01492** | female | SAME124137 | CLM | Colombian | AMR | American Ancestry | CLM | 1000 Genomes on GRCh38,1000 Genomes 30x on GRCh38,1000 Genomes phase 3 release,1000 Genomes phase 1 release |
| **HG03456** | male | SAME1839778 | MSL | Mende | AFR | African Ancestry | MSL | 1000 Genomes 30x on GRCh38 |
| **HG03625** | female | SAME1839719 | PJL | Punjabi | SAS | South Asian Ancestry | PJL | 1000 Genomes on GRCh38,1000 Genomes 30x on GRCh38,1000 Genomes phase 3 release |
| **HG01497** | male | SAME124134 | CLM | Colombian | AMR | American Ancestry | CLM | 1000 Genomes on GRCh38,1000 Genomes 30x on GRCh38,1000 Genomes phase 3 release,1000 Genomes phase 1 release |
| **HG03343** | male | SAME1840354 | ESN | Esan | AFR | African Ancestry | ESN | 1000 Genomes on GRCh38,1000 Genomes 30x on GRCh38,1000 Genomes phase 3 release |
| **HG03699** | male | SAME1840153 | PJL | Punjabi | SAS | South Asian Ancestry | PJL | 1000 Genomes on GRCh38,1000 Genomes 30x on GRCh38 |
| **HG01500** | male | SAME124941 | IBS | Iberian | EUR | European Ancestry | IBS | 1000 Genomes on GRCh38,1000 Genomes 30x on GRCh38,1000 Genomes phase 3 release |
| **HG03350** | male | SAME1840225 | ESN | Esan | AFR | African Ancestry | ESN | 1000 Genomes 30x on GRCh38 |
| **HG03702** | male | SAME1840029 | PJL | Punjabi | SAS | South Asian Ancestry | PJL | 1000 Genomes on GRCh38,1000 Genomes 30x on GRCh38,1000 Genomes phase 3 release |
| **HG01618** | female | SAME123631 | IBS | Iberian | EUR | European Ancestry | IBS | 1000 Genomes on GRCh38,1000 Genomes 30x on GRCh38,1000 Genomes phase 3 release,1000 Genomes phase 1 release |
| **HG03707** | male | SAME1839761 | PJL | Punjabi | SAS | South Asian Ancestry | PJL | 1000 Genomes 30x on GRCh38 |
| **HG01620** | female | SAME123844 | IBS | Iberian | EUR | European Ancestry | IBS | 1000 Genomes on GRCh38,1000 Genomes 30x on GRCh38,1000 Genomes phase 3 release,1000 Genomes phase 1 release |
| **HG03367** | male | SAME1840068 | ESN | Esan | AFR | African Ancestry | ESN | 1000 Genomes on GRCh38,1000 Genomes 30x on GRCh38,1000 Genomes phase 3 release |
| **HG03714** | female | SAME1839809 | ITU | Telugu | SAS | South Asian Ancestry | ITU | 1000 Genomes on GRCh38,1000 Genomes 30x on GRCh38,1000 Genomes phase 3 release |
| **HG01625** | male | SAME123839 | IBS | Iberian | EUR | European Ancestry | IBS | 1000 Genomes on GRCh38,1000 Genomes 30x on GRCh38,1000 Genomes phase 3 release,1000 Genomes phase 1 release |
| **HG03374** | female | SAME1839730 | ESN | Esan | AFR | African Ancestry | ESN | 1000 Genomes 30x on GRCh38 |
| **HG03752** | female | SAME1839314 | STU | Tamil | SAS | South Asian Ancestry | STU | 1000 Genomes on GRCh38,1000 Genomes 30x on GRCh38,1000 Genomes phase 3 release |
| **HG01632** | female | SAME124032 | IBS | Iberian | EUR | European Ancestry | IBS | 1000 Genomes on GRCh38,1000 Genomes 30x on GRCh38,1000 Genomes phase 3 release |
| **HG03122** | male | SAME1839657 | ESN | Esan | AFR | African Ancestry | ESN | 1000 Genomes 30x on GRCh38 |
| **HG03757** | female | SAME1839089 | STU | Tamil | SAS | South Asian Ancestry | STU | 1000 Genomes on GRCh38,1000 Genomes 30x on GRCh38,1000 Genomes phase 3 release |
| **HG01668** | female | SAME123111 | IBS | Iberian | EUR | European Ancestry | IBS | 1000 Genomes on GRCh38,1000 Genomes 30x on GRCh38,1000 Genomes phase 3 release |
| **HG03127** | male | SAME1839686 | ESN | Esan | AFR | African Ancestry | ESN | 1000 Genomes on GRCh38,1000 Genomes 30x on GRCh38,1000 Genomes phase 3 release |
| **HG01670** | female | SAME123290 | IBS | Iberian | EUR | European Ancestry | IBS | 1000 Genomes on GRCh38,1000 Genomes 30x on GRCh38,1000 Genomes phase 3 release |
| **HG03134** | male | SAME1839380 | ESN | Esan | AFR | African Ancestry | ESN | 1000 Genomes 30x on GRCh38 |
| **HG03771** | male | SAME1840393 | ITU | Telugu | SAS | South Asian Ancestry | ITU | 1000 Genomes on GRCh38,1000 Genomes 30x on GRCh38,1000 Genomes phase 3 release |
| **HG01675** | male | SAME123287 | IBS | Iberian | EUR | European Ancestry | IBS | 1000 Genomes on GRCh38,1000 Genomes 30x on GRCh38,1000 Genomes phase 3 release |
| **HG03139** | male | SAME1839468 | ESN | Esan | AFR | African Ancestry | ESN | 1000 Genomes on GRCh38,1000 Genomes 30x on GRCh38,1000 Genomes phase 3 release |
| **HG01505** | male | SAME124946 | IBS | Iberian | EUR | European Ancestry | IBS | 1000 Genomes 30x on GRCh38,Human Genome Structural Variation Consortium, Phase 2,1000 Genomes phase 3 release |
| **HG03788** | male | SAME1840227 | ITU | Telugu | SAS | South Asian Ancestry | ITU | 1000 Genomes on GRCh38,1000 Genomes 30x on GRCh38,1000 Genomes phase 3 release |
| **HG01512** | male | SAME124769 | IBS | Iberian | EUR | European Ancestry | IBS | 1000 Genomes on GRCh38,1000 Genomes 30x on GRCh38,1000 Genomes phase 3 release |
| **HG03297** | female | SAME1839900 | ESN | Esan | AFR | African Ancestry | ESN | 1000 Genomes on GRCh38,1000 Genomes 30x on GRCh38,1000 Genomes phase 3 release |
| **HG03790** | male | SAME1839989 | ITU | Telugu | SAS | South Asian Ancestry | ITU | 1000 Genomes on GRCh38,1000 Genomes 30x on GRCh38,1000 Genomes phase 3 release |
| **HG01517** | female | SAME124772 | IBS | Iberian | EUR | European Ancestry | IBS | 1000 Genomes 30x on GRCh38,1000 Genomes phase 3 release |
| **HG03300** | female | SAME1839467 | ESN | Esan | AFR | African Ancestry | ESN | 1000 Genomes on GRCh38,1000 Genomes 30x on GRCh38,1000 Genomes phase 3 release |
| **HG03795** | male | SAME1839925 | BEB | Bengali | SAS | South Asian Ancestry | BEB | 1000 Genomes 30x on GRCh38 |
| **HG01524** | male | SAME125327 | IBS | Iberian | EUR | European Ancestry | IBS | 1000 Genomes on GRCh38,1000 Genomes 30x on GRCh38,1000 Genomes phase 3 release |
| **HG03305** | female | SAME1839497 | ESN | Esan | AFR | African Ancestry | ESN | 1000 Genomes 30x on GRCh38 |
| **HG03579** | male | SAME1839247 | MSL | Mende | AFR | African Ancestry | MSL | 1000 Genomes 30x on GRCh38 |
| **HG01815** | female | SAME124524 | CDX | Dai Chinese | EAS | East Asian Ancestry | CDX | 1000 Genomes on GRCh38,1000 Genomes 30x on GRCh38,1000 Genomes phase 3 release |
| **HG03312** | female | SAME1839209 | ESN | Esan | AFR | African Ancestry | ESN | 1000 Genomes on GRCh38,1000 Genomes 30x on GRCh38 |
| **HG03593** | male | SAME1839171 | BEB | Bengali | SAS | South Asian Ancestry | BEB | 1000 Genomes on GRCh38,1000 Genomes 30x on GRCh38,1000 Genomes phase 3 release |
| **HG01841** | female | SAME123551 | KHV | Kinh Vietnamese | EAS | East Asian Ancestry | KHV | 1000 Genomes on GRCh38,1000 Genomes 30x on GRCh38,1000 Genomes phase 3 release |
| **HG03160** | male | SAME1839192 | ESN | Esan | AFR | African Ancestry | ESN | 1000 Genomes on GRCh38,1000 Genomes 30x on GRCh38,1000 Genomes phase 3 release |
| **HG03598** | female | SAME1839194 | BEB | Bengali | SAS | South Asian Ancestry | BEB | 1000 Genomes on GRCh38,1000 Genomes 30x on GRCh38,1000 Genomes phase 3 release |
| **HG01846** | male | SAME123558 | KHV | Kinh,Kinh Vietnamese | EAS | East Asia (SGDP),East Asian Ancestry | KinhSGDP,KHV | 1000 Genomes on GRCh38,Simons Genome Diversity Project,1000 Genomes 30x on GRCh38,1000 Genomes phase 3 release |
| **HG03172** | male | SAME1840392 | ESN | Esan | AFR | African Ancestry | ESN | 1000 Genomes on GRCh38,1000 Genomes 30x on GRCh38,1000 Genomes phase 3 release |
| **HG03548** | female | SAME1839883 | MSL | Mende | AFR | African Ancestry | MSL | 1000 Genomes on GRCh38,1000 Genomes 30x on GRCh38,1000 Genomes phase 3 release |
| **HG01853** | female | SAME123751 | KHV | Kinh Vietnamese | EAS | East Asian Ancestry | KHV | 1000 Genomes on GRCh38,1000 Genomes 30x on GRCh38,1000 Genomes phase 3 release |
| **HG03189** | female | SAME1840226 | ESN | Esan | AFR | African Ancestry | ESN | 1000 Genomes on GRCh38,1000 Genomes 30x on GRCh38,1000 Genomes phase 3 release |
| **HG03567** | female | SAME1839364 | MSL | Mende | AFR | African Ancestry | MSL | 1000 Genomes on GRCh38,1000 Genomes 30x on GRCh38,1000 Genomes phase 3 release |
| **HG01858** | female | SAME123754 | KHV | Kinh Vietnamese | EAS | East Asian Ancestry | KHV | 1000 Genomes on GRCh38,1000 Genomes 30x on GRCh38,1000 Genomes phase 3 release |
| **HG03191** | male | SAME1839988 | ESN | Esan | AFR | African Ancestry | ESN | 1000 Genomes 30x on GRCh38 |
| **HG03574** | male | SAME1839229 | MSL | Mende | AFR | African Ancestry | MSL | 1000 Genomes on GRCh38,1000 Genomes 30x on GRCh38 |
| **HG01860** | male | SAME123201 | KHV | Kinh Vietnamese | EAS | East Asian Ancestry | KHV | 1000 Genomes on GRCh38,1000 Genomes 30x on GRCh38,1000 Genomes phase 3 release |
| **HG03682** | male | SAME1840323 | STU | Tamil | SAS | South Asian Ancestry | STU | 1000 Genomes 30x on GRCh38 |
| **HG03687** | male | SAME1840337 | STU | Tamil | SAS | South Asian Ancestry | STU | 1000 Genomes on GRCh38,1000 Genomes 30x on GRCh38,1000 Genomes phase 3 release |
| **HG01771** | male | SAME124637 | IBS | Iberian | EUR | European Ancestry | IBS | 1000 Genomes on GRCh38,1000 Genomes 30x on GRCh38,1000 Genomes phase 3 release |
| **HG03694** | male | SAME1840375 | STU | Tamil | SAS | South Asian Ancestry | STU | 1000 Genomes on GRCh38,1000 Genomes 30x on GRCh38,1000 Genomes phase 3 release |
| **HG01776** | female | SAME124632 | IBS | Iberian | EUR | European Ancestry | IBS | 1000 Genomes on GRCh38,1000 Genomes 30x on GRCh38,1000 Genomes phase 3 release |
| **HG03401** | female | SAME1839299 | MSL | Mende | AFR | African Ancestry | MSL | 1000 Genomes on GRCh38,1000 Genomes 30x on GRCh38,1000 Genomes phase 3 release |
| **HG03973** | female | SAME1839728 | ITU | Telugu | SAS | South Asian Ancestry | ITU | 1000 Genomes on GRCh38,1000 Genomes 30x on GRCh38,1000 Genomes phase 3 release |
| **HG01783** | male | SAME124427 | IBS | Iberian | EUR | European Ancestry | IBS | 1000 Genomes on GRCh38,1000 Genomes 30x on GRCh38,1000 Genomes phase 3 release |
| **HG03978** | male | SAME1839748 | ITU | Telugu | SAS | South Asian Ancestry | ITU | 1000 Genomes on GRCh38,1000 Genomes 30x on GRCh38,1000 Genomes phase 3 release |
| **HG01790** | female | SAME124222 | GBR | British | EUR | European Ancestry | GBR | 1000 Genomes on GRCh38,1000 Genomes 30x on GRCh38,1000 Genomes phase 3 release,Geuvadis |
| **HG03470** | female | SAME1839670 | MSL | Mende | AFR | African Ancestry | MSL | 1000 Genomes on GRCh38,1000 Genomes 30x on GRCh38,1000 Genomes phase 3 release |
| **HG03985** | male | SAME1839601 | STU | Tamil | SAS | South Asian Ancestry | STU | 1000 Genomes on GRCh38,1000 Genomes 30x on GRCh38,1000 Genomes phase 3 release |
| **HG01947** | male | SAME124155 | PEL | Peruvian | AMR | American Ancestry | PEL | 1000 Genomes on GRCh38,1000 Genomes 30x on GRCh38,1000 Genomes phase 3 release |
| **HG03538** | male | SAME1840007 | GWD | Gambian Mandinka | AFR | African Ancestry | GWD | 1000 Genomes on GRCh38,1000 Genomes 30x on GRCh38,1000 Genomes phase 3 release,Gambian Genome Variation Project (GRCh38) |
| **HG03992** | male | SAME1839646 | STU | Tamil | SAS | South Asian Ancestry | STU | 1000 Genomes 30x on GRCh38 |
| **HG01954** | female | SAME123980 | PEL | Peruvian | AMR | American Ancestry | PEL | 1000 Genomes on GRCh38,1000 Genomes 30x on GRCh38,1000 Genomes phase 3 release |
| **HG03540** | female | SAME1839745 | GWD | Gambian Mandinka | AFR | African Ancestry | GWD | 1000 Genomes 30x on GRCh38 |
| **HG03909** | male | SAME1839438 | BEB | Bengali | SAS | South Asian Ancestry | BEB | 1000 Genomes 30x on GRCh38 |
| **HG01959** | female | SAME1839343 | ACB | African Caribbean | AFR | African Ancestry | ACB | 1000 Genomes 30x on GRCh38,1000 Genomes phase 3 release |
| **HG03557** | female | SAME1839556 | MSL | Mende | AFR | African Ancestry | MSL | 1000 Genomes on GRCh38,1000 Genomes 30x on GRCh38,1000 Genomes phase 3 release |
| **HG03911** | male | SAME1839212 | BEB | Bengali | SAS | South Asian Ancestry | BEB | 1000 Genomes on GRCh38,1000 Genomes 30x on GRCh38,1000 Genomes phase 3 release |
| **HG01961** | male | SAME1839067 | PEL | Peruvian | AMR | American Ancestry | PEL | 1000 Genomes on GRCh38,1000 Genomes 30x on GRCh38,1000 Genomes phase 3 release |
| **HG03916** | female | SAME1839316 | BEB | Bengali | SAS | South Asian Ancestry | BEB | 1000 Genomes on GRCh38,1000 Genomes 30x on GRCh38,1000 Genomes phase 3 release |
| **HG01682** | male | SAME124654 | IBS | Iberian | EUR | European Ancestry | IBS | 1000 Genomes on GRCh38,1000 Genomes 30x on GRCh38,1000 Genomes phase 3 release |
| **HG03928** | female | SAME1839042 | BEB | Bengali | SAS | South Asian Ancestry | BEB | 1000 Genomes on GRCh38,1000 Genomes 30x on GRCh38,1000 Genomes phase 3 release |
| **HG01687** | female | SAME124657 | IBS | Iberian | EUR | European Ancestry | IBS | 1000 Genomes 30x on GRCh38,1000 Genomes phase 3 release |
| **HG03571** | male | SAME1839458 | MSL | Mende | AFR | African Ancestry | MSL | 1000 Genomes on GRCh38,1000 Genomes 30x on GRCh38,1000 Genomes phase 3 release |
| **HG03884** | female | SAME1840005 | STU | Tamil | SAS | South Asian Ancestry | STU | 1000 Genomes on GRCh38,1000 Genomes 30x on GRCh38,1000 Genomes phase 3 release |
| **HG01694** | male | SAME124844 | IBS | Iberian | EUR | European Ancestry | IBS | 1000 Genomes on GRCh38,1000 Genomes 30x on GRCh38,1000 Genomes phase 3 release |
| **HG03576** | female | SAME1839238 | MSL | Mende | AFR | African Ancestry | MSL | 1000 Genomes 30x on GRCh38 |
| **HG03896** | male | SAME1839898 | STU | Tamil | SAS | South Asian Ancestry | STU | 1000 Genomes on GRCh38,1000 Genomes 30x on GRCh38,1000 Genomes phase 3 release |
| **HG01699** | male | SAME1839210 | IBS | Iberian | EUR | European Ancestry | IBS | 1000 Genomes on GRCh38,1000 Genomes 30x on GRCh38,1000 Genomes phase 3 release |
| **HG03696** | male | SAME1840135 | STU | Tamil | SAS | South Asian Ancestry | STU | 1000 Genomes on GRCh38,1000 Genomes 30x on GRCh38,1000 Genomes phase 3 release |

| | | | | | | | | |
|---|---|---|---|---|---|---|---|---|
| **HG03904** | female | SAME1839501 | BEB | Bengali | SAS | South Asian Ancestry | BEB | 1000 Genomes on GRCh38,1000 Genomes 30x on GRCh38 |
| **HG01702** | female | SAME123259 | IBS | Iberian | EUR | European Ancestry | IBS | 1000 Genomes on GRCh38,1000 Genomes 30x on GRCh38,1000 Genomes phase 3 release |
| **HG03704** | female | SAME1840040 | PJL | Punjabi | SAS | South Asian Ancestry | PJL | 1000 Genomes 30x on GRCh38 |
| **HG03803** | male | SAME1839527 | BEB | Bengali | SAS | South Asian Ancestry | BEB | 1000 Genomes on GRCh38,1000 Genomes 30x on GRCh38,1000 Genomes phase 3 release |
| **HG01865** | male | SAME123196 | KHV | Kinh Vietnamese | EAS | East Asian Ancestry | KHV | 1000 Genomes on GRCh38,1000 Genomes 30x on GRCh38,1000 Genomes phase 3 release |
| **HG03709** | female | SAME1839780 | PJL | Punjabi | SAS | South Asian Ancestry | PJL | 1000 Genomes on GRCh38,1000 Genomes 30x on GRCh38,1000 Genomes phase 3 release |
| **HG03808** | female | SAME1839552 | BEB | Bengali | SAS | South Asian Ancestry | BEB | 1000 Genomes on GRCh38,1000 Genomes 30x on GRCh38,1000 Genomes phase 3 release |
| **HG01872** | male | SAME123995 | KHV | Kinh Vietnamese | EAS | East Asian Ancestry | KHV | 1000 Genomes on GRCh38,1000 Genomes 30x on GRCh38,1000 Genomes phase 3 release |
| **HG03711** | male | SAME1839894 | STU | Tamil | SAS | South Asian Ancestry | STU | 1000 Genomes on GRCh38,1000 Genomes 30x on GRCh38,1000 Genomes phase 3 release |
| **HG03815** | male | SAME1839419 | BEB | Bengali | SAS | South Asian Ancestry | BEB | 1000 Genomes on GRCh38,1000 Genomes 30x on GRCh38,1000 Genomes phase 3 release |
| **HG01884** | female | SAME122845 | ACB | African Caribbean | AFR | African Ancestry | ACB | 1000 Genomes 30x on GRCh38,1000 Genomes phase 3 release |
| **HG03716** | male | SAME1839825 | ITU | Telugu | SAS | South Asian Ancestry | ITU | 1000 Genomes on GRCh38,1000 Genomes 30x on GRCh38,1000 Genomes phase 3 release |
| **HG01923** | male | SAME123788 | PEL | Peruvian | AMR | American Ancestry | PEL | 1000 Genomes on GRCh38,1000 Genomes 30x on GRCh38,1000 Genomes phase 3 release |
| **HG03487** | male | SAME1839480 | PJL | Punjabi | SAS | South Asian Ancestry | PJL | 1000 Genomes on GRCh38,1000 Genomes 30x on GRCh38 |
| **HG03930** | female | SAME1839156 | BEB | Bengali | SAS | South Asian Ancestry | BEB | 1000 Genomes 30x on GRCh38 |
| **HG01928** | male | SAME123782 | PEL | Peruvian | AMR | American Ancestry | PEL | 1000 Genomes 30x on GRCh38,1000 Genomes phase 3 release |
| **HG03499** | female | SAME1839219 | ESN | Esan | AFR | African Ancestry | ESN | 1000 Genomes on GRCh38,1000 Genomes 30x on GRCh38,1000 Genomes phase 3 release |
| **HG03942** | male | SAME1840356 | BEB | Bengali | SAS | South Asian Ancestry | BEB | 1000 Genomes 30x on GRCh38 |
| **HG01935** | male | SAME123588 | PEL | Peruvian | AMR | American Ancestry | PEL | 1000 Genomes on GRCh38,1000 Genomes 30x on GRCh38,1000 Genomes phase 3 release |
| **HG03514** | female | SAME1840150 | ESN | Esan | AFR | African Ancestry | ESN | 1000 Genomes on GRCh38,1000 Genomes 30x on GRCh38,1000 Genomes phase 3 release |
| **HG03947** | female | SAME1840378 | STU | Tamil | SAS | South Asian Ancestry | STU | 1000 Genomes on GRCh38,1000 Genomes 30x on GRCh38,1000 Genomes phase 3 release |
| **HG01942** | female | SAME124150 | PEL | Peruvian | AMR | American Ancestry | PEL | 1000 Genomes on GRCh38,1000 Genomes 30x on GRCh38,1000 Genomes phase 3 release |
| **HG03519** | male | SAME1840176 | ESN | Esan | AFR | African Ancestry | ESN | 1000 Genomes 30x on GRCh38 |
| **HG03834** | female | SAME1839269 | BEB | Bengali | SAS | South Asian Ancestry | BEB | 1000 Genomes 30x on GRCh38 |
| **HG01889** | female | SAME122851 | ACB | African Caribbean | AFR | African Ancestry | ACB | 1000 Genomes on GRCh38,1000 Genomes 30x on GRCh38,1000 Genomes phase 3 release |
| **HG03521** | male | SAME1840190 | ESN | Esan | AFR | African Ancestry | ESN | 1000 Genomes on GRCh38,1000 Genomes 30x on GRCh38,1000 Genomes phase 3 release |
| **HG03846** | male | SAME1839172 | STU | Tamil | SAS | South Asian Ancestry | STU | 1000 Genomes on GRCh38,1000 Genomes 30x on GRCh38,1000 Genomes phase 3 release |
| **HG01891** | female | SAME123029 | ACB | African Caribbean | AFR | African Ancestry | ACB | 1000 Genomes 30x on GRCh38,1000 Genomes phase 3 release |
| **HG03639** | male | SAME1839603 | PJL | Punjabi | SAS | South Asian Ancestry | PJL | 1000 Genomes on GRCh38,1000 Genomes 30x on GRCh38 |
| **HG04003** | male | SAME1839775 | STU | Tamil | SAS | South Asian Ancestry | STU | 1000 Genomes on GRCh38,1000 Genomes 30x on GRCh38,1000 Genomes phase 3 release |
| **HG01896** | female | SAME1840348 | ACB | African Caribbean | AFR | African Ancestry | ACB | 1000 Genomes on GRCh38,1000 Genomes 30x on GRCh38,1000 Genomes phase 3 release |
| **HG03641** | male | SAME1839626 | PJL | Punjabi | SAS | South Asian Ancestry | PJL | 1000 Genomes 30x on GRCh38 |
| **HG04015** | male | SAME1839847 | ITU | Telugu | SAS | South Asian Ancestry | ITU | 1000 Genomes on GRCh38,1000 Genomes 30x on GRCh38,1000 Genomes phase 3 release |
| **HG01916** | female | SAME1839853 | ACB | African Caribbean | AFR | African Ancestry | ACB | 1000 Genomes 30x on GRCh38,1000 Genomes phase 3 release |
| **HG03646** | male | SAME1839643 | STU | Tamil | SAS | South Asian Ancestry | STU | 1000 Genomes on GRCh38,1000 Genomes 30x on GRCh38,1000 Genomes phase 3 release |
| **HG04022** | male | SAME1839691 | ITU | Telugu | SAS | South Asian Ancestry | ITU | 1000 Genomes on GRCh38,1000 Genomes 30x on GRCh38,1000 Genomes phase 3 release |
| **HG01707** | female | SAME123264 | IBS | Iberian | EUR | European Ancestry | IBS | 1000 Genomes on GRCh38,1000 Genomes 30x on GRCh38,1000 Genomes phase 3 release |
| **HG03653** | female | SAME1839508 | PJL | Punjabi | SAS | South Asian Ancestry | PJL | 1000 Genomes on GRCh38,1000 Genomes 30x on GRCh38,1000 Genomes phase 3 release |
| **HG04039** | male | SAME1839506 | STU | Tamil | SAS | South Asian Ancestry | STU | 1000 Genomes on GRCh38,1000 Genomes 30x on GRCh38,1000 Genomes phase 3 release |
| **HG01757** | female | SAME124987 | IBS | Iberian | EUR | European Ancestry | IBS | 1000 Genomes on GRCh38,1000 Genomes 30x on GRCh38,1000 Genomes phase 3 release |
| **HG03660** | male | SAME1839218 | PJL | Punjabi | SAS | South Asian Ancestry | PJL | 1000 Genomes on GRCh38,1000 Genomes 30x on GRCh38,1000 Genomes phase 3 release |
| **HG03754** | female | SAME1839073 | STU | Tamil | SAS | South Asian Ancestry | STU | 1000 Genomes on GRCh38,1000 Genomes 30x on GRCh38,1000 Genomes phase 3 release |
| **HG01795** | female | SAME124217 | CDX | Dai Chinese | EAS | East Asian Ancestry | CDX | 1000 Genomes on GRCh38,1000 Genomes 30x on GRCh38,1000 Genomes phase 3 release |
| **HG03761** | male | SAME1839114 | PJL | Punjabi | SAS | South Asian Ancestry | PJL | 1000 Genomes on GRCh38,1000 Genomes 30x on GRCh38 |
| **HG04060** | male | SAME1839126 | ITU | Telugu | SAS | South Asian Ancestry | ITU | 1000 Genomes on GRCh38,1000 Genomes 30x on GRCh38,1000 Genomes phase 3 release |
| **HG01808** | female | SAME124300 | CDX | Dai Chinese | EAS | East Asian Ancestry | CDX | 1000 Genomes on GRCh38,1000 Genomes 30x on GRCh38,1000 Genomes phase 3 release |
| **HG03858** | female | SAME1840301 | STU | Tamil | SAS | South Asian Ancestry | STU | 1000 Genomes on GRCh38,1000 Genomes 30x on GRCh38,1000 Genomes phase 3 release |
| **HG01810** | male | SAME124519 | CDX | Dai Chinese | EAS | East Asian Ancestry | CDX | 1000 Genomes on GRCh38,1000 Genomes 30x on GRCh38,1000 Genomes phase 3 release |
| **HG03773** | male | SAME1840406 | ITU | Telugu | SAS | South Asian Ancestry | ITU | 1000 Genomes on GRCh38,1000 Genomes 30x on GRCh38,1000 Genomes phase 3 release |
| **HG03872** | male | SAME1840202 | ITU | Telugu | SAS | South Asian Ancestry | ITU | 1000 Genomes on GRCh38,1000 Genomes 30x on GRCh38,1000 Genomes phase 3 release |
| **HG01992** | female | SAME124742 | PEL | Peruvian | AMR | American Ancestry | PEL | 1000 Genomes on GRCh38,1000 Genomes 30x on GRCh38,1000 Genomes phase 3 release |
| **HG03583** | female | SAME1839283 | MSL | Mende | AFR | African Ancestry | MSL | 1000 Genomes on GRCh38,1000 Genomes 30x on GRCh38,1000 Genomes phase 3 release |
| **NA12146** | male | SAME124158 | CEU | CEPH | EUR | European Ancestry | CEU | 1000 Genomes 30x on GRCh38,1000 Genomes phase 3 release |
| **HG01997** | female | SAME124740 | PEL | Peruvian | AMR | American Ancestry | PEL | 1000 Genomes on GRCh38,1000 Genomes 30x on GRCh38,1000 Genomes phase 3 release |
| **NA12273** | female | SAME123256 | CEU | CEPH | EUR | European Ancestry | CEU | 1000 Genomes on GRCh38,1000 Genomes 30x on GRCh38,1000 Genomes phase 3 release,1000 Genomes phase 1 release,Geuvadis |
| **HG02003** | female | SAME123619 | PEL | Peruvian | AMR | American Ancestry | PEL | 1000 Genomes on GRCh38,1000 Genomes 30x on GRCh38,1000 Genomes phase 3 release |
| **HG03595** | female | SAME1839185 | BEB | Bengali | SAS | South Asian Ancestry | BEB | 1000 Genomes on GRCh38,1000 Genomes 30x on GRCh38,1000 Genomes phase 3 release |
| **NA12329** | female | SAME123531 | CEU | CEPH | EUR | European Ancestry | CEU | 1000 Genomes 30x on GRCh38,Human Genome Structural Variation Consortium, Phase 2,1000 Genomes phase 3 release |
| **HG02008** | male | SAME123621 | PEL | Peruvian | AMR | American Ancestry | PEL | 1000 Genomes on GRCh38,1000 Genomes 30x on GRCh38,1000 Genomes phase 3 release |
| **HG03603** | male | SAME1840229 | BEB | Bengali | SAS | South Asian Ancestry | BEB | 1000 Genomes on GRCh38,1000 Genomes 30x on GRCh38,1000 Genomes phase 3 release |
| **NA12336** | female | SAME125068 | CEU | CEPH | EUR | European Ancestry | CEU | 1000 Genomes 30x on GRCh38,1000 Genomes phase 3 release |
| **HG02010** | female | SAME123444 | ACB | African Caribbean | AFR | African Ancestry | ACB | 1000 Genomes on GRCh38,1000 Genomes 30x on GRCh38,1000 Genomes phase 3 release |
| **HG03672** | male | SAME1839020 | STU | Tamil | SAS | South Asian Ancestry | STU | 1000 Genomes on GRCh38,1000 Genomes 30x on GRCh38,1000 Genomes phase 3 release |
| **NA12343** | female | SAME125156 | CEU | CEPH | EUR | European Ancestry | CEU | 1000 Genomes 30x on GRCh38,1000 Genomes phase 3 release |
| **HG02015** | male | SAME123447 | KHV | Kinh Vietnamese | EAS | East Asian Ancestry | KHV | 1000 Genomes 30x on GRCh38,1000 Genomes phase 3 release |
| **HG03684** | female | SAME1840329 | STU | Tamil | SAS | South Asian Ancestry | STU | 1000 Genomes on GRCh38,1000 Genomes 30x on GRCh38,1000 Genomes phase 3 release |
| **NA12348** | female | SAME125163 | CEU | CEPH | EUR | European Ancestry | CEU | 1000 Genomes on GRCh38,1000 Genomes 30x on GRCh38,1000 Genomes phase 3 release,1000 Genomes phase 1 release,Geuvadis |
| **HG02077** | male | SAME123108 | KHV | Kinh Vietnamese | EAS | East Asian Ancestry | KHV | 1000 Genomes 30x on GRCh38,1000 Genomes phase 3 release |
| **HG03689** | female | SAME1840268 | STU | Tamil | SAS | South Asian Ancestry | STU | 1000 Genomes on GRCh38,1000 Genomes 30x on GRCh38,1000 Genomes phase 3 release |
| **NA12386** | female | SAME124386 | CEU | CEPH | EUR | European Ancestry | CEU | 1000 Genomes 30x on GRCh38,1000 Genomes phase 3 release |
| **HG02084** | female | SAME124973 | KHV | Kinh Vietnamese | EAS | East Asian Ancestry | KHV | 1000 Genomes on GRCh38,1000 Genomes 30x on GRCh38,1000 Genomes phase 3 release |
| **HG03691** | male | SAME1840366 | STU | Tamil | SAS | South Asian Ancestry | STU | 1000 Genomes on GRCh38,1000 Genomes 30x on GRCh38,1000 Genomes phase 3 release |
| **NA12413** | male | SAME124055 | CEU | CEPH | EUR | European Ancestry | CEU | 1000 Genomes on GRCh38,1000 Genomes 30x on GRCh38,1000 Genomes phase 3 release,1000 Genomes phase 1 release,Geuvadis |
| **HG02089** | female | SAME124821 | PEL | Peruvian | AMR | American Ancestry | PEL | 1000 Genomes on GRCh38,1000 Genomes 30x on GRCh38,1000 Genomes phase 3 release |
| **HG03615** | male | SAME1839951 | BEB | Bengali | SAS | South Asian Ancestry | BEB | 1000 Genomes on GRCh38,1000 Genomes 30x on GRCh38,1000 Genomes phase 3 release |
| **NA12716** | male | SAME123912 | CEU | CEPH | EUR | European Ancestry | CEU | 1000 Genomes on GRCh38,1000 Genomes 30x on GRCh38,1000 Genomes phase 3 release,1000 Genomes phase 1 release,Geuvadis |
| **HG02091** | female | SAME124650 | PEL | Peruvian | AMR | American Ancestry | PEL | 1000 Genomes 30x on GRCh38,1000 Genomes phase 3 release |
| **HG03634** | female | SAME1839575 | PJL | Punjabi | SAS | South Asian Ancestry | PJL | 1000 Genomes on GRCh38,1000 Genomes 30x on GRCh38,1000 Genomes phase 3 release |
| **NA12761** | female | SAME124466 | CEU | CEPH | EUR | European Ancestry | CEU | 1000 Genomes on GRCh38,1000 Genomes 30x on GRCh38,1000 Genomes phase 3 release,1000 Genomes phase 1 release,Geuvadis |
| **HG02104** | male | SAME123874 | PEL | Peruvian | AMR | American Ancestry | PEL | 1000 Genomes on GRCh38,1000 Genomes 30x on GRCh38,1000 Genomes phase 3 release |
| **NA12766** | male | SAME124471 | CEU | CEPH | EUR | European Ancestry | CEU | 1000 Genomes 30x on GRCh38,1000 Genomes phase 3 release |
| **HG02111** | female | SAME1839922 | ACB | African Caribbean | AFR | African Ancestry | ACB | 1000 Genomes on GRCh38,1000 Genomes 30x on GRCh38,1000 Genomes phase 3 release |
| **HG03730** | female | SAME1839369 | ITU | Telugu | SAS | South Asian Ancestry | ITU | 1000 Genomes on GRCh38,1000 Genomes 30x on GRCh38,1000 Genomes phase 3 release |
| **NA12778** | female | SAME124249 | CEU | CEPH | EUR | European Ancestry | CEU | 1000 Genomes on GRCh38,1000 Genomes 30x on GRCh38,1000 Genomes phase 3 release,1000 Genomes phase 1 release,Geuvadis |
| **HG02116** | male | SAME1839990 | KHV | Kinh Vietnamese | EAS | East Asian Ancestry | KHV | 1000 Genomes on GRCh38,1000 Genomes 30x on GRCh38,1000 Genomes phase 3 release |
| **HG03742** | male | SAME1839255 | ITU | Telugu | SAS | South Asian Ancestry | ITU | 1000 Genomes on GRCh38,1000 Genomes 30x on GRCh38,1000 Genomes phase 3 release |
| **NA12812** | male | SAME123023 | CEU | CEPH | EUR | European Ancestry | CEU | 1000 Genomes on GRCh38,1000 Genomes 30x on GRCh38,1000 Genomes phase 3 release,1000 Genomes phase 1 release,Geuvadis |
| **HG02128** | male | SAME123475 | KHV | Kinh Vietnamese | EAS | East Asian Ancestry | KHV | 1000 Genomes on GRCh38,1000 Genomes 30x on GRCh38,1000 Genomes phase 3 release |
| **HG03999** | male | SAME1839416 | STU | Tamil | SAS | South Asian Ancestry | STU | 1000 Genomes on GRCh38,1000 Genomes 30x on GRCh38,1000 Genomes phase 3 release |
| **NA12817** | male | SAME123020 | CEU | CEPH | EUR | European Ancestry | CEU | 1000 Genomes 30x on GRCh38,1000 Genomes phase 3 release |
| **HG02130** | female | SAME1839904 | KHV | Kinh Vietnamese | EAS | East Asian Ancestry | KHV | 1000 Genomes on GRCh38,1000 Genomes 30x on GRCh38,1000 Genomes phase 3 release |
| **HG04017** | male | SAME1839851 | ITU | Telugu | SAS | South Asian Ancestry | ITU | 1000 Genomes on GRCh38,1000 Genomes 30x on GRCh38,1000 Genomes phase 3 release |
| **NA12829** | male | SAME125075 | CEU | CEPH | EUR | European Ancestry | CEU | 1000 Genomes on GRCh38,1000 Genomes 30x on GRCh38,1000 Genomes phase 3 release,1000 Genomes phase 1 release,Geuvadis |
| **HG02135** | male | SAME1839872 | KHV | Kinh Vietnamese | EAS | East Asian Ancestry | KHV | 1000 Genomes 30x on GRCh38,1000 Genomes phase 3 release |
| **NA12843** | female | SAME123541 | CEU | CEPH | EUR | European Ancestry | CEU | 1000 Genomes on GRCh38,1000 Genomes 30x on GRCh38,1000 Genomes phase 3 release,1000 Genomes phase 1 release,Geuvadis |
| **HG02166** | female | SAME124207 | CDX | Dai Chinese | EAS | East Asian Ancestry | CDX | 1000 Genomes on GRCh38,1000 Genomes 30x on GRCh38,1000 Genomes phase 3 release |
| **HG04029** | female | SAME1839630 | STU | Tamil | SAS | South Asian Ancestry | STU | 1000 Genomes on GRCh38,1000 Genomes 30x on GRCh38,1000 Genomes phase 3 release |
| **NA12874** | male | SAME123389 | CEU | CEPH | EUR | European Ancestry | CEU | 1000 Genomes on GRCh38,1000 Genomes 30x on GRCh38,1000 Genomes phase 3 release,1000 Genomes phase 1 release,Geuvadis |
| **HG04036** | female | SAME1839492 | STU | Tamil | SAS | South Asian Ancestry | STU | 1000 Genomes 30x on GRCh38 |
| **NA18485** | male | SAME124100 | YRI | Yoruba | AFR | African Ancestry | YRI | 1000 Genomes 30x on GRCh38,1000 Genomes phase 3 release |
| **HG02178** | female | SAME124414 | CDX | Dai Chinese | EAS | East Asian Ancestry | CDX | 1000 Genomes on GRCh38,1000 Genomes 30x on GRCh38,1000 Genomes phase 3 release |
| **HG04229** | male | SAME1839165 | STU | Tamil | SAS | South Asian Ancestry | STU | 1000 Genomes on GRCh38,1000 Genomes 30x on GRCh38,1000 Genomes phase 3 release |
| **HG02180** | female | SAME123208 | CDX | Dai Chinese | EAS | East Asian Ancestry | CDX | 1000 Genomes on GRCh38,1000 Genomes 30x on GRCh38,1000 Genomes phase 3 release |
| **HG03886** | female | SAME1840021 | STU | Tamil | SAS | South Asian Ancestry | STU | 1000 Genomes on GRCh38,1000 Genomes 30x on GRCh38,1000 Genomes phase 3 release |
| **HG02185** | female | SAME123215 | CDX | Dai Chinese | EAS | East Asian Ancestry | CDX | 1000 Genomes on GRCh38,1000 Genomes 30x on GRCh38,1000 Genomes phase 3 release |
| **HG03898** | female | SAME1839806 | STU | Tamil | SAS | South Asian Ancestry | STU | 1000 Genomes on GRCh38,1000 Genomes 30x on GRCh38,1000 Genomes phase 3 release |
| **NA06985** | female | SAME124762 | CEU | CEPH | EUR | European Ancestry | CEU | 1000 Genomes on GRCh38,1000 Genomes 30x on GRCh38,1000 Genomes phase 3 release,1000 Genomes phase 1 release,Geuvadis |
| **NA06997** | female | SAME124930 | CEU | CEPH | EUR | European Ancestry | CEU | 1000 Genomes 30x on GRCh38,1000 Genomes phase 3 release |
| **HG02224** | male | SAME1839579 | IBS | Iberian | EUR | European Ancestry | IBS | 1000 Genomes on GRCh38,1000 Genomes 30x on GRCh38,1000 Genomes phase 3 release |
| **HG03848** | male | SAME1839187 | STU | Tamil | SAS | South Asian Ancestry | STU | 1000 Genomes on GRCh38,1000 Genomes 30x on GRCh38,1000 Genomes phase 3 release |
| **NA07022** | male | SAME125336 | CEU | CEPH | EUR | European Ancestry | CEU | 1000 Genomes 30x on GRCh38,1000 Genomes phase 3 release |

| | | | | | | | | |
|---|---|---|---|---|---|---|---|---|
| **HG02229** | female | SAME1839649 | IBS | Iberian | EUR | European Ancestry | IBS | 1000 Genomes 30x on GRCh38,1000 Genomes phase 3 release |
| **HG03850** | male | SAME1840270 | STU | Tamil | SAS | South Asian Ancestry | STU | 1000 Genomes on GRCh38,1000 Genomes 30x on GRCh38,1000 Genomes phase 3 release |
| **NA07034** | male | SAME122863 | CEU | CEPH | EUR | European Ancestry | CEU | 1000 Genomes 30x on GRCh38,1000 Genomes phase 3 release |
| **HG02231** | male | SAME1839376 | IBS | Iberian | EUR | European Ancestry | IBS | 1000 Genomes on GRCh38,1000 Genomes 30x on GRCh38,1000 Genomes phase 3 release |
| **HG03862** | female | SAME1840158 | ITU | Telugu | SAS | South Asian Ancestry | ITU | 1000 Genomes on GRCh38,1000 Genomes 30x on GRCh38,1000 Genomes phase 3 release |
| **NA07349** | male | SAME123568 | CEU | CEPH | EUR | European Ancestry | CEU | 1000 Genomes 30x on GRCh38,1000 Genomes phase 3 release |
| **HG02236** | male | SAME123520 | IBS | Iberian | EUR | European Ancestry | IBS | 1000 Genomes on GRCh38,1000 Genomes 30x on GRCh38,1000 Genomes phase 3 release |
| **HG03867** | male | SAME1840095 | ITU | Telugu | SAS | South Asian Ancestry | ITU | 1000 Genomes on GRCh38,1000 Genomes 30x on GRCh38,1000 Genomes phase 3 release |
| **NA10830** | male | SAME124548 | CEU | CEPH | EUR | European Ancestry | CEU | 1000 Genomes 30x on GRCh38,1000 Genomes phase 3 release |
| **HG02250** | male | SAME1839268 | CDX | Dai Chinese | EAS | East Asian Ancestry | CDX | 1000 Genomes on GRCh38,1000 Genomes 30x on GRCh38,1000 Genomes phase 3 release |
| **HG03874** | female | SAME1839961 | ITU | Telugu | SAS | South Asian Ancestry | ITU | 1000 Genomes on GRCh38,1000 Genomes 30x on GRCh38,1000 Genomes phase 3 release |
| **NA10835** | male | SAME124547 | CEU | CEPH | EUR | European Ancestry | CEU | 1000 Genomes 30x on GRCh38,1000 Genomes phase 3 release |
| **HG02142** | female | SAME124614 | KHV | Kinh Vietnamese | EAS | East Asian Ancestry | KHV | 1000 Genomes on GRCh38,1000 Genomes 30x on GRCh38,1000 Genomes phase 3 release |
| **HG03824** | male | SAME1839459 | BEB | Bengali | SAS | South Asian Ancestry | BEB | 1000 Genomes on GRCh38,1000 Genomes 30x on GRCh38,1000 Genomes phase 3 release |
| **NA10842** | male | SAME124755 | CEU | CEPH | EUR | European Ancestry | CEU | 1000 Genomes 30x on GRCh38,1000 Genomes phase 3 release |
| **HG02147** | female | SAME1839700 | PEL | Peruvian | AMR | American Ancestry | PEL | 1000 Genomes on GRCh38,1000 Genomes 30x on GRCh38,1000 Genomes phase 3 release |
| **HG03829** | female | SAME1839241 | BEB | Bengali | SAS | South Asian Ancestry | BEB | 1000 Genomes on GRCh38,1000 Genomes 30x on GRCh38,1000 Genomes phase 3 release |
| **NA10847** | female | SAME124757 | CEU | CEPH | EUR | European Ancestry | CEU | 1000 Genomes on GRCh38,1000 Genomes 30x on GRCh38,1000 Genomes phase 3 release,1000 Genomes phase 1 release,Geuvadis |
| **HG02154** | female | SAME124794 | CDX | Dai Chinese | EAS | East Asian Ancestry | CDX | 1000 Genomes on GRCh38,1000 Genomes 30x on GRCh38,1000 Genomes phase 3 release |
| **HG03831** | female | SAME1839348 | BEB | Bengali | SAS | South Asian Ancestry | BEB | 1000 Genomes 30x on GRCh38 |
| **NA10854** | female | SAME124933 | CEU | CEPH | EUR | European Ancestry | CEU | 1000 Genomes 30x on GRCh38,1000 Genomes phase 3 release |
| **HG01973** | female | SAME124328 | PEL | Peruvian | AMR | American Ancestry | PEL | 1000 Genomes on GRCh38,1000 Genomes 30x on GRCh38,1000 Genomes phase 3 release |
| **HG03836** | female | SAME1839280 | STU | Tamil | SAS | South Asian Ancestry | STU | 1000 Genomes on GRCh38,1000 Genomes 30x on GRCh38,1000 Genomes phase 3 release |
| **NA10859** | female | SAME124937 | CEU | CEPH | EUR | European Ancestry | CEU | 1000 Genomes 30x on GRCh38,1000 Genomes phase 3 release |
| **HG01978** | female | SAME124332 | PEL | Peruvian | AMR | American Ancestry | PEL | 1000 Genomes 30x on GRCh38,1000 Genomes phase 3 release |
| **HG03937** | female | SAME1840330 | BEB | Bengali | SAS | South Asian Ancestry | BEB | 1000 Genomes on GRCh38,1000 Genomes 30x on GRCh38,1000 Genomes phase 3 release |
| **NA10861** | female | SAME125130 | CEU | CEPH | EUR | European Ancestry | CEU | 1000 Genomes 30x on GRCh38,1000 Genomes phase 3 release |
| **HG01980** | female | SAME124917 | PEL | Peruvian | AMR | American Ancestry | PEL | 1000 Genomes on GRCh38,1000 Genomes 30x on GRCh38,1000 Genomes phase 3 release |
| **HG03944** | female | SAME1840368 | STU | Tamil | SAS | South Asian Ancestry | STU | 1000 Genomes on GRCh38,1000 Genomes 30x on GRCh38,1000 Genomes phase 3 release |
| **NA11840** | female | SAME125139 | CEU | CEPH | EUR | European Ancestry | CEU | 1000 Genomes on GRCh38,1000 Genomes 30x on GRCh38,1000 Genomes phase 3 release,1000 Genomes phase 1 release,Geuvadis |
| **HG01985** | female | SAME1840099 | ACB | African Caribbean | AFR | African Ancestry | ACB | 1000 Genomes on GRCh38,1000 Genomes 30x on GRCh38,1000 Genomes phase 3 release |
| **HG03949** | female | SAME1840132 | STU | Tamil | SAS | South Asian Ancestry | STU | 1000 Genomes on GRCh38,1000 Genomes 30x on GRCh38,1000 Genomes phase 3 release |
| **NA12045** | male | SAME123211 | CEU | CEPH | EUR | European Ancestry | CEU | 1000 Genomes on GRCh38,1000 Genomes 30x on GRCh38,1000 Genomes phase 3 release,1000 Genomes phase 1 release,Geuvadis |
| **HG02053** | male | SAME1839543 | ACB | African Caribbean | AFR | African Ancestry | ACB | 1000 Genomes on GRCh38,1000 Genomes 30x on GRCh38,1000 Genomes phase 3 release |
| **HG03951** | female | SAME1840239 | STU | Tamil | SAS | South Asian Ancestry | STU | 1000 Genomes on GRCh38,1000 Genomes 30x on GRCh38,1000 Genomes phase 3 release |
| **NA12057** | female | SAME123415 | CEU | CEPH | EUR | European Ancestry | CEU | 1000 Genomes 30x on GRCh38,1000 Genomes phase 3 release |
| **HG02058** | male | SAME125319 | KHV | Kinh Vietnamese | EAS | East Asian Ancestry | KHV | 1000 Genomes on GRCh38,1000 Genomes 30x on GRCh38,1000 Genomes phase 3 release |
| **HG03797** | male | SAME1839931 | BEB | Bengali | SAS | South Asian Ancestry | BEB | 1000 Genomes on GRCh38,1000 Genomes 30x on GRCh38 |
| **HG04173** | male | SAME1839977 | BEB | Bengali | SAS | South Asian Ancestry | BEB | 1000 Genomes on GRCh38,1000 Genomes 30x on GRCh38,1000 Genomes phase 3 release |
| **HG02060** | female | SAME1839679 | KHV | Kinh Vietnamese | EAS | East Asian Ancestry | KHV | 1000 Genomes on GRCh38,1000 Genomes 30x on GRCh38,1000 Genomes phase 3 release |
| **HG03800** | male | SAME1839595 | BEB | Bengali | SAS | South Asian Ancestry | BEB | 1000 Genomes on GRCh38,1000 Genomes 30x on GRCh38,1000 Genomes phase 3 release |
| **HG04180** | female | SAME1840012 | BEB | Bengali | SAS | South Asian Ancestry | BEB | 1000 Genomes on GRCh38,1000 Genomes 30x on GRCh38,1000 Genomes phase 3 release |
| **HG02072** | female | SAME123105 | KHV | Kinh Vietnamese | EAS | East Asian Ancestry | KHV | 1000 Genomes on GRCh38,1000 Genomes 30x on GRCh38,1000 Genomes phase 3 release |
| **HG03805** | female | SAME1839540 | BEB | Bengali | SAS | South Asian Ancestry | BEB | 1000 Genomes on GRCh38,1000 Genomes 30x on GRCh38,1000 Genomes phase 3 release |
| **HG04185** | male | SAME1839756 | BEB | Bengali | SAS | South Asian Ancestry | BEB | 1000 Genomes on GRCh38,1000 Genomes 30x on GRCh38,1000 Genomes phase 3 release |
| **HG02027** | male | SAME125318 | KHV | Kinh Vietnamese | EAS | East Asian Ancestry | KHV | 1000 Genomes 30x on GRCh38,1000 Genomes phase 3 release |
| **HG03812** | male | SAME1839405 | BEB | Bengali | SAS | South Asian Ancestry | BEB | 1000 Genomes on GRCh38,1000 Genomes 30x on GRCh38,1000 Genomes phase 3 release |
| **HG04192** | female | SAME1839902 | BEB | Bengali | SAS | South Asian Ancestry | BEB | 1000 Genomes on GRCh38,1000 Genomes 30x on GRCh38 |
| **HG03817** | female | SAME1839426 | BEB | Bengali | SAS | South Asian Ancestry | BEB | 1000 Genomes on GRCh38,1000 Genomes 30x on GRCh38,1000 Genomes phase 3 release |
| **HG02419** | female | SAME1839244 | ACB | African Caribbean | AFR | African Ancestry | ACB | 1000 Genomes on GRCh38,1000 Genomes 30x on GRCh38,1000 Genomes phase 3 release |
| **HG03906** | male | SAME1839512 | BEB | Bengali | SAS | South Asian Ancestry | BEB | 1000 Genomes 30x on GRCh38 |
| **HG04200** | female | SAME1839511 | ITU | Telugu | SAS | South Asian Ancestry | ITU | 1000 Genomes on GRCh38,1000 Genomes 30x on GRCh38,1000 Genomes phase 3 release |
| **HG02433** | male | SAME1839175 | ACB | African Caribbean | AFR | African Ancestry | ACB | 1000 Genomes on GRCh38,1000 Genomes 30x on GRCh38,1000 Genomes phase 3 release |
| **HG03913** | female | SAME1839216 | BEB | Bengali | SAS | South Asian Ancestry | BEB | 1000 Genomes on GRCh38,1000 Genomes 30x on GRCh38,1000 Genomes phase 3 release |
| **HG04212** | female | SAME1839325 | ITU | Telugu | SAS | South Asian Ancestry | ITU | 1000 Genomes on GRCh38,1000 Genomes 30x on GRCh38,1000 Genomes phase 3 release |
| **HG02445** | male | SAME1840341 | ACB | African Caribbean | AFR | African Ancestry | ACB | 1000 Genomes on GRCh38,1000 Genomes 30x on GRCh38,1000 Genomes phase 3 release |
| **HG03920** | male | SAME1839093 | BEB | Bengali | SAS | South Asian Ancestry | BEB | 1000 Genomes on GRCh38,1000 Genomes 30x on GRCh38,1000 Genomes phase 3 release |
| **HG04217** | female | SAME1839350 | ITU | Telugu | SAS | South Asian Ancestry | ITU | 1000 Genomes 30x on GRCh38,Human Genome Structural Variation Consortium, Phase 2 |
| **HG02464** | male | SAME1839978 | GWD | Gambian Mandinka,Gar | AFR | African Ancestry,Africa (SGDP) | GWD,GambianSGDP | 1000 Genomes on GRCh38,Simons Genome Diversity Project,1000 Genomes 30x on GRCh38,1000 Genomes phase 3 release,Gambian Genome Variation Project (GRCh3 |
| **HG03925** | female | SAME1839025 | BEB | Bengali | SAS | South Asian Ancestry | BEB | 1000 Genomes on GRCh38,1000 Genomes 30x on GRCh38,1000 Genomes phase 3 release |
| **HG04142** | female | SAME1839177 | BEB | Bengali | SAS | South Asian Ancestry | BEB | 1000 Genomes 30x on GRCh38 |
| **HG02558** | female | SAME1840057 | ACB | African Caribbean | AFR | African Ancestry | ACB | 1000 Genomes on GRCh38,1000 Genomes 30x on GRCh38,1000 Genomes phase 3 release |
| **HG03963** | male | SAME1840055 | ITU | Telugu | SAS | South Asian Ancestry | ITU | 1000 Genomes on GRCh38,1000 Genomes 30x on GRCh38,1000 Genomes phase 3 release |
| **HG04147** | female | SAME1839116 | BEB | Bengali | SAS | South Asian Ancestry | BEB | 1000 Genomes on GRCh38,1000 Genomes 30x on GRCh38 |
| **HG02572** | male | SAME1839570 | GWD | Gambian Mandinka | AFR | African Ancestry | GWD | 1000 Genomes 30x on GRCh38 |
| **HG03968** | female | SAME1840077 | ITU | Telugu | SAS | South Asian Ancestry | ITU | 1000 Genomes on GRCh38,1000 Genomes 30x on GRCh38,1000 Genomes phase 3 release |
| **HG04159** | female | SAME1840399 | BEB | Bengali | SAS | South Asian Ancestry | BEB | 1000 Genomes on GRCh38,1000 Genomes 30x on GRCh38,1000 Genomes phase 3 release |
| **HG02577** | female | SAME1839642 | ACB | African Caribbean | AFR | African Ancestry | ACB | 1000 Genomes on GRCh38,1000 Genomes 30x on GRCh38,1000 Genomes phase 3 release |
| **HG04161** | male | SAME1840092 | BEB | Bengali | SAS | South Asian Ancestry | BEB | 1000 Genomes on GRCh38,1000 Genomes 30x on GRCh38,1000 Genomes phase 3 release |
| **HG02584** | male | SAME1839425 | GWD | Gambian Mandinka | AFR | African Ancestry | GWD | 1000 Genomes 30x on GRCh38 |
| **HG03778** | male | SAME1840344 | ITU | Telugu | SAS | South Asian Ancestry | ITU | 1000 Genomes on GRCh38,1000 Genomes 30x on GRCh38,1000 Genomes phase 3 release |
| **HG04096** | male | SAME1840054 | ITU | Telugu | SAS | South Asian Ancestry | ITU | 1000 Genomes on GRCh38,1000 Genomes 30x on GRCh38,1000 Genomes phase 3 release |
| **HG02589** | female | SAME1839401 | GWD | Gambian Mandinka | AFR | African Ancestry | GWD | 1000 Genomes on GRCh38,1000 Genomes 30x on GRCh38,1000 Genomes phase 3 release,Gambian Genome Variation Project (GRCh38) |
| **HG03780** | female | SAME1840112 | ITU | Telugu | SAS | South Asian Ancestry | ITU | 1000 Genomes on GRCh38,1000 Genomes 30x on GRCh38,1000 Genomes phase 3 release |
| **HG02596** | male | SAME1839201 | GWD | Gambian Mandinka | AFR | African Ancestry | GWD | 1000 Genomes 30x on GRCh38 |
| **HG03785** | male | SAME1840212 | ITU | Telugu | SAS | South Asian Ancestry | ITU | 1000 Genomes on GRCh38,1000 Genomes 30x on GRCh38,1000 Genomes phase 3 release |
| **HG04135** | female | SAME1839305 | BEB | Bengali | SAS | South Asian Ancestry | BEB | 1000 Genomes on GRCh38,1000 Genomes 30x on GRCh38 |
| **HG02279** | male | SAME1840362 | PEL | Peruvian | AMR | American Ancestry | PEL | 1000 Genomes 30x on GRCh38,1000 Genomes phase 3 release |
| **HG03792** | male | SAME1839912 | ITU | Telugu | SAS | South Asian Ancestry | ITU | 1000 Genomes on GRCh38,1000 Genomes 30x on GRCh38,1000 Genomes phase 3 release |
| **NA18555** | female | SAME124090 | CHB | Han Chinese | EAS | East Asian Ancestry | CHB | 1000 Genomes on GRCh38,1000 Genomes 30x on GRCh38,1000 Genomes phase 3 release,1000 Genomes phase 1 release,90 Han Chinese high coverage genomes |
| **HG02281** | male | SAME1840094 | ACB | African Caribbean | AFR | African Ancestry | ACB | 1000 Genomes on GRCh38,1000 Genomes 30x on GRCh38,1000 Genomes phase 3 release |
| **NA12155** | male | SAME124955 | CEU | CEPH | EUR | European Ancestry | CEU | 1000 Genomes on GRCh38,1000 Genomes 30x on GRCh38,1000 Genomes phase 3 release,1000 Genomes phase 1 release,Geuvadis |
| **NA18562** | male | SAME123927 | CHB | Han Chinese | EAS | East Asian Ancestry | CHB | 1000 Genomes on GRCh38,1000 Genomes 30x on GRCh38,1000 Genomes phase 3 release,1000 Genomes phase 1 release,90 Han Chinese high coverage genomes |
| **HG02286** | female | SAME122912 | PEL | Peruvian | AMR | American Ancestry | PEL | 1000 Genomes on GRCh38,1000 Genomes 30x on GRCh38,1000 Genomes phase 3 release |
| **NA12249** | female | SAME125371 | CEU | CEPH | EUR | European Ancestry | CEU | 1000 Genomes on GRCh38,1000 Genomes 30x on GRCh38,1000 Genomes phase 3 release,1000 Genomes phase 1 release,Geuvadis |
| **NA18567** | female | SAME123924 | CHB | Han Chinese | EAS | East Asian Ancestry | CHB | 1000 Genomes on GRCh38,1000 Genomes 30x on GRCh38,1000 Genomes phase 3 release,1000 Genomes phase 1 release |
| **HG02293** | female | SAME1839934 | PEL | Peruvian | AMR | American Ancestry | PEL | 1000 Genomes 30x on GRCh38,1000 Genomes phase 3 release |
| **NA12275** | female | SAME125263 | CEU | CEPH | EUR | European Ancestry | CEU | 1000 Genomes on GRCh38,1000 Genomes 30x on GRCh38,1000 Genomes phase 3 release,1000 Genomes phase 1 release,Geuvadis |
| **NA18574** | female | SAME123737 | CHB | Han Chinese | EAS | East Asian Ancestry | CHB | 1000 Genomes on GRCh38,1000 Genomes 30x on GRCh38,1000 Genomes phase 3 release,1000 Genomes phase 1 release |
| **HG02298** | female | SAME1839913 | PEL | Peruvian | AMR | American Ancestry | PEL | 1000 Genomes on GRCh38,1000 Genomes 30x on GRCh38,1000 Genomes phase 3 release |
| **NA12282** | male | SAME124640 | CEU | CEPH | EUR | European Ancestry | CEU | 1000 Genomes on GRCh38,1000 Genomes 30x on GRCh38,1000 Genomes phase 3 release,1000 Genomes phase 1 release,Geuvadis |
| **NA18579** | female | SAME123742 | CHB | Han Chinese | EAS | East Asian Ancestry | CHB | 1000 Genomes on GRCh38,1000 Genomes 30x on GRCh38,1000 Genomes phase 3 release,1000 Genomes phase 1 release,90 Han Chinese high coverage genomes |
| **HG02301** | female | SAME124280 | PEL | Peruvian | AMR | American Ancestry | PEL | 1000 Genomes on GRCh38,1000 Genomes 30x on GRCh38,1000 Genomes phase 3 release |
| **NA12287** | female | SAME124641 | CEU | CEPH | EUR | European Ancestry | CEU | 1000 Genomes on GRCh38,1000 Genomes 30x on GRCh38,1000 Genomes phase 3 release,1000 Genomes phase 1 release,Geuvadis |
| **NA18593** | female | SAME123720 | CHB | Han Chinese | EAS | East Asian Ancestry | CHB | 1000 Genomes on GRCh38,1000 Genomes 30x on GRCh38,1000 Genomes phase 3 release,1000 Genomes phase 1 release,90 Han Chinese high coverage genomes |
| **HG02471** | female | SAME1839741 | ACB | African Caribbean | AFR | African Ancestry | ACB | 1000 Genomes on GRCh38,1000 Genomes 30x on GRCh38,1000 Genomes phase 3 release |
| **NA12340** | male | SAME125157 | CEU | CEPH | EUR | European Ancestry | CEU | 1000 Genomes on GRCh38,1000 Genomes 30x on GRCh38,1000 Genomes phase 3 release,1000 Genomes phase 1 release,Geuvadis |
| **NA18606** | male | SAME124879 | CHB | Han Chinese | EAS | East Asian Ancestry | CHB | 1000 Genomes on GRCh38,1000 Genomes 30x on GRCh38,1000 Genomes phase 3 release,1000 Genomes phase 1 release |
| **HG02476** | female | SAME1839717 | ACB | African Caribbean | AFR | African Ancestry | ACB | 1000 Genomes on GRCh38,1000 Genomes 30x on GRCh38,1000 Genomes phase 3 release |
| **NA12376** | female | SAME123275 | CEU | CEPH | EUR | European Ancestry | CEU | 1000 Genomes 30x on GRCh38,1000 Genomes phase 3 release |
| **NA18613** | male | SAME125037 | CHB | Han Chinese | EAS | East Asian Ancestry | CHB | 1000 Genomes on GRCh38,1000 Genomes 30x on GRCh38,1000 Genomes phase 3 release,1000 Genomes phase 1 release |
| **HG02490** | male | SAME1839625 | PJL | Punjabi | SAS | South Asian Ancestry | PJL | 1000 Genomes on GRCh38,1000 Genomes 30x on GRCh38,1000 Genomes phase 3 release |
| **NA12383** | female | SAME124384 | CEU | CEPH | EUR | European Ancestry | CEU | 1000 Genomes on GRCh38,1000 Genomes 30x on GRCh38,1000 Genomes phase 3 release,1000 Genomes phase 1 release,Geuvadis |
| **NA18618** | female | SAME125042 | CHB | Han Chinese | EAS | East Asian Ancestry | CHB | 1000 Genomes on GRCh38,1000 Genomes 30x on GRCh38,1000 Genomes phase 3 release,1000 Genomes phase 1 release |
| **HG02495** | female | SAME1839694 | PJL | Punjabi | SAS | South Asian Ancestry | PJL | 1000 Genomes 30x on GRCh38 |
| **NA12489** | female | SAME125358 | CEU | CEPH | EUR | European Ancestry | CEU | 1000 Genomes on GRCh38,1000 Genomes 30x on GRCh38,1000 Genomes phase 3 release,1000 Genomes phase 1 release,Geuvadis |
| **NA18620** | male | SAME124510 | CHB | Han Chinese | EAS | East Asian Ancestry | CHB | 1000 Genomes on GRCh38,1000 Genomes 30x on GRCh38,1000 Genomes phase 3 release,1000 Genomes phase 1 release,90 Han Chinese high coverage genomes |
| **HG02318** | female | SAME1839391 | ACB | African Caribbean | AFR | African Ancestry | ACB | 1000 Genomes on GRCh38,1000 Genomes 30x on GRCh38,1000 Genomes phase 3 release |
| **NA10851** | male | SAME125317 | CEU | CEPH | EUR | European Ancestry | CEU | 1000 Genomes on GRCh38,1000 Genomes 30x on GRCh38,1000 Genomes phase 3 release,1000 Genomes phase 1 release,Geuvadis |
| **NA18625** | female | SAME125253 | CHB | Han Chinese | EAS | East Asian Ancestry | CHB | 1000 Genomes on GRCh38,1000 Genomes 30x on GRCh38,1000 Genomes phase 3 release,1000 Genomes phase 1 release |
| **HG02325** | female | SAME1839521 | ACB | African Caribbean | AFR | African Ancestry | ACB | 1000 Genomes on GRCh38,1000 Genomes 30x on GRCh38,1000 Genomes phase 3 release |

| | | | | | | | | |
|---|---|---|---|---|---|---|---|---|
| **NA10856** | male | SAME124935 | CEU | CEPH | EUR | European Ancestry | CEU | 1000 Genomes 30x on GRCh38,1000 Genomes phase 3 release |
| **NA18632** | male | SAME124703 | CHB | Han Chinese | EAS | East Asian Ancestry | CHB | 1000 Genomes on GRCh38,1000 Genomes 30x on GRCh38,1000 Genomes phase 3 release,1000 Genomes phase 1 release,90 Han Chinese high coverage genomes |
| **HG02332** | male | SAME1839315 | ACB | African Caribbean | AFR | African Ancestry | ACB | 1000 Genomes on GRCh38,1000 Genomes 30x on GRCh38,1000 Genomes phase 3 release |
| **NA10863** | female | SAME125132 | CEU | CEPH | EUR | European Ancestry | CEU | 1000 Genomes 30x on GRCh38,1000 Genomes phase 3 release |
| **NA18637** | male | SAME124700 | CHB | Han Chinese | EAS | East Asian Ancestry | CHB | 1000 Genomes on GRCh38,1000 Genomes 30x on GRCh38,1000 Genomes phase 3 release,1000 Genomes phase 1 release,90 Han Chinese high coverage genomes |
| **HG02337** | female | SAME1839296 | ACB | African Caribbean | AFR | African Ancestry | ACB | 1000 Genomes on GRCh38,1000 Genomes 30x on GRCh38,1000 Genomes phase 3 release |
| **NA11830** | female | SAME124553 | CEU | CEPH | EUR | European Ancestry | CEU | 1000 Genomes on GRCh38,1000 Genomes 30x on GRCh38,1000 Genomes phase 3 release,1000 Genomes phase 1 release,Geuvadis |
| **NA18644** | female | SAME122976 | CHB | Han Chinese | EAS | East Asian Ancestry | CHB | 1000 Genomes on GRCh38,1000 Genomes 30x on GRCh38,1000 Genomes phase 3 release |
| **NA11892** | female | SAME123045 | CEU | CEPH | EUR | European Ancestry | CEU | 1000 Genomes on GRCh38,1000 Genomes 30x on GRCh38,1000 Genomes phase 3 release,1000 Genomes phase 1 release,Geuvadis |
| **NA18740** | male | SAME123917 | CHB | Han Chinese | EAS | East Asian Ancestry | CHB | 1000 Genomes on GRCh38,1000 Genomes 30x on GRCh38,1000 Genomes phase 3 release,1000 Genomes phase 1 release |
| **HG02508** | female | SAME1839334 | ACB | African Caribbean | AFR | African Ancestry | ACB | 1000 Genomes on GRCh38,1000 Genomes 30x on GRCh38,1000 Genomes phase 3 release |
| **NA11917** | male | SAME122838 | CEU | CEPH | EUR | European Ancestry | CEU | 1000 Genomes 30x on GRCh38,1000 Genomes phase 3 release |
| **NA18745** | male | SAME123914 | CHB | Han Chinese | EAS | East Asian Ancestry | CHB | 1000 Genomes on GRCh38,1000 Genomes 30x on GRCh38,1000 Genomes phase 3 release,1000 Genomes phase 1 release |
| **HG02522** | female | SAME1840326 | KHV | Kinh Vietnamese | EAS | East Asian Ancestry | KHV | 1000 Genomes on GRCh38,1000 Genomes 30x on GRCh38,1000 Genomes phase 3 release |
| **NA11931** | female | SAME125096 | CEU | CEPH | EUR | European Ancestry | CEU | 1000 Genomes on GRCh38,1000 Genomes 30x on GRCh38,1000 Genomes phase 3 release,1000 Genomes phase 1 release,Geuvadis |
| **NA18757** | male | SAME123727 | CHB | Han Chinese | EAS | East Asian Ancestry | CHB | 1000 Genomes on GRCh38,1000 Genomes 30x on GRCh38,1000 Genomes phase 3 release,1000 Genomes phase 1 release |
| **HG02541** | male | SAME1840214 | ACB | African Caribbean | AFR | African Ancestry | ACB | 1000 Genomes on GRCh38,1000 Genomes 30x on GRCh38,1000 Genomes phase 3 release |
| **NA11993** | female | SAME123965 | CEU | CEPH | EUR | European Ancestry | CEU | 1000 Genomes on GRCh38,1000 Genomes 30x on GRCh38,1000 Genomes phase 3 release,1000 Genomes phase 1 release,Geuvadis |
| **HG02546** | female | SAME1839932 | ACB | African Caribbean | AFR | African Ancestry | ACB | 1000 Genomes on GRCh38,1000 Genomes 30x on GRCh38,1000 Genomes phase 3 release |
| **NA12004** | female | SAME125144 | CEU | CEPH | EUR | European Ancestry | CEU | 1000 Genomes on GRCh38,1000 Genomes 30x on GRCh38,1000 Genomes phase 3 release,1000 Genomes phase 1 release,Geuvadis |
| **NA18853** | male | SAME124733 | YRI | Yoruba | AFR | African Ancestry | YRI | 1000 Genomes on GRCh38,1000 Genomes 30x on GRCh38,1000 Genomes phase 3 release,1000 Genomes phase 1 release |
| **HG02351** | male | SAME1840284 | CDX | Dai Chinese | EAS | East Asian Ancestry | CDX | 1000 Genomes on GRCh38,1000 Genomes 30x on GRCh38,1000 Genomes phase 3 release |
| **HG04238** | male | SAME1840274 | ITU | Telugu | SAS | South Asian Ancestry | ITU | 1000 Genomes on GRCh38,1000 Genomes 30x on GRCh38,1000 Genomes phase 3 release |
| **NA18916** | female | SAME123408 | YRI | Yoruba | AFR | African Ancestry | YRI | 1000 Genomes on GRCh38,1000 Genomes 30x on GRCh38,1000 Genomes phase 3 release,1000 Genomes phase 1 release,Geuvadis |
| **HG02356** | male | SAME1840261 | CDX | Dai Chinese | EAS | East Asian Ancestry | CDX | 1000 Genomes on GRCh38,1000 Genomes 30x on GRCh38,1000 Genomes phase 3 release |
| **NA18923** | male | SAME123205 | YRI | Yoruba | AFR | African Ancestry | YRI | 1000 Genomes on GRCh38,1000 Genomes 30x on GRCh38,1000 Genomes phase 3 release,1000 Genomes phase 1 release,Geuvadis |
| **NA06994** | male | SAME124931 | CEU | CEPH | EUR | European Ancestry | CEU | 1000 Genomes on GRCh38,1000 Genomes 30x on GRCh38,1000 Genomes phase 3 release,1000 Genomes phase 1 release,Geuvadis |
| **NA18930** | female | SAME123040 | YRI | Yoruba | AFR | African Ancestry | YRI | 1000 Genomes 30x on GRCh38,1000 Genomes phase 3 release |
| **HG02375** | male | SAME1840244 | CDX | Dai Chinese | EAS | East Asian Ancestry | CDX | 1000 Genomes on GRCh38,1000 Genomes 30x on GRCh38,1000 Genomes phase 3 release |
| **NA07000** | female | SAME124963 | CEU | CEPH | EUR | European Ancestry | CEU | 1000 Genomes on GRCh38,1000 Genomes 30x on GRCh38,1000 Genomes phase 3 release,1000 Genomes phase 1 release,Geuvadis |
| **NA18935** | male | SAME123035 | YRI | Yoruba | AFR | African Ancestry | YRI | 1000 Genomes 30x on GRCh38,1000 Genomes phase 3 release |
| **HG02382** | male | SAME1840025 | CDX | Dai Chinese | EAS | East Asian Ancestry | CDX | 1000 Genomes on GRCh38,1000 Genomes 30x on GRCh38,1000 Genomes phase 3 release |
| **NA07029** | male | SAME125338 | CEU | CEPH | EUR | European Ancestry | CEU | 1000 Genomes 30x on GRCh38,1000 Genomes phase 3 release |
| **NA18942** | female | SAME124308 | JPT | Japanese | EAS | East Asian Ancestry | JPT | 1000 Genomes on GRCh38,1000 Genomes 30x on GRCh38,1000 Genomes phase 3 release,1000 Genomes phase 1 release |
| **NA07031** | female | SAME122860 | CEU | CEPH | EUR | European Ancestry | CEU | 1000 Genomes 30x on GRCh38,1000 Genomes phase 3 release |
| **NA18947** | female | SAME124313 | JPT | Japanese | EAS | East Asian Ancestry | JPT | 1000 Genomes on GRCh38,1000 Genomes 30x on GRCh38,1000 Genomes phase 3 release,1000 Genomes phase 1 release |
| **HG02394** | male | SAME123305 | CDX | Dai Chinese | EAS | East Asian Ancestry | CDX | 1000 Genomes on GRCh38,1000 Genomes 30x on GRCh38,1000 Genomes phase 3 release |
| **NA07048** | male | SAME124188 | CEU | CEPH | EUR | European Ancestry | CEU | 1000 Genomes on GRCh38,1000 Genomes 30x on GRCh38,1000 Genomes phase 3 release,1000 Genomes phase 1 release,Geuvadis |
| **NA18954** | female | SAME124108 | JPT | Japanese | EAS | East Asian Ancestry | JPT | 1000 Genomes on GRCh38,1000 Genomes 30x on GRCh38,1000 Genomes phase 3 release,1000 Genomes phase 1 release |
| **HG02399** | male | SAME1839842 | CDX | Dai Chinese | EAS | East Asian Ancestry | CDX | 1000 Genomes on GRCh38,1000 Genomes 30x on GRCh38,1000 Genomes phase 3 release |
| **NA07055** | female | SAME124767 | CEU | CEPH | EUR | European Ancestry | CEU | 1000 Genomes 30x on GRCh38,1000 Genomes phase 3 release |
| **NA18959** | male | SAME124115 | JPT | Japanese | EAS | East Asian Ancestry | JPT | 1000 Genomes on GRCh38,1000 Genomes 30x on GRCh38,1000 Genomes phase 3 release,1000 Genomes phase 1 release |
| **HG02402** | male | SAME1839412 | CDX | Dai Chinese | EAS | East Asian Ancestry | CDX | 1000 Genomes on GRCh38,1000 Genomes 30x on GRCh38,1000 Genomes phase 3 release |
| **NA07346** | female | SAME123571 | CEU | CEPH | EUR | European Ancestry | CEU | 1000 Genomes on GRCh38,1000 Genomes 30x on GRCh38,1000 Genomes phase 3 release,1000 Genomes phase 1 release,Geuvadis |
| **NA18961** | male | SAME123957 | JPT | Japanese | EAS | East Asian Ancestry | JPT | 1000 Genomes on GRCh38,1000 Genomes 30x on GRCh38,1000 Genomes phase 3 release,1000 Genomes phase 1 release |
| **HG02407** | male | SAME1839464 | CDX | Dai Chinese | EAS | East Asian Ancestry | CDX | 1000 Genomes on GRCh38,1000 Genomes 30x on GRCh38,1000 Genomes phase 3 release |
| **NA07435** | male | SAME124342 | CEU | CEPH | EUR | European Ancestry | CEU | 1000 Genomes 30x on GRCh38,1000 Genomes phase 3 release |
| **NA18966** | male | SAME123962 | JPT | Japanese | EAS | East Asian Ancestry | JPT | 1000 Genomes on GRCh38,1000 Genomes 30x on GRCh38,1000 Genomes phase 3 release,1000 Genomes phase 1 release |
| **HG02255** | male | SAME1839337 | ACB | African Caribbean | AFR | African Ancestry | ACB | 1000 Genomes on GRCh38,1000 Genomes 30x on GRCh38,1000 Genomes phase 3 release |
| **NA10837** | male | SAME124546 | CEU | CEPH | EUR | European Ancestry | CEU | 1000 Genomes 30x on GRCh38,1000 Genomes phase 3 release |
| **NA18973** | female | SAME123761 | JPT | Japanese | EAS | East Asian Ancestry | JPT | 1000 Genomes on GRCh38,1000 Genomes 30x on GRCh38,1000 Genomes phase 3 release,1000 Genomes phase 1 release |
| **HG02262** | male | SAME1839128 | PEL | Peruvian | AMR | American Ancestry | PEL | 1000 Genomes on GRCh38,1000 Genomes 30x on GRCh38,1000 Genomes phase 3 release |
| **HG04194** | male | SAME1839818 | BEB | Bengali | SAS | South Asian Ancestry | BEB | 1000 Genomes on GRCh38,1000 Genomes 30x on GRCh38,1000 Genomes phase 3 release |
| **NA18978** | female | SAME123760 | JPT | Japanese | EAS | East Asian Ancestry | JPT | 1000 Genomes on GRCh38,1000 Genomes 30x on GRCh38,1000 Genomes phase 3 release,1000 Genomes phase 1 release |
| **HG04199** | male | SAME1839840 | STU | Tamil | SAS | South Asian Ancestry | STU | 1000 Genomes 30x on GRCh38 |
| **HG02274** | male | SAME1840306 | PEL | Peruvian | AMR | American Ancestry | PEL | 1000 Genomes on GRCh38,1000 Genomes 30x on GRCh38,1000 Genomes phase 3 release |
| **HG04202** | female | SAME1839519 | ITU | Telugu | SAS | South Asian Ancestry | ITU | 1000 Genomes on GRCh38,1000 Genomes 30x on GRCh38,1000 Genomes phase 3 release |
| **NA19058** | male | SAME125024 | JPT | Japanese | EAS | East Asian Ancestry | JPT | 1000 Genomes on GRCh38,1000 Genomes 30x on GRCh38,1000 Genomes phase 3 release,1000 Genomes phase 1 release |
| **HG02604** | female | SAME1840282 | PJL | Punjabi | SAS | South Asian Ancestry | PJL | 1000 Genomes on GRCh38,1000 Genomes 30x on GRCh38,1000 Genomes phase 3 release |
| **HG04214** | female | SAME1839328 | ITU | Telugu | SAS | South Asian Ancestry | ITU | 1000 Genomes on GRCh38,1000 Genomes 30x on GRCh38,1000 Genomes phase 3 release |
| **NA19060** | male | SAME123007 | JPT | Japanese | EAS | East Asian Ancestry | JPT | 1000 Genomes on GRCh38,1000 Genomes 30x on GRCh38,1000 Genomes phase 3 release,1000 Genomes phase 1 release |
| **HG02611** | female | SAME1840401 | GWD | Gambian Mandinka | AFR | African Ancestry | GWD | 1000 Genomes on GRCh38,1000 Genomes 30x on GRCh38,1000 Genomes phase 3 release,Gambian Genome Variation Project (GRCh38) |
| **HG04219** | male | SAME1839359 | ITU | Telugu | SAS | South Asian Ancestry | ITU | 1000 Genomes on GRCh38,1000 Genomes 30x on GRCh38,1000 Genomes phase 3 release |
| **NA19065** | female | SAME123011 | JPT | Japanese | EAS | East Asian Ancestry | JPT | 1000 Genomes on GRCh38,1000 Genomes 30x on GRCh38,1000 Genomes phase 3 release,1000 Genomes phase 1 release |
| **HG02623** | male | SAME1840182 | GWD | Gambian Mandinka | AFR | African Ancestry | GWD | 1000 Genomes on GRCh38,1000 Genomes 30x on GRCh38,1000 Genomes phase 3 release,Gambian Genome Variation Project (GRCh38) |
| **HG04093** | male | SAME1839936 | ITU | Telugu | SAS | South Asian Ancestry | ITU | 1000 Genomes on GRCh38,1000 Genomes 30x on GRCh38,1000 Genomes phase 3 release |
| **NA19072** | male | SAME122822 | JPT | Japanese | EAS | East Asian Ancestry | JPT | 1000 Genomes on GRCh38,1000 Genomes 30x on GRCh38,1000 Genomes phase 3 release,1000 Genomes phase 1 release |
| **HG02628** | male | SAME1840238 | GWD | Gambian Mandinka | AFR | African Ancestry | GWD | 1000 Genomes on GRCh38,1000 Genomes 30x on GRCh38,1000 Genomes phase 3 release,Gambian Genome Variation Project (GRCh38) |
| **HG04098** | male | SAME1840059 | ITU | Telugu | SAS | South Asian Ancestry | ITU | 1000 Genomes on GRCh38,1000 Genomes 30x on GRCh38,1000 Genomes phase 3 release |
| **NA19077** | female | SAME122818 | JPT | Japanese | EAS | East Asian Ancestry | JPT | 1000 Genomes on GRCh38,1000 Genomes 30x on GRCh38,1000 Genomes phase 3 release,1000 Genomes phase 1 release |
| **HG02630** | female | SAME1839963 | GWD | Gambian Mandinka | AFR | African Ancestry | GWD | 1000 Genomes 30x on GRCh38 |
| **HG04106** | female | SAME1839559 | STU | Tamil | SAS | South Asian Ancestry | STU | 1000 Genomes on GRCh38,1000 Genomes 30x on GRCh38,1000 Genomes phase 3 release |
| **NA19084** | female | SAME123364 | JPT | Japanese | EAS | East Asian Ancestry | JPT | 1000 Genomes on GRCh38,1000 Genomes 30x on GRCh38,1000 Genomes phase 3 release,1000 Genomes phase 1 release |
| **HG02635** | female | SAME1840023 | GWD | Gambian Mandinka | AFR | African Ancestry | GWD | 1000 Genomes on GRCh38,1000 Genomes 30x on GRCh38,1000 Genomes phase 3 release,Gambian Genome Variation Project (GRCh38) |
| **HG04118** | female | SAME1839379 | ITU | Telugu | SAS | South Asian Ancestry | ITU | 1000 Genomes on GRCh38,1000 Genomes 30x on GRCh38,1000 Genomes phase 3 release |
| **NA19089** | male | SAME123649 | JPT | Japanese | EAS | East Asian Ancestry | JPT | 1000 Genomes on GRCh38,1000 Genomes 30x on GRCh38,1000 Genomes phase 3 release |
| **HG02642** | male | SAME1839792 | GWD | Gambian Mandinka | AFR | African Ancestry | GWD | 1000 Genomes on GRCh38,1000 Genomes 30x on GRCh38,1000 Genomes phase 3 release,Gambian Genome Variation Project (GRCh38) |
| **HG04132** | female | SAME1839287 | BEB | Bengali | SAS | South Asian Ancestry | BEB | 1000 Genomes on GRCh38,1000 Genomes 30x on GRCh38 |
| **NA19091** | male | SAME123174 | JPT | Japanese | EAS | East Asian Ancestry | JPT | 1000 Genomes on GRCh38,1000 Genomes 30x on GRCh38,1000 Genomes phase 3 release |
| **HG02647** | male | SAME1839762 | GWD | Gambian Mandinka | AFR | African Ancestry | GWD | 1000 Genomes 30x on GRCh38 |
| **NA19096** | male | SAME123662 | YRI | Yoruba | AFR | African Ancestry | YRI | 1000 Genomes on GRCh38,1000 Genomes 30x on GRCh38,1000 Genomes phase 3 release,1000 Genomes phase 1 release,Geuvadis |
| **HG02786** | male | SAME1840292 | PJL | Punjabi | SAS | South Asian Ancestry | PJL | 1000 Genomes on GRCh38,1000 Genomes 30x on GRCh38,1000 Genomes phase 3 release |
| **HG04062** | female | SAME1839147 | ITU | Telugu | SAS | South Asian Ancestry | ITU | 1000 Genomes on GRCh38,1000 Genomes 30x on GRCh38,1000 Genomes phase 3 release |
| **NA19109** | female | SAME124544 | YRI | Yoruba | AFR | African Ancestry | YRI | 1000 Genomes 30x on GRCh38,1000 Genomes phase 3 release |
| **HG02793** | female | SAME1840415 | PJL | Punjabi | SAS | South Asian Ancestry | PJL | 1000 Genomes on GRCh38,1000 Genomes 30x on GRCh38,1000 Genomes phase 3 release |
| **NA18497** | male | SAME124277 | YRI | Yoruba | AFR | African Ancestry | YRI | 1000 Genomes 30x on GRCh38,1000 Genomes phase 3 release |
| **HG02798** | male | SAME1840387 | GWD | Gambian Mandinka | AFR | African Ancestry | GWD | 1000 Genomes on GRCh38,1000 Genomes 30x on GRCh38,1000 Genomes phase 3 release,Gambian Genome Variation Project (GRCh38) |
| **HG04144** | female | SAME1839189 | BEB | Bengali | SAS | South Asian Ancestry | BEB | 1000 Genomes on GRCh38,1000 Genomes 30x on GRCh38,1000 Genomes phase 3 release |
| **NA18500** | male | SAME123386 | YRI | Yoruba | AFR | African Ancestry | YRI | 1000 Genomes 30x on GRCh38,1000 Genomes phase 3 release |
| **HG02806** | female | SAME1840079 | GWD | Gambian Mandinka | AFR | African Ancestry | GWD | 1000 Genomes 30x on GRCh38 |
| **HG04149** | male | SAME1839120 | BEB | Bengali | SAS | South Asian Ancestry | BEB | 1000 Genomes on GRCh38,1000 Genomes 30x on GRCh38 |
| **NA18505** | female | SAME125062 | YRI | Yoruba | AFR | African Ancestry | YRI | 1000 Genomes on GRCh38,1000 Genomes 30x on GRCh38,1000 Genomes phase 3 release,1000 Genomes phase 1 release,Geuvadis |
| **HG02813** | male | SAME1839841 | GWD | Gambian Mandinka | AFR | African Ancestry | GWD | 1000 Genomes on GRCh38,1000 Genomes 30x on GRCh38,1000 Genomes phase 3 release,Gambian Genome Variation Project (GRCh38) |
| **HG04151** | female | SAME1840295 | BEB | Bengali | SAS | South Asian Ancestry | BEB | 1000 Genomes 30x on GRCh38 |
| **NA18517** | female | SAME124883 | YRI | Yoruba | AFR | African Ancestry | YRI | 1000 Genomes on GRCh38,1000 Genomes 30x on GRCh38,1000 Genomes phase 3 release,1000 Genomes phase 1 release,Geuvadis |
| **HG02818** | female | SAME1839810 | GWD | Gambian Mandinka | AFR | African Ancestry | GWD | 1000 Genomes 30x on GRCh38,Human Genome Structural Variation Consortium, Phase 2 |
| **HG04156** | female | SAME1840389 | BEB | Bengali | SAS | South Asian Ancestry | BEB | 1000 Genomes on GRCh38,1000 Genomes 30x on GRCh38,1000 Genomes phase 3 release |
| **NA18531** | female | SAME124483 | CHB | Han Chinese | EAS | East Asian Ancestry | CHB | 1000 Genomes on GRCh38,1000 Genomes 30x on GRCh38,1000 Genomes phase 3 release,1000 Genomes phase 1 release |
| **HG02971** | male | SAME1840255 | ESN | Esan | AFR | African Ancestry | ESN | 1000 Genomes on GRCh38,1000 Genomes 30x on GRCh38,1000 Genomes phase 3 release |
| **HG04175** | female | SAME1839980 | BEB | Bengali | SAS | South Asian Ancestry | BEB | 1000 Genomes 30x on GRCh38 |
| **NA18536** | male | SAME124486 | CHB | Han Chinese | EAS | East Asian Ancestry | CHB | 1000 Genomes on GRCh38,1000 Genomes 30x on GRCh38,1000 Genomes phase 3 release,1000 Genomes phase 1 release |
| **HG02976** | female | SAME1840223 | ESN | Esan | AFR | African Ancestry | ESN | 1000 Genomes on GRCh38,1000 Genomes 30x on GRCh38,1000 Genomes phase 3 release |
| **HG04182** | male | SAME1840036 | BEB | Bengali | SAS | South Asian Ancestry | BEB | 1000 Genomes on GRCh38,1000 Genomes 30x on GRCh38,1000 Genomes phase 3 release |
| **NA18543** | male | SAME124268 | CHB | Han Chinese | EAS | East Asian Ancestry | CHB | 1000 Genomes on GRCh38,1000 Genomes 30x on GRCh38,1000 Genomes phase 3 release,1000 Genomes phase 1 release |
| **HG02983** | female | SAME1840009 | GWD | Gambian Mandinka | AFR | African Ancestry | GWD | 1000 Genomes on GRCh38,1000 Genomes 30x on GRCh38,1000 Genomes phase 3 release,Gambian Genome Variation Project (GRCh38) |
| **HG04187** | male | SAME1839763 | BEB | Bengali | SAS | South Asian Ancestry | BEB | 1000 Genomes 30x on GRCh38 |
| **NA18548** | male | SAME124261 | CHB | Han Chinese | EAS | East Asian Ancestry | CHB | 1000 Genomes on GRCh38,1000 Genomes 30x on GRCh38,1000 Genomes phase 3 release,1000 Genomes phase 1 release |
| **HG03006** | male | SAME1840242 | BEB | Bengali,Bengali | SAS | South Asia (SGDP),South Asian Ancestry | BengaliSGDP,BEB | 1000 Genomes on GRCh38,Simons Genome Diversity Project,1000 Genomes 30x on GRCh38,1000 Genomes phase 3 release |
| **NA18538** | female | SAME123774 | CHB | Han Chinese | EAS | East Asian Ancestry | CHB | 1000 Genomes on GRCh38,1000 Genomes 30x on GRCh38,1000 Genomes phase 3 release,1000 Genomes phase 1 release |
| **NA18550** | female | SAME124087 | CHB | Han Chinese | EAS | East Asian Ancestry | CHB | 1000 Genomes on GRCh38,1000 Genomes 30x on GRCh38,1000 Genomes phase 3 release,1000 Genomes phase 1 release,90 Han Chinese high coverage genomes |
| **HG03018** | male | SAME1840053 | PJL | Punjabi | SAS | South Asian Ancestry | PJL | 1000 Genomes on GRCh38,1000 Genomes 30x on GRCh38,1000 Genomes phase 3 release |

| | | | | | | | | |
|---|---|---|---|---|---|---|---|---|
| **NA18545** | female | SAME124266 | CHB | Han Chinese | EAS | East Asian Ancestry | CHB | 1000 Genomes on GRCh38,1000 Genomes 30x on GRCh38,1000 Genomes phase 3 release,1000 Genomes phase 1 release,90 Han Chinese high coverage genomes |
| **NA19116** | female | SAME124911 | YRI | Yoruba | AFR | African Ancestry | YRI | 1000 Genomes on GRCh38,1000 Genomes 30x on GRCh38,1000 Genomes phase 3 release,1000 Genomes phase 1 release,Geuvadis |
| **HG02654** | male | SAME1839530 | PJL | Punjabi | SAS | South Asian Ancestry | PJL | 1000 Genomes on GRCh38,1000 Genomes 30x on GRCh38,1000 Genomes phase 3 release |
| **NA18552** | female | SAME124089 | CHB | Han Chinese | EAS | East Asian Ancestry | CHB | 1000 Genomes on GRCh38,1000 Genomes 30x on GRCh38,1000 Genomes phase 3 release,1000 Genomes phase 1 release,90 Han Chinese high coverage genomes |
| **NA19123** | male | SAME125013 | YRI | Yoruba | AFR | African Ancestry | YRI | 1000 Genomes 30x on GRCh38,1000 Genomes phase 3 release |
| **HG02659** | female | SAME1839590 | PJL | Punjabi | SAS | South Asian Ancestry | PJL | 1000 Genomes 30x on GRCh38,1000 Genomes phase 3 release |
| **NA18557** | male | SAME124092 | CHB | Han Chinese | EAS | East Asian Ancestry | CHB | 1000 Genomes on GRCh38,1000 Genomes 30x on GRCh38,1000 Genomes phase 3 release,1000 Genomes phase 1 release |
| **NA19128** | male | SAME125022 | YRI | Yoruba | AFR | African Ancestry | YRI | 1000 Genomes 30x on GRCh38,1000 Genomes phase 3 release |
| **HG02661** | female | SAME1839667 | PJL | Punjabi | SAS | South Asian Ancestry | PJL | 1000 Genomes on GRCh38,1000 Genomes 30x on GRCh38,1000 Genomes phase 3 release |
| **NA18564** | female | SAME123921 | CHB | Han Chinese | EAS | East Asian Ancestry | CHB | 1000 Genomes on GRCh38,1000 Genomes 30x on GRCh38,1000 Genomes phase 3 release,1000 Genomes phase 1 release,90 Han Chinese high coverage genomes |
| **NA19130** | male | SAME125229 | YRI | Yoruba | AFR | African Ancestry | YRI | 1000 Genomes on GRCh38,1000 Genomes 30x on GRCh38,1000 Genomes phase 3 release,1000 Genomes phase 1 release,Geuvadis |
| **HG02666** | male | SAME1839386 | GWD | Gambian Mandinka | AFR | African Ancestry | GWD | 1000 Genomes on GRCh38,1000 Genomes 30x on GRCh38,1000 Genomes phase 3 release,Gambian Genome Variation Project (GRCh38) |
| **NA18571** | female | SAME123740 | CHB | Han Chinese | EAS | East Asian Ancestry | CHB | 1000 Genomes on GRCh38,1000 Genomes 30x on GRCh38,1000 Genomes phase 3 release,1000 Genomes phase 1 release,90 Han Chinese high coverage genomes |
| **NA19142** | male | SAME124610 | YRI | Yoruba | AFR | African Ancestry | YRI | 1000 Genomes 30x on GRCh38,1000 Genomes phase 3 release |
| **HG02678** | male | SAME1839477 | GWD | Gambian Mandinka | AFR | African Ancestry | GWD | 1000 Genomes on GRCh38,1000 Genomes 30x on GRCh38,1000 Genomes phase 3 release,Gambian Genome Variation Project (GRCh38) |
| **NA19147** | female | SAME124613 | YRI | Yoruba | AFR | African Ancestry | YRI | 1000 Genomes on GRCh38,1000 Genomes 30x on GRCh38,1000 Genomes phase 3 release,1000 Genomes phase 1 release,Geuvadis |
| **HG02680** | female | SAME1839308 | GWD | Gambian Mandinka | AFR | African Ancestry | GWD | 1000 Genomes 30x on GRCh38 |
| **NA18595** | female | SAME123228 | CHB | Han Chinese | EAS | East Asian Ancestry | CHB | 1000 Genomes on GRCh38,1000 Genomes 30x on GRCh38,1000 Genomes phase 3 release,1000 Genomes phase 1 release |
| **NA19154** | male | SAME124835 | YRI | Yoruba | AFR | African Ancestry | YRI | 1000 Genomes 30x on GRCh38,1000 Genomes phase 3 release |
| **HG02685** | female | SAME1839286 | PJL | Punjabi | SAS | South Asian Ancestry | PJL | 1000 Genomes on GRCh38,1000 Genomes 30x on GRCh38,1000 Genomes phase 3 release |
| **NA18603** | male | SAME124875 | CHB | Han Chinese | EAS | East Asian Ancestry | CHB | 1000 Genomes on GRCh38,1000 Genomes 30x on GRCh38,1000 Genomes phase 3 release,1000 Genomes phase 1 release,90 Han Chinese high coverage genomes |
| **NA19159** | female | SAME124716 | YRI | Yoruba | AFR | African Ancestry | YRI | 1000 Genomes on GRCh38,1000 Genomes 30x on GRCh38,1000 Genomes phase 3 release,1000 Genomes phase 1 release,Geuvadis |
| **HG02692** | female | SAME1839078 | PJL | Punjabi | SAS | South Asian Ancestry | PJL | 1000 Genomes 30x on GRCh38 |
| **NA18608** | male | SAME124779 | CHB | Han Chinese | EAS | East Asian Ancestry | CHB | 1000 Genomes on GRCh38,1000 Genomes 30x on GRCh38,1000 Genomes phase 3 release,1000 Genomes phase 1 release,90 Han Chinese high coverage genomes |
| **NA19161** | male | SAME123757 | YRI | Yoruba | AFR | African Ancestry | YRI | 1000 Genomes 30x on GRCh38,1000 Genomes phase 3 release |
| **HG02697** | female | SAME1839142 | PJL | Punjabi | SAS | South Asian Ancestry | PJL | 1000 Genomes on GRCh38,1000 Genomes 30x on GRCh38,1000 Genomes phase 3 release |
| **NA18610** | female | SAME123222 | CHB | Han Chinese | EAS | East Asian Ancestry | CHB | 1000 Genomes on GRCh38,1000 Genomes 30x on GRCh38,1000 Genomes phase 3 release,1000 Genomes phase 1 release |
| **HG02700** | female | SAME1840159 | PJL | Punjabi | SAS | South Asian Ancestry | PJL | 1000 Genomes on GRCh38,1000 Genomes 30x on GRCh38,1000 Genomes phase 3 release |
| **NA18615** | female | SAME123653 | CHB | Han Chinese | EAS | East Asian Ancestry | CHB | 1000 Genomes on GRCh38,1000 Genomes 30x on GRCh38,1000 Genomes phase 3 release,1000 Genomes phase 1 release |
| **NA19173** | male | SAME124421 | YRI | Yoruba | AFR | African Ancestry | YRI | 1000 Genomes 30x on GRCh38,1000 Genomes phase 3 release |
| **HG02717** | male | SAME1839976 | GWD | Gambian Mandinka | AFR | African Ancestry | GWD | 1000 Genomes 30x on GRCh38 |
| **NA18622** | male | SAME125260 | CHB | Han Chinese | EAS | East Asian Ancestry | CHB | 1000 Genomes on GRCh38,1000 Genomes 30x on GRCh38,1000 Genomes phase 3 release,1000 Genomes phase 1 release,90 Han Chinese high coverage genomes |
| **NA18858** | female | SAME124730 | YRI | Yoruba | AFR | African Ancestry | YRI | 1000 Genomes on GRCh38,1000 Genomes 30x on GRCh38,1000 Genomes phase 3 release,1000 Genomes phase 1 release,Geuvadis |
| **HG02724** | male | SAME1839740 | PJL | Punjabi,Punjabi | SAS | South Asia (SGDP),South Asian Ancestry | PunjabiSGDP,PJL | 1000 Genomes on GRCh38,Simons Genome Diversity Project,1000 Genomes 30x on GRCh38,1000 Genomes phase 3 release |
| **NA18627** | female | SAME124106 | CHB | Han Chinese | EAS | East Asian Ancestry | CHB | 1000 Genomes on GRCh38,1000 Genomes 30x on GRCh38,1000 Genomes phase 3 release,1000 Genomes phase 1 release |
| **NA18860** | male | SAME124906 | YRI | Yoruba | AFR | African Ancestry | YRI | 1000 Genomes 30x on GRCh38,1000 Genomes phase 3 release |
| **HG02729** | male | SAME1839715 | PJL | Punjabi | SAS | South Asian Ancestry | PJL | 1000 Genomes 30x on GRCh38,1000 Genomes phase 3 release |
| **NA18634** | female | SAME124697 | CHB | Han Chinese | EAS | East Asian Ancestry | CHB | 1000 Genomes on GRCh38,1000 Genomes 30x on GRCh38,1000 Genomes phase 3 release,1000 Genomes phase 1 release |
| **NA18865** | male | SAME124902 | YRI | Yoruba | AFR | African Ancestry | YRI | 1000 Genomes on GRCh38,1000 Genomes 30x on GRCh38,1000 Genomes phase 3 release |
| **HG02731** | female | SAME1839886 | PJL | Punjabi | SAS | South Asian Ancestry | PJL | 1000 Genomes on GRCh38,1000 Genomes 30x on GRCh38,1000 Genomes phase 3 release |
| **NA18639** | male | SAME124694 | CHB | Han Chinese | EAS | East Asian Ancestry | CHB | 1000 Genomes on GRCh38,1000 Genomes 30x on GRCh38,1000 Genomes phase 3 release,1000 Genomes phase 1 release |
| **NA18872** | male | SAME125099 | YRI | Yoruba | AFR | African Ancestry | YRI | 1000 Genomes 30x on GRCh38,1000 Genomes phase 3 release |
| **HG02736** | male | SAME1839860 | PJL | Punjabi | SAS | South Asian Ancestry | PJL | 1000 Genomes on GRCh38,1000 Genomes 30x on GRCh38,1000 Genomes phase 3 release |
| **NA18641** | female | SAME122979 | CHB | Han Chinese | EAS | East Asian Ancestry | CHB | 1000 Genomes on GRCh38,1000 Genomes 30x on GRCh38,1000 Genomes phase 3 release,1000 Genomes phase 1 release |
| **NA18877** | male | SAME125104 | YRI | Yoruba | AFR | African Ancestry | YRI | 1000 Genomes on GRCh38,1000 Genomes 30x on GRCh38,1000 Genomes phase 3 release |
| **HG02762** | male | SAME1839258 | GWD | Gambian Mandinka | AFR | African Ancestry | GWD | 1000 Genomes on GRCh38,1000 Genomes 30x on GRCh38,Gambian Genome Variation Project (GRCh38) |
| **NA18646** | female | SAME122978 | CHB | Han Chinese | EAS | East Asian Ancestry | CHB | 1000 Genomes on GRCh38,1000 Genomes 30x on GRCh38,1000 Genomes phase 3 release |
| **NA18909** | female | SAME123561 | YRI | Yoruba | AFR | African Ancestry | YRI | 1000 Genomes on GRCh38,1000 Genomes 30x on GRCh38,1000 Genomes phase 3 release,1000 Genomes phase 1 release,Geuvadis |
| **HG02774** | male | SAME1839028 | PJL | Punjabi | SAS | South Asian Ancestry | PJL | 1000 Genomes on GRCh38,1000 Genomes 30x on GRCh38,1000 Genomes phase 3 release |
| **NA18747** | male | SAME123916 | CHB | Han Chinese | EAS | East Asian Ancestry | CHB | 1000 Genomes on GRCh38,1000 Genomes 30x on GRCh38,1000 Genomes phase 3 release,1000 Genomes phase 1 release |
| **NA18911** | male | SAME123401 | YRI | Yoruba | AFR | African Ancestry | YRI | 1000 Genomes 30x on GRCh38,1000 Genomes phase 3 release |
| **NA19031** | male | SAME123543 | LWK | Luhya | AFR | African Ancestry | LWK | 1000 Genomes on GRCh38,1000 Genomes 30x on GRCh38,1000 Genomes phase 3 release,1000 Genomes phase 1 release |
| **NA19248** | male | SAME123681 | YRI | Yoruba | AFR | African Ancestry | YRI | 1000 Genomes on GRCh38,1000 Genomes 30x on GRCh38,1000 Genomes phase 3 release,1000 Genomes phase 1 release,Geuvadis |
| **HG02820** | female | SAME1839591 | GWD | Gambian Mandinka | AFR | African Ancestry | GWD | 1000 Genomes on GRCh38,1000 Genomes 30x on GRCh38,1000 Genomes phase 3 release,Gambian Genome Variation Project (GRCh38) |
| **NA19036** | female | SAME124666 | LWK | Luhya | AFR | African Ancestry | LWK | 1000 Genomes on GRCh38,1000 Genomes 30x on GRCh38,Human Genome Structural Variation Consortium, Phase 2,1000 Genomes phase 3 release,1000 Genomes ph |
| **HG02837** | female | SAME1839422 | GWD | Gambian Mandinka | AFR | African Ancestry | GWD | 1000 Genomes on GRCh38,1000 Genomes 30x on GRCh38,1000 Genomes phase 3 release,Gambian Genome Variation Project (GRCh38) |
| **NA19043** | male | SAME125243 | LWK | Luhya | AFR | African Ancestry | LWK | 1000 Genomes on GRCh38,1000 Genomes 30x on GRCh38,1000 Genomes phase 3 release |
| **NA19318** | male | SAME124753 | LWK | Luhya | AFR | African Ancestry | LWK | 1000 Genomes on GRCh38,1000 Genomes 30x on GRCh38,1000 Genomes phase 3 release,1000 Genomes phase 1 release |
| **HG02851** | male | SAME1839352 | GWD | Gambian Mandinka | AFR | African Ancestry | GWD | 1000 Genomes on GRCh38,1000 Genomes 30x on GRCh38,1000 Genomes phase 3 release,Gambian Genome Variation Project (GRCh38) |
| **NA19055** | male | SAME125027 | JPT | Japanese | EAS | East Asian Ancestry | JPT | 1000 Genomes on GRCh38,1000 Genomes 30x on GRCh38,1000 Genomes phase 3 release,1000 Genomes phase 1 release |
| **NA19320** | female | SAME124570 | LWK | Luhya | AFR | African Ancestry | LWK | 1000 Genomes on GRCh38,1000 Genomes 30x on GRCh38,1000 Genomes phase 3 release |
| **HG02856** | male | SAME1839327 | GWD | Gambian Mandinka | AFR | African Ancestry | GWD | 1000 Genomes 30x on GRCh38 |
| **NA19062** | male | SAME123005 | JPT | Japanese | EAS | East Asian Ancestry | JPT | 1000 Genomes on GRCh38,1000 Genomes 30x on GRCh38,1000 Genomes phase 3 release,1000 Genomes phase 1 release |
| **NA19332** | female | SAME123723 | LWK | Luhya | AFR | African Ancestry | LWK | 1000 Genomes on GRCh38,1000 Genomes 30x on GRCh38,1000 Genomes phase 3 release,1000 Genomes phase 1 release |
| **HG02870** | female | SAME1840320 | GWD | Gambian Mandinka | AFR | African Ancestry | GWD | 1000 Genomes on GRCh38,1000 Genomes 30x on GRCh38,1000 Genomes phase 3 release,Gambian Genome Variation Project (GRCh38) |
| **NA19067** | male | SAME123835 | JPT | Japanese | EAS | East Asian Ancestry | JPT | 1000 Genomes on GRCh38,1000 Genomes 30x on GRCh38,1000 Genomes phase 3 release,1000 Genomes phase 1 release |
| **NA19351** | female | SAME122854 | LWK | Luhya | AFR | African Ancestry | LWK | 1000 Genomes on GRCh38,1000 Genomes 30x on GRCh38,1000 Genomes phase 3 release,1000 Genomes phase 1 release |
| **HG02882** | female | SAME1840171 | GWD | Gambian Mandinka | AFR | African Ancestry | GWD | 1000 Genomes on GRCh38,1000 Genomes 30x on GRCh38,1000 Genomes phase 3 release,Gambian Genome Variation Project (GRCh38) |
| **NA19074** | female | SAME122817 | JPT | Japanese | EAS | East Asian Ancestry | JPT | 1000 Genomes on GRCh38,1000 Genomes 30x on GRCh38,1000 Genomes phase 3 release,1000 Genomes phase 1 release |
| **NA19375** | male | SAME125116 | LWK | Luhya | AFR | African Ancestry | LWK | 1000 Genomes on GRCh38,1000 Genomes 30x on GRCh38,1000 Genomes phase 3 release,1000 Genomes phase 1 release |
| **HG02887** | male | SAME1840147 | GWD | Gambian Mandinka | AFR | African Ancestry | GWD | 1000 Genomes on GRCh38,1000 Genomes 30x on GRCh38,1000 Genomes phase 3 release,Gambian Genome Variation Project (GRCh38) |
| **NA19079** | male | SAME122823 | JPT | Japanese | EAS | East Asian Ancestry | JPT | 1000 Genomes on GRCh38,1000 Genomes 30x on GRCh38,1000 Genomes phase 3 release,1000 Genomes phase 1 release |
| **NA18980** | female | SAME123527 | JPT | Japanese | EAS | East Asian Ancestry | JPT | 1000 Genomes on GRCh38,1000 Genomes 30x on GRCh38,1000 Genomes phase 3 release,1000 Genomes phase 1 release |
| **HG03025** | female | SAME1839711 | GWD | Gambian Mandinka | AFR | African Ancestry | GWD | 1000 Genomes on GRCh38,1000 Genomes 30x on GRCh38,1000 Genomes phase 3 release,Gambian Genome Variation Project (GRCh38) |
| **NA19081** | female | SAME123369 | JPT | Japanese | EAS | East Asian Ancestry | JPT | 1000 Genomes on GRCh38,1000 Genomes 30x on GRCh38,1000 Genomes phase 3 release,1000 Genomes phase 1 release |
| **NA18985** | male | SAME123413 | JPT | Japanese | EAS | East Asian Ancestry | JPT | 1000 Genomes on GRCh38,1000 Genomes 30x on GRCh38,1000 Genomes phase 3 release,1000 Genomes phase 1 release |
| **HG03049** | female | SAME1839653 | GWD | Gambian Mandinka | AFR | African Ancestry | GWD | 1000 Genomes on GRCh38,1000 Genomes 30x on GRCh38,1000 Genomes phase 3 release,Gambian Genome Variation Project (GRCh38) |
| **NA19086** | male | SAME123366 | JPT | Japanese | EAS | East Asian Ancestry | JPT | 1000 Genomes on GRCh38,1000 Genomes 30x on GRCh38,1000 Genomes phase 3 release,1000 Genomes phase 1 release |
| **NA18992** | female | SAME124983 | JPT | Japanese | EAS | East Asian Ancestry | JPT | 1000 Genomes on GRCh38,1000 Genomes 30x on GRCh38,1000 Genomes phase 3 release,1000 Genomes phase 1 release |
| **HG03056** | male | SAME1839518 | MSL | Mende | AFR | African Ancestry | MSL | 1000 Genomes 30x on GRCh38 |
| **NA19093** | female | SAME123172 | YRI | Yoruba | AFR | African Ancestry | YRI | 1000 Genomes on GRCh38,1000 Genomes 30x on GRCh38,1000 Genomes phase 3 release,1000 Genomes phase 1 release,Geuvadis |
| **NA18997** | female | SAME124898 | JPT | Japanese | EAS | East Asian Ancestry | JPT | 1000 Genomes on GRCh38,1000 Genomes 30x on GRCh38,1000 Genomes phase 3 release,1000 Genomes phase 1 release |
| **HG02938** | male | SAME1839284 | ESN | Esan | AFR | African Ancestry | ESN | 1000 Genomes on GRCh38,1000 Genomes 30x on GRCh38,1000 Genomes phase 3 release |
| **NA19098** | male | SAME123633 | YRI | Yoruba | AFR | African Ancestry | YRI | 1000 Genomes on GRCh38,1000 Genomes 30x on GRCh38,1000 Genomes phase 3 release,1000 Genomes phase 1 release,Geuvadis |
| **NA19003** | female | SAME124480 | JPT | Japanese | EAS | East Asian Ancestry | JPT | 1000 Genomes on GRCh38,1000 Genomes 30x on GRCh38,1000 Genomes phase 3 release,1000 Genomes phase 1 release |
| **HG02945** | female | SAME1839075 | ESN | Esan | AFR | African Ancestry | ESN | 1000 Genomes 30x on GRCh38 |
| **NA12890** | female | SAME124116 | CEU | CEPH | EUR | European Ancestry | CEU | 1000 Genomes on GRCh38,1000 Genomes 30x on GRCh38,Illumina Platinum pedigree,1000 Genomes phase 3 release,1000 Genomes phase 1 release,Geuvadis |
| **NA19010** | female | SAME123989 | JPT | Japanese | EAS | East Asian Ancestry | JPT | 1000 Genomes on GRCh38,1000 Genomes 30x on GRCh38,1000 Genomes phase 3 release,1000 Genomes phase 1 release |
| **HG02952** | female | SAME1840275 | ESN | Esan | AFR | African Ancestry | ESN | 1000 Genomes on GRCh38,1000 Genomes 30x on GRCh38,1000 Genomes phase 3 release |
| **NA18487** | male | SAME124098 | YRI | Yoruba | AFR | African Ancestry | YRI | 1000 Genomes on GRCh38,1000 Genomes 30x on GRCh38,1000 Genomes phase 3 release,1000 Genomes phase 1 release,Geuvadis |
| **NA19027** | male | SAME124850 | LWK | Luhya | AFR | African Ancestry | LWK | 1000 Genomes on GRCh38,1000 Genomes 30x on GRCh38,1000 Genomes phase 3 release,1000 Genomes phase 1 release |
| **HG02964** | female | SAME1840122 | ESN | Esan | AFR | African Ancestry | ESN | 1000 Genomes on GRCh38,1000 Genomes 30x on GRCh38 |
| **NA18499** | female | SAME124059 | YRI | Yoruba | AFR | African Ancestry | YRI | 1000 Genomes on GRCh38,1000 Genomes 30x on GRCh38,1000 Genomes phase 3 release,1000 Genomes phase 1 release,Geuvadis |
| **NA19041** | male | SAME125241 | LWK | Luhya | AFR | African Ancestry | LWK | 1000 Genomes on GRCh38,1000 Genomes 30x on GRCh38,1000 Genomes phase 3 release,1000 Genomes phase 1 release |
| **NA18502** | female | SAME125055 | YRI | Yoruba | AFR | African Ancestry | YRI | 1000 Genomes on GRCh38,1000 Genomes 30x on GRCh38,1000 Genomes phase 3 release,1000 Genomes phase 1 release,Geuvadis |
| **NA19185** | female | SAME123858 | YRI | Yoruba | AFR | African Ancestry | YRI | 1000 Genomes on GRCh38,1000 Genomes 30x on GRCh38,1000 Genomes phase 3 release,1000 Genomes phase 1 release,Geuvadis |
| **HG03239** | female | SAME1839295 | PJL | Punjabi | SAS | South Asian Ancestry | PJL | 1000 Genomes 30x on GRCh38,1000 Genomes phase 3 release |
| **NA18507** | male | SAME125060 | YRI | Yoruba | AFR | African Ancestry | YRI | 1000 Genomes on GRCh38,1000 Genomes 30x on GRCh38,1000 Genomes phase 3 release,1000 Genomes phase 1 release |
| **NA19197** | female | SAME125309 | YRI | Yoruba | AFR | African Ancestry | YRI | 1000 Genomes on GRCh38,1000 Genomes 30x on GRCh38,1000 Genomes phase 3 release,1000 Genomes phase 1 release,Geuvadis |
| **HG03241** | female | SAME1839053 | GWD | Gambian Mandinka | AFR | African Ancestry | GWD | 1000 Genomes on GRCh38,1000 Genomes 30x on GRCh38,1000 Genomes phase 3 release,Gambian Genome Variation Project (GRCh38) |
| **NA18519** | male | SAME124888 | YRI | Yoruba | AFR | African Ancestry | YRI | 1000 Genomes on GRCh38,1000 Genomes 30x on GRCh38,1000 Genomes phase 3 release,1000 Genomes phase 1 release,Geuvadis |
| **NA19200** | male | SAME122989 | YRI | Yoruba | AFR | African Ancestry | YRI | 1000 Genomes on GRCh38,1000 Genomes 30x on GRCh38,1000 Genomes phase 3 release,1000 Genomes phase 1 release,Geuvadis |
| **HG03246** | male | SAME1839168 | GWD | Gambian Mandinka | AFR | African Ancestry | GWD | 1000 Genomes on GRCh38,1000 Genomes 30x on GRCh38,1000 Genomes phase 3 release,Gambian Genome Variation Project (GRCh38) |
| **NA18521** | male | SAME123566 | YRI | Yoruba | AFR | African Ancestry | YRI | 1000 Genomes 30x on GRCh38,1000 Genomes phase 3 release |
| **NA19205** | male | SAME122986 | YRI | Yoruba | AFR | African Ancestry | YRI | 1000 Genomes 30x on GRCh38,1000 Genomes phase 3 release |
| **HG03258** | male | SAME1840296 | GWD | Gambian Mandinka | AFR | African Ancestry | GWD | 1000 Genomes on GRCh38,1000 Genomes 30x on GRCh38,1000 Genomes phase 3 release,Gambian Genome Variation Project (GRCh38) |
| **NA18526** | female | SAME124673 | CHB | Han Chinese | EAS | East Asian Ancestry | CHB | 1000 Genomes on GRCh38,1000 Genomes 30x on GRCh38,1000 Genomes phase 3 release,1000 Genomes phase 1 release,90 Han Chinese high coverage genomes |
| **NA19224** | male | SAME125217 | YRI | Yoruba | AFR | African Ancestry | YRI | 1000 Genomes 30x on GRCh38,1000 Genomes phase 3 release |
| **HG03260** | male | SAME1840140 | GWD | Gambian Mandinka | AFR | African Ancestry | GWD | 1000 Genomes 30x on GRCh38 |
| **NA18533** | female | SAME124481 | CHB | Han Chinese | EAS | East Asian Ancestry | CHB | 1000 Genomes on GRCh38,1000 Genomes 30x on GRCh38,1000 Genomes phase 3 release |
| **NA19236** | male | SAME125001 | YRI | Yoruba | AFR | African Ancestry | YRI | 1000 Genomes on GRCh38,1000 Genomes 30x on GRCh38,1000 Genomes phase 3 release,1000 Genomes phase 1 release,Geuvadis |

| | | | | | | | | |
|---|---|---|---|---|---|---|---|---|
| **HG03265** | male | SAME1840167 | ESN | Esan | AFR | African Ancestry | ESN | 1000 Genomes on GRCh38,1000 Genomes 30x on GRCh38,1000 Genomes phase 3 release |
| **NA18968** | female | SAME123953 | JPT | Japanese | EAS | East Asian Ancestry | JPT | 1000 Genomes on GRCh38,1000 Genomes 30x on GRCh38,1000 Genomes phase 3 release,1000 Genomes phase 1 release |
| **NA20517** | female | SAME124557 | TSI | Toscani | EUR | European Ancestry | TSI | 1000 Genomes on GRCh38,1000 Genomes 30x on GRCh38,1000 Genomes phase 3 release,1000 Genomes phase 1 release,Geuvadis |
| **HG03272** | female | SAME1840199 | ESN | Esan | AFR | African Ancestry | ESN | 1000 Genomes 30x on GRCh38 |
| **NA18970** | male | SAME123764 | JPT | Japanese | EAS | East Asian Ancestry | JPT | 1000 Genomes on GRCh38,1000 Genomes 30x on GRCh38,1000 Genomes phase 3 release,1000 Genomes phase 1 release |
| **NA20524** | male | SAME124016 | TSI | Toscani | EUR | European Ancestry | TSI | 1000 Genomes on GRCh38,1000 Genomes 30x on GRCh38,1000 Genomes phase 3 release,1000 Genomes phase 1 release,Geuvadis |
| **HG03291** | female | SAME1839861 | ESN | Esan | AFR | African Ancestry | ESN | 1000 Genomes on GRCh38,1000 Genomes 30x on GRCh38,1000 Genomes phase 3 release |
| **NA18975** | female | SAME123767 | JPT | Japanese | EAS | East Asian Ancestry | JPT | 1000 Genomes on GRCh38,1000 Genomes 30x on GRCh38,1000 Genomes phase 3 release,1000 Genomes phase 1 release |
| **NA20529** | female | SAME124011 | TSI | Toscani | EUR | European Ancestry | TSI | 1000 Genomes on GRCh38,1000 Genomes 30x on GRCh38,1000 Genomes phase 3 release,1000 Genomes phase 1 release,Geuvadis |
| **HG03342** | female | SAME1840350 | ESN | Esan | AFR | African Ancestry | ESN | 1000 Genomes on GRCh38,1000 Genomes 30x on GRCh38,1000 Genomes phase 3 release |
| **NA18982** | male | SAME125073 | JPT | Japanese | EAS | East Asian Ancestry | JPT | 1000 Genomes on GRCh38,1000 Genomes 30x on GRCh38,1000 Genomes phase 3 release,1000 Genomes phase 1 release |
| **NA20531** | female | SAME124170 | TSI | Toscani | EUR | European Ancestry | TSI | 1000 Genomes on GRCh38,1000 Genomes 30x on GRCh38,1000 Genomes phase 3 release,1000 Genomes phase 1 release,Geuvadis |
| **HG03354** | female | SAME1840249 | ESN | Esan | AFR | African Ancestry | ESN | 1000 Genomes on GRCh38,1000 Genomes 30x on GRCh38,1000 Genomes phase 3 release |
| **NA18987** | female | SAME125070 | JPT | Japanese | EAS | East Asian Ancestry | JPT | 1000 Genomes on GRCh38,1000 Genomes 30x on GRCh38,1000 Genomes phase 3 release,1000 Genomes phase 1 release |
| **NA20536** | male | SAME124167 | TSI | Toscani | EUR | European Ancestry | TSI | 1000 Genomes on GRCh38,1000 Genomes 30x on GRCh38,1000 Genomes phase 3 release,1000 Genomes phase 1 release,Geuvadis |
| **NA18994** | male | SAME124897 | JPT | Japanese | EAS | East Asian Ancestry | JPT | 1000 Genomes on GRCh38,1000 Genomes 30x on GRCh38,1000 Genomes phase 3 release,1000 Genomes phase 1 release |
| **NA20543** | male | SAME125149 | TSI | Toscani | EUR | European Ancestry | TSI | 1000 Genomes on GRCh38,1000 Genomes 30x on GRCh38,1000 Genomes phase 3 release,1000 Genomes phase 1 release,Geuvadis |
| **HG03366** | female | SAME1840066 | ESN | Esan | AFR | African Ancestry | ESN | 1000 Genomes on GRCh38,1000 Genomes 30x on GRCh38,1000 Genomes phase 3 release |
| **NA18999** | female | SAME124890 | JPT | Japanese | EAS | East Asian Ancestry | JPT | 1000 Genomes on GRCh38,1000 Genomes 30x on GRCh38,1000 Genomes phase 3 release,1000 Genomes phase 1 release |
| **NA20581** | male | SAME122859 | TSI | Toscani | EUR | European Ancestry | TSI | 1000 Genomes on GRCh38,1000 Genomes 30x on GRCh38,1000 Genomes phase 3 release,1000 Genomes phase 1 release,Geuvadis |
| **HG03373** | male | SAME1839720 | ESN | Esan | AFR | African Ancestry | ESN | 1000 Genomes on GRCh38,1000 Genomes 30x on GRCh38 |
| **NA19000** | male | SAME124477 | JPT | Japanese | EAS | East Asian Ancestry | JPT | 1000 Genomes on GRCh38,1000 Genomes 30x on GRCh38,1000 Genomes phase 3 release,1000 Genomes phase 1 release |
| **NA20586** | male | SAME122864 | TSI | Toscani | EUR | European Ancestry | TSI | 1000 Genomes on GRCh38,1000 Genomes 30x on GRCh38,1000 Genomes phase 3 release,1000 Genomes phase 1 release,Geuvadis |
| **HG03164** | female | SAME1839121 | ESN | Esan | AFR | African Ancestry | ESN | 1000 Genomes 30x on GRCh38 |
| **NA19005** | male | SAME124474 | JPT | Japanese | EAS | East Asian Ancestry | JPT | 1000 Genomes on GRCh38,1000 Genomes 30x on GRCh38,1000 Genomes phase 3 release,1000 Genomes phase 1 release |
| **NA20752** | male | SAME124177 | TSI | Toscani | EUR | European Ancestry | TSI | 1000 Genomes on GRCh38,1000 Genomes 30x on GRCh38,1000 Genomes phase 3 release,1000 Genomes phase 1 release,Geuvadis |
| **HG03169** | male | SAME1839148 | ESN | Esan | AFR | African Ancestry | ESN | 1000 Genomes on GRCh38,1000 Genomes 30x on GRCh38,1000 Genomes phase 3 release |
| **NA19012** | male | SAME124260 | JPT | Japanese | EAS | East Asian Ancestry | JPT | 1000 Genomes on GRCh38,1000 Genomes 30x on GRCh38,1000 Genomes phase 3 release,1000 Genomes phase 1 release |
| **NA20757** | female | SAME124176 | TSI | Toscani | EUR | European Ancestry | TSI | 1000 Genomes on GRCh38,1000 Genomes 30x on GRCh38,1000 Genomes phase 3 release,1000 Genomes phase 1 release,Geuvadis |
| **NA19017** | female | SAME124259 | LWK | Luhya | AFR | African Ancestry | LWK | 1000 Genomes on GRCh38,1000 Genomes 30x on GRCh38,1000 Genomes phase 3 release |
| **NA20764** | female | SAME124368 | TSI | Toscani | EUR | European Ancestry | TSI | 1000 Genomes on GRCh38,1000 Genomes 30x on GRCh38,1000 Genomes phase 3 release |
| **NA19024** | female | SAME124851 | LWK | Luhya | AFR | African Ancestry | LWK | 1000 Genomes on GRCh38,1000 Genomes 30x on GRCh38,1000 Genomes phase 3 release |
| **NA20769** | female | SAME124371 | TSI | Toscani | EUR | European Ancestry | TSI | 1000 Genomes on GRCh38,1000 Genomes 30x on GRCh38,1000 Genomes phase 3 release,1000 Genomes phase 1 release,Geuvadis |
| **HG03190** | male | SAME1839984 | ESN | Esan | AFR | African Ancestry | ESN | 1000 Genomes on GRCh38,1000 Genomes 30x on GRCh38,1000 Genomes phase 3 release |
| **NA12718** | female | SAME124454 | CEU | CEPH | EUR | European Ancestry | CEU | 1000 Genomes on GRCh38,1000 Genomes 30x on GRCh38,1000 Genomes phase 3 release,1000 Genomes phase 1 release,Geuvadis |
| **NA20771** | female | SAME124577 | TSI | Toscani | EUR | European Ancestry | TSI | 1000 Genomes on GRCh38,1000 Genomes 30x on GRCh38,1000 Genomes phase 3 release,1000 Genomes phase 1 release,Geuvadis |
| **HG03121** | female | SAME1839563 | ESN | Esan | AFR | African Ancestry | ESN | 1000 Genomes on GRCh38,1000 Genomes 30x on GRCh38,1000 Genomes phase 3 release |
| **NA12749** | female | SAME124828 | CEU | CEPH | EUR | European Ancestry | CEU | 1000 Genomes on GRCh38,1000 Genomes 30x on GRCh38,1000 Genomes phase 3 release,1000 Genomes phase 1 release,Geuvadis |
| **HG03126** | female | SAME1839683 | ESN | Esan | AFR | African Ancestry | ESN | 1000 Genomes on GRCh38,1000 Genomes 30x on GRCh38,1000 Genomes phase 3 release |
| **NA12751** | female | SAME123515 | CEU | CEPH | EUR | European Ancestry | CEU | 1000 Genomes on GRCh38,1000 Genomes 30x on GRCh38,1000 Genomes phase 3 release,1000 Genomes phase 1 release,Geuvadis |
| **NA19818** | male | SAME124677 | ASW | African Ancestry SW | AFR | African Ancestry | ASW | 1000 Genomes on GRCh38,1000 Genomes 30x on GRCh38,1000 Genomes phase 3 release,1000 Genomes phase 1 release |
| **HG03133** | male | SAME1839375 | ESN | Esan | AFR | African Ancestry | ESN | 1000 Genomes on GRCh38,1000 Genomes 30x on GRCh38,1000 Genomes phase 3 release |
| **NA12763** | female | SAME123872 | CEU | CEPH | EUR | European Ancestry | CEU | 1000 Genomes on GRCh38,1000 Genomes 30x on GRCh38,1000 Genomes phase 3 release,1000 Genomes phase 1 release,Geuvadis |
| **NA19902** | female | SAME125051 | ASW | African Ancestry SW | AFR | African Ancestry | ASW | 1000 Genomes 30x on GRCh38,1000 Genomes phase 3 release |
| **NA12775** | male | SAME124257 | CEU | CEPH | EUR | European Ancestry | CEU | 1000 Genomes on GRCh38,1000 Genomes 30x on GRCh38,1000 Genomes phase 3 release,1000 Genomes phase 1 release,Geuvadis |
| **NA19914** | female | SAME125273 | ASW | African Ancestry SW | AFR | African Ancestry | ASW | 1000 Genomes on GRCh38,1000 Genomes 30x on GRCh38,1000 Genomes phase 3 release,1000 Genomes phase 1 release |
| **HG03157** | male | SAME1839083 | ESN | Esan | AFR | African Ancestry | ESN | 1000 Genomes on GRCh38,1000 Genomes 30x on GRCh38,1000 Genomes phase 3 release |
| **NA12802** | female | SAME122828 | CEU | CEPH | EUR | European Ancestry | CEU | 1000 Genomes 30x on GRCh38,1000 Genomes phase 3 release |
| **NA19919** | male | SAME125276 | ASW | African Ancestry SW | AFR | African Ancestry | ASW | 1000 Genomes 30x on GRCh38,1000 Genomes phase 3 release |
| **HG03296** | male | SAME1839887 | ESN | Esan | AFR | African Ancestry | ESN | 1000 Genomes 30x on GRCh38 |
| **NA12814** | male | SAME123021 | CEU | CEPH | EUR | European Ancestry | CEU | 1000 Genomes on GRCh38,1000 Genomes 30x on GRCh38,1000 Genomes phase 3 release,1000 Genomes phase 1 release,Geuvadis |
| **NA19921** | female | SAME124712 | ASW | African Ancestry SW | AFR | African Ancestry | ASW | 1000 Genomes on GRCh38,1000 Genomes 30x on GRCh38,1000 Genomes phase 3 release,1000 Genomes phase 1 release |
| **HG03304** | male | SAME1839490 | ESN | Esan | AFR | African Ancestry | ESN | 1000 Genomes on GRCh38,1000 Genomes 30x on GRCh38,1000 Genomes phase 3 release |
| **NA12864** | male | SAME123191 | CEU | CEPH | EUR | European Ancestry | CEU | 1000 Genomes 30x on GRCh38,1000 Genomes phase 3 release |
| **NA19983** | female | SAME123902 | ASW | African Ancestry SW | AFR | African Ancestry | ASW | 1000 Genomes 30x on GRCh38,Human Genome Structural Variation Consortium, Phase 2,1000 Genomes phase 3 release |
| **HG03309** | female | SAME1839520 | ESN | Esan | AFR | African Ancestry | ESN | 1000 Genomes on GRCh38,1000 Genomes 30x on GRCh38 |
| **NA18879** | male | SAME125108 | YRI | Yoruba | AFR | African Ancestry | YRI | 1000 Genomes on GRCh38,1000 Genomes 30x on GRCh38,1000 Genomes phase 3 release |
| **NA20276** | female | SAME123614 | ASW | African Ancestry SW | AFR | African Ancestry | ASW | 1000 Genomes on GRCh38,1000 Genomes 30x on GRCh38,1000 Genomes phase 3 release,1000 Genomes phase 1 release |
| **HG03311** | male | SAME1839206 | ESN | Esan | AFR | African Ancestry | ESN | 1000 Genomes on GRCh38,1000 Genomes 30x on GRCh38,1000 Genomes phase 3 release |
| **NA18881** | female | SAME125308 | YRI | Yoruba | AFR | African Ancestry | YRI | 1000 Genomes on GRCh38,1000 Genomes 30x on GRCh38,1000 Genomes phase 3 release |
| **NA20783** | male | SAME123607 | TSI | Toscani | EUR | European Ancestry | TSI | 1000 Genomes on GRCh38,1000 Genomes 30x on GRCh38,1000 Genomes phase 3 release,1000 Genomes phase 1 release,Geuvadis |
| **HG03195** | female | SAME1839919 | ESN | Esan | AFR | African Ancestry | ESN | 1000 Genomes on GRCh38,1000 Genomes 30x on GRCh38,1000 Genomes phase 3 release |
| **NA18906** | female | SAME123562 | YRI | Yoruba | AFR | African Ancestry | YRI | 1000 Genomes 30x on GRCh38,1000 Genomes phase 3 release |
| **NA20790** | female | SAME123815 | TSI | Toscani | EUR | European Ancestry | TSI | 1000 Genomes on GRCh38,1000 Genomes 30x on GRCh38,1000 Genomes phase 3 release,1000 Genomes phase 1 release,Geuvadis |
| **NA18913** | male | SAME123403 | YRI | Yoruba | AFR | African Ancestry | YRI | 1000 Genomes 30x on GRCh38,1000 Genomes phase 3 release |
| **NA20795** | female | SAME123818 | TSI | Toscani | EUR | European Ancestry | TSI | 1000 Genomes on GRCh38,1000 Genomes 30x on GRCh38,1000 Genomes phase 3 release,1000 Genomes phase 1 release,Geuvadis |
| **HG03234** | male | SAME1839357 | PJL | Punjabi | SAS | South Asian Ancestry | PJL | 1000 Genomes on GRCh38,1000 Genomes 30x on GRCh38,1000 Genomes phase 3 release |
| **NA18925** | male | SAME123203 | YRI | Yoruba | AFR | African Ancestry | YRI | 1000 Genomes 30x on GRCh38,1000 Genomes phase 3 release |
| **NA20803** | male | SAME125367 | TSI | Toscani | EUR | European Ancestry | TSI | 1000 Genomes on GRCh38,1000 Genomes 30x on GRCh38,1000 Genomes phase 3 release,1000 Genomes phase 1 release,Geuvadis |
| **HG03099** | female | SAME1840146 | ESN | Esan | AFR | African Ancestry | ESN | 1000 Genomes on GRCh38,1000 Genomes 30x on GRCh38,1000 Genomes phase 3 release |
| **NA18944** | male | SAME124314 | JPT | Japanese | EAS | East Asian Ancestry | JPT | 1000 Genomes on GRCh38,1000 Genomes 30x on GRCh38,1000 Genomes phase 3 release,1000 Genomes phase 1 release |
| **NA20808** | female | SAME122813 | TSI | Toscani | EUR | European Ancestry | TSI | 1000 Genomes on GRCh38,1000 Genomes 30x on GRCh38,1000 Genomes phase 3 release,1000 Genomes phase 1 release,Geuvadis |
| **NA18949** | female | SAME124317 | JPT | Japanese | EAS | East Asian Ancestry | JPT | 1000 Genomes on GRCh38,1000 Genomes 30x on GRCh38,1000 Genomes phase 3 release,1000 Genomes phase 1 release |
| **NA20810** | male | SAME125192 | TSI | Toscani | EUR | European Ancestry | TSI | 1000 Genomes on GRCh38,1000 Genomes 30x on GRCh38,1000 Genomes phase 3 release,1000 Genomes phase 1 release,Geuvadis |
| **HG03114** | female | SAME1839907 | ESN | Esan | AFR | African Ancestry | ESN | 1000 Genomes on GRCh38,1000 Genomes 30x on GRCh38,1000 Genomes phase 3 release |
| **NA18951** | female | SAME124111 | JPT | Japanese | EAS | East Asian Ancestry | JPT | 1000 Genomes on GRCh38,1000 Genomes 30x on GRCh38,1000 Genomes phase 3 release,1000 Genomes phase 1 release |
| **NA20815** | male | SAME125187 | TSI | Toscani | EUR | European Ancestry | TSI | 1000 Genomes on GRCh38,1000 Genomes 30x on GRCh38,1000 Genomes phase 3 release,1000 Genomes phase 1 release,Geuvadis |
| **HG03119** | female | SAME1839833 | ESN | Esan | AFR | African Ancestry | ESN | 1000 Genomes 30x on GRCh38 |
| **NA18956** | female | SAME124110 | JPT | Japanese | EAS | East Asian Ancestry | JPT | 1000 Genomes on GRCh38,1000 Genomes 30x on GRCh38,1000 Genomes phase 3 release,1000 Genomes phase 1 release |
| **NA20822** | female | SAME123073 | TSI | Toscani | EUR | European Ancestry | TSI | 1000 Genomes on GRCh38,1000 Genomes 30x on GRCh38,1000 Genomes phase 3 release |
| **HG03378** | female | SAME1839743 | MSL | Mende | AFR | African Ancestry | MSL | 1000 Genomes on GRCh38,1000 Genomes 30x on GRCh38,1000 Genomes phase 3 release |
| **NA18963** | female | SAME123959 | JPT | Japanese | EAS | East Asian Ancestry | JPT | 1000 Genomes on GRCh38,1000 Genomes 30x on GRCh38,1000 Genomes phase 3 release,1000 Genomes phase 1 release |
| **NA20827** | male | SAME123071 | TSI | Toscani | EUR | European Ancestry | TSI | 1000 Genomes on GRCh38,1000 Genomes 30x on GRCh38,1000 Genomes phase 3 release |
| **HG03380** | female | SAME1839569 | MSL | Mende | AFR | African Ancestry | MSL | 1000 Genomes on GRCh38,1000 Genomes 30x on GRCh38,1000 Genomes phase 3 release |
| **NA19101** | male | SAME123955 | YRI | Yoruba | AFR | African Ancestry | YRI | 1000 Genomes 30x on GRCh38,1000 Genomes phase 3 release |
| **NA20846** | male | SAME123431 | GIH | Gujarati | SAS | South Asian Ancestry | GIH | 1000 Genomes on GRCh38,1000 Genomes 30x on GRCh38,1000 Genomes phase 3 release |
| **HG03385** | male | SAME1839594 | MSL | Mende | AFR | African Ancestry | MSL | 1000 Genomes on GRCh38,1000 Genomes 30x on GRCh38,1000 Genomes phase 3 release |
| **NA19113** | male | SAME123838 | YRI | Yoruba | AFR | African Ancestry | YRI | 1000 Genomes on GRCh38,1000 Genomes 30x on GRCh38,1000 Genomes phase 3 release,1000 Genomes phase 1 release,Geuvadis |
| **NA20853** | female | SAME123243 | GIH | Gujarati | SAS | South Asian Ancestry | GIH | 1000 Genomes on GRCh38,1000 Genomes 30x on GRCh38,1000 Genomes phase 3 release |
| **HG03397** | male | SAME1839400 | MSL | Mende | AFR | African Ancestry | MSL | 1000 Genomes on GRCh38,1000 Genomes 30x on GRCh38,1000 Genomes phase 3 release |
| **NA19118** | female | SAME122943 | YRI | Yoruba | AFR | African Ancestry | YRI | 1000 Genomes on GRCh38,1000 Genomes 30x on GRCh38,1000 Genomes phase 3 release,1000 Genomes phase 1 release,Geuvadis |
| **HG03063** | male | SAME1839226 | MSL | Mende | AFR | African Ancestry | MSL | 1000 Genomes on GRCh38,1000 Genomes 30x on GRCh38,1000 Genomes phase 3 release |
| **NA19120** | male | SAME125016 | YRI | Yoruba | AFR | African Ancestry | YRI | 1000 Genomes 30x on GRCh38,1000 Genomes phase 3 release |
| **NA19471** | female | SAME123117 | LWK | Luhya | AFR | African Ancestry | LWK | 1000 Genomes on GRCh38,1000 Genomes 30x on GRCh38,1000 Genomes phase 3 release,1000 Genomes phase 1 release |
| **HG03082** | female | SAME1840316 | MSL | Mende | AFR | African Ancestry | MSL | 1000 Genomes on GRCh38,1000 Genomes 30x on GRCh38,1000 Genomes phase 3 release |
| **NA19132** | female | SAME125231 | YRI | Yoruba | AFR | African Ancestry | YRI | 1000 Genomes 30x on GRCh38,1000 Genomes phase 3 release |
| **NA19654** | female | SAME123128 | MXL | Mexican Ancestry | AMR | American Ancestry | MXL | 1000 Genomes on GRCh38,1000 Genomes 30x on GRCh38,1000 Genomes phase 3 release,1000 Genomes phase 1 release |
| **HG03486** | female | SAME1839474 | MSL | Mende | AFR | African Ancestry | MSL | 1000 Genomes 30x on GRCh38,Human Genome Structural Variation Consortium, Phase 2,1000 Genomes phase 3 release |
| **NA19137** | female | SAME125235 | YRI | Yoruba | AFR | African Ancestry | YRI | 1000 Genomes on GRCh38,1000 Genomes 30x on GRCh38,1000 Genomes phase 3 release,1000 Genomes phase 1 release,Geuvadis |
| **NA19659** | female | SAME123131 | MXL | Mexican Ancestry | AMR | American Ancestry | MXL | 1000 Genomes 30x on GRCh38,1000 Genomes phase 3 release |
| **NA19144** | male | SAME123373 | YRI | Yoruba | AFR | African Ancestry | YRI | 1000 Genomes on GRCh38,1000 Genomes 30x on GRCh38,1000 Genomes phase 3 release,1000 Genomes phase 1 release,Geuvadis |
| **NA19661** | male | SAME122959 | MXL | Mexican Ancestry | AMR | American Ancestry | MXL | 1000 Genomes on GRCh38,1000 Genomes 30x on GRCh38,1000 Genomes phase 3 release,1000 Genomes phase 1 release |
| **NA19149** | female | SAME124605 | YRI | Yoruba | AFR | African Ancestry | YRI | 1000 Genomes on GRCh38,1000 Genomes 30x on GRCh38,1000 Genomes phase 3 release,1000 Genomes phase 1 release,Geuvadis |
| **NA19678** | female | SAME122797 | MXL | Mexican Ancestry | AMR | American Ancestry | MXL | 1000 Genomes on GRCh38,1000 Genomes 30x on GRCh38,1000 Genomes phase 3 release,1000 Genomes phase 1 release |
| **HG03518** | male | SAME1840169 | ESN | Esan | AFR | African Ancestry | ESN | 1000 Genomes on GRCh38,1000 Genomes 30x on GRCh38,1000 Genomes phase 3 release |
| **NA19151** | female | SAME124839 | YRI | Yoruba | AFR | African Ancestry | YRI | 1000 Genomes 30x on GRCh38,1000 Genomes phase 3 release |
| **NA19680** | female | SAME122937 | MXL | Mexican Ancestry | AMR | American Ancestry | MXL | 1000 Genomes 30x on GRCh38,1000 Genomes phase 3 release |
| **HG03520** | female | SAME1840186 | ESN | Esan | AFR | African Ancestry | ESN | 1000 Genomes on GRCh38,1000 Genomes 30x on GRCh38,1000 Genomes phase 3 release |
| **NA19175** | male | SAME124887 | YRI | Yoruba | AFR | African Ancestry | YRI | 1000 Genomes on GRCh38,1000 Genomes 30x on GRCh38,1000 Genomes phase 3 release,1000 Genomes phase 1 release,Geuvadis |
| **NA19685** | male | SAME122942 | MXL | Mexican Ancestry | AMR | American Ancestry | MXL | 1000 Genomes on GRCh38,1000 Genomes 30x on GRCh38,1000 Genomes phase 3 release,1000 Genomes phase 1 release |
| **HG03607** | female | SAME1840254 | BEB | Bengali | SAS | South Asian Ancestry | BEB | 1000 Genomes on GRCh38,1000 Genomes 30x on GRCh38,1000 Genomes phase 3 release |

| | | | | | | | | |
|---|---|---|---|---|---|---|---|---|
| **NA19199** | female | SAME124050 | YRI | Yoruba | AFR | African Ancestry | YRI | 1000 Genomes 30x on GRCh38,1000 Genomes phase 3 release |
| **NA19700** | male | SAME124233 | ASW | African Ancestry SW | AFR | African Ancestry | ASW | 1000 Genomes on GRCh38,1000 Genomes 30x on GRCh38,1000 Genomes phase 3 release,1000 Genomes phase 1 release |
| **HG03619** | female | SAME1840064 | PJL | Punjabi | SAS | South Asian Ancestry | PJL | 1000 Genomes on GRCh38,1000 Genomes 30x on GRCh38,1000 Genomes phase 3 release |
| **NA19202** | female | SAME122987 | YRI | Yoruba | AFR | African Ancestry | YRI | 1000 Genomes 30x on GRCh38,1000 Genomes phase 3 release |
| **NA19705** | male | SAME124235 | ASW | African Ancestry SW | AFR | African Ancestry | ASW | 1000 Genomes 30x on GRCh38,1000 Genomes phase 3 release |
| **NA19207** | male | SAME122984 | YRI | Yoruba | AFR | African Ancestry | YRI | 1000 Genomes on GRCh38,1000 Genomes 30x on GRCh38,1000 Genomes phase 3 release,1000 Genomes phase 1 release,Geuvadis |
| **NA19712** | female | SAME124447 | ASW | African Ancestry SW | AFR | African Ancestry | ASW | 1000 Genomes on GRCh38,1000 Genomes 30x on GRCh38,1000 Genomes phase 3 release,1000 Genomes phase 1 release |
| **HG03626** | female | SAME1839723 | PJL | Punjabi | SAS | South Asian Ancestry | PJL | 1000 Genomes 30x on GRCh38 |
| **NA19214** | female | SAME125386 | YRI | Yoruba | AFR | African Ancestry | YRI | 1000 Genomes on GRCh38,1000 Genomes 30x on GRCh38,1000 Genomes phase 3 release,1000 Genomes phase 1 release,Geuvadis |
| **NA19717** | male | SAME124451 | MXL | Mexican Ancestry | AMR | American Ancestry | MXL | 1000 Genomes on GRCh38,1000 Genomes 30x on GRCh38,1000 Genomes phase 3 release,1000 Genomes phase 1 release |
| **HG03633** | male | SAME1839571 | PJL | Punjabi | SAS | South Asian Ancestry | PJL | 1000 Genomes on GRCh38,1000 Genomes 30x on GRCh38,1000 Genomes phase 3 release |
| **NA18855** | female | SAME124735 | YRI | Yoruba | AFR | African Ancestry | YRI | 1000 Genomes 30x on GRCh38,1000 Genomes phase 3 release |
| **NA19724** | male | SAME124690 | MXL | Mexican Ancestry | AMR | American Ancestry | MXL | 1000 Genomes 30x on GRCh38,1000 Genomes phase 3 release |
| **NA18862** | male | SAME125118 | YRI | Yoruba | AFR | African Ancestry | YRI | 1000 Genomes 30x on GRCh38,1000 Genomes phase 3 release |
| **NA19729** | male | SAME124701 | MXL | Mexican Ancestry | AMR | American Ancestry | MXL | 1000 Genomes on GRCh38,1000 Genomes 30x on GRCh38,1000 Genomes phase 3 release,1000 Genomes phase 1 release |
| **HG03683** | female | SAME1840325 | STU | Tamil | SAS | South Asian Ancestry | STU | 1000 Genomes 30x on GRCh38,Human Genome Structural Variation Consortium, Phase 2 |
| **NA18867** | female | SAME124904 | YRI | Yoruba | AFR | African Ancestry | YRI | 1000 Genomes on GRCh38,1000 Genomes 30x on GRCh38,1000 Genomes phase 3 release,1000 Genomes phase 1 release,Geuvadis |
| **NA19731** | female | SAME124870 | MXL | Mexican Ancestry | AMR | American Ancestry | MXL | 1000 Genomes on GRCh38,1000 Genomes 30x on GRCh38,1000 Genomes phase 3 release,1000 Genomes phase 1 release |
| **HG03688** | male | SAME1840265 | STU | Tamil | SAS | South Asian Ancestry | STU | 1000 Genomes 30x on GRCh38 |
| **NA18874** | male | SAME125105 | YRI | Yoruba | AFR | African Ancestry | YRI | 1000 Genomes on GRCh38,1000 Genomes 30x on GRCh38,1000 Genomes phase 3 release,1000 Genomes phase 1 release |
| **NA19748** | female | SAME124729 | MXL | Mexican Ancestry | AMR | American Ancestry | MXL | 1000 Genomes 30x on GRCh38,1000 Genomes phase 3 release |
| **HG03690** | female | SAME1840365 | STU | Tamil | SAS | South Asian Ancestry | STU | 1000 Genomes on GRCh38,1000 Genomes 30x on GRCh38,1000 Genomes phase 3 release |
| **NA19221** | female | SAME125221 | YRI | Yoruba | AFR | African Ancestry | YRI | 1000 Genomes 30x on GRCh38,1000 Genomes phase 3 release |
| **NA19750** | male | SAME123698 | MXL | Mexican Ancestry | AMR | American Ancestry | MXL | 1000 Genomes on GRCh38,1000 Genomes 30x on GRCh38,1000 Genomes phase 3 release,1000 Genomes phase 1 release |
| **HG03695** | male | SAME1840128 | STU | Tamil | SAS | South Asian Ancestry | STU | 1000 Genomes on GRCh38,1000 Genomes 30x on GRCh38,1000 Genomes phase 3 release |
| **NA19226** | male | SAME125219 | YRI | Yoruba | AFR | African Ancestry | YRI | 1000 Genomes 30x on GRCh38,1000 Genomes phase 3 release |
| **NA20339** | female | SAME124201 | ASW | African Ancestry SW | AFR | African Ancestry | ASW | 1000 Genomes on GRCh38,1000 Genomes 30x on GRCh38,1000 Genomes phase 3 release,1000 Genomes phase 1 release |
| **HG03640** | female | SAME1839621 | PJL | Punjabi | SAS | South Asian Ancestry | PJL | 1000 Genomes on GRCh38,1000 Genomes 30x on GRCh38,1000 Genomes phase 3 release |
| **NA19238** | female | SAME124999 | YRI | Yoruba | AFR | African Ancestry | YRI | 1000 Genomes on GRCh38,1000 Genomes 30x on GRCh38,Human Genome Structural Variation Consortium, Phase 2,The Human Genome Structural Variation Consortiu |
| **HG03645** | female | SAME1839641 | STU | Tamil | SAS | South Asian Ancestry | STU | 1000 Genomes on GRCh38,1000 Genomes 30x on GRCh38,1000 Genomes phase 3 release |
| **NA19240** | female | SAME123688 | YRI | Yoruba | AFR | African Ancestry | YRI | 1000 Genomes on GRCh38,1000 Genomes 30x on GRCh38,Human Genome Structural Variation Consortium, Phase 2,The Human Genome Structural Variation Consortiu |
| **NA20346** | male | SAME124402 | ASW | African Ancestry SW | AFR | African Ancestry | ASW | 1000 Genomes on GRCh38,1000 Genomes 30x on GRCh38,1000 Genomes phase 3 release,1000 Genomes phase 1 release |
| **HG03652** | male | SAME1839505 | PJL | Punjabi | SAS | South Asian Ancestry | PJL | 1000 Genomes on GRCh38,1000 Genomes 30x on GRCh38,1000 Genomes phase 3 release |
| **NA20358** | male | SAME124600 | ASW | African Ancestry SW | AFR | African Ancestry | ASW | 1000 Genomes 30x on GRCh38,1000 Genomes phase 3 release |
| **NA19257** | female | SAME123495 | YRI | Yoruba | AFR | African Ancestry | YRI | 1000 Genomes on GRCh38,1000 Genomes 30x on GRCh38,1000 Genomes phase 3 release,1000 Genomes phase 1 release,Geuvadis |
| **HG03669** | female | SAME1839345 | PJL | Punjabi | SAS | South Asian Ancestry | PJL | 1000 Genomes 30x on GRCh38 |
| **NA19308** | male | SAME124985 | LWK | Luhya | AFR | African Ancestry | LWK | 1000 Genomes on GRCh38,1000 Genomes 30x on GRCh38,1000 Genomes phase 3 release,1000 Genomes phase 1 release |
| **NA20505** | female | SAME124359 | TSI | Toscani | EUR | European Ancestry | TSI | 1000 Genomes on GRCh38,1000 Genomes 30x on GRCh38,1000 Genomes phase 3 release,1000 Genomes phase 1 release,Geuvadis |
| **HG03582** | male | SAME1839271 | MSL | Mende | AFR | African Ancestry | MSL | 1000 Genomes on GRCh38,1000 Genomes 30x on GRCh38,1000 Genomes phase 3 release |
| **NA19310** | female | SAME123449 | LWK | Luhya | AFR | African Ancestry | LWK | 1000 Genomes on GRCh38,1000 Genomes 30x on GRCh38,1000 Genomes phase 3 release,1000 Genomes phase 1 release |
| **NA20512** | male | SAME124560 | TSI | Toscani | EUR | European Ancestry | TSI | 1000 Genomes on GRCh38,1000 Genomes 30x on GRCh38,1000 Genomes phase 3 release,1000 Genomes phase 1 release,Geuvadis |
| **NA19315** | female | SAME123310 | LWK | Luhya | AFR | African Ancestry | LWK | 1000 Genomes on GRCh38,1000 Genomes 30x on GRCh38,1000 Genomes phase 3 release,1000 Genomes phase 1 release |
| **HG03594** | male | SAME1839181 | BEB | Bengali | SAS | South Asian Ancestry | BEB | 1000 Genomes on GRCh38,1000 Genomes 30x on GRCh38,1000 Genomes phase 3 release |
| **NA19327** | female | SAME123473 | LWK | Luhya | AFR | African Ancestry | LWK | 1000 Genomes on GRCh38,1000 Genomes 30x on GRCh38,1000 Genomes phase 3 release,1000 Genomes phase 1 release |
| **NA19394** | male | SAME124489 | LWK | Luhya | AFR | African Ancestry | LWK | 1000 Genomes on GRCh38,1000 Genomes 30x on GRCh38,1000 Genomes phase 3 release,1000 Genomes phase 1 release |
| **NA19334** | male | SAME123722 | LWK | Luhya | AFR | African Ancestry | LWK | 1000 Genomes on GRCh38,1000 Genomes 30x on GRCh38,1000 Genomes phase 3 release,1000 Genomes phase 1 release |
| **NA19399** | female | SAME124498 | LWK | Luhya | AFR | African Ancestry | LWK | 1000 Genomes on GRCh38,1000 Genomes 30x on GRCh38,1000 Genomes phase 3 release,1000 Genomes phase 1 release |
| **NA19438** | female | SAME123848 | LWK | Luhya | AFR | African Ancestry | LWK | 1000 Genomes on GRCh38,1000 Genomes 30x on GRCh38,1000 Genomes phase 3 release,1000 Genomes phase 1 release |
| **HG03703** | female | SAME1840032 | PJL | Punjabi | SAS | South Asian Ancestry | PJL | 1000 Genomes on GRCh38,1000 Genomes 30x on GRCh38,1000 Genomes phase 3 release |
| **NA20502** | female | SAME124358 | TSI | Toscani | EUR | European Ancestry | TSI | 1000 Genomes on GRCh38,1000 Genomes 30x on GRCh38,1000 Genomes phase 3 release,1000 Genomes phase 1 release,Geuvadis |
| **NA19440** | female | SAME122951 | LWK | Luhya | AFR | African Ancestry | LWK | 1000 Genomes on GRCh38,1000 Genomes 30x on GRCh38,1000 Genomes phase 3 release,1000 Genomes phase 1 release |
| **HG03708** | male | SAME1839773 | PJL | Punjabi | SAS | South Asian Ancestry | PJL | 1000 Genomes on GRCh38,1000 Genomes 30x on GRCh38,1000 Genomes phase 3 release |
| **NA20507** | female | SAME124361 | TSI | Toscani | EUR | European Ancestry | TSI | 1000 Genomes on GRCh38,1000 Genomes 30x on GRCh38,1000 Genomes phase 3 release,1000 Genomes phase 1 release,Geuvadis |
| **NA19445** | female | SAME122948 | LWK | Luhya | AFR | African Ancestry | LWK | 1000 Genomes on GRCh38,1000 Genomes 30x on GRCh38,1000 Genomes phase 3 release,1000 Genomes phase 1 release |
| **HG03710** | male | SAME1839891 | PJL | Punjabi | SAS | South Asian Ancestry | PJL | 1000 Genomes 30x on GRCh38 |
| **NA20514** | female | SAME124558 | TSI | Toscani | EUR | European Ancestry | TSI | 1000 Genomes on GRCh38,1000 Genomes 30x on GRCh38,1000 Genomes phase 3 release,1000 Genomes phase 1 release,Geuvadis |
| **NA19452** | male | SAME125414 | LWK | Luhya | AFR | African Ancestry | LWK | 1000 Genomes on GRCh38,1000 Genomes 30x on GRCh38,1000 Genomes phase 3 release,1000 Genomes phase 1 release |
| **NA20519** | male | SAME123179 | TSI | Toscani | EUR | European Ancestry | TSI | 1000 Genomes on GRCh38,1000 Genomes 30x on GRCh38,1000 Genomes phase 3 release,1000 Genomes phase 1 release,Geuvadis |
| **NA19457** | female | SAME125409 | LWK | Luhya | AFR | African Ancestry | LWK | 1000 Genomes on GRCh38,1000 Genomes 30x on GRCh38,1000 Genomes phase 3 release,1000 Genomes phase 1 release |
| **NA20521** | male | SAME124018 | TSI | Toscani | EUR | European Ancestry | TSI | 1000 Genomes on GRCh38,1000 Genomes 30x on GRCh38,1000 Genomes phase 3 release,1000 Genomes phase 1 release,Geuvadis |
| **NA19755** | female | SAME123695 | MXL | Mexican Ancestry | AMR | American Ancestry | MXL | 1000 Genomes on GRCh38,1000 Genomes 30x on GRCh38,1000 Genomes phase 3 release,1000 Genomes phase 1 release |
| **HG03556** | male | SAME1839553 | MSL | Mende | AFR | African Ancestry | MSL | 1000 Genomes on GRCh38,1000 Genomes 30x on GRCh38,1000 Genomes phase 3 release |
| **NA19762** | male | SAME123897 | MXL | Mexican Ancestry | AMR | American Ancestry | MXL | 1000 Genomes on GRCh38,1000 Genomes 30x on GRCh38,1000 Genomes phase 3 release,1000 Genomes phase 1 release |
| **HG03563** | female | SAME1839421 | MSL | Mende | AFR | African Ancestry | MSL | 1000 Genomes on GRCh38,1000 Genomes 30x on GRCh38,1000 Genomes phase 3 release |
| **NA20533** | female | SAME124172 | TSI | Toscani | EUR | European Ancestry | TSI | 1000 Genomes on GRCh38,1000 Genomes 30x on GRCh38,1000 Genomes phase 3 release,1000 Genomes phase 1 release |
| **NA19774** | male | SAME124064 | MXL | Mexican Ancestry | AMR | American Ancestry | MXL | 1000 Genomes on GRCh38,1000 Genomes 30x on GRCh38,1000 Genomes phase 3 release,1000 Genomes phase 1 release |
| **HG03575** | female | SAME1839236 | MSL | Mende | AFR | African Ancestry | MSL | 1000 Genomes on GRCh38,1000 Genomes 30x on GRCh38,1000 Genomes phase 3 release |
| **NA20538** | male | SAME124163 | TSI | Toscani | EUR | European Ancestry | TSI | 1000 Genomes on GRCh38,1000 Genomes 30x on GRCh38,1000 Genomes phase 3 release,1000 Genomes phase 1 release,Geuvadis |
| **NA19779** | female | SAME124061 | MXL | Mexican Ancestry | AMR | American Ancestry | MXL | 1000 Genomes on GRCh38,1000 Genomes 30x on GRCh38,1000 Genomes phase 3 release,1000 Genomes phase 1 release |
| **NA20540** | female | SAME125148 | TSI | Toscani | EUR | European Ancestry | TSI | 1000 Genomes on GRCh38,1000 Genomes 30x on GRCh38,1000 Genomes phase 3 release,1000 Genomes phase 1 release,Geuvadis |
| **NA19781** | female | SAME123146 | MXL | Mexican Ancestry | AMR | American Ancestry | MXL | 1000 Genomes 30x on GRCh38,1000 Genomes phase 3 release |
| **HG03436** | male | SAME1840120 | MSL | Mende | AFR | African Ancestry | MSL | 1000 Genomes on GRCh38,1000 Genomes 30x on GRCh38,1000 Genomes phase 3 release |
| **NA20588** | male | SAME122866 | TSI | Toscani | EUR | European Ancestry | TSI | 1000 Genomes on GRCh38,1000 Genomes 30x on GRCh38,1000 Genomes phase 3 release,1000 Genomes phase 1 release,Geuvadis |
| **NA19786** | male | SAME123149 | MXL | Mexican Ancestry | AMR | American Ancestry | MXL | 1000 Genomes on GRCh38,1000 Genomes 30x on GRCh38,1000 Genomes phase 3 release,1000 Genomes phase 1 release |
| **NA20754** | male | SAME124173 | TSI | Toscani | EUR | European Ancestry | TSI | 1000 Genomes on GRCh38,1000 Genomes 30x on GRCh38,1000 Genomes phase 3 release,1000 Genomes phase 1 release,Geuvadis |
| **HG03455** | female | SAME1839766 | MSL | Mende | AFR | African Ancestry | MSL | 1000 Genomes on GRCh38,1000 Genomes 30x on GRCh38,1000 Genomes phase 3 release |
| **NA20759** | male | SAME125094 | TSI | Toscani | EUR | European Ancestry | TSI | 1000 Genomes on GRCh38,1000 Genomes 30x on GRCh38,1000 Genomes phase 3 release,1000 Genomes phase 1 release,Geuvadis |
| **NA19346** | male | SAME124341 | LWK | Luhya | AFR | African Ancestry | LWK | 1000 Genomes on GRCh38,1000 Genomes 30x on GRCh38,1000 Genomes phase 3 release,1000 Genomes phase 1 release |
| **NA19360** | male | SAME124915 | LWK | Luhya | AFR | African Ancestry | LWK | 1000 Genomes on GRCh38,1000 Genomes 30x on GRCh38,1000 Genomes phase 3 release,1000 Genomes phase 1 release |
| **HG03479** | female | SAME1839623 | MSL | Mende | AFR | African Ancestry | MSL | 1000 Genomes on GRCh38,1000 Genomes 30x on GRCh38,1000 Genomes phase 3 release |
| **NA19372** | male | SAME125100 | LWK | Luhya | AFR | African Ancestry | LWK | 1000 Genomes on GRCh38,1000 Genomes 30x on GRCh38,1000 Genomes phase 3 release,1000 Genomes phase 1 release |
| **NA20334** | female | SAME124196 | ASW | African Ancestry SW | AFR | African Ancestry | ASW | 1000 Genomes on GRCh38,1000 Genomes 30x on GRCh38,1000 Genomes phase 3 release,1000 Genomes phase 1 release |
| **HG03931** | female | SAME1840305 | BEB | Bengali | SAS | South Asian Ancestry | BEB | 1000 Genomes on GRCh38,1000 Genomes 30x on GRCh38,1000 Genomes phase 3 release |
| **NA19377** | female | SAME124470 | LWK | Luhya | AFR | African Ancestry | LWK | 1000 Genomes on GRCh38,1000 Genomes 30x on GRCh38,1000 Genomes phase 3 release,1000 Genomes phase 1 release |
| **NA21142** | female | SAME123344 | GIH | Gujarati | SAS | South Asian Ancestry | GIH | 1000 Genomes on GRCh38,1000 Genomes 30x on GRCh38,1000 Genomes phase 3 release |
| **HG03943** | male | SAME1840361 | STU | Tamil | SAS | South Asian Ancestry | STU | 1000 Genomes on GRCh38,1000 Genomes 30x on GRCh38,1000 Genomes phase 3 release |
| **NA19384** | male | SAME124272 | LWK | Luhya | AFR | African Ancestry | LWK | 1000 Genomes on GRCh38,1000 Genomes 30x on GRCh38,1000 Genomes phase 3 release,1000 Genomes phase 1 release |
| **NA19391** | female | SAME124494 | LWK | Luhya | AFR | African Ancestry | LWK | 1000 Genomes on GRCh38,1000 Genomes 30x on GRCh38,1000 Genomes phase 3 release,1000 Genomes phase 1 release |
| **HG03950** | male | SAME1840233 | STU | Tamil | SAS | South Asian Ancestry | STU | 1000 Genomes on GRCh38,1000 Genomes 30x on GRCh38,1000 Genomes phase 3 release |
| **HG03955** | female | SAME1840181 | STU | Tamil | SAS | South Asian Ancestry | STU | 1000 Genomes on GRCh38,1000 Genomes 30x on GRCh38,1000 Genomes phase 3 release |
| **NA19404** | female | SAME123637 | LWK | Luhya | AFR | African Ancestry | LWK | 1000 Genomes on GRCh38,1000 Genomes 30x on GRCh38,1000 Genomes phase 3 release,1000 Genomes phase 1 release |
| **HG03823** | female | SAME1839456 | BEB | Bengali | SAS | South Asian Ancestry | BEB | 1000 Genomes on GRCh38,1000 Genomes 30x on GRCh38,1000 Genomes phase 3 release |
| **NA19428** | male | SAME124036 | LWK | Luhya | AFR | African Ancestry | LWK | 1000 Genomes on GRCh38,1000 Genomes 30x on GRCh38,1000 Genomes phase 3 release,1000 Genomes phase 1 release |
| **HG03830** | male | SAME1839341 | BEB | Bengali | SAS | South Asian Ancestry | BEB | 1000 Genomes on GRCh38,1000 Genomes 30x on GRCh38,1000 Genomes phase 3 release |
| **NA19430** | male | SAME123854 | LWK | Luhya | AFR | African Ancestry | LWK | 1000 Genomes on GRCh38,1000 Genomes 30x on GRCh38,1000 Genomes phase 3 release,1000 Genomes phase 1 release |
| **NA19435** | female | SAME123849 | LWK | Luhya | AFR | African Ancestry | LWK | 1000 Genomes on GRCh38,1000 Genomes 30x on GRCh38,1000 Genomes phase 3 release,1000 Genomes phase 1 release |
| **NA19454** | male | SAME125412 | LWK | Luhya | AFR | African Ancestry | LWK | 1000 Genomes on GRCh38,1000 Genomes 30x on GRCh38,1000 Genomes phase 3 release |
| **NA19461** | male | SAME123298 | LWK | Luhya | AFR | African Ancestry | LWK | 1000 Genomes on GRCh38,1000 Genomes 30x on GRCh38,1000 Genomes phase 3 release,1000 Genomes phase 1 release |
| **HG03753** | male | SAME1839062 | STU | Tamil | SAS | South Asian Ancestry | STU | 1000 Genomes on GRCh38,1000 Genomes 30x on GRCh38,1000 Genomes phase 3 release |
| **NA19466** | male | SAME123294 | LWK | Luhya | AFR | African Ancestry | LWK | 1000 Genomes on GRCh38,1000 Genomes 30x on GRCh38,1000 Genomes phase 3 release,1000 Genomes phase 1 release |
| **HG03760** | female | SAME1839105 | STU | Tamil | SAS | South Asian Ancestry | STU | 1000 Genomes on GRCh38,1000 Genomes 30x on GRCh38,1000 Genomes phase 3 release |
| **NA19473** | female | SAME123119 | LWK | Luhya | AFR | African Ancestry | LWK | 1000 Genomes on GRCh38,1000 Genomes 30x on GRCh38,1000 Genomes phase 3 release,1000 Genomes phase 1 release |
| **HG03765** | female | SAME1839131 | PJL | Punjabi | SAS | South Asian Ancestry | PJL | 1000 Genomes on GRCh38,1000 Genomes 30x on GRCh38,1000 Genomes phase 3 release |
| **NA19625** | female | SAME123655 | ASW | African Ancestry SW | AFR | African Ancestry | ASW | 1000 Genomes on GRCh38,1000 Genomes 30x on GRCh38,1000 Genomes phase 3 release,1000 Genomes phase 1 release |
| **HG03772** | female | SAME1840398 | ITU | Telugu | SAS | South Asian Ancestry | ITU | 1000 Genomes on GRCh38,1000 Genomes 30x on GRCh38,1000 Genomes phase 3 release |
| **NA19649** | male | SAME123442 | MXL | Mexican Ancestry | AMR | American Ancestry | MXL | 1000 Genomes on GRCh38,1000 Genomes 30x on GRCh38,1000 Genomes phase 3 release,1000 Genomes phase 1 release |
| **NA20904** | male | SAME122916 | GIH | Gujarati | SAS | South Asian Ancestry | GIH | 1000 Genomes on GRCh38,1000 Genomes 30x on GRCh38,1000 Genomes phase 3 release |
| **HG03722** | female | SAME1839664 | ITU | Telugu | SAS | South Asian Ancestry | ITU | 1000 Genomes on GRCh38,1000 Genomes 30x on GRCh38,1000 Genomes phase 3 release |
| **NA19651** | female | SAME123123 | MXL | Mexican Ancestry | AMR | American Ancestry | MXL | 1000 Genomes on GRCh38,1000 Genomes 30x on GRCh38,1000 Genomes phase 3 release,1000 Genomes phase 1 release |
| **HG03727** | male | SAME1839695 | ITU | Telugu | SAS | South Asian Ancestry | ITU | 1000 Genomes on GRCh38,1000 Genomes 30x on GRCh38,1000 Genomes phase 3 release |
| **NA19656** | female | SAME123130 | MXL | Mexican Ancestry | AMR | American Ancestry | MXL | 1000 Genomes 30x on GRCh38,1000 Genomes phase 3 release |

| | | | | | | | | |
|---|---|---|---|---|---|---|---|---|
| **NA20911** | male | SAME123101 | GIH | Gujarati | SAS | South Asian Ancestry | GIH | 1000 Genomes on GRCh38,1000 Genomes 30x on GRCh38,1000 Genomes phase 3 release |
| **HG03741** | female | SAME1839251 | STU | Tamil | SAS | South Asian Ancestry | STU | 1000 Genomes on GRCh38,1000 Genomes 30x on GRCh38,1000 Genomes phase 3 release |
| **NA19663** | female | SAME122961 | MXL | Mexican Ancestry | AMR | American Ancestry | MXL | 1000 Genomes on GRCh38,1000 Genomes 30x on GRCh38,1000 Genomes phase 3 release,1000 Genomes phase 1 release |
| **NA21089** | female | SAME123142 | GIH | Gujarati | SAS | South Asian Ancestry | GIH | 1000 Genomes on GRCh38,1000 Genomes 30x on GRCh38,1000 Genomes phase 3 release |
| **HG03746** | male | SAME1839200 | STU | Tamil | SAS | South Asian Ancestry | STU | 1000 Genomes on GRCh38,1000 Genomes 30x on GRCh38,1000 Genomes phase 3 release |
| **NA19670** | male | SAME122795 | MXL | Mexican Ancestry | AMR | American Ancestry | MXL | 1000 Genomes on GRCh38,1000 Genomes 30x on GRCh38,1000 Genomes phase 3 release,1000 Genomes phase 1 release |
| **NA21091** | male | SAME123321 | GIH | Gujarati | SAS | South Asian Ancestry | GIH | 1000 Genomes on GRCh38,1000 Genomes 30x on GRCh38,1000 Genomes phase 3 release |
| **HG03777** | male | SAME1840340 | ITU | Telugu | SAS | South Asian Ancestry | ITU | 1000 Genomes on GRCh38,1000 Genomes 30x on GRCh38,1000 Genomes phase 3 release |
| **NA19788** | female | SAME123153 | MXL | Mexican Ancestry | AMR | American Ancestry | MXL | 1000 Genomes on GRCh38,1000 Genomes 30x on GRCh38,1000 Genomes phase 3 release,1000 Genomes phase 1 release |
| **NA20884** | male | SAME124514 | GIH | Gujarati | SAS | South Asian Ancestry | GIH | 1000 Genomes on GRCh38,1000 Genomes 30x on GRCh38,1000 Genomes phase 3 release |
| **HG03784** | female | SAME1840210 | ITU | Telugu | SAS | South Asian Ancestry | ITU | 1000 Genomes on GRCh38,1000 Genomes 30x on GRCh38,1000 Genomes phase 3 release |
| **NA19790** | female | SAME123326 | MXL | Mexican Ancestry | AMR | American Ancestry | MXL | 1000 Genomes 30x on GRCh38,1000 Genomes phase 3 release |
| **NA20889** | male | SAME125305 | GIH | Gujarati | SAS | South Asian Ancestry | GIH | 1000 Genomes on GRCh38,1000 Genomes 30x on GRCh38,1000 Genomes phase 3 release |
| **HG03789** | female | SAME1840237 | ITU | Telugu | SAS | South Asian Ancestry | ITU | 1000 Genomes on GRCh38,1000 Genomes 30x on GRCh38,1000 Genomes phase 3 release |
| **NA19795** | male | SAME123329 | MXL | Mexican Ancestry | AMR | American Ancestry | MXL | 1000 Genomes on GRCh38,1000 Genomes 30x on GRCh38,1000 Genomes phase 3 release,1000 Genomes phase 1 release |
| **NA20891** | male | SAME123670 | GIH | Gujarati | SAS | South Asian Ancestry | GIH | 1000 Genomes on GRCh38,1000 Genomes 30x on GRCh38,1000 Genomes phase 3 release |
| **HG03796** | female | SAME1839928 | BEB | Bengali | SAS | South Asian Ancestry | BEB | 1000 Genomes on GRCh38,1000 Genomes 30x on GRCh38,1000 Genomes phase 3 release |
| **NA19834** | male | SAME125030 | ASW | African Ancestry SW | AFR | African Ancestry | ASW | 1000 Genomes on GRCh38,1000 Genomes 30x on GRCh38,1000 Genomes phase 3 release,1000 Genomes phase 1 release |
| **NA20896** | female | SAME123665 | GIH | Gujarati | SAS | South Asian Ancestry | GIH | 1000 Genomes on GRCh38,1000 Genomes 30x on GRCh38,1000 Genomes phase 3 release |
| **HG03804** | female | SAME1839533 | BEB | Bengali | SAS | South Asian Ancestry | BEB | 1000 Genomes 30x on GRCh38 |
| **NA19904** | male | SAME125047 | ASW | African Ancestry SW | AFR | African Ancestry | ASW | 1000 Genomes on GRCh38,1000 Genomes 30x on GRCh38,1000 Genomes phase 3 release,1000 Genomes phase 1 release |
| **NA21116** | male | SAME124186 | GIH | Gujarati | SAS | South Asian Ancestry | GIH | 1000 Genomes on GRCh38,1000 Genomes 30x on GRCh38,1000 Genomes phase 3 release |
| **HG03809** | male | SAME1839554 | BEB | Bengali | SAS | South Asian Ancestry | BEB | 1000 Genomes on GRCh38,1000 Genomes 30x on GRCh38,1000 Genomes phase 3 release |
| **NA19909** | female | SAME125052 | ASW | African Ancestry SW | AFR | African Ancestry | ASW | 1000 Genomes on GRCh38,1000 Genomes 30x on GRCh38,1000 Genomes phase 3 release,1000 Genomes phase 1 release |
| **NA21123** | male | SAME124031 | GIH | Gujarati | SAS | South Asian Ancestry | GIH | 1000 Genomes on GRCh38,1000 Genomes 30x on GRCh38,1000 Genomes phase 3 release |
| **NA19916** | male | SAME125271 | ASW | African Ancestry SW | AFR | African Ancestry | ASW | 1000 Genomes on GRCh38,1000 Genomes 30x on GRCh38,1000 Genomes phase 3 release,1000 Genomes phase 1 release |
| **NA21128** | male | SAME124022 | GIH | Gujarati | SAS | South Asian Ancestry | GIH | 1000 Genomes on GRCh38,1000 Genomes 30x on GRCh38,1000 Genomes phase 3 release |
| **HG03816** | female | SAME1839424 | BEB | Bengali | SAS | South Asian Ancestry | BEB | 1000 Genomes 30x on GRCh38 |
| **NA19923** | female | SAME124710 | ASW | African Ancestry SW | AFR | African Ancestry | ASW | 1000 Genomes on GRCh38,1000 Genomes 30x on GRCh38,1000 Genomes phase 3 release,1000 Genomes phase 1 release |
| **NA21130** | male | SAME123834 | GIH | Gujarati | SAS | South Asian Ancestry | GIH | 1000 Genomes on GRCh38,1000 Genomes 30x on GRCh38,1000 Genomes phase 3 release |
| **HG03905** | male | SAME1839503 | BEB | Bengali | SAS | South Asian Ancestry | BEB | 1000 Genomes on GRCh38,1000 Genomes 30x on GRCh38,1000 Genomes phase 3 release |
| **NA21135** | male | SAME123829 | GIH | Gujarati | SAS | South Asian Ancestry | GIH | 1000 Genomes on GRCh38,1000 Genomes 30x on GRCh38,1000 Genomes phase 3 release |
| **HG03917** | male | SAME1839323 | BEB | Bengali | SAS | South Asian Ancestry | BEB | 1000 Genomes on GRCh38,1000 Genomes 30x on GRCh38,1000 Genomes phase 3 release |
| **NA20127** | female | SAME122884 | ASW | African Ancestry SW | AFR | African Ancestry | ASW | 1000 Genomes on GRCh38,1000 Genomes 30x on GRCh38,1000 Genomes phase 3 release,1000 Genomes phase 1 release |
| **NA20858** | male | SAME123207 | GIH | Gujarati | SAS | South Asian Ancestry | GIH | 1000 Genomes on GRCh38,1000 Genomes 30x on GRCh38,1000 Genomes phase 3 release |
| **NA19675** | female | SAME122791 | MXL | Mexican Ancestry | AMR | American Ancestry | MXL | 1000 Genomes on GRCh38,1000 Genomes 30x on GRCh38,1000 Genomes phase 3 release,1000 Genomes phase 1 release |
| **NA20872** | female | SAME122842 | GIH | Gujarati | SAS | South Asian Ancestry | GIH | 1000 Genomes on GRCh38,1000 Genomes 30x on GRCh38,1000 Genomes phase 3 release |
| **HG03929** | male | SAME1839046 | BEB | Bengali | SAS | South Asian Ancestry | BEB | 1000 Genomes on GRCh38,1000 Genomes 30x on GRCh38 |
| **NA19682** | male | SAME122935 | MXL | Mexican Ancestry | AMR | American Ancestry | MXL | 1000 Genomes on GRCh38,1000 Genomes 30x on GRCh38,1000 Genomes phase 3 release,1000 Genomes phase 1 release |
| **NA20877** | female | SAME123598 | GIH | Gujarati | SAS | South Asian Ancestry | GIH | 1000 Genomes on GRCh38,1000 Genomes 30x on GRCh38,1000 Genomes phase 3 release |
| **HG03885** | male | SAME1840013 | STU | Tamil | SAS | South Asian Ancestry | STU | 1000 Genomes on GRCh38,1000 Genomes 30x on GRCh38,1000 Genomes phase 3 release |
| **NA19702** | male | SAME124231 | ASW | African Ancestry SW | AFR | African Ancestry | ASW | 1000 Genomes 30x on GRCh38,1000 Genomes phase 3 release |
| **NA21104** | male | SAME124436 | GIH | Gujarati | SAS | South Asian Ancestry | GIH | 1000 Genomes on GRCh38,1000 Genomes 30x on GRCh38,1000 Genomes phase 3 release |
| **HG03897** | female | SAME1839906 | STU | Tamil | SAS | South Asian Ancestry | STU | 1000 Genomes on GRCh38,1000 Genomes 30x on GRCh38,1000 Genomes phase 3 release |
| **NA19707** | female | SAME124234 | ASW | African Ancestry SW | AFR | African Ancestry | ASW | 1000 Genomes on GRCh38,1000 Genomes 30x on GRCh38,1000 Genomes phase 3 release,1000 Genomes phase 1 release |
| **NA21109** | male | SAME124443 | GIH | Gujarati | SAS | South Asian Ancestry | GIH | 1000 Genomes on GRCh38,1000 Genomes 30x on GRCh38,1000 Genomes phase 3 release |
| **HG03900** | male | SAME1839476 | STU | Tamil | SAS | South Asian Ancestry | STU | 1000 Genomes on GRCh38,1000 Genomes 30x on GRCh38,1000 Genomes phase 3 release |
| **NA21111** | male | SAME124179 | GIH | Gujarati | SAS | South Asian Ancestry | GIH | 1000 Genomes on GRCh38,1000 Genomes 30x on GRCh38,1000 Genomes phase 3 release |
| **HG03854** | male | SAME1840283 | STU | Tamil | SAS | South Asian Ancestry | STU | 1000 Genomes on GRCh38,1000 Genomes 30x on GRCh38,1000 Genomes phase 3 release |
| **NA19719** | female | SAME124445 | MXL | Mexican Ancestry | AMR | American Ancestry | MXL | 1000 Genomes on GRCh38,1000 Genomes 30x on GRCh38,1000 Genomes phase 3 release,1000 Genomes phase 1 release |
| **HG03861** | female | SAME1840156 | ITU | Telugu | SAS | South Asian Ancestry | ITU | 1000 Genomes on GRCh38,1000 Genomes 30x on GRCh38,1000 Genomes phase 3 release |
| **NA19721** | female | SAME125311 | MXL | Mexican Ancestry | AMR | American Ancestry | MXL | 1000 Genomes 30x on GRCh38,1000 Genomes phase 3 release |
| **HG03866** | male | SAME1840087 | ITU | Telugu | SAS | South Asian Ancestry | ITU | 1000 Genomes on GRCh38,1000 Genomes 30x on GRCh38,1000 Genomes phase 3 release |
| **NA19726** | male | SAME124692 | MXL | Mexican Ancestry | AMR | American Ancestry | MXL | 1000 Genomes on GRCh38,1000 Genomes 30x on GRCh38,1000 Genomes phase 3 release,1000 Genomes phase 1 release |
| **HG03873** | female | SAME1840205 | ITU | Telugu | SAS | South Asian Ancestry | ITU | 1000 Genomes on GRCh38,1000 Genomes 30x on GRCh38,1000 Genomes phase 3 release |
| **NA19733** | female | SAME124868 | MXL | Mexican Ancestry | AMR | American Ancestry | MXL | 1000 Genomes 30x on GRCh38,1000 Genomes phase 3 release |
| **NA07345** | female | SAME123570 | CEU | CEPH | EUR | European Ancestry | CEU | 1000 Genomes 30x on GRCh38,1000 Genomes phase 3 release |
| **NA19740** | female | SAME124719 | MXL | Mexican Ancestry | AMR | American Ancestry | MXL | 1000 Genomes on GRCh38,1000 Genomes 30x on GRCh38,1000 Genomes phase 3 release,1000 Genomes phase 1 release |
| **NA07357** | male | SAME123412 | CEU | CEPH | EUR | European Ancestry | CEU | 1000 Genomes on GRCh38,1000 Genomes 30x on GRCh38,1000 Genomes phase 3 release,1000 Genomes phase 1 release,Geuvadis |
| **NA19752** | female | SAME123700 | MXL | Mexican Ancestry | AMR | American Ancestry | MXL | 1000 Genomes on GRCh38,1000 Genomes 30x on GRCh38,1000 Genomes phase 3 release,1000 Genomes phase 1 release |
| **NA10831** | female | SAME124549 | CEU | CEPH | EUR | European Ancestry | CEU | 1000 Genomes 30x on GRCh38,1000 Genomes phase 3 release |
| **NA19757** | male | SAME123697 | MXL | Mexican Ancestry | AMR | American Ancestry | MXL | 1000 Genomes 30x on GRCh38,1000 Genomes phase 3 release |
| **NA10836** | female | SAME124545 | CEU | CEPH | EUR | European Ancestry | CEU | 1000 Genomes 30x on GRCh38,1000 Genomes phase 3 release |
| **NA19764** | female | SAME123898 | MXL | Mexican Ancestry | AMR | American Ancestry | MXL | 1000 Genomes on GRCh38,1000 Genomes 30x on GRCh38,1000 Genomes phase 3 release,1000 Genomes phase 1 release |
| **NA10843** | female | SAME124754 | CEU | CEPH | EUR | European Ancestry | CEU | 1000 Genomes 30x on GRCh38,1000 Genomes phase 3 release |
| **NA19771** | male | SAME124069 | MXL | Mexican Ancestry | AMR | American Ancestry | MXL | 1000 Genomes on GRCh38,1000 Genomes 30x on GRCh38,1000 Genomes phase 3 release,1000 Genomes phase 1 release |
| **NA19776** | female | SAME124062 | MXL | Mexican Ancestry | AMR | American Ancestry | MXL | 1000 Genomes on GRCh38,1000 Genomes 30x on GRCh38,1000 Genomes phase 3 release,1000 Genomes phase 1 release |
| **NA19783** | male | SAME123144 | MXL | Mexican Ancestry | AMR | American Ancestry | MXL | 1000 Genomes on GRCh38,1000 Genomes 30x on GRCh38,1000 Genomes phase 3 release,1000 Genomes phase 1 release |
| **NA06986** | male | SAME124764 | CEU | CEPH | EUR | European Ancestry | CEU | 1000 Genomes on GRCh38,1000 Genomes 30x on GRCh38,1000 Genomes phase 3 release,1000 Genomes phase 1 release,Geuvadis |
| **NA20278** | male | SAME123617 | ASW | African Ancestry SW | AFR | African Ancestry | ASW | 1000 Genomes on GRCh38,1000 Genomes 30x on GRCh38,1000 Genomes phase 3 release,1000 Genomes phase 1 release |
| **NA06993** | male | SAME124934 | CEU | CEPH | EUR | European Ancestry | CEU | 1000 Genomes 30x on GRCh38,1000 Genomes phase 3 release |
| **HG04042** | female | SAME1839203 | STU | Tamil | SAS | South Asian Ancestry | STU | 1000 Genomes on GRCh38,1000 Genomes 30x on GRCh38,1000 Genomes phase 3 release |
| **HG04047** | female | SAME1839221 | STU | Tamil | SAS | South Asian Ancestry | STU | 1000 Genomes on GRCh38,1000 Genomes 30x on GRCh38,1000 Genomes phase 3 release |
| **HG04054** | female | SAME1839094 | ITU | Telugu | SAS | South Asian Ancestry | ITU | 1000 Genomes on GRCh38,1000 Genomes 30x on GRCh38,1000 Genomes phase 3 release |
| **HG04059** | female | SAME1839031 | ITU | Telugu | SAS | South Asian Ancestry | ITU | 1000 Genomes on GRCh38,1000 Genomes 30x on GRCh38,1000 Genomes phase 3 release |
| **HG04061** | male | SAME1839140 | ITU | Telugu | SAS | South Asian Ancestry | ITU | 1000 Genomes on GRCh38,1000 Genomes 30x on GRCh38,1000 Genomes phase 3 release |
| **NA20317** | female | SAME123943 | ASW | African Ancestry SW | AFR | African Ancestry | ASW | 1000 Genomes on GRCh38,1000 Genomes 30x on GRCh38,1000 Genomes phase 3 release,1000 Genomes phase 1 release |
| **HG04080** | male | SAME1840211 | ITU | Telugu | SAS | South Asian Ancestry | ITU | 1000 Genomes on GRCh38,1000 Genomes 30x on GRCh38,1000 Genomes phase 3 release |
| **HG04148** | female | SAME1839115 | BEB | Bengali | SAS | South Asian Ancestry | BEB | 1000 Genomes 30x on GRCh38 |
| **HG04150** | female | SAME1840289 | BEB | Bengali | SAS | South Asian Ancestry | BEB | 1000 Genomes on GRCh38,1000 Genomes 30x on GRCh38 |
| **NA20348** | male | SAME124407 | ASW | African Ancestry SW | AFR | African Ancestry | ASW | 1000 Genomes on GRCh38,1000 Genomes 30x on GRCh38,1000 Genomes phase 3 release,1000 Genomes phase 1 release |
| **HG04155** | male | SAME1840381 | BEB | Bengali | SAS | South Asian Ancestry | BEB | 1000 Genomes on GRCh38,1000 Genomes 30x on GRCh38,1000 Genomes phase 3 release |
| **HG04162** | female | SAME1840098 | BEB | Bengali | SAS | South Asian Ancestry | BEB | 1000 Genomes on GRCh38,1000 Genomes 30x on GRCh38,1000 Genomes phase 3 release |
| **NA20355** | female | SAME124597 | ASW | African Ancestry SW | AFR | African Ancestry | ASW | 1000 Genomes on GRCh38,1000 Genomes 30x on GRCh38,1000 Genomes phase 3 release |
| **HG04174** | female | SAME1839982 | BEB | Bengali | SAS | South Asian Ancestry | BEB | 1000 Genomes on GRCh38,1000 Genomes 30x on GRCh38 |
| **NA20362** | male | SAME124788 | ASW | African Ancestry SW | AFR | African Ancestry | ASW | 1000 Genomes on GRCh38,1000 Genomes 30x on GRCh38,1000 Genomes phase 3 release |
| **NA20761** | female | SAME124363 | TSI | Toscani | EUR | European Ancestry | TSI | 1000 Genomes on GRCh38,1000 Genomes 30x on GRCh38,1000 Genomes phase 3 release,1000 Genomes phase 1 release,Geuvadis |
| **HG04186** | female | SAME1839753 | BEB | Bengali | SAS | South Asian Ancestry | BEB | 1000 Genomes on GRCh38,1000 Genomes 30x on GRCh38,1000 Genomes phase 3 release |
| **NA20766** | female | SAME124370 | TSI | Toscani | EUR | European Ancestry | TSI | 1000 Genomes on GRCh38,1000 Genomes 30x on GRCh38,1000 Genomes phase 3 release,1000 Genomes phase 1 release,Geuvadis |
| **HG04193** | male | SAME1839811 | BEB | Bengali | SAS | South Asian Ancestry | BEB | 1000 Genomes 30x on GRCh38 |
| **NA20773** | female | SAME124575 | TSI | Toscani | EUR | European Ancestry | TSI | 1000 Genomes on GRCh38,1000 Genomes 30x on GRCh38,1000 Genomes phase 3 release,1000 Genomes phase 1 release,Geuvadis |
| **HG04100** | male | SAME1839536 | STU | Tamil | SAS | South Asian Ancestry | STU | 1000 Genomes on GRCh38,1000 Genomes 30x on GRCh38,1000 Genomes phase 3 release |
| **NA20778** | male | SAME124571 | TSI | Toscani | EUR | European Ancestry | TSI | 1000 Genomes on GRCh38,1000 Genomes 30x on GRCh38,1000 Genomes phase 3 release,1000 Genomes phase 1 release,Geuvadis |
| **HG04131** | male | SAME1839279 | BEB | Bengali | SAS | South Asian Ancestry | BEB | 1000 Genomes on GRCh38,1000 Genomes 30x on GRCh38,1000 Genomes phase 3 release |
| **NA20785** | male | SAME123611 | TSI | Toscani | EUR | European Ancestry | TSI | 1000 Genomes on GRCh38,1000 Genomes 30x on GRCh38,1000 Genomes phase 3 release,1000 Genomes phase 1 release,Geuvadis |
| **HG04136** | female | SAME1839310 | BEB | Bengali | SAS | South Asian Ancestry | BEB | 1000 Genomes 30x on GRCh38 |
| **NA20792** | male | SAME123817 | TSI | Toscani | EUR | European Ancestry | TSI | 1000 Genomes on GRCh38,1000 Genomes 30x on GRCh38,1000 Genomes phase 3 release,1000 Genomes phase 1 release,Geuvadis |
| **HG04198** | male | SAME1839835 | ITU | Telugu | SAS | South Asian Ancestry | ITU | 1000 Genomes on GRCh38,1000 Genomes 30x on GRCh38,1000 Genomes phase 3 release |
| **NA20797** | female | SAME123820 | TSI | Toscani | EUR | European Ancestry | TSI | 1000 Genomes on GRCh38,1000 Genomes 30x on GRCh38,1000 Genomes phase 3 release,1000 Genomes phase 1 release,Geuvadis |
| **HG04206** | male | SAME1839447 | ITU | Telugu | SAS | South Asian Ancestry | ITU | 1000 Genomes on GRCh38,1000 Genomes 30x on GRCh38,1000 Genomes phase 3 release |
| **NA20800** | female | SAME125364 | TSI | Toscani | EUR | European Ancestry | TSI | 1000 Genomes on GRCh38,1000 Genomes 30x on GRCh38,1000 Genomes phase 3 release,1000 Genomes phase 1 release,Geuvadis |
| **HG04225** | male | SAME1839048 | ITU | Telugu | SAS | South Asian Ancestry | ITU | 1000 Genomes on GRCh38,1000 Genomes 30x on GRCh38,1000 Genomes phase 3 release |
| **NA20805** | male | SAME125361 | TSI | Toscani | EUR | European Ancestry | TSI | 1000 Genomes on GRCh38,1000 Genomes 30x on GRCh38,1000 Genomes phase 3 release,1000 Genomes phase 1 release,Geuvadis |
| **HG03967** | male | SAME1840072 | ITU | Telugu | SAS | South Asian Ancestry | ITU | 1000 Genomes on GRCh38,1000 Genomes 30x on GRCh38,1000 Genomes phase 3 release |
| **NA20812** | male | SAME125190 | TSI | Toscani | EUR | European Ancestry | TSI | 1000 Genomes on GRCh38,1000 Genomes 30x on GRCh38,1000 Genomes phase 3 release,1000 Genomes phase 1 release,Geuvadis |
| **HG03974** | male | SAME1839735 | ITU | Telugu | SAS | South Asian Ancestry | ITU | 1000 Genomes on GRCh38,1000 Genomes 30x on GRCh38,1000 Genomes phase 3 release |
| **HG03986** | female | SAME1839605 | STU | Tamil | SAS | South Asian Ancestry | STU | 1000 Genomes on GRCh38,1000 Genomes 30x on GRCh38,1000 Genomes phase 3 release |
| **HG03998** | male | SAME1839408 | STU | Tamil | SAS | South Asian Ancestry | STU | 1000 Genomes on GRCh38,1000 Genomes 30x on GRCh38,1000 Genomes phase 3 release |
| **NA21137** | female | SAME124038 | GIH | Gujarati | SAS | South Asian Ancestry | GIH | 1000 Genomes on GRCh38,1000 Genomes 30x on GRCh38,1000 Genomes phase 3 release |
| **HG04023** | male | SAME1839696 | ITU | Telugu | SAS | South Asian Ancestry | ITU | 1000 Genomes on GRCh38,1000 Genomes 30x on GRCh38,1000 Genomes phase 3 release |
| **NA21144** | female | SAME124511 | GIH | Gujarati | SAS | South Asian Ancestry | GIH | 1000 Genomes on GRCh38,1000 Genomes 30x on GRCh38,1000 Genomes phase 3 release |

| | | | | | | | | |
|---|---|---|---|---|---|---|---|---|
| **HG04035** | female | SAME1839487 | STU | Tamil | SAS | South Asian Ancestry | STU | 1000 Genomes on GRCh38,1000 Genomes 30x on GRCh38,1000 Genomes phase 3 release |
| **NA18486** | male | SAME124099 | YRI | Yoruba | AFR | African Ancestry | YRI | 1000 Genomes on GRCh38,1000 Genomes 30x on GRCh38,1000 Genomes phase 3 release,1000 Genomes phase 1 release,Geuvadis |
| **NA18498** | male | SAME124058 | YRI | Yoruba | AFR | African Ancestry | YRI | 1000 Genomes on GRCh38,1000 Genomes 30x on GRCh38,1000 Genomes phase 3 release,1000 Genomes phase 1 release,Geuvadis |
| **NA18501** | male | SAME124991 | YRI | Yoruba | AFR | African Ancestry | YRI | 1000 Genomes on GRCh38,1000 Genomes 30x on GRCh38,1000 Genomes phase 3 release,1000 Genomes phase 1 release |
| **NA20874** | female | SAME123597 | GIH | Gujarati | SAS | South Asian Ancestry | GIH | 1000 Genomes on GRCh38,1000 Genomes 30x on GRCh38,1000 Genomes phase 3 release |
| **NA18506** | male | SAME125059 | YRI | Yoruba | AFR | African Ancestry | YRI | 1000 Genomes 30x on GRCh38,1000 Genomes phase 3 release |
| **NA18518** | female | SAME124889 | YRI | Yoruba | AFR | African Ancestry | YRI | 1000 Genomes 30x on GRCh38,1000 Genomes phase 3 release |
| **NA20881** | female | SAME124531 | GIH | Gujarati | SAS | South Asian Ancestry | GIH | 1000 Genomes on GRCh38,1000 Genomes 30x on GRCh38,1000 Genomes phase 3 release |
| **NA18520** | female | SAME123565 | YRI | Yoruba | AFR | African Ancestry | YRI | 1000 Genomes on GRCh38,1000 Genomes 30x on GRCh38,1000 Genomes phase 3 release,1000 Genomes phase 1 release,Geuvadis |
| **NA20886** | female | SAME123878 | GIH | Gujarati | SAS | South Asian Ancestry | GIH | 1000 Genomes on GRCh38,1000 Genomes 30x on GRCh38,1000 Genomes phase 3 release |
| **NA18525** | female | SAME124672 | CHB | Han Chinese | EAS | East Asian Ancestry | CHB | 1000 Genomes on GRCh38,1000 Genomes 30x on GRCh38,1000 Genomes phase 3 release,1000 Genomes phase 1 release |
| **NA18532** | female | SAME124482 | CHB | Han Chinese | EAS | East Asian Ancestry | CHB | 1000 Genomes on GRCh38,1000 Genomes 30x on GRCh38,1000 Genomes phase 3 release,1000 Genomes phase 1 release,90 Han Chinese high coverage genomes |
| **NA18537** | female | SAME124485 | CHB | Han Chinese | EAS | East Asian Ancestry | CHB | 1000 Genomes on GRCh38,1000 Genomes 30x on GRCh38,1000 Genomes phase 3 release,1000 Genomes phase 1 release,90 Han Chinese high coverage genomes |
| **NA20901** | male | SAME122919 | GIH | Gujarati | SAS | South Asian Ancestry | GIH | 1000 Genomes on GRCh38,1000 Genomes 30x on GRCh38,1000 Genomes phase 3 release |
| **NA18544** | male | SAME124265 | CHB | Han Chinese | EAS | East Asian Ancestry | CHB | 1000 Genomes on GRCh38,1000 Genomes 30x on GRCh38,1000 Genomes phase 3 release,1000 Genomes phase 1 release |
| **NA20906** | female | SAME122918 | GIH | Gujarati | SAS | South Asian Ancestry | GIH | 1000 Genomes on GRCh38,1000 Genomes 30x on GRCh38,1000 Genomes phase 3 release |
| **NA18549** | male | SAME124262 | CHB | Han Chinese | EAS | East Asian Ancestry | CHB | 1000 Genomes on GRCh38,1000 Genomes 30x on GRCh38,1000 Genomes phase 3 release,1000 Genomes phase 1 release |
| **NA21086** | female | SAME123139 | GIH | Gujarati | SAS | South Asian Ancestry | GIH | 1000 Genomes on GRCh38,1000 Genomes 30x on GRCh38,1000 Genomes phase 3 release |
| **NA18563** | male | SAME123928 | CHB | Han Chinese | EAS | East Asian Ancestry | CHB | 1000 Genomes on GRCh38,1000 Genomes 30x on GRCh38,1000 Genomes phase 3 release,1000 Genomes phase 1 release,90 Han Chinese high coverage genomes |
| **NA18570** | female | SAME123741 | CHB | Han Chinese | EAS | East Asian Ancestry | CHB | 1000 Genomes on GRCh38,1000 Genomes 30x on GRCh38,1000 Genomes phase 3 release,1000 Genomes phase 1 release,90 Han Chinese high coverage genomes |
| **NA18582** | female | SAME123905 | CHB | Han Chinese | EAS | East Asian Ancestry | CHB | 1000 Genomes on GRCh38,1000 Genomes 30x on GRCh38,1000 Genomes phase 3 release,1000 Genomes phase 1 release,90 Han Chinese high coverage genomes |
| **NA18599** | female | SAME123713 | CHB | Han Chinese | EAS | East Asian Ancestry | CHB | 1000 Genomes on GRCh38,1000 Genomes 30x on GRCh38,1000 Genomes phase 3 release,1000 Genomes phase 1 release |
| **NA18602** | female | SAME124876 | CHB | Han Chinese | EAS | East Asian Ancestry | CHB | 1000 Genomes on GRCh38,1000 Genomes 30x on GRCh38,1000 Genomes phase 3 release,1000 Genomes phase 1 release |
| **NA18614** | female | SAME125034 | CHB | Han Chinese | EAS | East Asian Ancestry | CHB | 1000 Genomes on GRCh38,1000 Genomes 30x on GRCh38,1000 Genomes phase 3 release,1000 Genomes phase 1 release |
| **NA21113** | male | SAME124181 | GIH | Gujarati | SAS | South Asian Ancestry | GIH | 1000 Genomes on GRCh38,1000 Genomes 30x on GRCh38,1000 Genomes phase 3 release |
| **NA18619** | female | SAME125043 | CHB | Han Chinese | EAS | East Asian Ancestry | CHB | 1000 Genomes on GRCh38,1000 Genomes 30x on GRCh38,1000 Genomes phase 3 release,1000 Genomes phase 1 release |
| **NA21118** | male | SAME123585 | GIH | Gujarati | SAS | South Asian Ancestry | GIH | 1000 Genomes on GRCh38,1000 Genomes 30x on GRCh38,1000 Genomes phase 3 release |
| **NA18621** | male | SAME124509 | CHB | Han Chinese | EAS | East Asian Ancestry | CHB | 1000 Genomes on GRCh38,1000 Genomes 30x on GRCh38,1000 Genomes phase 3 release,1000 Genomes phase 1 release,90 Han Chinese high coverage genomes |
| **NA21120** | female | SAME124028 | GIH | Gujarati | SAS | South Asian Ancestry | GIH | 1000 Genomes on GRCh38,1000 Genomes 30x on GRCh38,1000 Genomes phase 3 release |
| **NA18626** | female | SAME125250 | CHB | Han Chinese | EAS | East Asian Ancestry | CHB | 1000 Genomes on GRCh38,1000 Genomes 30x on GRCh38,1000 Genomes phase 3 release,1000 Genomes phase 1 release |
| **NA21125** | female | SAME124025 | GIH | Gujarati | SAS | South Asian Ancestry | GIH | 1000 Genomes on GRCh38,1000 Genomes 30x on GRCh38,1000 Genomes phase 3 release |
| **NA18633** | male | SAME124704 | CHB | Han Chinese | EAS | East Asian Ancestry | CHB | 1000 Genomes on GRCh38,1000 Genomes 30x on GRCh38,1000 Genomes phase 3 release,1000 Genomes phase 1 release,90 Han Chinese high coverage genomes |
| **NA21093** | male | SAME123323 | GIH | Gujarati | SAS | South Asian Ancestry | GIH | 1000 Genomes on GRCh38,1000 Genomes 30x on GRCh38,1000 Genomes phase 3 release |
| **NA18638** | male | SAME124693 | CHB | Han Chinese | EAS | East Asian Ancestry | CHB | 1000 Genomes on GRCh38,1000 Genomes 30x on GRCh38,1000 Genomes phase 3 release,1000 Genomes phase 1 release |
| **NA21098** | male | SAME123324 | GIH | Gujarati | SAS | South Asian Ancestry | GIH | 1000 Genomes on GRCh38,1000 Genomes 30x on GRCh38,1000 Genomes phase 3 release |
| **NA18640** | female | SAME122980 | CHB | Han Chinese | EAS | East Asian Ancestry | CHB | 1000 Genomes on GRCh38,1000 Genomes 30x on GRCh38,1000 Genomes phase 3 release,1000 Genomes phase 1 release |
| **NA21101** | female | SAME124441 | GIH | Gujarati | SAS | South Asian Ancestry | GIH | 1000 Genomes on GRCh38,1000 Genomes 30x on GRCh38,1000 Genomes phase 3 release |
| **NA18645** | male | SAME122975 | CHB | Han Chinese | EAS | East Asian Ancestry | CHB | 1000 Genomes on GRCh38,1000 Genomes 30x on GRCh38,1000 Genomes phase 3 release,1000 Genomes phase 1 release |
| **NA21106** | female | SAME124434 | GIH | Gujarati | SAS | South Asian Ancestry | GIH | 1000 Genomes on GRCh38,1000 Genomes 30x on GRCh38,1000 Genomes phase 3 release |
| **NA20850** | male | SAME123246 | GIH | Gujarati | SAS | South Asian Ancestry | GIH | 1000 Genomes on GRCh38,1000 Genomes 30x on GRCh38,1000 Genomes phase 3 release |
| **NA18979** | female | SAME123759 | JPT | Japanese | EAS | East Asian Ancestry | JPT | 1000 Genomes on GRCh38,1000 Genomes 30x on GRCh38,1000 Genomes phase 3 release,1000 Genomes phase 1 release |
| **NA20862** | female | SAME123804 | GIH | Gujarati | SAS | South Asian Ancestry | GIH | 1000 Genomes on GRCh38,1000 Genomes 30x on GRCh38,1000 Genomes phase 3 release |
| **NA18981** | female | SAME123426 | JPT | Japanese | EAS | East Asian Ancestry | JPT | 1000 Genomes on GRCh38,1000 Genomes 30x on GRCh38,1000 Genomes phase 3 release,1000 Genomes phase 1 release |
| **NA20867** | male | SAME123809 | GIH | Gujarati | SAS | South Asian Ancestry | GIH | 1000 Genomes on GRCh38,1000 Genomes 30x on GRCh38,1000 Genomes phase 3 release |
| **NA18986** | male | SAME125069 | JPT | Japanese | EAS | East Asian Ancestry | JPT | 1000 Genomes on GRCh38,1000 Genomes 30x on GRCh38,1000 Genomes phase 3 release,1000 Genomes phase 1 release |
| **NA18993** | female | SAME124894 | JPT | Japanese | EAS | East Asian Ancestry | JPT | 1000 Genomes on GRCh38,1000 Genomes 30x on GRCh38,1000 Genomes phase 3 release,1000 Genomes phase 1 release |
| **NA18998** | female | SAME124891 | JPT | Japanese | EAS | East Asian Ancestry | JPT | 1000 Genomes on GRCh38,1000 Genomes 30x on GRCh38,1000 Genomes phase 3 release,1000 Genomes phase 1 release |
| **NA19004** | male | SAME124473 | JPT | Japanese | EAS | East Asian Ancestry | JPT | 1000 Genomes on GRCh38,1000 Genomes 30x on GRCh38,1000 Genomes phase 3 release,1000 Genomes phase 1 release |
| **NA19009** | male | SAME125133 | JPT | Japanese | EAS | East Asian Ancestry | JPT | 1000 Genomes on GRCh38,1000 Genomes 30x on GRCh38,1000 Genomes phase 3 release,1000 Genomes phase 1 release |
| **NA19011** | female | SAME124512 | JPT | Japanese | EAS | East Asian Ancestry | JPT | 1000 Genomes on GRCh38,1000 Genomes 30x on GRCh38,1000 Genomes phase 3 release |
| **NA19023** | female | SAME124854 | LWK | Luhya,Luhya | AFR | African Ancestry,Africa (SGDP) | LWK,LuhyaSGDP | 1000 Genomes on GRCh38,Simons Genome Diversity Project,1000 Genomes 30x on GRCh38,1000 Genomes phase 3 release |
| **NA19028** | male | SAME124848 | LWK | Luhya | AFR | African Ancestry | LWK | 1000 Genomes on GRCh38,1000 Genomes 30x on GRCh38,1000 Genomes phase 3 release,1000 Genomes phase 1 release |
| **NA19030** | female | SAME123544 | LWK | Luhya | AFR | African Ancestry | LWK | 1000 Genomes on GRCh38,1000 Genomes 30x on GRCh38,1000 Genomes phase 3 release |
| **NA19035** | male | SAME124664 | LWK | Luhya | AFR | African Ancestry | LWK | 1000 Genomes on GRCh38,1000 Genomes 30x on GRCh38,1000 Genomes phase 3 release,1000 Genomes phase 1 release |
| **NA19042** | female | SAME125242 | LWK | Luhya | AFR | African Ancestry | LWK | 1000 Genomes on GRCh38,1000 Genomes 30x on GRCh38,1000 Genomes phase 3 release |
| **NA19054** | female | SAME125028 | JPT | Japanese | EAS | East Asian Ancestry | JPT | 1000 Genomes on GRCh38,1000 Genomes 30x on GRCh38,1000 Genomes phase 3 release,1000 Genomes phase 1 release |
| **NA19059** | female | SAME125023 | JPT | Japanese | EAS | East Asian Ancestry | JPT | 1000 Genomes on GRCh38,1000 Genomes 30x on GRCh38,1000 Genomes phase 3 release,1000 Genomes phase 1 release |
| **NA19066** | male | SAME123008 | JPT | Japanese | EAS | East Asian Ancestry | JPT | 1000 Genomes on GRCh38,1000 Genomes 30x on GRCh38,1000 Genomes phase 3 release,1000 Genomes phase 1 release |
| **NA19078** | female | SAME122824 | JPT | Japanese | EAS | East Asian Ancestry | JPT | 1000 Genomes on GRCh38,1000 Genomes 30x on GRCh38,1000 Genomes phase 3 release,1000 Genomes phase 1 release |
| **NA19080** | female | SAME125313 | JPT | Japanese | EAS | East Asian Ancestry | JPT | 1000 Genomes on GRCh38,1000 Genomes 30x on GRCh38,1000 Genomes phase 3 release,1000 Genomes phase 1 release |
| **NA19085** | male | SAME123365 | JPT | Japanese | EAS | East Asian Ancestry | JPT | 1000 Genomes on GRCh38,1000 Genomes 30x on GRCh38,1000 Genomes phase 3 release,1000 Genomes phase 1 release |
| **NA19092** | male | SAME123173 | YRI | Yoruba | AFR | African Ancestry | YRI | 1000 Genomes on GRCh38,1000 Genomes 30x on GRCh38,1000 Genomes phase 3 release,1000 Genomes phase 1 release,Geuvadis |
| **NA19097** | female | SAME123168 | YRI | Yoruba | AFR | African Ancestry | YRI | 1000 Genomes 30x on GRCh38,1000 Genomes phase 3 release |
| **NA19100** | female | SAME125401 | YRI | Yoruba | AFR | African Ancestry | YRI | 1000 Genomes 30x on GRCh38,1000 Genomes phase 3 release |
| **NA12154** | male | SAME124959 | CEU | CEPH | EUR | European Ancestry | CEU | 1000 Genomes on GRCh38,1000 Genomes 30x on GRCh38,1000 Genomes phase 3 release,1000 Genomes phase 1 release,Geuvadis |
| **NA12248** | male | SAME125372 | CEU | CEPH | EUR | European Ancestry | CEU | 1000 Genomes 30x on GRCh38,1000 Genomes phase 3 release |
| **NA12274** | male | SAME125249 | CEU | CEPH | EUR | European Ancestry | CEU | 1000 Genomes 30x on GRCh38,1000 Genomes phase 3 release |
| **NA12286** | male | SAME124642 | CEU | CEPH | EUR | European Ancestry | CEU | 1000 Genomes on GRCh38,1000 Genomes 30x on GRCh38,1000 Genomes phase 3 release,1000 Genomes phase 1 release,Geuvadis |
| **NA12344** | male | SAME123946 | CEU | CEPH | EUR | European Ancestry | CEU | 1000 Genomes 30x on GRCh38,1000 Genomes phase 3 release |
| **NA12399** | male | SAME124193 | CEU | CEPH | EUR | European Ancestry | CEU | 1000 Genomes on GRCh38,1000 Genomes 30x on GRCh38,1000 Genomes phase 3 release,1000 Genomes phase 1 release,Geuvadis |
| **NA12414** | female | SAME124057 | CEU | CEPH | EUR | European Ancestry | CEU | 1000 Genomes on GRCh38,1000 Genomes 30x on GRCh38,1000 Genomes phase 3 release,1000 Genomes phase 1 release |
| **NA12546** | male | SAME124777 | CEU | CEPH | EUR | European Ancestry | CEU | 1000 Genomes on GRCh38,1000 Genomes 30x on GRCh38,1000 Genomes phase 3 release,1000 Genomes phase 1 release,Geuvadis |
| **NA12717** | female | SAME123911 | CEU | CEPH | EUR | European Ancestry | CEU | 1000 Genomes on GRCh38,1000 Genomes 30x on GRCh38,1000 Genomes phase 3 release,1000 Genomes phase 1 release,Geuvadis |
| **NA12748** | male | SAME124827 | CEU | CEPH | EUR | European Ancestry | CEU | 1000 Genomes on GRCh38,1000 Genomes 30x on GRCh38,1000 Genomes phase 3 release,1000 Genomes phase 1 release |
| **NA12750** | male | SAME124646 | CEU | CEPH | EUR | European Ancestry | CEU | 1000 Genomes on GRCh38,1000 Genomes 30x on GRCh38,1000 Genomes phase 3 release,1000 Genomes phase 1 release,Geuvadis |
| **NA12762** | male | SAME125109 | CEU | CEPH | EUR | European Ancestry | CEU | 1000 Genomes on GRCh38,1000 Genomes 30x on GRCh38,1000 Genomes phase 3 release,1000 Genomes phase 1 release,Geuvadis |
| **NA12767** | female | SAME124472 | CEU | CEPH | EUR | European Ancestry | CEU | 1000 Genomes 30x on GRCh38,1000 Genomes phase 3 release |
| **NA12801** | male | SAME122827 | CEU | CEPH | EUR | European Ancestry | CEU | 1000 Genomes 30x on GRCh38,1000 Genomes phase 3 release |
| **NA12813** | female | SAME123024 | CEU | CEPH | EUR | European Ancestry | CEU | 1000 Genomes on GRCh38,1000 Genomes 30x on GRCh38,1000 Genomes phase 3 release,1000 Genomes phase 1 release,Geuvadis |
| **NA12818** | female | SAME123025 | CEU | CEPH | EUR | European Ancestry | CEU | 1000 Genomes 30x on GRCh38,1000 Genomes phase 3 release |
| **NA12832** | female | SAME125290 | CEU | CEPH | EUR | European Ancestry | CEU | 1000 Genomes 30x on GRCh38,1000 Genomes phase 3 release |
| **NA12875** | female | SAME123390 | CEU | CEPH | EUR | European Ancestry | CEU | 1000 Genomes 30x on GRCh38,1000 Genomes phase 3 release |
| **NA18912** | female | SAME123404 | YRI | Yoruba | AFR | African Ancestry | YRI | 1000 Genomes on GRCh38,1000 Genomes 30x on GRCh38,1000 Genomes phase 3 release,1000 Genomes phase 1 release,Geuvadis |
| **NA18917** | male | SAME123407 | YRI | Yoruba | AFR | African Ancestry | YRI | 1000 Genomes on GRCh38,1000 Genomes 30x on GRCh38,1000 Genomes phase 3 release,1000 Genomes phase 1 release,Geuvadis |
| **NA18924** | female | SAME123202 | YRI | Yoruba | AFR | African Ancestry | YRI | 1000 Genomes on GRCh38,1000 Genomes 30x on GRCh38,1000 Genomes phase 3 release,1000 Genomes phase 1 release |
| **NA18943** | male | SAME124309 | JPT | Japanese | EAS | East Asian Ancestry | JPT | 1000 Genomes on GRCh38,1000 Genomes 30x on GRCh38,1000 Genomes phase 3 release,1000 Genomes phase 1 release |
| **NA18948** | male | SAME124316 | JPT | Japanese | EAS | East Asian Ancestry | JPT | 1000 Genomes on GRCh38,1000 Genomes 30x on GRCh38,1000 Genomes phase 3 release,1000 Genomes phase 1 release |
| **NA18950** | female | SAME124112 | JPT | Japanese | EAS | East Asian Ancestry | JPT | 1000 Genomes on GRCh38,1000 Genomes 30x on GRCh38,1000 Genomes phase 3 release,1000 Genomes phase 1 release |
| **NA18962** | male | SAME123958 | JPT | Japanese | EAS | East Asian Ancestry | JPT | 1000 Genomes on GRCh38,1000 Genomes 30x on GRCh38,1000 Genomes phase 3 release,1000 Genomes phase 1 release |
| **NA18967** | male | SAME123963 | JPT | Japanese | EAS | East Asian Ancestry | JPT | 1000 Genomes on GRCh38,1000 Genomes 30x on GRCh38,1000 Genomes phase 3 release,1000 Genomes phase 1 release |
| **NA18974** | male | SAME123768 | JPT | Japanese | EAS | East Asian Ancestry | JPT | 1000 Genomes on GRCh38,1000 Genomes 30x on GRCh38,1000 Genomes phase 3 release,1000 Genomes phase 1 release |
| **NA10855** | female | SAME125331 | CEU | CEPH | EUR | European Ancestry | CEU | 1000 Genomes 30x on GRCh38,1000 Genomes phase 3 release |
| **NA11839** | male | SAME124555 | CEU | CEPH | EUR | European Ancestry | CEU | 1000 Genomes 30x on GRCh38,1000 Genomes phase 3 release |
| **NA11891** | male | SAME123046 | CEU | CEPH | EUR | European Ancestry | CEU | 1000 Genomes 30x on GRCh38,1000 Genomes phase 3 release |
| **NA11930** | male | SAME125095 | CEU | CEPH | EUR | European Ancestry | CEU | 1000 Genomes on GRCh38,1000 Genomes 30x on GRCh38,1000 Genomes phase 3 release,1000 Genomes phase 1 release,Geuvadis |
| **NA11992** | male | SAME123964 | CEU | CEPH | EUR | European Ancestry | CEU | 1000 Genomes on GRCh38,1000 Genomes 30x on GRCh38,1000 Genomes phase 3 release,1000 Genomes phase 1 release,Geuvadis |
| **NA12003** | male | SAME125142 | CEU | CEPH | EUR | European Ancestry | CEU | 1000 Genomes on GRCh38,1000 Genomes 30x on GRCh38,1000 Genomes phase 3 release,1000 Genomes phase 1 release |
| **NA12046** | female | SAME123213 | CEU | CEPH | EUR | European Ancestry | CEU | 1000 Genomes on GRCh38,1000 Genomes 30x on GRCh38,1000 Genomes phase 3 release,1000 Genomes phase 1 release |
| **NA12058** | female | SAME123416 | CEU | CEPH | EUR | European Ancestry | CEU | 1000 Genomes on GRCh38,1000 Genomes 30x on GRCh38,1000 Genomes phase 3 release,1000 Genomes phase 1 release,Geuvadis |
| **NA18854** | male | SAME124734 | YRI | Yoruba | AFR | African Ancestry | YRI | 1000 Genomes 30x on GRCh38,1000 Genomes phase 3 release |
| **NA18859** | male | SAME124731 | YRI | Yoruba | AFR | African Ancestry | YRI | 1000 Genomes 30x on GRCh38,1000 Genomes phase 3 release |
| **NA18861** | female | SAME124905 | YRI | Yoruba | AFR | African Ancestry | YRI | 1000 Genomes on GRCh38,1000 Genomes 30x on GRCh38,1000 Genomes phase 3 release,1000 Genomes phase 1 release,Geuvadis |
| **NA18873** | female | SAME123536 | YRI | Yoruba | AFR | African Ancestry | YRI | 1000 Genomes on GRCh38,1000 Genomes 30x on GRCh38,1000 Genomes phase 3 release,1000 Genomes phase 1 release,Geuvadis |
| **NA18878** | female | SAME125107 | YRI | Yoruba | AFR | African Ancestry | YRI | 1000 Genomes on GRCh38,1000 Genomes 30x on GRCh38,1000 Genomes phase 3 release |
| **NA19472** | female | SAME123120 | LWK | Luhya | AFR | African Ancestry | LWK | 1000 Genomes on GRCh38,1000 Genomes 30x on GRCh38,1000 Genomes phase 3 release,1000 Genomes phase 1 release |
| **NA19648** | female | SAME123307 | MXL | Mexican Ancestry | AMR | American Ancestry | MXL | 1000 Genomes on GRCh38,1000 Genomes 30x on GRCh38,1000 Genomes phase 3 release,1000 Genomes phase 1 release |
| **NA19650** | male | SAME124976 | MXL | Mexican Ancestry | AMR | American Ancestry | MXL | 1000 Genomes 30x on GRCh38,Human Genome Structural Variation Consortium, Phase 2,1000 Genomes phase 3 release |
| **NA19655** | male | SAME124980 | MXL | Mexican Ancestry | AMR | American Ancestry | MXL | 1000 Genomes on GRCh38,1000 Genomes 30x on GRCh38,1000 Genomes phase 3 release,1000 Genomes phase 1 release |
| **NA19662** | female | SAME122960 | MXL | Mexican Ancestry | AMR | American Ancestry | MXL | 1000 Genomes 30x on GRCh38,1000 Genomes phase 3 release |
| **NA19679** | male | SAME122796 | MXL | Mexican Ancestry | AMR | American Ancestry | MXL | 1000 Genomes on GRCh38,1000 Genomes 30x on GRCh38,1000 Genomes phase 3 release,1000 Genomes phase 1 release |
| **NA19681** | female | SAME122938 | MXL | Mexican Ancestry | AMR | American Ancestry | MXL | 1000 Genomes on GRCh38,1000 Genomes 30x on GRCh38,1000 Genomes phase 3 release,1000 Genomes phase 1 release |

| | | | | | | | | |
|---|---|---|---|---|---|---|---|---|
| **NA19249** | male | SAME123682 | YRI | Yoruba | AFR | African Ancestry | YRI | 1000 Genomes 30x on GRCh38,1000 Genomes phase 3 release |
| **NA19256** | male | SAME124707 | YRI | Yoruba | AFR | African Ancestry | YRI | 1000 Genomes on GRCh38,1000 Genomes 30x on GRCh38,1000 Genomes phase 3 release,1000 Genomes phase 1 release,Geuvadis |
| **NA19307** | male | SAME124981 | LWK | Luhya | AFR | African Ancestry | LWK | 1000 Genomes on GRCh38,1000 Genomes 30x on GRCh38,1000 Genomes phase 3 release,1000 Genomes phase 1 release |
| **NA19314** | female | SAME123309 | LWK | Luhya | AFR | African Ancestry | LWK | 1000 Genomes on GRCh38,1000 Genomes 30x on GRCh38,1000 Genomes phase 3 release,1000 Genomes phase 1 release |
| **NA19319** | male | SAME123438 | LWK | Luhya | AFR | African Ancestry | LWK | 1000 Genomes on GRCh38,1000 Genomes 30x on GRCh38,1000 Genomes phase 3 release,1000 Genomes phase 1 release |
| **NA19321** | female | SAME124569 | LWK | Luhya | AFR | African Ancestry | LWK | 1000 Genomes on GRCh38,1000 Genomes 30x on GRCh38,1000 Genomes phase 3 release,1000 Genomes phase 1 release |
| **NA19338** | female | SAME123274 | LWK | Luhya | AFR | African Ancestry | LWK | 1000 Genomes on GRCh38,1000 Genomes 30x on GRCh38,1000 Genomes phase 3 release,1000 Genomes phase 1 release |
| **NA19376** | male | SAME124469 | LWK | Luhya | AFR | African Ancestry | LWK | 1000 Genomes on GRCh38,1000 Genomes 30x on GRCh38,1000 Genomes phase 3 release,1000 Genomes phase 1 release |
| **NA19686** | female | SAME122939 | MXL | Mexican Ancestry | AMR | American Ancestry | MXL | 1000 Genomes 30x on GRCh38,1000 Genomes phase 3 release |
| **NA19701** | female | SAME124232 | ASW | African Ancestry SW | AFR | African Ancestry | ASW | 1000 Genomes on GRCh38,1000 Genomes 30x on GRCh38,1000 Genomes phase 3 release,1000 Genomes phase 1 release |
| **NA19713** | female | SAME124448 | ASW | African Ancestry SW | AFR | African Ancestry | ASW | 1000 Genomes on GRCh38,1000 Genomes 30x on GRCh38,1000 Genomes phase 3 release,1000 Genomes phase 1 release |
| **NA19718** | female | SAME124444 | MXL | Mexican Ancestry | AMR | American Ancestry | MXL | 1000 Genomes 30x on GRCh38,1000 Genomes phase 3 release |
| **NA19720** | male | SAME124118 | MXL | Mexican Ancestry | AMR | American Ancestry | MXL | 1000 Genomes on GRCh38,1000 Genomes 30x on GRCh38,1000 Genomes phase 3 release,1000 Genomes phase 1 release |
| **NA19725** | female | SAME124689 | MXL | Mexican Ancestry | AMR | American Ancestry | MXL | 1000 Genomes on GRCh38,1000 Genomes 30x on GRCh38,1000 Genomes phase 3 release,1000 Genomes phase 1 release |
| **NA19732** | male | SAME124778 | MXL | Mexican Ancestry | AMR | American Ancestry | MXL | 1000 Genomes on GRCh38,1000 Genomes 30x on GRCh38,1000 Genomes phase 3 release,1000 Genomes phase 1 release |
| **NA19749** | female | SAME124726 | MXL | Mexican Ancestry | AMR | American Ancestry | MXL | 1000 Genomes on GRCh38,1000 Genomes 30x on GRCh38,1000 Genomes phase 3 release,1000 Genomes phase 1 release |
| **NA19174** | male | SAME124882 | YRI | Yoruba | AFR | African Ancestry | YRI | 1000 Genomes 30x on GRCh38,1000 Genomes phase 3 release |
| **NA19186** | male | SAME123861 | YRI | Yoruba | AFR | African Ancestry | YRI | 1000 Genomes 30x on GRCh38,1000 Genomes phase 3 release |
| **NA19198** | male | SAME124049 | YRI | Yoruba | AFR | African Ancestry | YRI | 1000 Genomes on GRCh38,1000 Genomes 30x on GRCh38,1000 Genomes phase 3 release,1000 Genomes phase 1 release,Geuvadis |
| **NA19201** | female | SAME122990 | YRI | Yoruba | AFR | African Ancestry | YRI | 1000 Genomes on GRCh38,1000 Genomes 30x on GRCh38,1000 Genomes phase 3 release,1000 Genomes phase 1 release,Geuvadis |
| **NA19206** | female | SAME122983 | YRI | Yoruba | AFR | African Ancestry | YRI | 1000 Genomes on GRCh38,1000 Genomes 30x on GRCh38,1000 Genomes phase 3 release,1000 Genomes phase 1 release,Geuvadis |
| **NA19213** | male | SAME125384 | YRI | Yoruba | AFR | African Ancestry | YRI | 1000 Genomes on GRCh38,1000 Genomes 30x on GRCh38,1000 Genomes phase 3 release,1000 Genomes phase 1 release,Geuvadis |
| **NA19225** | female | SAME125218 | YRI | Yoruba | AFR | African Ancestry | YRI | 1000 Genomes on GRCh38,1000 Genomes 30x on GRCh38,1000 Genomes phase 3 release,1000 Genomes phase 1 release,Geuvadis |
| **NA19237** | female | SAME125000 | YRI | Yoruba | AFR | African Ancestry | YRI | 1000 Genomes 30x on GRCh38,1000 Genomes phase 3 release |
| **NA19383** | male | SAME124273 | LWK | Luhya | AFR | African Ancestry | LWK | 1000 Genomes on GRCh38,1000 Genomes 30x on GRCh38,1000 Genomes phase 3 release,1000 Genomes phase 1 release |
| **NA19390** | female | SAME124493 | LWK | Luhya | AFR | African Ancestry | LWK | 1000 Genomes on GRCh38,1000 Genomes 30x on GRCh38,1000 Genomes phase 3 release,1000 Genomes phase 1 release |
| **NA19395** | female | SAME124490 | LWK | Luhya | AFR | African Ancestry | LWK | 1000 Genomes on GRCh38,1000 Genomes 30x on GRCh38,1000 Genomes phase 3 release,1000 Genomes phase 1 release |
| **NA19403** | female | SAME123636 | LWK | Luhya | AFR | African Ancestry | LWK | 1000 Genomes on GRCh38,1000 Genomes 30x on GRCh38,1000 Genomes phase 3 release,1000 Genomes phase 1 release |
| **NA19434** | female | SAME123850 | LWK | Luhya | AFR | African Ancestry | LWK | 1000 Genomes on GRCh38,1000 Genomes 30x on GRCh38,Human Genome Structural Variation Consortium, Phase 2,1000 Genomes phase 3 release,1000 Genomes ph[illegible] |
| **NA19439** | female | SAME123847 | LWK | Luhya | AFR | African Ancestry | LWK | 1000 Genomes on GRCh38,1000 Genomes 30x on GRCh38,1000 Genomes phase 3 release,1000 Genomes phase 1 release |
| **NA19446** | female | SAME122949 | LWK | Luhya | AFR | African Ancestry | LWK | 1000 Genomes on GRCh38,1000 Genomes 30x on GRCh38,1000 Genomes phase 3 release,1000 Genomes phase 1 release |
| **NA19117** | male | SAME124914 | YRI | Yoruba | AFR | African Ancestry | YRI | 1000 Genomes on GRCh38,1000 Genomes 30x on GRCh38,1000 Genomes phase 3 release,1000 Genomes phase 1 release,Geuvadis |
| **NA19129** | female | SAME125021 | YRI | Yoruba | AFR | African Ancestry | YRI | 1000 Genomes on GRCh38,1000 Genomes 30x on GRCh38,1000 Genomes phase 3 release,1000 Genomes phase 1 release,Geuvadis |
| **NA19131** | female | SAME125230 | YRI | Yoruba | AFR | African Ancestry | YRI | 1000 Genomes on GRCh38,1000 Genomes 30x on GRCh38,1000 Genomes phase 3 release,1000 Genomes phase 1 release,Geuvadis |
| **NA19143** | female | SAME124609 | YRI | Yoruba | AFR | African Ancestry | YRI | 1000 Genomes on GRCh38,1000 Genomes 30x on GRCh38,1000 Genomes phase 3 release,1000 Genomes phase 1 release,Geuvadis |
| **NA19148** | female | SAME124606 | YRI | Yoruba | AFR | African Ancestry | YRI | 1000 Genomes 30x on GRCh38,1000 Genomes phase 3 release |
| **NA19150** | male | SAME124838 | YRI | Yoruba | AFR | African Ancestry | YRI | 1000 Genomes on GRCh38,1000 Genomes 30x on GRCh38,1000 Genomes phase 3 release,1000 Genomes phase 1 release,Geuvadis |
| **NA19751** | male | SAME123699 | MXL | Mexican Ancestry | AMR | American Ancestry | MXL | 1000 Genomes 30x on GRCh38,1000 Genomes phase 3 release |
| **NA19756** | male | SAME123696 | MXL | Mexican Ancestry | AMR | American Ancestry | MXL | 1000 Genomes on GRCh38,1000 Genomes 30x on GRCh38,1000 Genomes phase 3 release,1000 Genomes phase 1 release |
| **NA19763** | female | SAME123896 | MXL | Mexican Ancestry | AMR | American Ancestry | MXL | 1000 Genomes 30x on GRCh38,1000 Genomes phase 3 release |
| **NA19770** | female | SAME124068 | MXL | Mexican Ancestry | AMR | American Ancestry | MXL | 1000 Genomes on GRCh38,1000 Genomes 30x on GRCh38,1000 Genomes phase 3 release,1000 Genomes phase 1 release |
| **NA19775** | female | SAME124065 | MXL | Mexican Ancestry | AMR | American Ancestry | MXL | 1000 Genomes 30x on GRCh38,1000 Genomes phase 3 release |
| **NA19782** | female | SAME123145 | MXL | Mexican Ancestry | AMR | American Ancestry | MXL | 1000 Genomes on GRCh38,1000 Genomes 30x on GRCh38,1000 Genomes phase 3 release,1000 Genomes phase 1 release |
| **NA19787** | male | SAME123148 | MXL | Mexican Ancestry | AMR | American Ancestry | MXL | 1000 Genomes 30x on GRCh38,1000 Genomes phase 3 release |
| **NA19794** | female | SAME123328 | MXL | Mexican Ancestry | AMR | American Ancestry | MXL | 1000 Genomes on GRCh38,1000 Genomes 30x on GRCh38,1000 Genomes phase 3 release,1000 Genomes phase 1 release |
| **NA19819** | female | SAME124676 | ASW | African Ancestry SW | AFR | African Ancestry | ASW | 1000 Genomes on GRCh38,1000 Genomes 30x on GRCh38,1000 Genomes phase 3 release,1000 Genomes phase 1 release |
| **NA19908** | male | SAME125053 | ASW | African Ancestry SW | AFR | African Ancestry | ASW | 1000 Genomes on GRCh38,1000 Genomes 30x on GRCh38,1000 Genomes phase 3 release,1000 Genomes phase 1 release |
| **NA19922** | male | SAME124711 | ASW | African Ancestry SW | AFR | African Ancestry | ASW | 1000 Genomes on GRCh38,1000 Genomes 30x on GRCh38,1000 Genomes phase 3 release,1000 Genomes phase 1 release |
| **NA19984** | male | SAME124287 | ASW | African Ancestry SW | AFR | African Ancestry | ASW | 1000 Genomes on GRCh38,1000 Genomes 30x on GRCh38,1000 Genomes phase 3 release,1000 Genomes phase 1 release |
| **NA20126** | male | SAME122885 | ASW | African Ancestry SW | AFR | African Ancestry | ASW | 1000 Genomes on GRCh38,1000 Genomes 30x on GRCh38,1000 Genomes phase 3 release,1000 Genomes phase 1 release |
| **NA20289** | female | SAME124579 | ASW | African Ancestry SW | AFR | African Ancestry | ASW | 1000 Genomes on GRCh38,1000 Genomes 30x on GRCh38,1000 Genomes phase 3 release,1000 Genomes phase 1 release |
| **NA20291** | male | SAME124373 | ASW | African Ancestry SW | AFR | African Ancestry | ASW | 1000 Genomes on GRCh38,1000 Genomes 30x on GRCh38,1000 Genomes phase 3 release,1000 Genomes phase 1 release |
| **NA20296** | female | SAME124380 | ASW | African Ancestry SW | AFR | African Ancestry | ASW | 1000 Genomes on GRCh38,1000 Genomes 30x on GRCh38,1000 Genomes phase 3 release,1000 Genomes phase 1 release |
| **NA20513** | male | SAME124561 | TSI | Toscani | EUR | European Ancestry | TSI | 1000 Genomes on GRCh38,1000 Genomes 30x on GRCh38,1000 Genomes phase 3 release,1000 Genomes phase 1 release,Geuvadis |
| **NA20518** | male | SAME123178 | TSI | Toscani | EUR | European Ancestry | TSI | 1000 Genomes on GRCh38,1000 Genomes 30x on GRCh38,1000 Genomes phase 3 release,1000 Genomes phase 1 release,Geuvadis |
| **NA20520** | male | SAME124019 | TSI | Toscani | EUR | European Ancestry | TSI | 1000 Genomes on GRCh38,1000 Genomes 30x on GRCh38,1000 Genomes phase 3 release,1000 Genomes phase 1 release,Geuvadis |
| **NA20525** | male | SAME124015 | TSI | Toscani | EUR | European Ancestry | TSI | 1000 Genomes on GRCh38,1000 Genomes 30x on GRCh38,1000 Genomes phase 3 release,1000 Genomes phase 1 release,Geuvadis |
| **NA20532** | male | SAME124171 | TSI | Toscani | EUR | European Ancestry | TSI | 1000 Genomes on GRCh38,1000 Genomes 30x on GRCh38,1000 Genomes phase 3 release,1000 Genomes phase 1 release,Geuvadis |
| **NA20544** | male | SAME125146 | TSI | Toscani | EUR | European Ancestry | TSI | 1000 Genomes on GRCh38,1000 Genomes 30x on GRCh38,1000 Genomes phase 3 release,1000 Genomes phase 1 release,Geuvadis |
| **NA20582** | female | SAME123474 | TSI | Toscani | EUR | European Ancestry | TSI | 1000 Genomes on GRCh38,1000 Genomes 30x on GRCh38,1000 Genomes phase 3 release,1000 Genomes phase 1 release,Geuvadis |
| **NA20587** | female | SAME123481 | TSI | Toscani | EUR | European Ancestry | TSI | 1000 Genomes on GRCh38,1000 Genomes 30x on GRCh38,1000 Genomes phase 3 release |
| **NA20753** | female | SAME124178 | TSI | Toscani | EUR | European Ancestry | TSI | 1000 Genomes on GRCh38,1000 Genomes 30x on GRCh38,1000 Genomes phase 3 release,1000 Genomes phase 1 release |
| **NA20758** | male | SAME125117 | TSI | Toscani | EUR | European Ancestry | TSI | 1000 Genomes on GRCh38,1000 Genomes 30x on GRCh38,1000 Genomes phase 3 release,1000 Genomes phase 1 release,Geuvadis |
| **NA20760** | female | SAME124364 | TSI | Toscani | EUR | European Ancestry | TSI | 1000 Genomes on GRCh38,1000 Genomes 30x on GRCh38,1000 Genomes phase 3 release,1000 Genomes phase 1 release,Geuvadis |
| **NA20765** | male | SAME124367 | TSI | Toscani | EUR | European Ancestry | TSI | 1000 Genomes on GRCh38,1000 Genomes 30x on GRCh38,1000 Genomes phase 3 release,1000 Genomes phase 1 release,Geuvadis |
| **NA20772** | female | SAME124574 | TSI | Toscani | EUR | European Ancestry | TSI | 1000 Genomes on GRCh38,1000 Genomes 30x on GRCh38,1000 Genomes phase 3 release,1000 Genomes phase 1 release,Geuvadis |
| **NA21143** | female | SAME123359 | GIH | Gujarati | SAS | South Asian Ancestry | GIH | 1000 Genomes on GRCh38,1000 Genomes 30x on GRCh38,1000 Genomes phase 3 release |
| **NA20796** | male | SAME123819 | TSI | Toscani | EUR | European Ancestry | TSI | 1000 Genomes on GRCh38,1000 Genomes 30x on GRCh38,1000 Genomes phase 3 release,1000 Genomes phase 1 release,Geuvadis |
| **NA20804** | female | SAME125224 | TSI | Toscani | EUR | European Ancestry | TSI | 1000 Genomes on GRCh38,1000 Genomes 30x on GRCh38,1000 Genomes phase 3 release,1000 Genomes phase 1 release,Geuvadis |
| **NA20809** | male | SAME125359 | TSI | Toscani | EUR | European Ancestry | TSI | 1000 Genomes on GRCh38,1000 Genomes 30x on GRCh38,1000 Genomes phase 3 release,1000 Genomes phase 1 release,Geuvadis |
| **NA20811** | male | SAME123069 | TSI | Toscani | EUR | European Ancestry | TSI | 1000 Genomes on GRCh38,1000 Genomes 30x on GRCh38,1000 Genomes phase 3 release,1000 Genomes phase 1 release,Geuvadis |
| **NA20828** | female | SAME123076 | TSI | Toscani | EUR | European Ancestry | TSI | 1000 Genomes on GRCh38,1000 Genomes 30x on GRCh38,1000 Genomes phase 3 release,1000 Genomes phase 1 release,Geuvadis |
| **NA20847** | female | SAME123432 | GIH | Gujarati | SAS | South Asian Ancestry | GIH | 1000 Genomes on GRCh38,1000 Genomes 30x on GRCh38,Human Genome Structural Variation Consortium, Phase 2,1000 Genomes phase 3 release |
| **NA20854** | female | SAME123248 | GIH | Gujarati | SAS | South Asian Ancestry | GIH | 1000 Genomes on GRCh38,1000 Genomes 30x on GRCh38,1000 Genomes phase 3 release |
| **NA20342** | male | SAME124398 | ASW | African Ancestry SW | AFR | African Ancestry | ASW | 1000 Genomes on GRCh38,1000 Genomes 30x on GRCh38,1000 Genomes phase 3 release,1000 Genomes phase 1 release |
| **NA20359** | female | SAME124601 | ASW | African Ancestry SW | AFR | African Ancestry | ASW | 1000 Genomes on GRCh38,1000 Genomes 30x on GRCh38,1000 Genomes phase 3 release,1000 Genomes phase 1 release |
| **NA20412** | female | SAME124002 | ASW | African Ancestry SW | AFR | African Ancestry | ASW | 1000 Genomes on GRCh38,1000 Genomes 30x on GRCh38,1000 Genomes phase 3 release,1000 Genomes phase 1 release |
| **NA20506** | female | SAME124362 | TSI | Toscani | EUR | European Ancestry | TSI | 1000 Genomes on GRCh38,1000 Genomes 30x on GRCh38,1000 Genomes phase 3 release,1000 Genomes phase 1 release,Geuvadis |
| **NA20859** | female | SAME123249 | GIH | Gujarati | SAS | South Asian Ancestry | GIH | 1000 Genomes on GRCh38,1000 Genomes 30x on GRCh38,1000 Genomes phase 3 release |
| **NA20861** | male | SAME123807 | GIH | Gujarati | SAS | South Asian Ancestry | GIH | 1000 Genomes on GRCh38,1000 Genomes 30x on GRCh38,1000 Genomes phase 3 release |
| **NA20866** | male | SAME123808 | GIH | Gujarati | SAS | South Asian Ancestry | GIH | 1000 Genomes on GRCh38,1000 Genomes 30x on GRCh38,1000 Genomes phase 3 release |
| **NA20878** | female | SAME123595 | GIH | Gujarati | SAS | South Asian Ancestry | GIH | 1000 Genomes on GRCh38,1000 Genomes 30x on GRCh38,1000 Genomes phase 3 release |
| **NA20885** | male | SAME124525 | GIH | Gujarati | SAS | South Asian Ancestry | GIH | 1000 Genomes on GRCh38,1000 Genomes 30x on GRCh38,1000 Genomes phase 3 release |
| **NA20892** | female | SAME123109 | GIH | Gujarati | SAS | South Asian Ancestry | GIH | 1000 Genomes on GRCh38,1000 Genomes 30x on GRCh38,1000 Genomes phase 3 release |
| **NA20897** | male | SAME123664 | GIH | Gujarati | SAS | South Asian Ancestry | GIH | 1000 Genomes on GRCh38,1000 Genomes 30x on GRCh38,1000 Genomes phase 3 release |
| **NA21097** | female | SAME123319 | GIH | Gujarati | SAS | South Asian Ancestry | GIH | 1000 Genomes on GRCh38,1000 Genomes 30x on GRCh38,1000 Genomes phase 3 release |
| **NA21100** | male | SAME124440 | GIH | Gujarati | SAS | South Asian Ancestry | GIH | 1000 Genomes on GRCh38,1000 Genomes 30x on GRCh38,1000 Genomes phase 3 release |
| **NA21105** | male | SAME124437 | GIH | Gujarati | SAS | South Asian Ancestry | GIH | 1000 Genomes on GRCh38,1000 Genomes 30x on GRCh38,1000 Genomes phase 3 release |
| **NA21112** | male | SAME124182 | GIH | Gujarati | SAS | South Asian Ancestry | GIH | 1000 Genomes on GRCh38,1000 Genomes 30x on GRCh38,1000 Genomes phase 3 release |
| **NA20900** | female | SAME122920 | GIH | Gujarati | SAS | South Asian Ancestry | GIH | 1000 Genomes on GRCh38,1000 Genomes 30x on GRCh38,1000 Genomes phase 3 release |
| **NA20905** | male | SAME122915 | GIH | Gujarati | SAS | South Asian Ancestry | GIH | 1000 Genomes on GRCh38,1000 Genomes 30x on GRCh38,1000 Genomes phase 3 release |
| **NA21092** | male | SAME123322 | GIH | Gujarati | SAS | South Asian Ancestry | GIH | 1000 Genomes on GRCh38,1000 Genomes 30x on GRCh38,1000 Genomes phase 3 release |
| **NA21117** | male | SAME124185 | GIH | Gujarati | SAS | South Asian Ancestry | GIH | 1000 Genomes on GRCh38,1000 Genomes 30x on GRCh38,1000 Genomes phase 3 release |
| **NA21124** | male | SAME124024 | GIH | Gujarati | SAS | South Asian Ancestry | GIH | 1000 Genomes on GRCh38,1000 Genomes 30x on GRCh38,1000 Genomes phase 3 release |
| **NA21129** | male | SAME124023 | GIH | Gujarati | SAS | South Asian Ancestry | GIH | 1000 Genomes on GRCh38,1000 Genomes 30x on GRCh38,1000 Genomes phase 3 release |
| **NA12236** | female | SAMEA6604124 | CEU | CEPH | EUR | European Ancestry | CEU | 1000 Genomes 30x on GRCh38 |
