## Supplemental Figure 1 for "High-throughput complement component 4 genomic sequence analysis with C4Investigator"

A

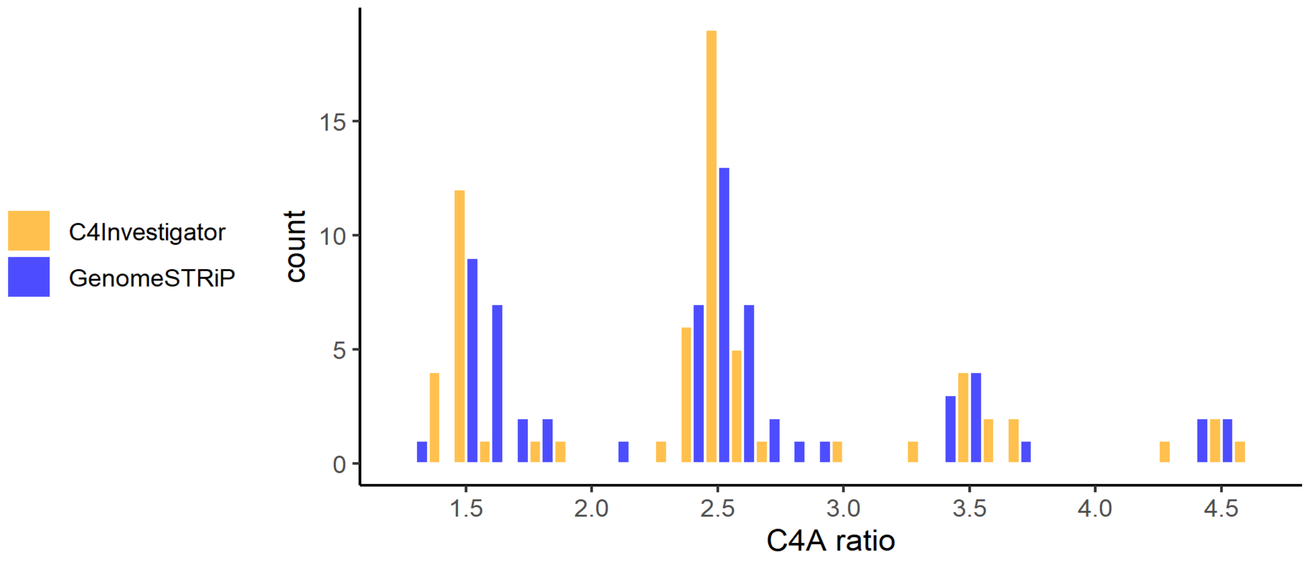


B

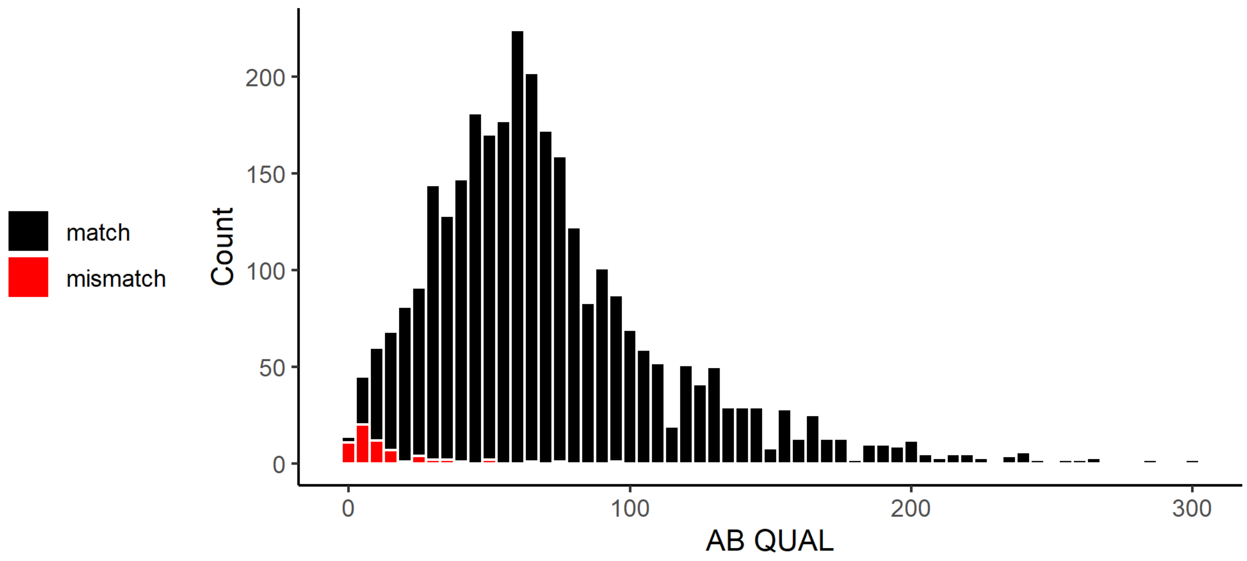


**Figure S1.** A. Histogram of normalized ratios for *C4A* read/k-mer counts for the C4Investigator and GenomeSTRiP workflows for discordant samples from the 1000 Genomes Project dataset. B. Histogram of quality scores for the concordant (match) and discordant (mismatch) samples when comparing results for the C4Investigator and Genome STRiP workflows for the 1000 Genomes Project dataset.
