## Supplemental Figure 2 for "High-throughput complement component 4 genomic sequence analysis with C4Investigator"

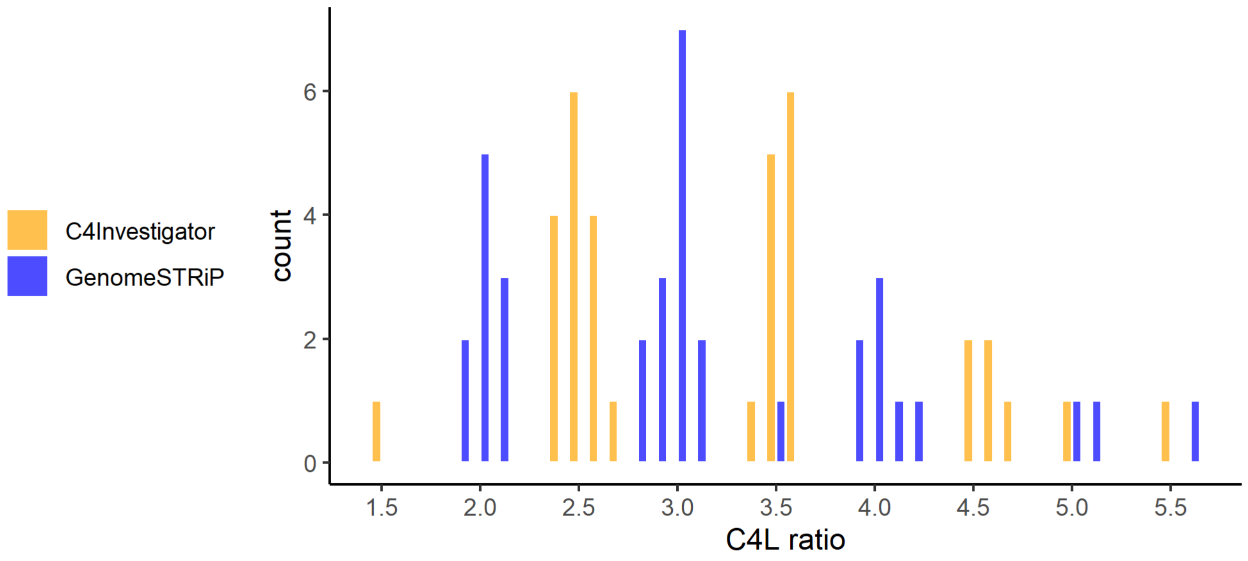


**Figure S2.** Histogram of normalized ratios for *C4(L)* read/k-mer counts for the C4Investigator and GenomeSTRiP workflows for discordant samples from the 1000 Genomes Project dataset.
